# DupyliCate: mining, classifying, and characterizing gene duplications

**DOI:** 10.1101/2025.10.10.681656

**Authors:** Shakunthala Natarajan, Boas Pucker

## Abstract

Paralogs, copies of a gene, form an important basis for novelty during evolution. Analysis of such gene duplications is important to understand the emergence of novel traits during evolution. DupyliCate is a Python tool that has been developed for this purpose. With the ability to process multiple datasets concurrently, flexible features, and parameters to set species-specific thresholds, DupyliCate offers a high-throughput method for gene copy identification and analysis. The different available parameters and modes are explored in detail based on the *Arabidopsis thaliana* datasets. Proof of concept for the tool is presented by characterizing well known duplications in different plants, and its broad applicability is demonstrated by running it on diverse datasets including complex plant genome sequences with high heterozygosity. Further, two case studies involving the evolution of *flavonol synthase* (*FLS*) genes in Brassicales, and the evolution of flavonol synthesis regulating myeloblastosis (MYB) transcription factors- *MYB12* and *MYB111* across a large number of plant species, are presented as exemplar use cases. The tool’s applicability beyond plants is demonstrated on *Escherichia coli, Saccharomyces cerevisiae, and Caenorhabditis elegans* datasets. DupyliCate is available at: https://github.com/ShakNat/DupyliCate.

## Introduction

Gene duplication products also called paralogs are homologous genes originating from a common ancestor gene. In contrast to orthologs, they arise through duplication events and not speciation events^1^. They are an important source of evolutionary novelty in all organisms and are major drivers for the diversification of biosynthetic pathways seen in plants and other organisms. Gene duplications can broadly be divided into small-scale duplications and large-scale duplications. The small-scale duplication group encompasses tandem duplications, proximal duplications, transposon-derived duplications, and retroduplications. Whereas, the large-scale duplication group includes whole genome duplications, and segmental duplications^2^. Identification of gene duplications is fraught with difficulties like functional redundancy of gene duplicates, and extensive small-scale duplication events. It is also compounded by gene loss, rearrangement, mutation events across evolutionary time scales and presence of multidomain protein structures^2,3^. Lastly, gene duplicates can be formed by a number of mechanisms, and have different ages and evolutionary fates, further increasing the difficulty of detection and analysis^1^.

Nevertheless, there have been huge research efforts in this direction resulting in a plethora of tools and methodologies to identify gene duplications. GenomeHistory^4^, was among the first tools to provide a basic framework for gene duplication detection with a combination of local and global alignment approaches followed by pairwise computation of Ka/Ks values to evaluate selection pressures. But this software’s web page no longer exists and is hence not accessible. Following this, several tools like OrthoMCL^5^, OrthoFinder^6^, MCScanX^7^ were developed to identify orthologs and as a result also distinguish them from paralogs using tree-based approaches and synteny approaches, respectively. All these tools focus more on the ortholog identification aspects. Next, DuplicationDetector^8^ was a Perl script developed to identify putative recent segmental duplicates in homozygous organisms from next-generation sequencing data. It focuses on identifying large scale duplications like segmental duplications, at the DNA level. Later, with a detailed study on gene duplications and evolution in plants, DupGen_finder^9^, a Perl based script was developed to identify gene duplicate pairs, and comprehensively classify gene duplicates into tandem duplicates, proximal duplicates, transposed duplicates and whole genome duplicates. The study also provided separate scripts for Ka/Ks analysis by integrating KaKs_Calculator 2.0^10^, a dedicated tool for Ka/Ks calculations. Ka/Ks is defined as the ratio of nonsynonymous mutation rate to that of synonymous mutation rate. Ka/Ks ratios are useful for analysing selection pressures and sequence divergence between genes. Thus, the Ka/Ks computation was integrated with the tool, to study the sequence divergence of the identified gene duplicates ^9,10^. The work further involved expression divergence analysis of the identified gene duplicates, but this analysis was not a part of the DupGen_finder main script, and is not expandable to wide-ranging user data. Zhang and co-workers later developed HSDFinder^11^, a BLAST-based web tool to annotate, categorize and visualize highly similar duplicates across species. More recently, an updated version of this tool, called HSDSnake^12^ was released as a user-friendly SnakeMake pipeline, with added features to classify duplicate pairs by integrating the DupGen_finder script and compute Ka/Ks ratios. WGDI^13^ was another Python-based command-line tool that was developed to identify and characterize whole genome duplication related events in species, thus focusing on large scale duplications. Another Python-based tool integrating machine learning framework to detect orthologs, annotate them, identify gene losses and duplications across alignable genomes is TOGA (Tool to infer Orthologs from Genome Alignments)^14^. TOGA leverages alignments between intergenic regions and introns to identify orthologs, and perform other auxiliary analyses. Following this, doubletrouble^15^ was released as an R package to classify and identify gene duplications. It works on a framework similar to DupGen_finder, identifies duplicate pairs, classifies them, calculates Ka/Ks ratios, but also goes one step ahead in helping identify retrotransposon derived duplicates and DNA transposon derived duplicates. Finally, SegmantX^16^ is another recent Python script that uses a novel approach of local alignment followed by a chaining algorithm to identify segmental duplications at the DNA level and also provides a docker container for ease of use.

From the varying tools discussed above, the following aspects are evident - first, most of the gene duplication identification tools identify duplicates in pairs. But in a biological context there can be arrays of duplicates, and identifying them all and maintaining the link between them is crucial for further evolutionary understanding. Next, most tools have been adapted for use on standardized data from databases like Ensembl and NCBI, limiting the use cases on widely varying bioinformatics data formats. Given the heterogeneity of General Feature Format (GFF) files from different sources, and databases, it is necessary to come up with tools that can work across different GFF file flavours. Furthermore, many tools focus on either identifying large scale duplications and characterizing relevant evolutionary events or classifying duplications. But it is also important to integrate expression analysis, to understand the functional divergence of identified duplicates and also study the nature of the identified duplicates by adding details on genomic order conservation with synteny, in cases of comparative genomic studies between two species. Finally, from a biological perspective, different species have different duplication landscapes in their genomes. However, existing methods work based on defined thresholds necessitating the use of species-specific thresholds^2^.

In this context, DupyliCate is presented here as a solution for gene duplication array identification, and broad classification of the duplicates. It provides options for downstream expression analysis of the identified duplicates to understand the expression divergence, and gene-level Ka/Ks computations implemented independently as a part of the script, to infer selection pressures. It has been deployed here on datasets from NCBI, Phytozome, and varying additional sources, to demonstrate its ability to process widely varying input file formats. It also leverages BUSCO-based metrics to determine species-specific thresholds for gene duplicates identification, and offers the users a wide range of parameters that can be fine-tuned according to the objectives at hand.

## Results

### Benchmarking results

DupyliCate was compared against the tools doubletrouble and DupGen_finder. The results of DupyliCate runs using the default BUSCO-based threshold, and user-specified cutoffs for singleton-duplicate segregation, were different from one another as well from the other tools compared (Table 1). The run with user-specified cutoffs (a) gave gene duplicate numbers comparable to the results output by both doubletrouble and DupGen_finder. In this baseline run with the DupyliCate-specific parameter flags turned off, the number of gene duplicates classified by DupyliCate (a) as tandems and proximals is comparable to that output by doubletrouble, while DupGen_finder gives a lower number of tandem and proximal duplicates. When comparing the total number of duplicates, DupyliCate and DupGen_finder have close estimates while doubletrouble gives a lower number of total duplicates. Next, the BUSCO-based threshold run that was chosen for the actual benchmarking. This run inferred a self-normalized bit score cutoff of 0.696 from the BUSCO analysis and resulted in approximately two-thirds of the genes being classified as singletons. The numbers of tandem duplicates are similar in the DupyliCate and DupGen_finder results, while doubletrouble reported more tandem duplicates. Proximal and dispersed duplicates are lower in the DupyliCate output, while both of the other tools reported similar numbers. Singleton genes showed the maximum difference across the three tools. Both doubletrouble and DupGen_finder have remnant unclassified genes, while DupyliCate classified all the input genes.

**Table 1:** Consolidated results of benchmarking analysis on the *A. thaliana* Col-0 dataset. ‘Misc.’ gene duplicates in the table indicate all gene duplicates that are not tandem and proximal duplicates.

| Tool | Tandem | Proximal | *Misc. | Singleton | Non-recurrent tandem and proximal duplicates | Non-recurrent total gene duplicates | Non-recurrent total duplicates and singletons | Total no. of coding genes input for classification | Total no. of unclassified genes |
| --- | --- | --- | --- | --- | --- | --- | --- | --- | --- |
| DupyliCate (a) (baseline check run) | 3584 | 1530 | 20092 | 3829 | 4844 | 23615 | 27444 | 27444 | 0 |
| DupyliCate (b) (default BUSCO-based threshold run for benchmarking) | 1756 | 705 | 5387 | 19909 | 2374 | 7535 | 27444 | 27444 | 0 |
| doubletrouble | 3045 | 3581 | 22224 | - | 4970 | 22869 | 22869 | 27444 | 4575 |
| DupGen_finder | 1316 | 2978 | 20921 | 19 | 4294 | 23340 | 23359 | 27444 | 4085 |

When compared to the other tools, DupyliCate additionally produces a duplication landscape plot from the distribution of the self-normalized bit scores in the genome of a sample analyzed for gene duplications (described in workflow sub-section ‘Step 5: Threshold determination for singleton-duplicate segregation and self-alignment’). The plot of *A. thaliana* Columbia-0 (Col-0) is shown in Fig. 1. More details on the plot interpretation are presented in the discussion section.

**Figure 1:**
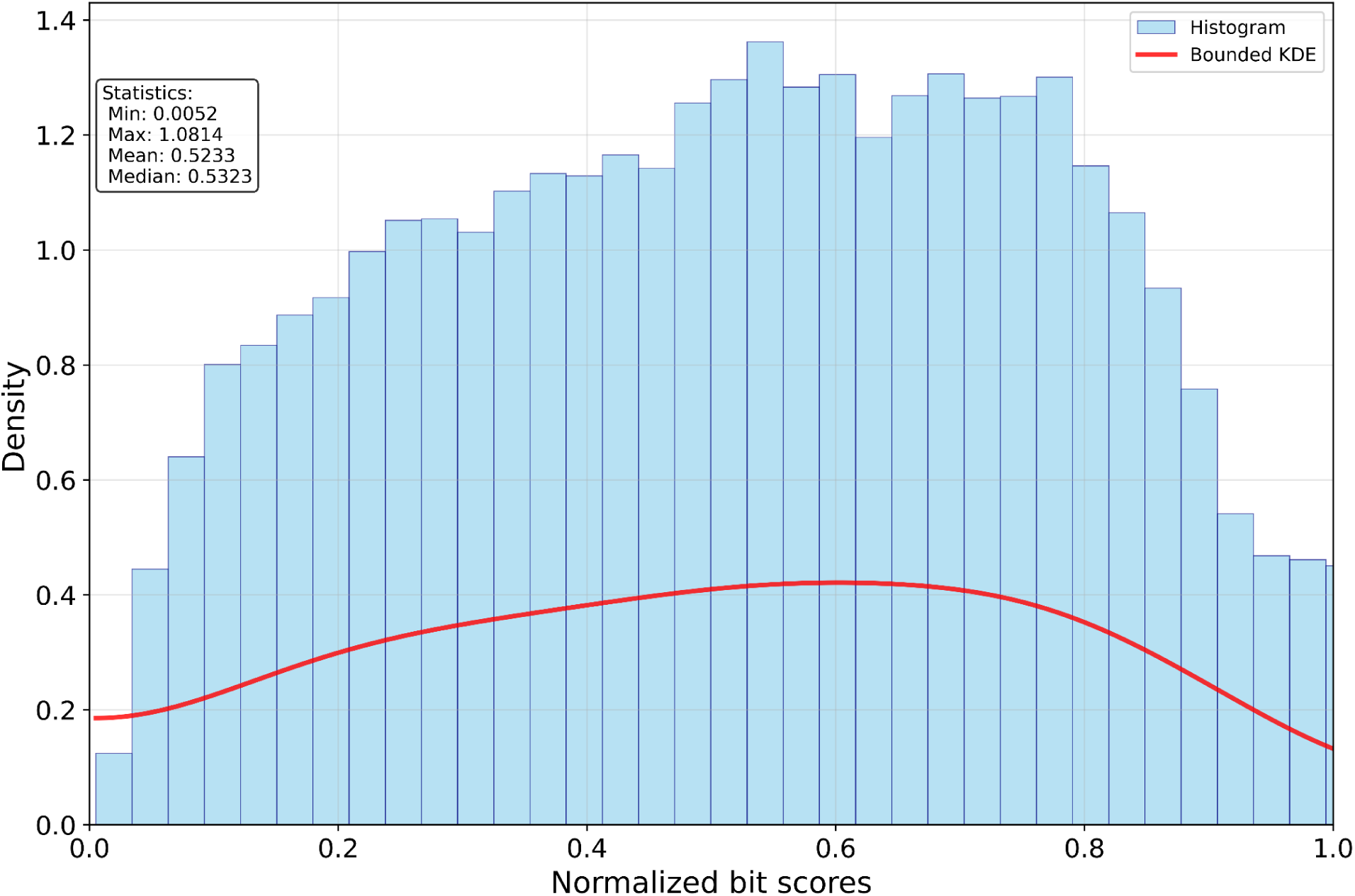
Duplication landscape plot of *A. thaliana* Col-0 output by DupyliCate - shows the distribution of self-normalized bit scores from the sequence alignments. The plot can help get an idea of the genome-wide duplications in the sample being analyzed and can also help decide optimal normalized bit score thresholds in case a user would like to set them manually without using the auto option. Left and right skewed distributions in the plot could suggest genomes with extremely low gene duplications and genomes with extensive gene duplications or polyploidization, respectively. KDE stands for the Kernel Density Estimate. It is a smoothed, continuous probability density curve estimated from the discrete histogram data. The KDE reveals the underlying distribution shape independent of the histogram bin width, enabling clearer detection of skewness and multiple peaks.

In terms of runtime performance, doubletrouble and DupyliCate without BUSCO had comparable runtimes while DupGen_finder and DupyliCate with BUSCO had comparable runtimes (Fig. 2).

**Figure 2:**
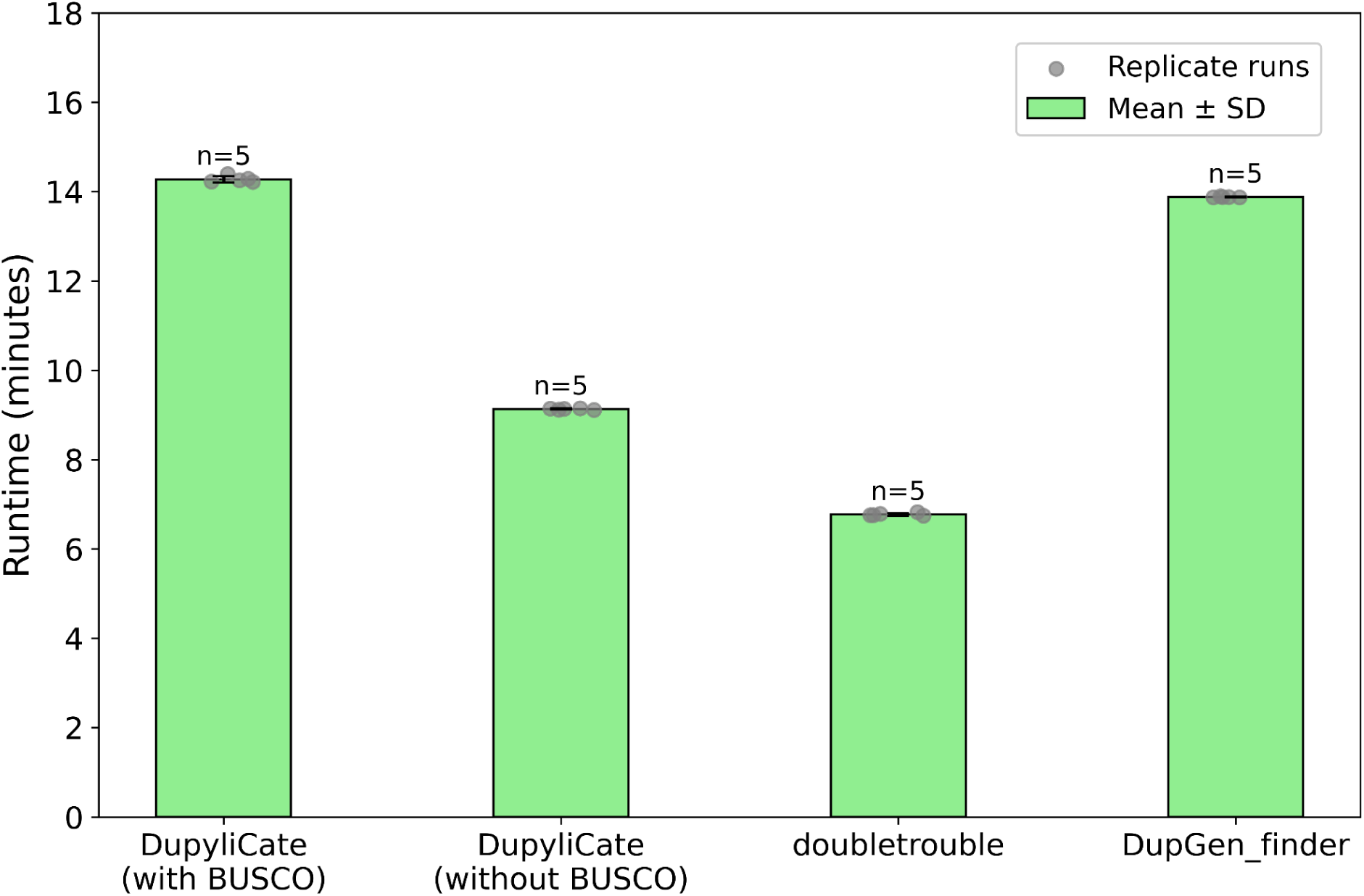
Runtime comparison across DupyliCate, doubletrouble, and DupGen_finder

### Runtime analyses

The different parameters tunable in DupyliCate were explored in different runs of the tool using *A. thaliana* Col-0 dataset on an Ubuntu 24.04 LTS machine with 30 cores. For runs involving comparative analysis with a reference, *A. thaliana* Niederzenz-1 (Nd-1) was used as the sample and *A. thaliana* Col-0 was used as the reference. All the runs were performed using the default BUSCO-based thresholding method. The runtimes for the different runs are shown in Table 2.

**Table 2:**
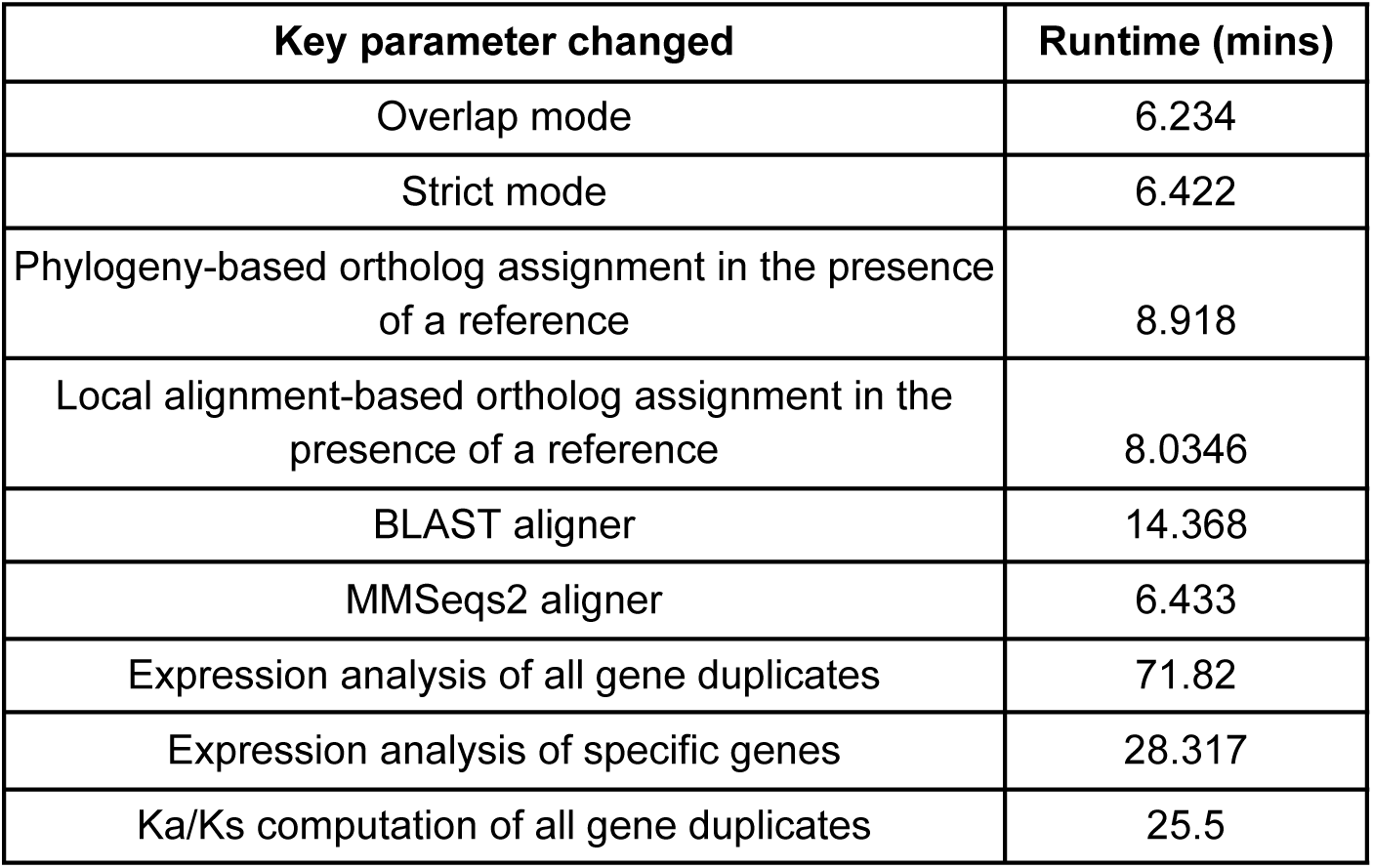
Runtime compilation when using different modes and parameters of DupyliCate on *A. thaliana* datasets

The specific gene expression analysis of the *Rapid Alkalinization Factor* (*RALF*) array^17^, *Xeroderma pigmentosum* (*XPB*), and *β-L arabinofuranosidases* (*BLA*) duplicates resulted in the plots shown in Fig. 3. Subfunctionalization can be achieved through expression in different parts of the plant or under different stress conditions. One such example of subfunctionalization is the spatially different gene expression patterns of the subgroup 7 myeloblastosis (*MYB*) transcription factor (TF) gene copies - *MYB11*, *MYB12,* and *MYB111*, in *Arabidopsis thaliana* seedlings. Each of these TFs, *MYB11*, *MYB12,* and *MYB111,* are expressed in specific domains of the seedling, roots, and cotyledons, respectively^18^. However, the expression pattern of the *XPB* duplicates shows a correlation with both copies being active (Fig. 3(b)). This correlation in expression suggests functional redundancy. DupyliCate would require tissue-specific expression data to discover a subfunctionalization (Supplementary Table S4) that has been previously reported by Masuda and co-workers^19^. Whereas, the expression plot of the *BLA* duplicates indicates a potential neofunctionalization fate, where only one of the duplicate copies is highly expressed while the other shows very low expression (Fig. 3(c))^20^. Neofunctionalization is defined as the fate where one of the copies diverges and develops a new function. While the low expression of one of the gene copies can be attributed to pseudogenization as well, it is important again to note that overall expression is considered here and spatial gene expression information is not included. But the experimental evidence from the work of Tao and co-workers shows that the copy *AT5G12960 is pollen-specific* while the other copy *AT5G12950* is ubiquitously expressed in all other tissues^20^. This supports the hypothesis that *AT5G12950* was the original copy while *AT5G12960* diverged after duplication and acquired new tissue-specific functions not found in the original copy leading to their fate being called neofunctionalization. The correlation analysis of these duplicates computed with DupyliCate also indicate expression divergence (Supplementary Table S4) and the low overall expression of *AT5G12960* supports its pollen-specific function. Further, the gene expression plots can be obtained in the compact mode as shown in Fig. 3, as well as matrix mode wherein the genes in a duplicate array are arranged in the form of a matrix and the respective expression plots are displayed at the intersection of the correct gene pair in the matrix representation (Supplementary Figure 11). Moving on, the runtime trajectory analyses were performed using the Brassicales plant datasets used for the *FLS* case study. The runtime trajectory for duplicates mining (reference-free) (Fig. 4(a)), and duplicates mining combined with orthology assignments for comparative gene copy number analysis (with phylogeny) showed a linear time increase trend (Fig. 4(b)). The runtime for the large-scale study involving plant datasets from Phytozome with *A. thaliana* as the reference took 34.8 hours to complete. This demonstrates the suitability of DupyliCate for high-throughput use cases involving gene duplicates mining as well as comparative genomic studies needing orthology assignments with respect to a reference.

**Figure 3:**
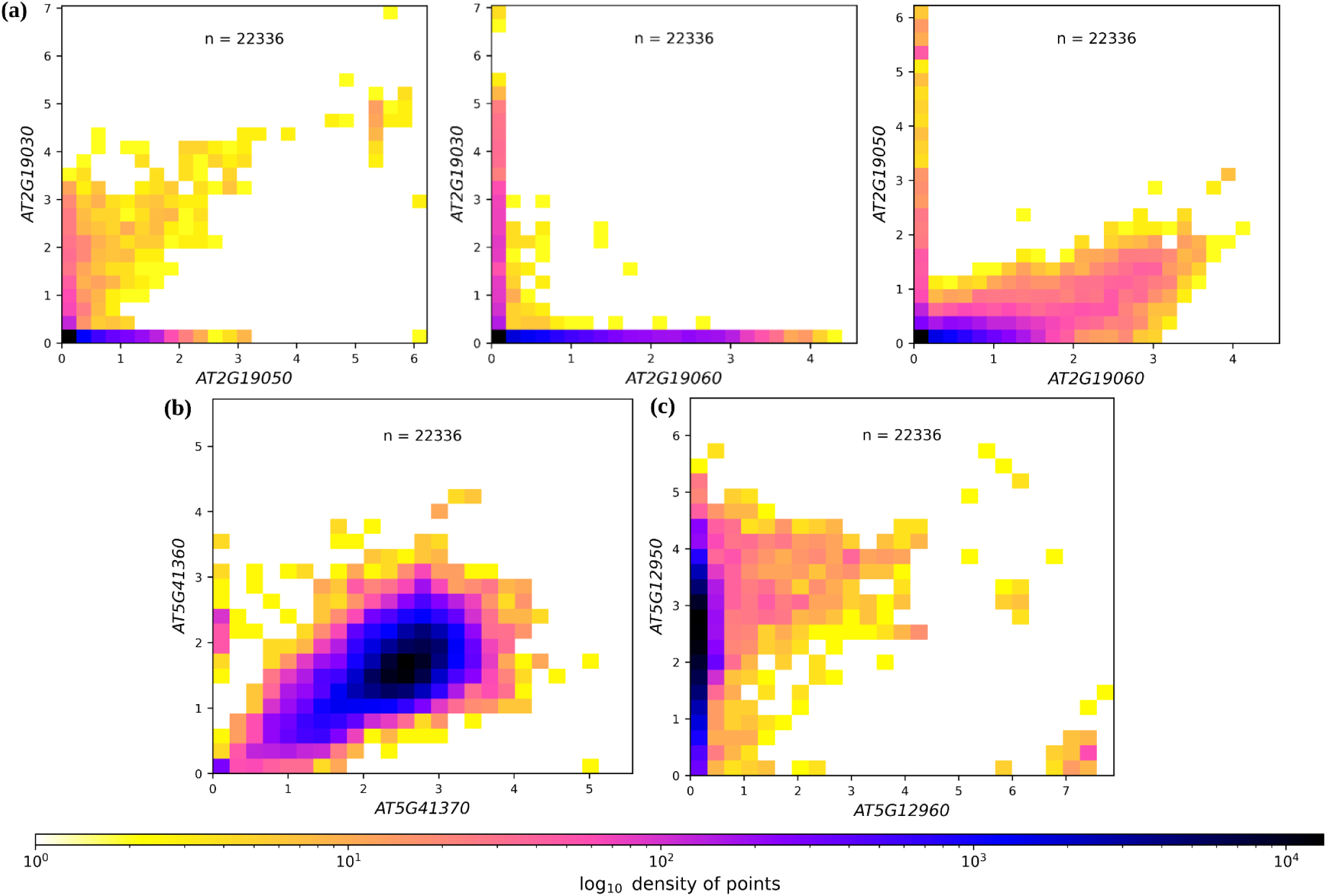
Pairwise gene expression plots of *A. thaliana* Col-0 tandem genes encoding (a) Rapid Alkalinization Factors (b) Xeroderma pigmentosum, and (c) β-L arabinofuranosidases. ‘n’ represents the number of samples used to create the gene expression plot. The x and y axes show the gene expression in log(1+TPM). TPM stands for Transcripts Per Million.

**Figure 4:**
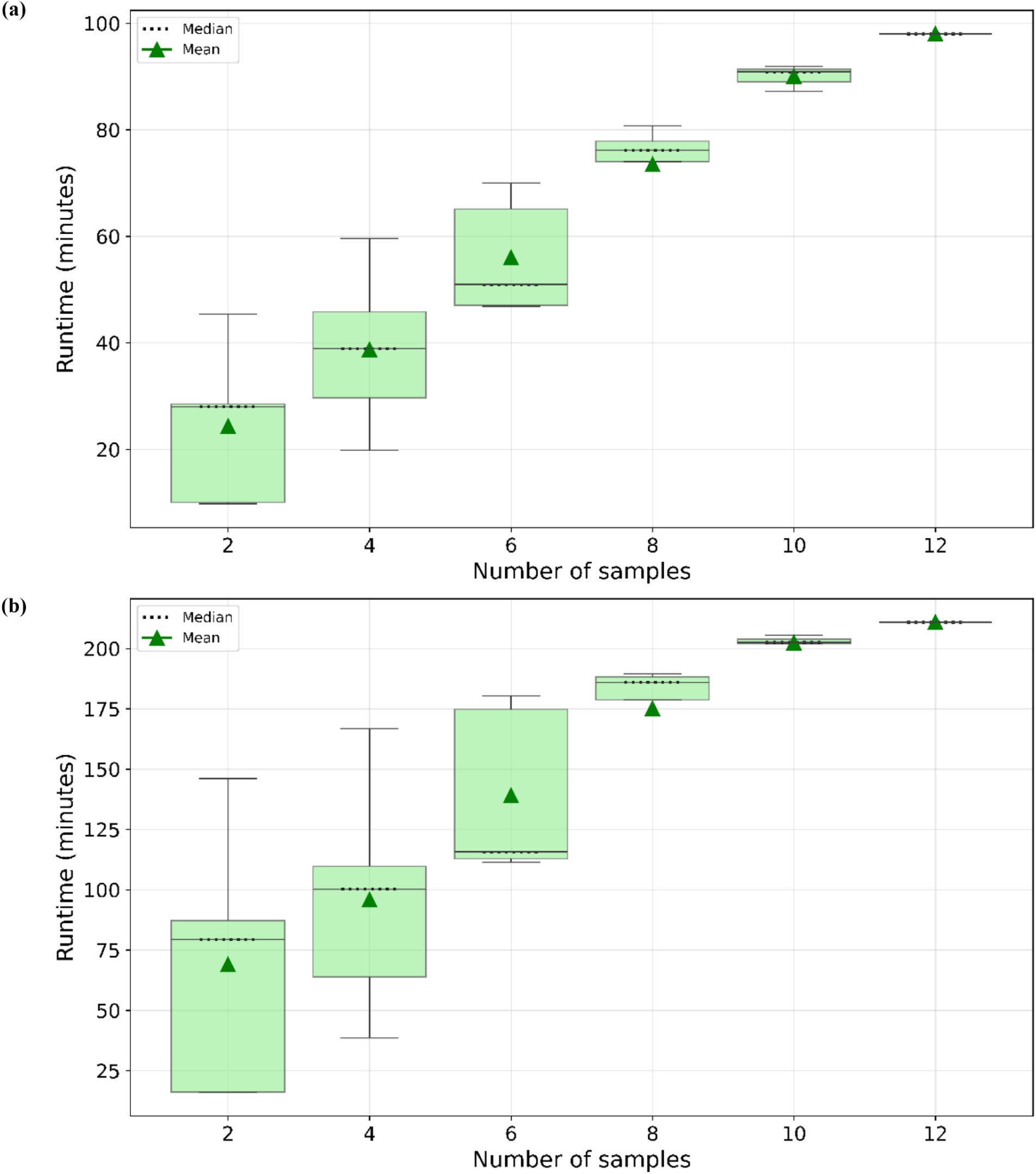
Runtime trajectory analysis for (a) Gene duplicates mining across twelve Brassicales genome sequences (b) Gene duplicates mining, and comparative genomics involving ortholog assignments across twelve Brassicales genome sequences with respect to the reference *A. thaliana*.

### Proof of Concept analyses

Proof of Concept (POC) analyses of DupyliCate were performed using well-known gene duplications reported in different plants. DupyliCate correctly identified tandem duplication of the *SEC10* gene in *A. thaliana* Nd-1 with respect to *A. thaliana* Col-0^21,22^. Next, tandem duplication of a gene in *Gardenia jasminoides* with respect to *Coffea canephora*^23^ was correctly mined as a tandem duplicate array of four genes. A proximal duplication of genes was detected in the nematode tolerance locus of the nematode tolerant sugarbeet line StrubeU2Bv whereas the susceptible line KWS2320 had just a single copy of the gene^24^. DupyliCate confirmed the dispersed duplication of the sterol Δ^24^-isomerase (24ISO) gene in *W. somnifera* responsible for withanolides biosynthesis^25^. These genes were also found to show up as copies along with the closely related sterol side chain reductase (SSR) genes in a proximal-dispersed gene duplicate group in the duplicate relationships file.

GeMoMa activating flag in the tool produced annotations for *V. amurensis* and *V. rotundifolia* with *V. vinifera* as the reference, and helped identify the gene duplicates for species with missing annotation files. *V. amurensis* had the highest number of gene duplicates (Table 3). Among the analyzed rice and related genomes, *Echinochloa colona* and *Echinochloa oryzicola, which* are rice weeds, had the highest number of gene duplicates. All the other rice varieties and their relatives had a comparative number of duplicates (Table 3). The copy number table (Supplementary Table S5) gave a visual presence-absence matrix of the orthologs of the different stress genes in *O. sativa* indica across the analyzed rice relatives.

**Table 3:**
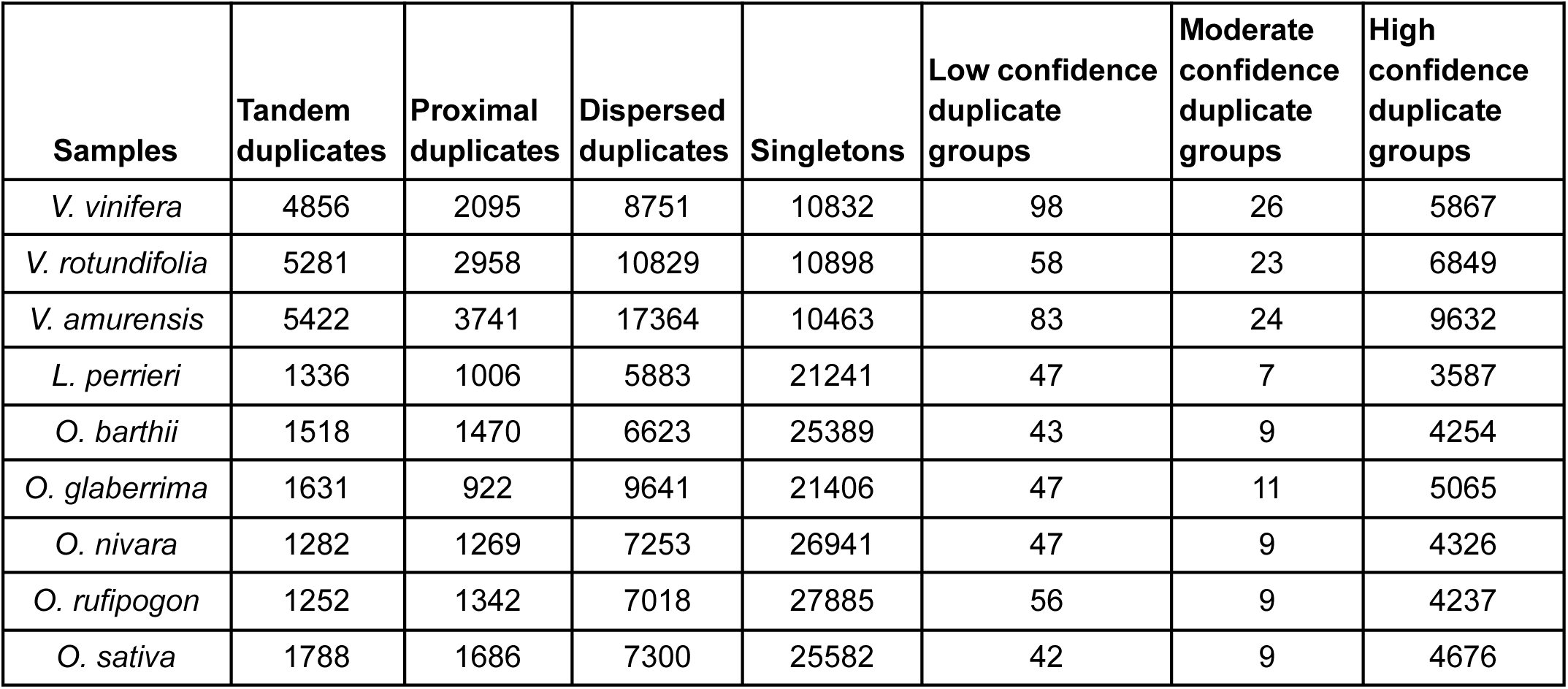
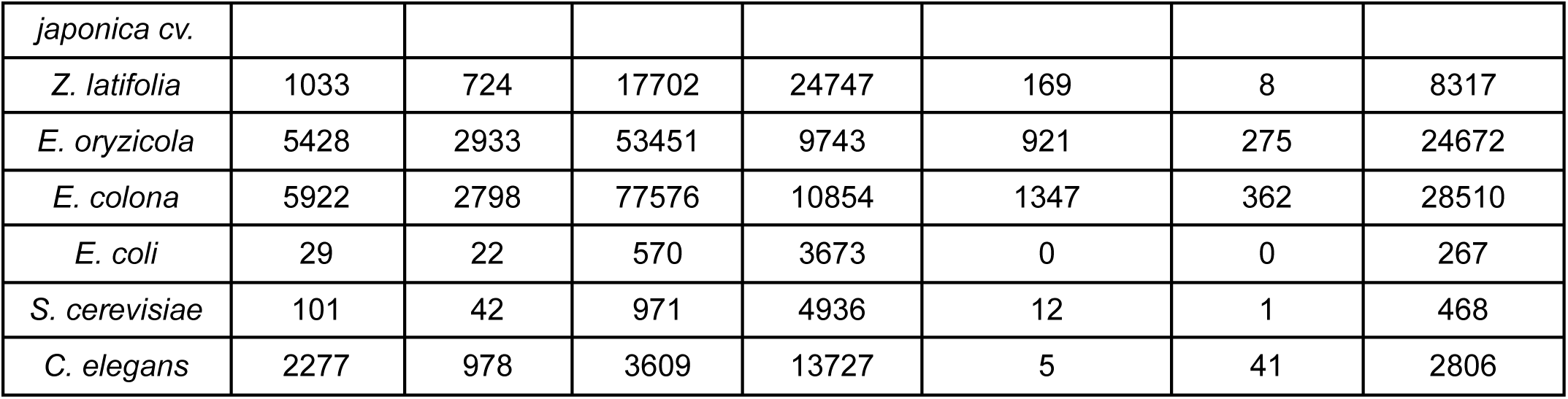
Consolidated results of gene duplicates identified and classified in *Vitis* species, *Oryza* species, rice relatives, and non-plant samples

| Samples | Tandem duplicates | Proximal duplicates | Dispersed duplicates | Singletons | Low confidence duplicate groups | Moderate confidence duplicate groups | High confidence duplicate groups |
| --- | --- | --- | --- | --- | --- | --- | --- |
| *V. vinifera* | 4856 | 2095 | 8751 | 10832 | 98 | 26 | 5867 |
| *V. rotundifolia* | 5281 | 2958 | 10829 | 10898 | 58 | 23 | 6849 |
| *V. amurensis* | 5422 | 3741 | 17364 | 10463 | 83 | 24 | 9632 |
| *L. perrieri* | 1336 | 1006 | 5883 | 21241 | 47 | 7 | 3587 |
| *O. barthii* | 1518 | 1470 | 6623 | 25389 | 43 | 9 | 4254 |
| *O. glaberrima* | 1631 | 922 | 9641 | 21406 | 47 | 11 | 5065 |
| *O. nivara* | 1282 | 1269 | 7253 | 26941 | 47 | 9 | 4326 |
| *O. rufipogon* | 1252 | 1342 | 7018 | 27885 | 56 | 9 | 4237 |
| *O. sativa* | 1788 | 1686 | 7300 | 25582 | 42 | 9 | 4676 |
| *japonica cv.* |  |  |  |  |  |  |  |
| *Z. latifolia* | 1033 | 724 | 17702 | 24747 | 169 | 8 | 8317 |
| *E. oryzicola* | 5428 | 2933 | 53451 | 9743 | 921 | 275 | 24672 |
| *E. colona* | 5922 | 2798 | 77576 | 10854 | 1347 | 362 | 28510 |
| *E. coli* | 29 | 22 | 570 | 3673 | 0 | 0 | 267 |
| *S. cerevisiae* | 101 | 42 | 971 | 4936 | 12 | 1 | 468 |
| *C. elegans* | 2277 | 978 | 3609 | 13727 | 5 | 41 | 2806 |

DupyliCate also identified gene duplicates and singletons in the non-plant species - *E. coli, S. cerevisiae, and C. elegans*. As expected, *E. coli* and *S. cerevisiae* had a low number of gene duplicates whereas *C. elegans* showed a relatively high number of gene duplicates (Table 3).

### FLS evolution in Brassicales

Among the Brassicales plants analyzed in the *FLS* (flavonol synthase) case study*, I. tinctoria, E. vesicaria, S. alba* and *B. napus* showed a high number of gene duplications. Tables consolidating the gene duplicates mined and the orthologs and copy numbers of the *Arabidopsis thaliana FLS* (*AtFLS*) genes in the other Brassicales plants are included in Supplementary Table S3 and Supplementary Table S6, respectively. The presence-absence variation in the copy number table showed the presence of *FLS1* - the functional *FLS* in all the Brassicales plants analyzed. But all of the Brassicales plants were missing at least one or more of the other *FLS* genes. The phylogenetic tree used to validate the ortholog assignments is shown in Fig. 5. Most of the ortholog assignments in DupyliCate and the clade grouping in the phylogenetic tree match, except for a few sequences (coloured with red strips in the tree (Fig. 5). The *C. papaya* gene identified as *FLS1* (evm.TU.supercontig_37.77) does not exactly group in the *AtFLS1* clade but is a sister to this clade. No other *FLS* genes were identified in *C. papaya.* The *B. oleracea* gene that groups in the *AtFLS2* clade (Bol019362.v1.0) shows up as an *FLS1* ortholog in DupyliCate. On a similar note, a group of genes marked by the red strips in the *FLS4* clade in the tree are assigned as orthologs of *FLS1* or *FLS3* in DupyliCate results. A consistent occurrence of this pattern among these genes that form a separate subclade with respect to *AtFLS4* is interesting. The phylogenetic tree also shows a high number of sequences in the *FLS3* and *FLS4* clades, indicating a potential expansion of these genes in Brassicales.

**Figure 5:**
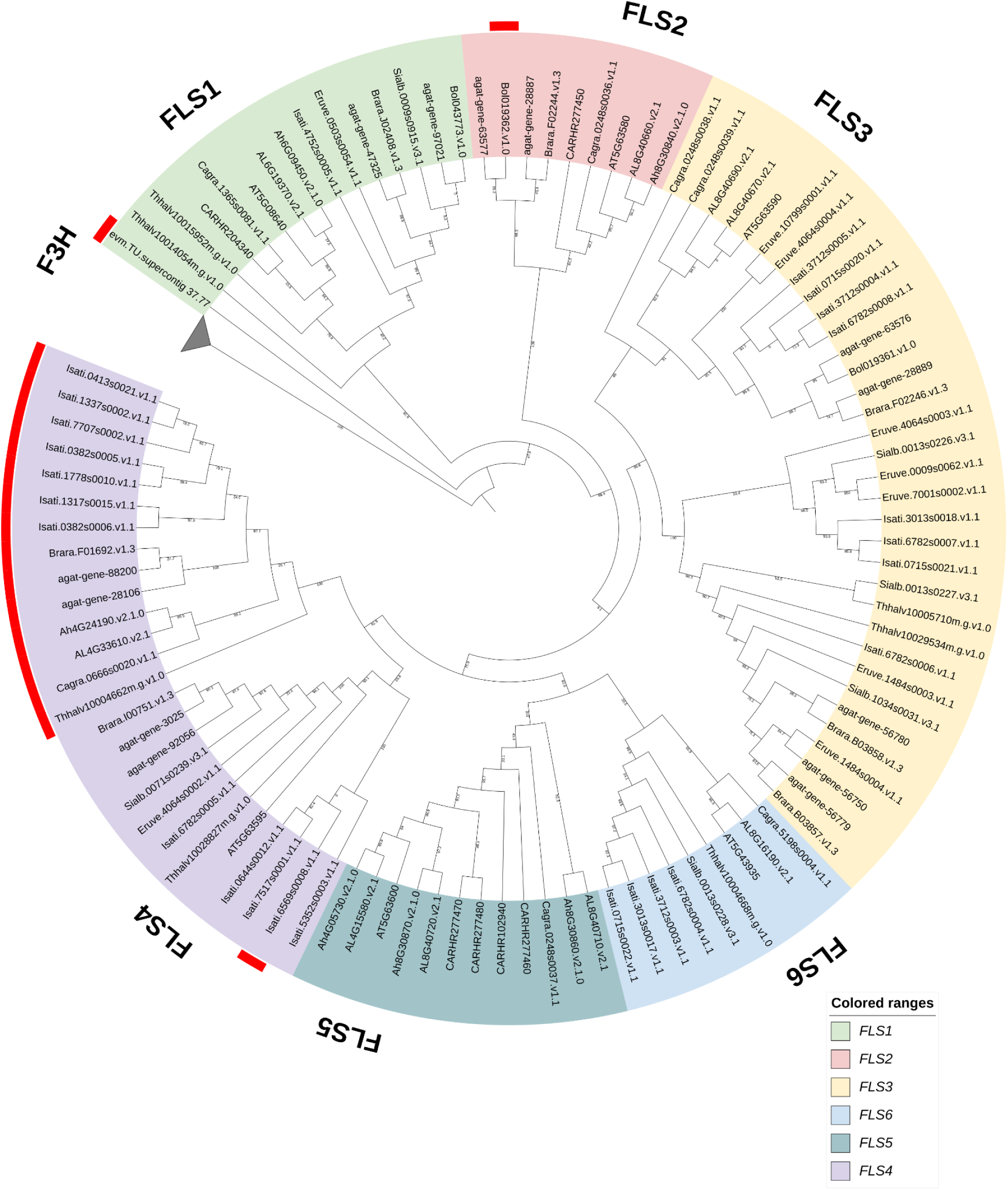
Phylogenetic analysis of *FLS* genes across twelve Brassicales plant genomes. The red striped sequences have different ortholog assignments in DupyliCate and the phylogenetic tree. The collapsed clade contains flavanone 3-hydroxylase (F3H) sequences as outgroups

### MYB evolution across a broad phylogenetic range of plant species

A large-scale analysis of plant datasets from Phytozome facilitated the understanding of *MYB* (Myeloblastosis) transcription factors - *AtMYB12,* and *AtMYB111* evolution across a wide range of plant species. A summary of the number of gene duplicates mined across the 153 plant species along with the copy number table representing the *A. thaliana MYB12* and *MYB111* orthologs across the analyzed datasets were obtained (Supplementary Table S7). Among the analyzed plants, orthologs for *MYB12* and *MYB111* were not found in algae, mosses, liverworts, and ferns. Interestingly, the *MYB12,* and *MYB111* orthologs were found in lycophytes (club mosses). In the other plants, orthologs were found for at least one or both of *AtMYB12* and *AtMYB111*. The phylogenetic tree used to validate the ortholog assignments is shown in Fig. 6. The phylogenetic analysis also reveals an interesting evolutionary pattern of *MYB12* and *MYB111*. As seen in Fig. 6, *AtMYB12* and *AtMYB111* fall in separate clades that consist of sequences from *A. thaliana* and other dicot plants (pink and purple clades in Fig. 6). Monocots, a few other dicots, and some basal angiosperms on the other hand do not show separate lineages of these flavonol biosynthesis-regulating MYBs.

**Figure 6:**
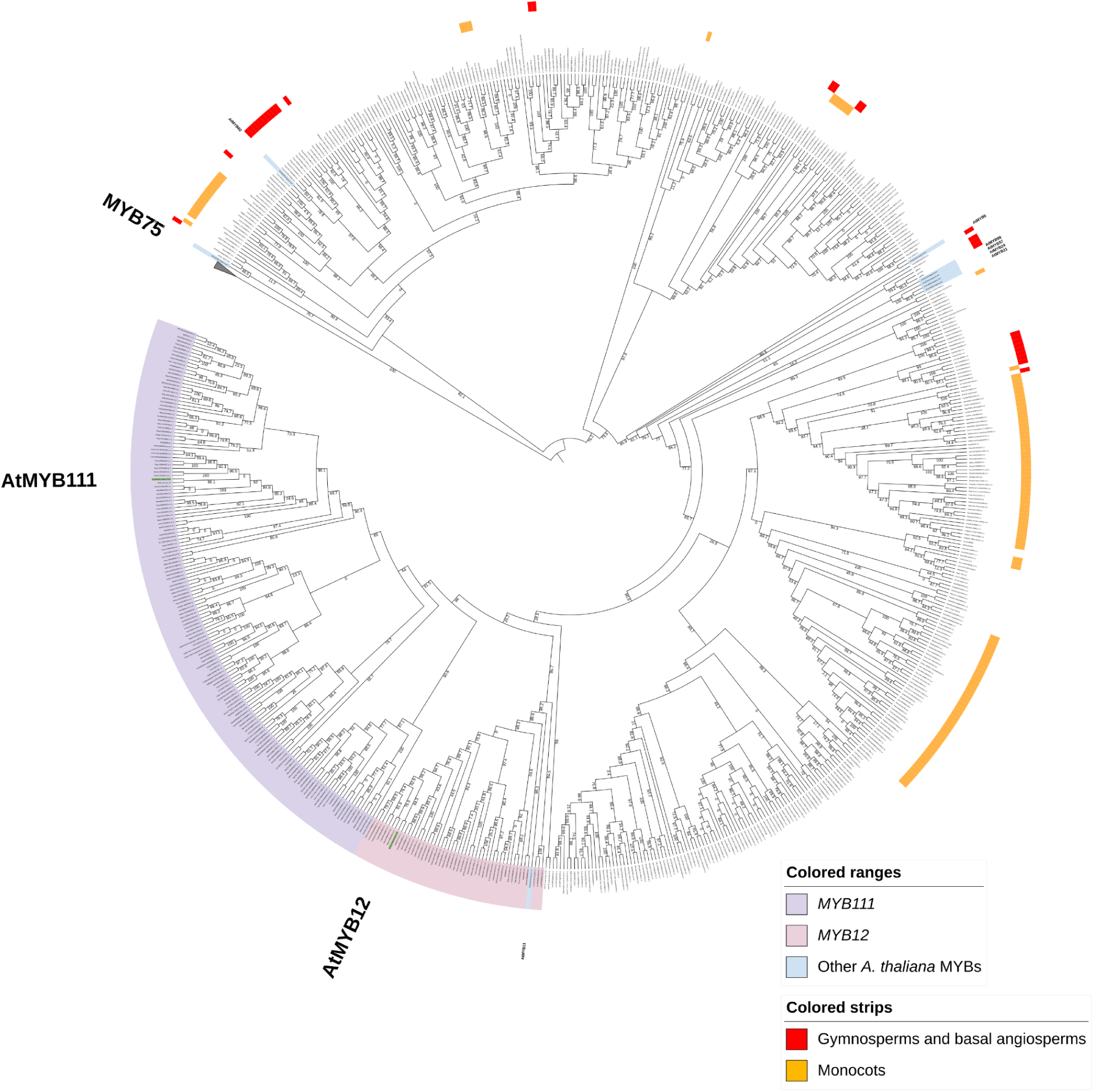
Phylogenetic analysis of *MYB12* and *MYB111* genes across a broad phylogenetic range of plants. All the sequences not covered by the yellow strips belong to dicots. The collapsed clade contains *MYB75* sequences as outgroups.

Further, the tree revealed that there are *MYB111* and *MYB12* orthologs captured correctly, as well as false positive sequences (uncoloured sequences bounded by the outgroup MYB75 clade and the *A. thaliana* flavonol MYB clade coloured in blue). The ortholog assignments in DupyliCate and the clade grouping in the tree were compared (Fig. 7).

**Figure 7:**
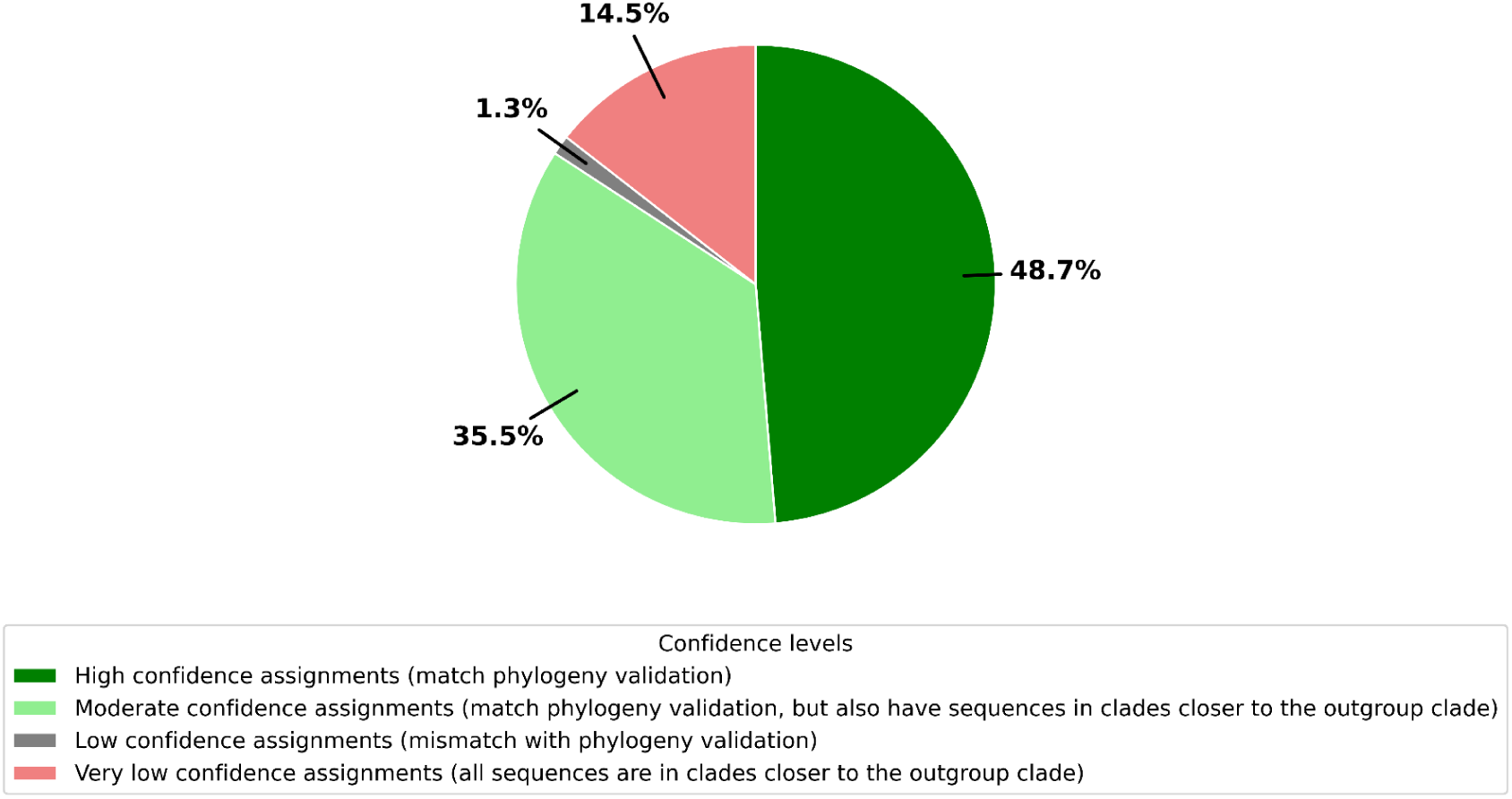
Pie chart representing the validation of DupyliCate’s ortholog assignments with respect to clade grouping in the phylogenetic analysis

The validation analysis shows that nearly 50% of the ortholog assignments in DupyliCate matched exactly with the phylogenetic tree. More details on the validation analysis can be found in Supplementary Methods.

### Parameter tuning results

The parameter tuning analyses revealed that --score, --self_simcut, and --seq_aligner are the top sensitive parameter flags that influenced the final results the most (Supplementary Table S10). A parameter sensitivity ranking table was created based on these tuning trials and a total of five parameter flags were found to be the most sensitive influencers of the final results (Supplementary Table S11).

The BUSCO vs manual threshold comparative analyses were performed and the results were consolidated in Supplementary Table S12. While the total number of gene duplicates were higher in the manual threshold method for *A. thaliana*, *D. purpurea,* and *D. dumetorum*, for *V. cruziana*, the BUSCO threshold-based method gave more gene duplicates. But the overall proportion of gene duplicates and singletons in the BUSCO-based threshold approach corresponded well with the respective duplication landscape plots of the different plant species (Supplementary Results 1).

## Discussion

The baseline run (a) (Table 1) with the DupyliCate flags turned off, gave comparable gene duplicates output to doubletrouble^15^ and DupGen_finder^9^, with DupyliCate identifying more singleton genes and classifying all input genes unlike the other tools. This confirmed that the core gene duplicate classification framework is consistent with the compared tools, with DupyliCate’s singleton gene identification approach helping classify all protein-encoding genes in the genome. The benchmarking run (b) (Table 1) with default BUSCO-based normalized bit score threshold classified nearly two thirds of the *A. thaliana* genes (∼19900) as singletons, with DupGen_finder giving as few as only ∼20 singletons and doubletrouble not having a separate singleton classification. Further, in the benchmarking run results, the number of tandem duplicates in DupyliCate were comparable to that of DupGen_finder, but the number of proximal and dispersed duplicates were lower than that of the other tools, which had comparable numbers. The difference in DupyliCate’s benchmark run can be explained by the use of BUSCO-derived threshold as a biological boundary, to obtain singleton and duplicate gene spaces that are species-specific, which is a fundamental design distinction from the other tools. Also, DupyliCate classifies all input genes into one of singleton or duplicate categories, while DupGen_finder and doubletrouble have a remnant number of unclassified genes (Table 1). doubletrouble does not specify or produce a singleton output file, hence it can be explained that it inherently considers the genes not assigned to the duplicate groups as singletons. DupGen_finder on the other hand, outputs only those genes that do not have self-local alignment hits as singletons. This ignores genes that hit up only against themselves in the local self-alignment. In the manual threshold specification to DupyliCate, genes that do not have self hits, genes that have only themselves as self hits, and genes whose second best hit within the genome fail to pass the threshold are classified as singleton genes, resulting in a few thousand singleton genes. But when using a BUSCO-based threshold for obtaining gene duplicates, genes that do not have self hits, genes that have only themselves as self hits, and genes whose second best hit within the genome fail to pass the BUSCO-derived threshold are classified as singleton genes. This causes the results to significantly differ giving around 27.5 % of gene duplicates in *A. thaliana*. The relatively lower number of gene duplicates in *A. thaliana* is supported by a low BUSCO duplication percentage (Supplementary Table S2). A potential biological explanation could be that only a certain proportion of genes were retained as duplicates after the polyploidization events reported in *A. thaliana*^26^. These estimates are also close to the estimate of 28% gene duplicates in *A. thaliana* previously reported^27^ after the most recent alpha polyploidy event (Supplementary Figure 1)^26,28^. This puts forth the biological implication of the BUSCO-based thresholding method that distinguishes genes that escaped ancient duplication events and those that were retained in gene families or underwent more recent duplications. It thus allows for species-specific gene duplicates identification and classification. But it is also important to note that there is no clear boundary between singleton genes and duplicated genes. Both are rather extremes on a gradient. Therefore, the identification is imperfect, influenced by the cutoffs and other selection criteria determined by the users. Further the different number of singleton gene outputs in the different existing tools could as well be attributed to the fluidity of the singleton-duplicate segregation criteria. However, the BUSCO-based approach developed in this study offers one possible way of segregating singleton and duplicated genes with a reliable biological support. Additionally, the duplication landscape plot of *A. thaliana* output by DupyliCate (Fig. 1) also offers a visual inference on the genome-wide duplication status. Left and right skewed distributions in the plot could suggest genomes with extremely low gene duplications (as seen for *E. coli, S. cerevisiae* (Supplementary Results 1)) and genomes with extensive gene duplications or polyploidization (Supplementary Results 1), respectively. The approximately central, bell-shaped curve for *A. thaliana* (Fig. 1) corresponds well with the polyploidization-gene duplications loss events described for *A. thaliana*^27,29^. Looking at the performance metrics of the three tools (Fig. 2), DupGen_finder had the highest runtime, while doubletrouble had the lowest runtime. However, since an outgroup was specified in DupGen_finder, the inclusion of a forward alignment step leads to an increase in runtime. Thus doubletrouble and DupGen_finder can be considered to be on par with respect to runtimes. Since DupyliCate incorporates FASTA cleaning, GFF validation, and duplicate group clustering steps, the runtime is slightly longer, especially when BUSCO analysis is included. Also steps like isoform removal and execution of local alignment, which need to be performed separately by the user prior to running doubletrouble and DupGen_finder, are integrated in DupyliCate’s run. Although this increases the runtime, it could lead to possibly reduced hands-on time when using DupyliCate. Moreover, it is expected that the availability of DupyliCate via docker and conda would further simplify the installation and use of the tool, and expand its usability for users with different levels of bioinformatics expertise.

The runtime analyses involving different parameter and mode selection with the *Arabidopsis* datasets give an idea of the runtimes for the different functionalities. As seen from Table 2, a complete expression analysis, and Ka/Ks analysis of gene duplicates take longer. Hence, when such analyses are intended, it is recommended to use as many cores as available to complete the run in the shortest possible time. The expression analysis plots output by the use of the specific duplicate analysis flag in the absence of a reference (Fig. 3) help analyze the functional fates of the duplicated genes and determine expression divergence among them. The matrix pattern of the plot helps visualize the gene expression trends among all the genes in the same duplicate group at once. The entire RALF tandem duplicates array reported by Cao and Shi^17^ - *AT2G19020, AT2G19030, AT2G19040, AT2G19045,* was identified in the RALF tandem group by DupyliCate along with the genes *AT2G19050* and *AT2G19060.* However, *AT2G19020, AT2G19040*, and *AT2G19045* were found to be perceived (putative) pseudogenes based on the expression analysis. Thus Fig. 3(a) shows the pairwise expression plots of the other genes in the identified tandem duplicates array. But it is important to note that including information on the spatial aspects of gene expression in the different tissues and under different conditions is necessary to confidently ascertain the subfunctionalization, neofunctionalization, or pseudogenization fates of the gene duplicates. From Fig. 4(b) it is evident that dataset size and inclusion of phylogeny in the presence of a reference, are factors that influence the runtime. Large datasets and inclusion of phylogeny for ortholog assignment in the presence of a reference lead to an increase in the runtime. But incorporating phylogeny during ortholog assignments will produce better ortholog assignments, as local alignment-based approaches alone are not enough to identify orthologs correctly^6^. Hence phylogeny is recommended when a reference is used. In case the user just wants to have a first hand assessment of gene copy numbers with comparative analysis, runtime can be prioritized over accuracy by deactivating the phylogeny analysis.

The *SEC10* gene is incorrectly collapsed in the Col-0 genome sequence, but is properly represented in the *A. thaliana* Nd-1 assembly and annotation^21,22,30^. This was confirmed through the DupyliCate analysis where the *SEC10* gene duplication was identified as a tandem duplication in the Nd-1 dataset compared to TAIR (Col-0). The first dedicated gene in the crocine biosynthesis pathway of *Gardenia jasminoides* is a part of a four-gene tandem array that shares just one ortholog with the *Coffea canephora* genome. This tandem duplication-led expansion of the gene cluster is responsible for the initial critical steps in crocine biosynthesis that is not found in coffee^23^. The entire tandem cluster was correctly identified by DupyliCate demonstrating the usefulness of representing gene duplicates in cluster groups instead of gene pairs. In the analysis of nematode resistance locus between tolerant and susceptible sugarbeet lines^24^, DupyliCate identified that the crucial genes from the locus belong to a proximal duplicate cluster. This shows that proximal duplication was responsible for the nematode tolerance phenotype seen in the tolerant line StrubeU2Bv while exemplifying the application of DupyliCate to answer biological questions. Finally, the 24ISO gene responsible for the first steps of withanolide biosynthesis in *W. somnifera* was identified to be present in a dispersed duplicate group. It was also found to appear in the duplicates relationships results along with the closely related SSR genes^25^ providing a PoC for the usefulness of the duplicates relationships file in the overlap mode and the mixed duplicates file in the strict mode. These different PoC analyses show the ability of DupyliCate to correctly identify gene duplicates and classify them into the different duplication classes. The gene duplicates identification in the *Vitis* species lacking annotation was facilitated through the GeMoMa flag of DupyliCate (Table 3). This option could be useful in cases where the samples of interest lack structural annotation or a good quality structural annotation, but have a closely related species with a good structural annotation that can serve as the hint for gene prediction with GeMoMa. It is also helpful to note that the hint for GeMoMa annotation and the reference for comparative analysis with DupyliCate could be the same or different. This means, the GeMoMa flag can be used in the presence as well as in the absence of a reference for DupyliCate analysis, but requires the specification of a hint for creating the annotation for the sample. In either case, the species whose annotation is to be used by GeMoMa for the new annotation must be specified in the GeMoMa config TXT file along with the names of the samples to be annotated. More details are provided in the GitHub repository. In the rice genome sequence analysis, the rice weeds showed a high number of gene duplicates (Table 3). The BUSCO analyses (Supplementary_Table_S2) showed that these rice weeds have a high percentage of duplicated BUSCO genes (>80%), and the pseudo-ploidy numbers (apparent indicators of ploidy of the sample based on BUSCO gene frequencies) of *E. colona* and *E. oryzicola* were ∼3 and ∼2, respectively. Their duplication landscape plots (Supplementary_Results_1) were also distinctively right skewed suggesting a recent polyploidization event. The other rice varieties had a comparable number of gene duplicates and showed a very similar trend of duplication landscape plots except *Oryza glaberrima.* The plot of *O*. *glaberrima* was almost flat except for a marked high in the number of genes with a self-normalized bit score of 1 indicating that the duplicates in the genome are recent or have not diverged greatly (Supplementary Results 1). DupyliCate’s applicability to non-plant samples was shown by mining duplicate genes in *E. coli, S. cerevisiae*, and *C. elegans*. The high number of gene duplicates identified in *C. elegans* corroborates the widespread occurrence of gene duplications in *C. elegans*, when compared to other non-plant eukaryotes like yeast and fruit fly^31^.

Among the analyzed Brassicales plants for the *FLS* case study, *I. tinctoria, E. vesicaria*, and *S. alba* had a pseudo-ploidy number of ∼2 indicating their diploidy, while *B. napus* with a pseudo-ploidy of ∼3 was classified as triploid (Supplementary Table S2). This explains their high gene duplication numbers. Coming to the ortholog assignments in the copy number analysis, *C. papaya FLS1* (evm.TU.supercontig_37.77) groups as a sister of the *AtFLS1* clade and is the only *FLS* ortholog identified. It also has a good BUSCO completeness (>80%) (Supplementary Table S2) indicating a biological phenomenon. *C. papaya* belongs to Caricaceae, one of the most basal families in Brassicales^32^. This along with the phylogenetic tree (Fig. 5) suggests a possible hypothesis on *FLS* evolution in the Brassicales - the true *FLS* had appeared in the early Brassicales plants followed by an expansion of the *FLS* gene family in Brassicaceae after the divergence of Caricaceae and Brassicaceae. The *B. oleracea* gene showing up in the *AtFLS2* clade (Bol019362.v1.0) could probably be an ancient *FLS2* duplicate that cannot be placed correctly only based on local alignment methods. Similarly, almost all the sequences falling in the later subclade of *AtFLS4* (coloured by the red strips in *AtFLS4* clade in Fig. 5), have *AtFLS4* appear as only the fourth or the fifth best forward hits of these genes in the forward local alignment. These genes could also be ancient duplications of *FLS4* and hence difficult to assign orthologs based on candidates from similarity based methods. Such difficult cases of ortholog detection can be resolved with higher phylogenetic resolution by including more sequences that give better placement and context to the query gene within its gene family supported by better taxon sampling and global alignment cues^33^. Further, from the phylogenetic analysis, there is an evident expansion of the *FLS3* and *FLS4* lineages happening in Brassicaceae, which is quite intriguing. This is also observed in the *FLS* phylogenetic tree presented by Schilbert et al.^34^. Expression analyses performed in DupyliCate, found a majority of genes in the *FLS3*/*5* clades to be pseudogenes. In the *FLS4* clade most of the *I. tinctoria* genes turned out to be pseudogenes. A potential explanation for the *FLS3/4* expansion could come from the expression analysis. Since these are non-functional *FLS* genes, they could be under relaxed selection pressure. Expansion might be tolerated because additional copies do not impose strong fitness costs, allowing them to accumulate mutations and gradually become pseudogenes.

The transcriptional regulation of flavonol biosynthesis in plants is achieved by the action of *MYB11, MYB12*, and *MYB111* transcription factors (TFs)^35^. Schilbert and Glover analyzed the evolution of flavonol regulators in the family Brassicaceae and reported the duplication of *MYB111* and *MYB12* TFs in the family^35^. The MYB analysis presented here serves as a case study of these flavonol regulators’ evolution across a large phylogenetic range of plants. Flavonoid biosynthesis is well known and established in land plants. Flavonols, a class of flavonoids that play a major role in protecting against abiotic stress factors like UV exposure, are also widespread in land plants^36–38^. These factors provide a basis for the reason behind the absence of flavonol regulating *MYB12* and *MYB111* orthologs in non-land plants and early land plants like mosses. The *MYB12,* and *MYB111* orthologs mined in club mosses were examined and were found to belong to very low confidence ortholog groups, which could mean that these species could also lack the *MYB12* and *MYB111* orthologs. An alternative explanation could also be that algae, mosses, and other basal plants might have other flavonol biosynthesis regulating MYBs. Looking at the other plants in the analyses, all assignments that had very high confidence orthologous group confidence scores (OGCS) in DupyliCate were found to match exactly with the phylogeny results. While groups with moderate and low OGCS were found to have mismatches with the tree or have false positive sequences. This demonstrates the usefulness of the OGCS scoring system developed that can help refine and obtain more reliable results. Further, nearly half of the examined plant species’ ortholog assignments are exact matches with the tree with nearly one third having matches with the presence of false positive sequences (Fig. 7). One possible reason for the ortholog misassignments observed could be the widely varying evolutionary distance differences between the reference *A. thaliana* and the analyzed sample species. This is also supported in the phylogenetic tree where most of the sequences near the outgroup clade belong to either gymnosperms, basal angiosperms, monocots, or polyploid plants like cotton. The phylogenetic tree (Fig. 6) also provides some biological insights. It indicates the duplication of *MYB12* and *MYB111* in some dicots (purple and pink clades) that could have been derived from an ancestral flavonol regulating MYB which is still observed in gymnosperms, basal angiosperms, monocots, and some dicots. In conclusion, the MYB case study demonstrates the usability of the tool for large scale analysis involving gene duplicates mining, gene copy number analysis in nominal time, and its ability to provide biological insights.

Further, the parameter tuning analyses helped analyse the effect of the different settings controlled by the user. As seen in Supplementary Table S10, --score and --self_simcut parameters influenced the results significantly. As the values of both these parameters were gradually increased, the number of identified gene duplicates lowered, while a decrease in these cutoffs increased the gene duplicate numbers identified due to the relaxed thresholds. Next, the type of sequence alignment tool used, influenced the results with an albeit low difference between the different runs. Since the duplicate and singleton segregation relies on the self-normalized bit score cutoff, differences in the bit scores put out by the different aligners can lead to slightly different results. Nevertheless, considering the low runtime and small difference between the gene duplicate numbers reported by the three aligners, the default DIAMOND aligner was found to work well. The next most important parameter is the --proximity flag that determines the maximum number of intervening genes between duplicates for them to be considered proximal duplications. As expected, when the proximity value was increased, the number of proximal duplicates increased and the number of dispersed duplicates decreased. The parameter --evalue is the third most sensitive parameter and it slightly influences the total number of singletons and duplicates identified. Low --evalue cutoffs led to slightly more gene duplicates being identified while higher cutoffs lowered the number of gene duplicates identified (Supplementary Table S11). The other parameter flags in the table --fwd_simcut, --scoreratio, --hits, --occupancy, --synteny_score, --flank, and --side do not impact the overall gene duplicates singletons segregation and classification. This is because all these parameter flags are relevant for forward local alignment in case a reference is provided, and hence play a role in finding orthologs for the sample gene duplicates in the reference organism. There is a detailed discussion on adjusting these parameters for ortholog detection in the presence of a reference organism along with some suggestions for parameter tuning of DupyliCate, in the Supplementary Discussion.

Moving on to the BUSCO vs manual threshold run results, there is an increase in gene duplicates for manual mode with *A. thaliana*, *D. purpurea*, and *D. dumetorum* and a decrease in gene duplicates in the manual mode with *V. cruziana*. This can be explained by looking at the manually set normalized bit score cutoff used and the BUSCO-based normalized bit score cutoff used. The BUSCO-derived self-normalized bit score threshold for *A. thaliana*, *D. purpurea*, *D. dumetorum*, and *V. cruziana* were 0.696, 0.726, 0.822, and 0.430, respectively. Since the first three species have a higher threshold when compared to the manual threshold of 0.5, the number of gene duplicates were lower in the BUSCO mode and higher in the manual mode due to the lowering of the cutoff. However, the scenario for *V. cruziana* is inverted since its species-specific threshold is lower than 0.5, explaining the decrease in the gene duplicate numbers in the manual mode due to an increase in the cutoff. The duplication landscape plots of these four species (Supplementary Results 1) reveal that *A. thaliana* has an almost bell-shaped curve, while the other three species show a right skewed curve, with *V. cruziana* having sufficient number of genes with low normalized bit scores as well. The shape of *A. thaliana*’s plot combined with duplication loss events reported in the species conform well with the low number of gene duplicates identified, as discussed before. The lower normalized bit score threshold for *V. cruziana*, combined with the almost similar proportions of gene duplicates and singletons, explain the right skewed shape of the curve with a significantly populated tail representing a good number of singletons too. The duplication plots for the other two plant species - *D. purpurea*, and *D. dumetorum* correspond well with the high proportion of gene duplicates identified. Thereby, these results show the applicability of DupyliCate to a wide taxonomic range of species with the parameters contributing to species-specific results that are reproducible.

In conclusion, DupyliCate is a high-throughput Python script developed for gene duplicates identification, classification, and integrated expression analysis. It has a number of adjustable parameters, species-specific singleton duplicates segregation, flexibility to handle a variety of GFF files, synteny analysis of small-scale duplicates (in the presence of a reference) and optional phylogeny-based ortholog detection for comparative gene copy number analysis. DupyliCate can thus be a valuable addition to the bioinformatics toolkit to support research in the domains of gene duplications research and comparative genomics for both plant and non-plant species.

## Materials and methods

### Datasets

Datasets of *Arabidopsis thaliana* Col-0 and Nd-1^22^ were used to demonstrate the available modes, different aligner options, possible Ka/Ks analysis and expression analysis functionalities of DupyliCate. The two accessions were also used to demonstrate the PoC for *SEC10* duplication in the *A. thaliana* Col-0 dataset. *Gardenia jasminoides* and *Coffea canephora* datasets^23^ were used to demonstrate the PoC for tandem duplication of the first gene in crocine biosynthesis in *G. jasminoides* with respect to *C. canephora.* Datasets of the *Beta vulgaris* lines StrubeU2Bv and KWSONT2320Bv^24^ were used as PoC for proximal duplication of the nematode tolerance gene in the tolerant line StrubeU2Bv with respect to KWS2320. Datasets of *Withania somnifera* and *Physalis pruinosa*^25^ were included to show PoC for dispersed duplication of the 24-Isomerase gene in *W. somnifera* with respect to *P. pruinosa*. DupyliCate was used on three *Vitis* datasets (*V. vinifera, V. rotundifolia,* and *V. amurensis*) to demonstrate applicability on complex genome sequences and also to demonstrate the use case of GeMoMa feature of the tool using the datasets of *V. rotundifolia* and *V. amurensis* that lacked structural annotation. A collection of 10 genome sequences of rice (*Oryza sativa*) and close relatives from the RiceRelativesGD database^39^ were leveraged to demonstrate the performance of DupyliCate on monocot plants. Gene duplicates were mined in *O. sativa indica* variety and its relatives. The stress related genes in *O. sativa indica* sp. were used to create a copy number table to analyse the number of the corresponding orthologs in the other rice relatives. *Escherichia coli*^40^*, Saccharomyces cerevisiae*^41^, and *Caenorhabditis elegans*^42^ datasets were used to show PoC for deployment on non-plant species.

### Application case studies

*Arabidopsis thaliana* is reported to have six copies of the flavonol synthase (*FLS1-6*) gene encoding a key enzyme in flavonol biosynthesis^43^. Of the copies, only *AtFLS1* is known to produce a catalytically functional protein whereas the functions of the other gene copies are yet to be uncovered fully. Comparative analyses of the *FLS* gene family in the Brassicales order combined with expression analysis can help better understand the evolutionary dynamics of this gene. A total of 12 datasets of the Brassicales family (detailed in Supplementary Table S1), for which expression data were available, were chosen to do an application case study on flavonol synthase evolution in the Brassicales using *Arabidopsis thaliana FLS1-6* as reference genes.

Next, a large collection of 153 plant datasets was retrieved from Phytozome (details specified in Supplementary Table S1). These datasets were used to undertake a large-scale gene duplicates mining study across plant datasets to demonstrate the high-throughput utility of the tool. Further, these datasets were also used to study the evolution of the key flavonol regulator MYBs (Myeloblastosis genes) - *MYB12* and *MYB111* across the plant kingdom with *A. thaliana MYB12* and *MYB111* as reference genes, to demonstrate the biological relevance of DupyliCate’s results. The forward normalized bit score and forward similarity cutoffs were lowered to 0.2 and 30%, respectively, since the run involved samples quite distant from the reference *A. thaliana* to avoid missing potential orthologs in the samples.

The results of DupyliCate from the two application case studies were also validated by building a phylogenetic tree. For the FLS tree, flavanone 3-hydroxylase (F3H) sequences were used as outgroups. For the MYB tree the ortholog results from DupyliCate were visualized in a phylogenetic tree with anthocyanin regulating *MYB75* sequences as outgroups. Other closely related *A. thaliana* MYBs *AtMYB5, AtMYB11, AtMYB21, AtMYB24, AtMYB57, AtMYB99, AtMYB82*, and *AtMYB123* were also included in the tree. MAFFTv7.5.2.6^44^ was used with the G-INS-i accuracy-oriented method for alignment of the amino acid sequences. pxclsq from PHYX^45^ was used to remove alignment columns with occupancy below 10%. Phylogenetic trees were inferred using IQ-TREE3^46^ with the parameters -m MFP -wsr and --alrt 1000 -B 1000^47^. The best-fit evolutionary model inferred with ModelFinder^48^ for the FLS case study was Q.PLANT+R5 and for the MYB case study the best fitting model was JTT+R8. The phylogenetic trees were visualized using iTOL v7.2.2^49^.

All the datasets were analyzed with DupyliCate v1.0 in the overlap mode with BUSCO-specific threshold option activated. In case, a reference was used, phylogeny was used to determine the ortholog relationships with the reference genes.

### Workflow

A pipeline for gene duplicates identification, classification, and further downstream analyses was implemented in Python3 (Fig. 8). The overall workflow involves validation of input files followed by self-alignment of the sample(s) (sample(s) - species in which gene duplicates are to be identified). Species-specific thresholds are then used to segregate singleton genes from duplicates and duplicates are categorized as tandem, proximal, dispersed, and mixed (a class where genes can belong to more than one duplication class; this helps retain the relationship across the different duplicate groups). In the presence of a reference organism (reference - the species with respect to which the sample’s genes are assigned orthologs and synteny analysis is performed), forward alignment is performed. The classified gene duplicates are assigned to orthologs in the reference and synteny analysis is performed for small-scale duplicate groups of tandem duplicates and proximal duplicates. Depending on the mode chosen, a fourth category of mixed duplicates is returned. DupyliCate also facilitates optional gene expression analysis of gene duplicates. Further, it provides the user with the option to calculate Ka/Ks values of the duplicates. The DupyliCate script was tested with its dependencies and tools whose version numbers are specified in the workflow steps detailed below.

**Figure 8:**
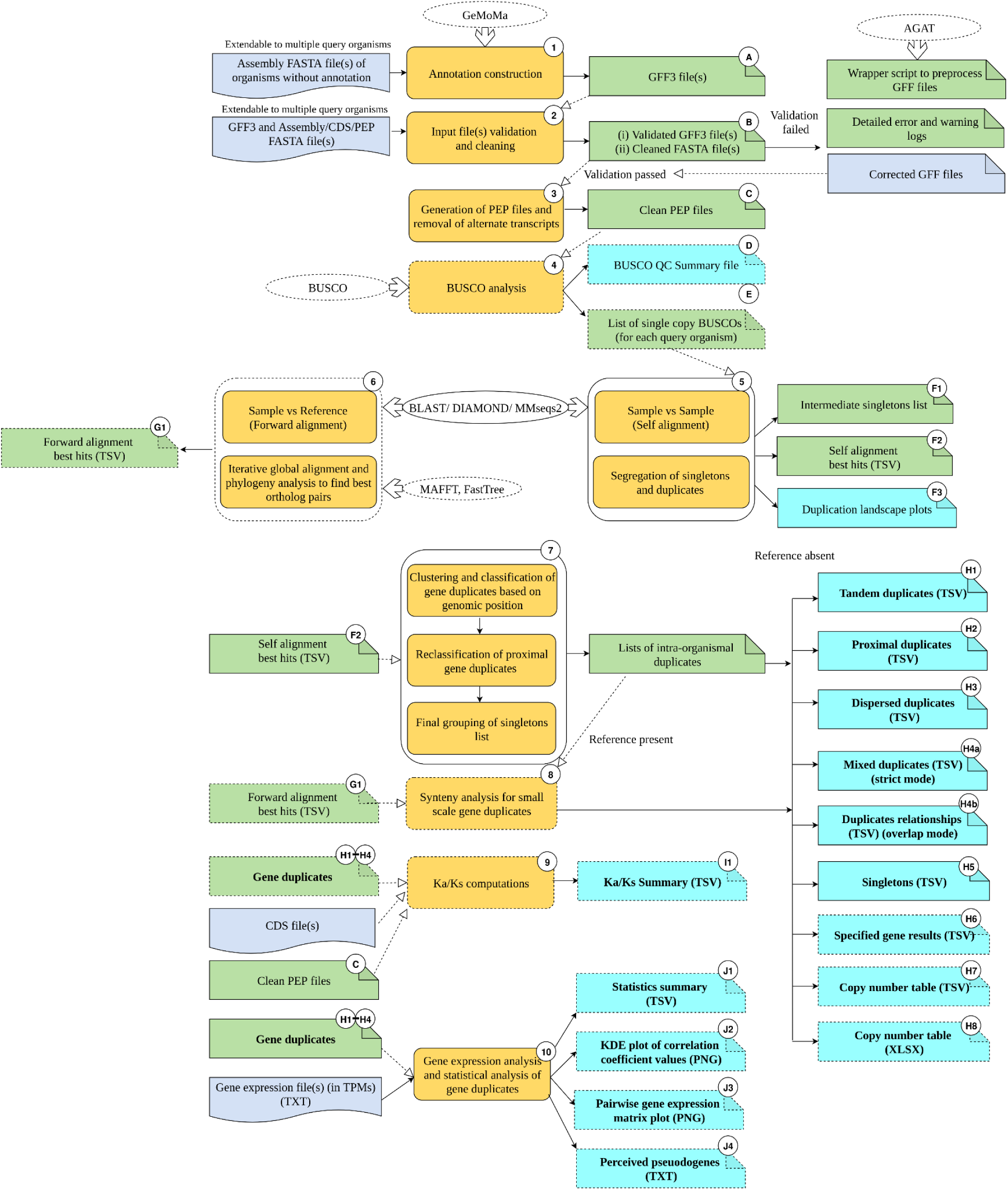
Schematic representation of DupyliCate’s workflow. The steps of the pipeline are numbered and the output files are alphanumerically labelled. The blue boxes indicate user provided input files and green boxes indicate intermediate output files produced by the pipeline that serve as inputs for further steps. Pipeline steps are colored yellow. Output files are shown by the cyan coloured boxes. External tools are represented by the ovals. Dotted outlines around the shapes indicate that they are not a part of the mandatory inputs, and dependencies or routine outputs of the pipeline.

**Step 1: Optional structural annotation:** DupyliCate requires the structural annotation file(s) (GFF3) along with one of the following (1) assembly sequence, or (2) coding sequences, or (3) polypeptide sequences in FASTA format. In case structural annotation is not available for a sample in which duplicates need to be analyzed, there is an option to provide the annotation of a related species as a hint for the gene prediction process with GeMoMa v1.9^50^.

**Step 2: Input file(s) check and validation:** The input files are first checked and validated before the actual run starts. If the check fails, the script exits and the errors are recorded and displayed sample-wise. In case of GFF errors, there is a helper script (AGAT_wrapper.py) provided along with the main script that uses AGAT v1.5.1^51^ to process and correct the GFF files.

**Step 3: GFF-FASTA files standardization, error correction, and polypeptide file generation:** GFF files have widely varying attribute fields denoting the information about the different genomic features. It is also necessary for the headers in the protein FASTA file to match correctly with the text denoted by these different attribute fields. This is a major limitation hampering the deployment of a number of existing tools on different GFF files. To address this issue, this work provides a two-step solution. First, it offers another helper script (FASTA_fix.py) for the users to process the FASTA file headers according to the attribute field text in the respective GFF files. Next, the main DupyliCate script offers an option to the user to indicate the specific GFF attributes to be used by the script when processing the GFF file of a particular species, through a simple config TXT file. This helps to circumvent GFF-FASTA mismatch errors, and expand the usability of DupyliCate on a number of GFF file flavours. The corrected and validated files then enter the main analysis and are processed to generate the FASTA files of polypeptide sequences without considering alternate transcripts.

**Step 4: Quality check of input polypeptide files:** Since the output depends on the quality of the input files and is also influenced by the ploidy of the samples being analyzed, a BUSCO^52^-based QC step is included. This provides detailed QC reports containing the BUSCO completeness, duplication, and BUSCO-derived pseudo-ploidy number that indicates the ploidy of the input sample(s).

**Step 5: Threshold determination for singleton-duplicate segregation and self-alignment:** Before moving on to the duplication analysis, it is important to segregate singletons and duplicates correctly. This in turn depends on the duplication landscape within each analyzed species. A new method for such species-specific threshold determination based on BUSCO v6.0.0 is implemented here. For this, two cutoffs - one based on self-normalized bit score (bit score of the hit of a polypeptide sequence with another polypeptide sequence within the genome (second best hit) relative to the bit score against itself) and self-similarity are offered. The complete BUSCO single copy genes are retrieved for each sample. If enough BUSCO single copy genes are present, the normalized bit score of the 95th percentile of the BUSCO single copy genes is picked as the threshold for duplicate and singleton identification. By default, this BUSCO-based auto threshold method is chosen. If BUSCO is not available or if the sample has a very low number of BUSCO single copy genes, then the fallback is to go for a self-similarity threshold instead of normalized bit score threshold. There is also an option to manually set the normalized bit score threshold and self-similarity threshold, allowing for fine grained analysis. The detailed steps of threshold determination can be found in Supplementary Figure 2.

Next, self-alignment of the sequences of a sample is performed. Options are provided to choose between DIAMOND v2.1.14^53^, BLAST v2.17.0^54^, and MMseqs2^55^ for the alignment step. Following self-alignment, duplicates and singletons are segregated using the previously determined threshold. Independent of the thresholding method, a plot folder called Duplication_landscape_plots is produced. It contains the normalized self-bit score distribution plots of samples. The overall pattern of the duplication landscape plot can be used to infer the genome-wide duplication status of the species.

**Step 6: Ortholog determination:** If sequences of a reference are provided, forward alignment of the sequences of each sample is performed against the sequences of this reference. Here again, two thresholds based on similarity value of the hits in the forward alignment and forward normalized bit score (bit score of a forward alignment hit of a polypeptide sequence relative to the bit score of the hit of the polypeptide sequence against itself in self-alignment) are applied to retain good quality forward hits for the further downstream steps. Next follows a comprehensive ortholog detection step in the presence of a reference. The top n candidates (default value of n=10) from local alignment algorithms like DIAMOND are subjected to a global alignment using MAFFT v7.5.2.6^44^, synteny analysis (the synteny analysis is performed by code developed as a part of DupyliCate and hence does not rely on external dependencies), and gene tree inference using FastTree2^56^. Phylogeny derived evolutionary distance assessments using DendroPy^57^ are utilized to obtain orthologs of genes in the samples among the reference genes. The detailed steps in ortholog detection are shown in Supplementary Figure 3. Since a multi-evidence approach is used for ortholog search, a confidence scoring scheme was developed to ascertain the confidence of each of the assigned orthologs based on the number of evidence points needed to determine orthology. The ortholog confidence scoring scheme is detailed in Supplementary Table S8.

**Step 7: Gene duplicates grouping and classification:** Next, the duplicates clustering and grouping step is performed per sample to give rise to the different gene duplicate groups shown in Fig. 9 below. There are two main modes of DupyliCate. Gene duplicates are classified as tandems, proximal, and dispersed duplicates in both modes. Differences arise between the modes in the following aspects -

**Figure 9:**
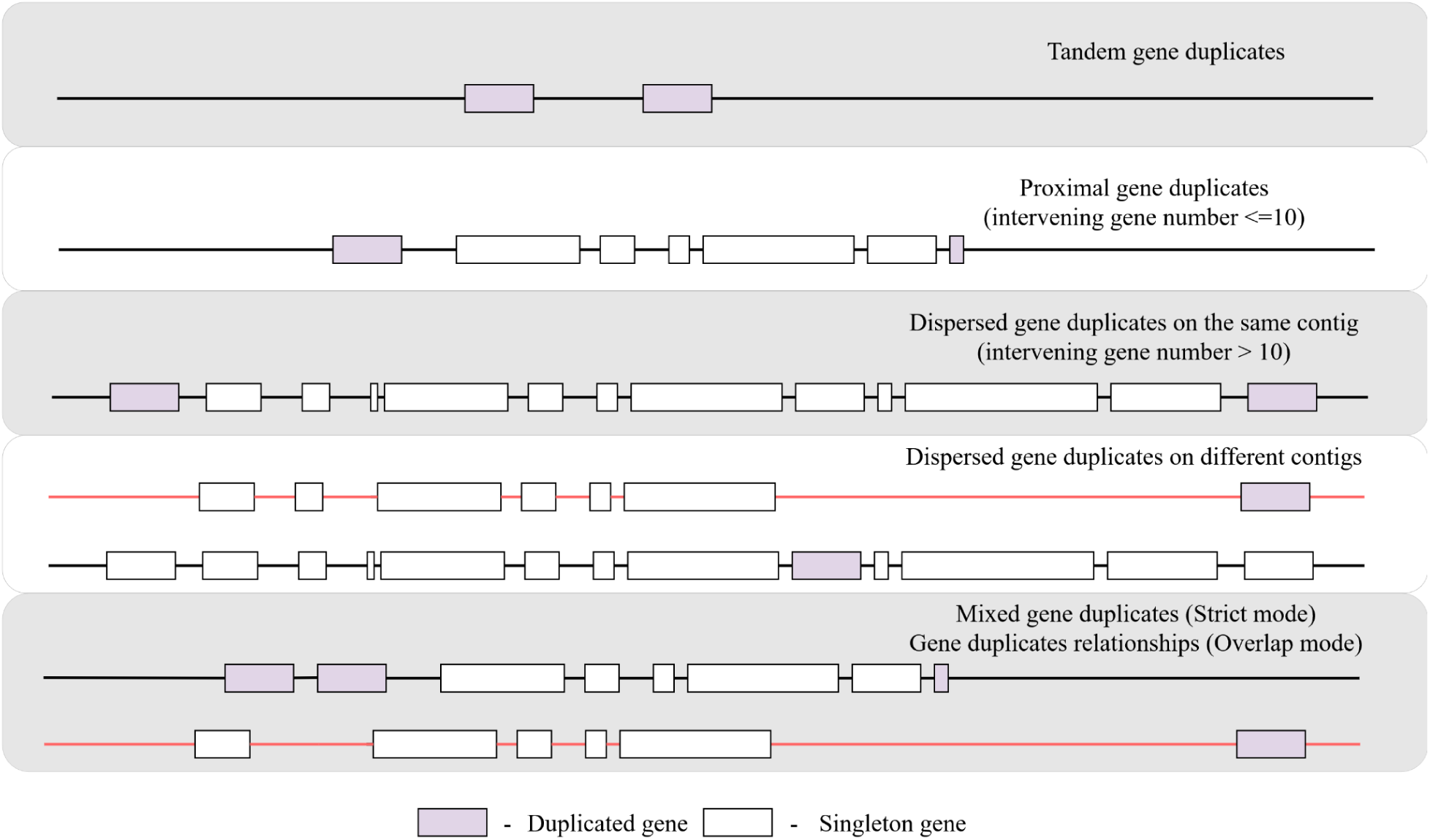
Different gene duplicate classifications utilized by DupyliCate. The red and black lines in the dispersed duplicates, mixed duplicates and gene duplicates relationships representations indicate different contigs/ scaffolds/ chromosomes.

**’Overlap’:** Some genes, repeat across groups and this mode also produces an output file called ’Duplicates_relationships’ that connects the different genes repeating across the duplicate groups

**’Strict’:** Gene duplicates do not repeat across groups and this mode produces a fourth class of duplicates type called ’Mixed_duplicates’ where the gene duplicates from different duplicate groups are merged together to retain their individual classification and also their relationship across the groups

The classification and clustering approach is detailed in Supplementary Figure 4. As a clustering approach is used to obtain gene duplicate groups/arrays, for all the duplicate groups in the output, there is an internal confidence scoring scheme used to classify the gene duplicate group as low, moderate and high confidence group (Supplementary Figure 5) that can help in interpreting the results accordingly. Along with the different duplicate group files, the singletons in a sample species are also provided per species.

In case the user has a set of desired genes in the reference whose copy number variation (CNV) needs to be assessed in the sample, a specific analysis flag option (--specific_genes) is available. Use of this flag produces two files. One is a comprehensive file providing information about the duplication and singleton status of the orthologs of the desired reference genes in the sample. Second is a copy number TSV file that provides the copy numbers of the specified reference genes across the different samples along with the orthologous group confidence score (OGCS) (OGCS scoring explained in Supplementary Table S9) to convey the confidence of the identified CNVs. This step also outputs a copy number XLSX file that has cells with no orthologs in the samples automatically colour coded in red, to allow for easy visual interpretation of presence-absence variation of the specified reference genes across the species analyzed.

**Step 8: Synteny analysis:** In the presence of a reference, synteny analysis is also carried out for small-scale gene duplicate groups of tandem and proximal duplicates, to add a confidence layer of genomic positional context with respect to the reference. If a reference is involved, detailed gene duplicate group nature details including gene group expansion and *de novo* duplication are provided for small-scale gene duplicate groups of tandem and proximal duplicates in the output. More information on the synteny analysis and gene duplicate group nature attributes have been described in Supplementary Figure 6 and Supplementary Figure 7, respectively. In the reference-free mode, ortholog detection, synteny analysis, and the gene group nature analysis steps are skipped.

**Step 9: Ka/Ks computation:** Optional downstream evolutionary analyses like Ka/Ks computation is performed for all gene duplicates on an individual gene basis using either Nei Gojobori (NG)^58^ or Modified Yang Nielsen (MYN)^59^ methods. These methods have been reimplemented from Ka_Ks_Calculator^10,60^. There is also in-built code facilitating amino-acid to codon-alignments and trimming of the multiple sequence alignments, thus removing the need for external tools in these steps. While an exact implementation of the NG method is available, the MYN method implemented is an approximation of the MYN method implemented in Ka_Ks_Calculator (Supplementary Figure 8 and Supplementary Results 1). Supplementary Figure 9 details the computational steps involved in the Ka/Ks analysis.

**Step 10: Gene duplicates expression analysis and statistical analysis**: If expression data is available for the sample(s), optional expression analysis of gene duplicates can also be performed by DupyliCate. This step gives out comprehensive information about the correlation among genes in duplicate groups, generates pairwise gene expression plots in a matrix figure and also helps determine divergent expression among duplicates as shown in Supplementary Figure 10. This step also outputs a list of perceived (putative) pseudogenes that have a low average expression value across samples. Since spatial information of gene expression in the different tissues is not involved in this analysis, genes with low average expression value across the samples are designated as perceived pseudogenes. The method used in this expression analysis was adopted and modified from the method implemented by Qiao and co-workers^9^.

### Benchmarking

*Arabidopsis thaliana* Col-0 and Nd-1 accessions were used to benchmark DupyliCate v1.0, DupGen_finder v1.0.0^9^, and doubletrouble v1.8.0^15^. DupyliCate was used in the overlap mode with *A. thaliana* Col-0 as the sample. DupyliCate additionally offers species-specific cutoff for segregating singletons and duplicates using BUSCO. Hence two runs of the tool were performed with *Arabidopsis thaliana* Col-0 dataset using the default BUSCO-based cutoff method, and user-specified cut-off method. There are no equivalent parameters for the normalized bit score and self similarity flags in DupGen_finder and doubletrouble. Therefore, a truly parameter equivalent comparison is not possible. In order to perform a baseline check for the consistency of results and core classification logic across the three tools when using only the local alignment-based filtering, in the user-specified cutoff run, 0 was specified as cutoff for both normalized bit score and self similarity parameter flags., The same e-value cutoff of 1e-5 was used for local alignment in all the three tools. Further, since the standard classification scheme of doubletrouble was the closest to the classification scheme of DupyliCate, it was adopted for the doubletrouble run with *A. thaliana* Col-0 as the target. Transposed duplicates (TDs) are not classified in the standard scheme, hence, the doubletrouble run involved the use of only the target species *A. thaliana* Col-0. For DupGen_finder, *A. thaliana* Nd-1 was the outgroup and *A. thaliana* Col-0 was the target, as DupGen_finder does not offer classification schemes without TD classification, necessitating the use of an outgroup species. BLAST was used as the aligner for all three tools with a resource allocation of 15 threads during the benchmarking runs to avoid differences due to different sensitivity parameters in DIAMOND.

DupyliCate mainly assigns duplicates to tandems, proximals, and dispersed groups. Therefore, all the other duplicate group classifications apart from tandem and proximal duplicates in DupGen_finder and doubletrouble were processed with custom Python scripts to obtain a single miscellaneous group (‘Misc.’ in Table. 1) that is comparable to the dispersed duplicates group in DupyliCate. The benchmarking was therefore done mainly with respect to the number of genes classified as tandem, proximal, and dispersed duplicates by the three tools. Also, the runtime of the three tools was compared. Five replicate runs were performed on the *A. thaliana* dataset using each of the three tools to estimate the runtime variability given the use of the same computational resources. Since local alignment is a step included within a run of DupyliCate, the local alignment times were also included in the runtime for the other two tools to standardize the comparison. The additional codes, and commands used for running doubletrouble, running DupGen_finder, and performing the benchmarking analysis can be found in the GitHub repository of DupyliCate (https://github.com/ShakNat/DupyliCate).

### Runtime analyses

The runtime of DupyliCate depends on the selected mode and parameter settings. The following approach was adopted to offer a comprehensive runtime analysis when using the different modes and parameter settings. The runtime for each run of the tool using *A. thaliana* datasets was recorded for the following run variations: 1) overlap and strict mode; 2) phylogeny and non-phylogeny-based ortholog detection in the presence of a reference; 3) expression analysis of all identified gene duplicates; 4) expression analysis of specified gene duplicates (XPB tandem duplication^19^, BLA^61^, and RALF tandem duplicates array^17^ reported in *A. thaliana* Col-0 were chosen for demonstrating the specific expression analysis feature) 5) Ka/Ks analysis of all identified gene duplicates; and 6) run with different aligners - BLAST/ DIAMOND/ MMseqs2. Next, to analyze the tool runtime increase with an increasing number of input datasets, a comprehensive runtime analysis was conducted with the Brassicales plants used for the FLS case study. The different datasets were randomly grouped into groups of sizes 2, 4, 6, 8, 10, and 12, respectively. Runtimes were recorded using each group as an input for DupyliCate. To account for the variation in the file sizes of the datasets in each group, there were five random sets of the same group length generated, and the runtime was recorded for every such replicate run.

Finally, to get an idea of the performance of DupyliCate in large-scale studies, the runtime was also analyzed for the tool run on the 153 plant datasets used for the MYB case study that was done in the overlap mode using DIAMOND as the aligner, *A. thaliana* as the reference, and integrated phylogeny for ortholog detection.

### Parameter tuning analysis

DupyliCate offers a number of qualitative and quantitative parameter flags that can be adjusted and tuned according to the specific needs of the user. Among these, most of the qualitative parameters take in the input file paths, and intermediate dependency tool paths and do not contribute to variability in the final results. Hence, twelve parameters that could significantly impact the results in a concrete quantitative manner were selected for tuning trials that were performed with the *A. thaliana* Col-0 dataset as the sample and *A. thaliana* Nd-1 dataset as the reference. One factor at a time (OFAT) approach was employed for these trials wherein only one parameter was changed at a time, and the other parameters remained constantly at default. For almost all the parameters, tuning was performed with three values - the default value, a value 25% lower than the default and a value 25% higher than the default. In case of parameters that accepted only integer values, the low and high values were rounded off to the next nearest integer. For the --score parameter alone, a slightly different approach was adopted. The default for this parameter flag is ‘auto’ which activates the BUSCO-based threshold method, and calculates a species-specific normalized bit score threshold for singleton-duplicate segregation. The other option for this --score flag is manual where the user needs to enter specific values. Since the auto mode does not depend on user input at all and is entirely species-dependent, the self-similarity threshold is switched off or is automatically set to 0 (--self_simcut is 0). But in the manual mode the --score flag must be provided by the user and the default value of 50% for the --self_simcut parameter flag is used. But this --self_simcut value can also be adjusted by the user in the manual mode. Thus the --self_simcut flag jointly influences the singleton-duplicates segregation along with the --score flag. Therefore the user needs to know the contributions of self-normalized bit score cutoff and self-similarity cutoff individually and taken together to select suitable values when setting them manually. Hence, a detailed combinatorial analysis was performed with these two parameters, where one flag was set to 0 and the other was set to the low, default, and high values, to demonstrate their individual effects. Another set of analyses that combined the low, default and high values from both these parameters were also performed to show their combined effects.

Further, after the tuning trials, a set of BUSCO-based threshold vs. manually set threshold runs were performed using DupyliCate on the plant species *Digitalis purpurea*^62^, *Victoria cruziana*^63^, and *Dioscorea dumetorum*^64^ along with *Arabidopsis thaliana* Col-0. The manual threshold runs were performed with --score of 0.5 (0.5 was chosen since it is a median-normalized bit score cutoff that is bounded by 0 and 1) and --self_simcut of 50% (default value of the --self_simcut flag). The BUSCO vs manual runs were performed to give a comparative analysis of the results between the two approaches that can be used for adjusting the parameters of a DupyliCate run. The above mentioned plant species were specifically chosen for this analysis in order to show the applicability of the method and the reproducibility of results over a wide variety of plant species.

## Data Availability

DupyliCate, the helper scripts, and test data sets can be accessed at: https://github.com/ShakNat/DupyliCate. The repository is also archived in Zenodo and can be accessed using https://doi.org/10.5281/zenodo.19202192. Underlying data sets are publicly available at the NCBI, Phytozome, and other databases as specified in Supplementary Table S1. Details of the supplementary data can be found in Supplementary Information.

The datasets used in the study can also be accessed here:

https://doi.org/10.1186/s12864-023-09823-2, https://doi.org/10.1038/s41467-025-61686-1, https://doi.org/10.1038/srep17394, https://phytozome.jgi.doe.gov/info/Mtilingiivar_LVR_v1_1, https://doi.org/10.1111/tpj.14993, https://doi.org/10.1186/s13059-020-1952-4, http://phytozome-next.jgi.doe.gov/info/AtequilanavarWeber';sBlueHAP1_v3_1, https://doi.org/10.1038/ng.3435, https://doi.org/10.1038/nature06148, https://doi.org/10.1093/g3journal/jkae262, https://doi.org/10.1111/tpj.16454, https://doi.org/10.1038/nbt.2196, https://doi.org/10.1186/gb-2012-13-5-r39, https://doi.org/10.1093/plcell/koab077, http://phytozome.jgi.doe.gov/info/KlaxifloraFTBG2000359A_v3_1, https://doi.org/10.1186/s12915-017-0412-4, https://doi.org/10.1038/nbt.2906, http://phytozome-next.jgi.doe.gov/info/Hquercifolia_v1_1, https://doi.org/10.1038/ncomms4311, https://doi.org/10.1038/s41467-017-01491-7, https://doi.org/10.1093/plcell/koac347, https://doi.org/10.1038/s41467-021-24328-w, https://phytozome.jgi.doe.gov/info/Mnasutusvar_SF_v2_1, https://doi.org/10.1126/science.1203810, https://doi.org/10.1101/gr.276358.121, https://doi.org/10.1038/s41467-019-14197-9, https://phytozome-next.jgi.doe.gov/info/PnigraxmaximowicziiNM6_v1_1, https://doi.org/10.1038/nature22971, https://doi.org/10.3389/fpls.2013.00046, https://doi.org/10.1038/s41559-017-0119, https://doi.org/10.1093/g3journal/jkad209, https://doi.org/10.1101/2021.07.23.453237, https://doi.org/10.1038/s41586-019-1852-5, https://doi.org/10.1093/g3journal/jkae021, https://doi.org/10.1073/pnas.1703088114, https://doi.org/10.12688/f1000research.38156.1), https://doi.org/10.1038/s42003-021-02009-0, https://doi.org/10.1002/tpg2.20319, https://doi.org/10.1073/pnas.1619928114, https://phytozome.jgi.doe.gov/info/EanacuaAZ03848HAP1_v1_1, https://doi.org/10.1126/science.1188800, https://doi.org/10.1101/2024.06.15.599162, https://doi.org/10.1038/s41467-017-01064-8, https://doi.org/10.1038/s41588-019-0356-4, https://doi.org/10.1073/pnas.1708621114, https://doi.org/10.1093/database/baz110, http://phytozome.jgi.doe.gov/, http://phytozome.jgi.doe.gov/info/CmollissimaMahoganyHAP1_v1_1, https://phytozome-next.jgi.doe.gov/info/Ufusca_v1_1, https://doi.org/10.3389/fpls.2020.00496, https://doi.org/10.1111/tpj.14500, https://doi.org/10.1038/ncomms4930, https://doi.org/10.1038/nbt.1674, https://doi.org/10.1038/s41467-020-18923-6, http://phytozome.jgi.doe.gov/info/Ccanadensis_V3_1, https://phytozome-next.jgi.doe.gov/info/Pvirgatumvar_AP13HAP1_v6_1, https://doi.org/10.1101/2024.02.14.580303, https://phytozome.jgi.doe.gov/info/HleucocephalaHAP1_v2_1, https://phytozome.jgi.doe.gov/info/Pcoccineus_v1_1, https://doi.org/10.1038/nature06856, https://doi.org/10.1073/pnas.0611046104, https://doi.org/10.1186/1471-2164-15-312, http://phytozome.jgi.doe.gov, https://doi.org/10.1038/nature13308, https://doi.org/10.1038/s41467-020-17302-5, https://www.ncbi.nlm.nih.gov/datasets/genome/GCA_044591495.1/, https://doi.org/10.1038/nature11798, https://doi.org/10.1038/s41587-020-0681-2, https://vandepoelelab.be/plaza/instances/dicots_05/organism/view/chi, https://doi.org/10.1111/j.1365-313x.2012.05093.x, https://doi.org/10.1038/s41438-021-00481-7, https://phytozome.jgi.doe.gov/info/YfilamentosaC3alt_v2_1, https://doi.org/10.1126/science.1251788, https://doi.org/10.1186/s12915-020-00795-3, https://doi.org/10.1093/gigascience/giaa100, https://doi.org/10.7554/elife.36426, https://doi.org/10.1111/pbi.14199, https://doi.org/10.1038/s41477-018-0337-0, https://doi.org/10.1126/sciadv.abh2488, https://doi.org/10.1038/s41588-020-0614-5, https://doi.org/10.1038/s41467-021-20921-1, https://doi.org/10.1093/nar/gkl976, https://doi.org/10.1126/science.1128691, https://doi.org/10.1371/journal.pone.0216233, https://doi.org/10.1016/j.algal.2020.101990, https://doi.org/10.1038/s41438-019-0142-6, https://phytozome.jgi.doe.gov/info/Brapassp_chinensisvar_communisPCGlu_v2_1, https://doi.org/10.1038/nbt.2491, http://bmap.jgi.doe.gov/, https://doi.org/10.1038/s41477-022-01226-7, https://doi.org/10.1038/ng.3886, https://doi.org/10.1038/s41467-018-07669-x, https://doi.org/10.1186/1471-2164-14-498, https://doi.org/10.1073/pnas.1109451108, https://phytozome-next.jgi.doe.gov/info/Mesculenta_v8_1, https://doi.org/10.1534/genetics.105.048900, https://doi.org/10.1038/s41467-023-38915-6, https://doi.org/10.1038/s41467-021-22858-x, https://phytozome-next.jgi.doe.gov/info/AgerardiHAP2_v1_1, https://doi.org/10.1038/s41586-022-04808-9, https://doi.org/10.1016/j.cell.2017.09.030, http://phytozome-next.jgi.doe.gov/info/Claxum_v1_1, https://doi.org/10.1093/hr/uhad061, https://doi.org/10.1038/nature22380, https://doi.org/10.1038/ncomms14953, https://doi.org/10.1002/tpg2.20101, https://doi.org/10.1038/s41586-023-05791-5, https://doi.org/10.3390/genes11050483, https://doi.org/10.1016/j.xgen.2023.100467, https://doi.org/10.1038/ng.2586, https://phytozome.jgi.doe.gov/info/CsativaAcsn_226_v1_1, https://doi.org/10.1093/g3journal/jkae208, https://phytozome.jgi.doe.gov/info/Uamericanavar_NA87034HAP1_v1_1, https://doi.org/10.1038/nature21370, https://doi.org/10.1038/ng.807, http://phytozome-next.jgi.doe.gov/, https://phytozome.jgi.doe.gov/info/Bjunceassp_integrifoliaMIZ_19_v1_1, https://www.ncbi.nlm.nih.gov/datasets/genome/GCA_037355405.1/, https://doi.org/10.1038/ng.889, https://doi.org/10.1093/dnares/dsac033, https://doi.org/10.1038/ng.2669, https://doi.org/10.1111/tpj.13415, http://phytozome.jgi.doe.gov/info/BhybridumBhyb26_v2_1, https://doi.org/10.1038/nature11241, https://doi.org/10.1038/s41477-023-01526-6, https://phytozome.jgi.doe.gov/info/NdensiflorusRogue1HAP1_v1_1, https://doi.org/10.1007/s00122-019-03311-6, https://doi.org/10.1038/nature15714, https://doi.org/10.3390/genes11030274, https://doi.org/10.1038/s41467-022-35507-8, https://doi.org/10.1093/plphys/kiac116, https://doi.org/10.1038/s41438-021-00641-9

## AI Statement

AI tools (ChatGPT and Claude) were used for optimizing, restructuring, and debugging the DupyliCate script. Claude was also used to check the language and grammar of the manuscript text.

## Supporting information

Supplementary Discussion

Supplementary Results 1

Supplementary Results 2

## Acknowledgements

The authors thank the Plant Biotechnology and Bioinformatics group (University of Bonn) for valuable inputs and inspiring discussions. SN was supported by the doctoral scholarship of the German Academic Exchange Service (DAAD). This work was supported by the de.NBI Cloud within the German Network for Bioinformatics Infrastructure (de.NBI) and ELIXIR-DE (Forschungszentrum Jülich and W-de.NBI-001, W-de.NBI-004, W-de.NBI-008, W-de.NBI-010, W-de.NBI-013, W-de.NBI-014, W-de.NBI-016, W-de.NBI-022). Open access funding was enabled by Project DEAL.

## Author information

### Contributions

S.N. and B.P. designed the study. S.N. wrote the software and conducted the bioinformatic analyses. S.N. and B.P. interpreted the results and wrote the manuscript. All authors read the final version of the manuscript and agreed to its submission.

## Ethics declarations

### Ethics approval and consent to participate

Not applicable.

### Consent for publication

Not applicable.

