## Supplementary Discussion for "DupyliCate: mining, classifying, and characterizing gene duplications"

### Parameter tuning

DupyliCate offers a number of tunable parameters and run modes for the user to adapt it to the specific use case at hand. Of the different parameters available, some are more crucial for the analysis results than others. Among these, the parameters can be divided as being auto-selectable by the tool itself and user tunable. Currently, the auto select option is available for the step involving duplicate or paralog identification using a BUSCO-derived self-normalized bit score, detailed previously. But the paralog detection can also be made completely user controlled by choosing to manually adjust the parameters of self-similarity cutoff and self-normalized bit score. It is generally recommended to utilize the auto detection of optimal settings to arrive at species-specific thresholds.

For the duplicates' classification, genes, exactly adjacent to each other are considered tandem duplicates. A proximity parameter with a default value of 10 is used to classify non-adjacent genes that have 10 or less than 10 intervening genes in between as proximal duplicates while the rest are classified as dispersed duplicates. This value can be adjusted depending on the level of proximity required for specific analyses.

Moving on to ortholog detection in the presence of a reference, both the forward alignment similarity score and forward alignment normalized bit score (bit score of a gene's forward hit relative to the bit score against itself) are user tunable. For cases where the evolutionary distance between the reference and samples are from low to moderate, the set similarity default of 40%, and normalized bit score default of 0.3 are recommended. When the distance becomes large, or when the analysis involves a mix of samples, some of which are close and some of which are farther away from the reference, it is recommended to lower the similarity and normalized bit score cutoffs accordingly. The ortholog detection is a comprehensive step involving confidence score assignments to predict the reliability of the identified orthologs. Low confidence ortholog assignments have an additional output, where other possible orthologs for the particular gene are listed along with the gene chosen as the ortholog for the particular sample gene. By default, the top 3 forward alignment hits are returned as the potential orthologs. When a phylogeny-based ortholog detection is conducted, iterative phylogenetic tree building is done with the top n forward local alignment hits in case the top hit is not distinct enough from the rest of the hits. The last of the top n hits is used as the root point for the tree, as described in Supplementary_Figure_3. The default value of n is 10 and it is recommended to keep this setting.. This is because an increase in this number would lead to increased phylogenetic noise, and a decrease would distort the phylogenetic signal. In case this parameter needs to be tweaked, it is recommended to increase the value rather than decrease it, because of the last top hits' influence on the tree root point. But it is important to keep in mind that this will also increase the computational time of the analysis due to the increased number of sequences for each tree construction step. Further, the parameters, flank, side, and synteny score play a role in influencing the synteny results reported between the identified orthologs. Flank takes the genes upstream and downstream of the target gene, and decides the synteny window size for the analysis. An increase in the flank parameter value, increases the synteny window size, and consequently leads to a longer runtime. Side is a parameter that defines the number of gene pairs that should show up as each other's best hits in the forward alignment on the upstream and downstream ends of a target gene. This parameter influences the stringency of synteny. This means, higher the value of the side parameter, more stringent will be the criteria to call a particular gene as being syntenic to its corresponding ortholog, as more number of gene pairs need to show up as best hits in the upstream and downstream regions. Lastly, synteny score is a ratio that combines the synteny window size and side values to act as a cutoff that decides whether the particular ortholog pair is syntenic or non- syntenic and shows a stringency level similar to the side parameter.

The default e-value cutoff for both forward and self alignments is 1e-5 that should work for most use cases. All the critical parameters have been looked at here. More details on other available options and flags can be found in the DupyliCate's GitHub repository.
