## Supplementary Figure 1 for "DupyliCate: mining, classifying, and characterizing gene duplications"

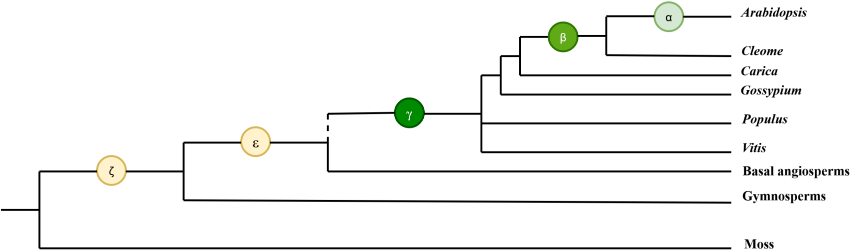


Phylogenetic tree depicting the major polyploidization events (γ,β,α) in the evolutionary history leading to *Arabidopsis* along with the ancient polyploid events ζ and ε reported in the evolutionary trajectory leading to seed plants and angiosperms, respectively. The dotted line before the γ is placed to show the other polyploidization events (τ, σ, ρ) not included in the representation [27, 29].
