## Supplementary Figure 2 for "DupyliCate: mining, classifying, and characterizing gene duplications"

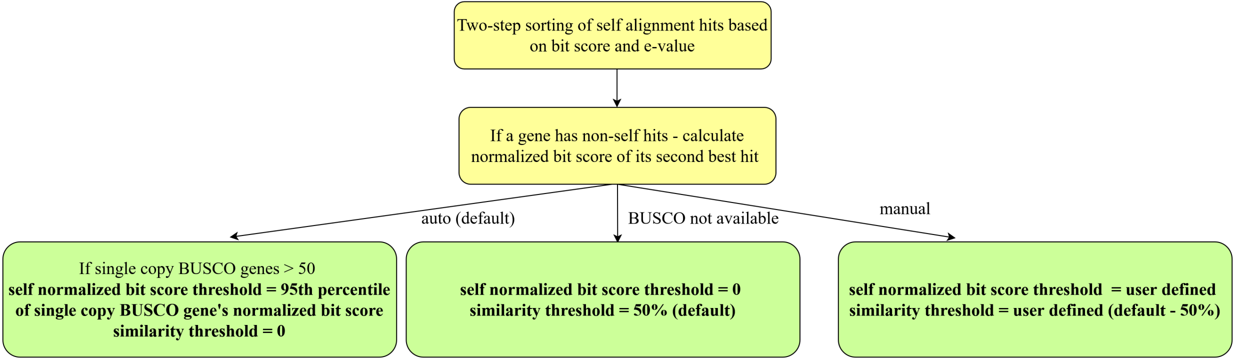


Steps involved in singleton-duplicate genes segregation based on different thresholding approaches
