## Supplementary Figure 3 for "DupyliCate: mining, classifying, and characterizing gene duplications"

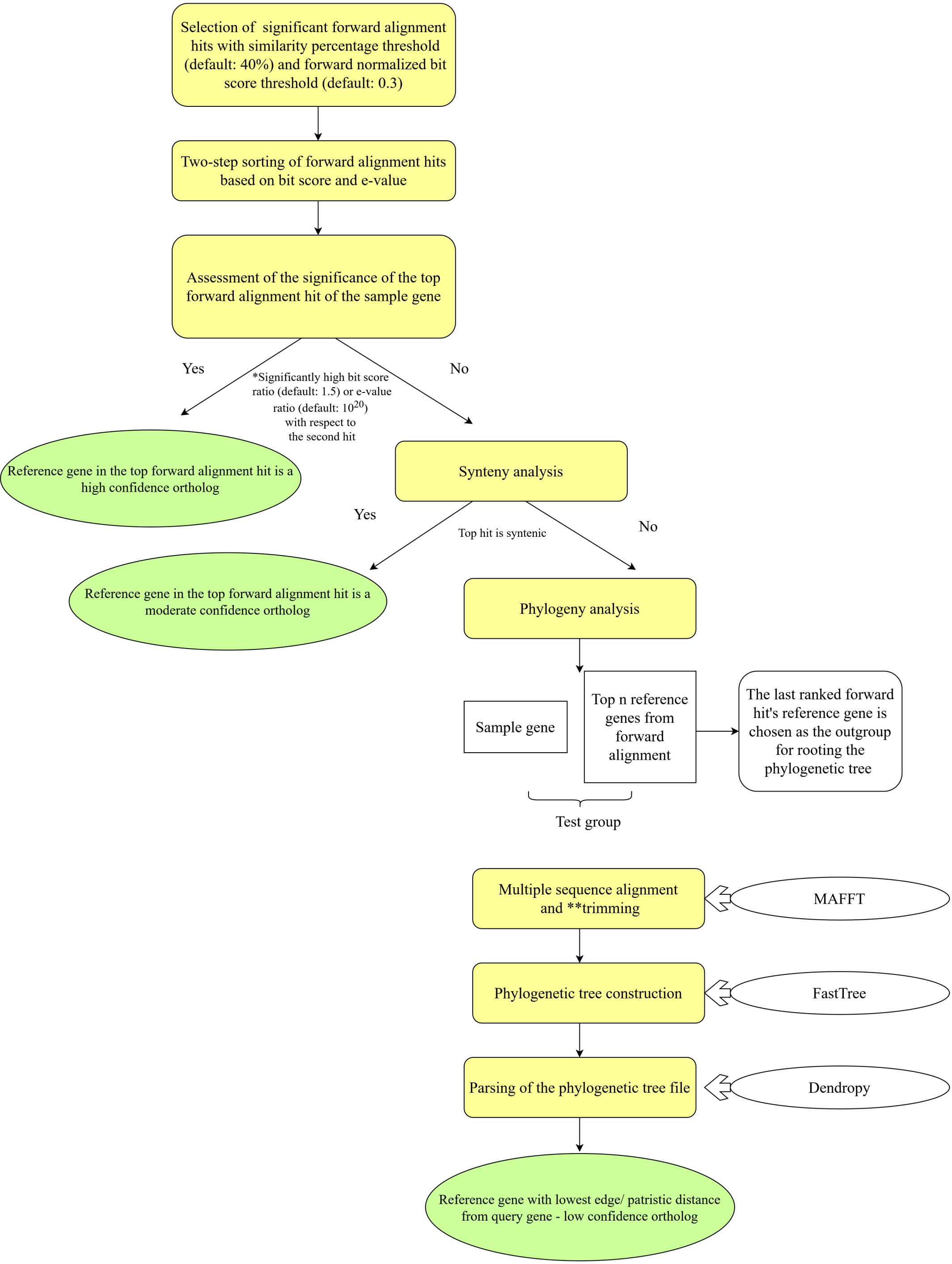


Ortholog assignment steps in the presence of a reference organism. *The bit score and e-value ratios used for checking if the top hit is significantly different from the second hit, were empirically determined. **Trimming of multiple sequence alignment files is performed by code in-built within DupyliCate and does not rely on any external tool for this step.
