## Supplementary Figure 4 for "DupyliCate: mining, classifying, and characterizing gene duplications"

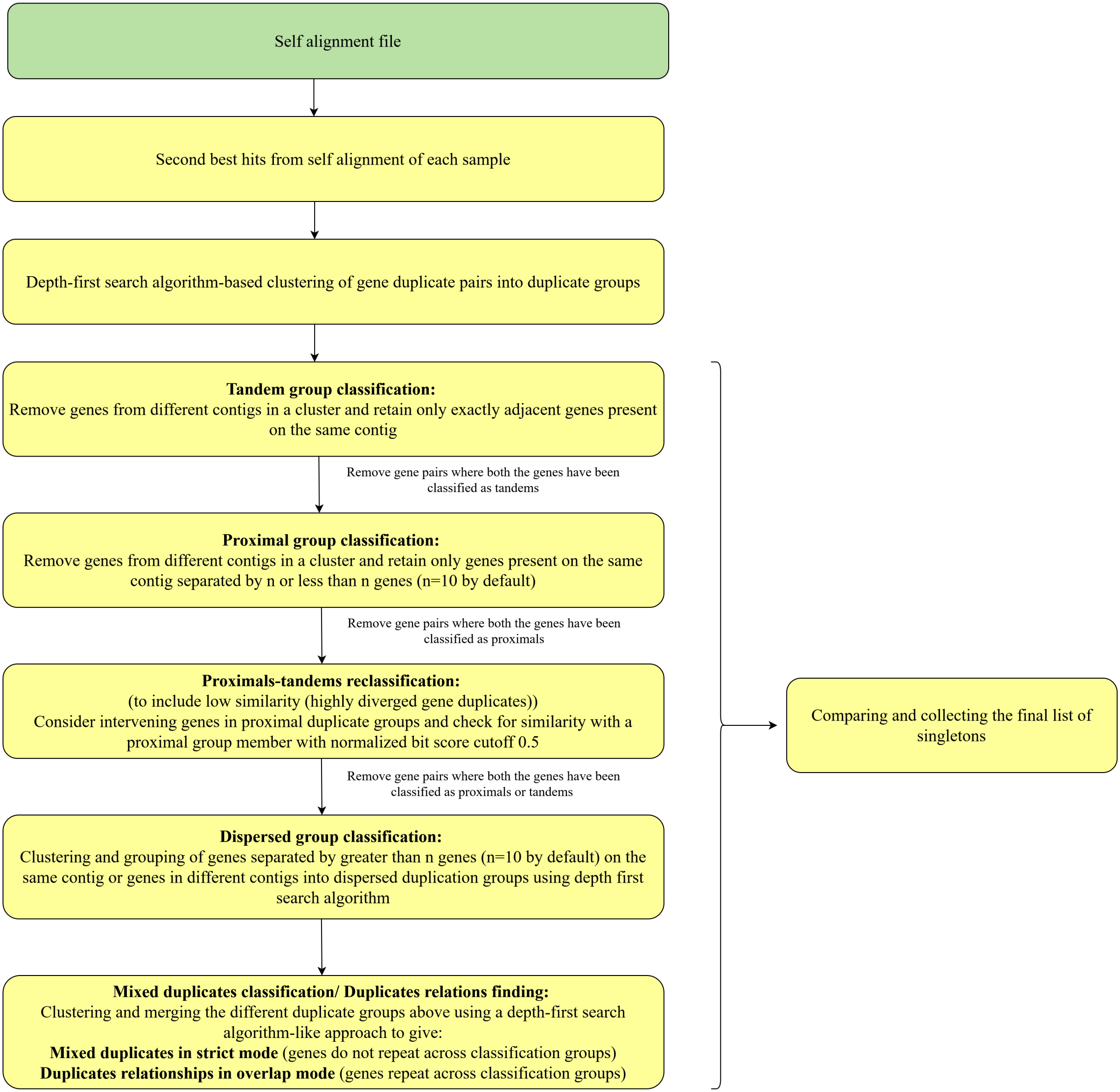


Flow chart of the algorithms and steps involved in gene duplicates array clustering, and classification
