## Supplementary figures and images for "DupyliCate: mining, classifying, and characterizing gene duplications"

### Supplementary Figure 5

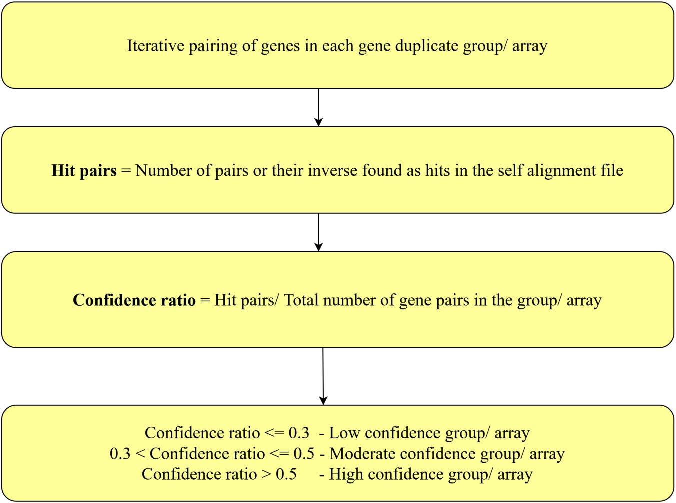


Confidence scoring scheme for gene duplicate groups output by DupyliCate

### Supplementary Figure 9

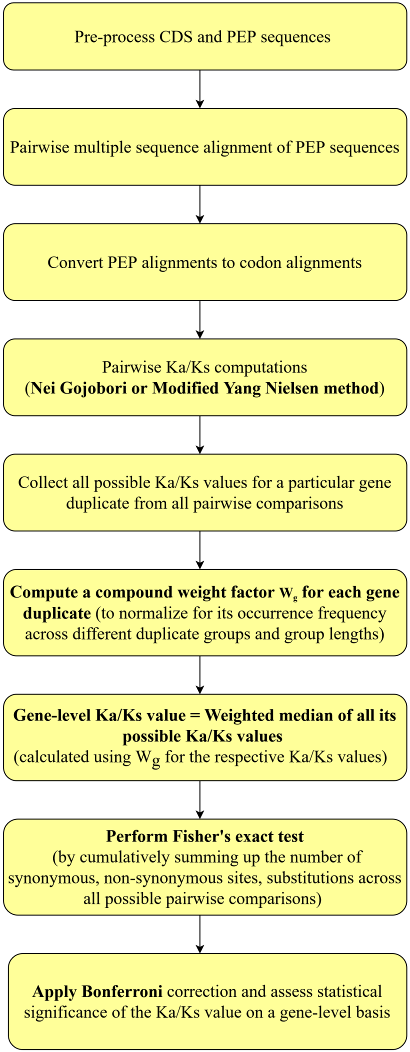


Steps involved in gene-level Ka/Ks computation in DupyliCate

### Supplementary Figure 10

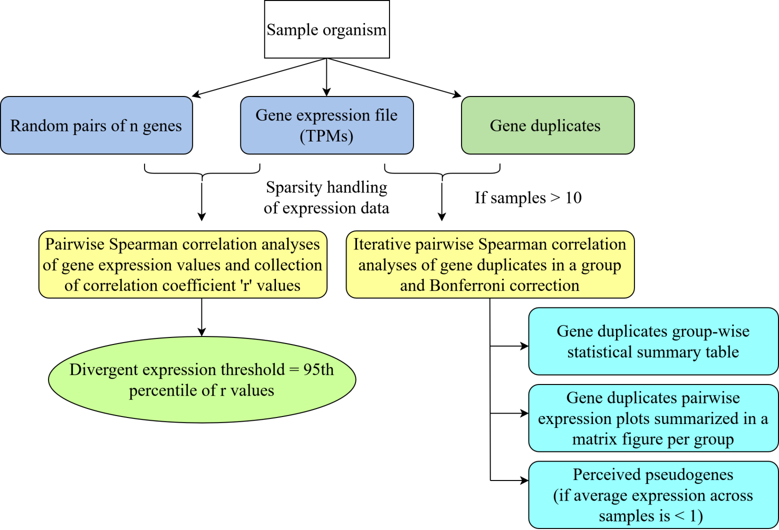


Methodology of statistical analysis and gene expression analysis of gene duplicates

### Supplementary Figure 11

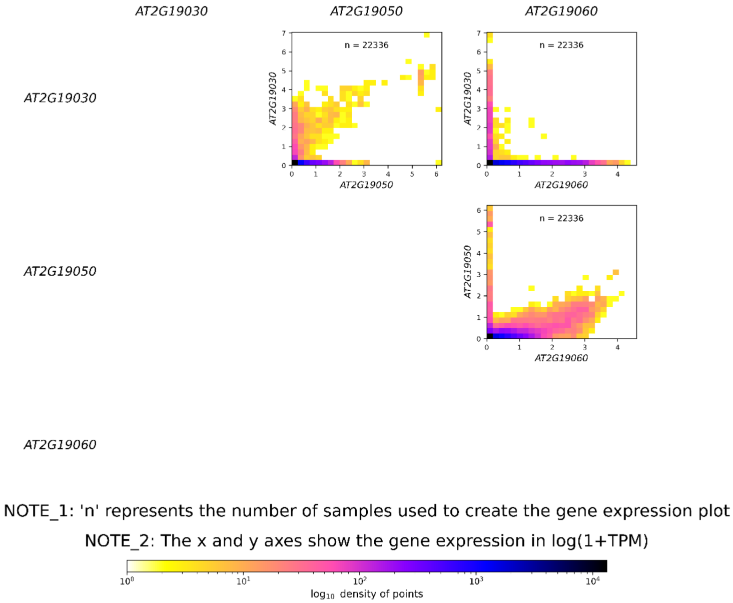


Matrix type gene expression plot
