## Supplementary Figure 6 for "DupyliCate: mining, classifying, and characterizing gene duplications"

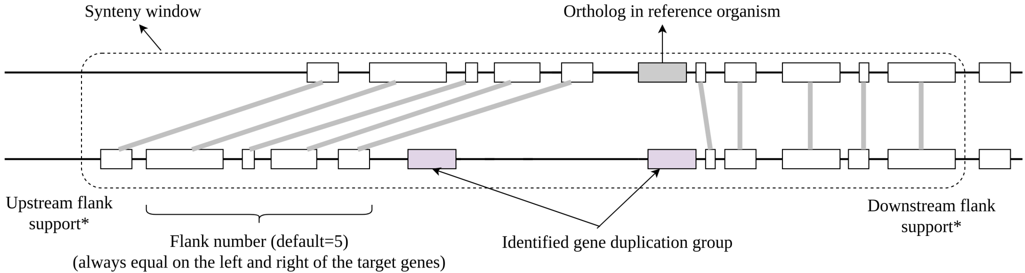


* - sufficient flanking genes must be best hits in forward local alignment
(default flank support is 1 each in the upstream and downstream flank regions)

Schematic representation of synteny analysis performed for small scale gene duplicates as well as ortholog assignment
