## Supplementary Figure 7 for "DupyliCate: mining, classifying, and characterizing gene duplications"

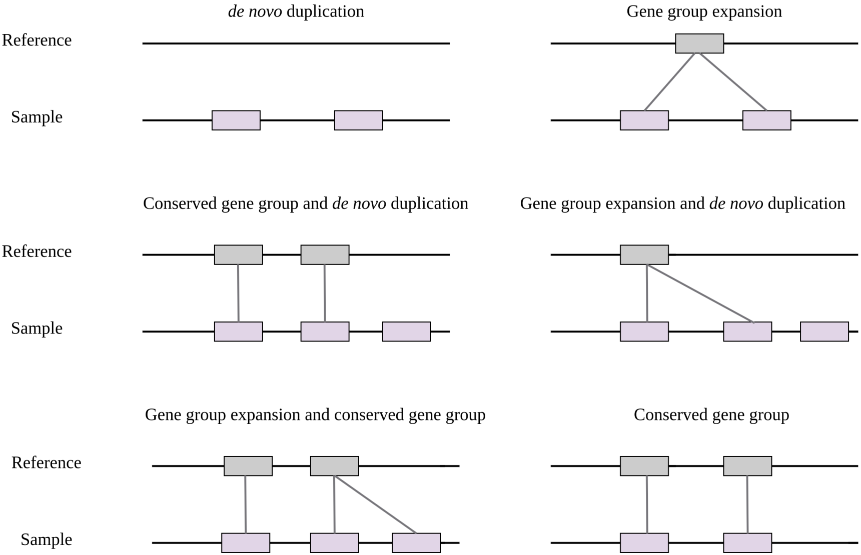


Schematic representation of the different possible types of the identified small scale gene duplicate groups with respect to orthologous genes in the reference organism
