## Supplementary Figure 8 for "DupyliCate: mining, classifying, and characterizing gene duplications"

**(a)**

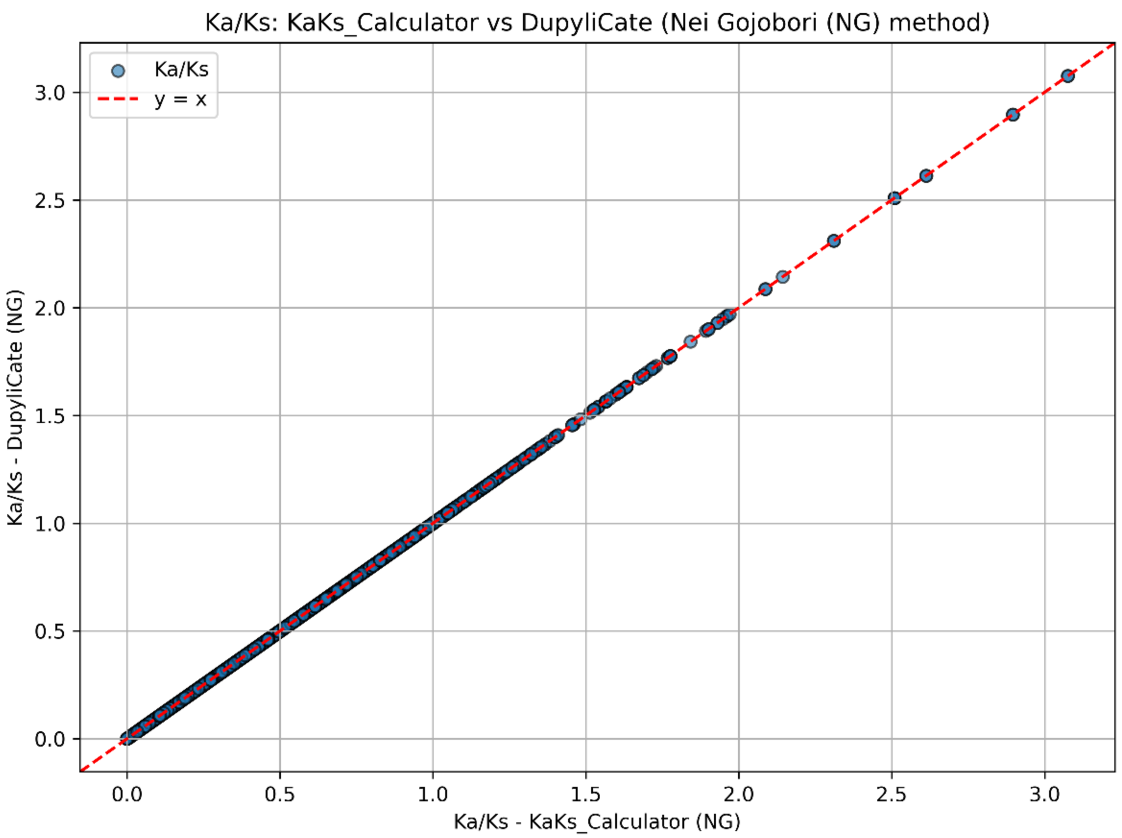


**(b)**

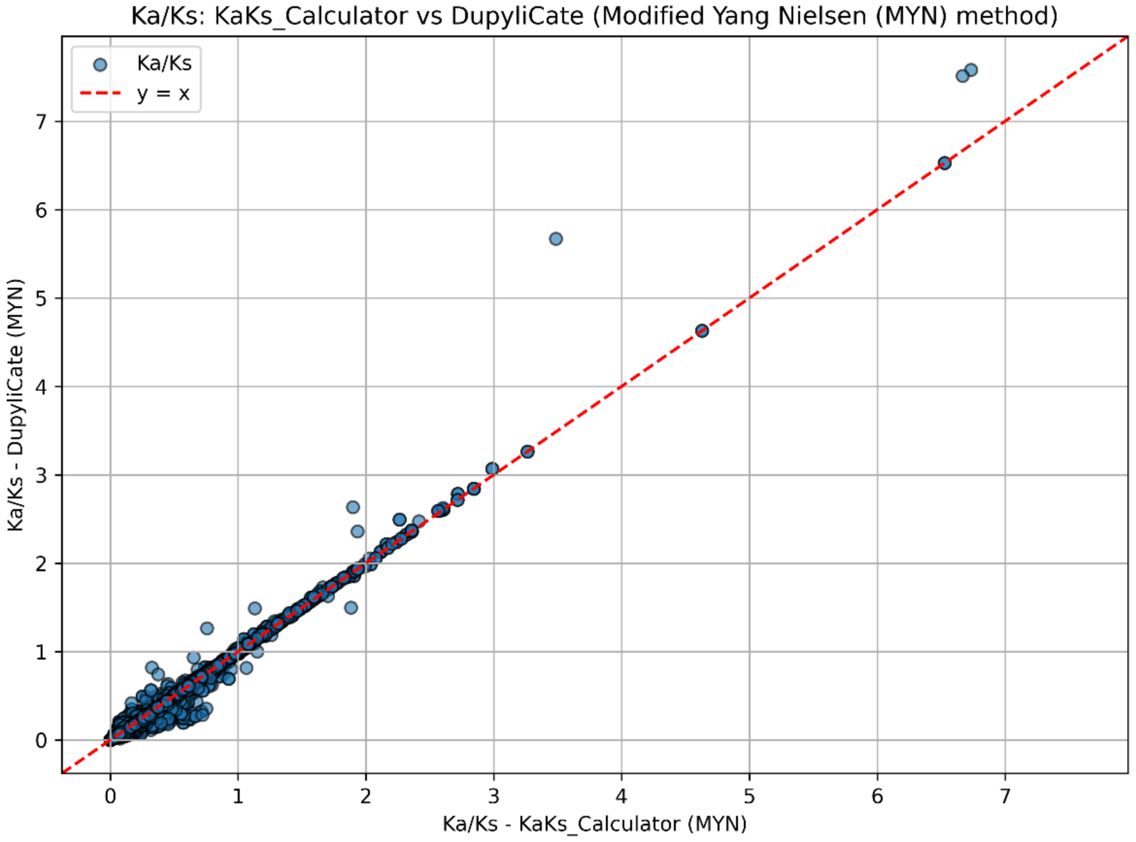


Correlation analysis of KaKs results using python implementations of the (a) Nei Gojobori (NG) method with the results of NG method in KaKs_Calculator2.0 and the (b) Modified Yang Nielsen (MYN) method with the results of MYN method in KaKs_Calculator2.0
