## Supplementary Information for "DupyliCate: mining, classifying, and characterizing gene duplications"

Supplementary_Table_S1: List of all datasets used in the study along with their respective references

Supplementary_Table_S2: Consolidated results of BUSCO analysis

Supplementary_Table_S3: Consolidated gene duplicates results summary

Supplementary_Table_S4: Statistical analysis results of specific tandem gene duplicates in *A. thaliana* Col-0

Supplementary_Table_S5: Orthologs and copy number variations of orthologs of stress genes in the genomes of *O. sativa* indica and related species (XLSX file)

Supplementary_Table_S6: Orthologs and copy number variations of orthologs of *FLS* genes in *A. thaliana* across other Brassicales plant species (XLSX file)

Supplementary_Table_S7: Orthologs and copy number variations of orthologs of *AtMYB12,* and *AtMYB111* genes in *A. thaliana* across a wide phylogenetic range of plant species (XLSX file)

Supplementary_Table_S8: Orthologs confidence scoring scheme

Supplementary_Table_S9: Orthologous groups confidence scoring scheme

Supplementary_Table_S10: Consolidated results of the parameter tuning trials of DupyliCate

Supplementary_Table_S11: DupyliCate parameter sensitivity ranking table

Supplementary_Table_S12: Comparative analyses results for BUSCO-based threshold and manual threshold DupyliCate runs

Supplementary_Figure_1: Representative phylogenetic tree depicting whole genome duplication events in *A. thaliana*

Supplementary_Figure_2: Threshold and cutoff determination methods for singleton duplication classification

Supplementary_Figure_3: Workflow of ortholog detection steps in the presence of a reference organism

Supplementary_Figure_4: Workflow of gene duplicates clustering and classification
Supplementary_Figure_5: Confidence scoring scheme of gene duplicate clusters
Supplementary_Figure_6: Synteny analysis concept diagram

Supplementary_Figure_7: Different types of gene duplicate group attributions based on synteny, gene copy number expansion or conservation

Supplementary_Figure_8: Correlation analysis plots of Ka/Ks values obtained using DupyliCate and KaKs_Calculator

Supplementary_Figure_9: Ka/Ks computation steps in DupyliCate

Supplementary_Figure_10: Gene expression and statistical analyses workflow

Supplementary_Figure_11: Matrix type gene expression plot

Supplementary_Results_1: Consolidated PDF file containing all the duplication landscape plots of plants analyzed in this study

Supplementary_Results_2: Detailed results of correlation analysis of Ka/Ks values obtained using DupyliCate and KaKs_Calculator

Supplementary_Methods: Details on methodology adopted for validation of the ortholog assignments in the MYB case study (Fig. 6)

Supplementary_Discussion: Detailed discussion on parameter tuning of DupyliCate
