## Supplementary Methods for "DupyliCate: mining, classifying, and characterizing gene duplications"

### Validation analysis for MYB case study

The following approach was undertaken for validating the orthology assignments in DupyliCate and the clade grouping in the phylogenetic tree (Fig. 6). Each plant species used for the MYB case study was taken and the respective *AtMYB111* and *AtMYB12* ortholog assignments were looked at in the copy number table produced in the analysis (Supplementary_Table_S7). If the genes placed as *MYB111* or *MYB12* orthologs in the table were also placed in the *AtMYB111* or *AtMYB12* clades in the tree, respectively, then it was taken as an exact match. For monocots, basal angiosperms, and few other dicots (that were placed in the sister clade to both *MYB12* and *MYB111*) since *MYB12* and *MYB111* were not found as separate lineages, a different approach was followed. If the ortholog assignments for these lineages in DupyliCate turned out to be *MYB12* or *MYB111*, and the assignments had negligible or no sequences near the outgroup clade, they were taken to exactly match. Species, in which the ortholog assignments had correct placements in the tree but also included false positives (like other closely related MYBs showing up as orthologs for *MYB111* or *MYB12* - the sequences close to the outgroup clade), were classified as showing moderate match. Species where the ortholog assignments had mismatches as in - if DupyliCate gave a gene as an ortholog for *MYB12* but the tree placed it in the *AtMYB111* clade, were designated as low level matches. Finally, species that only had sequences near the outgroup clade were designated as extremely low confidence matches. The species placed in the different levels of confidence were then counted and a percentage of species allocated to each of the different match confidence classes (from the total number of plant species used for the MYB case study) were computed and represented in the validation pie chart (Fig. 7).
