## Supplementary Results 1 for "DupyliCate: mining, classifying, and characterizing gene duplications"

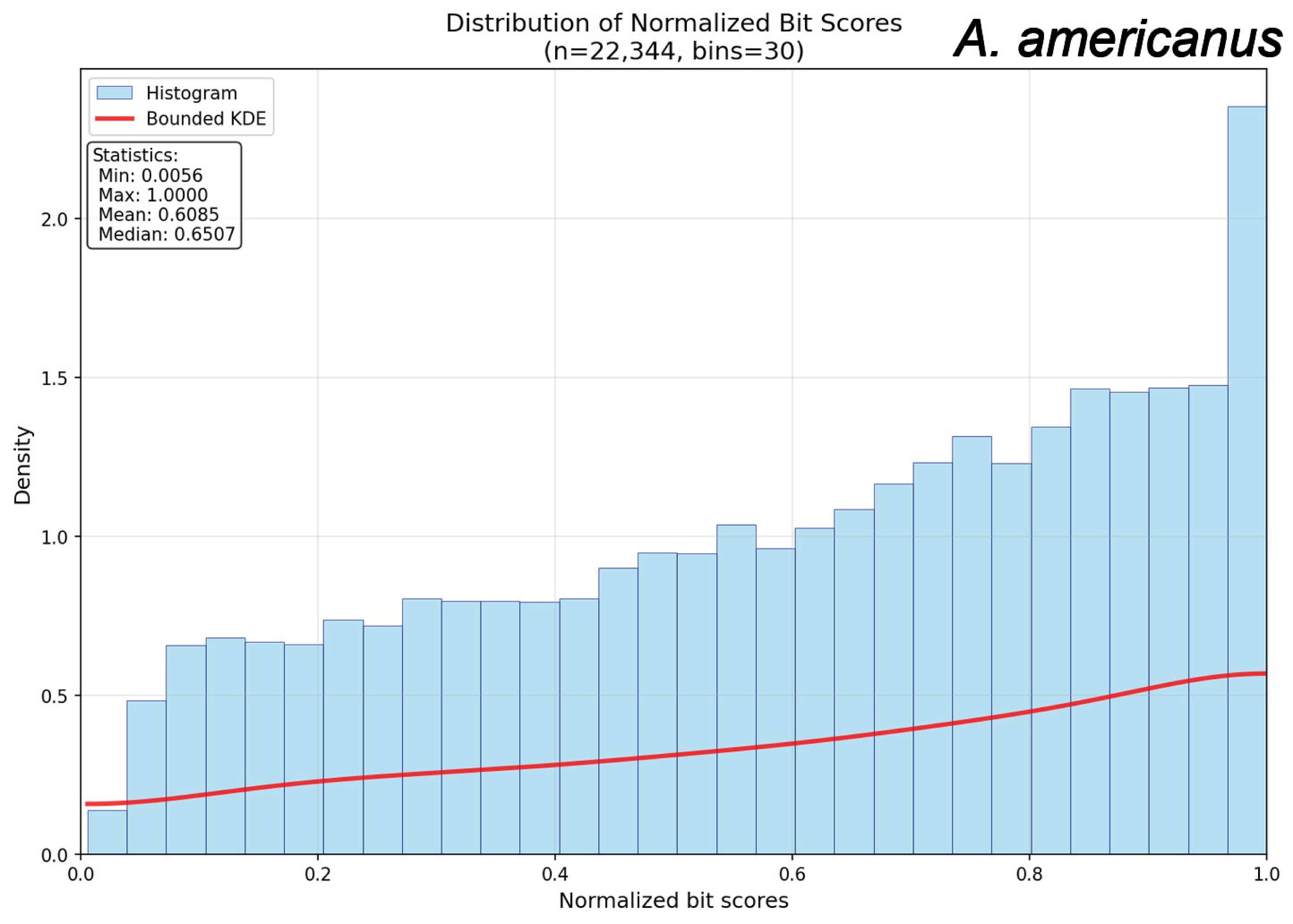
Distribution of Normalized Bit Scores
(n=22,344, bins=30)
A. americanus
Histogram
Bounded KDE
Statistics:
Min: 0.0056
Max: 1.0000
Mean: 0.6085
Median: 0.6507
Density
0.0
0.5
1.0
1.5
2.0
0.0
0.2
0.4
0.6
0.8
1.0
Normalized bit scores

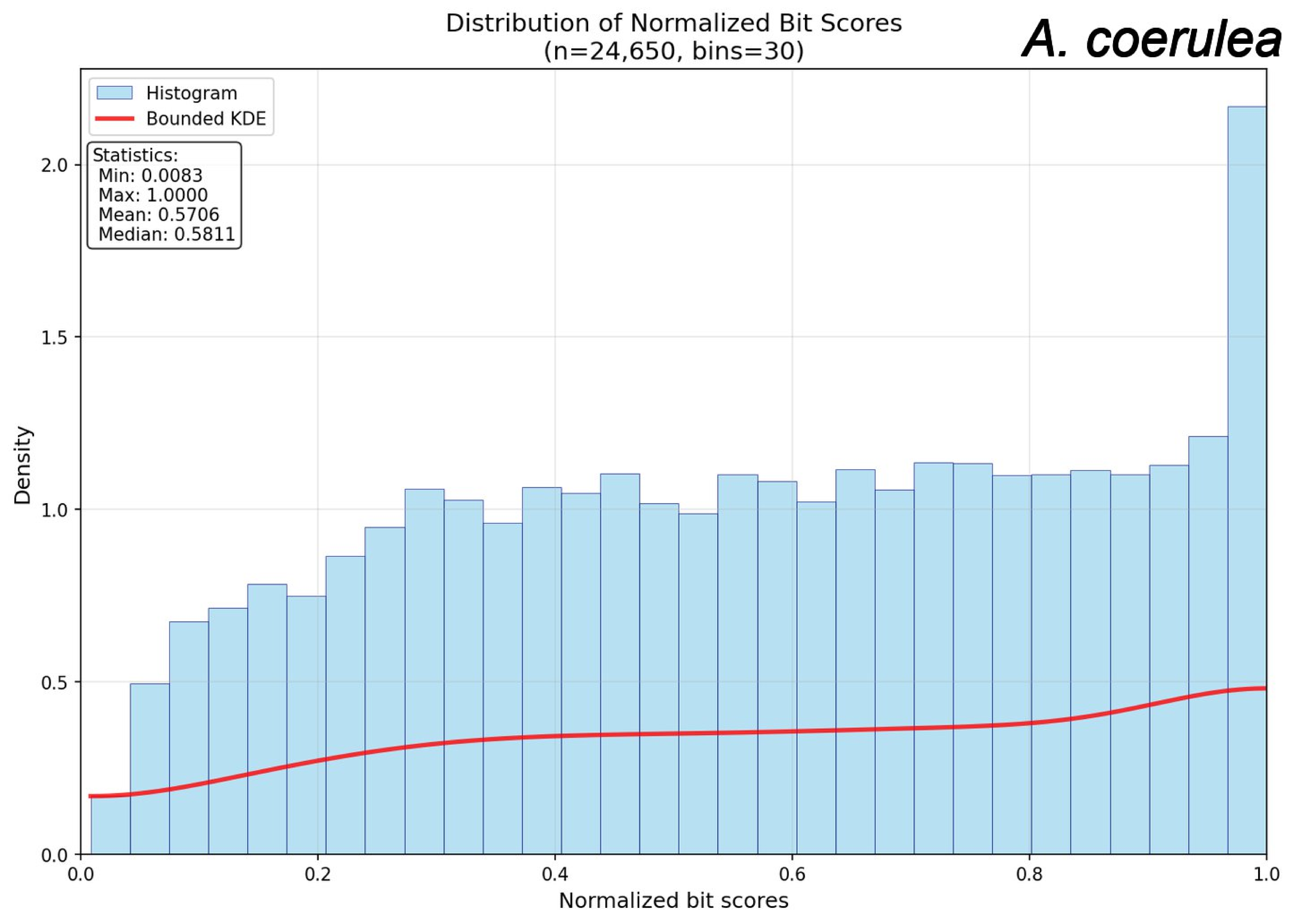
Distribution of Normalized Bit Scores
(n=24,650, bins=30)
A. coerulea
Histogram
Bounded KDE
Statistics:
Min: 0.0083
Max: 1.0000
Mean: 0.5706
Median: 0.5811
Density
0.0
0.5
1.0
1.5
2.0
0.0
0.2
0.4
0.6
0.8
1.0
Normalized bit scores

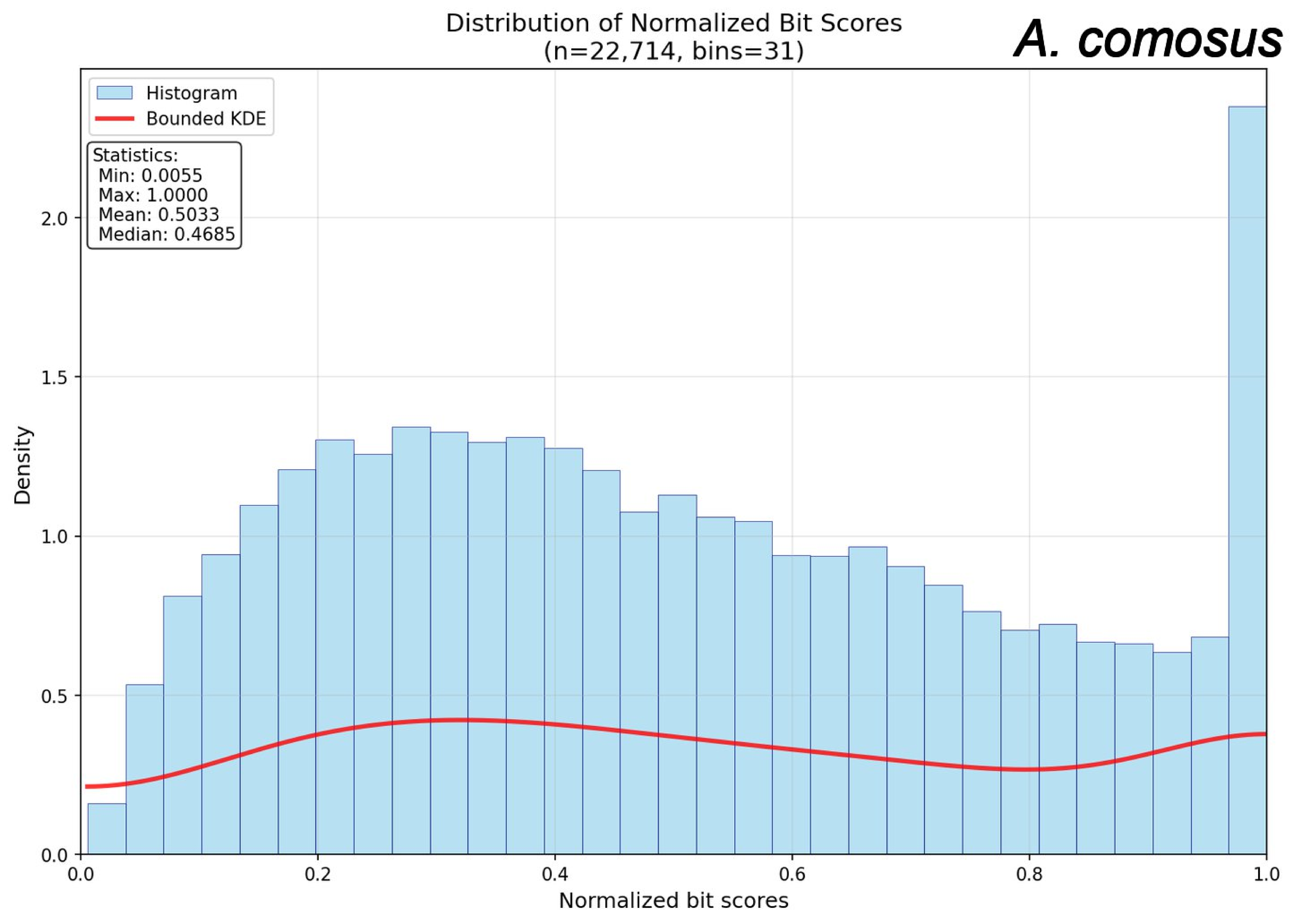
Distribution of Normalized Bit Scores
(n=22,714, bins=31)
A. comosus
Histogram
Bounded KDE
Statistics:
Min: 0.0055
Max: 1.0000
Mean: 0.5033
Median: 0.4685
Density
0.0
0.5
1.0
1.5
2.0
0.0
0.2
0.4
0.6
0.8
1.0
Normalized bit scores

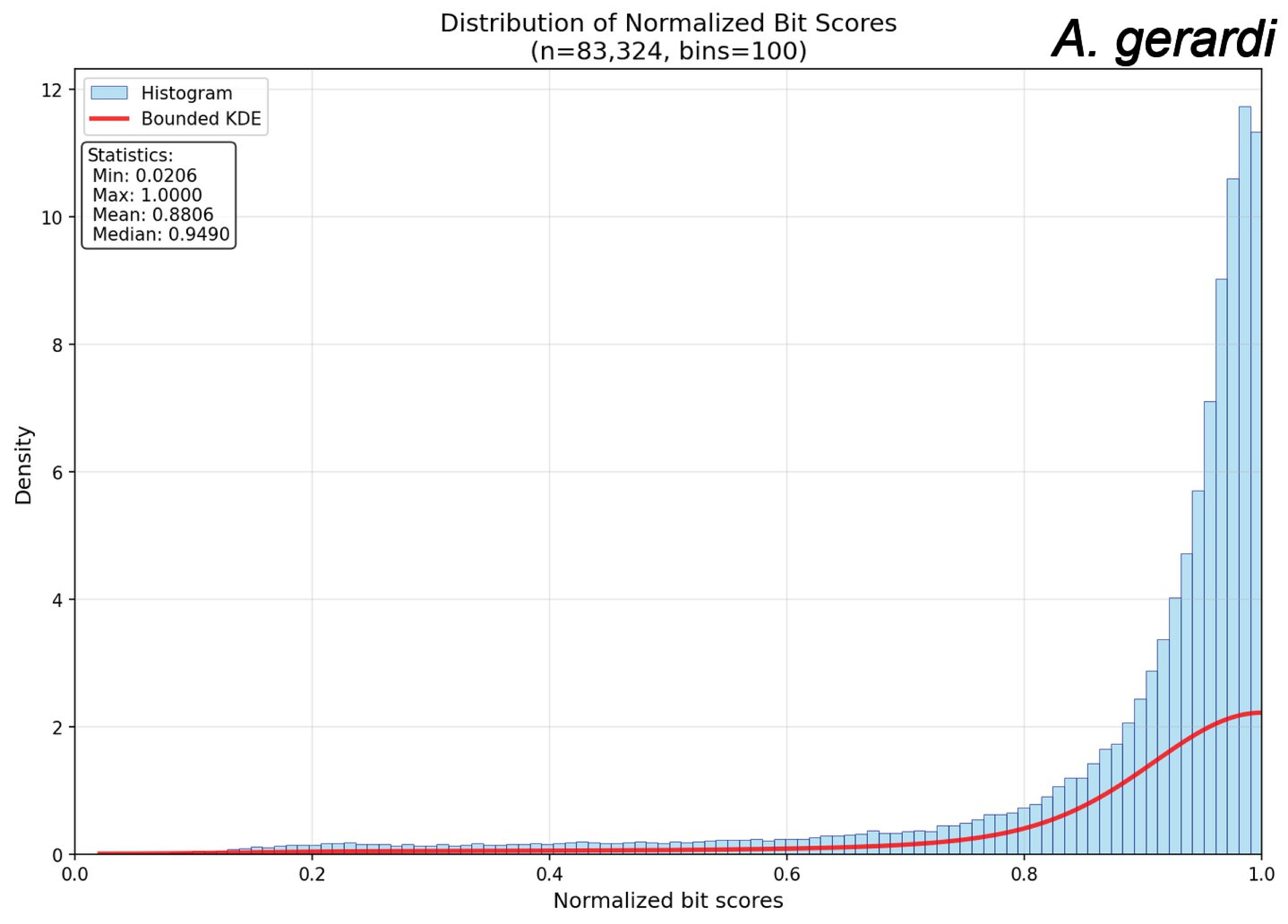
Distribution of Normalized Bit Scores
(n=83,324, bins=100)
A. gerardi
Histogram
Bounded KDE
Statistics:
Min: 0.0206
Max: 1.0000
Mean: 0.8806
Median: 0.9490
Density
Normalized bit scores
0
2
4
6
8
10
12
0.0
0.2
0.4
0.6
0.8
1.0

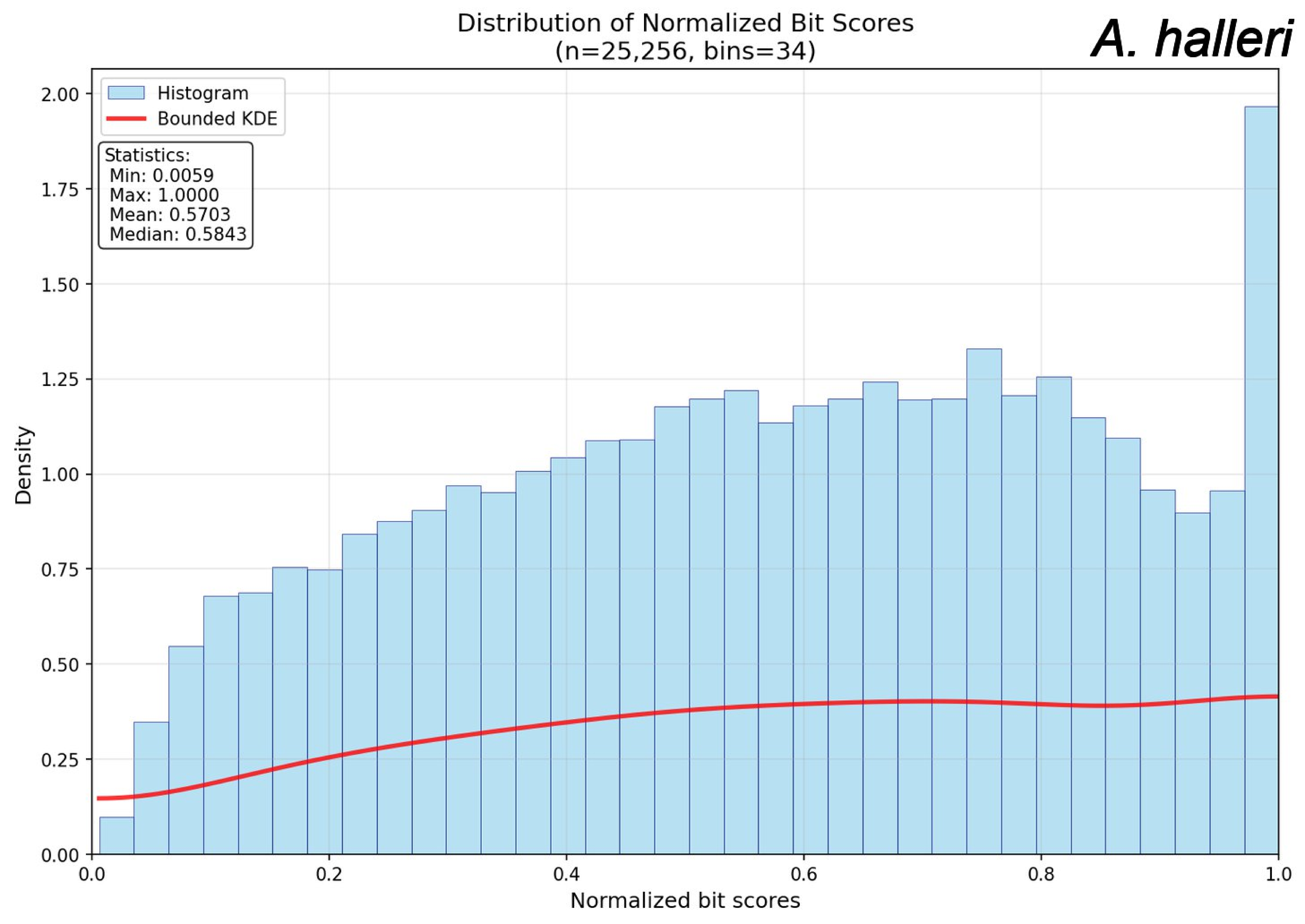
Distribution of Normalized Bit Scores
(n=25,256, bins=34)
A. halleri
Histogram
Bounded KDE
Statistics:
Min: 0.0059
Max: 1.0000
Mean: 0.5703
Median: 0.5843
Density
Normalized bit scores
0.00
0.25
0.50
0.75
1.00
1.25
1.50
1.75
2.00
0.0
0.2
0.4
0.6
0.8
1.0

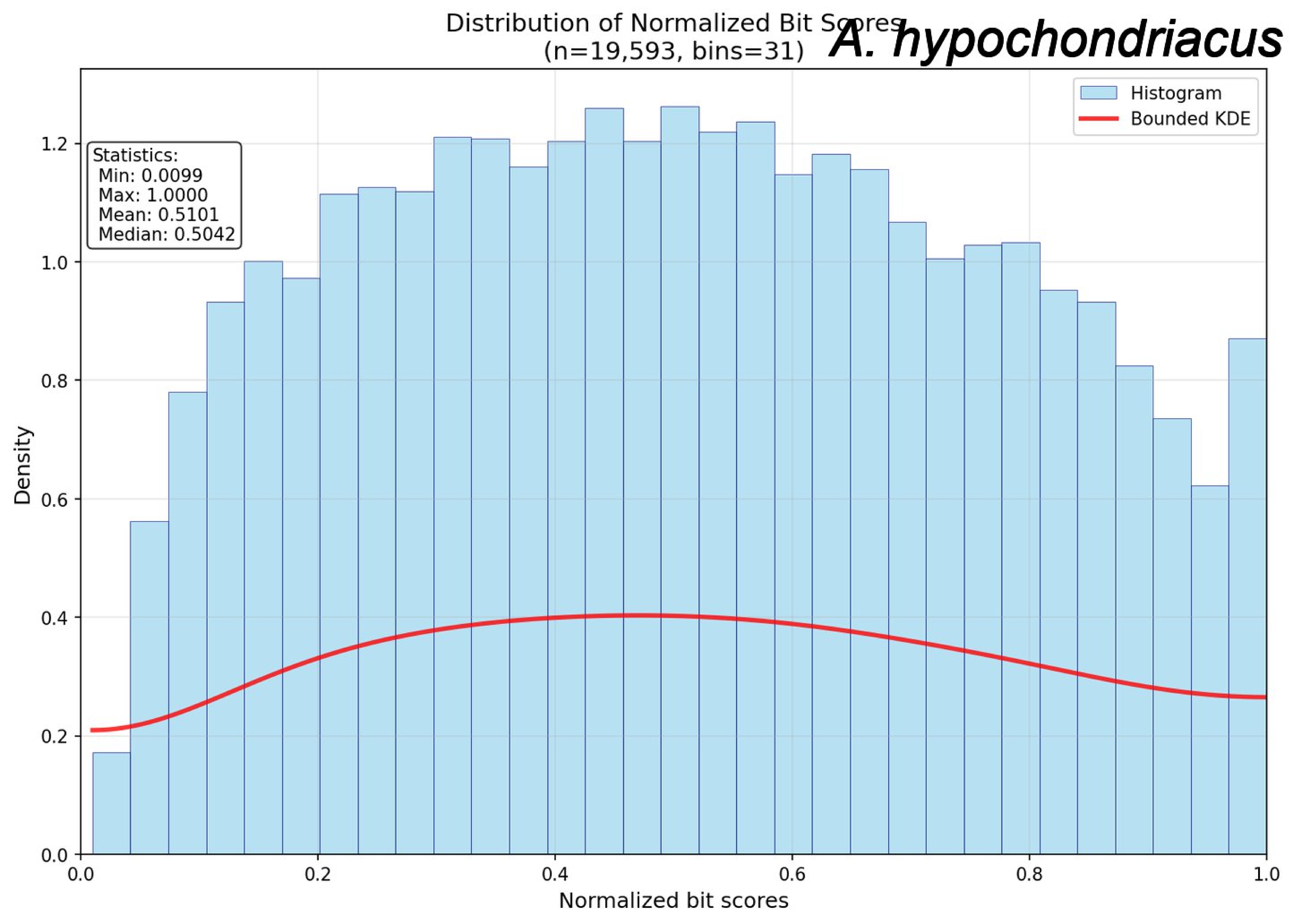

Distribution of Normalized Bit Scores
(n=19,593, bins=31)
A. hypochondriacus
Histogram
Bounded KDE
Statistics:
Min: 0.0099
Max: 1.0000
Mean: 0.5101
Median: 0.5042
Density
Normalized bit scores
0.0
0.2
0.4
0.6
0.8
1.0
1.2

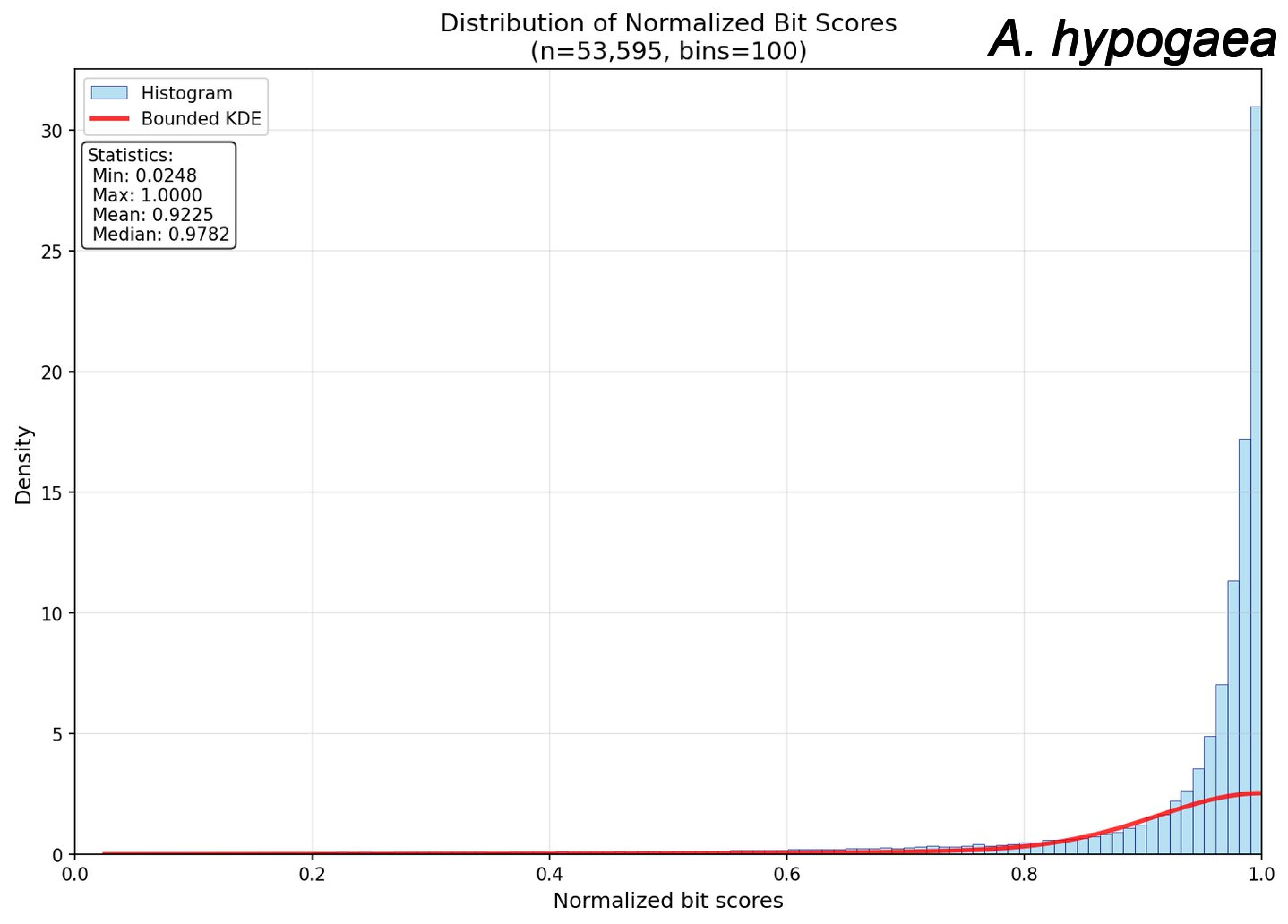
Distribution of Normalized Bit Scores
(n=53,595, bins=100)
A. hypogaea
Histogram
Bounded KDE
Statistics:
Min: 0.0248
Max: 1.0000
Mean: 0.9225
Median: 0.9782
Density
Normalized bit scores
0
5
10
15
20
25
30
0.0
0.2
0.4
0.6
0.8
1.0

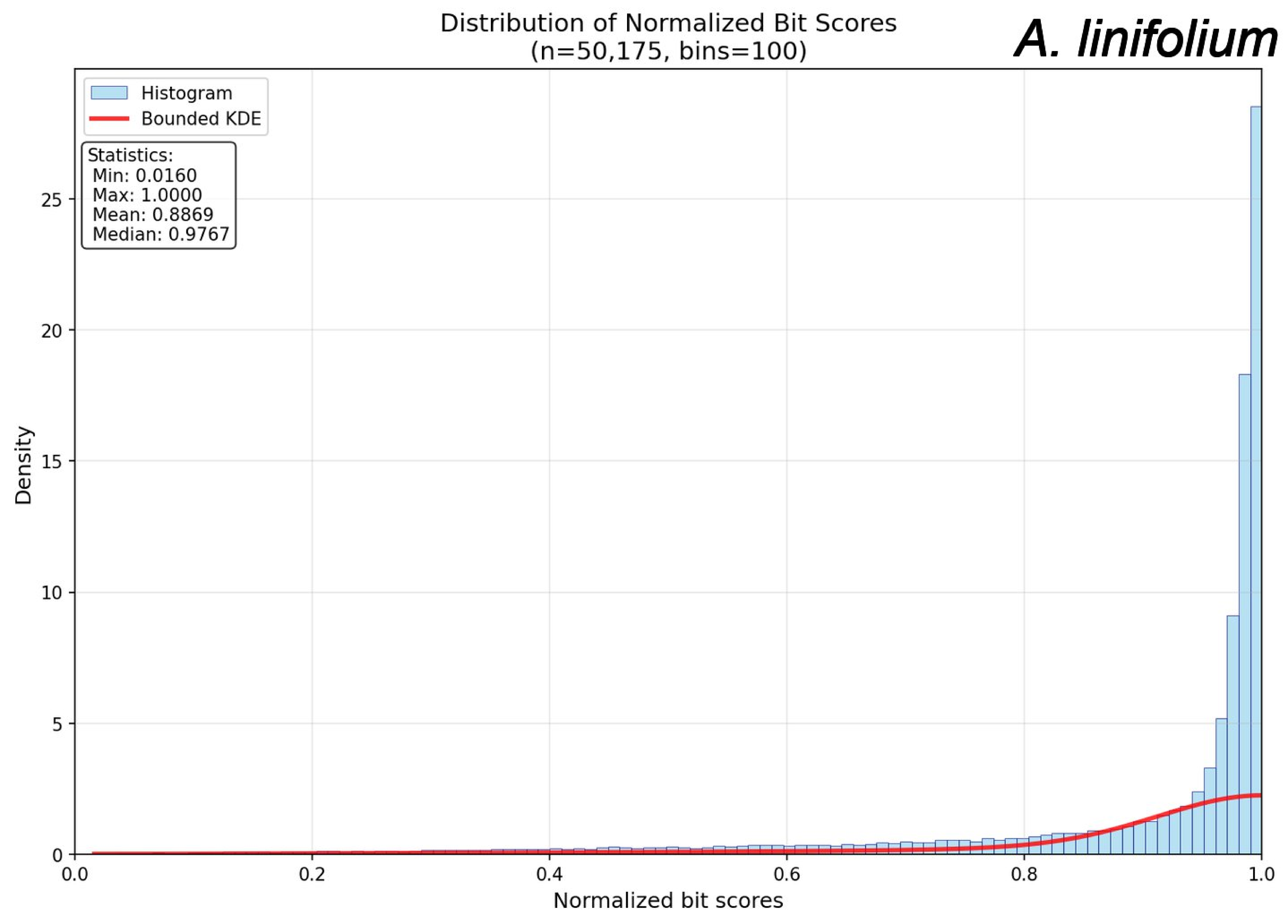

Distribution of Normalized Bit Scores
(n=50,175, bins=100)
A. linifolium
Histogram
Bounded KDE
Statistics:
Min: 0.0160
Max: 1.0000
Mean: 0.8869
Median: 0.9767
Density
Normalized bit scores
0
5
10
15
20
25
0.0
0.2
0.4
0.6
0.8
1.0

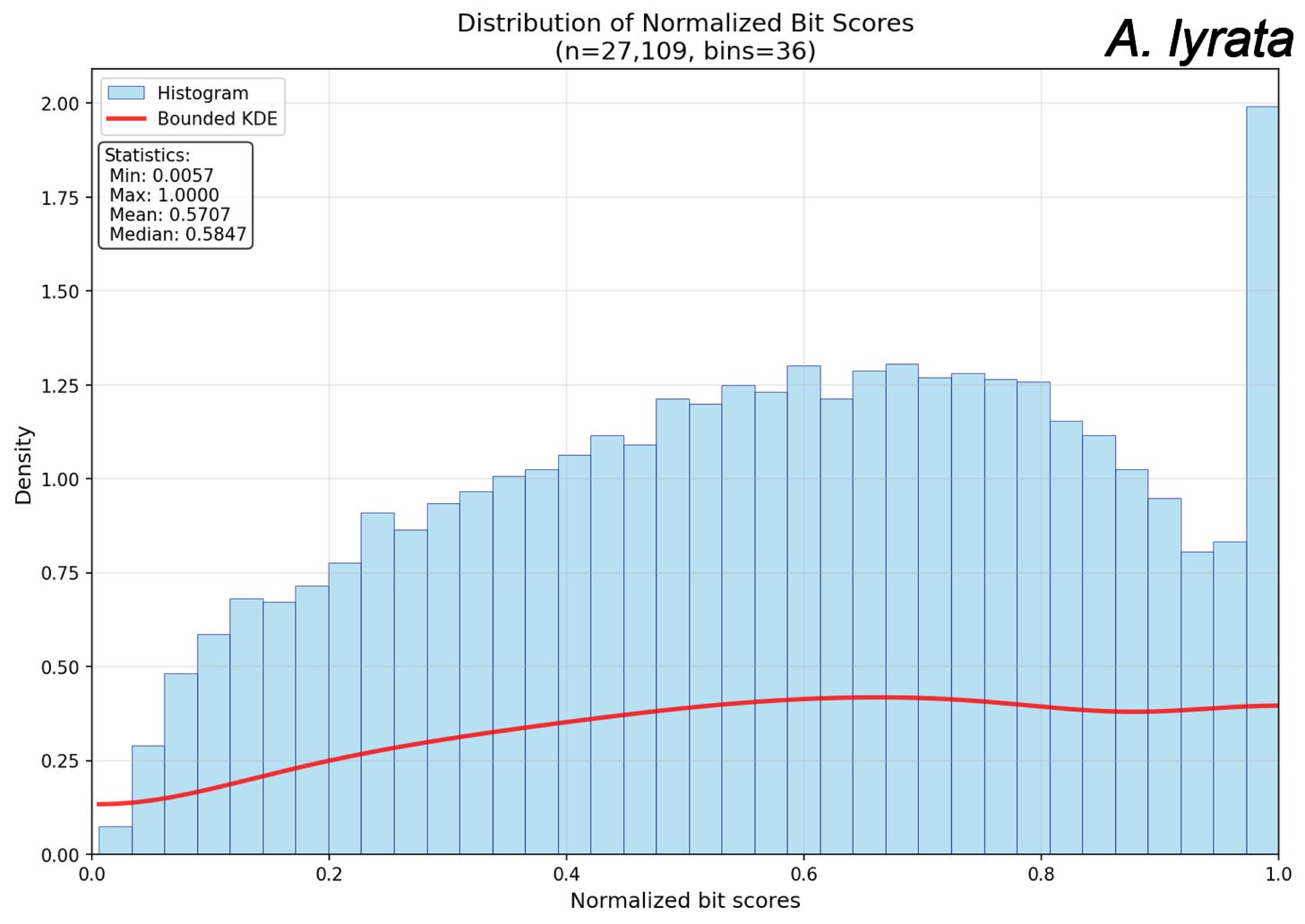
Distribution of Normalized Bit Scores
(n=27,109, bins=36)
A. lyrata
Histogram
Bounded KDE
Statistics:
Min: 0.0057
Max: 1.0000
Mean: 0.5707
Median: 0.5847
Density
Normalized bit scores
0.00
0.25
0.50
0.75
1.00
1.25
1.50
1.75
2.00
0.0
0.2
0.4
0.6
0.8
1.0

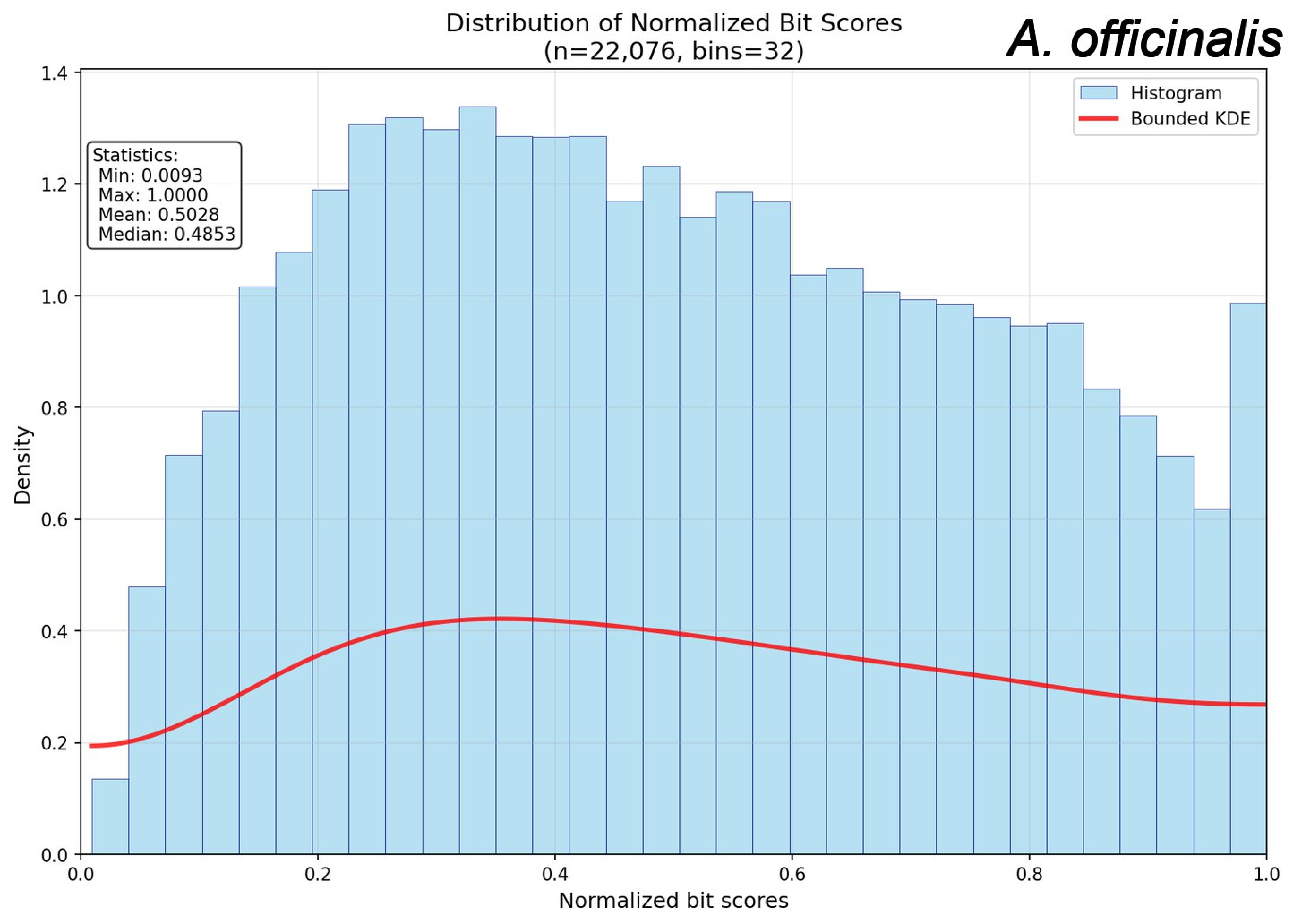

Distribution of Normalized Bit Scores
(n=22,076, bins=32)
A. officinalis
Statistics:
Min: 0.0093
Max: 1.0000
Mean: 0.5028
Median: 0.4853
Histogram
Bounded KDE
Density
Normalized bit scores
0.0
0.2
0.4
0.6
0.8
1.0
1.2
1.4

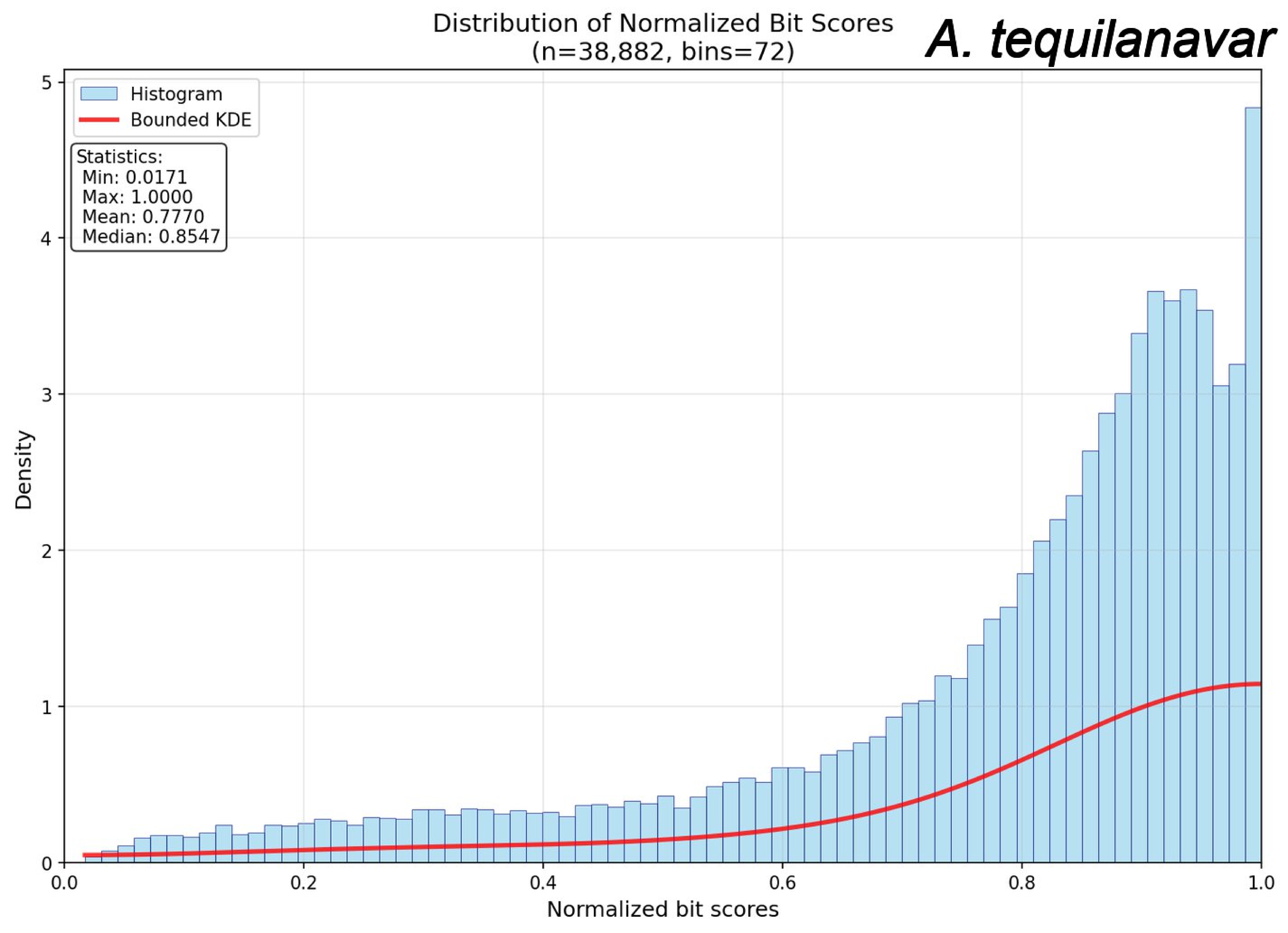

Distribution of Normalized Bit Scores
(n=38,882, bins=72)
A. tequilanavar
Histogram
Bounded KDE
Statistics:
Min: 0.0171
Max: 1.0000
Mean: 0.7770
Median: 0.8547
Density
Normalized bit scores
0.0
0.2
0.4
0.6
0.8
1.0
0
1
2
3
4
5

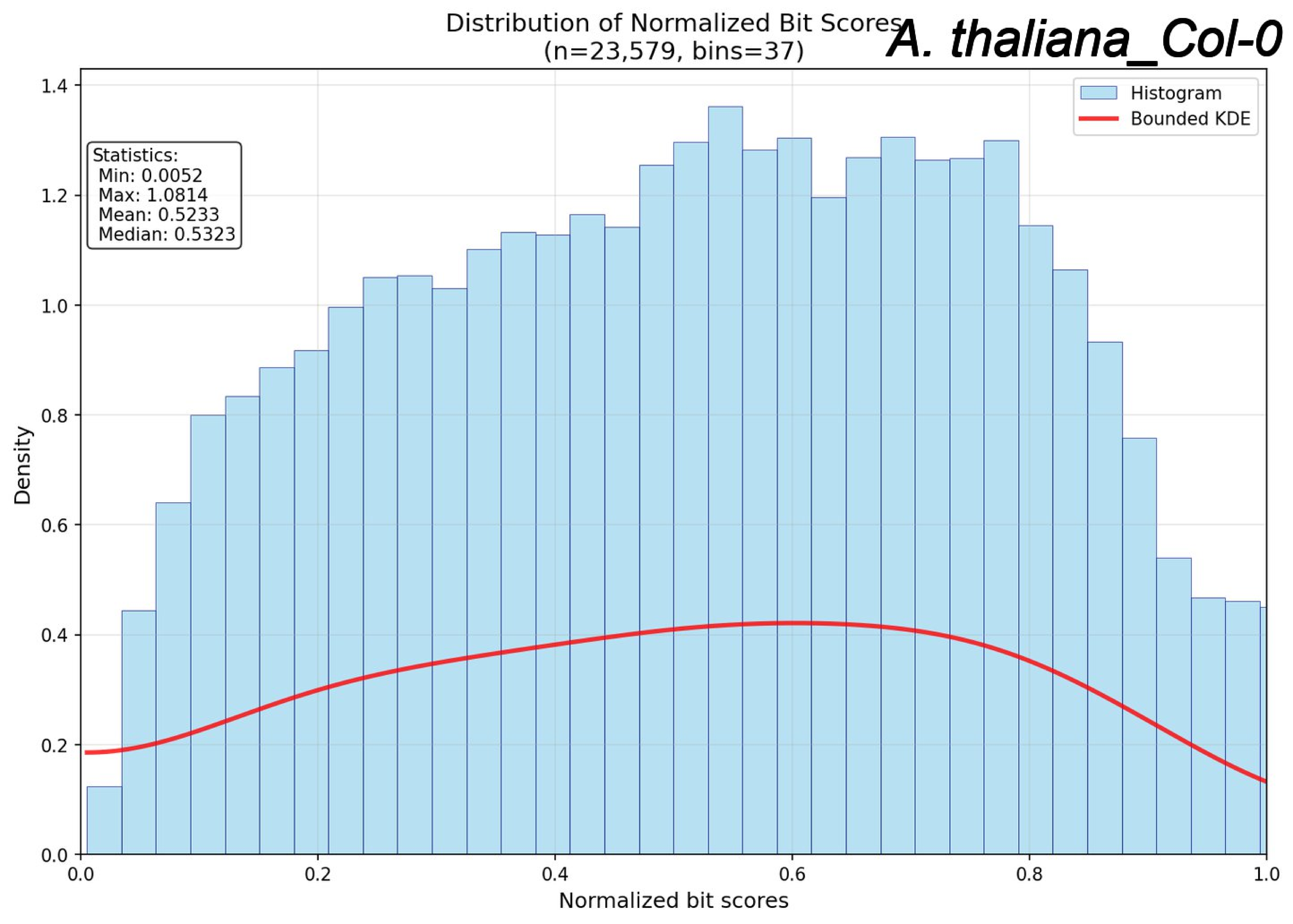

Distribution of Normalized Bit Scores
(n=23,579, bins=37)
A. thaliana_Col-0
Histogram
Bounded KDE
Statistics:
Min: 0.0052
Max: 1.0814
Mean: 0.5233
Median: 0.5323
Density
Normalized bit scores
0.0
0.2
0.4
0.6
0.8
1.0
1.2
1.4

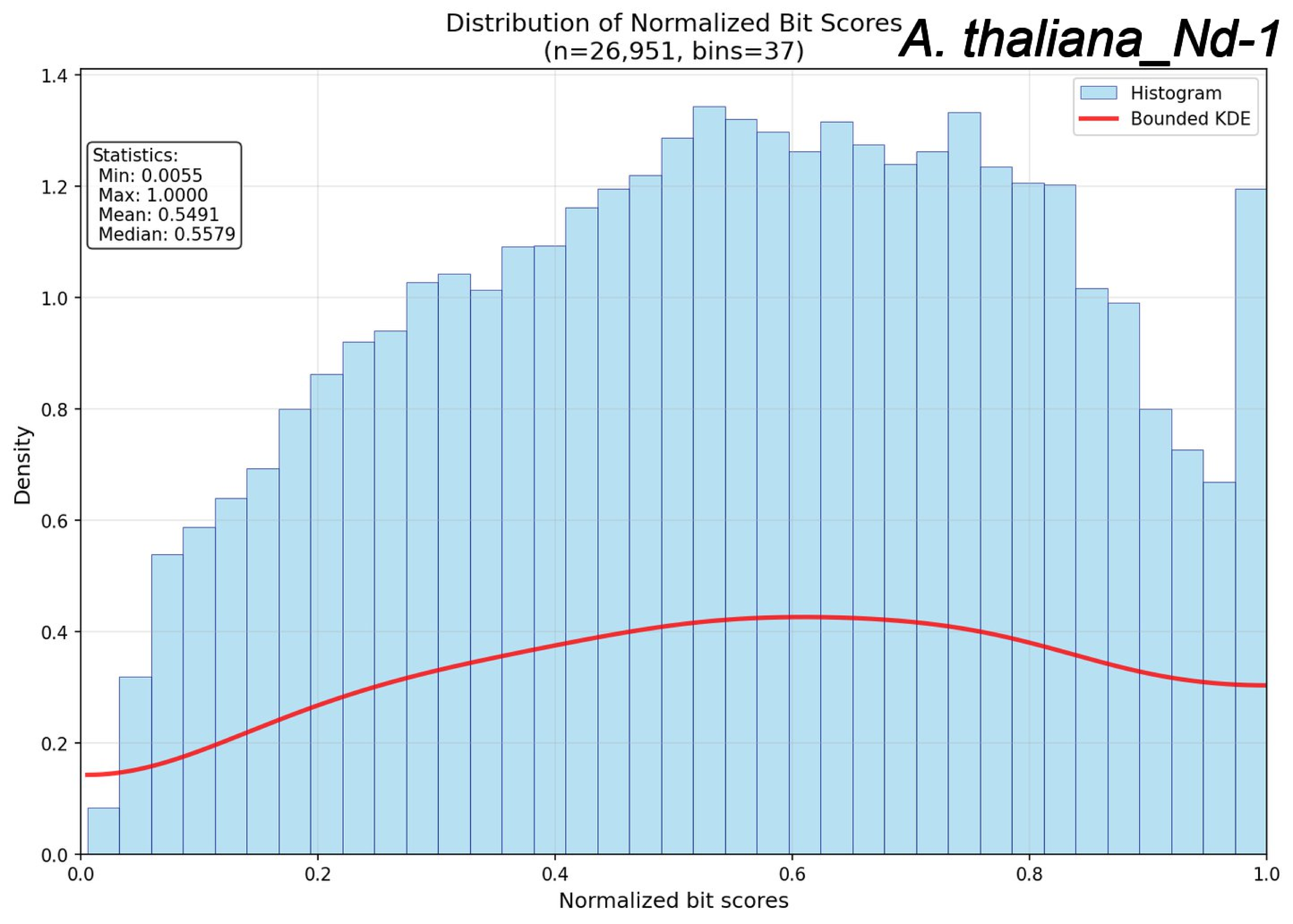
Distribution of Normalized Bit Scores
(n=26,951, bins=37)
A. thaliana_Nd-1
Histogram
Bounded KDE
Statistics:
Min: 0.0055
Max: 1.0000
Mean: 0.5491
Median: 0.5579
Density
Normalized bit scores
0.0
0.2
0.4
0.6
0.8
1.0
1.2
1.4

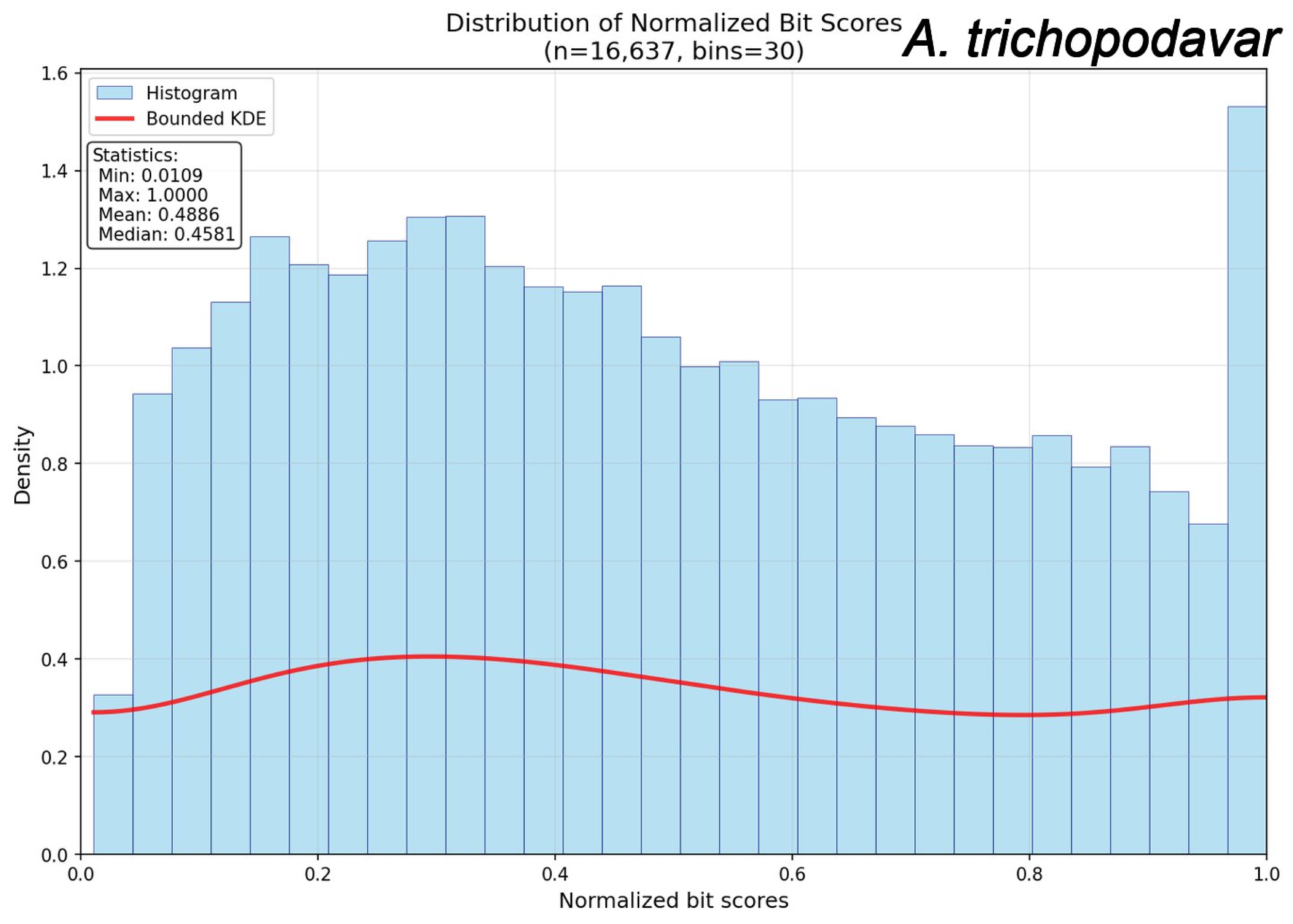

Distribution of Normalized Bit Scores
(n=16,637, bins=30)
A. trichopodavar
Histogram
Bounded KDE
Statistics:
Min: 0.0109
Max: 1.0000
Mean: 0.4886
Median: 0.4581
Density
Normalized bit scores
0.0
0.2
0.4
0.6
0.8
1.0
1.2
1.4
1.6

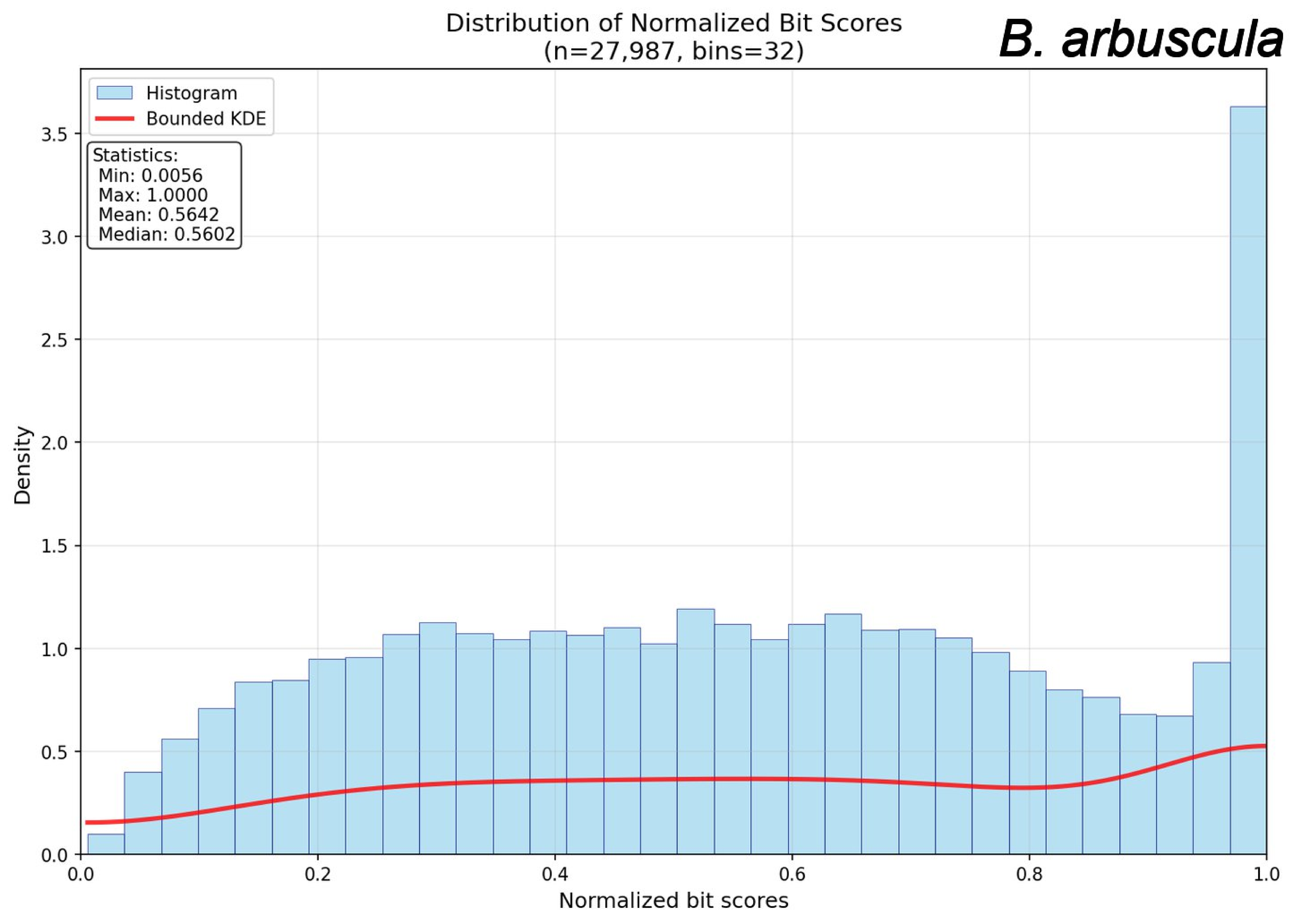
Distribution of Normalized Bit Scores
(n=27,987, bins=32)
B. arbuscula
Histogram
Bounded KDE
Statistics:
Min: 0.0056
Max: 1.0000
Mean: 0.5642
Median: 0.5602
Density
Normalized bit scores
0.0
0.5
1.0
1.5
2.0
2.5
3.0
3.5
0.0
0.2
0.4
0.6
0.8
1.0

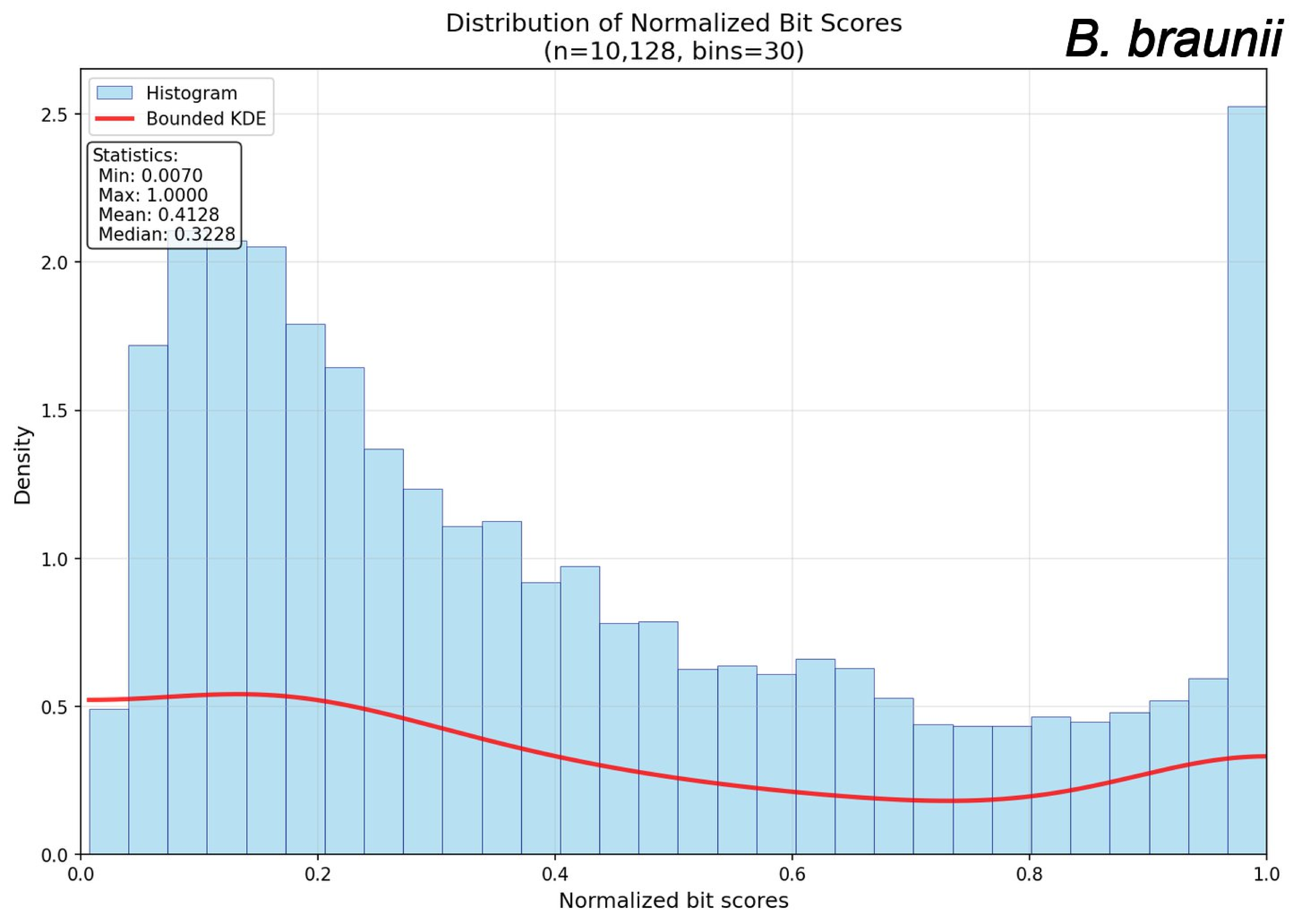

Distribution of Normalized Bit Scores
(n=10,128, bins=30)
B. braunii
Histogram
Bounded KDE
Statistics:
Min: 0.0070
Max: 1.0000
Mean: 0.4128
Median: 0.3228
Density
0.0
0.5
1.0
1.5
2.0
2.5
0.0
0.2
0.4
0.6
0.8
1.0
Normalized bit scores

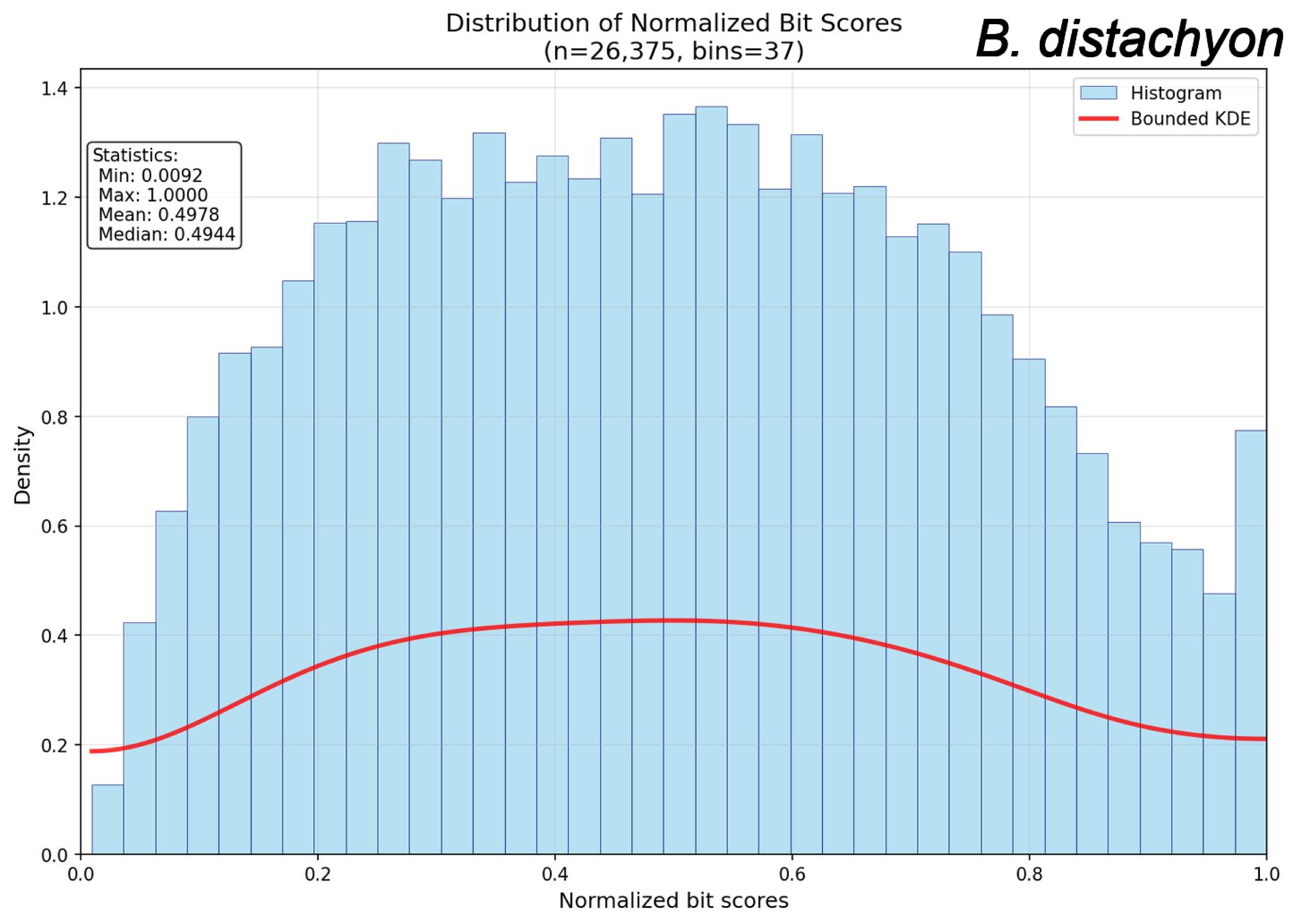

Distribution of Normalized Bit Scores
(n=26,375, bins=37)
B. distachyon
Histogram
Bounded KDE
Statistics:
Min: 0.0092
Max: 1.0000
Mean: 0.4978
Median: 0.4944
Density
Normalized bit scores
0.0
0.2
0.4
0.6
0.8
1.0
1.2
1.4

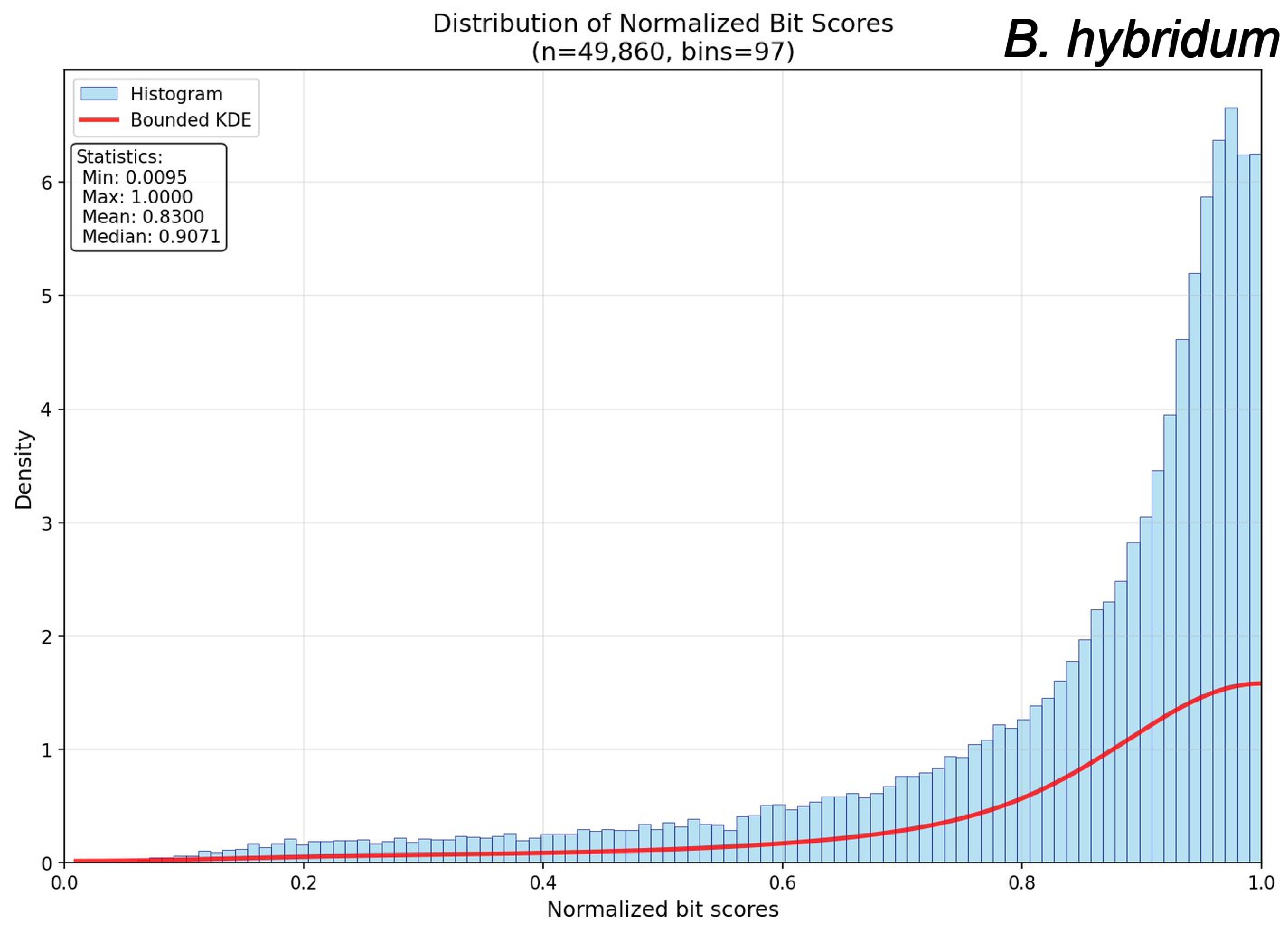
Distribution of Normalized Bit Scores
(n=49,860, bins=97)
B. hybridum
Histogram
Bounded KDE
Statistics:
Min: 0.0095
Max: 1.0000
Mean: 0.8300
Median: 0.9071
Density
Normalized bit scores
0
1
2
3
4
5
6
0.0
0.2
0.4
0.6
0.8
1.0

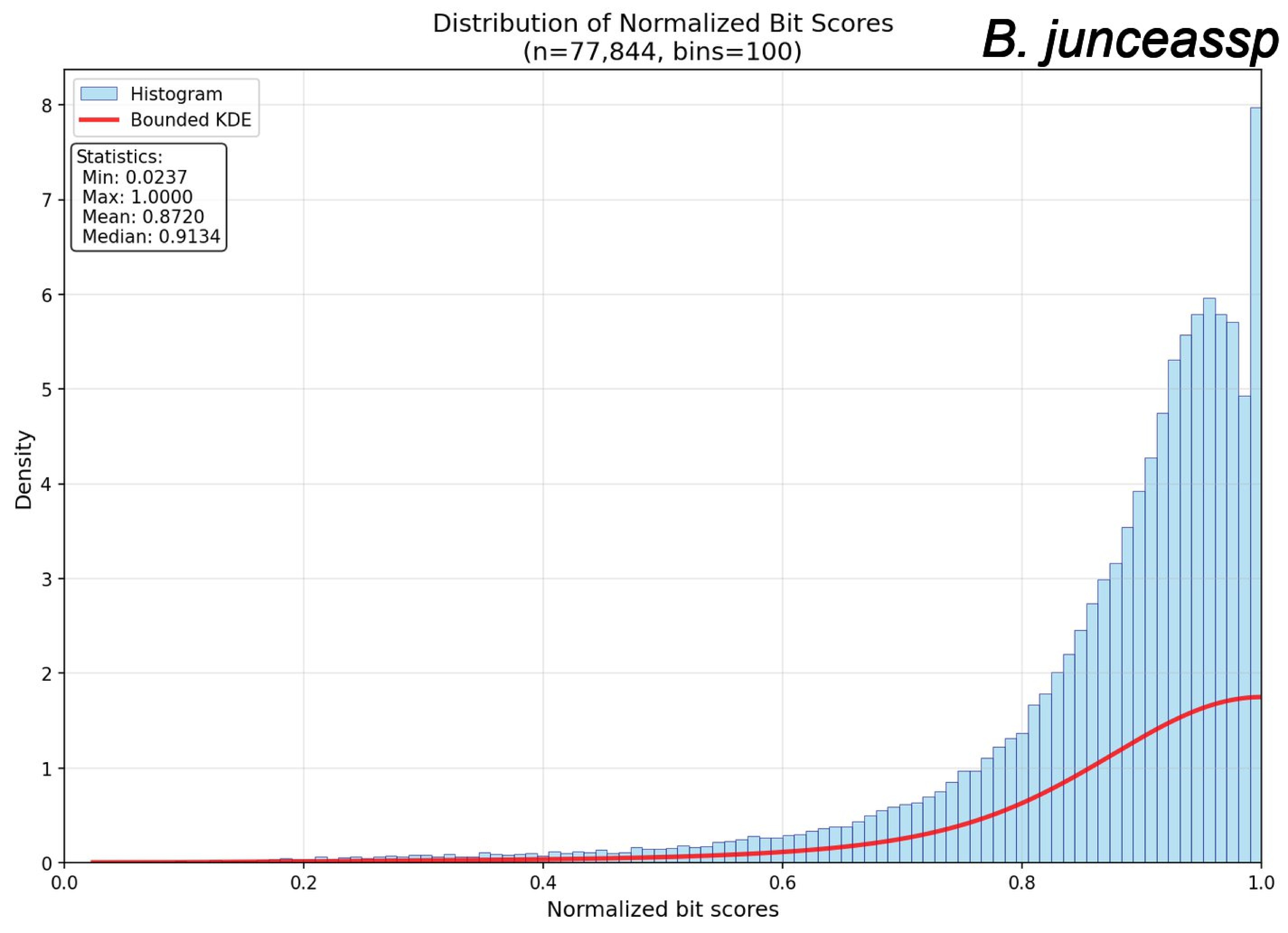

Distribution of Normalized Bit Scores
(n=77,844, bins=100)
B. junceassp
Histogram
Bounded KDE
Statistics:
Min: 0.0237
Max: 1.0000
Mean: 0.8720
Median: 0.9134
Density
Normalized bit scores
0
1
2
3
4
5
6
7
8
0.0
0.2
0.4
0.6
0.8
1.0

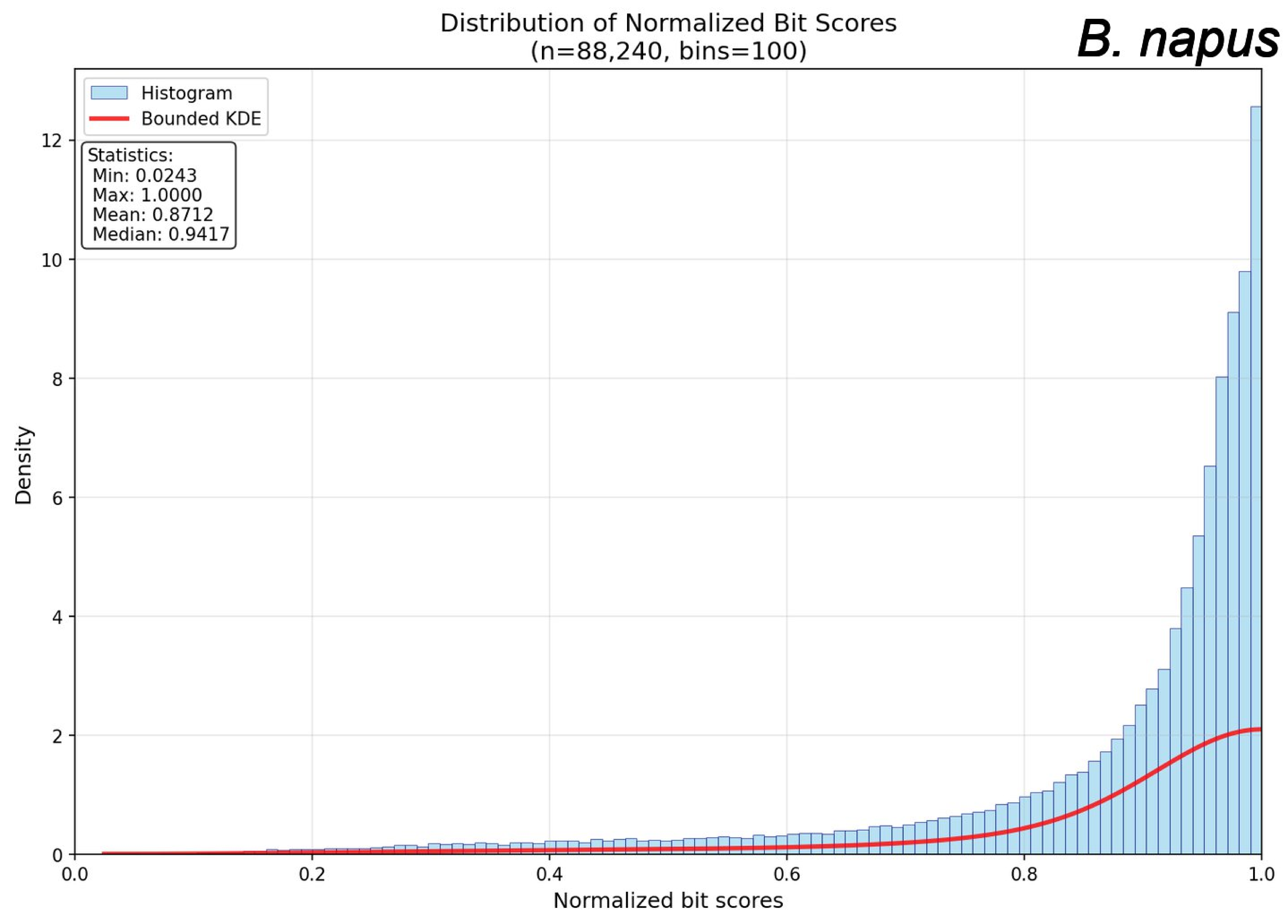

Distribution of Normalized Bit Scores
(n=88,240, bins=100)
B. napus
Histogram
Bounded KDE
Statistics:
Min: 0.0243
Max: 1.0000
Mean: 0.8712
Median: 0.9417
Density
Normalized bit scores
0
2
4
6
8
10
12
0.0
0.2
0.4
0.6
0.8
1.0

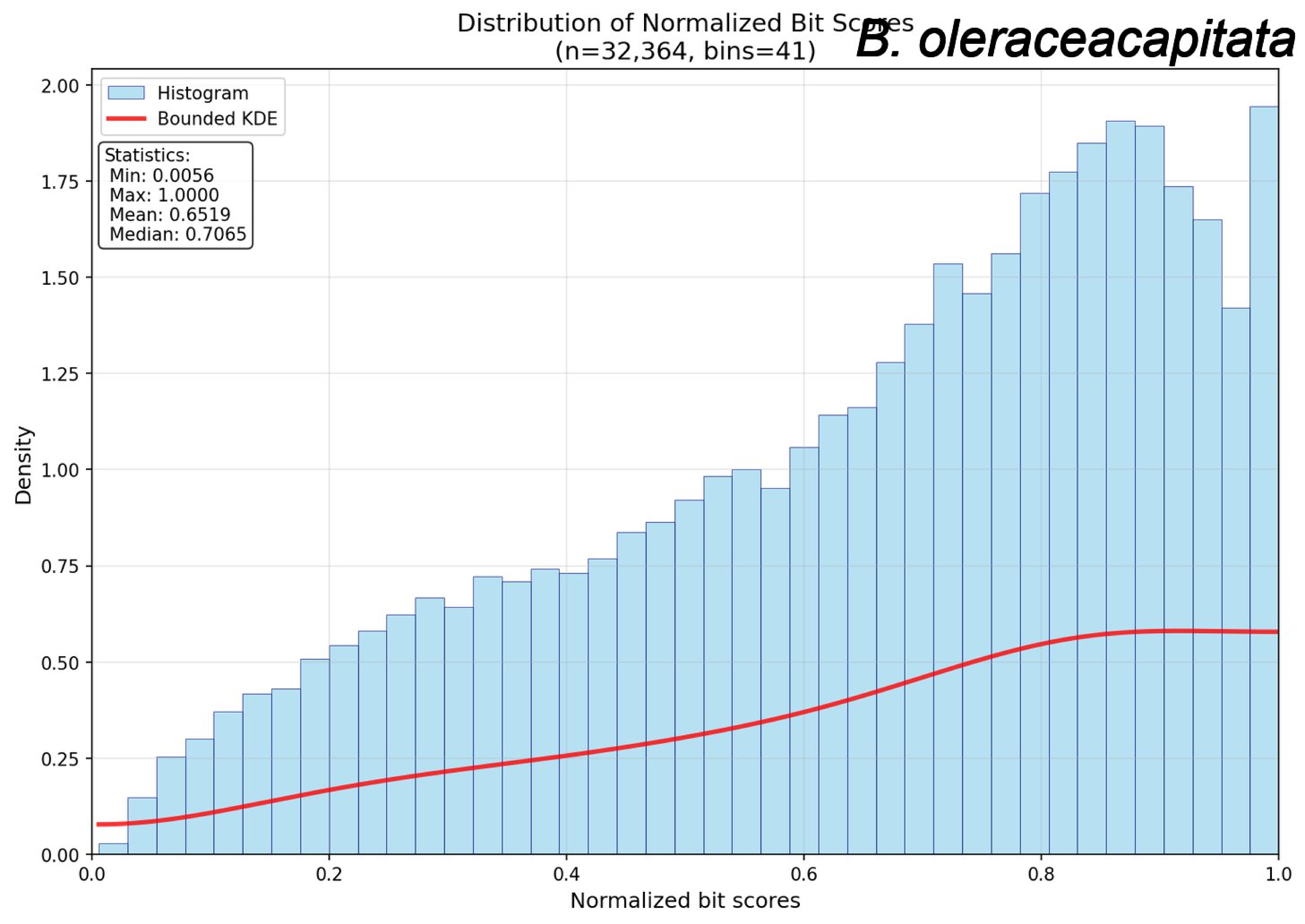
Distribution of Normalized Bit Scores
(n=32,364, bins=41)
B. oleraceacapitata
Histogram
Bounded KDE
Statistics:
Min: 0.0056
Max: 1.0000
Mean: 0.6519
Median: 0.7065
Density
Normalized bit scores
0.00
0.25
0.50
0.75
1.00
1.25
1.50
1.75
2.00
0.0
0.2
0.4
0.6
0.8
1.0

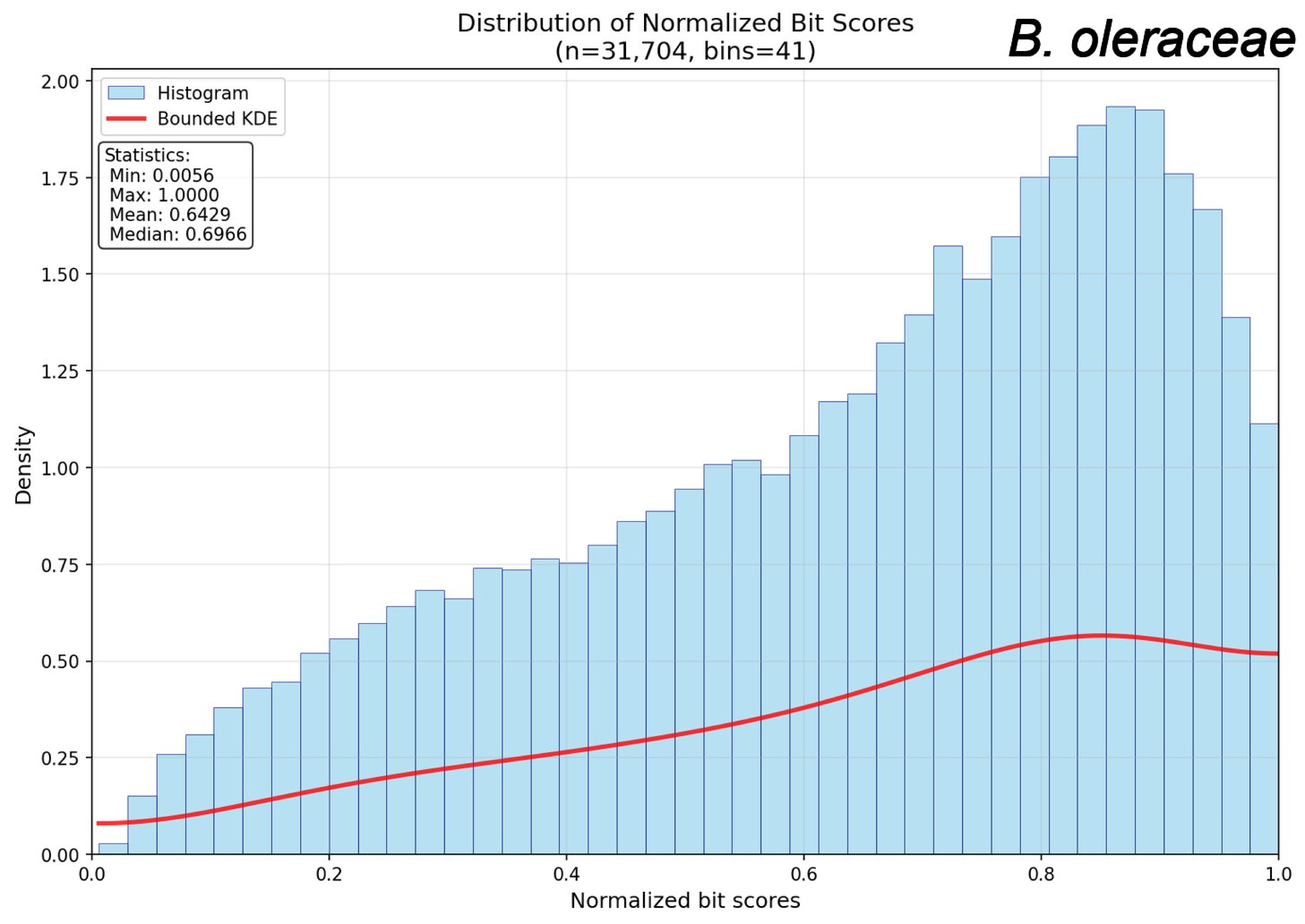

Distribution of Normalized Bit Scores
(n=31,704, bins=41)
B. oleraceae
Histogram
Bounded KDE
Statistics:
Min: 0.0056
Max: 1.0000
Mean: 0.6429
Median: 0.6966
Density
Normalized bit scores
0.00
0.25
0.50
0.75
1.00
1.25
1.50
1.75
2.00
0.0
0.2
0.4
0.6
0.8
1.0

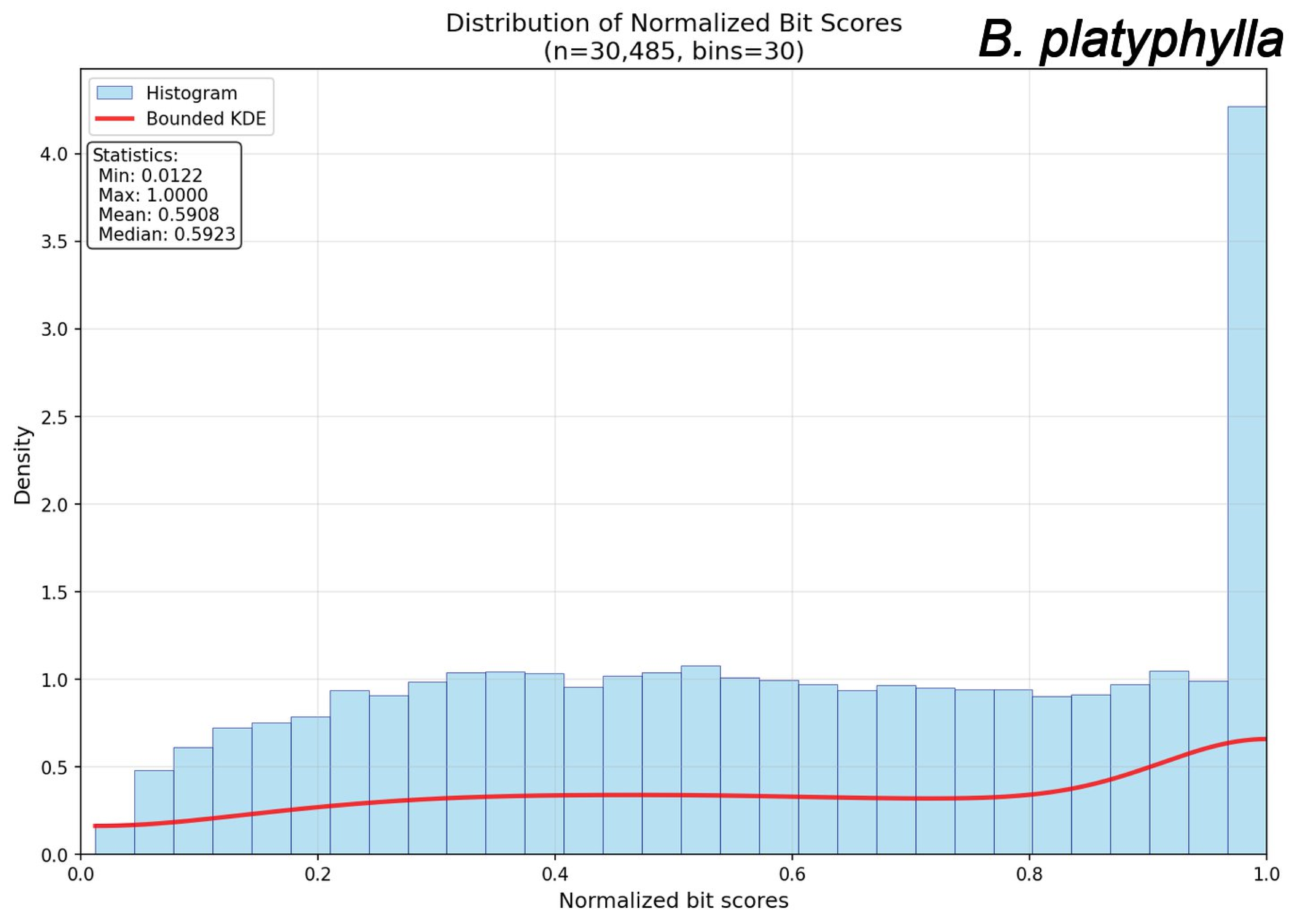

Distribution of Normalized Bit Scores
(n=30,485, bins=30)
B. platyphylla
Histogram
Bounded KDE
Statistics:
Min: 0.0122
Max: 1.0000
Mean: 0.5908
Median: 0.5923
Density
Normalized bit scores
0.0
0.5
1.0
1.5
2.0
2.5
3.0
3.5
4.0
0.2
0.4
0.6
0.8

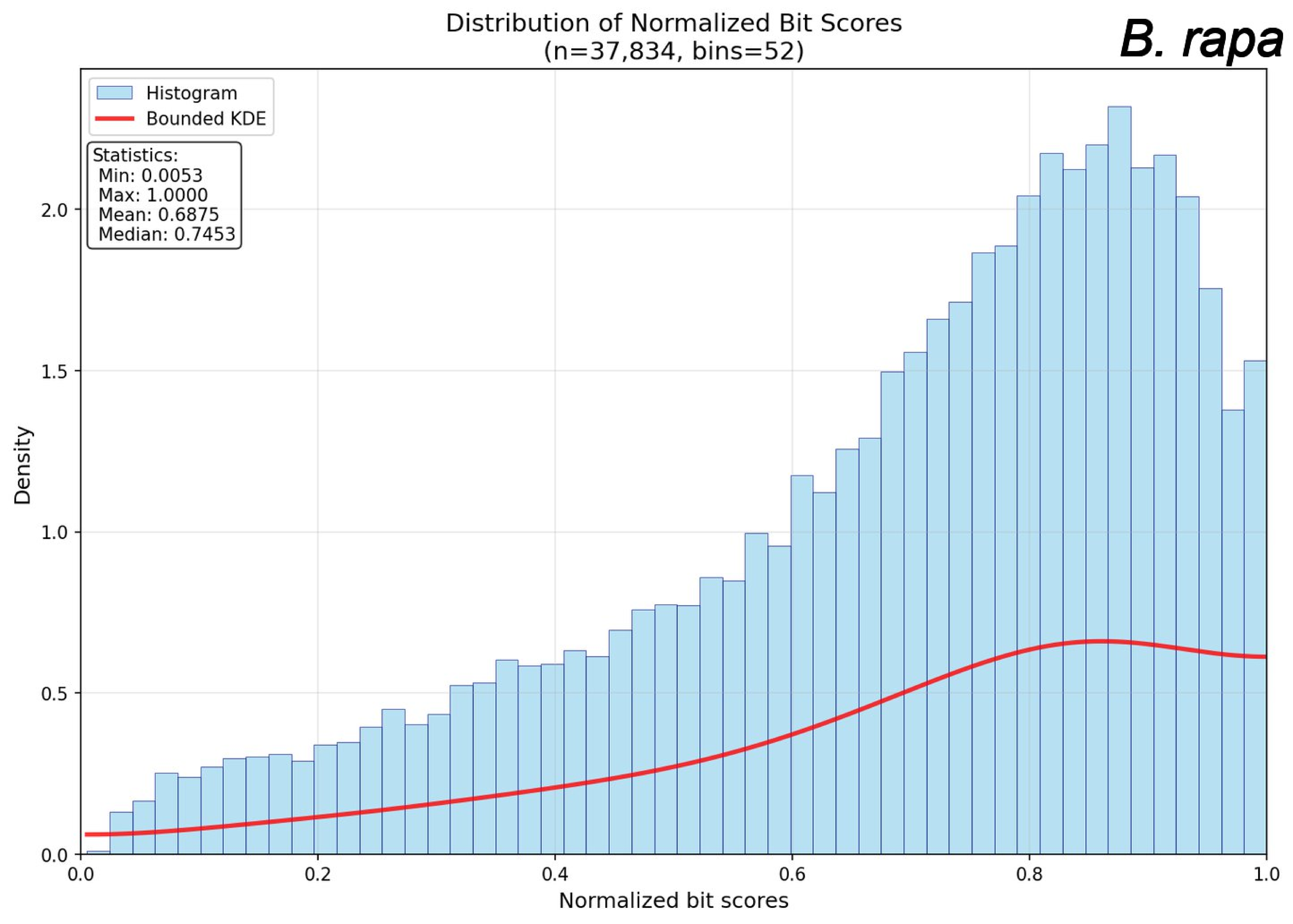
Distribution of Normalized Bit Scores
(n=37,834, bins=52)
B. rapa
Histogram
Bounded KDE
Statistics:
Min: 0.0053
Max: 1.0000
Mean: 0.6875
Median: 0.7453
Density
Normalized bit scores
0.0
0.5
1.0
1.5
2.0
0.2
0.4
0.6
0.8

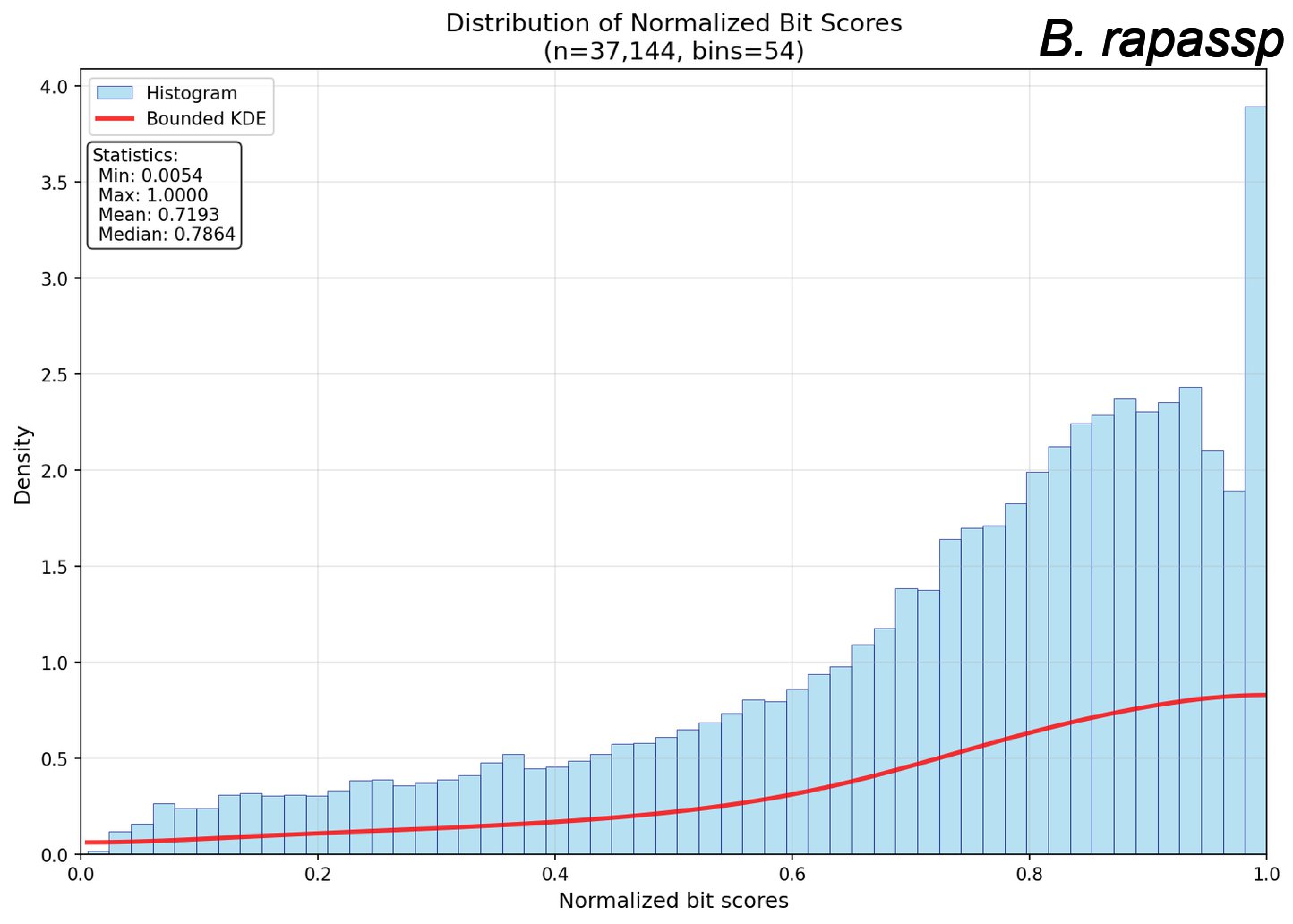

Distribution of Normalized Bit Scores
(n=37,144, bins=54)
B. rapassp
Histogram
Bounded KDE
Statistics:
Min: 0.0054
Max: 1.0000
Mean: 0.7193
Median: 0.7864
Density
Normalized bit scores
0.0
0.5
1.0
1.5
2.0
2.5
3.0
3.5
4.0
0.2
0.4
0.6
0.8

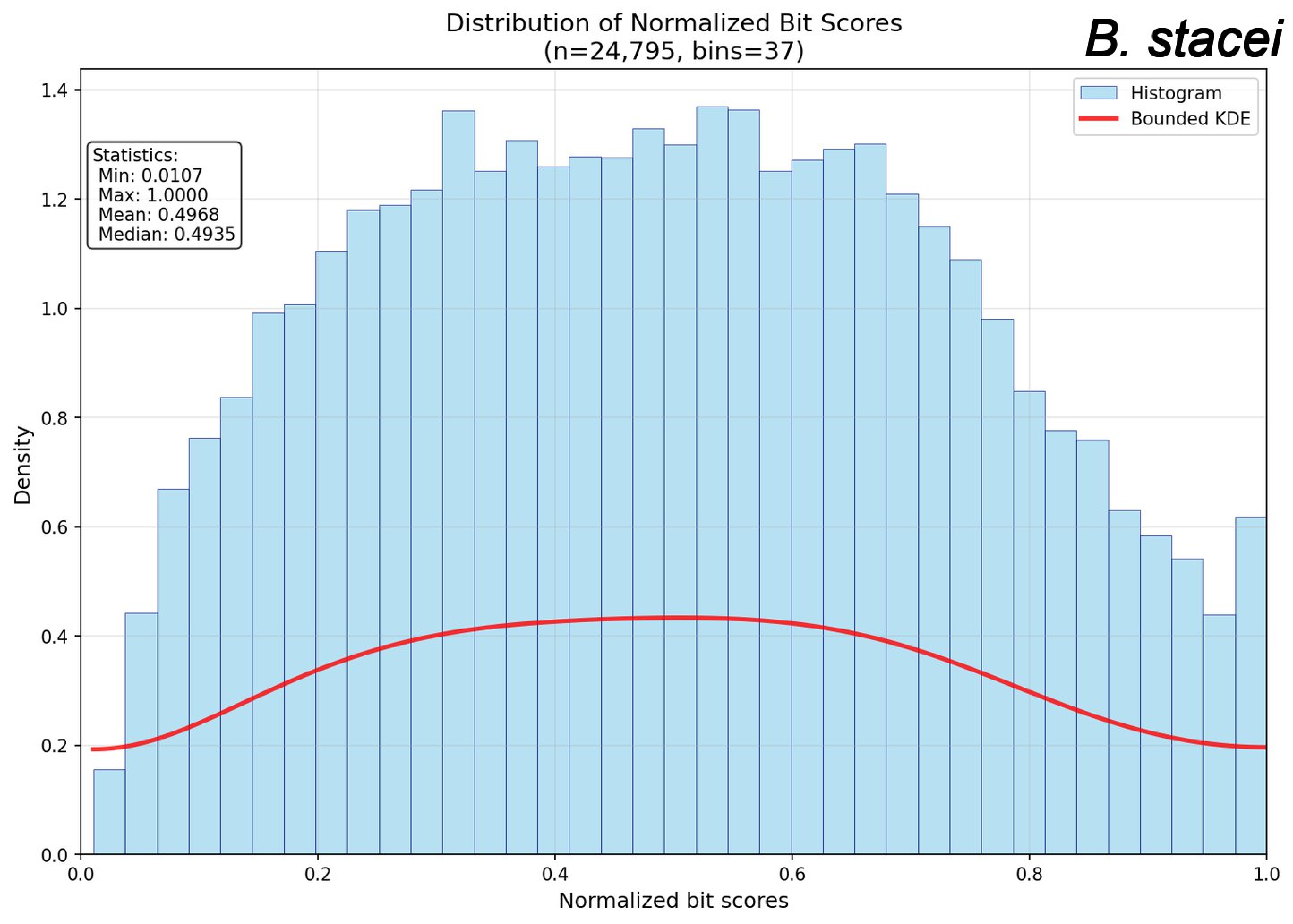
Distribution of Normalized Bit Scores
(n=24,795, bins=37)
B. stacei
Histogram
Bounded KDE
Statistics:
Min: 0.0107
Max: 1.0000
Mean: 0.4968
Median: 0.4935
Density
Normalized bit scores
0.0
0.2
0.4
0.6
0.8
1.0
1.2
1.4

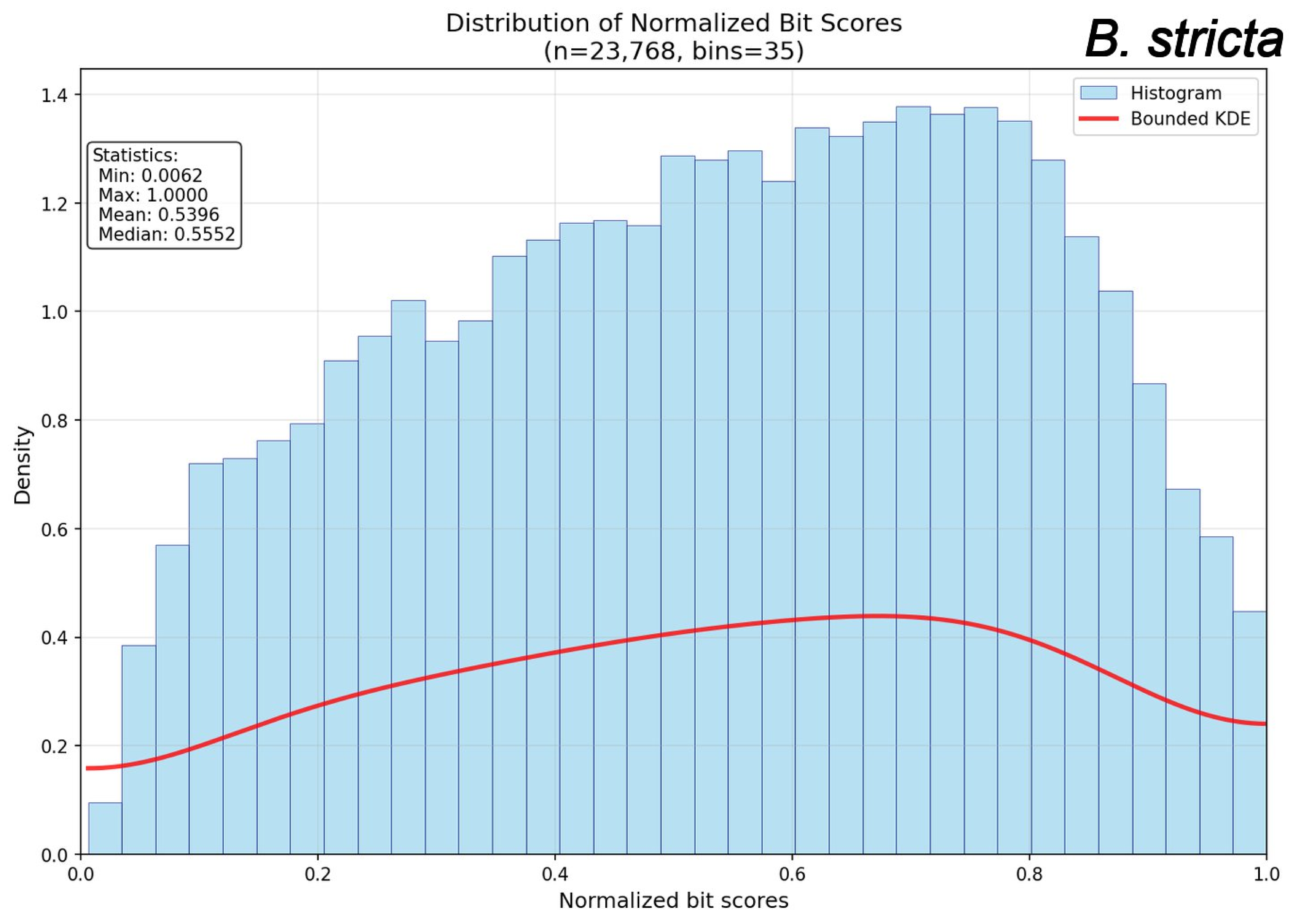
Distribution of Normalized Bit Scores
(n=23,768, bins=35)
B. stricta
Histogram
Bounded KDE
Statistics:
Min: 0.0062
Max: 1.0000
Mean: 0.5396
Median: 0.5552
Density
Normalized bit scores
0.0
0.2
0.4
0.6
0.8
1.0
1.2
1.4

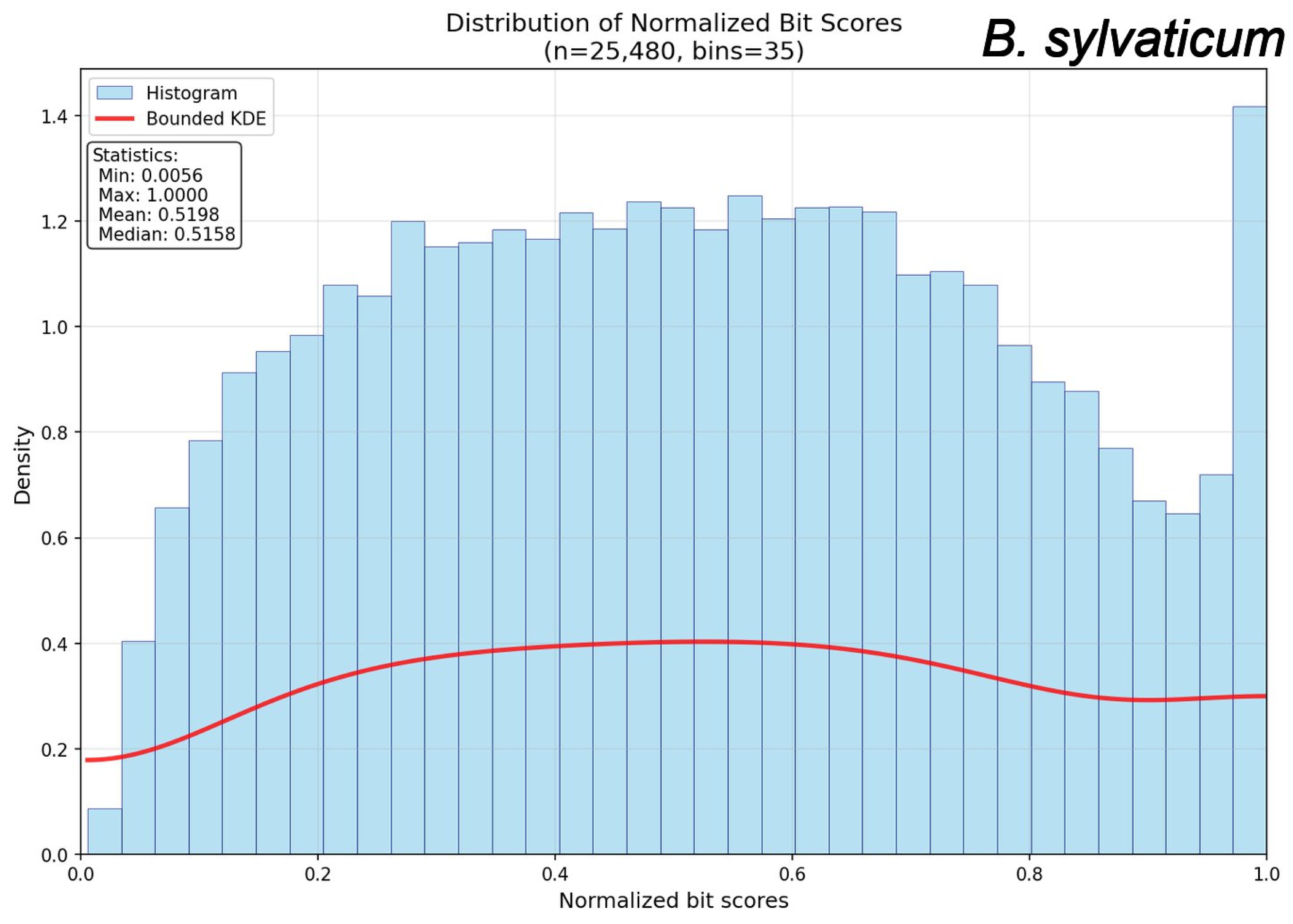

Distribution of Normalized Bit Scores
(n=25,480, bins=35)
B. sylvaticum
Histogram
Bounded KDE
Statistics:
Min: 0.0056
Max: 1.0000
Mean: 0.5198
Median: 0.5158
Density
Normalized bit scores
0.0
0.2
0.4
0.6
0.8
1.0
1.2
1.4

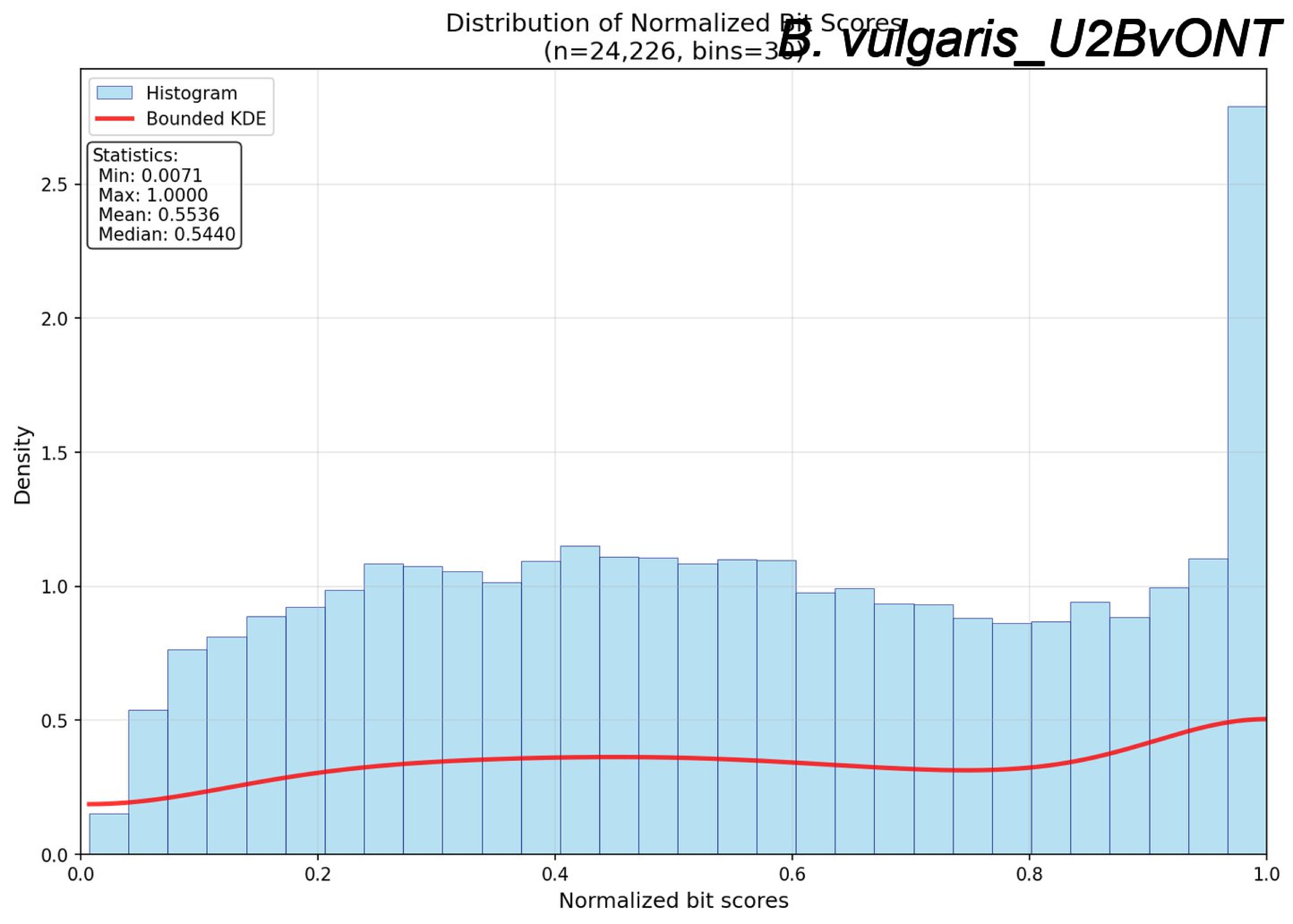
Distribution of Normalized Bit Scores
(n=24,226, bins=30)
B. vulgaris_U2BvONT
Histogram
Bounded KDE
Statistics:
Min: 0.0071
Max: 1.0000
Mean: 0.5536
Median: 0.5440
Density
Normalized bit scores
0.0
0.5
1.0
1.5
2.0
2.5
0.2
0.4
0.6
0.8

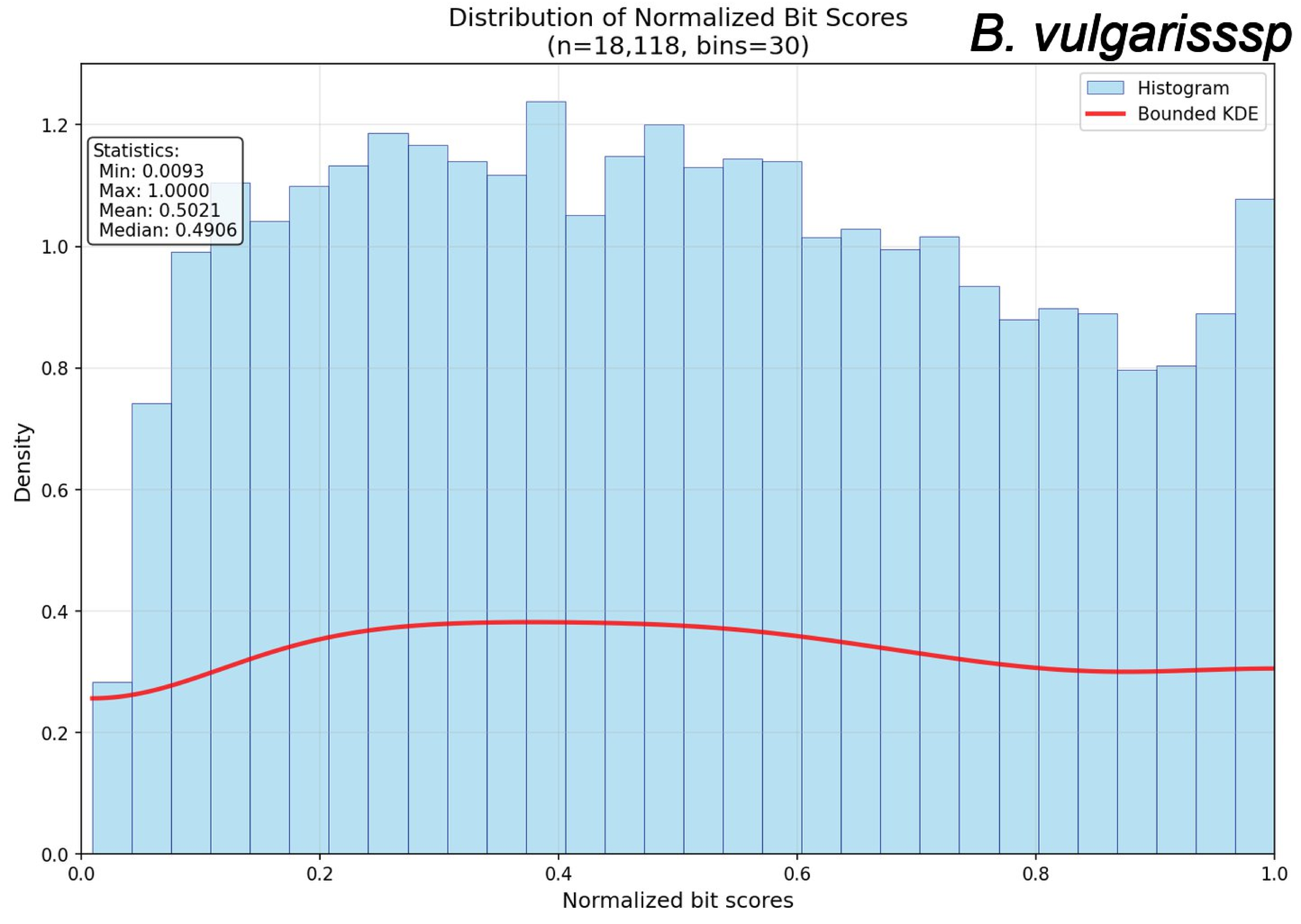
Distribution of Normalized Bit Scores
(n=18,118, bins=30)
B. vulgarisssp
Histogram
Bounded KDE
Statistics:
Min: 0.0093
Max: 1.0000
Mean: 0.5021
Median: 0.4906
Density
Normalized bit scores
0.0
0.2
0.4
0.6
0.8
1.0
1.2

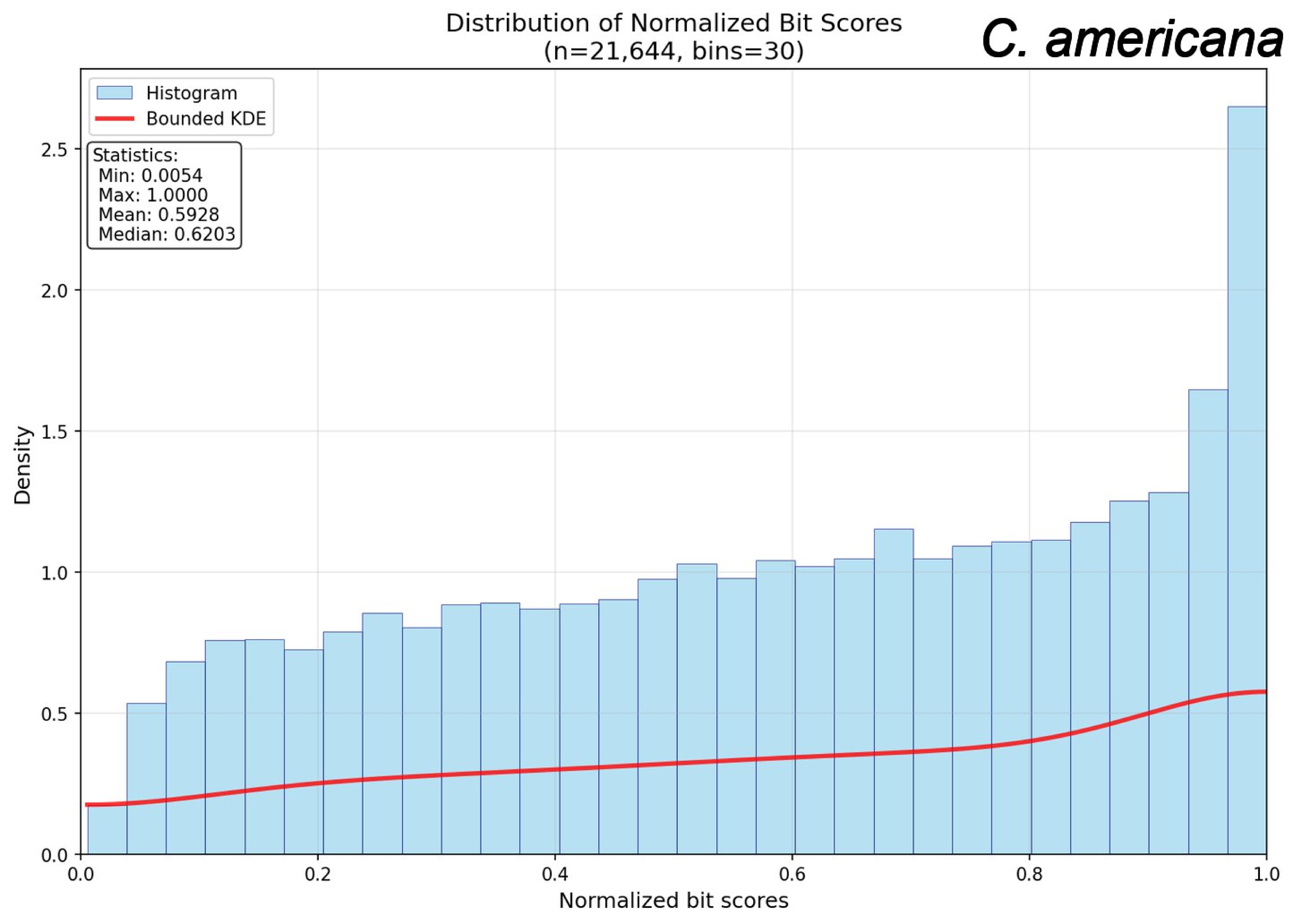

Distribution of Normalized Bit Scores
(n=21,644, bins=30)
C. americana
Histogram
Bounded KDE
Statistics:
Min: 0.0054
Max: 1.0000
Mean: 0.5928
Median: 0.6203
Density
Normalized bit scores
0.0
0.5
1.0
1.5
2.0
2.5
0.0
0.2
0.4
0.6
0.8
1.0

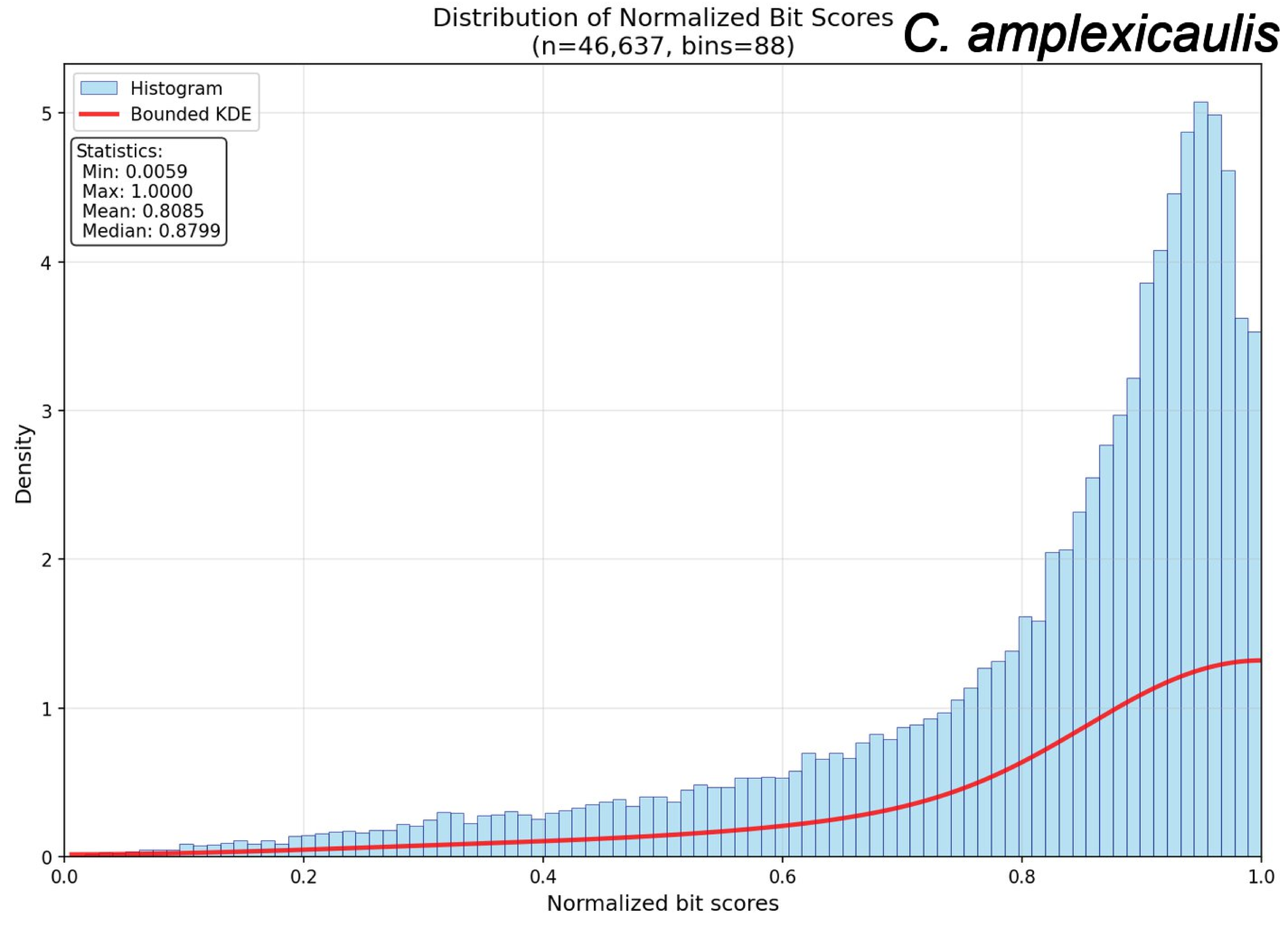

Distribution of Normalized Bit Scores
(n=46,637, bins=88)
C. amplexicaulis
Histogram
Bounded KDE
Statistics:
Min: 0.0059
Max: 1.0000
Mean: 0.8085
Median: 0.8799
Density
Normalized bit scores
0
1
2
3
4
5
0.0
0.2
0.4
0.6
0.8
1.0

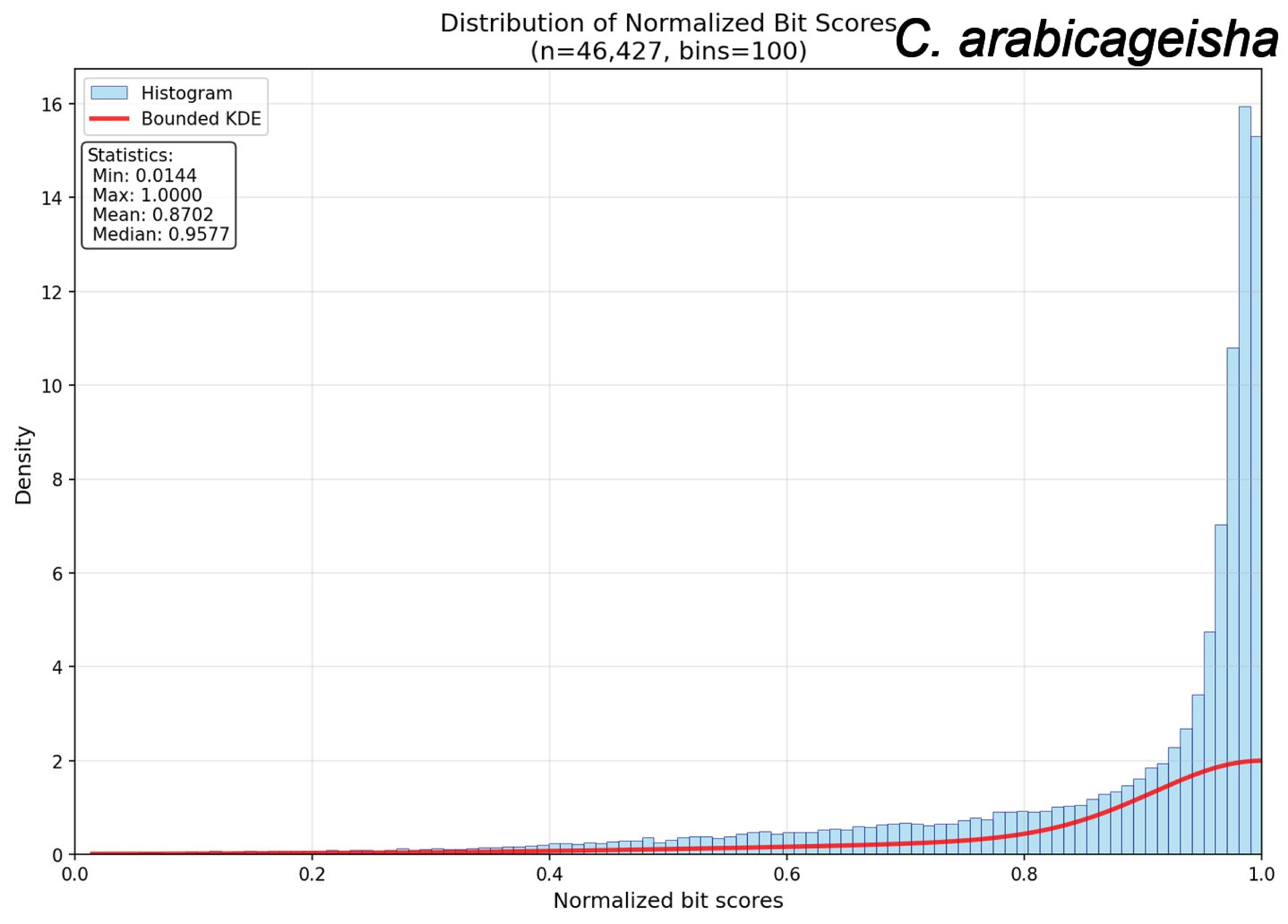

Distribution of Normalized Bit Scores
(n=46,427, bins=100)
C. arabicageisha
Histogram
Bounded KDE
Statistics:
Min: 0.0144
Max: 1.0000
Mean: 0.8702
Median: 0.9577
Density
Normalized bit scores
0
2
4
6
8
10
12
14
16
0.0
0.2
0.4
0.6
0.8
1.0

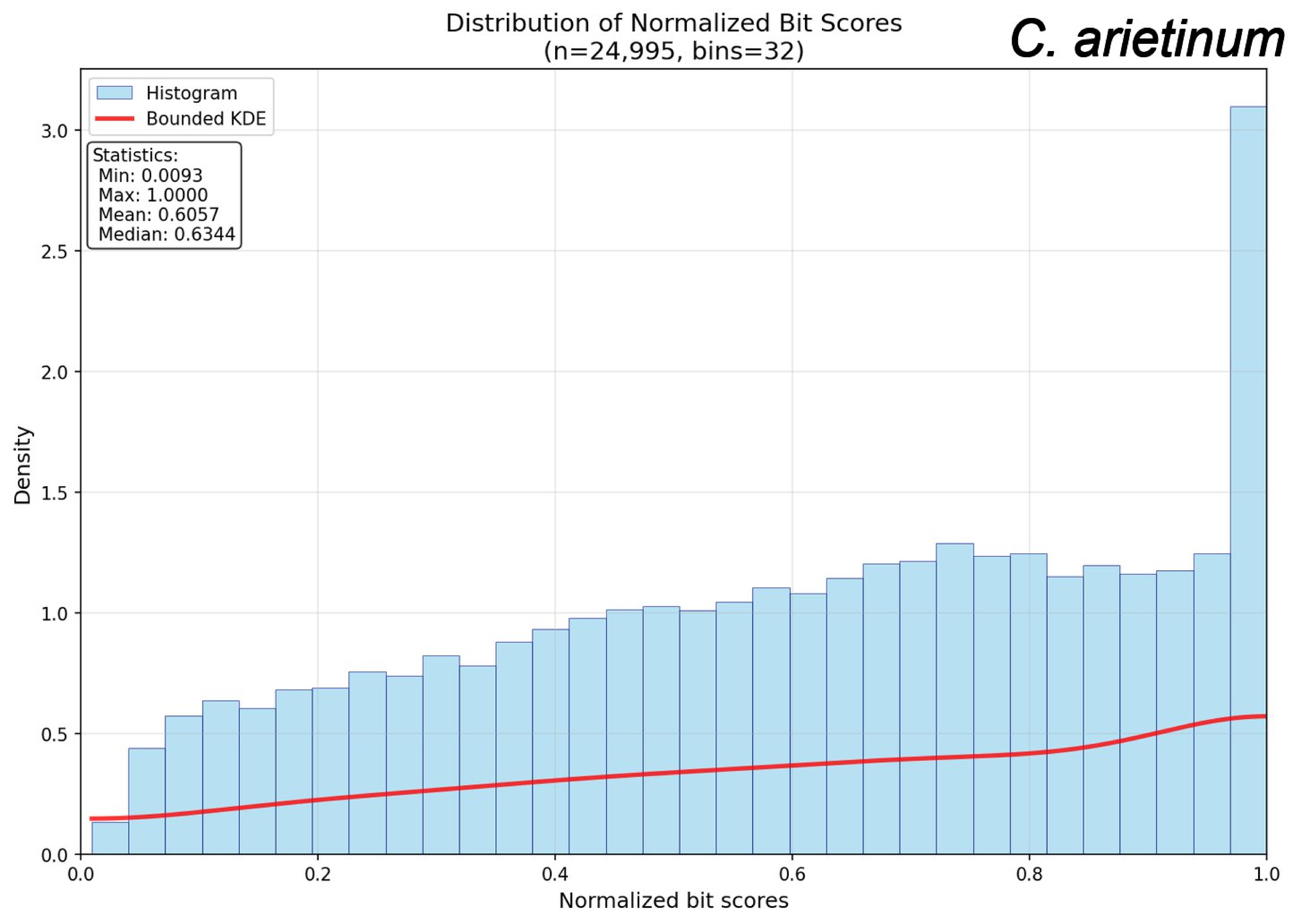
Distribution of Normalized Bit Scores
(n=24,995, bins=32)
C. arietinum
Histogram
Bounded KDE
Statistics:
Min: 0.0093
Max: 1.0000
Mean: 0.6057
Median: 0.6344
Density
Normalized bit scores
0.0
0.5
1.0
1.5
2.0
2.5
3.0
0.0
0.2
0.4
0.6
0.8
1.0

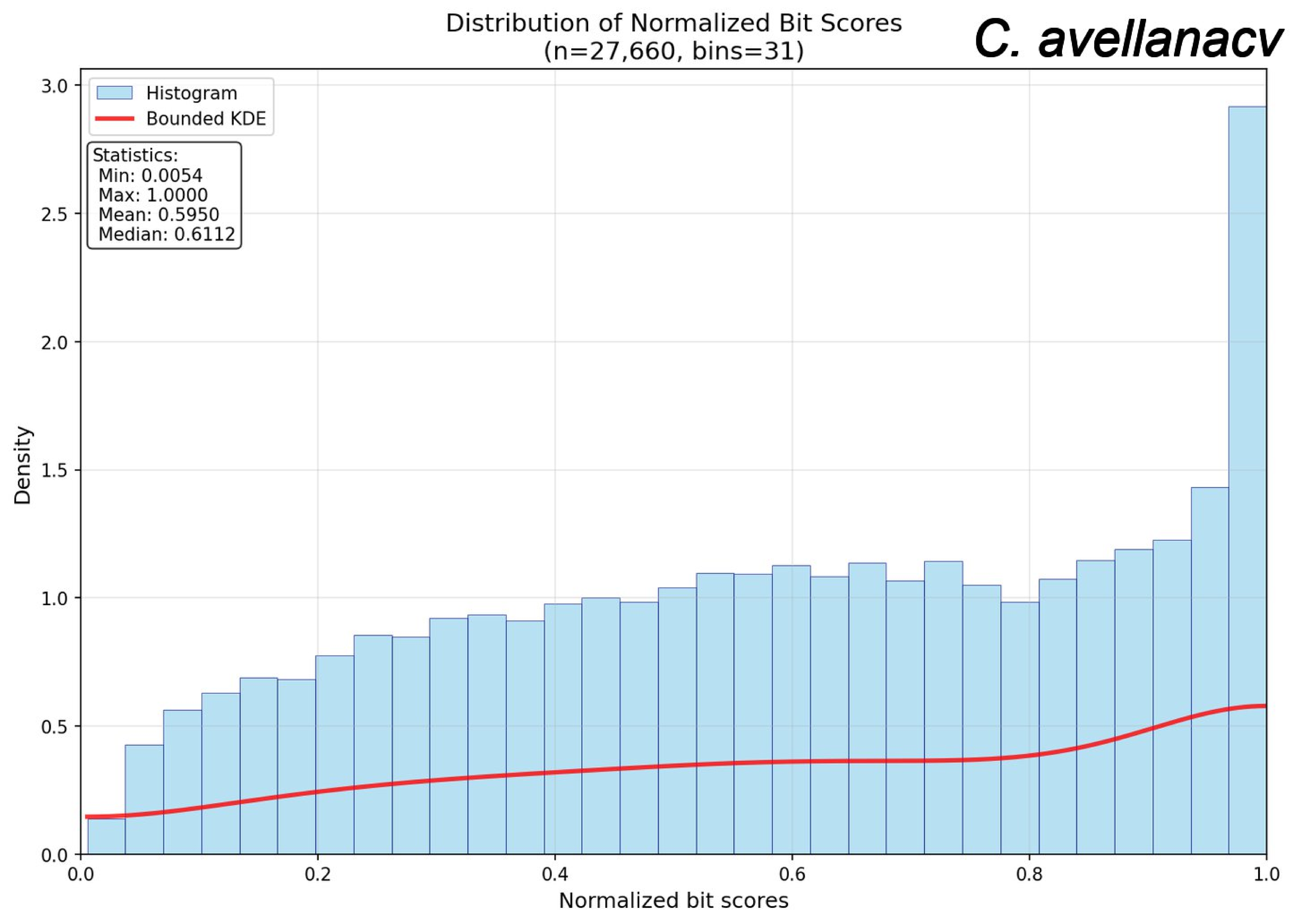
Distribution of Normalized Bit Scores
(n=27,660, bins=31)
C. avellanacv
Histogram
Bounded KDE
Statistics:
Min: 0.0054
Max: 1.0000
Mean: 0.5950
Median: 0.6112
Density
Normalized bit scores
0.0
0.5
1.0
1.5
2.0
2.5
3.0
0.0
0.2
0.4
0.6
0.8
1.0

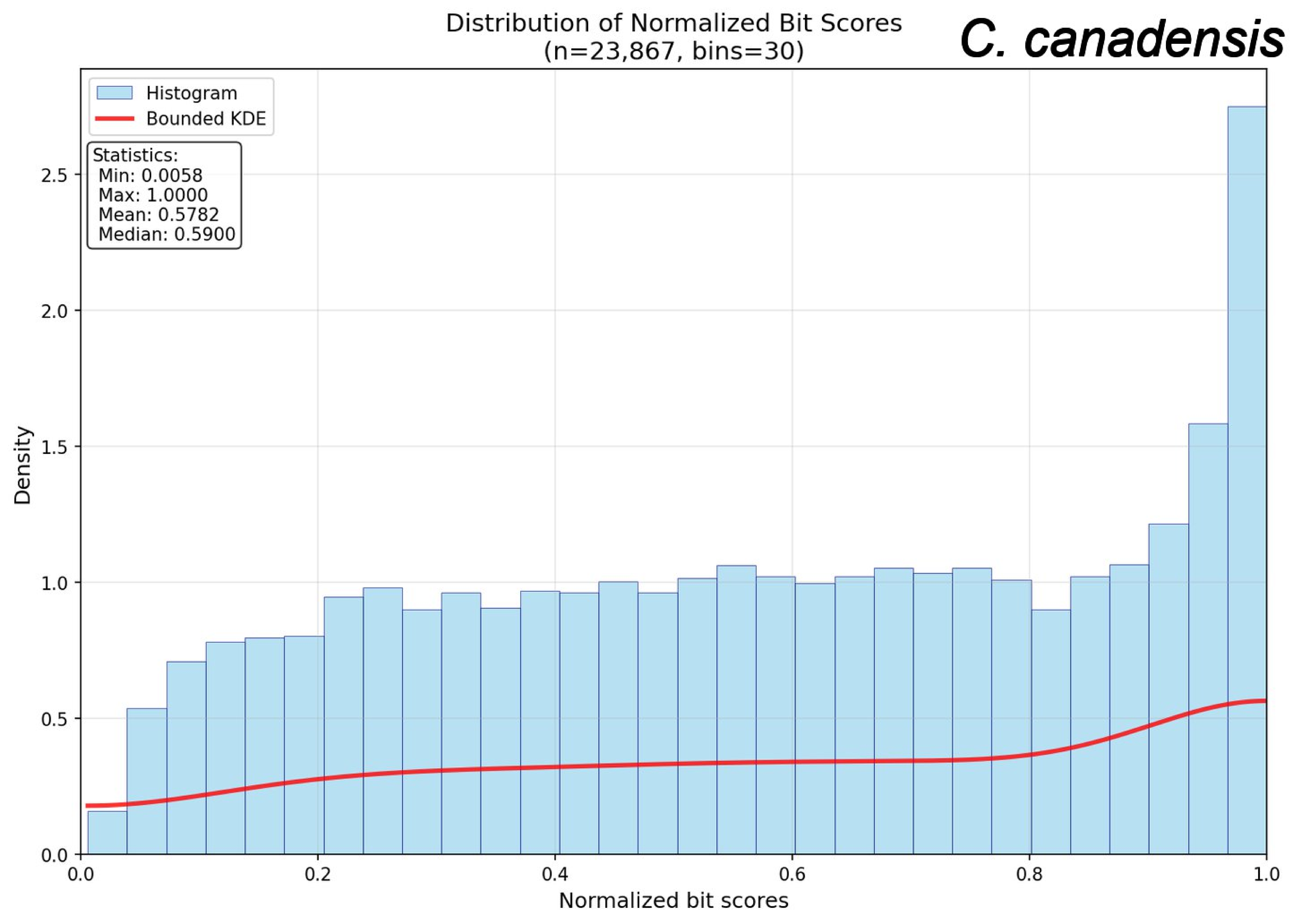
Distribution of Normalized Bit Scores
(n=23,867, bins=30)
C. canadensis
Histogram
Bounded KDE
Statistics:
Min: 0.0058
Max: 1.0000
Mean: 0.5782
Median: 0.5900
Density
0.0
0.5
1.0
1.5
2.0
2.5
0.0
0.2
0.4
0.6
0.8
1.0
Normalized bit scores

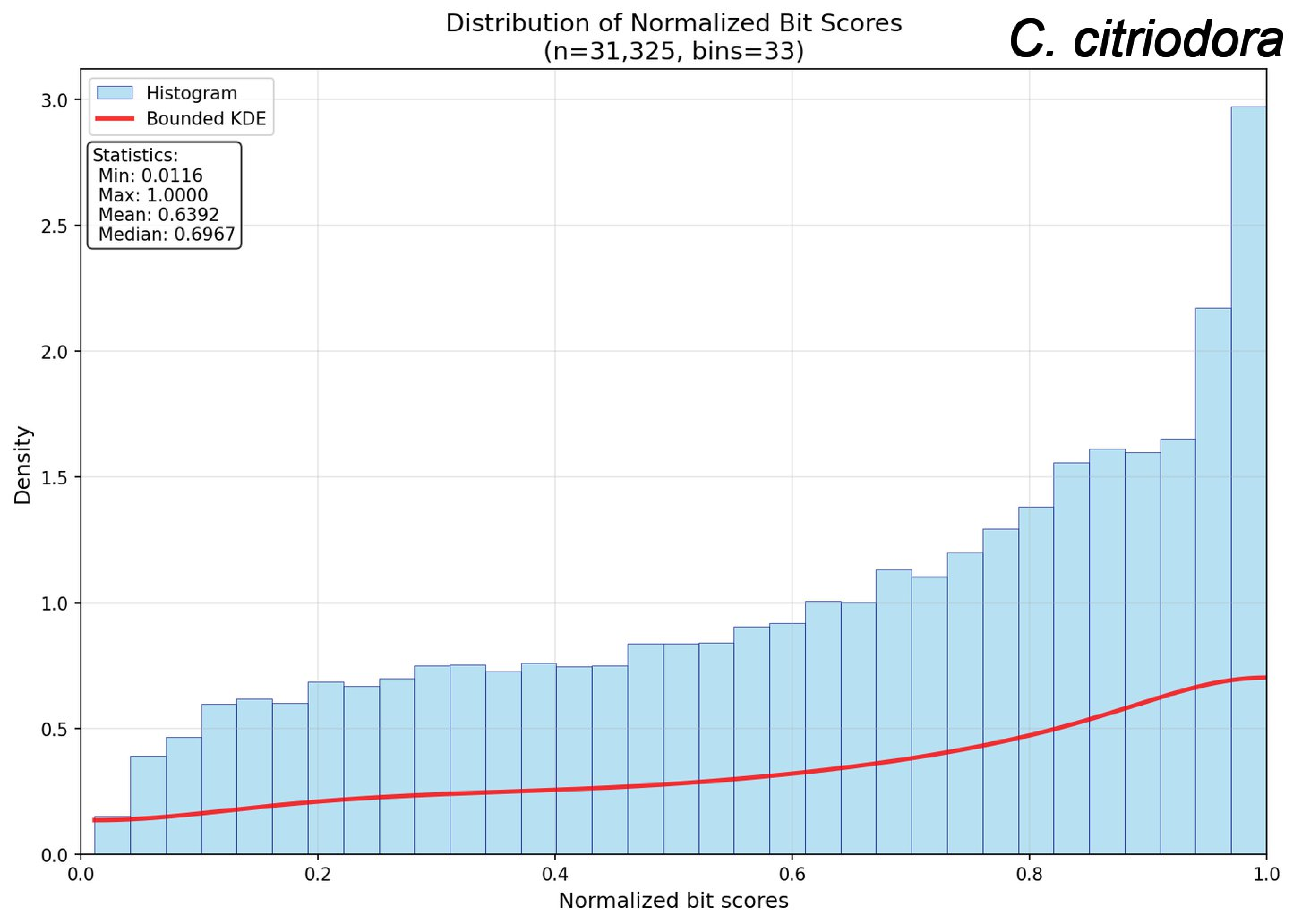

Distribution of Normalized Bit Scores
(n=31,325, bins=33)
C. citriodora
Histogram
Bounded KDE
Statistics:
Min: 0.0116
Max: 1.0000
Mean: 0.6392
Median: 0.6967
Density
Normalized bit scores
0.0
0.5
1.0
1.5
2.0
2.5
3.0
0.0
0.2
0.4
0.6
0.8
1.0

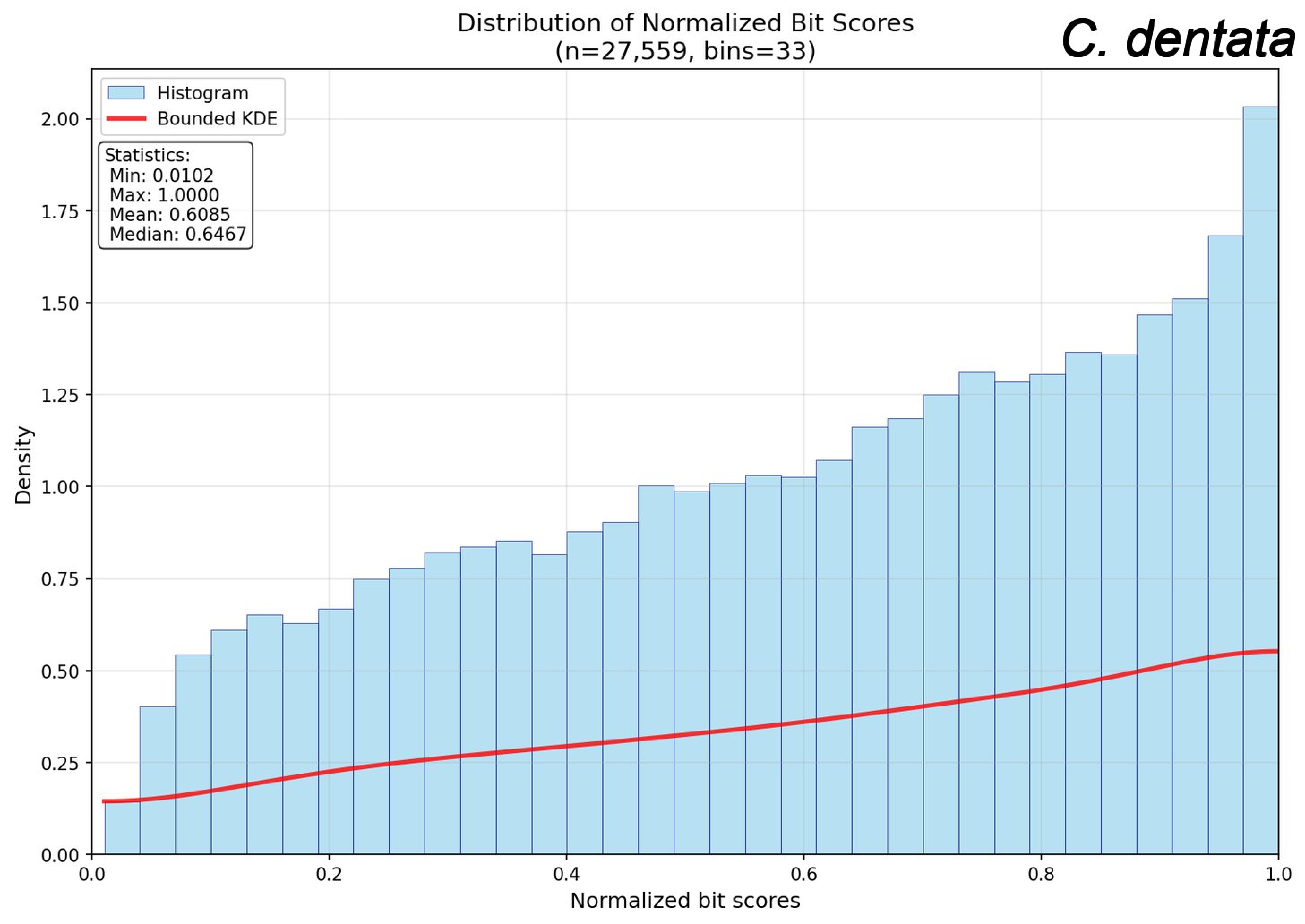
Distribution of Normalized Bit Scores
(n=27,559, bins=33)
C. dentata
Histogram
Bounded KDE
Statistics:
Min: 0.0102
Max: 1.0000
Mean: 0.6085
Median: 0.6467
Density
0.00
0.25
0.50
0.75
1.00
1.25
1.50
1.75
2.00
0.0
0.2
0.4
0.6
0.8
1.0
Normalized bit scores

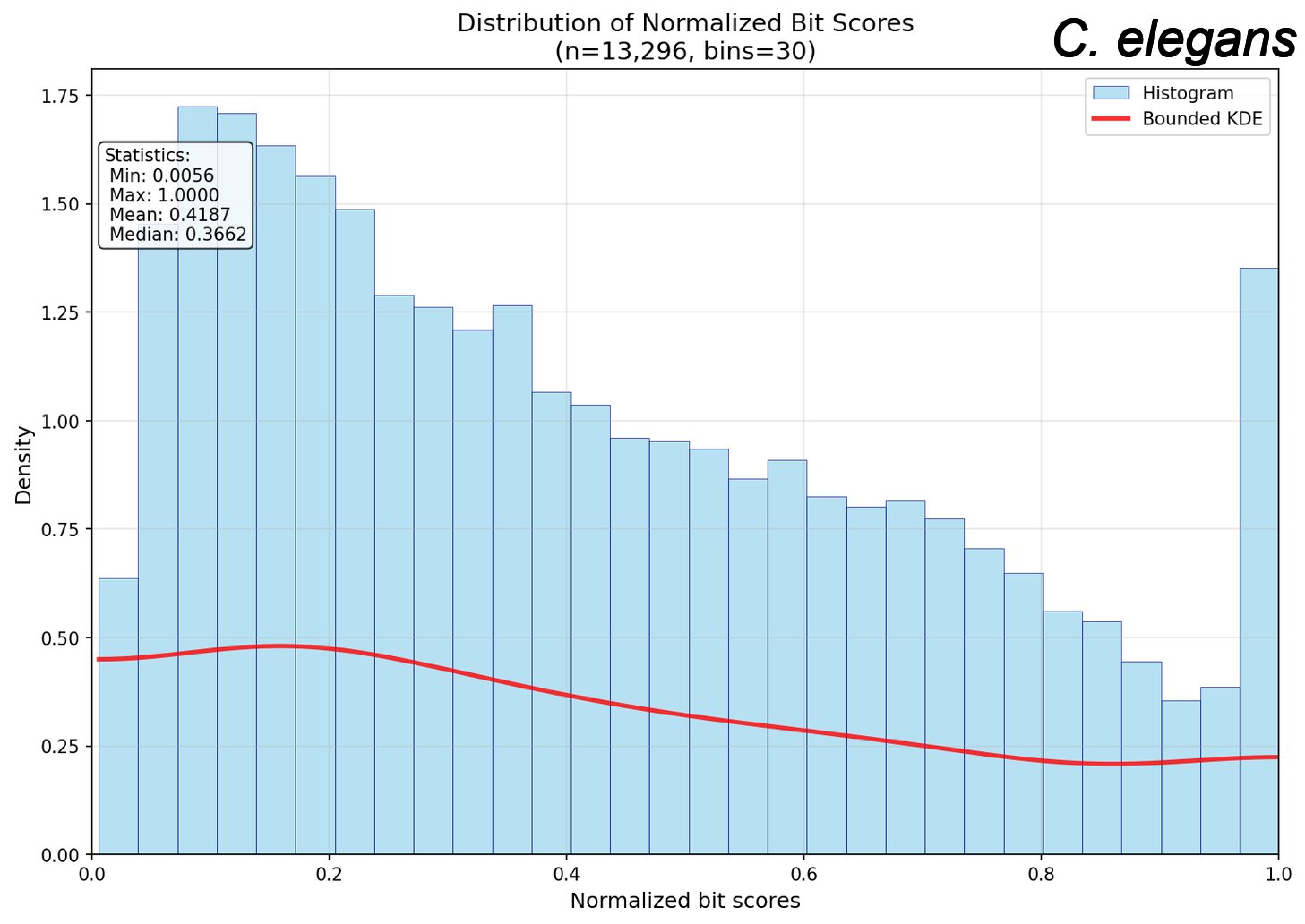
Distribution of Normalized Bit Scores
(n=13,296, bins=30)
C. elegans
Histogram
Bounded KDE
Statistics:
Min: 0.0056
Max: 1.0000
Mean: 0.4187
Median: 0.3662
Density
Normalized bit scores
0.00
0.25
0.50
0.75
1.00
1.25
1.50
1.75
0.0
0.2
0.4
0.6
0.8
1.0

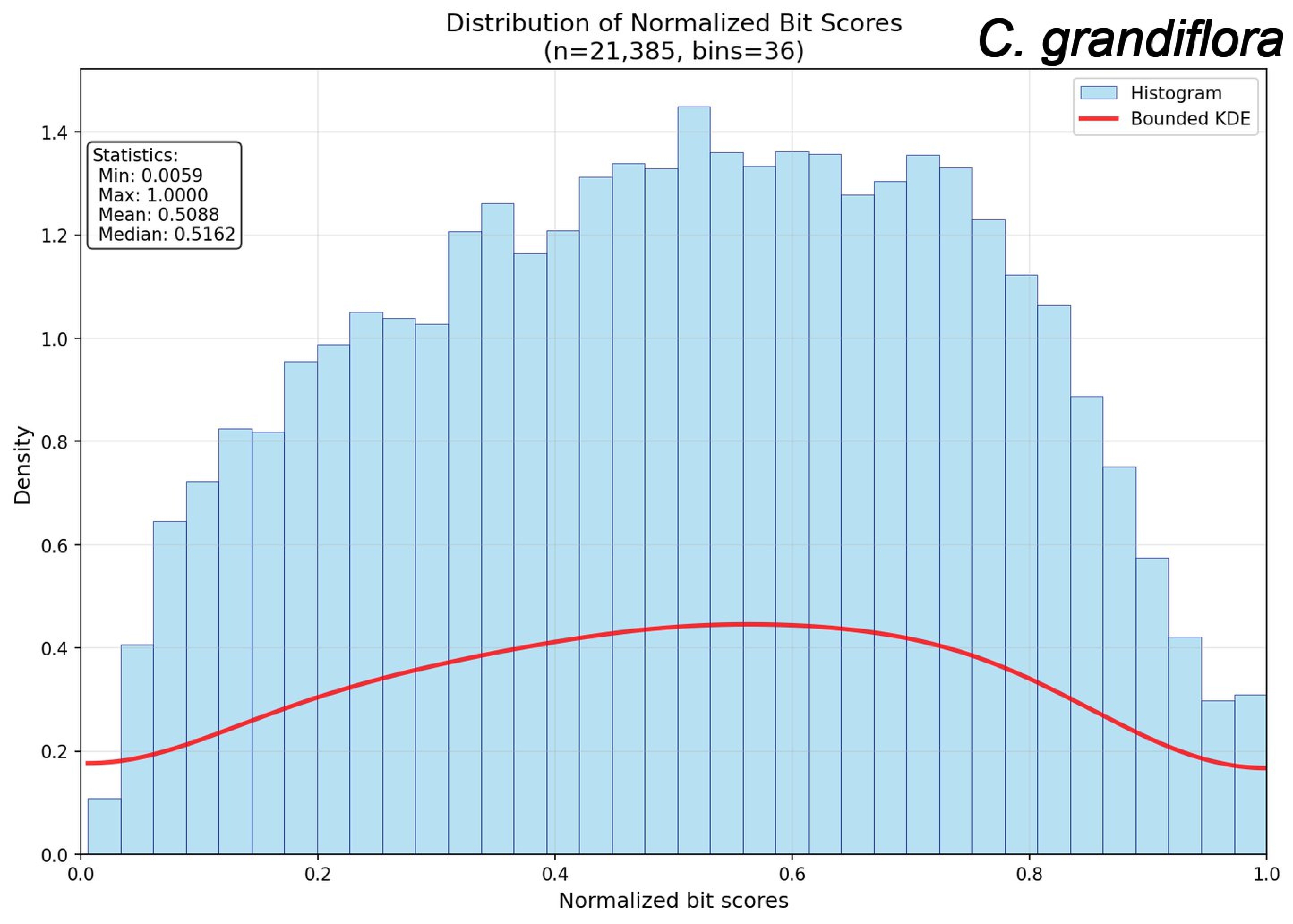
Distribution of Normalized Bit Scores
(n=21,385, bins=36)
C. grandiflora
Histogram
Bounded KDE
Statistics:
Min: 0.0059
Max: 1.0000
Mean: 0.5088
Median: 0.5162
Density
Normalized bit scores
0.0
0.2
0.4
0.6
0.8
1.0
1.2
1.4

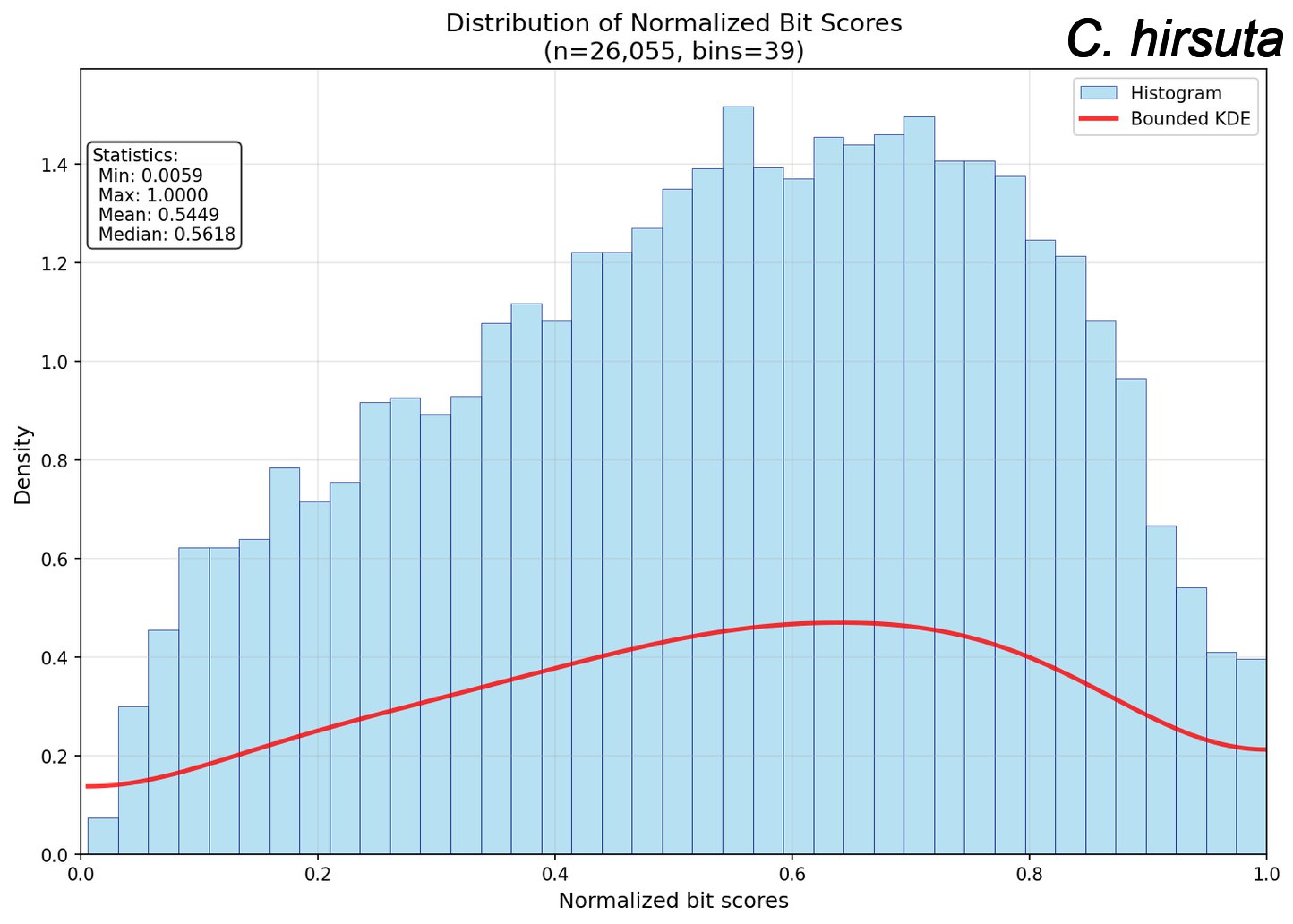
Distribution of Normalized Bit Scores
(n=26,055, bins=39)
C. hirsuta
Histogram
Bounded KDE
Statistics:
Min: 0.0059
Max: 1.0000
Mean: 0.5449
Median: 0.5618
Density
Normalized bit scores
0.0
0.2
0.4
0.6
0.8
1.0
1.2
1.4

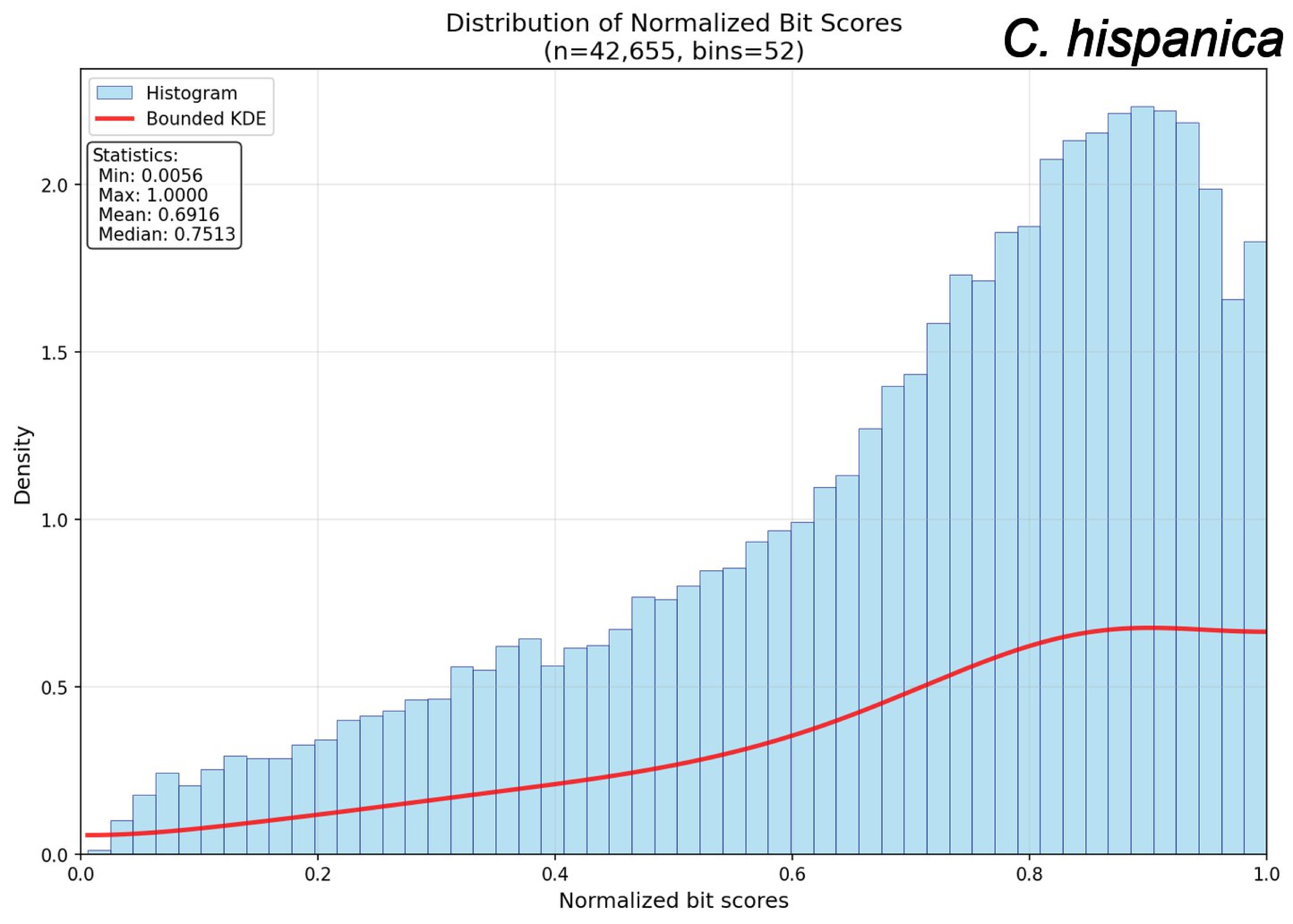

Distribution of Normalized Bit Scores
(n=42,655, bins=52)
C. hispanica
Histogram
Bounded KDE
Statistics:
Min: 0.0056
Max: 1.0000
Mean: 0.6916
Median: 0.7513
Density
Normalized bit scores
0.0
0.2
0.4
0.6
0.8
1.0
0.5
1.5
2.0

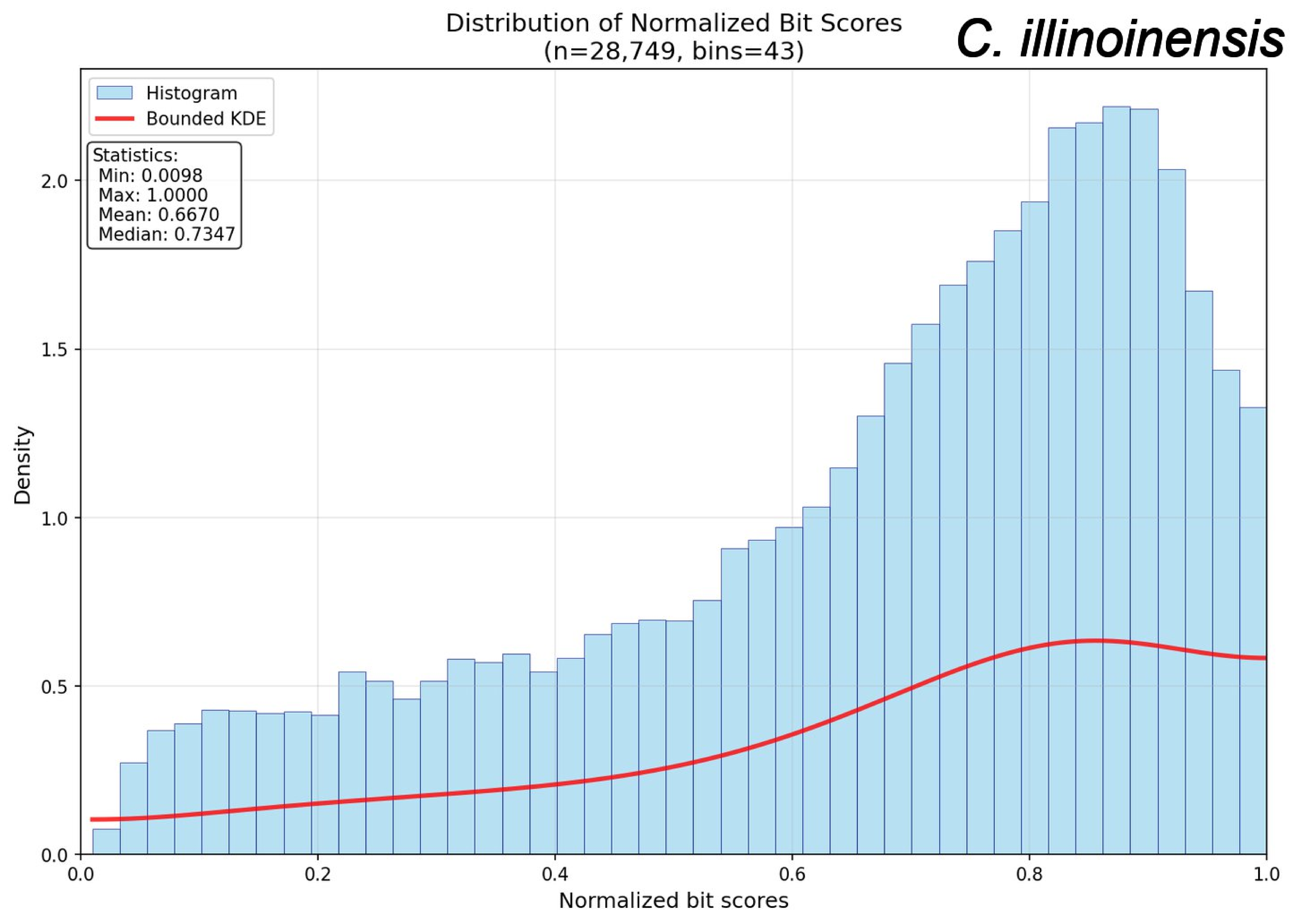

Distribution of Normalized Bit Scores
(n=28,749, bins=43)
C. illinoinensis
Histogram
Bounded KDE
Statistics:
Min: 0.0098
Max: 1.0000
Mean: 0.6670
Median: 0.7347
Density
Normalized bit scores
0.0
0.5
1.0
1.5
2.0
0.2
0.4
0.6
0.8

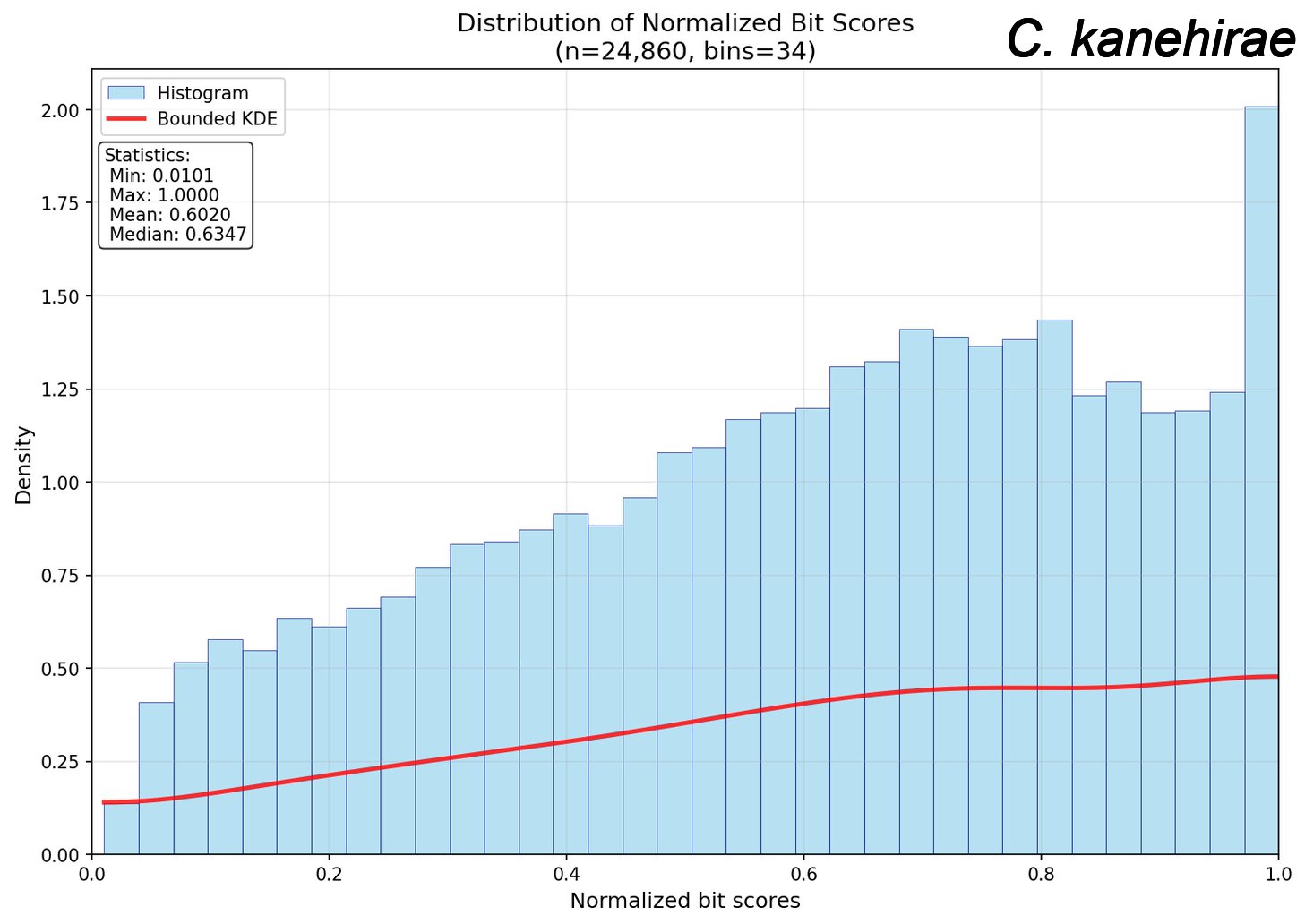

Distribution of Normalized Bit Scores
(n=24,860, bins=34)
C. kanehirae
Histogram
Bounded KDE
Statistics:
Min: 0.0101
Max: 1.0000
Mean: 0.6020
Median: 0.6347
Density
Normalized bit scores
0.00
0.25
0.50
0.75
1.00
1.25
1.50
1.75
2.00
0.0
0.2
0.4
0.6
0.8
1.0

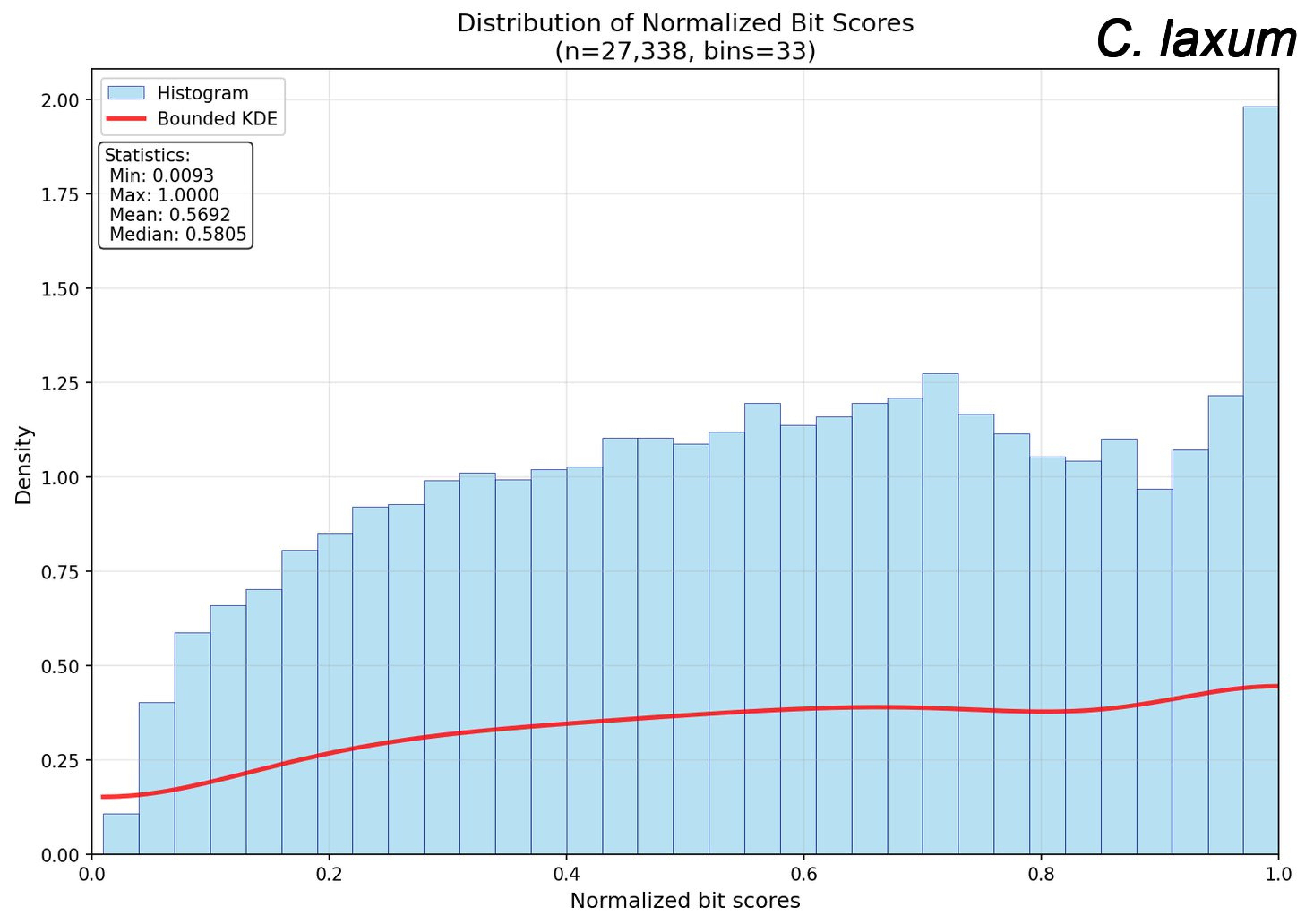

Distribution of Normalized Bit Scores
(n=27,338, bins=33)
C. laxum
Histogram
Bounded KDE
Statistics:
Min: 0.0093
Max: 1.0000
Mean: 0.5692
Median: 0.5805
Density
Normalized bit scores
0.00
0.25
0.50
0.75
1.00
1.25
1.50
1.75
2.00
0.0
0.2
0.4
0.6
0.8
1.0

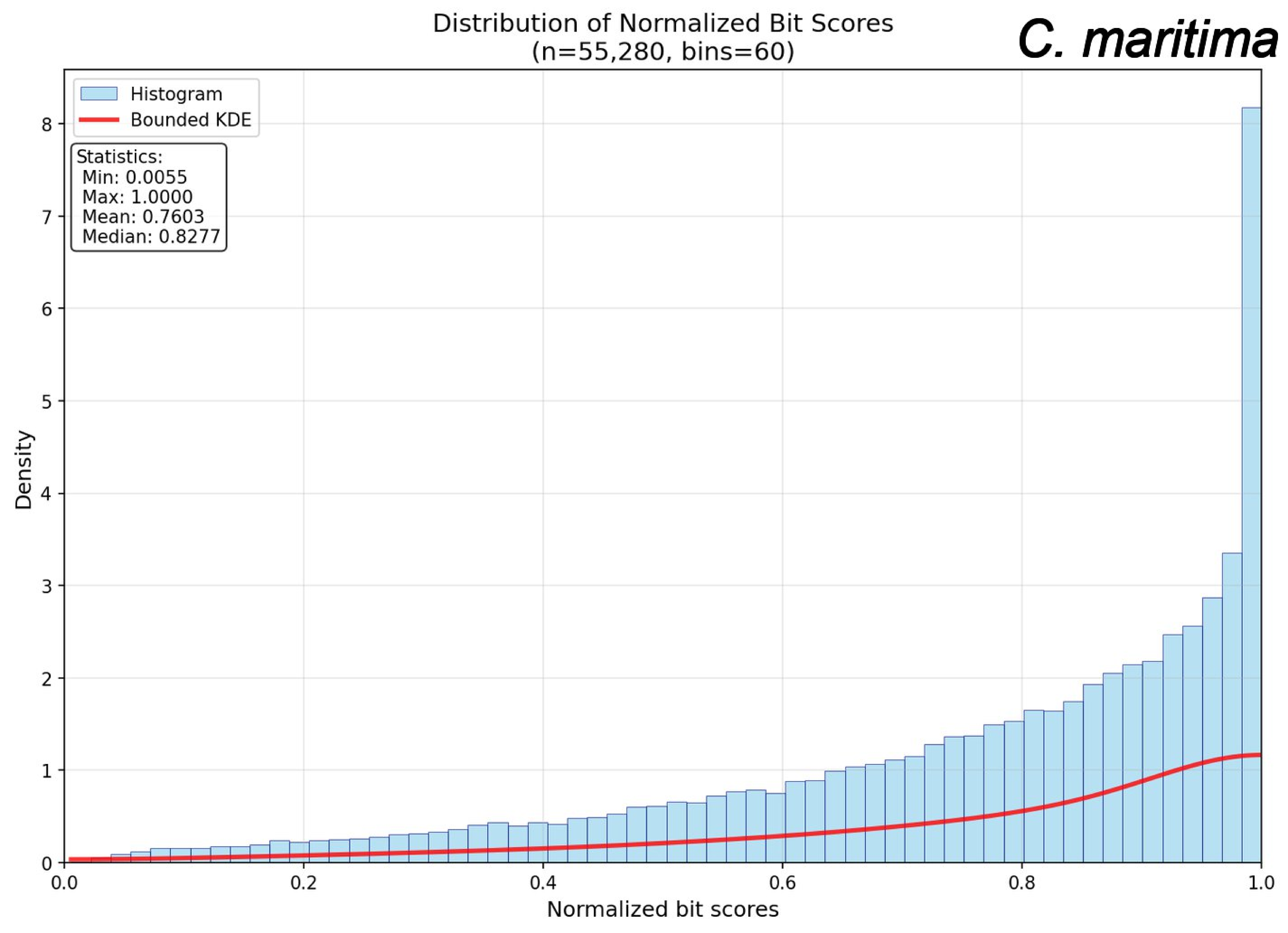
Distribution of Normalized Bit Scores
(n=55,280, bins=60)
C. maritima
Histogram
Bounded KDE
Statistics:
Min: 0.0055
Max: 1.0000
Mean: 0.7603
Median: 0.8277
Density
Normalized bit scores
0
1
2
3
4
5
6
7
8
0.0
0.2
0.4
0.6
0.8
1.0

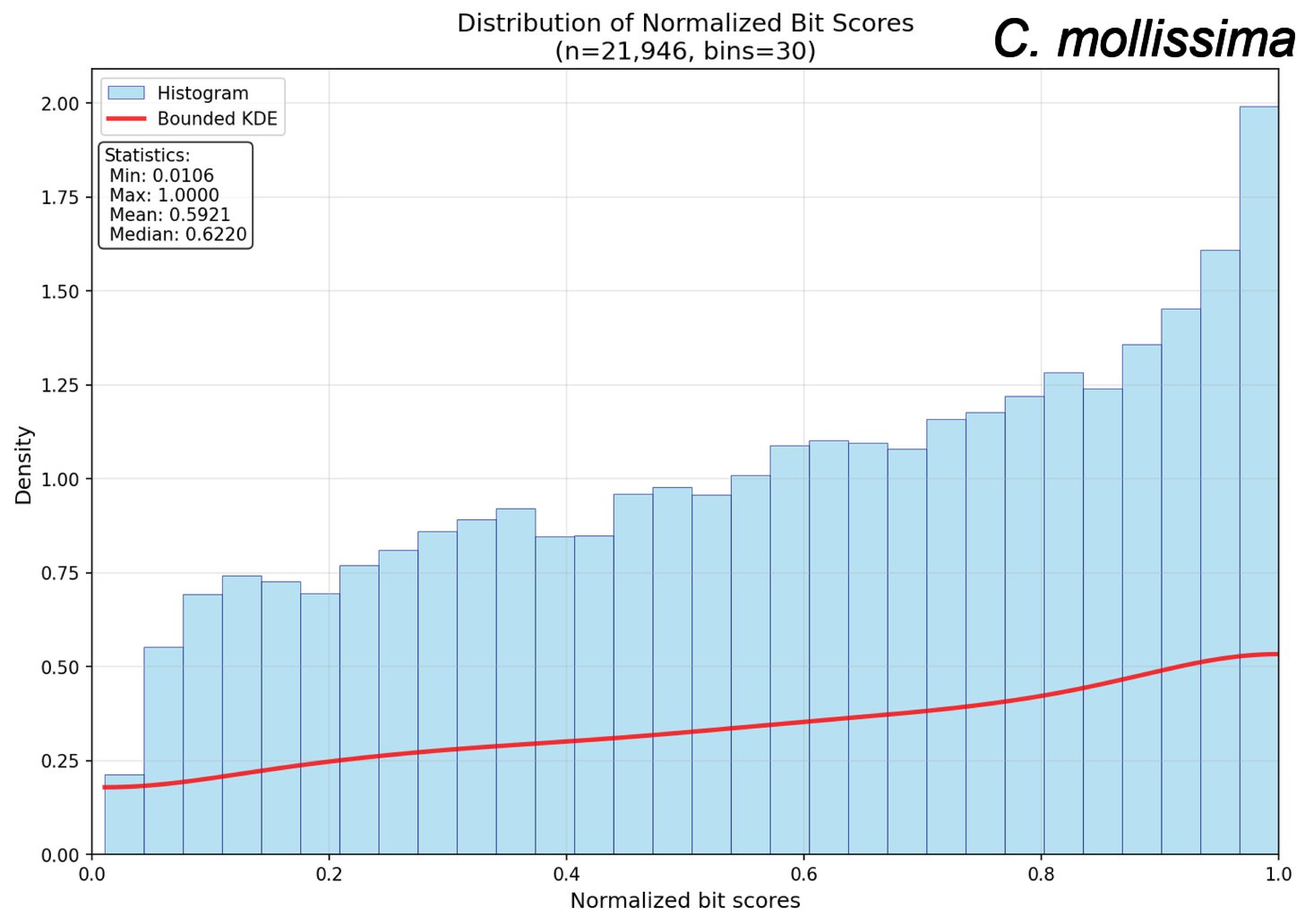

Distribution of Normalized Bit Scores
(n=21,946, bins=30)
C. mollissima
Histogram
Bounded KDE
Statistics:
Min: 0.0106
Max: 1.0000
Mean: 0.5921
Median: 0.6220
Density
Normalized bit scores
0.00
0.25
0.50
0.75
1.00
1.25
1.50
1.75
2.00
0.0
0.2
0.4
0.6
0.8
1.0

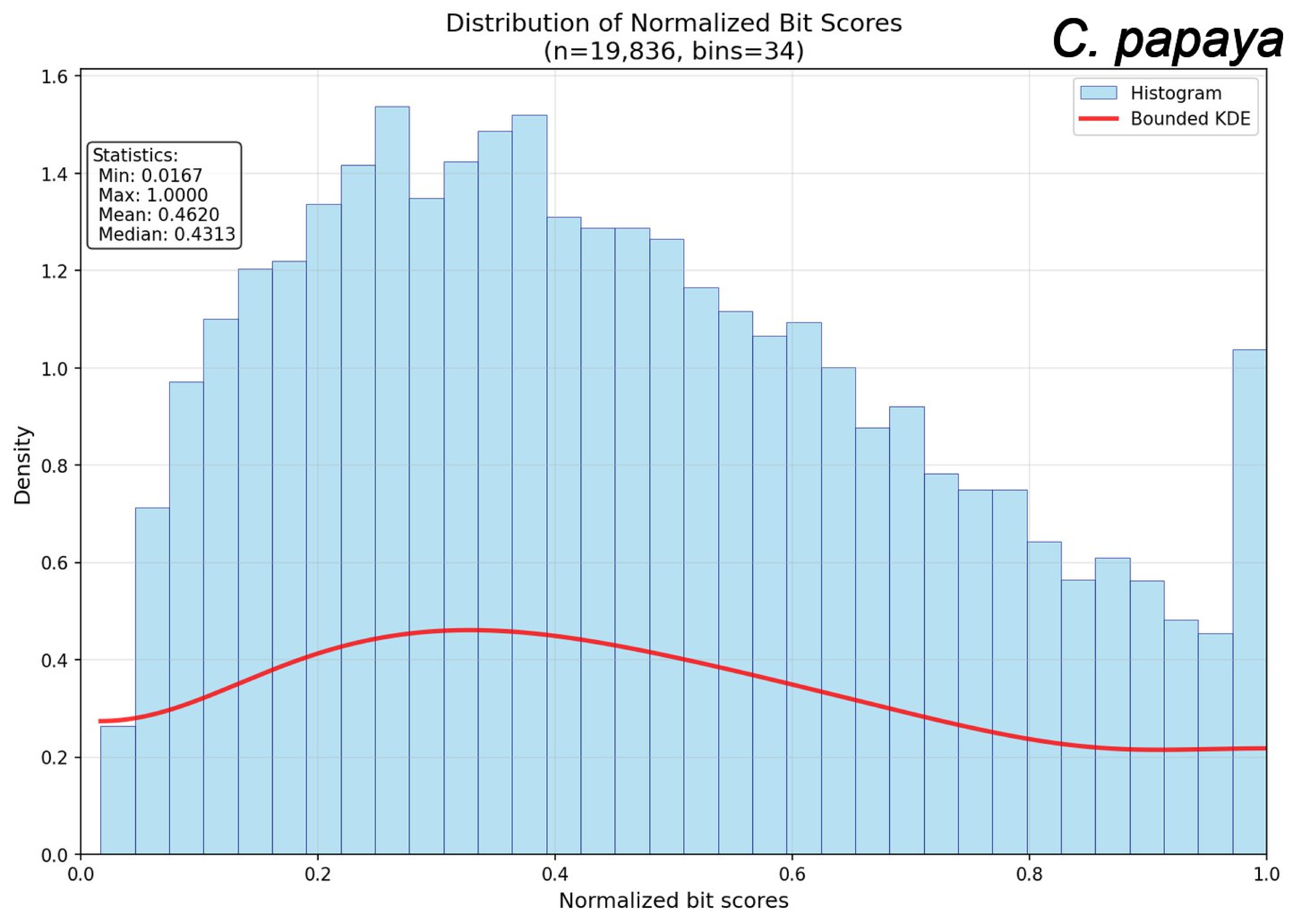
Distribution of Normalized Bit Scores
(n=19,836, bins=34)
C. papaya
Histogram
Bounded KDE
Statistics:
Min: 0.0167
Max: 1.0000
Mean: 0.4620
Median: 0.4313
Density
Normalized bit scores
0.0
0.2
0.4
0.6
0.8
1.0
1.2
1.4
1.6

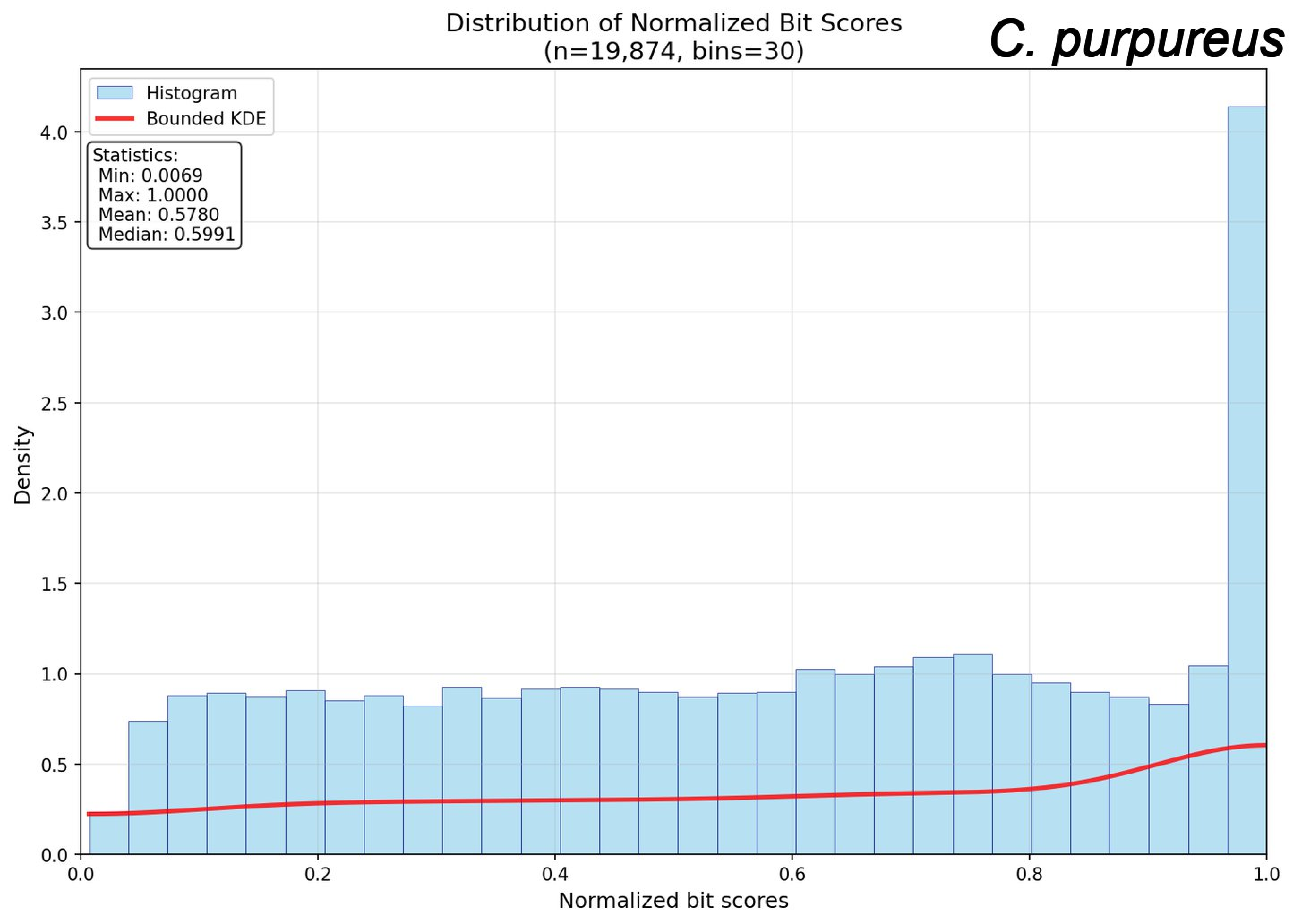
Distribution of Normalized Bit Scores
(n=19,874, bins=30)
C. purpureus
Histogram
Bounded KDE
Statistics:
Min: 0.0069
Max: 1.0000
Mean: 0.5780
Median: 0.5991
Density
Normalized bit scores
0.0
0.5
1.0
1.5
2.0
2.5
3.0
3.5
4.0
0.0
0.2
0.4
0.6
0.8
1.0

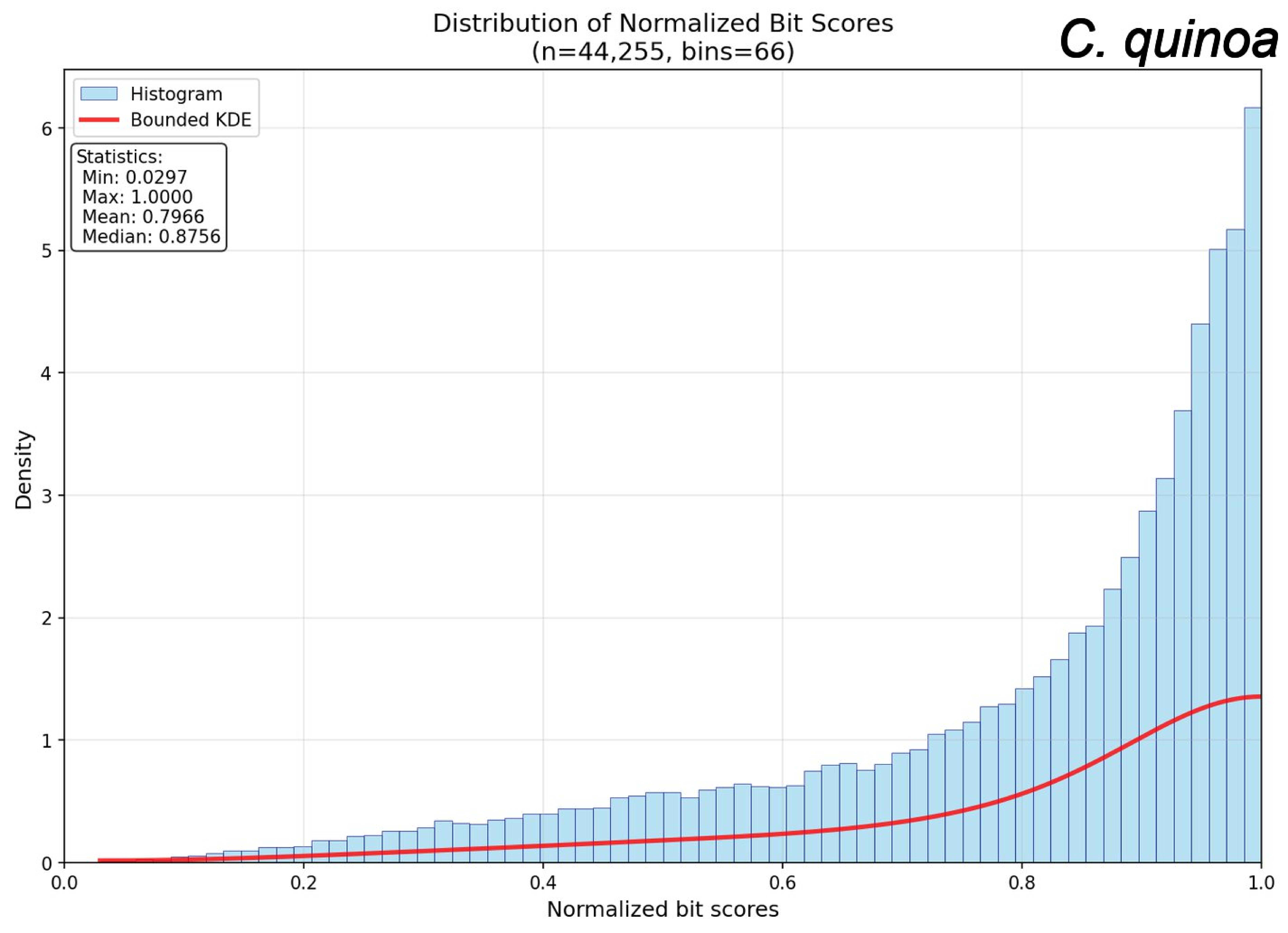
Distribution of Normalized Bit Scores
(n=44,255, bins=66)
C. quinoa
Histogram
Bounded KDE
Statistics:
Min: 0.0297
Max: 1.0000
Mean: 0.7966
Median: 0.8756
Density
Normalized bit scores
0
1
2
3
4
5
6
0.0
0.2
0.4
0.6
0.8
1.0

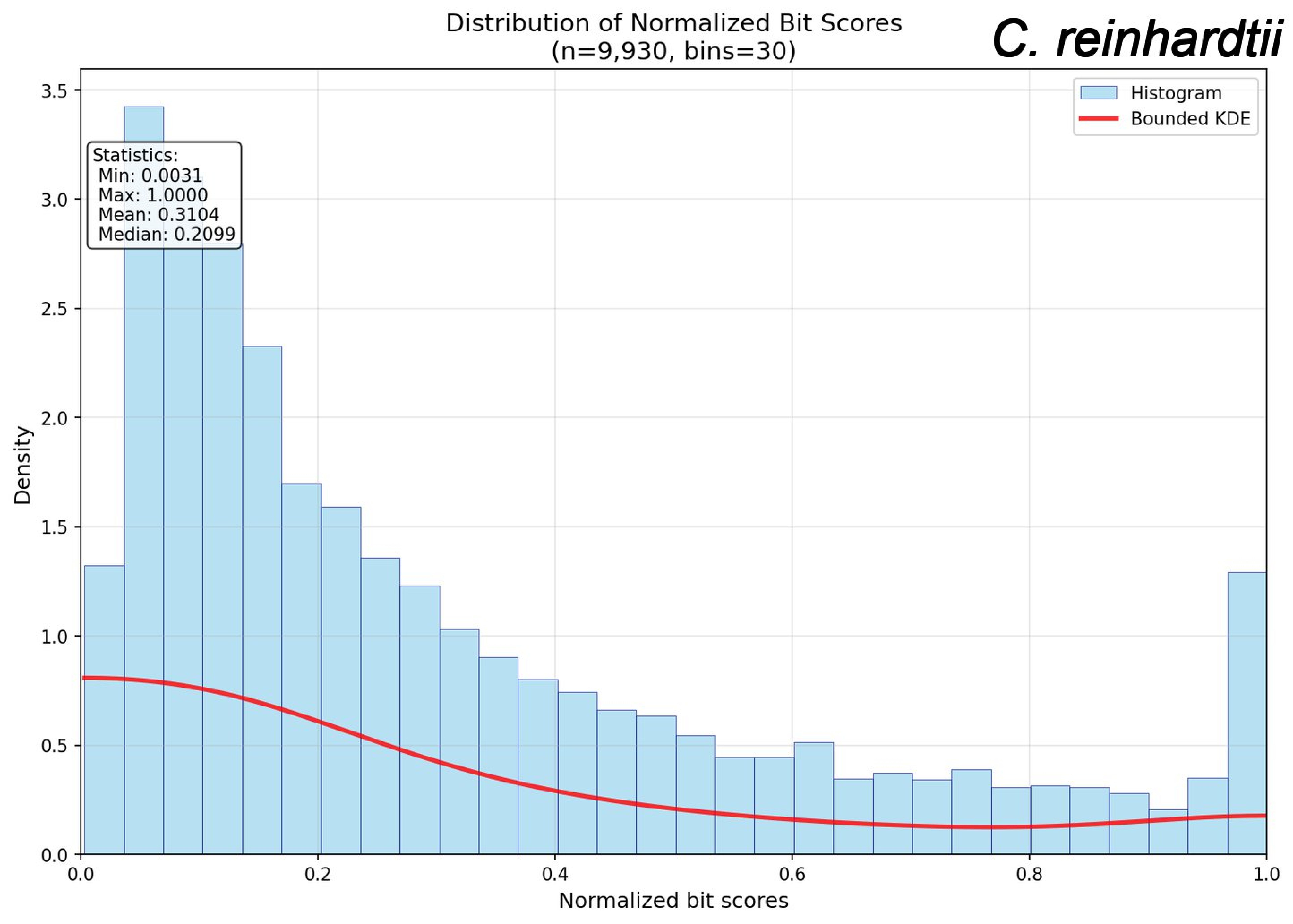

Distribution of Normalized Bit Scores
(n=9,930, bins=30)
C. reinhardtii
Histogram
Bounded KDE
Statistics:
Min: 0.0031
Max: 1.0000
Mean: 0.3104
Median: 0.2099
Density
Normalized bit scores
0.0
0.5
1.0
1.5
2.0
2.5
3.0
3.5
0.0
0.2
0.4
0.6
0.8
1.0

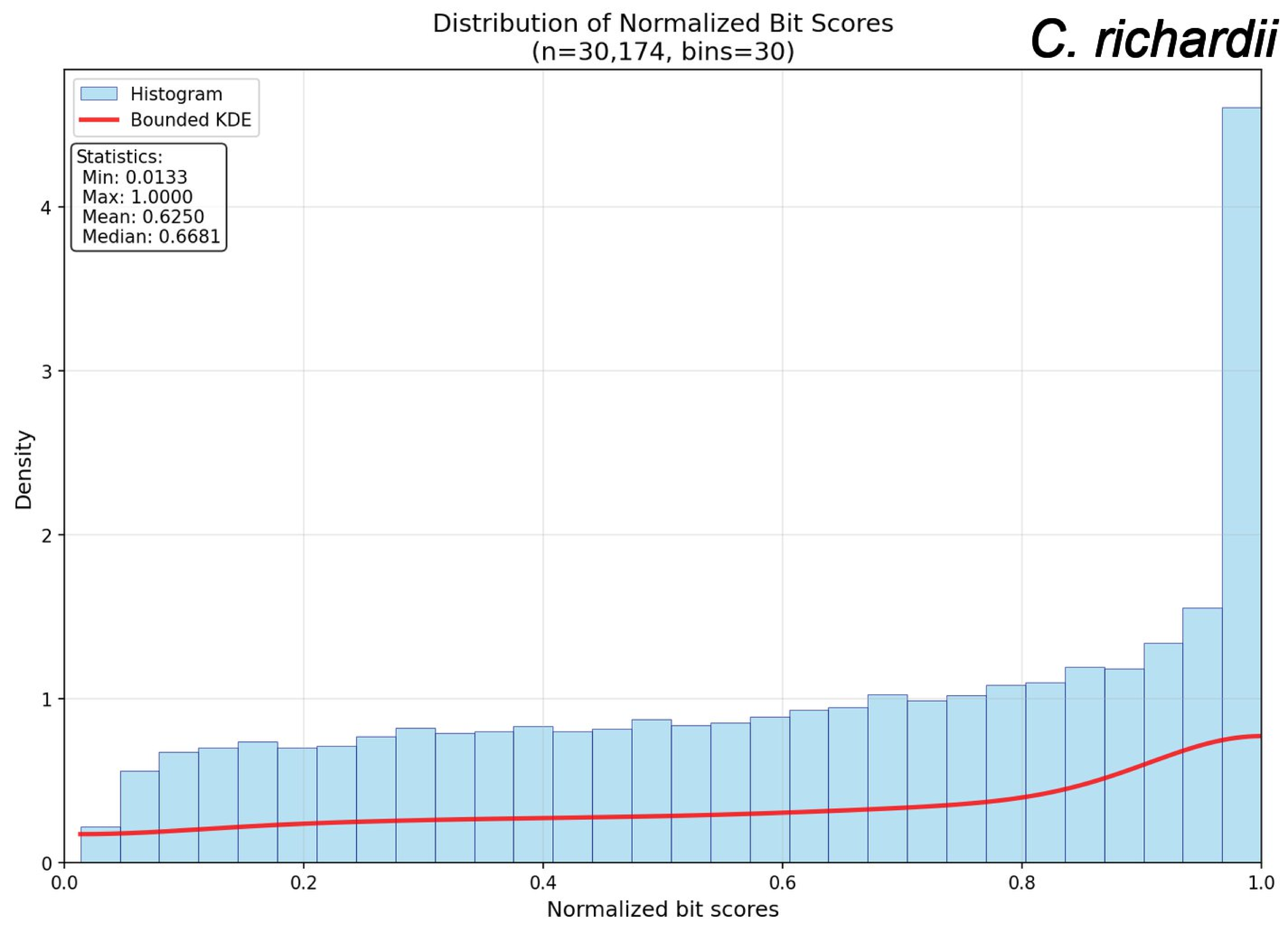
Distribution of Normalized Bit Scores
(n=30,174, bins=30)
C. richardii
Histogram
Bounded KDE
Statistics:
Min: 0.0133
Max: 1.0000
Mean: 0.6250
Median: 0.6681
Density
Normalized bit scores
0
1
2
3
4
0.0
0.2
0.4
0.6
0.8
1.0

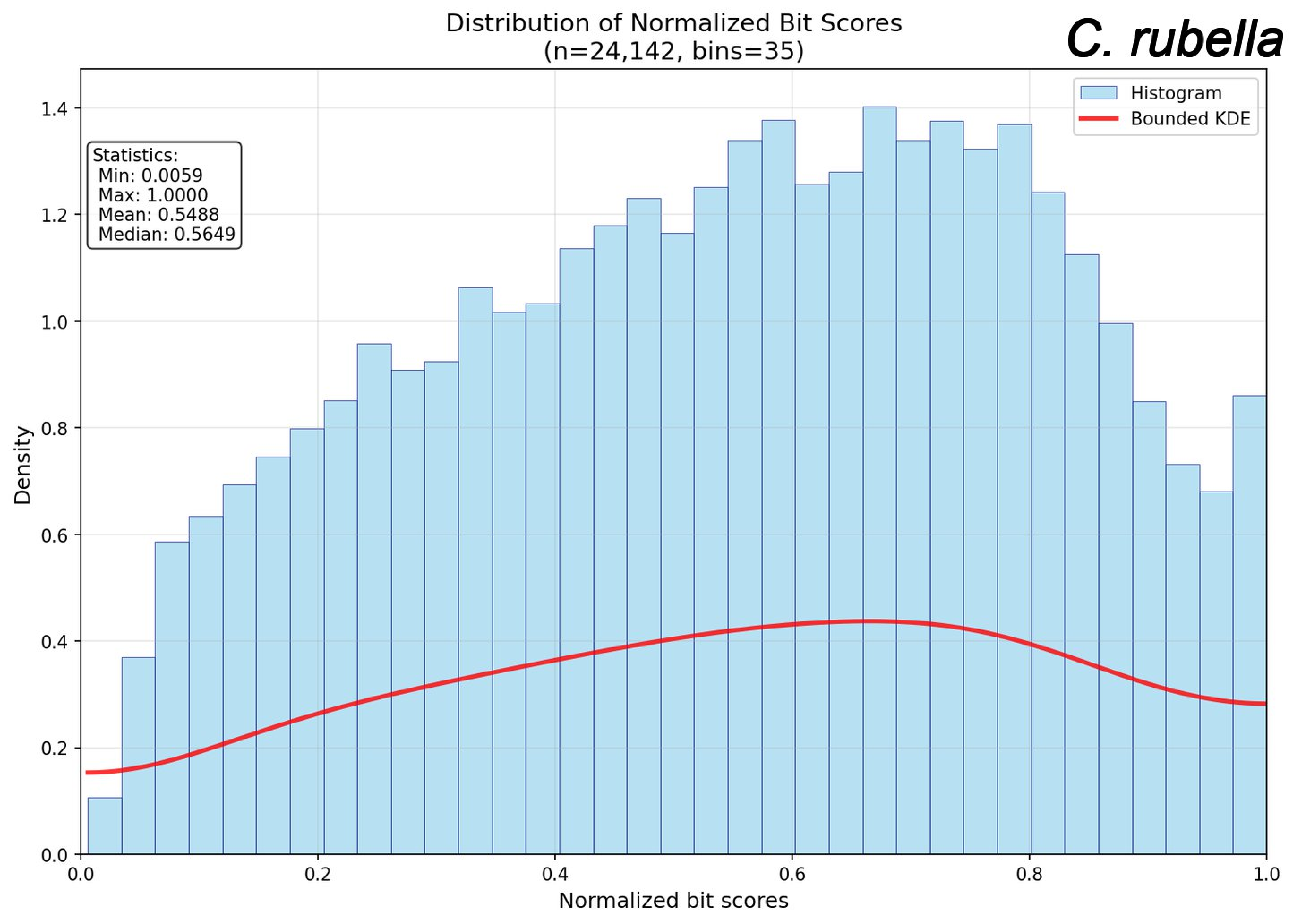
Distribution of Normalized Bit Scores
(n=24,142, bins=35)
C. rubella
Histogram
Bounded KDE
Statistics:
Min: 0.0059
Max: 1.0000
Mean: 0.5488
Median: 0.5649
Density
Normalized bit scores
0.0
0.2
0.4
0.6
0.8
1.0
1.2
1.4

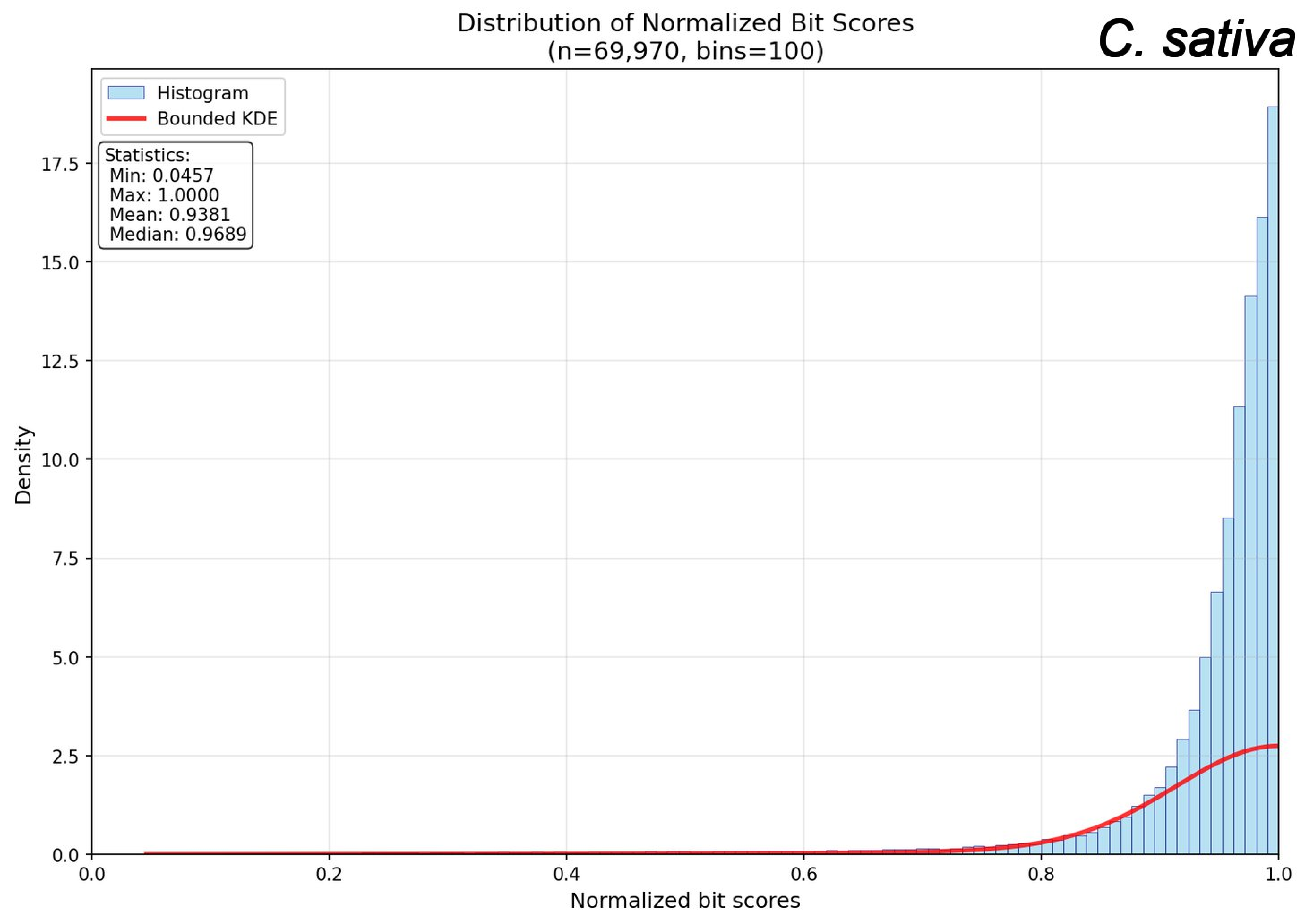

Distribution of Normalized Bit Scores
(n=69,970, bins=100)
C. sativa
Histogram
Bounded KDE
Statistics:
Min: 0.0457
Max: 1.0000
Mean: 0.9381
Median: 0.9689
Density
Normalized bit scores
0.0
2.5
5.0
7.5
10.0
12.5
15.0
17.5
0.0
0.2
0.4
0.6
0.8
1.0

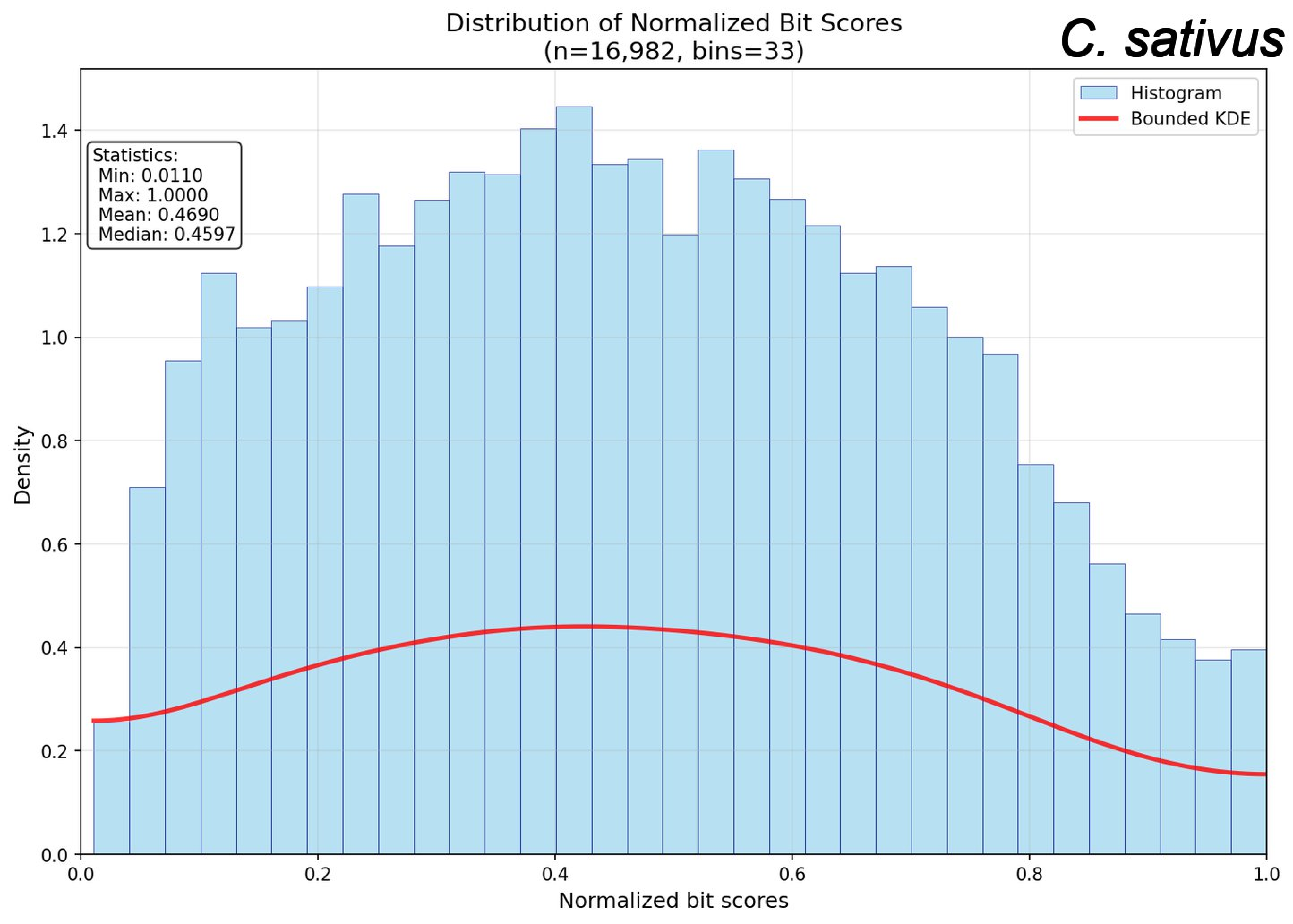

Distribution of Normalized Bit Scores
(n=16,982, bins=33)
C. sativus
Histogram
Bounded KDE
Statistics:
Min: 0.0110
Max: 1.0000
Mean: 0.4690
Median: 0.4597
Density
Normalized bit scores
0.0
0.2
0.4
0.6
0.8
1.0
1.2
1.4

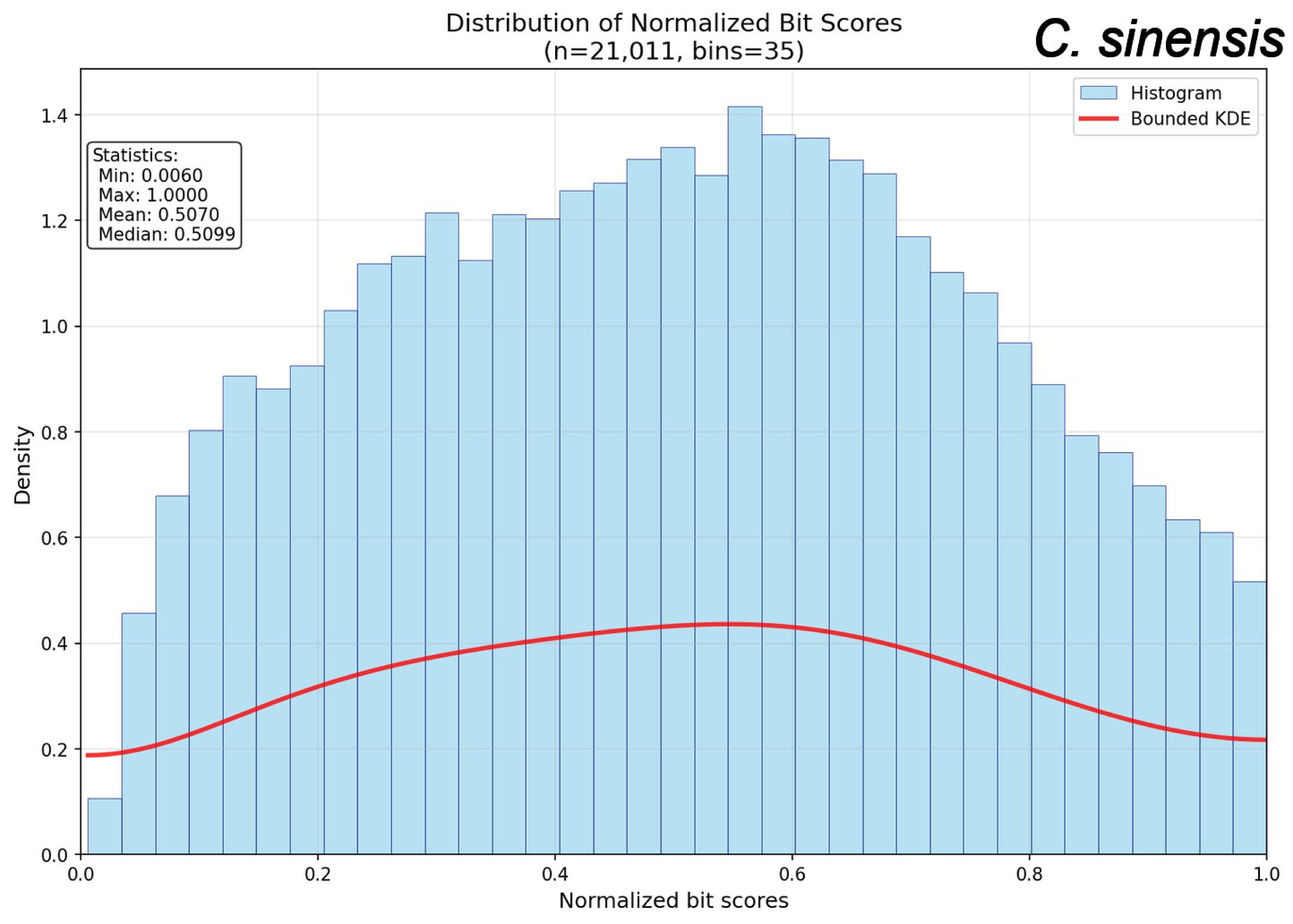
Distribution of Normalized Bit Scores
(n=21,011, bins=35)
C. sinensis
Histogram
Bounded KDE
Statistics:
Min: 0.0060
Max: 1.0000
Mean: 0.5070
Median: 0.5099
Density
Normalized bit scores
0.0
0.2
0.4
0.6
0.8
1.0
1.2
1.4

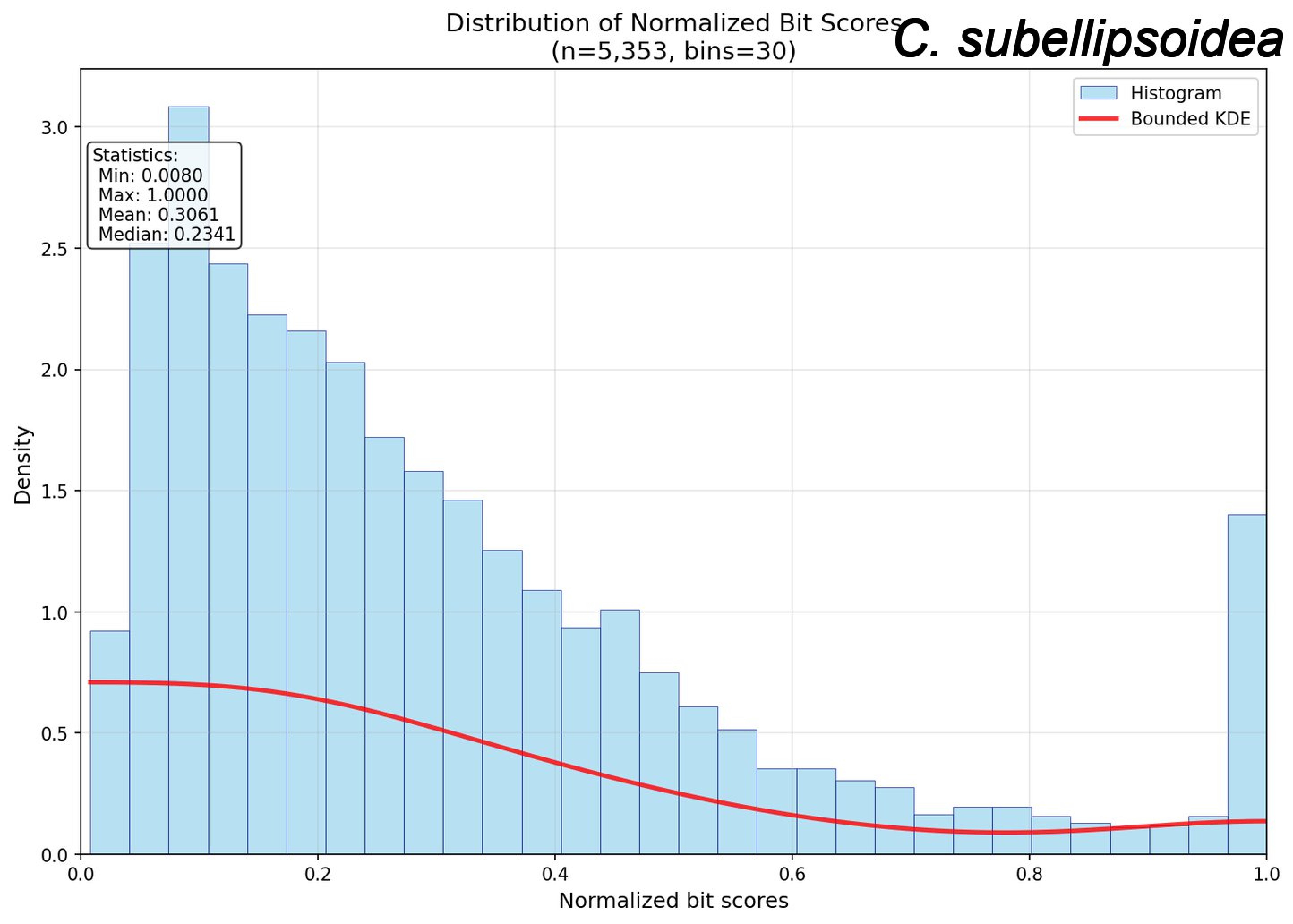
Distribution of Normalized Bit Scores
(n=5,353, bins=30)
C. subellipsoidea
Histogram
Bounded KDE
Statistics:
Min: 0.0080
Max: 1.0000
Mean: 0.3061
Median: 0.2341
Density
Normalized bit scores
0.0
0.5
1.0
1.5
2.0
2.5
3.0
0.0
0.2
0.4
0.6
0.8
1.0

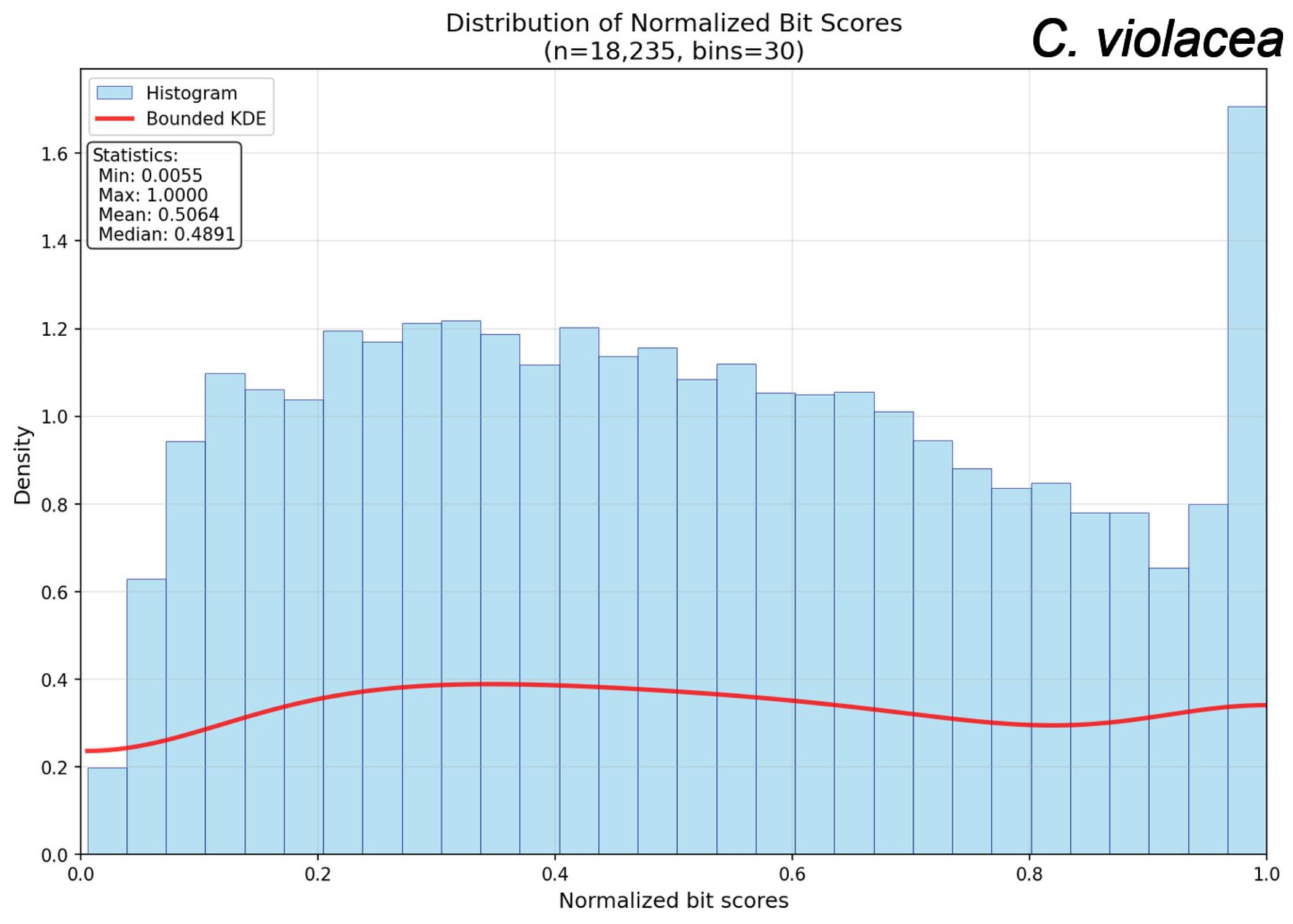

Distribution of Normalized Bit Scores
(n=18,235, bins=30)
C. violacea
Histogram
Bounded KDE
Statistics:
Min: 0.0055
Max: 1.0000
Mean: 0.5064
Median: 0.4891
Density
Normalized bit scores
0.0
0.2
0.4
0.6
0.8
1.0
1.2
1.4
1.6

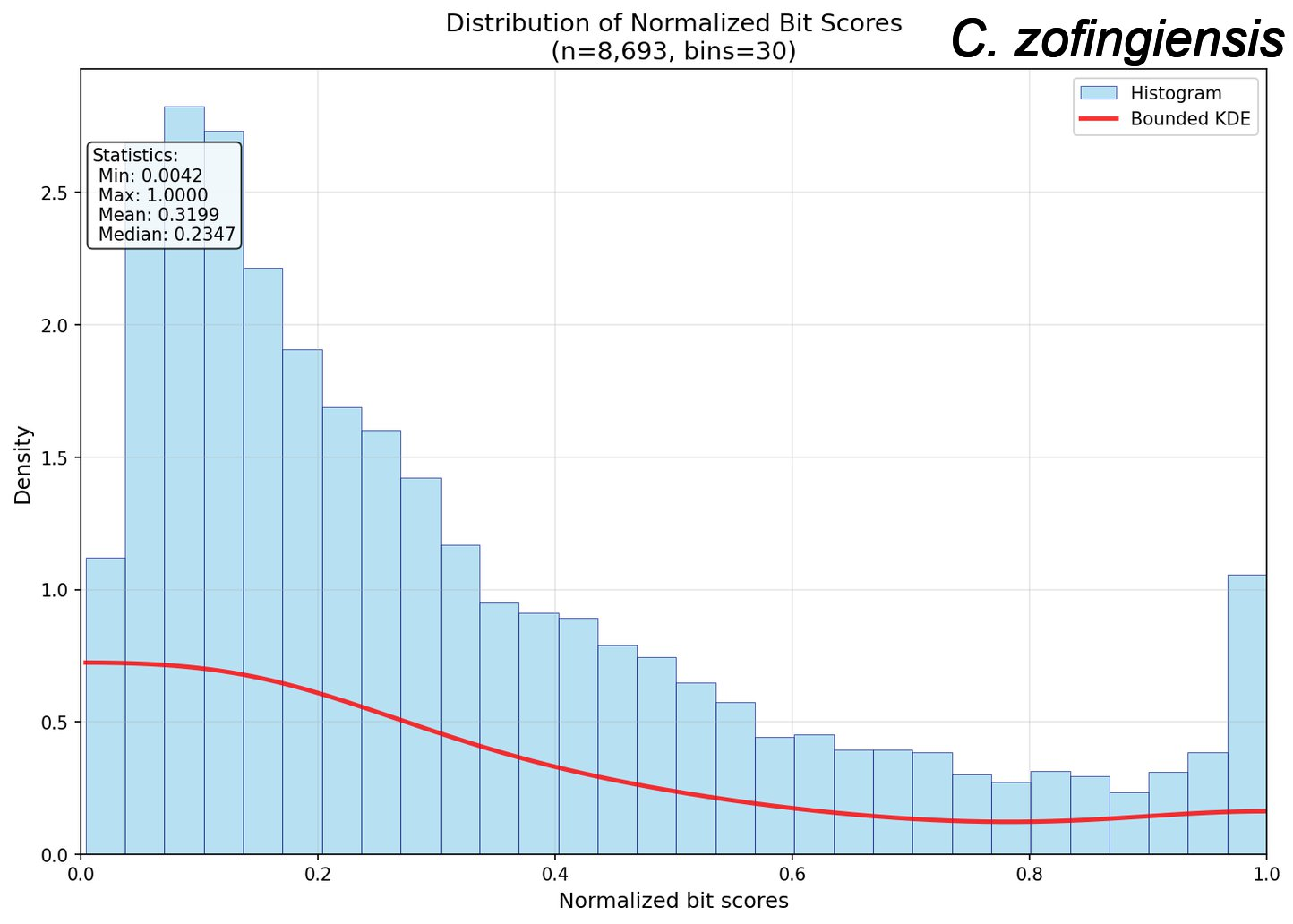
Distribution of Normalized Bit Scores
(n=8,693, bins=30)
C. zofingiensis
Histogram
Bounded KDE
Statistics:
Min: 0.0042
Max: 1.0000
Mean: 0.3199
Median: 0.2347
Density
Normalized bit scores
0.0
0.5
1.0
1.5
2.0
2.5
0.0
0.2
0.4
0.6
0.8
1.0

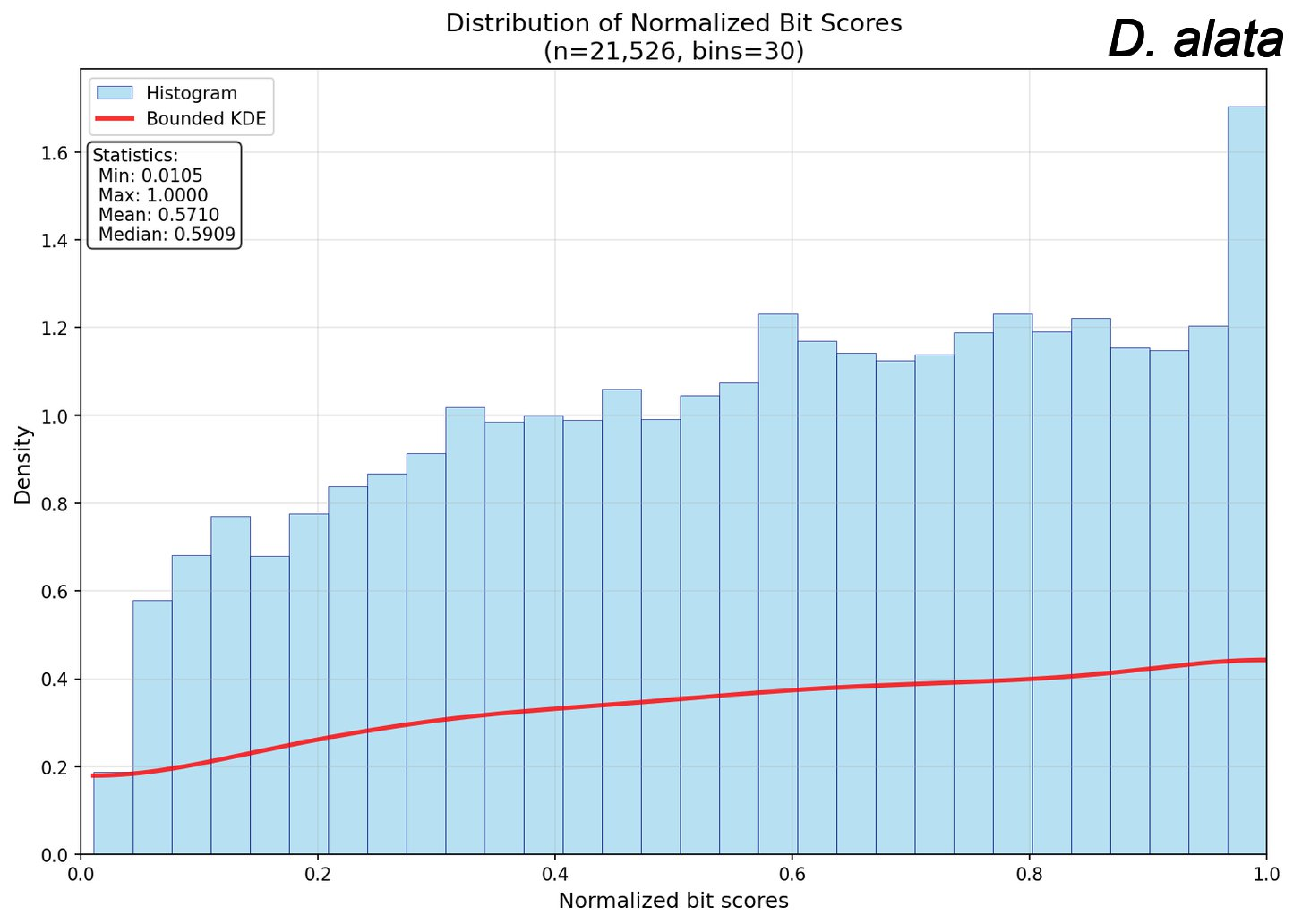

Distribution of Normalized Bit Scores
(n=21,526, bins=30)
D. alata
Histogram
Bounded KDE
Statistics:
Min: 0.0105
Max: 1.0000
Mean: 0.5710
Median: 0.5909
Density
Normalized bit scores
0.0
0.2
0.4
0.6
0.8
1.0
1.2
1.4
1.6

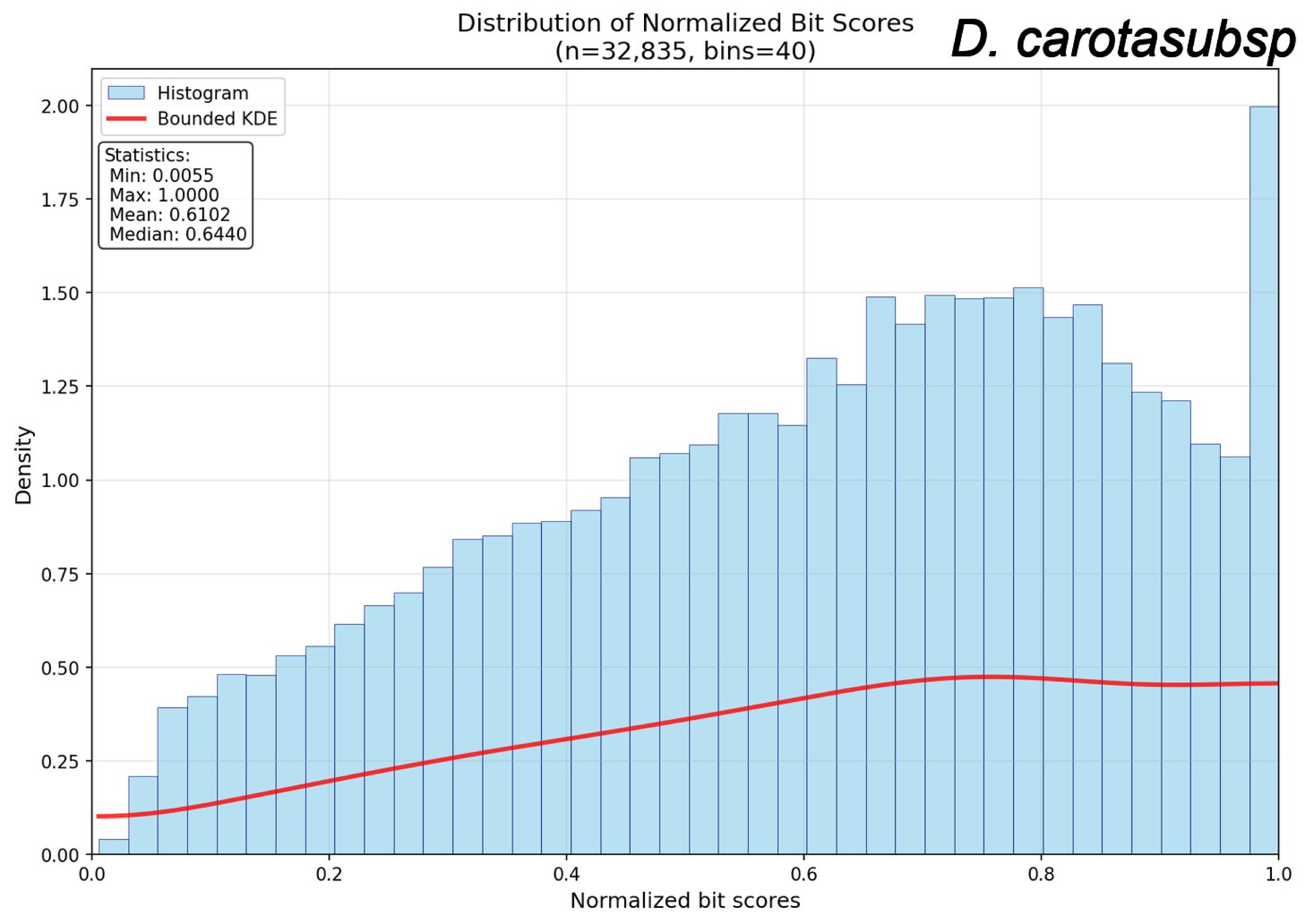
Distribution of Normalized Bit Scores
(n=32,835, bins=40)
D. carotasubsp
Histogram
Bounded KDE
Statistics:
Min: 0.0055
Max: 1.0000
Mean: 0.6102
Median: 0.6440
Density
Normalized bit scores
0.00
0.25
0.50
0.75
1.00
1.25
1.50
1.75
2.00
0.0
0.2
0.4
0.6
0.8
1.0

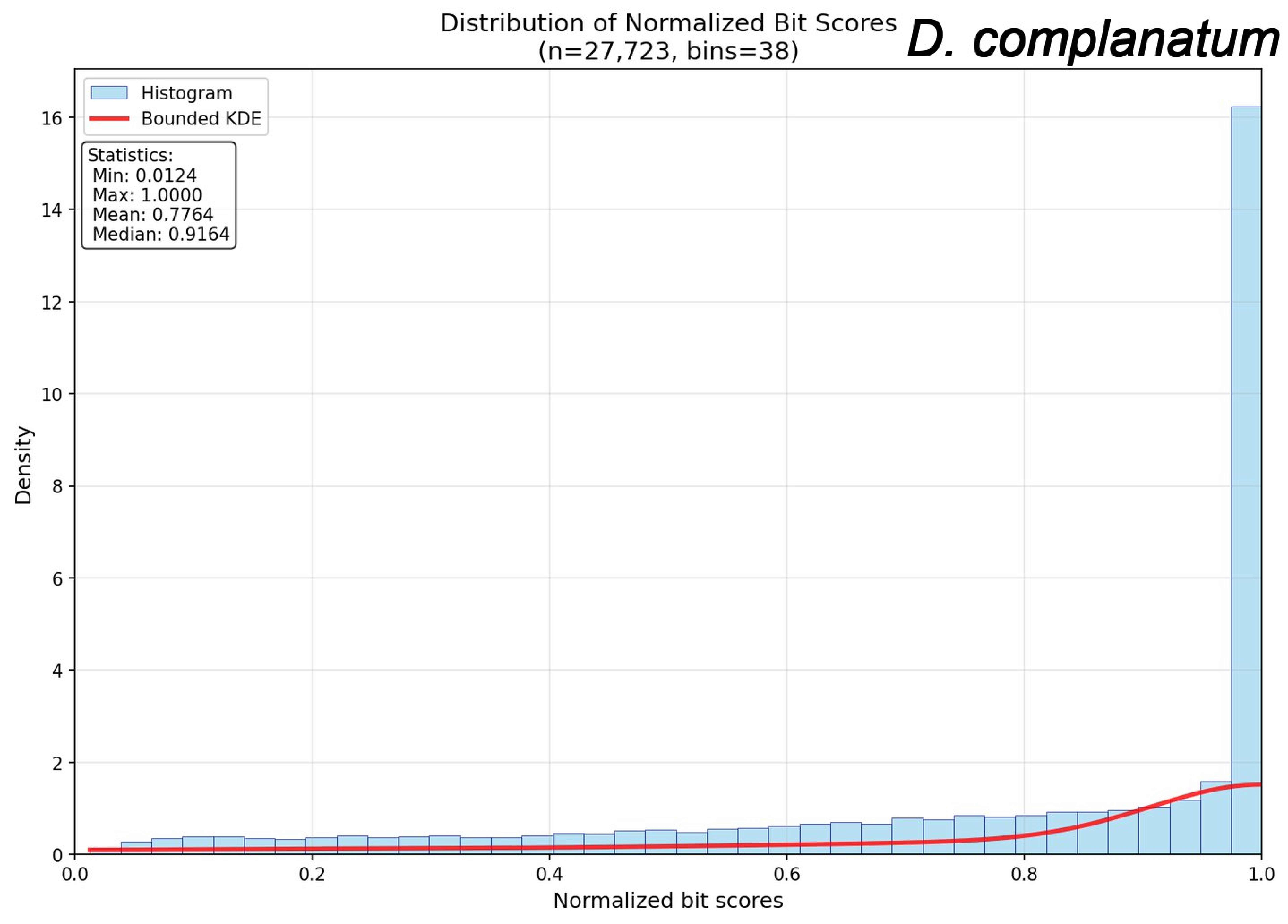

Distribution of Normalized Bit Scores
(n=27,723, bins=38)
D. complanatum
Histogram
Bounded KDE
Statistics:
Min: 0.0124
Max: 1.0000
Mean: 0.7764
Median: 0.9164
Density
Normalized bit scores
0
2
4
6
8
10
12
14
16
0.0
0.2
0.4
0.6
0.8
1.0

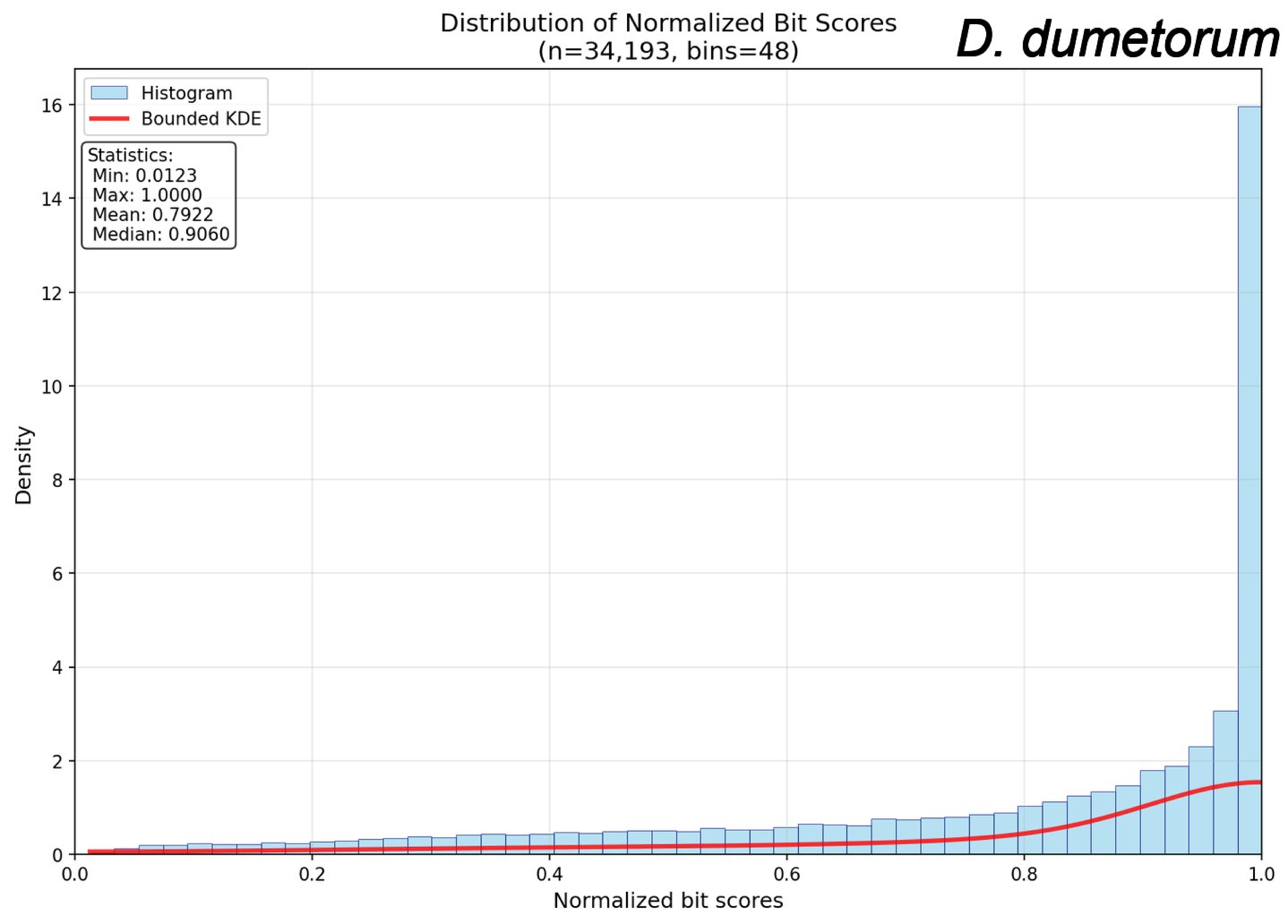
Distribution of Normalized Bit Scores
(n=34,193, bins=48)
D. dumetorum
Histogram
Bounded KDE
Statistics:
Min: 0.0123
Max: 1.0000
Mean: 0.7922
Median: 0.9060
Density
Normalized bit scores
0
2
4
6
8
10
12
14
16
0.0
0.2
0.4
0.6
0.8
1.0

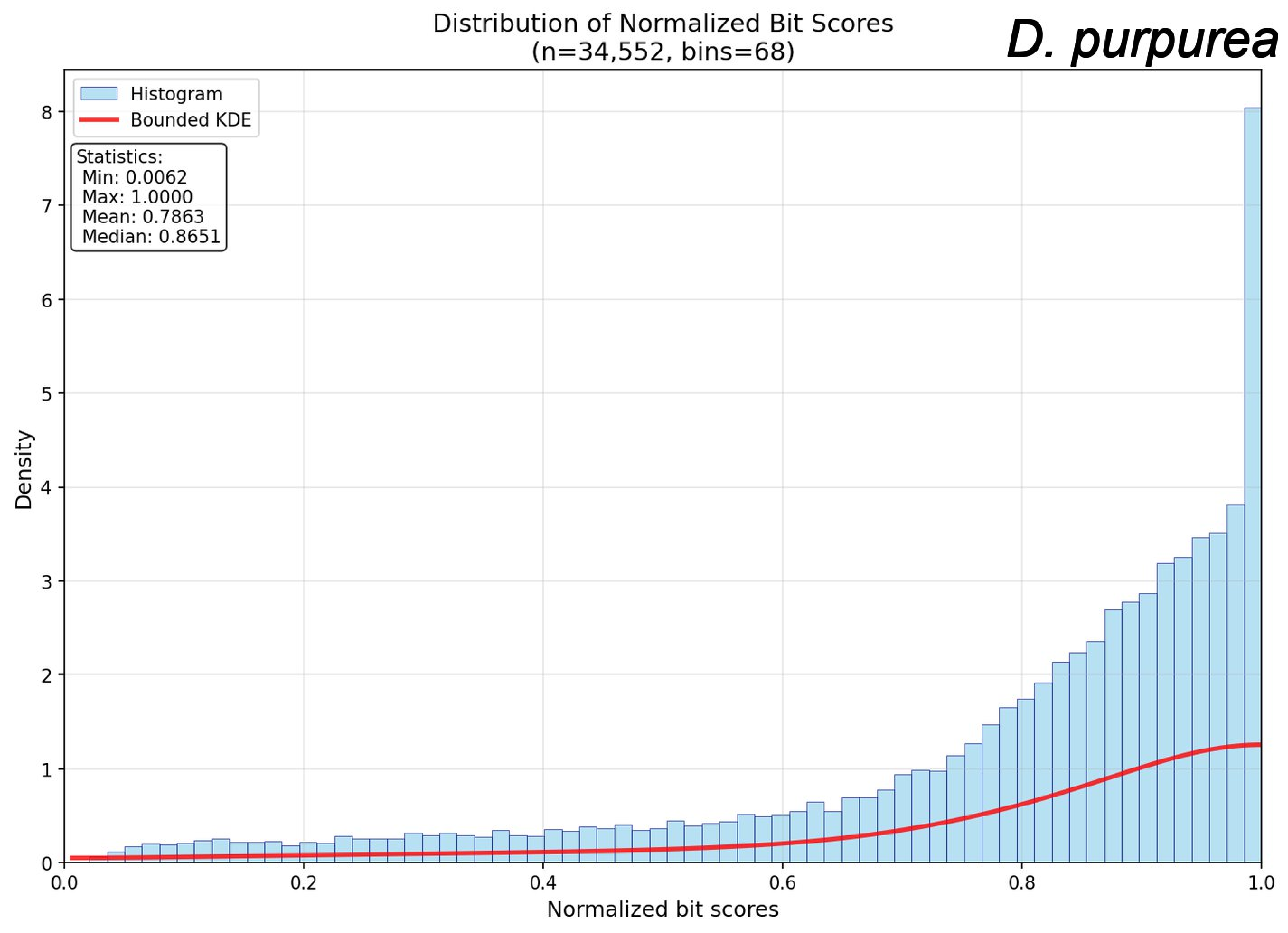
Distribution of Normalized Bit Scores
(n=34,552, bins=68)
D. purpurea
Histogram
Bounded KDE
Statistics:
Min: 0.0062
Max: 1.0000
Mean: 0.7863
Median: 0.8651
Density
Normalized bit scores
0
1
2
3
4
5
6
7
8
0.0
0.2
0.4
0.6
0.8
1.0

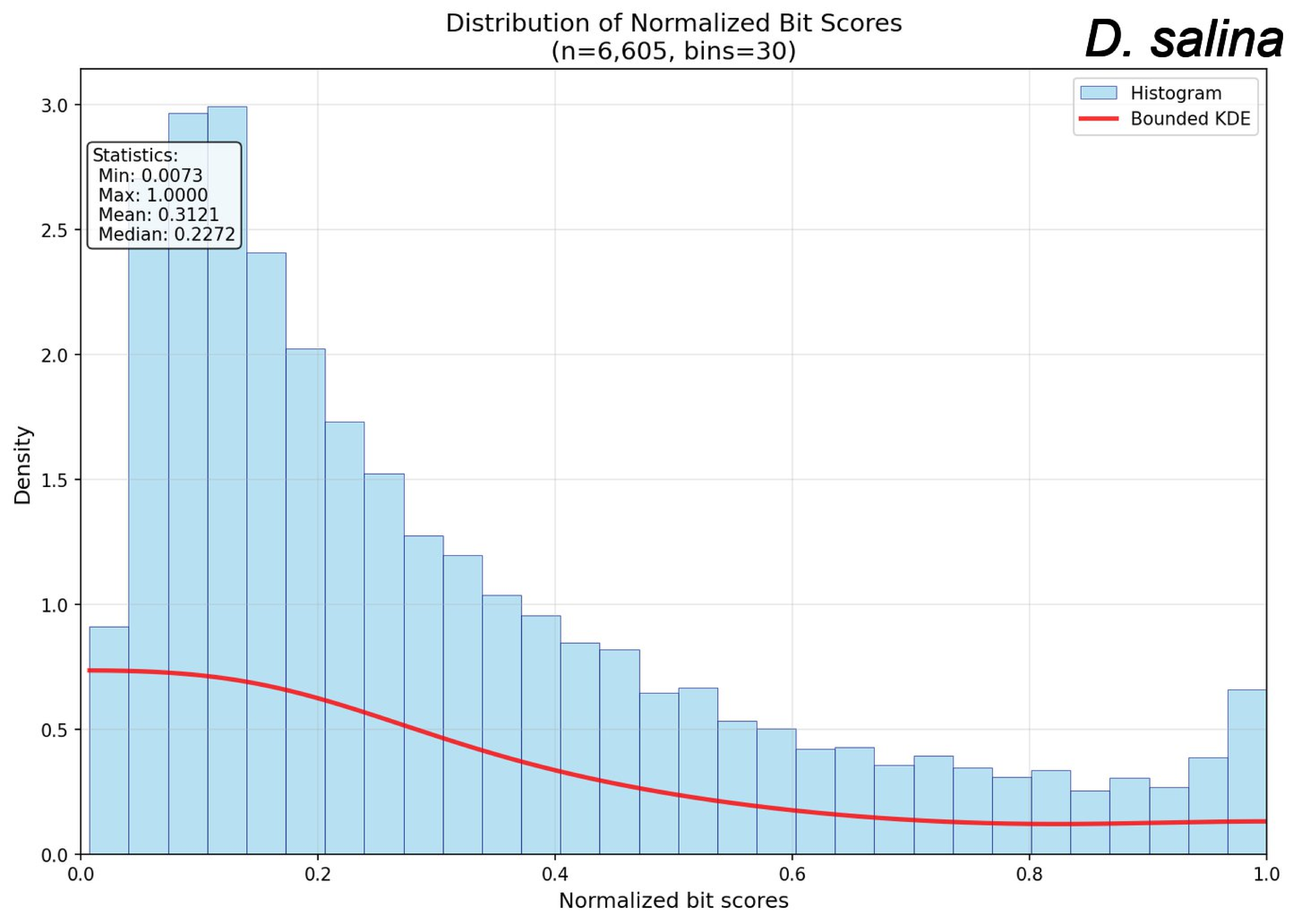

Distribution of Normalized Bit Scores
(n=6,605, bins=30)
D. salina
Histogram
Bounded KDE
Statistics:
Min: 0.0073
Max: 1.0000
Mean: 0.3121
Median: 0.2272
Density
Normalized bit scores
0.0
0.5
1.0
1.5
2.0
2.5
3.0
0.2
0.4
0.6
0.8
1.0

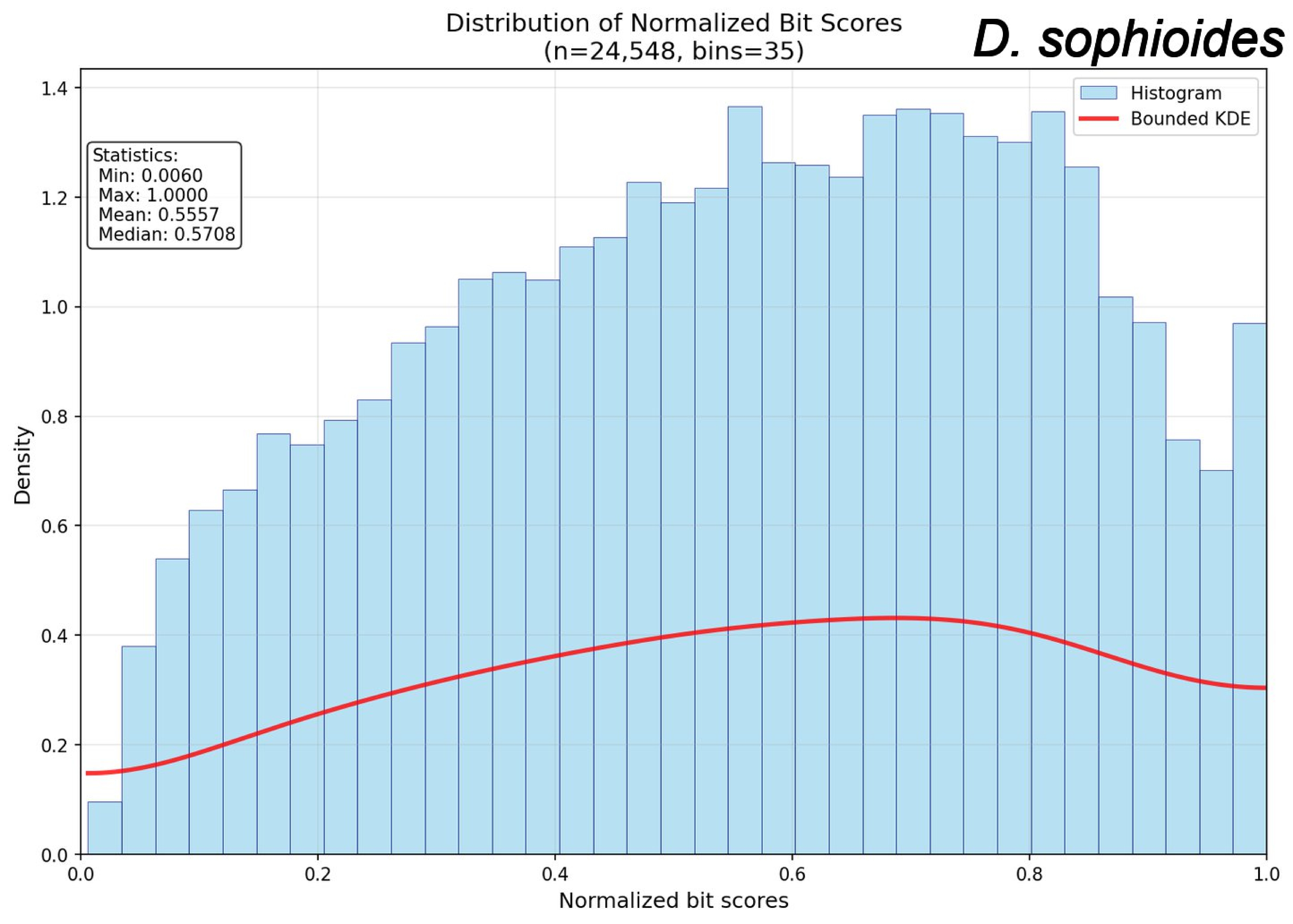

Distribution of Normalized Bit Scores
(n=24,548, bins=35)
D. sophioides
Histogram
Bounded KDE
Statistics:
Min: 0.0060
Max: 1.0000
Mean: 0.5557
Median: 0.5708
Density
Normalized bit scores
0.0
0.2
0.4
0.6
0.8
1.0
1.2
1.4

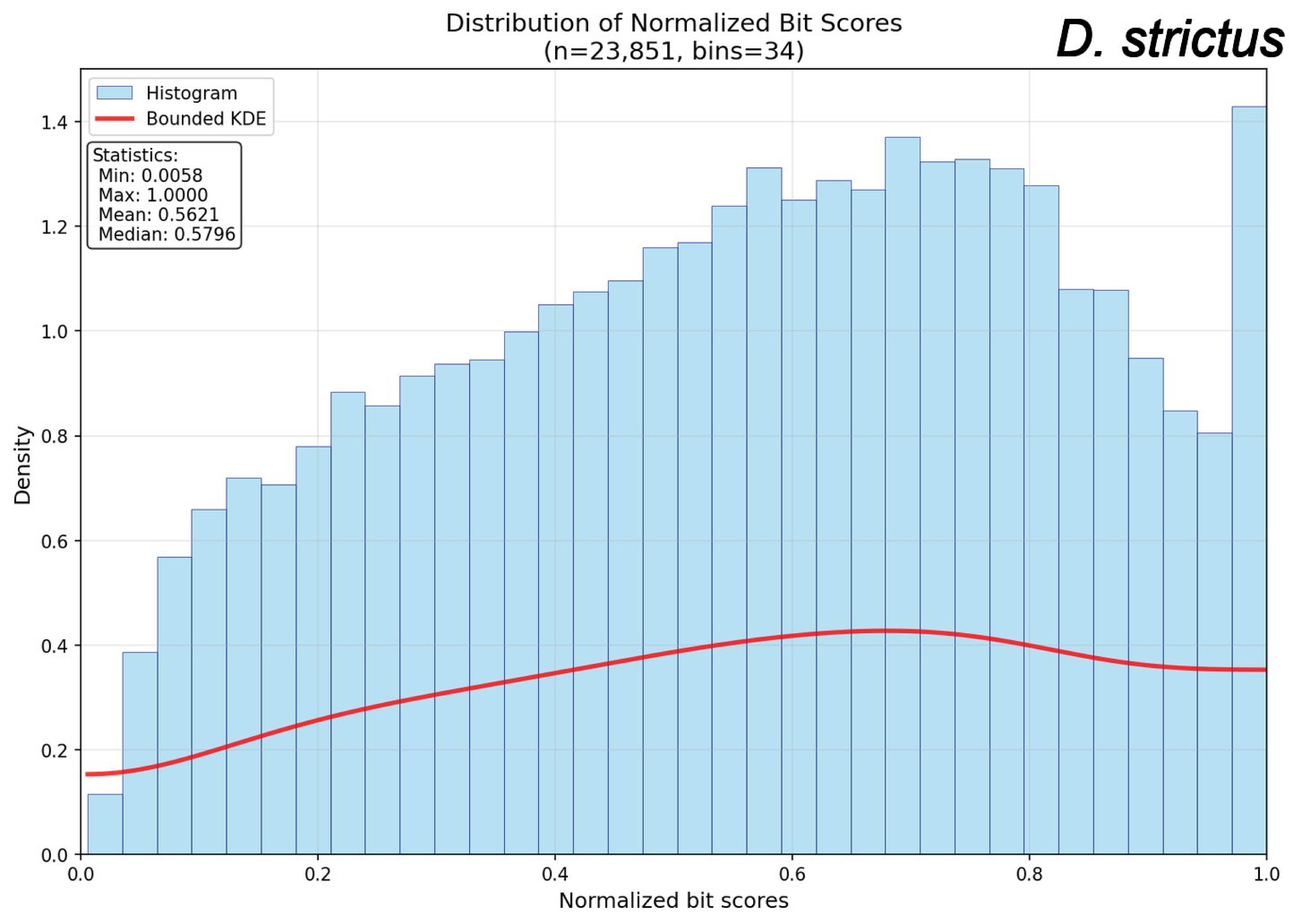

Distribution of Normalized Bit Scores
(n=23,851, bins=34)
D. strictus
Histogram
Bounded KDE
Statistics:
Min: 0.0058
Max: 1.0000
Mean: 0.5621
Median: 0.5796
Density
Normalized bit scores
0.0
0.2
0.4
0.6
0.8
1.0
1.2
1.4

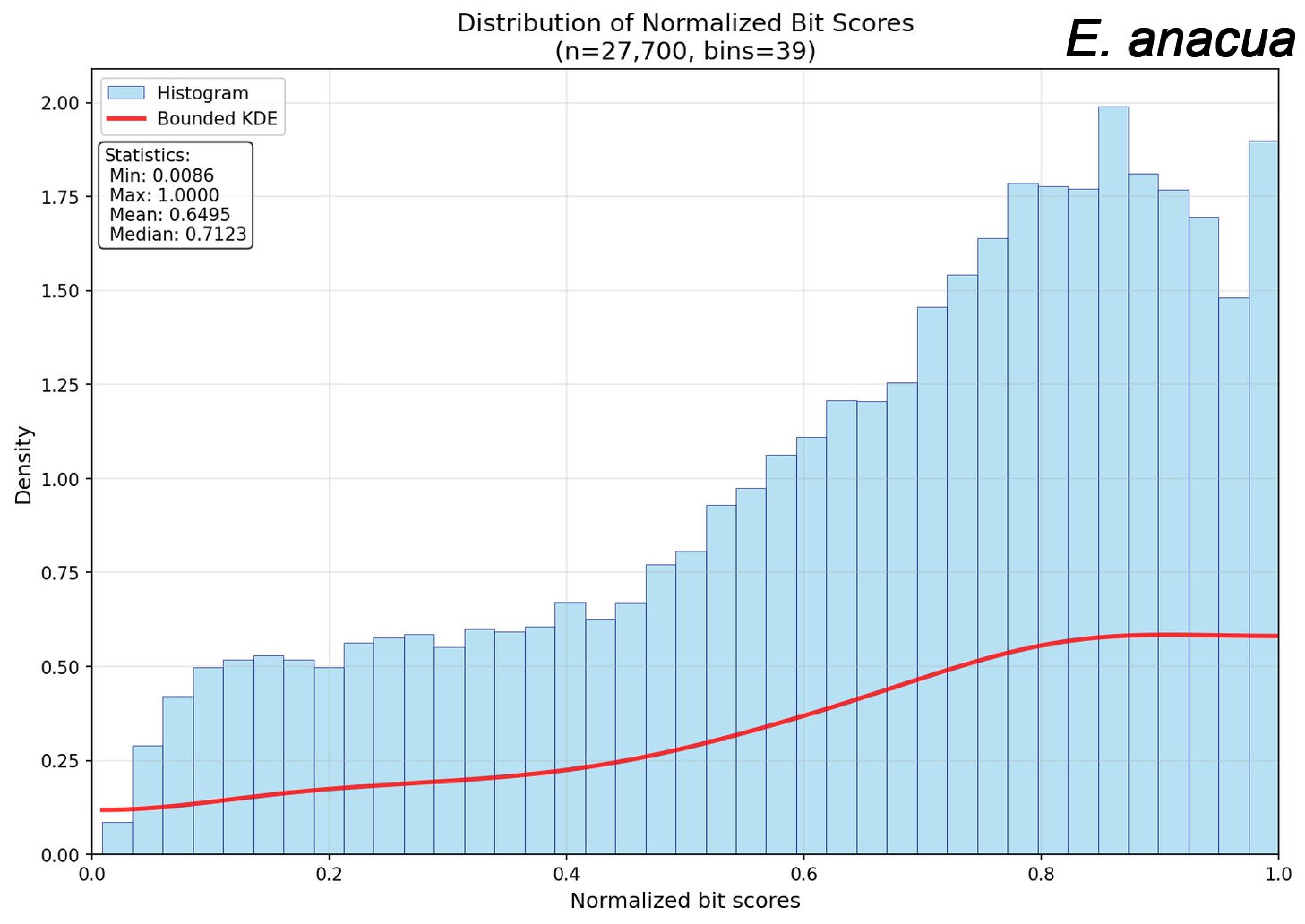
Distribution of Normalized Bit Scores
(n=27,700, bins=39)
E. anacua
Histogram
Bounded KDE
Statistics:
Min: 0.0086
Max: 1.0000
Mean: 0.6495
Median: 0.7123
Density
Normalized bit scores
0.00
0.25
0.50
0.75
1.00
1.25
1.50
1.75
2.00
0.0
0.2
0.4
0.6
0.8
1.0

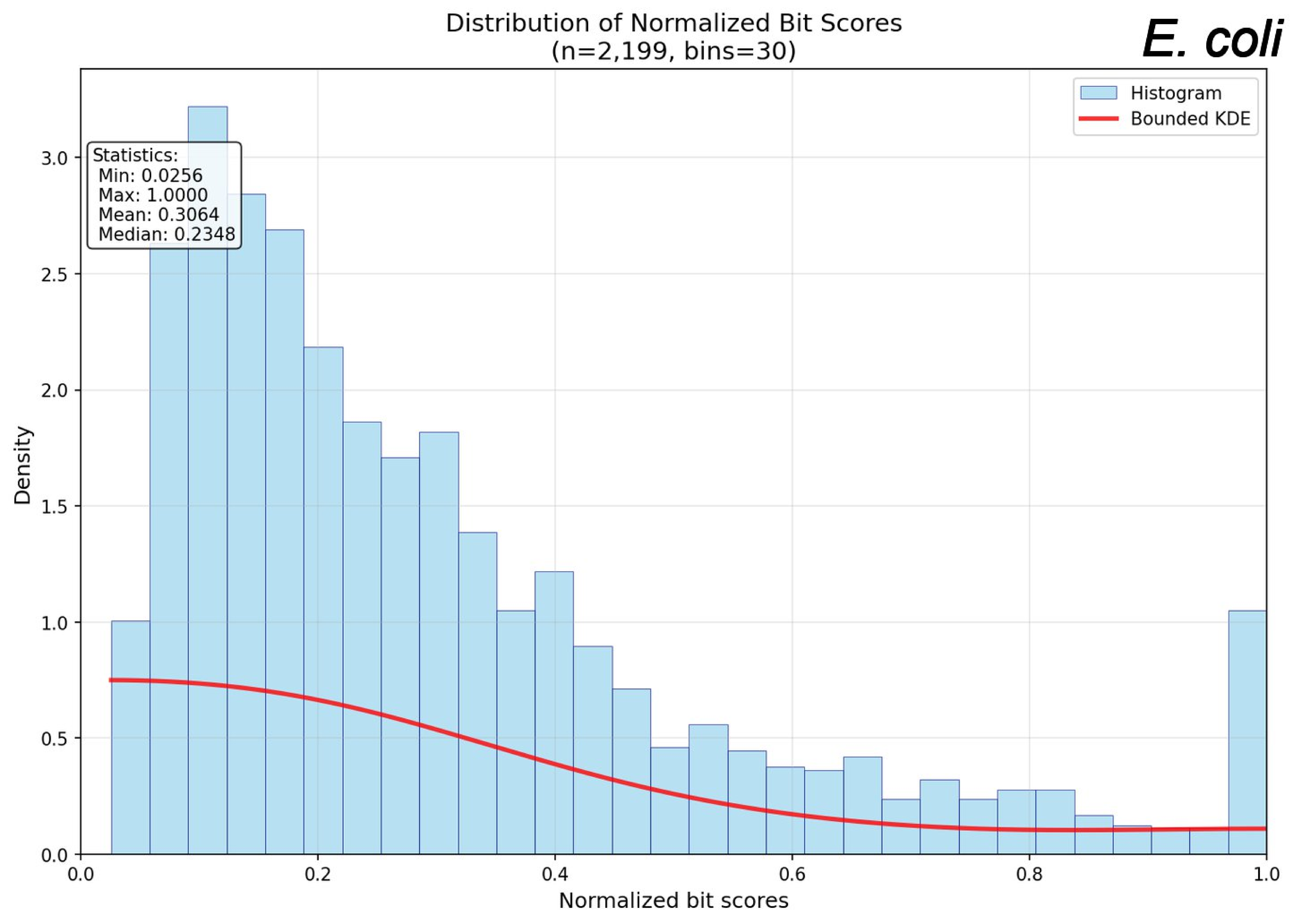

Distribution of Normalized Bit Scores
(n=2,199, bins=30)
E. coli
Histogram
Bounded KDE
Statistics:
Min: 0.0256
Max: 1.0000
Mean: 0.3064
Median: 0.2348
Density
Normalized bit scores
0.0
0.5
1.0
1.5
2.0
2.5
3.0
0.2
0.4
0.6
0.8

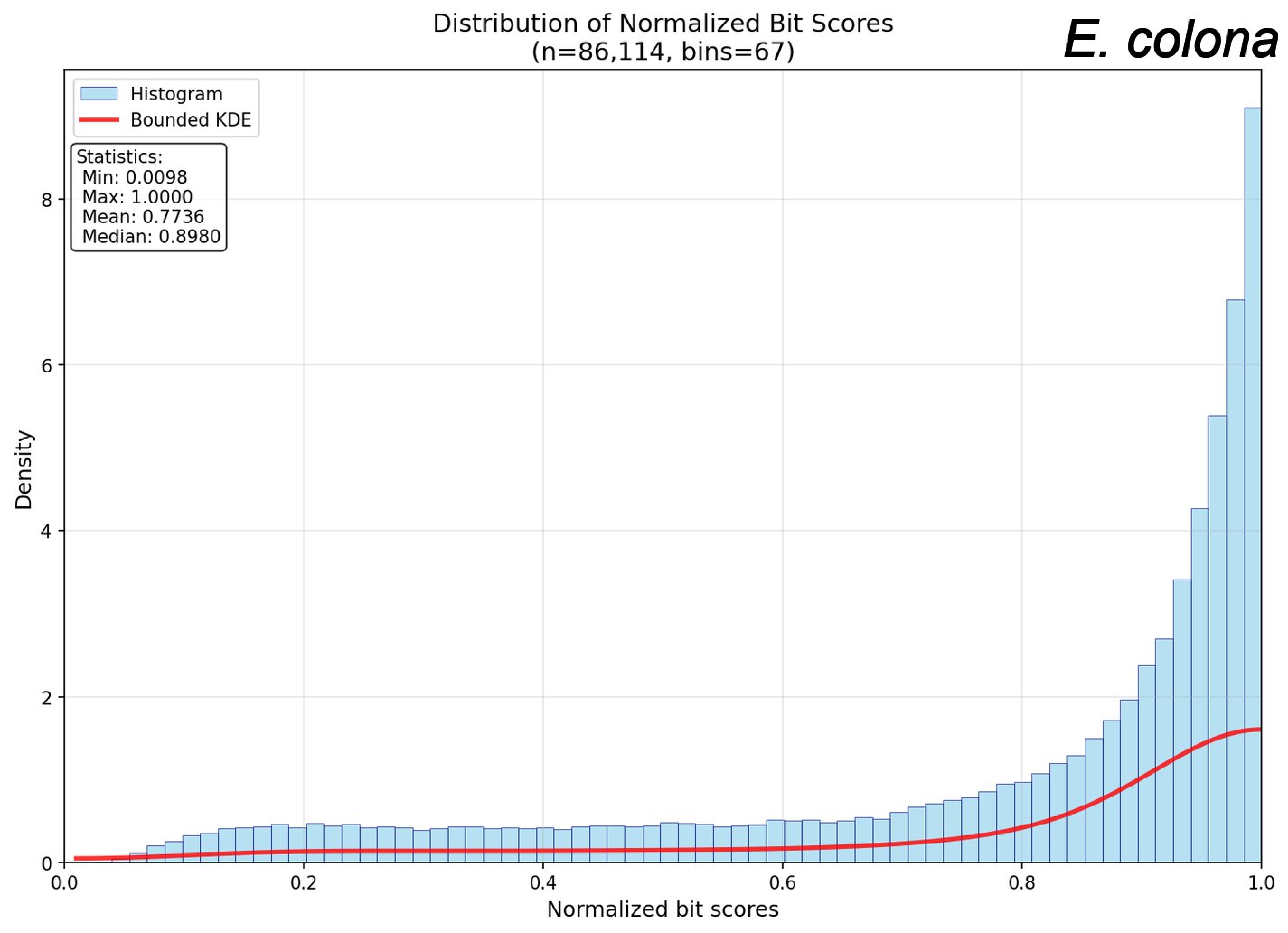

Distribution of Normalized Bit Scores
(n=86,114, bins=67)
E. colona
Histogram
Bounded KDE
Statistics:
Min: 0.0098
Max: 1.0000
Mean: 0.7736
Median: 0.8980
Density
Normalized bit scores
0
2
4
6
8
0.0
0.2
0.4
0.6
0.8
1.0

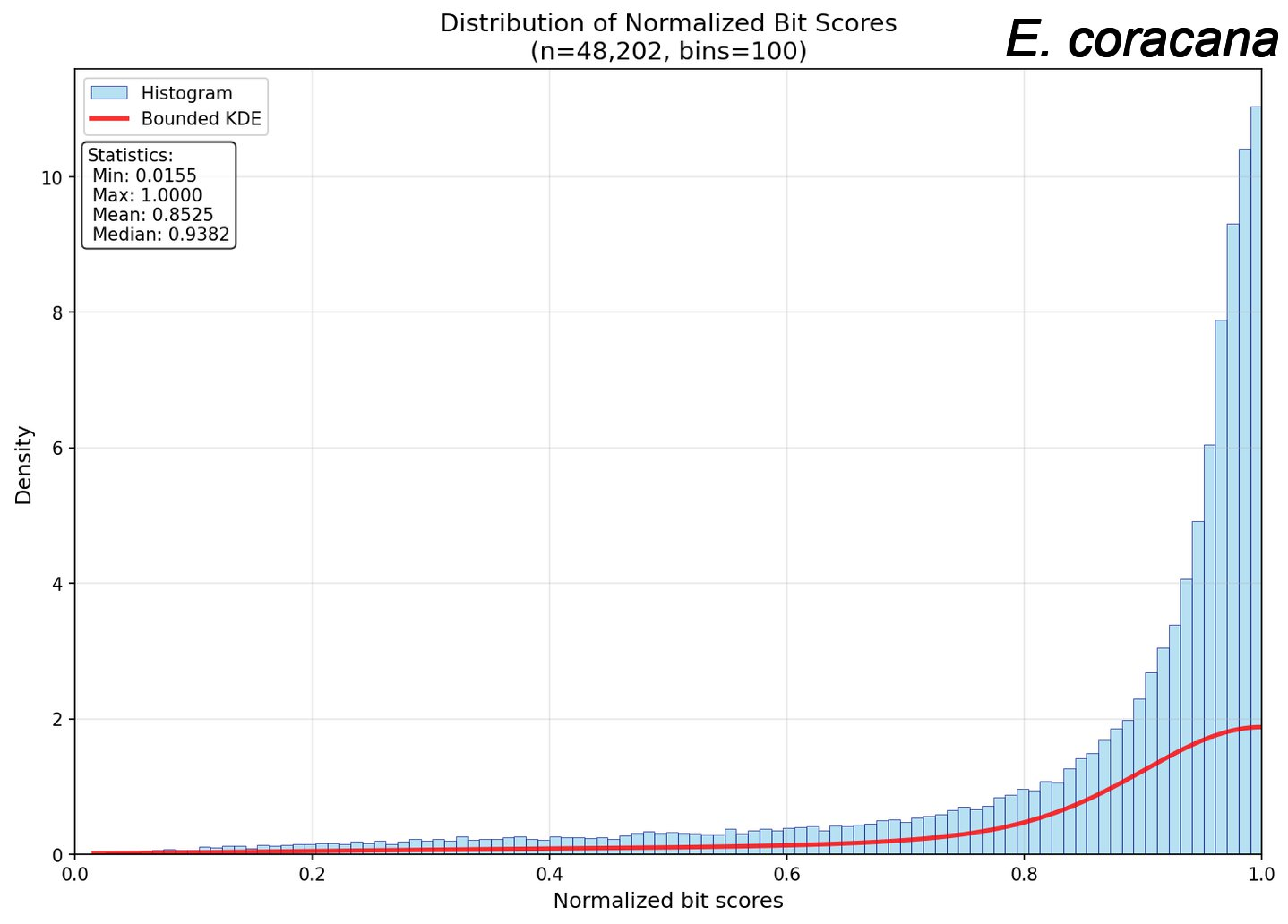
Distribution of Normalized Bit Scores
(n=48,202, bins=100)
E. coracana
Histogram
Bounded KDE
Statistics:
Min: 0.0155
Max: 1.0000
Mean: 0.8525
Median: 0.9382
Density
Normalized bit scores
0.0
0.2
0.4
0.6
0.8
1.0
0
2
4
6
8
10

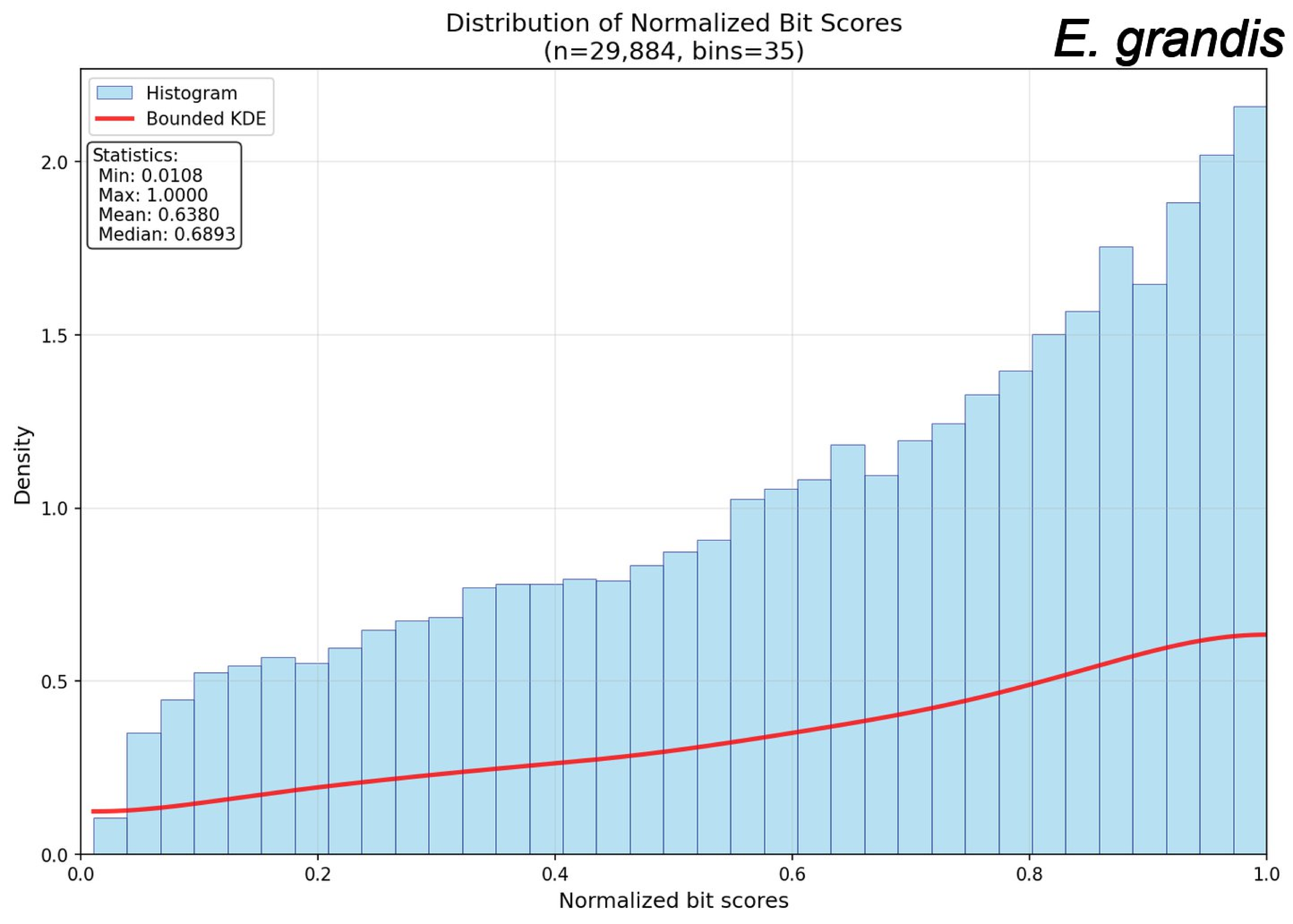

Distribution of Normalized Bit Scores
(n=29,884, bins=35)
E. grandis
Histogram
Bounded KDE
Statistics:
Min: 0.0108
Max: 1.0000
Mean: 0.6380
Median: 0.6893
Density
0.0
0.5
1.0
1.5
2.0
0.0
0.2
0.4
0.6
0.8
1.0
Normalized bit scores

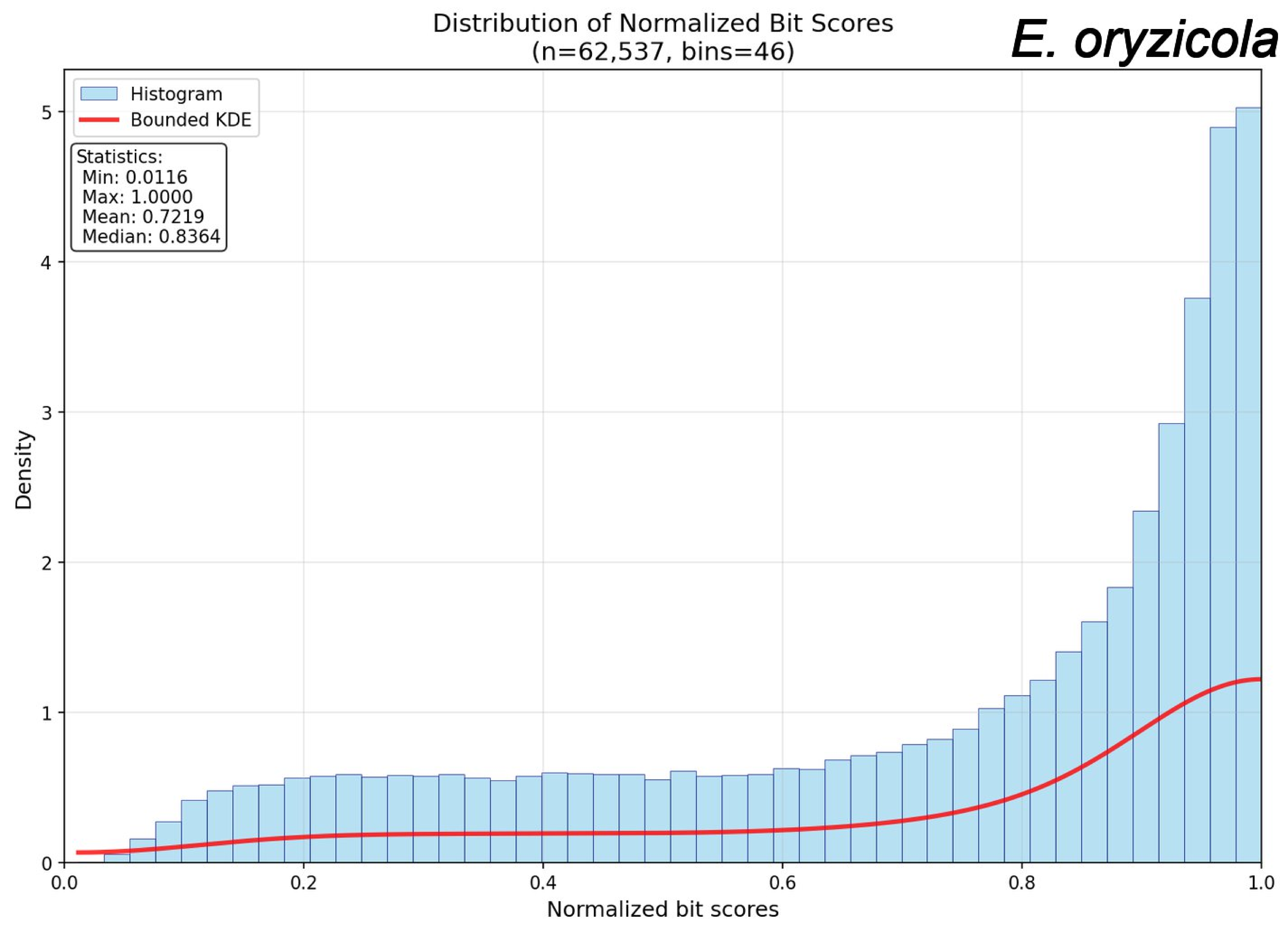
Distribution of Normalized Bit Scores
(n=62,537, bins=46)
E. oryzicola
Histogram
Bounded KDE
Statistics:
Min: 0.0116
Max: 1.0000
Mean: 0.7219
Median: 0.8364
Density
Normalized bit scores
0
1
2
3
4
5
0.0
0.2
0.4
0.6
0.8
1.0

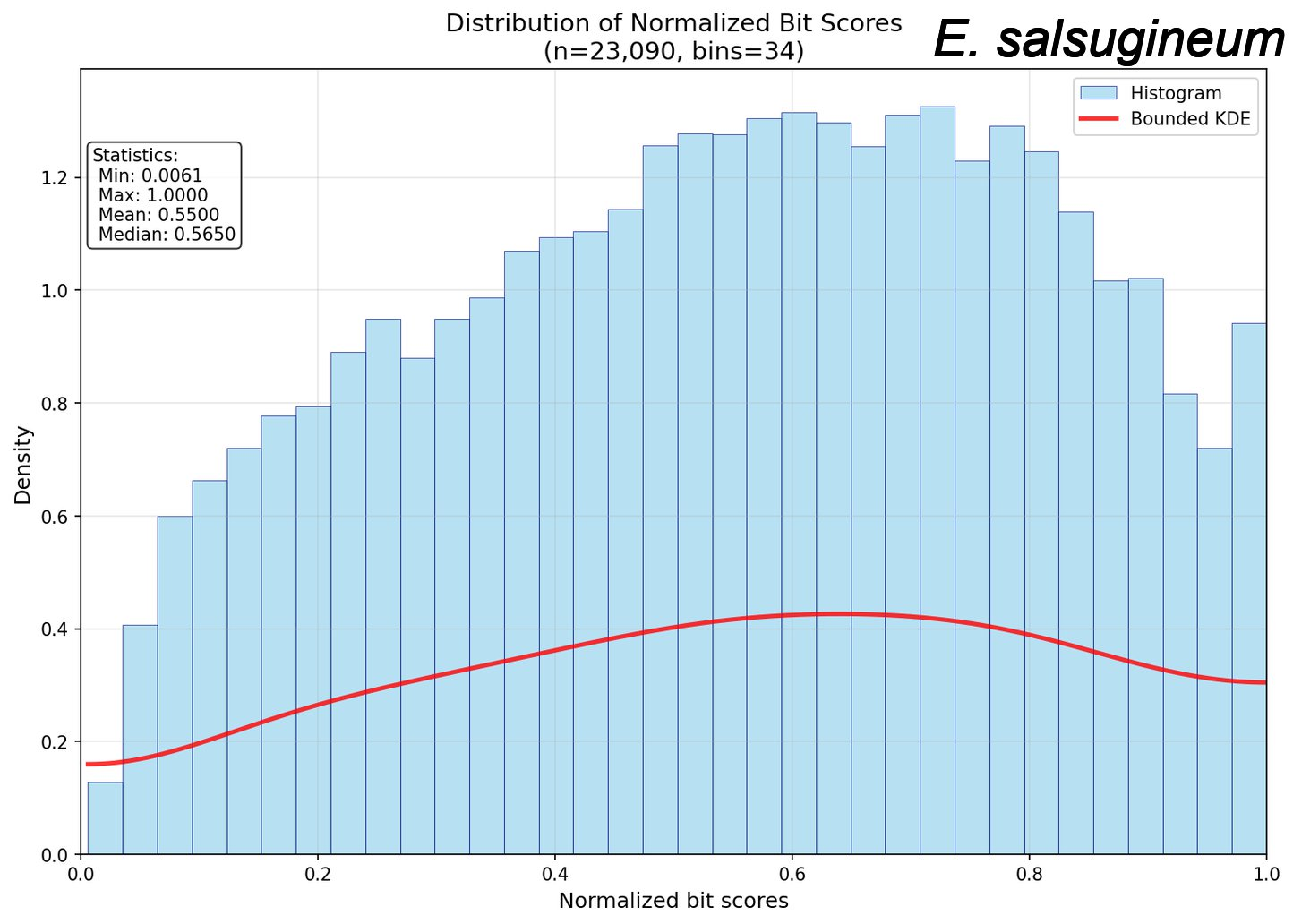
Distribution of Normalized Bit Scores
(n=23,090, bins=34)
E. salsugineum
Histogram
Bounded KDE
Statistics:
Min: 0.0061
Max: 1.0000
Mean: 0.5500
Median: 0.5650
Density
Normalized bit scores
0.0
0.2
0.4
0.6
0.8
1.0
1.2

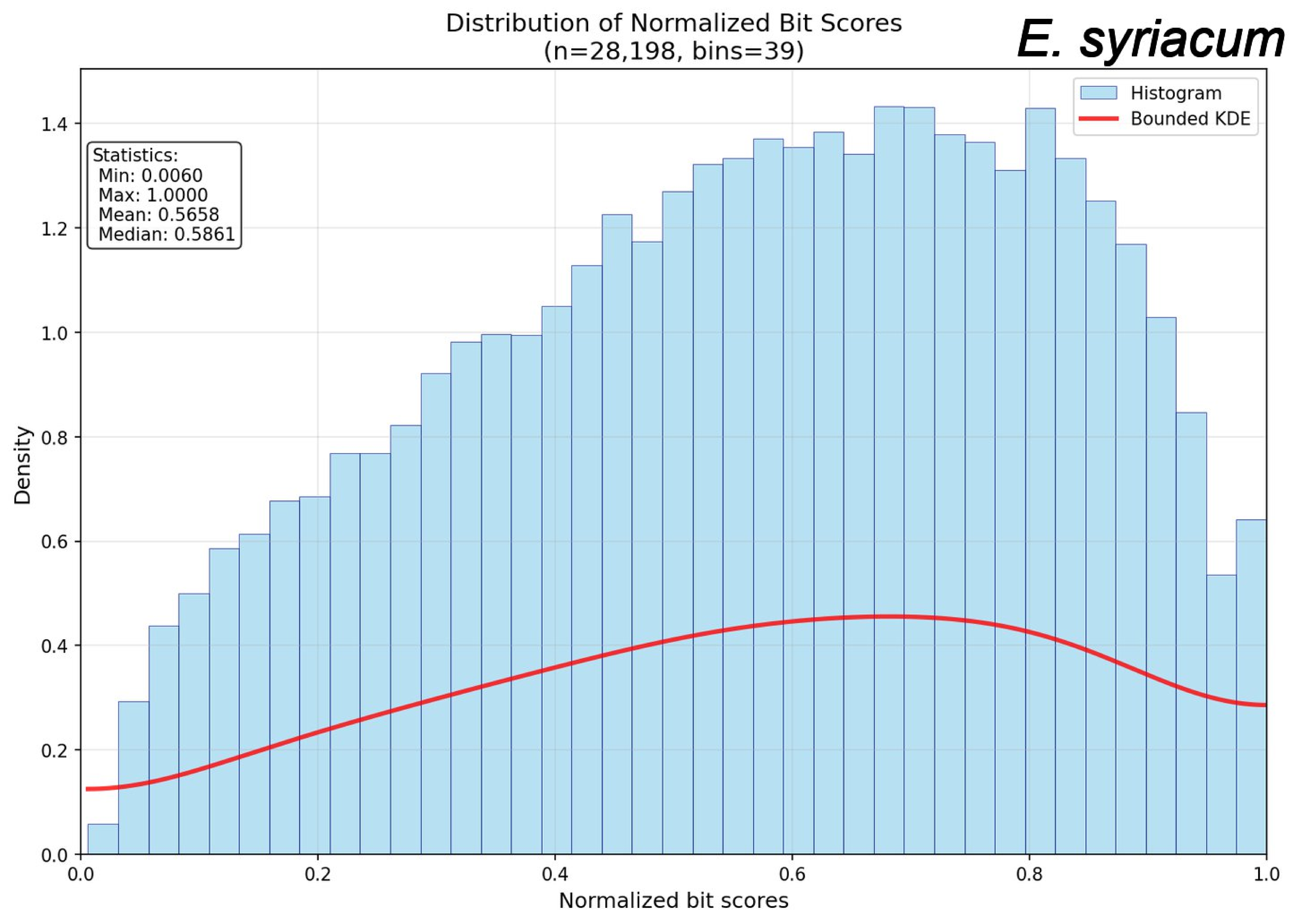

Distribution of Normalized Bit Scores
(n=28,198, bins=39)
E. syriacum
Histogram
Bounded KDE
Statistics:
Min: 0.0060
Max: 1.0000
Mean: 0.5658
Median: 0.5861
Density
Normalized bit scores
0.0
0.2
0.4
0.6
0.8
1.0
1.2
1.4

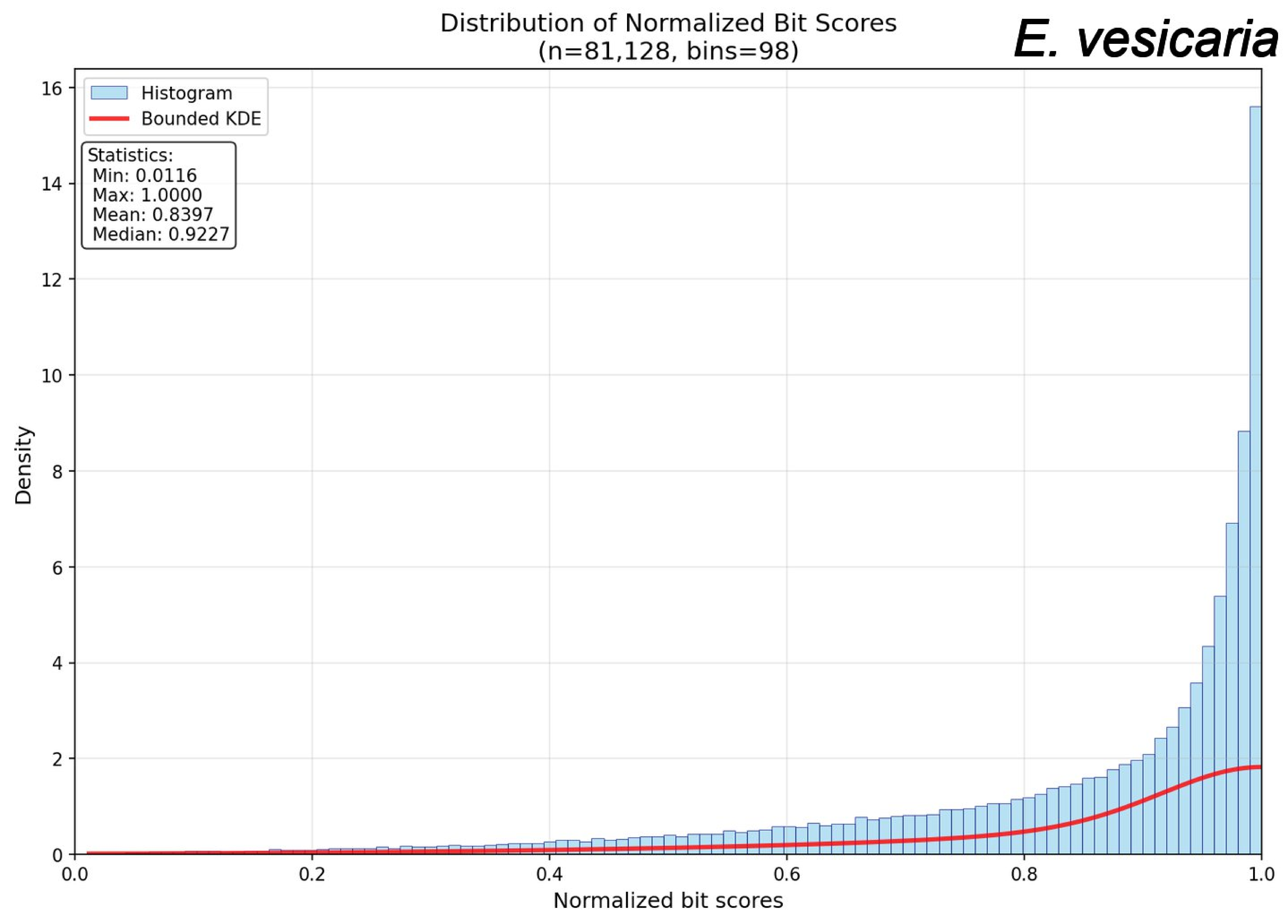

Distribution of Normalized Bit Scores
(n=81,128, bins=98)
E. vesicaria
Histogram
Bounded KDE
Statistics:
Min: 0.0116
Max: 1.0000
Mean: 0.8397
Median: 0.9227
Density
Normalized bit scores
0.0
0.2
0.4
0.6
0.8
1.0
0
2
4
6
8
10
12
14
16

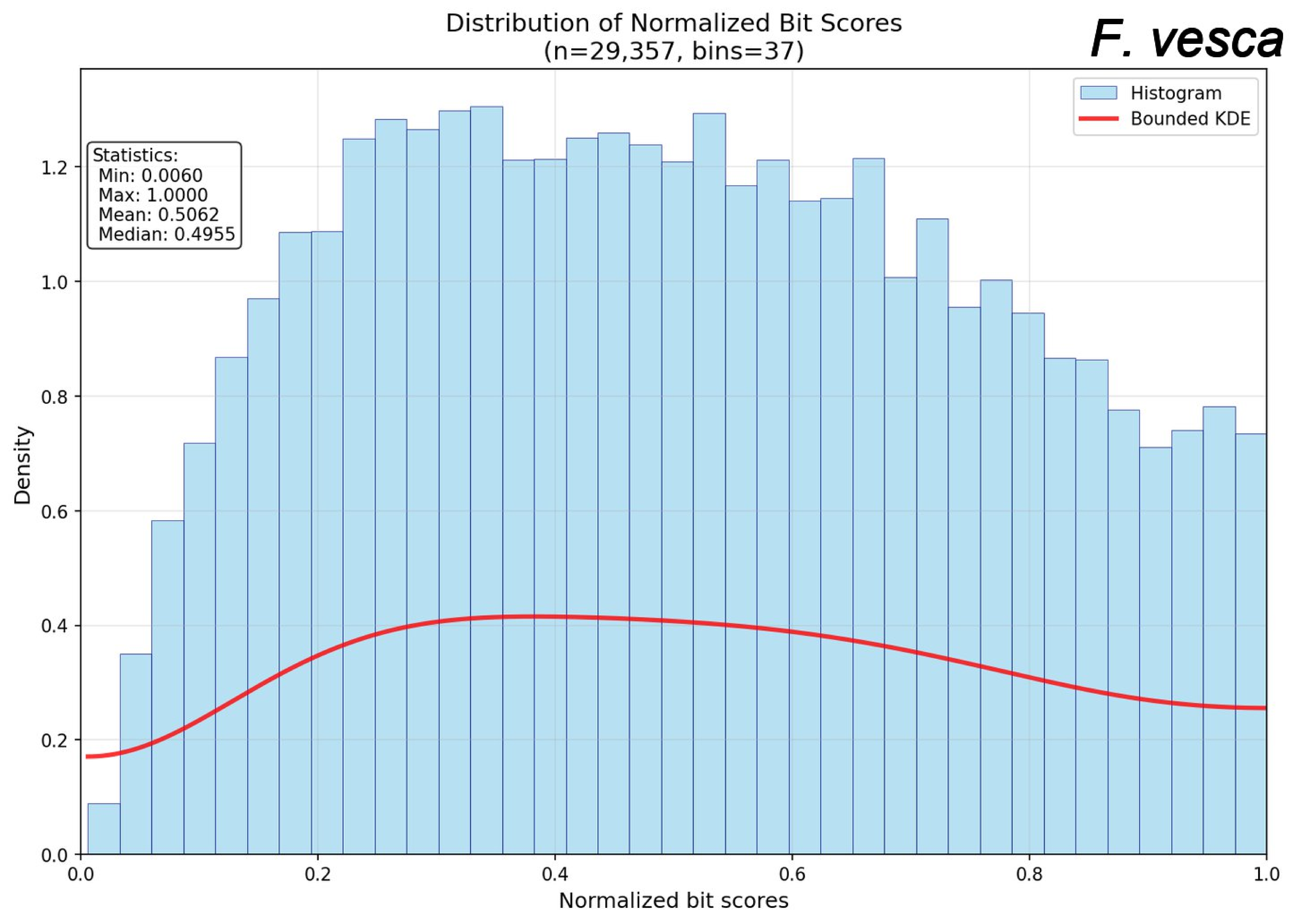

Distribution of Normalized Bit Scores
(n=29,357, bins=37)
F. vesca
Histogram
Bounded KDE
Statistics:
Min: 0.0060
Max: 1.0000
Mean: 0.5062
Median: 0.4955
Density
Normalized bit scores
0.0
0.2
0.4
0.6
0.8
1.0
1.2

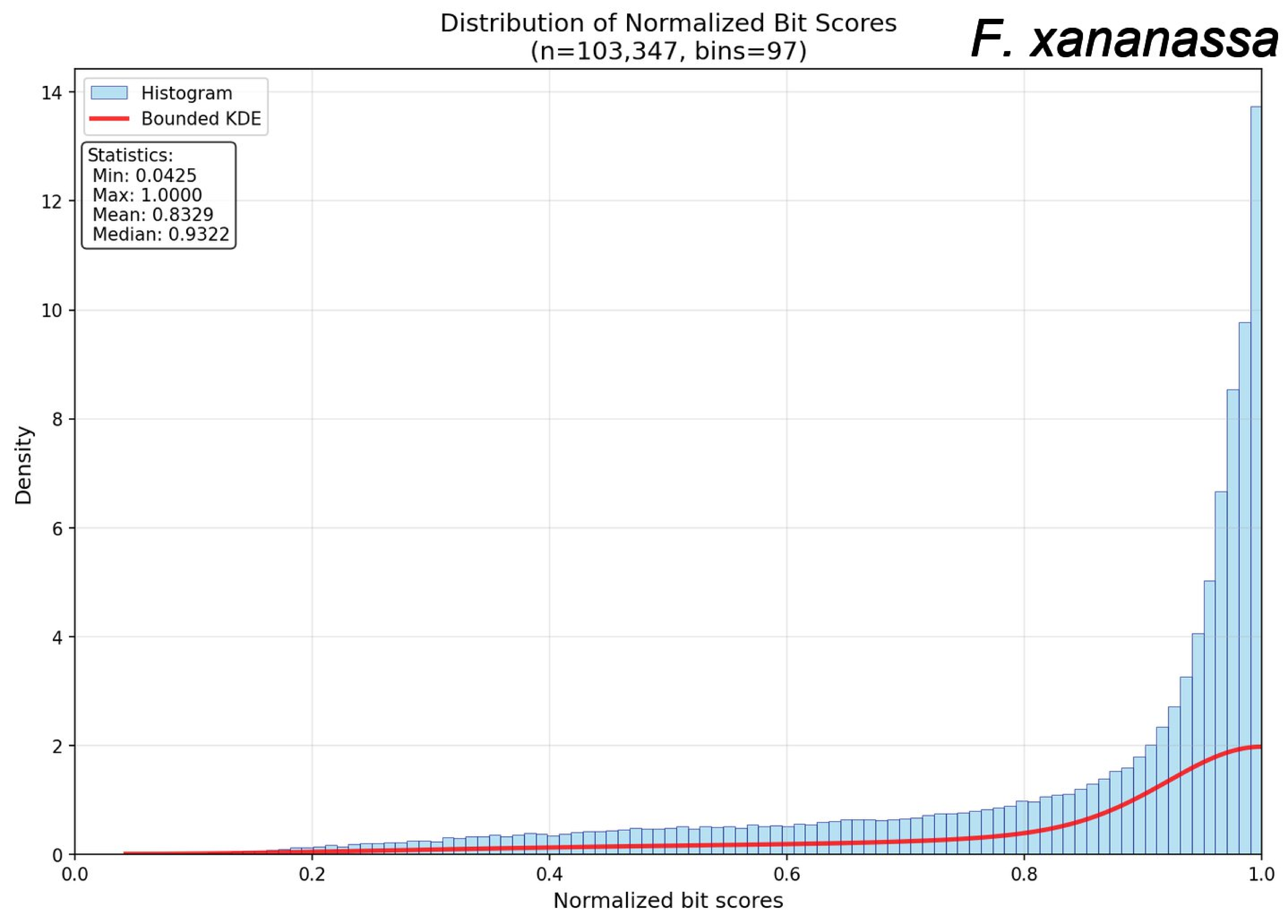

Distribution of Normalized Bit Scores
(n=103,347, bins=97)
F. xananassa
Histogram
Bounded KDE
Statistics:
Min: 0.0425
Max: 1.0000
Mean: 0.8329
Median: 0.9322
Density
Normalized bit scores
0.0
0.2
0.4
0.6
0.8
1.0
0
2
4
6
8
10
12
14

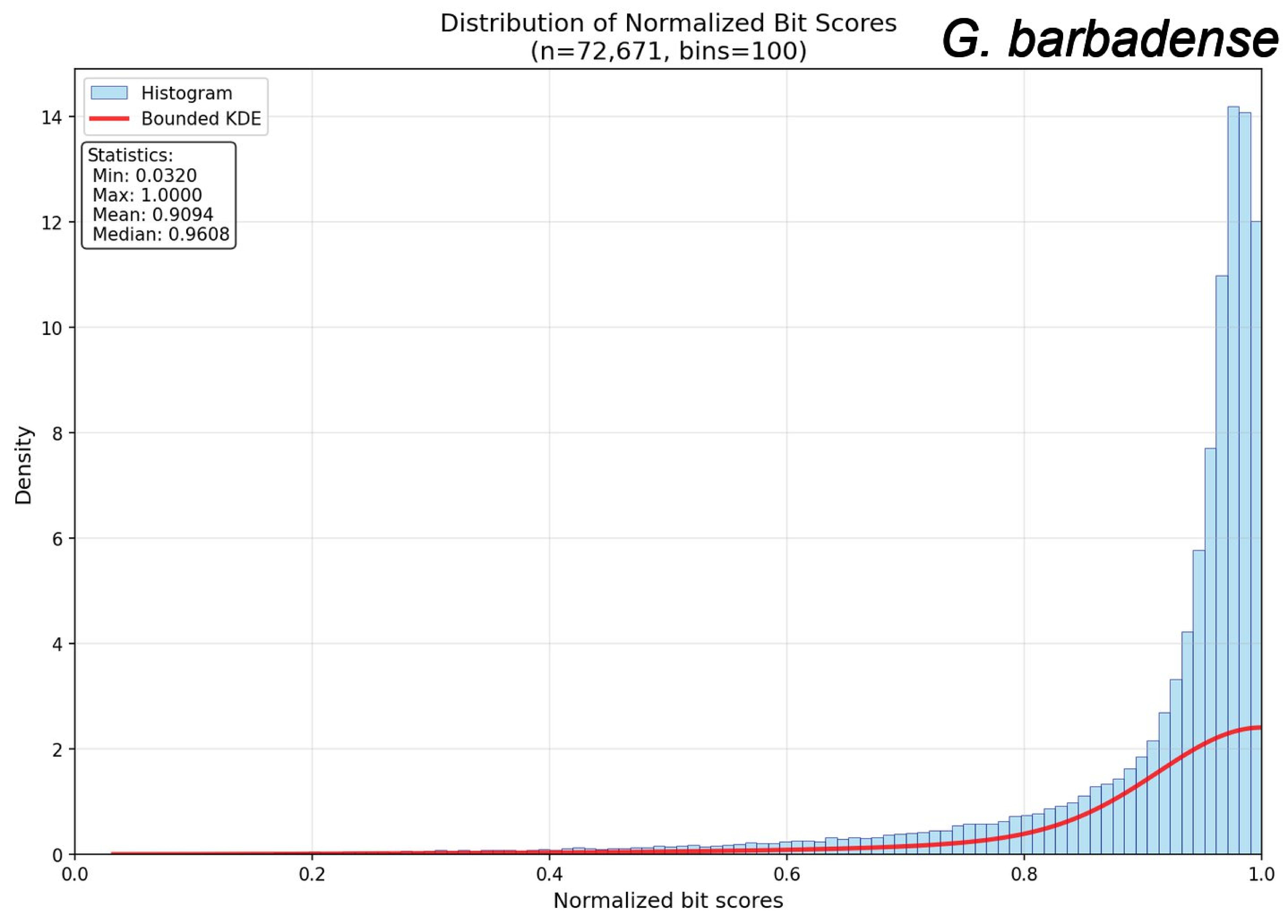

Distribution of Normalized Bit Scores
(n=72,671, bins=100)
G. barbadense
Histogram
Bounded KDE
Statistics:
Min: 0.0320
Max: 1.0000
Mean: 0.9094
Median: 0.9608
Density
Normalized bit scores
0.0
0.2
0.4
0.6
0.8
1.0
0
2
4
6
8
10
12
14

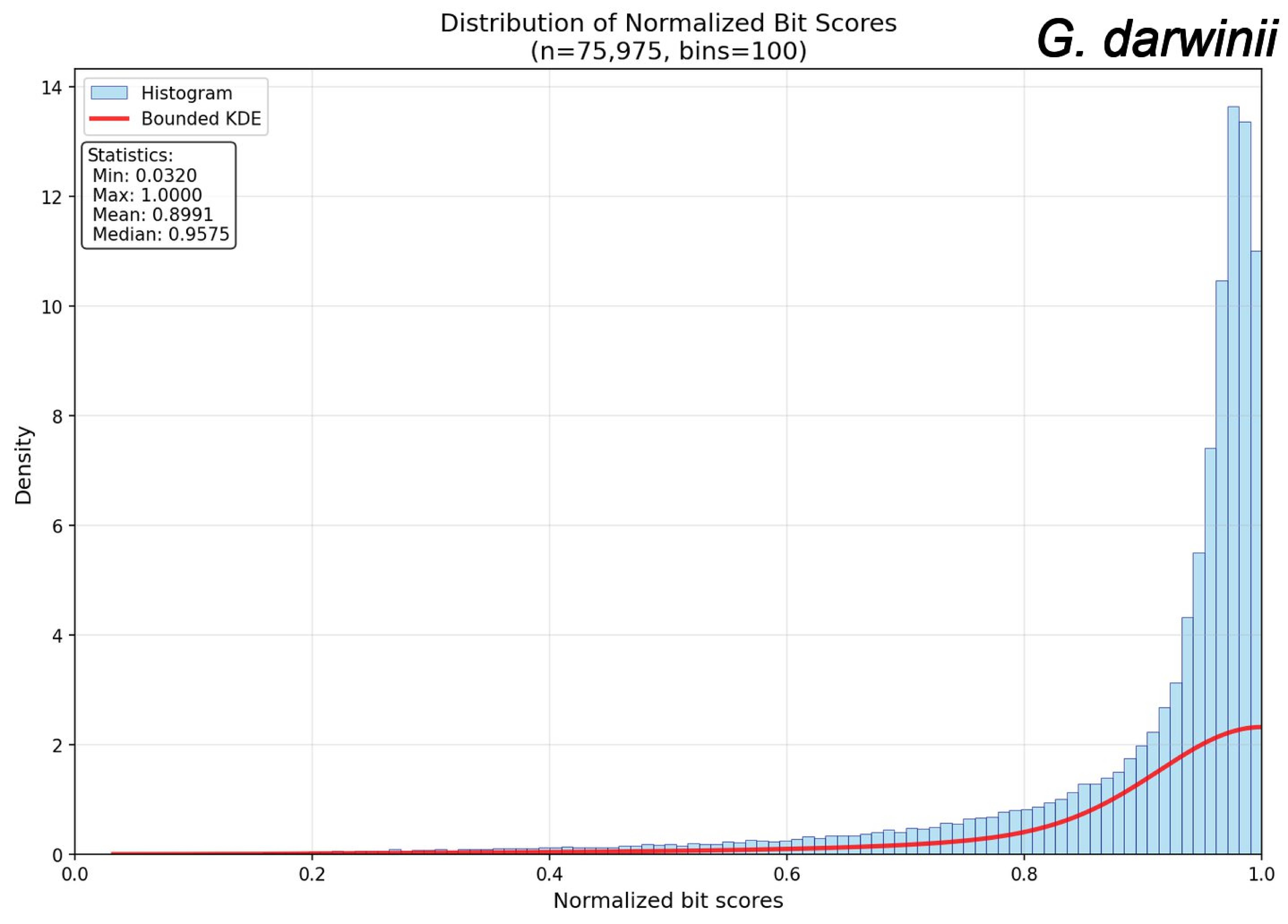

Distribution of Normalized Bit Scores
(n=75,975, bins=100)
G. darwinii
Histogram
Bounded KDE
Statistics:
Min: 0.0320
Max: 1.0000
Mean: 0.8991
Median: 0.9575
Density
Normalized bit scores

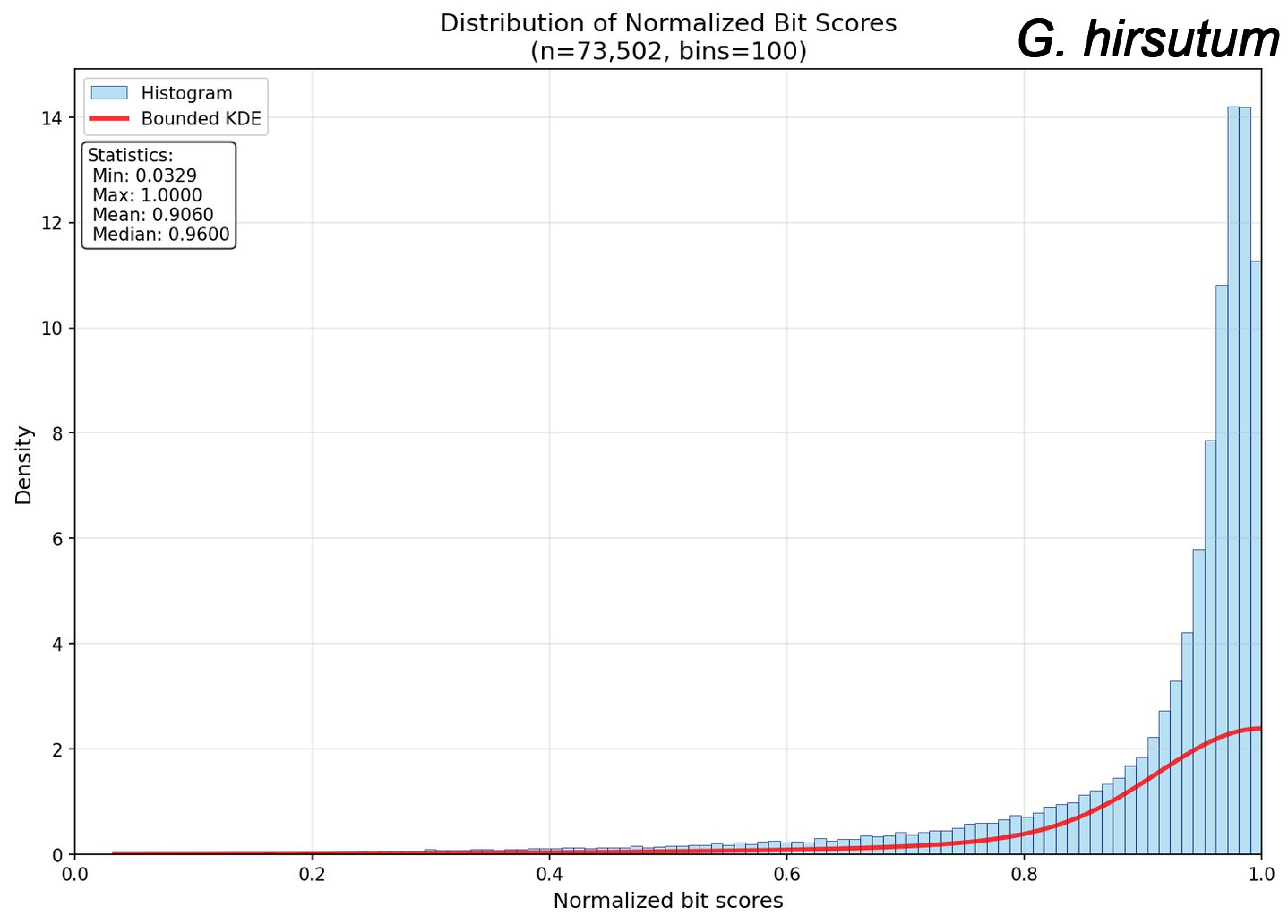
Distribution of Normalized Bit Scores
(n=73,502, bins=100)
G. hirsutum
Histogram
Bounded KDE
Statistics:
Min: 0.0329
Max: 1.0000
Mean: 0.9060
Median: 0.9600
Density
Normalized bit scores
0
2
4
6
8
10
12
14
0.0
0.2
0.4
0.6
0.8
1.0

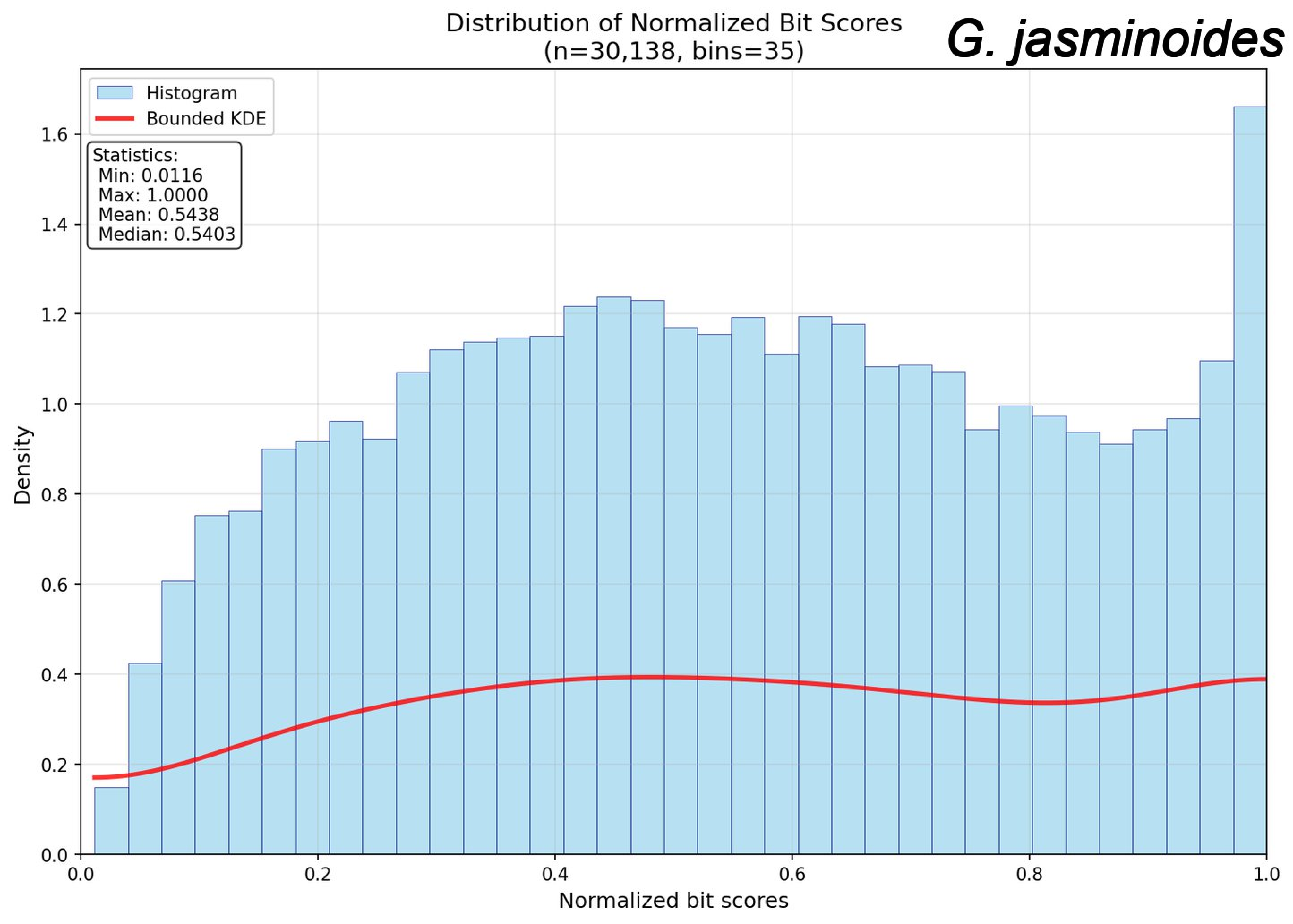

Distribution of Normalized Bit Scores
(n=30,138, bins=35)
G. jasminoides
Histogram
Bounded KDE
Statistics:
Min: 0.0116
Max: 1.0000
Mean: 0.5438
Median: 0.5403
Density
Normalized bit scores
0.0
0.2
0.4
0.6
0.8
1.0
1.2
1.4
1.6

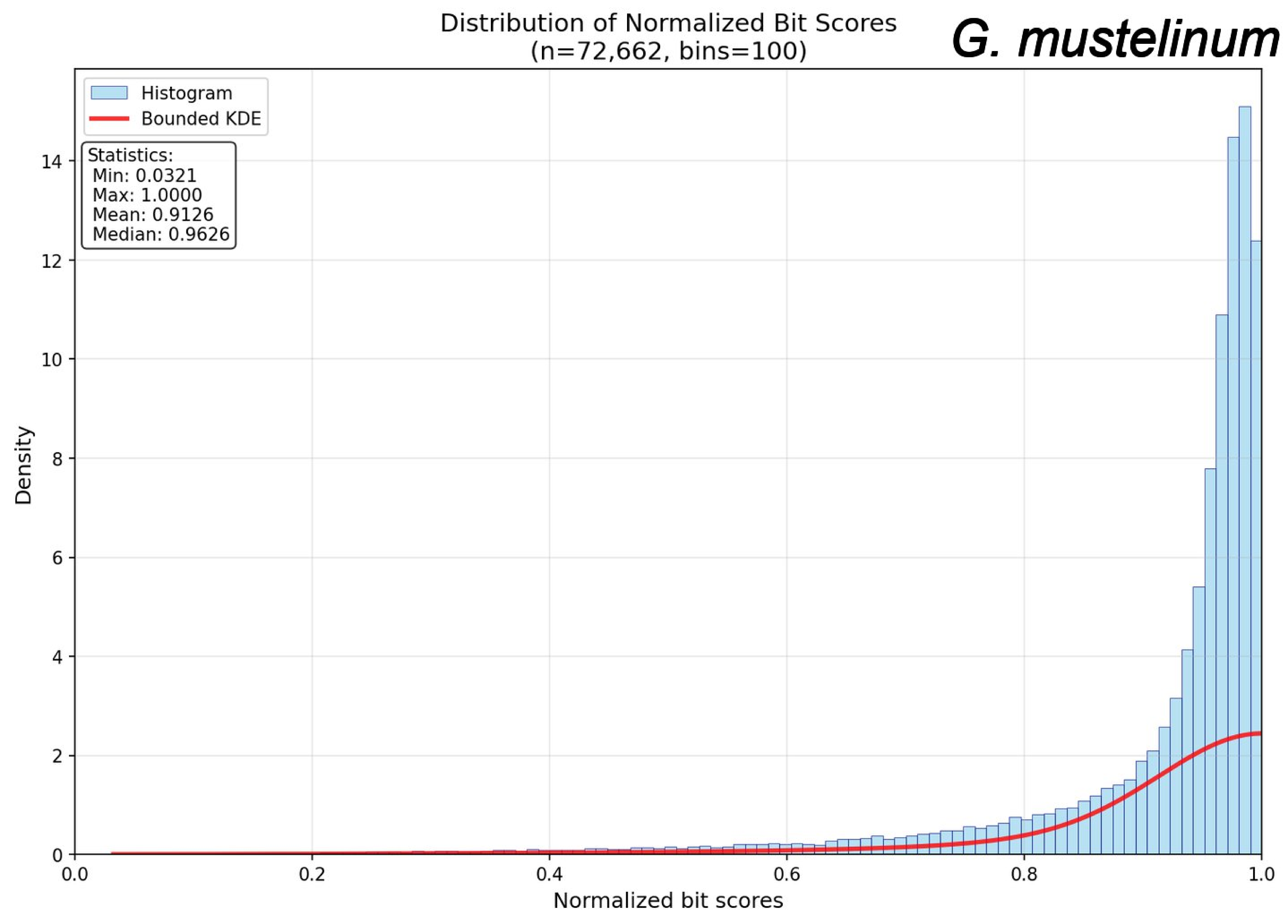

Distribution of Normalized Bit Scores
(n=72,662, bins=100)
G. mustelinum
Histogram
Bounded KDE
Statistics:
Min: 0.0321
Max: 1.0000
Mean: 0.9126
Median: 0.9626
Density
Normalized bit scores
0
2
4
6
8
10
12
14
0.0
0.2
0.4
0.6
0.8
1.0

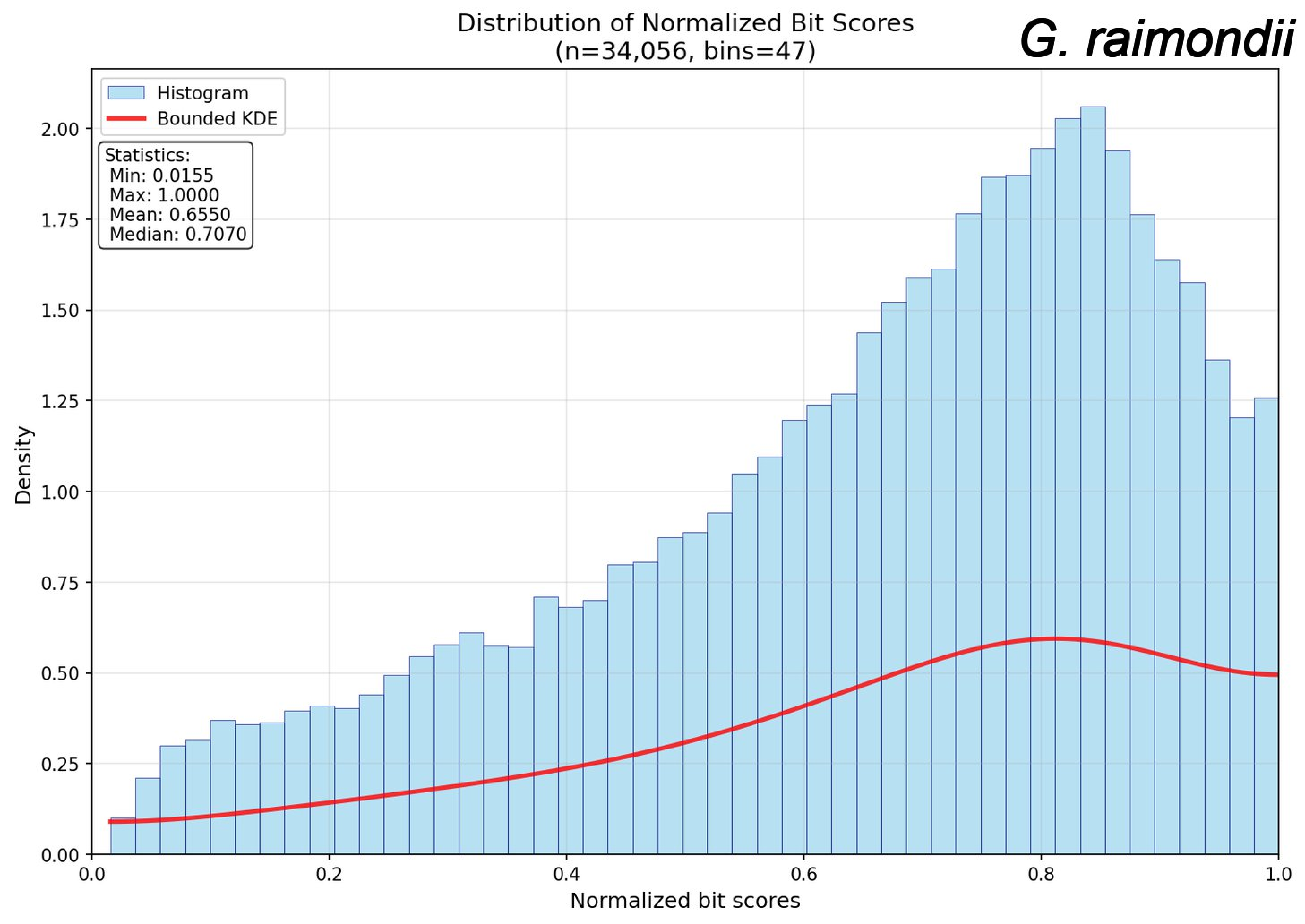

Distribution of Normalized Bit Scores
(n=34,056, bins=47)
G. raimondii
Histogram
Bounded KDE
Statistics:
Min: 0.0155
Max: 1.0000
Mean: 0.6550
Median: 0.7070
Density
Normalized bit scores
0.00
0.25
0.50
0.75
1.00
1.25
1.50
1.75
2.00
0.0
0.2
0.4
0.6
0.8
1.0

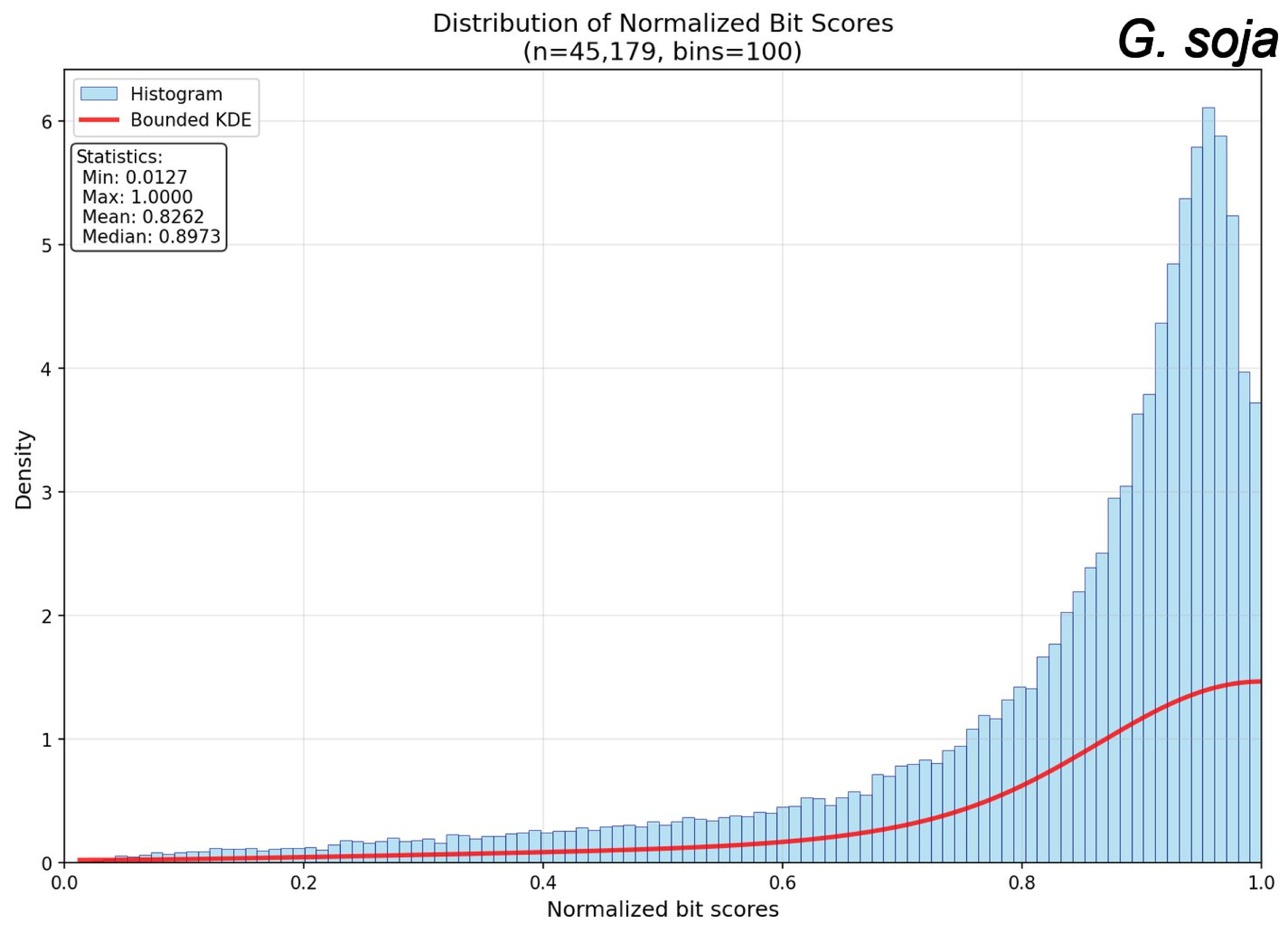

Distribution of Normalized Bit Scores
(n=45,179, bins=100)
G. soja
Histogram
Bounded KDE
Statistics:
Min: 0.0127
Max: 1.0000
Mean: 0.8262
Median: 0.8973
Density
Normalized bit scores
0
1
2
3
4
5
6
0.0
0.2
0.4
0.6
0.8
1.0

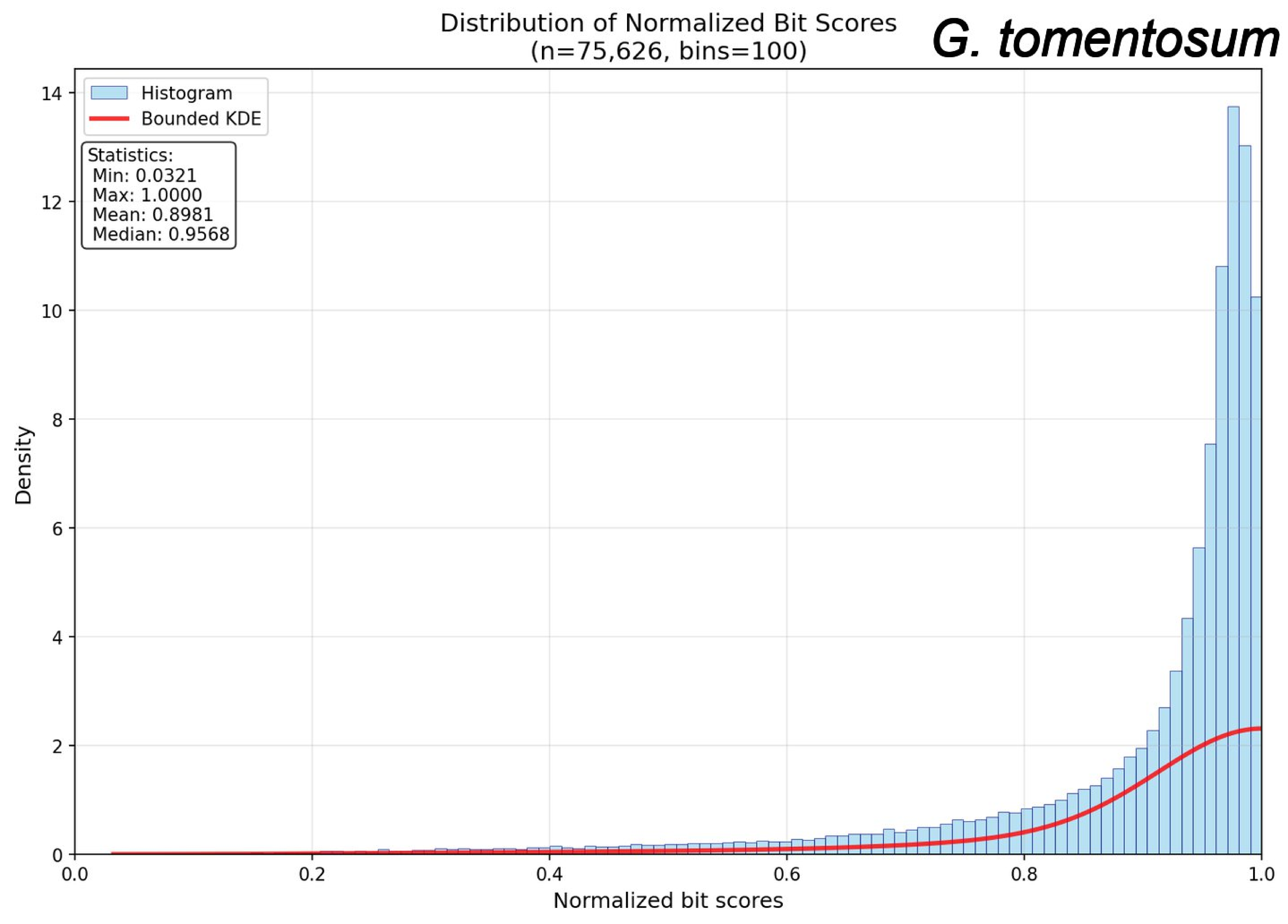

Distribution of Normalized Bit Scores
(n=75,626, bins=100)
G. tomentosum
Histogram
Bounded KDE
Statistics:
Min: 0.0321
Max: 1.0000
Mean: 0.8981
Median: 0.9568
Density
Normalized bit scores
0
2
4
6
8
10
12
14
0.0
0.2
0.4
0.6
0.8
1.0

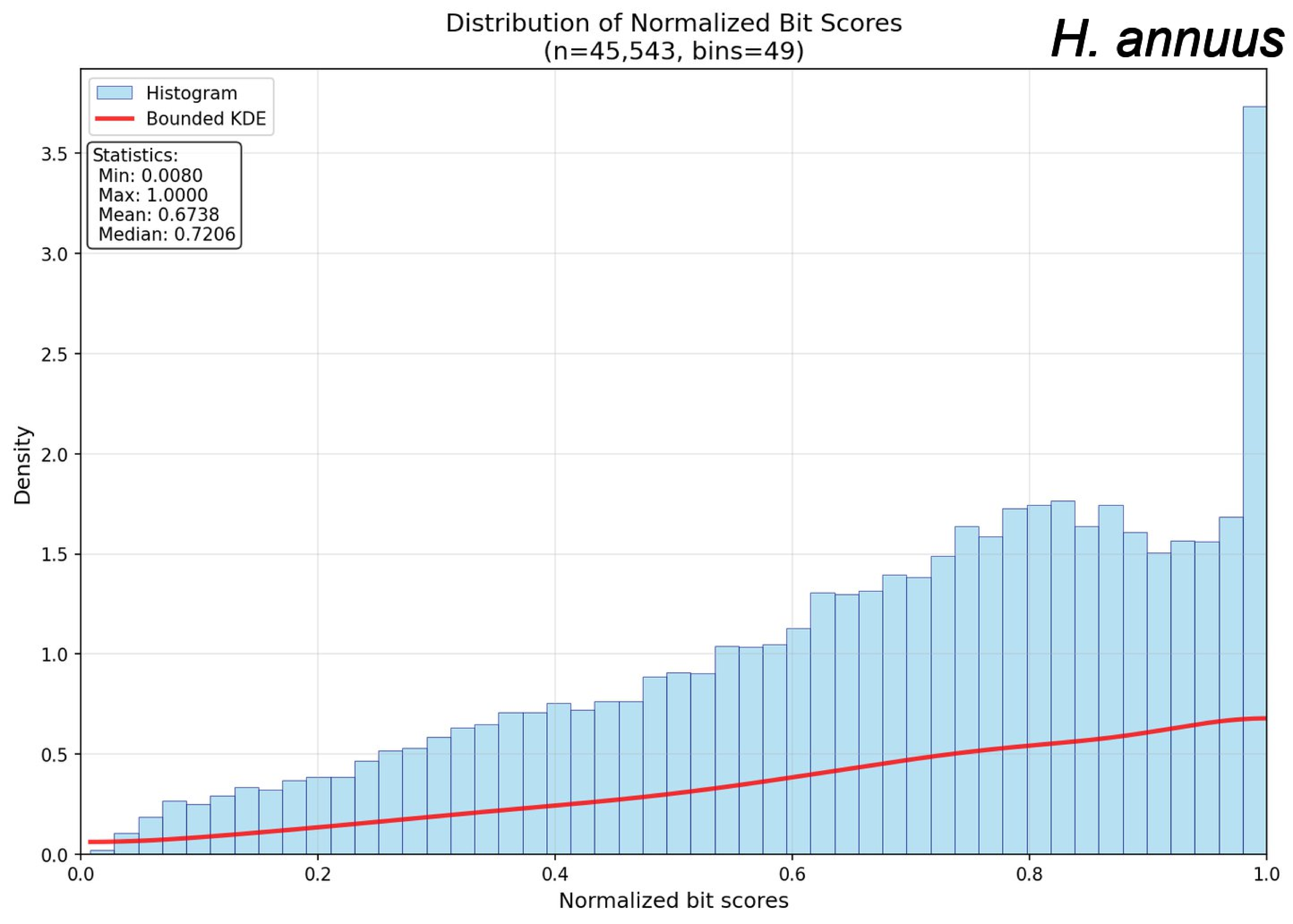
Distribution of Normalized Bit Scores
(n=45,543, bins=49)
H. annuus
Histogram
Bounded KDE
Statistics:
Min: 0.0080
Max: 1.0000
Mean: 0.6738
Median: 0.7206
Density
Normalized bit scores
0.0
0.5
1.0
1.5
2.0
2.5
3.0
3.5
0.0
0.2
0.4
0.6
0.8
1.0

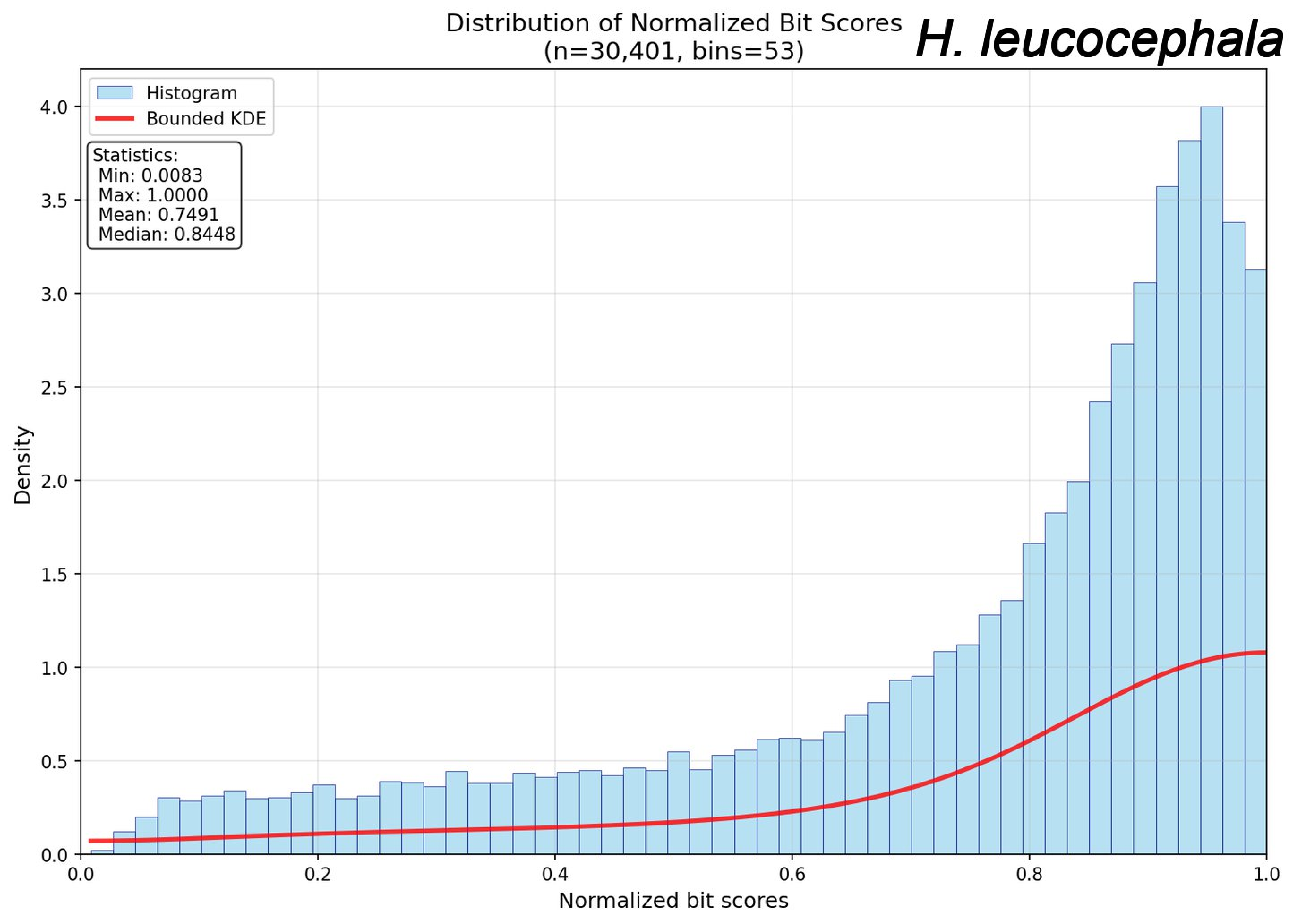
Distribution of Normalized Bit Scores
(n=30,401, bins=53)
H. leucocephala
Histogram
Bounded KDE
Statistics:
Min: 0.0083
Max: 1.0000
Mean: 0.7491
Median: 0.8448
Density
Normalized bit scores
0.0
0.5
1.0
1.5
2.0
2.5
3.0
3.5
4.0
0.0
0.2
0.4
0.6
0.8
1.0

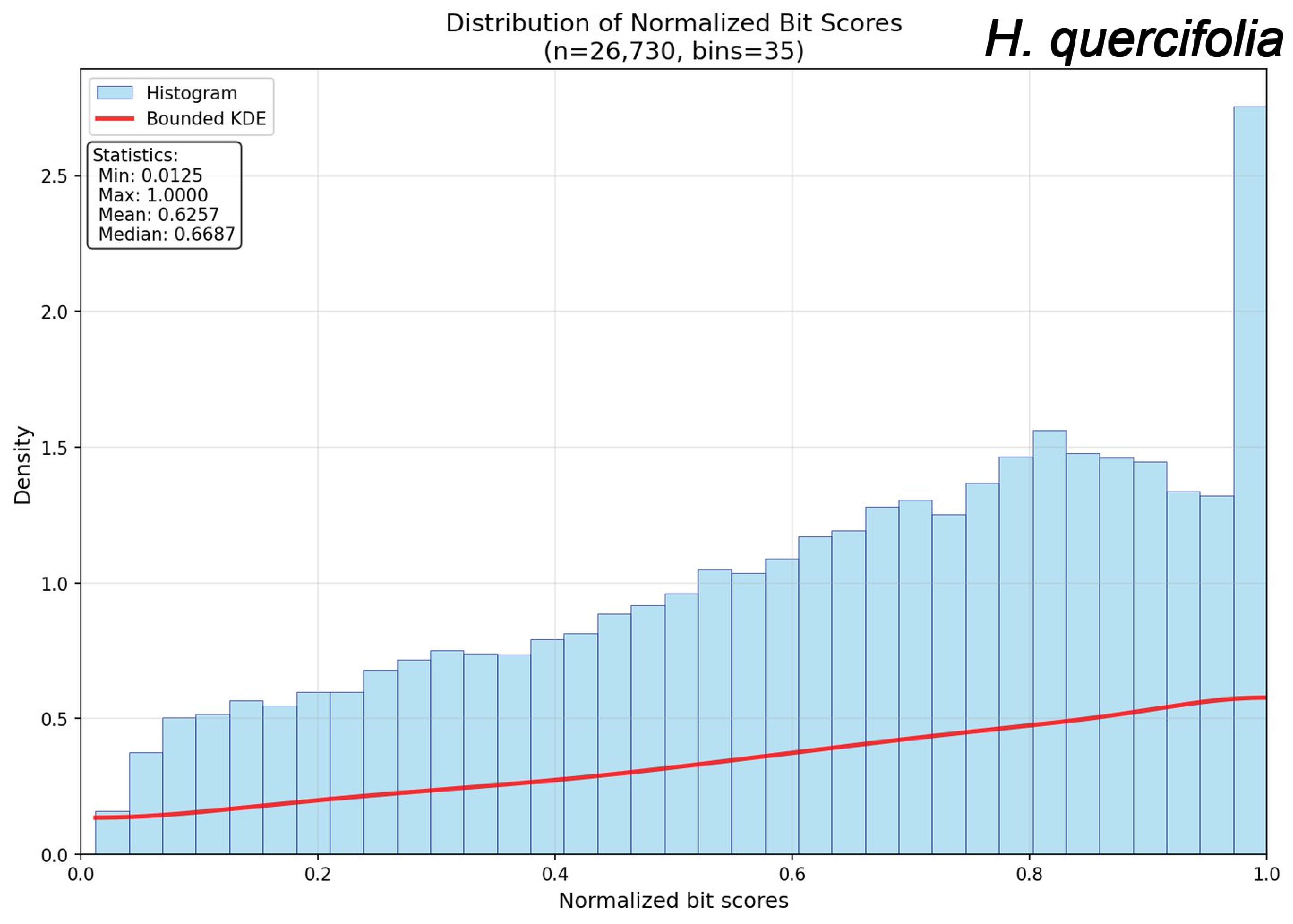
Distribution of Normalized Bit Scores
(n=26,730, bins=35)
H. quercifolia
Histogram
Bounded KDE
Statistics:
Min: 0.0125
Max: 1.0000
Mean: 0.6257
Median: 0.6687
Density
Normalized bit scores
0.0
0.5
1.0
1.5
2.0
2.5
0.0
0.2
0.4
0.6
0.8
1.0

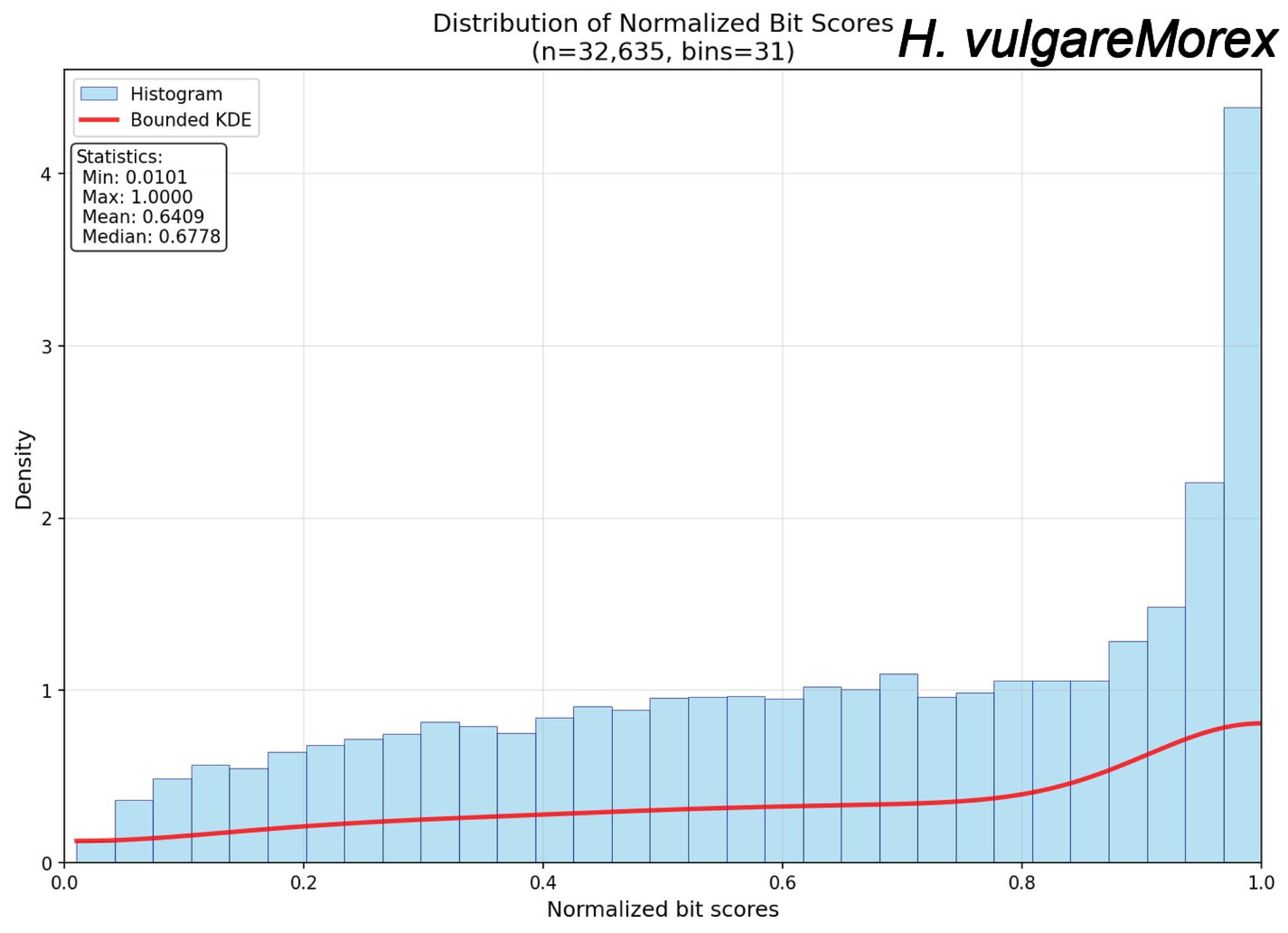

Distribution of Normalized Bit Scores
(n=32,635, bins=31)
H. vulgareMorex
Histogram
Bounded KDE
Statistics:
Min: 0.0101
Max: 1.0000
Mean: 0.6409
Median: 0.6778
Density
Normalized bit scores
0
1
2
3
4
0.0
0.2
0.4
0.6
0.8
1.0

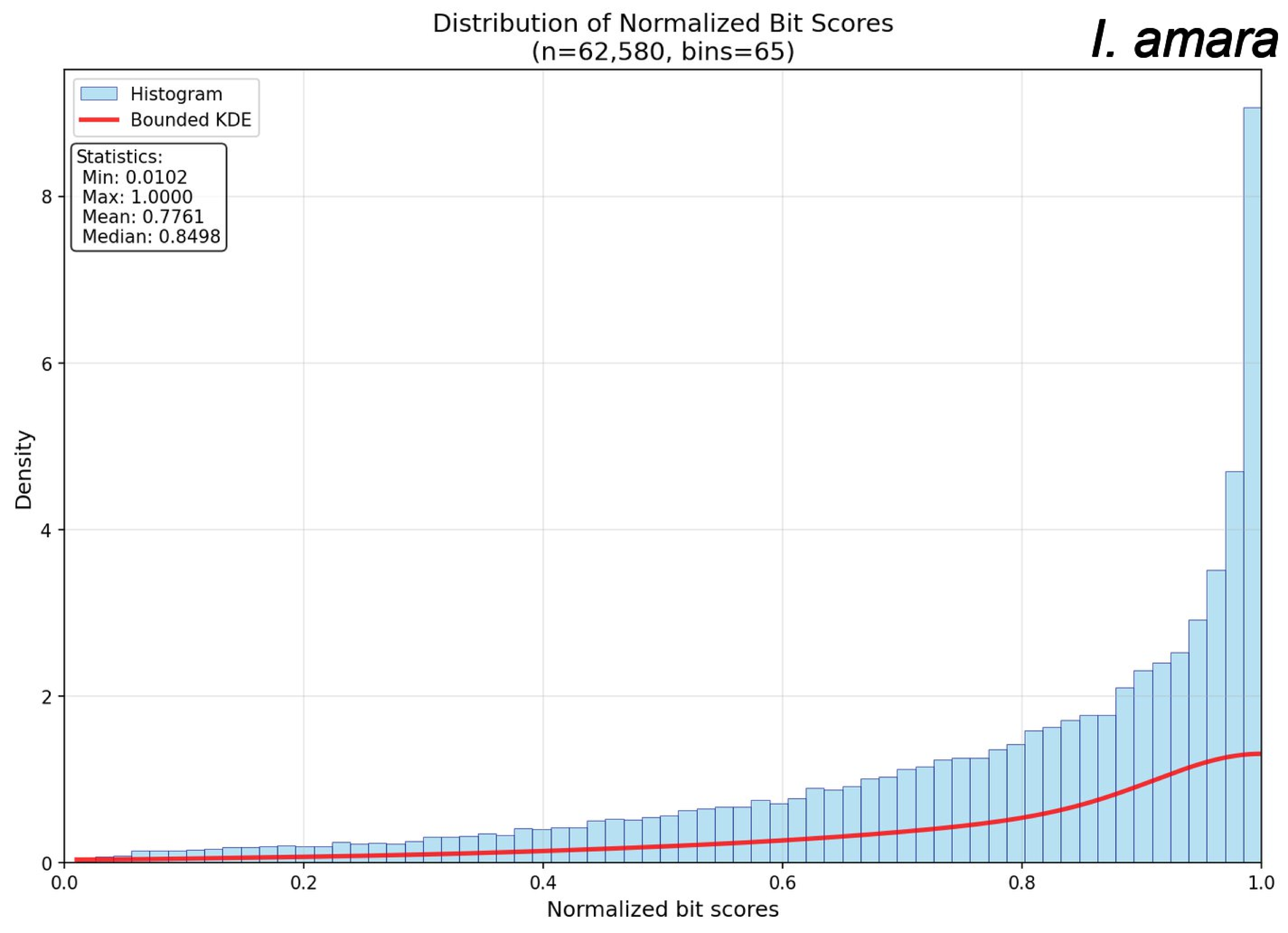

Distribution of Normalized Bit Scores
(n=62,580, bins=65)
I. amara
Histogram
Bounded KDE
Statistics:
Min: 0.0102
Max: 1.0000
Mean: 0.7761
Median: 0.8498
Density
Normalized bit scores
0
2
4
6
8
0.0
0.2
0.4
0.6
0.8
1.0

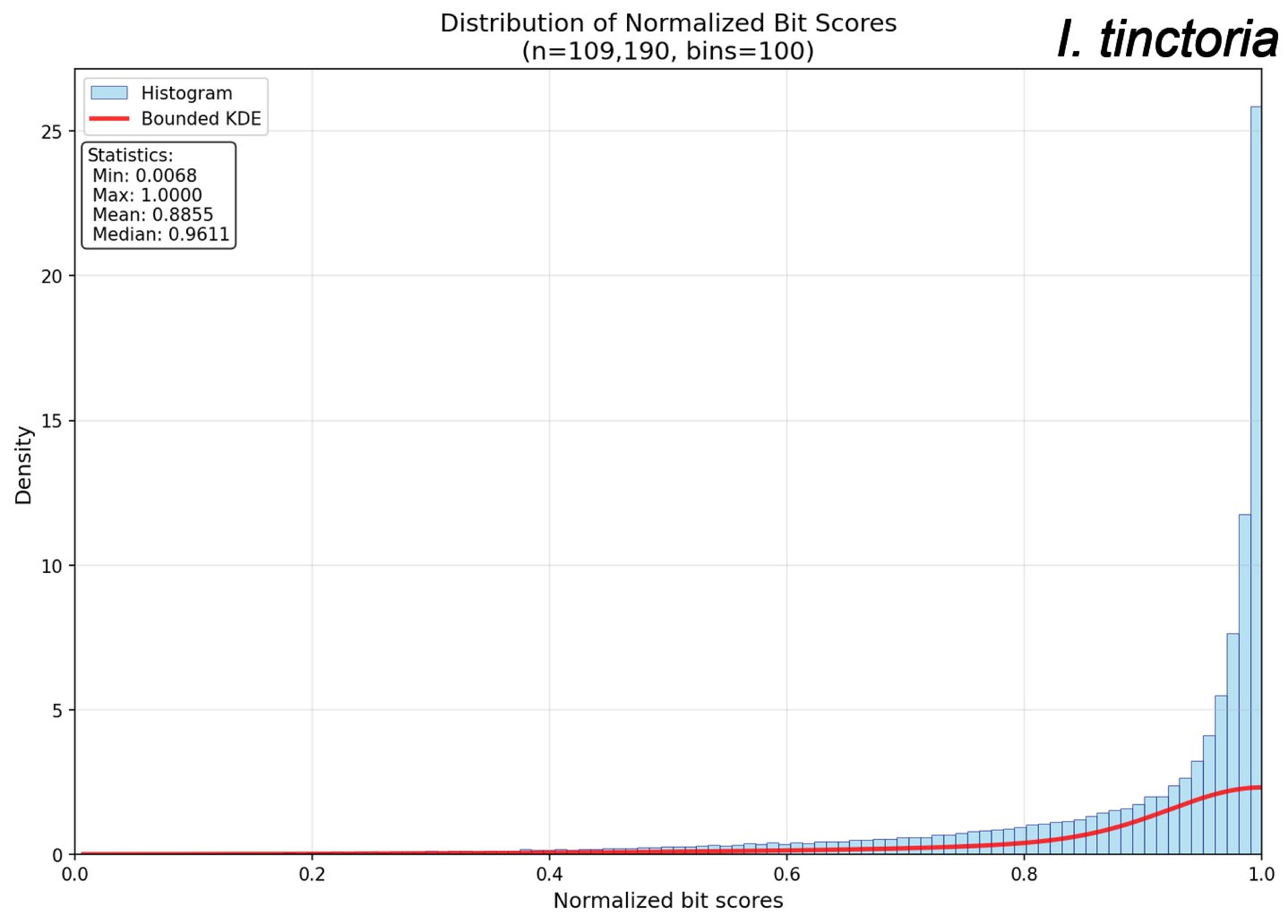
Distribution of Normalized Bit Scores
(n=109,190, bins=100)
I. tinctoria
Histogram
Bounded KDE
Statistics:
Min: 0.0068
Max: 1.0000
Mean: 0.8855
Median: 0.9611
Density
Normalized bit scores
0
5
10
15
20
25
0.0
0.2
0.4
0.6
0.8
1.0

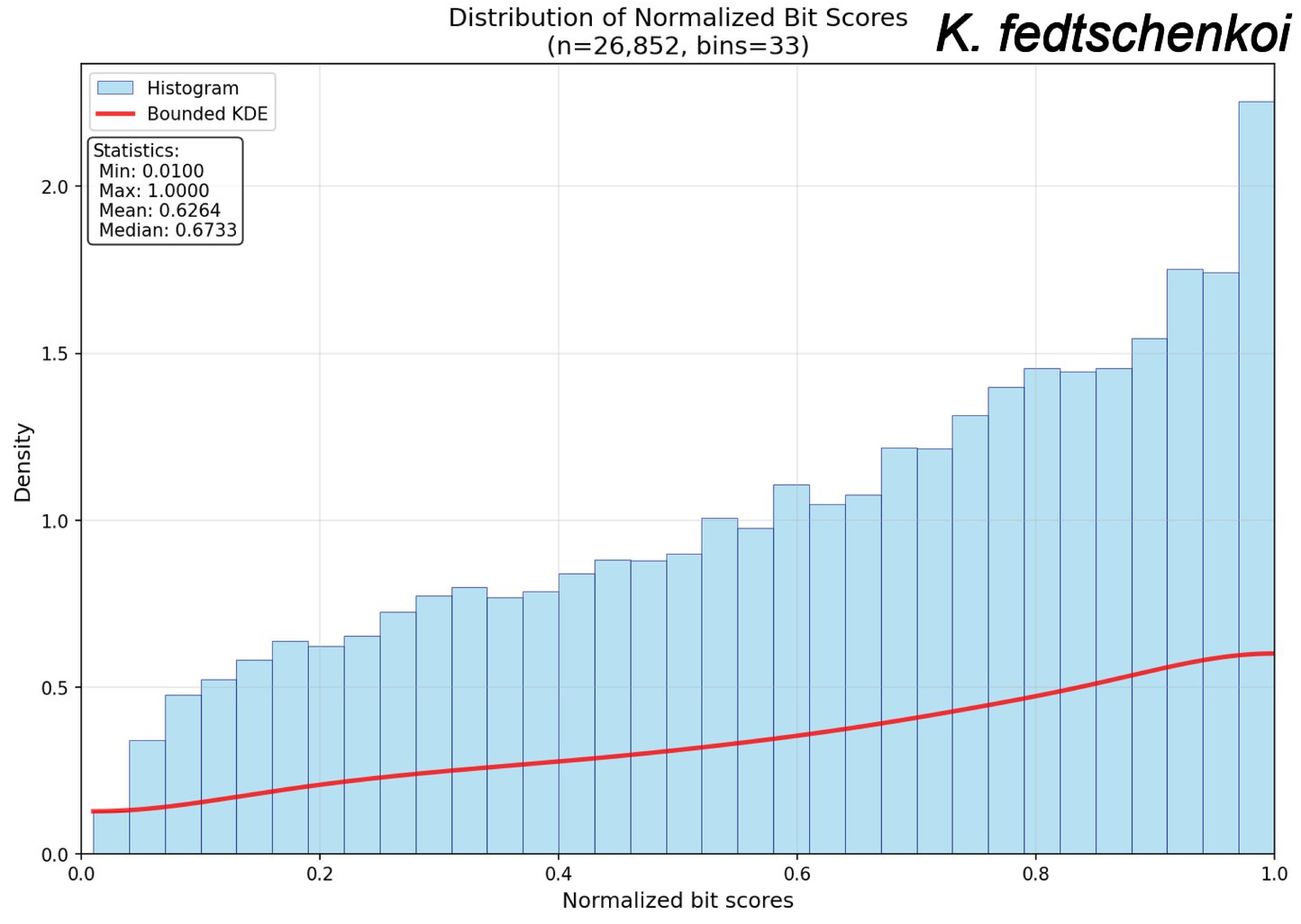
Distribution of Normalized Bit Scores
(n=26,852, bins=33)
K. fedtschenkoi
Histogram
Bounded KDE
Statistics:
Min: 0.0100
Max: 1.0000
Mean: 0.6264
Median: 0.6733
Density
Normalized bit scores
0.0
0.5
1.0
1.5
2.0
0.2
0.4
0.6
0.8
1.0

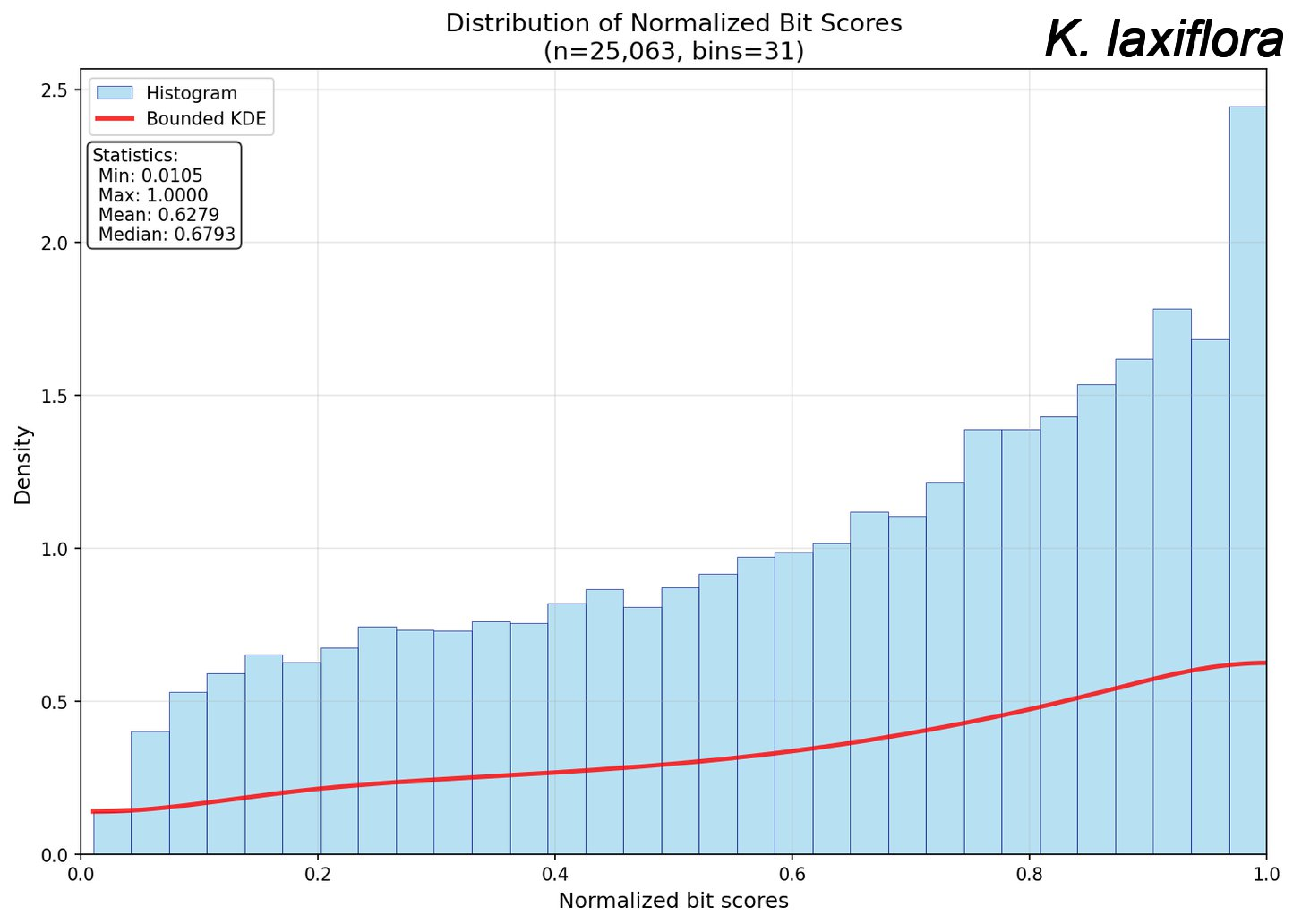
Distribution of Normalized Bit Scores
(n=25,063, bins=31)
K. laxiflora
Histogram
Bounded KDE
Statistics:
Min: 0.0105
Max: 1.0000
Mean: 0.6279
Median: 0.6793
Density
Normalized bit scores
0.0
0.5
1.0
1.5
2.0
2.5
0.0
0.2
0.4
0.6
0.8
1.0

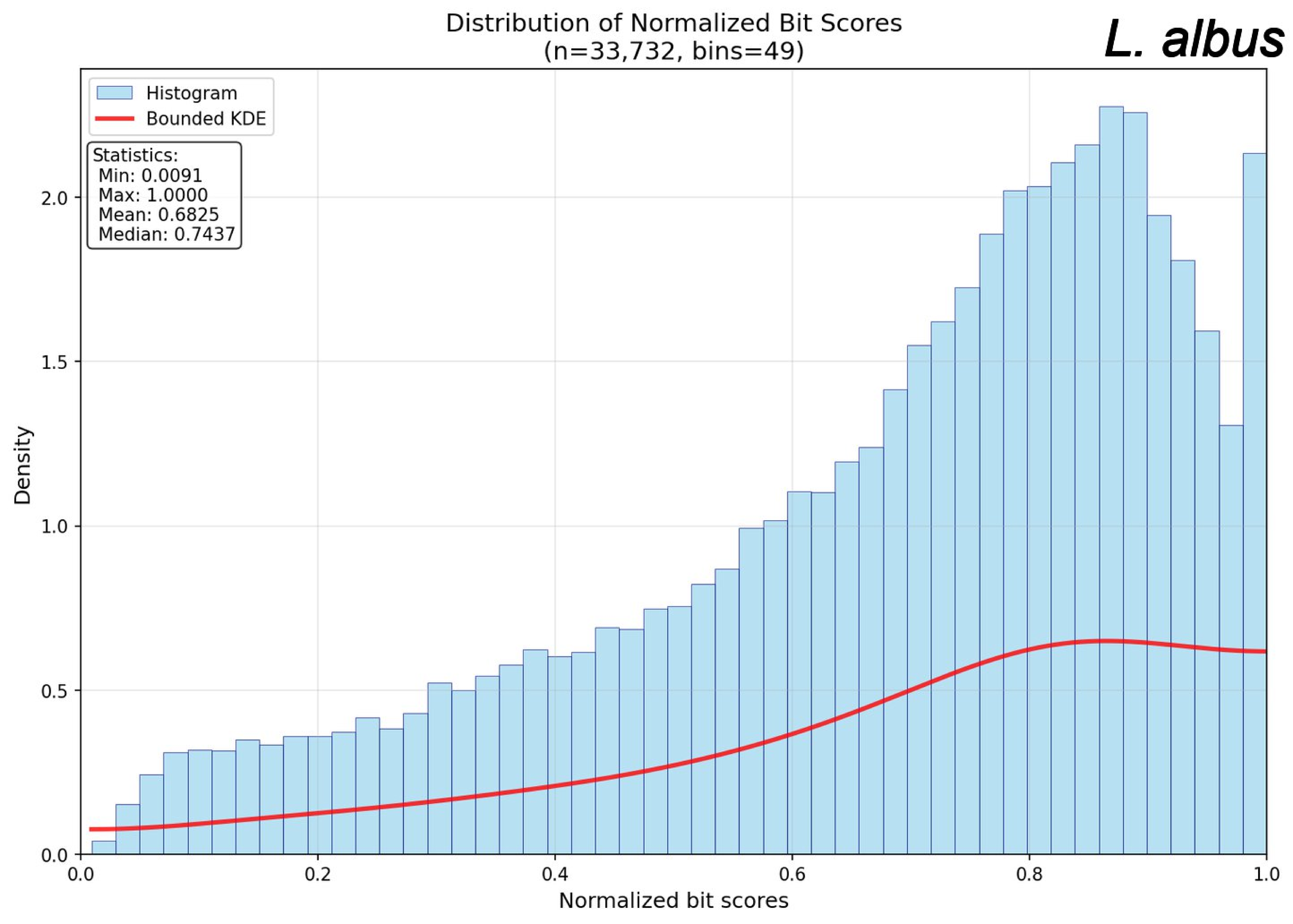

Distribution of Normalized Bit Scores
(n=33,732, bins=49)
L. albus
Histogram
Bounded KDE
Statistics:
Min: 0.0091
Max: 1.0000
Mean: 0.6825
Median: 0.7437
Density
Normalized bit scores
0.0
0.5
1.0
1.5
2.0
0.0
0.2
0.4
0.6
0.8
1.0

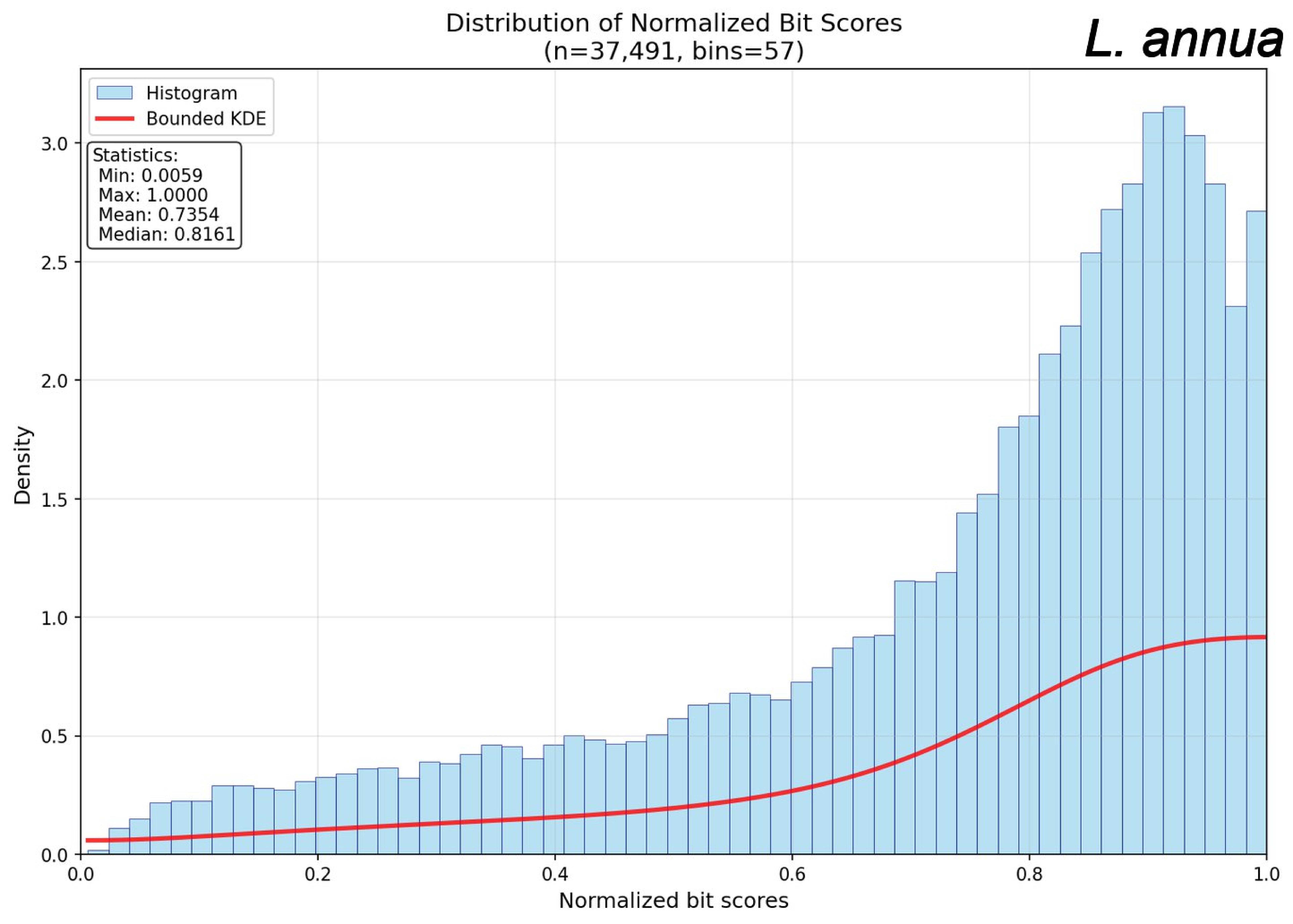
Distribution of Normalized Bit Scores
(n=37,491, bins=57)
L. annua
Histogram
Bounded KDE
Statistics:
Min: 0.0059
Max: 1.0000
Mean: 0.7354
Median: 0.8161
Density
Normalized bit scores
0.0
0.5
1.0
1.5
2.0
2.5
3.0
0.0
0.2
0.4
0.6
0.8
1.0

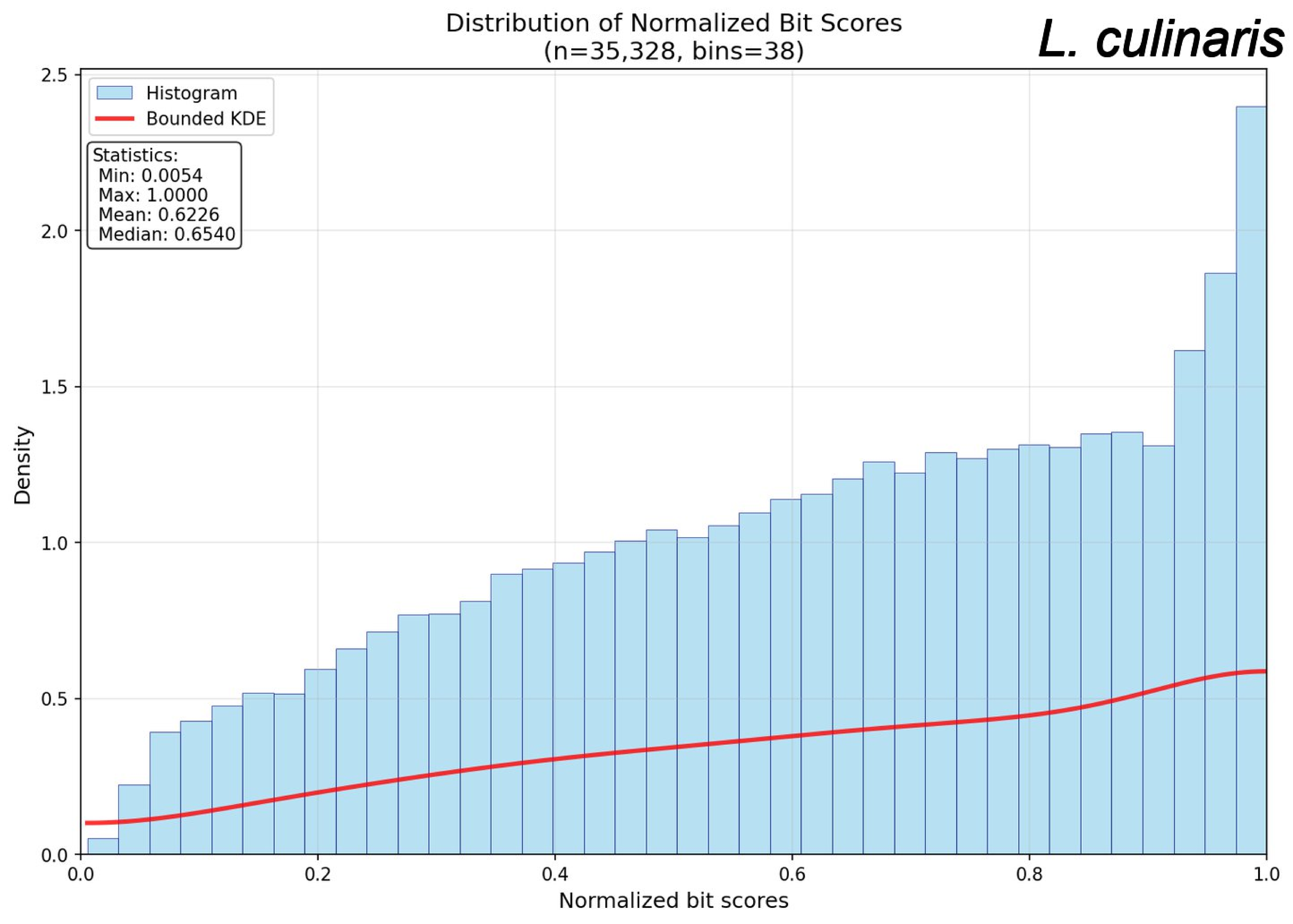

Distribution of Normalized Bit Scores
(n=35,328, bins=38)
L. culinaris
Histogram
Bounded KDE
Statistics:
Min: 0.0054
Max: 1.0000
Mean: 0.6226
Median: 0.6540
Density
Normalized bit scores
0.0
0.5
1.0
1.5
2.0
2.5
0.0
0.2
0.4
0.6
0.8
1.0

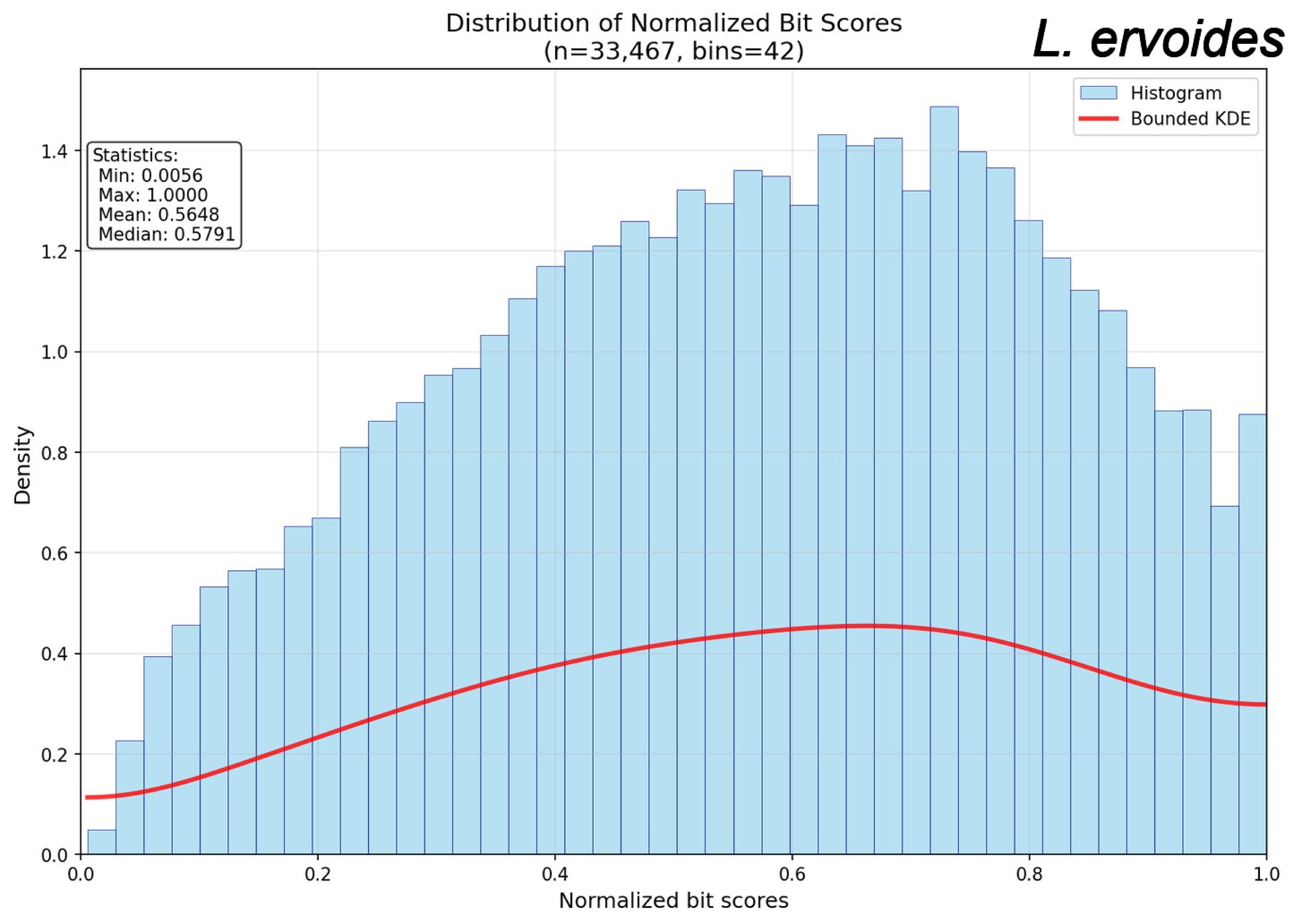

Distribution of Normalized Bit Scores
(n=33,467, bins=42)
L. ervoides
Histogram
Bounded KDE
Statistics:
Min: 0.0056
Max: 1.0000
Mean: 0.5648
Median: 0.5791
Density
Normalized bit scores
0.0
0.2
0.4
0.6
0.8
1.0
1.2
1.4

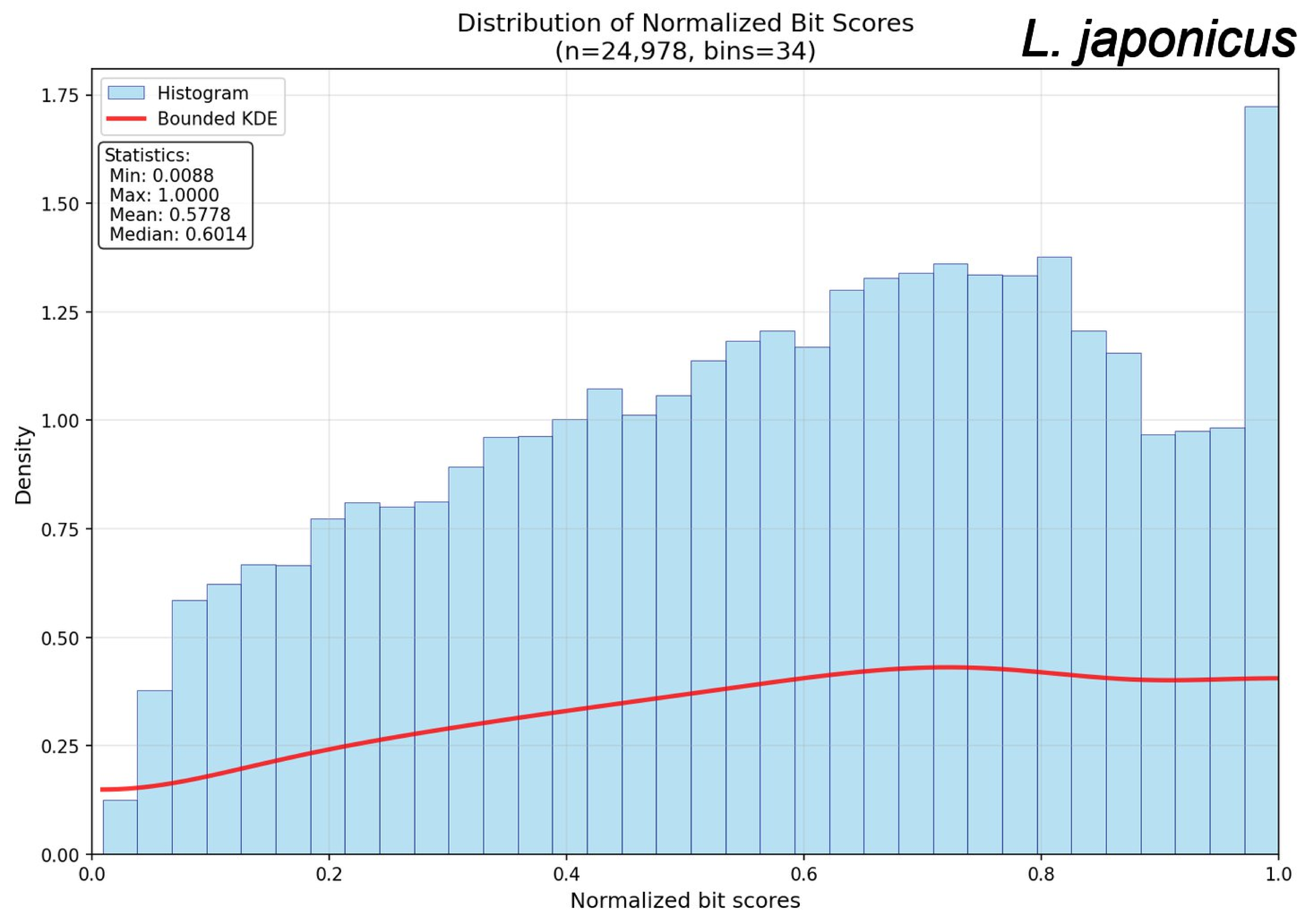
Distribution of Normalized Bit Scores
(n=24,978, bins=34)
L. japonicus
Histogram
Bounded KDE
Statistics:
Min: 0.0088
Max: 1.0000
Mean: 0.5778
Median: 0.6014
Density
Normalized bit scores
0.00
0.25
0.50
0.75
1.00
1.25
1.50
1.75
0.0
0.2
0.4
0.6
0.8
1.0

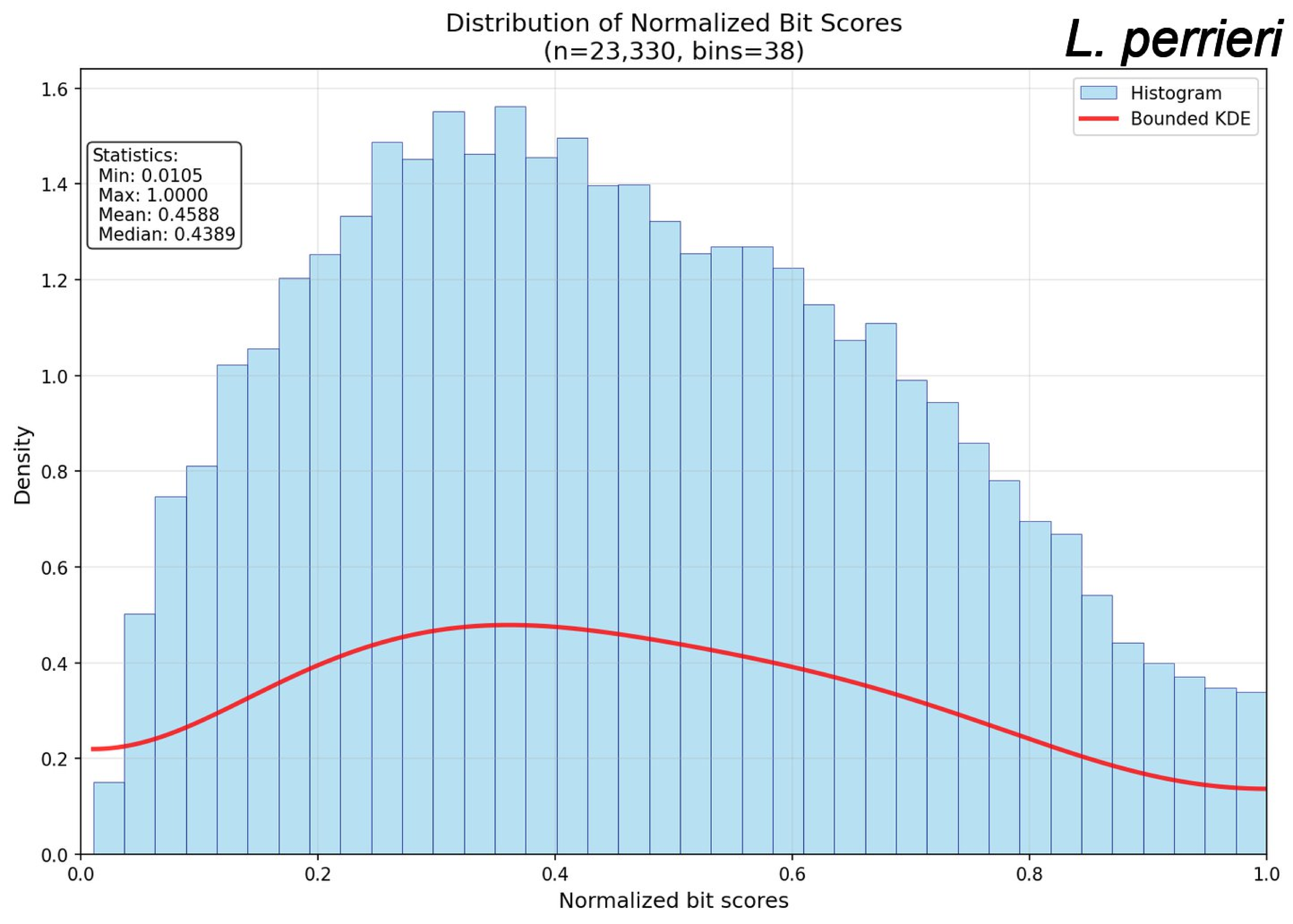
Distribution of Normalized Bit Scores
(n=23,330, bins=38)
L. perrieri
Histogram
Bounded KDE
Statistics:
Min: 0.0105
Max: 1.0000
Mean: 0.4588
Median: 0.4389
Density
Normalized bit scores
0.0
0.2
0.4
0.6
0.8
1.0
1.2
1.4
1.6

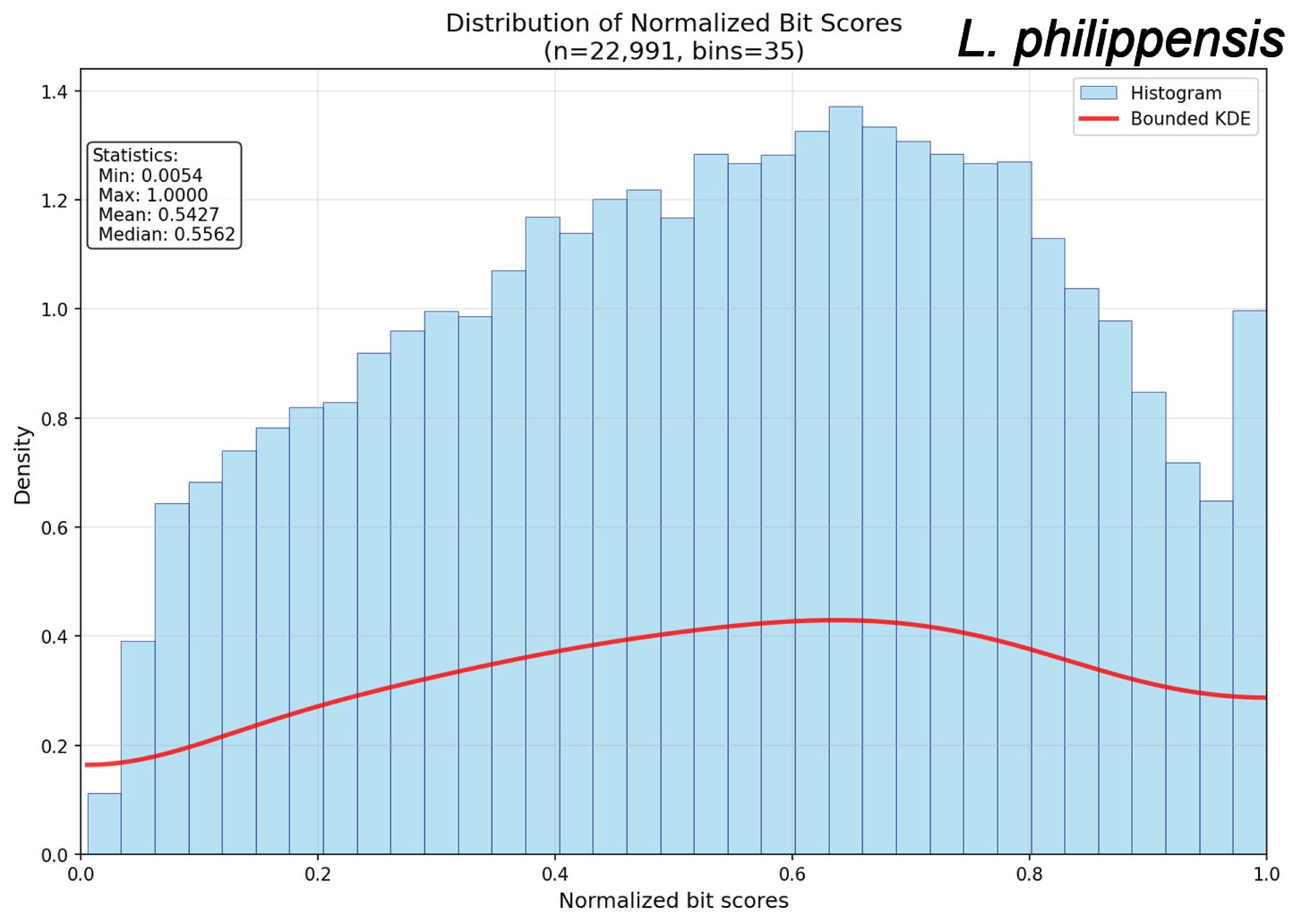
Distribution of Normalized Bit Scores
(n=22,991, bins=35)
L. philippensis
Histogram
Bounded KDE
Statistics:
Min: 0.0054
Max: 1.0000
Mean: 0.5427
Median: 0.5562
Density
Normalized bit scores
0.0
0.2
0.4
0.6
0.8
1.0
1.2
1.4

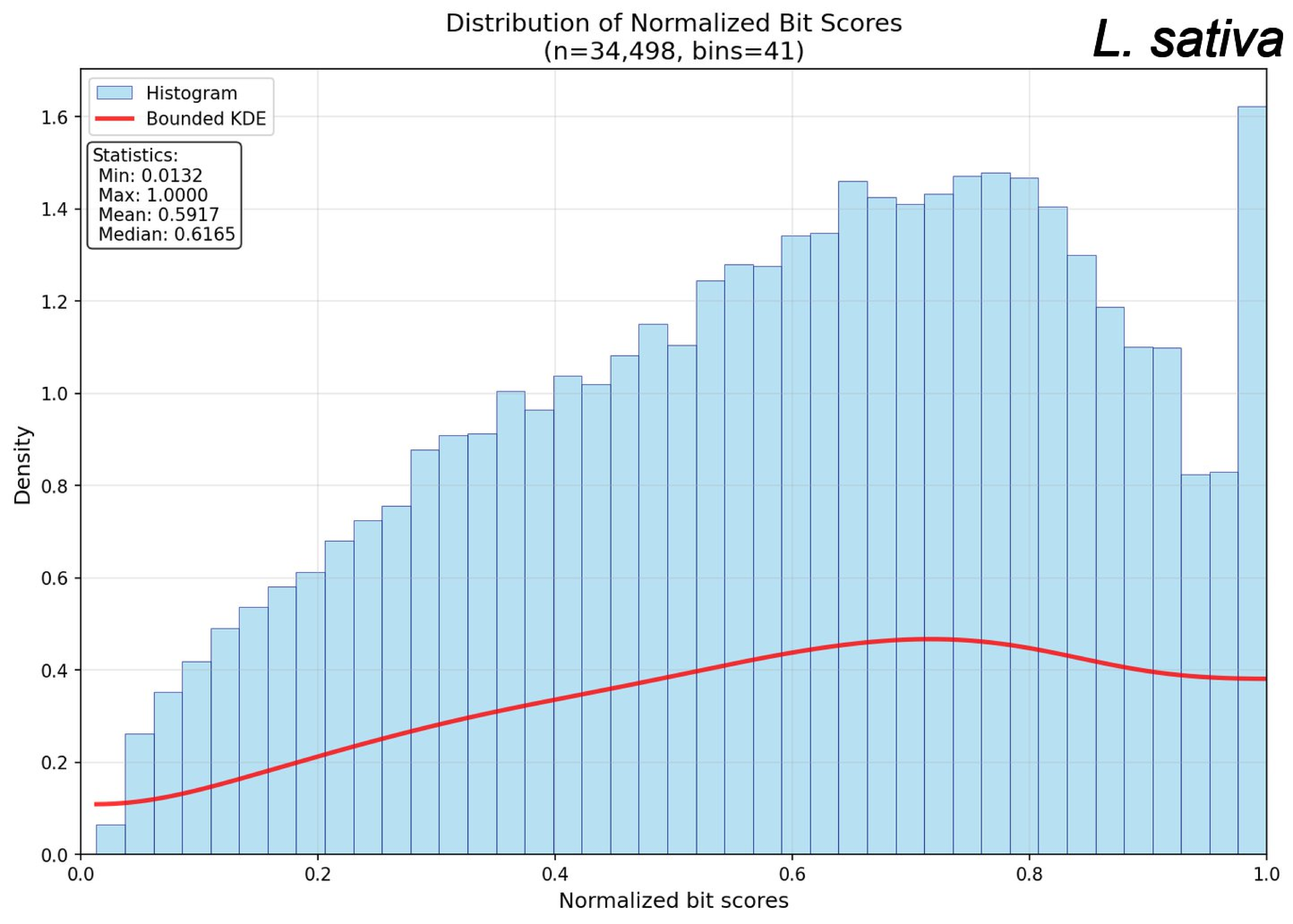
Distribution of Normalized Bit Scores
(n=34,498, bins=41)
L. sativa
Histogram
Bounded KDE
Statistics:
Min: 0.0132
Max: 1.0000
Mean: 0.5917
Median: 0.6165
Density
Normalized bit scores
0.0
0.2
0.4
0.6
0.8
1.0
1.2
1.4
1.6

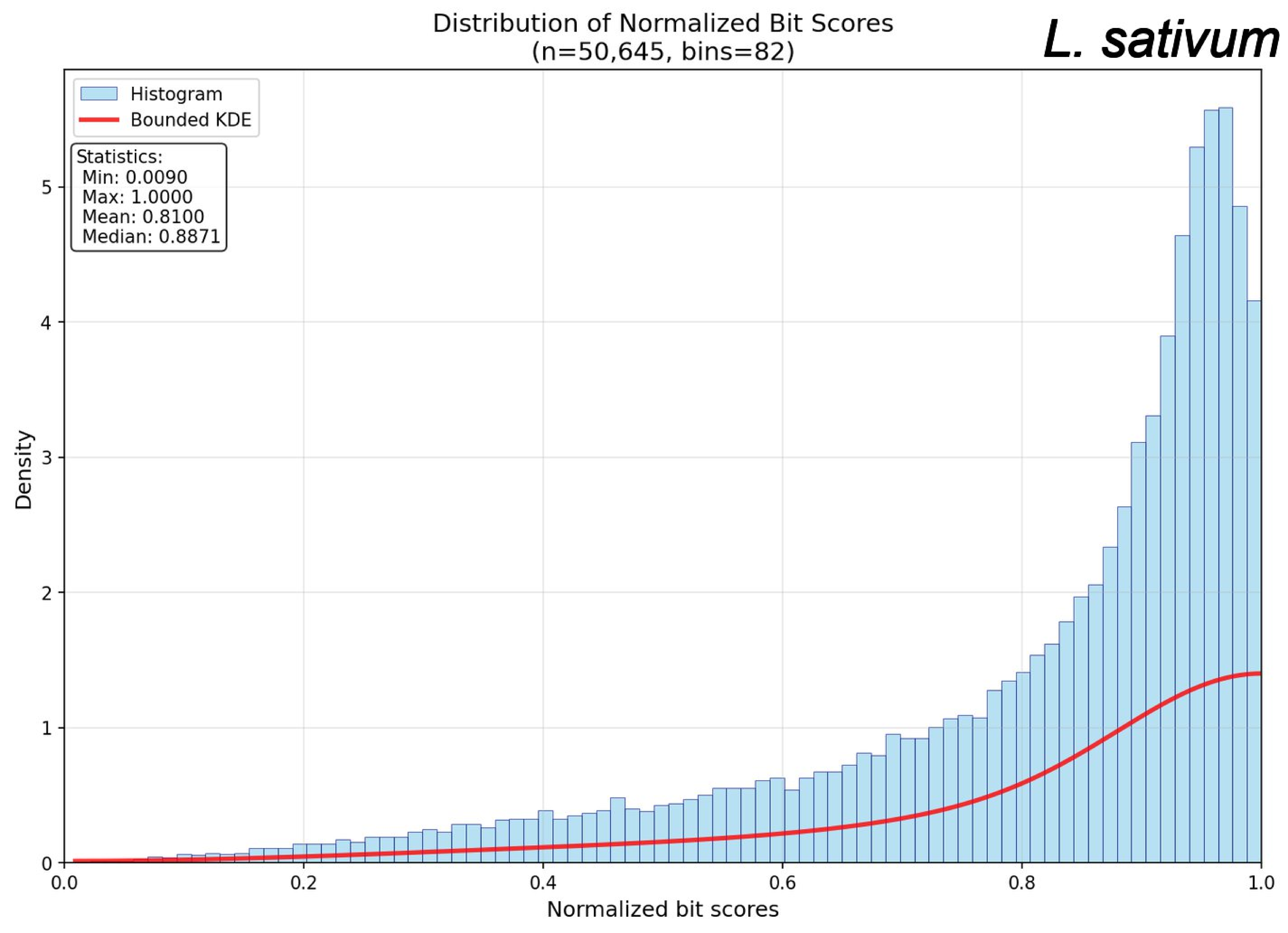

Distribution of Normalized Bit Scores
(n=50,645, bins=82)
L. sativum
Histogram
Bounded KDE
Statistics:
Min: 0.0090
Max: 1.0000
Mean: 0.8100
Median: 0.8871
Density
Normalized bit scores
0
1
2
3
4
5
0.0
0.2
0.4
0.6
0.8
1.0

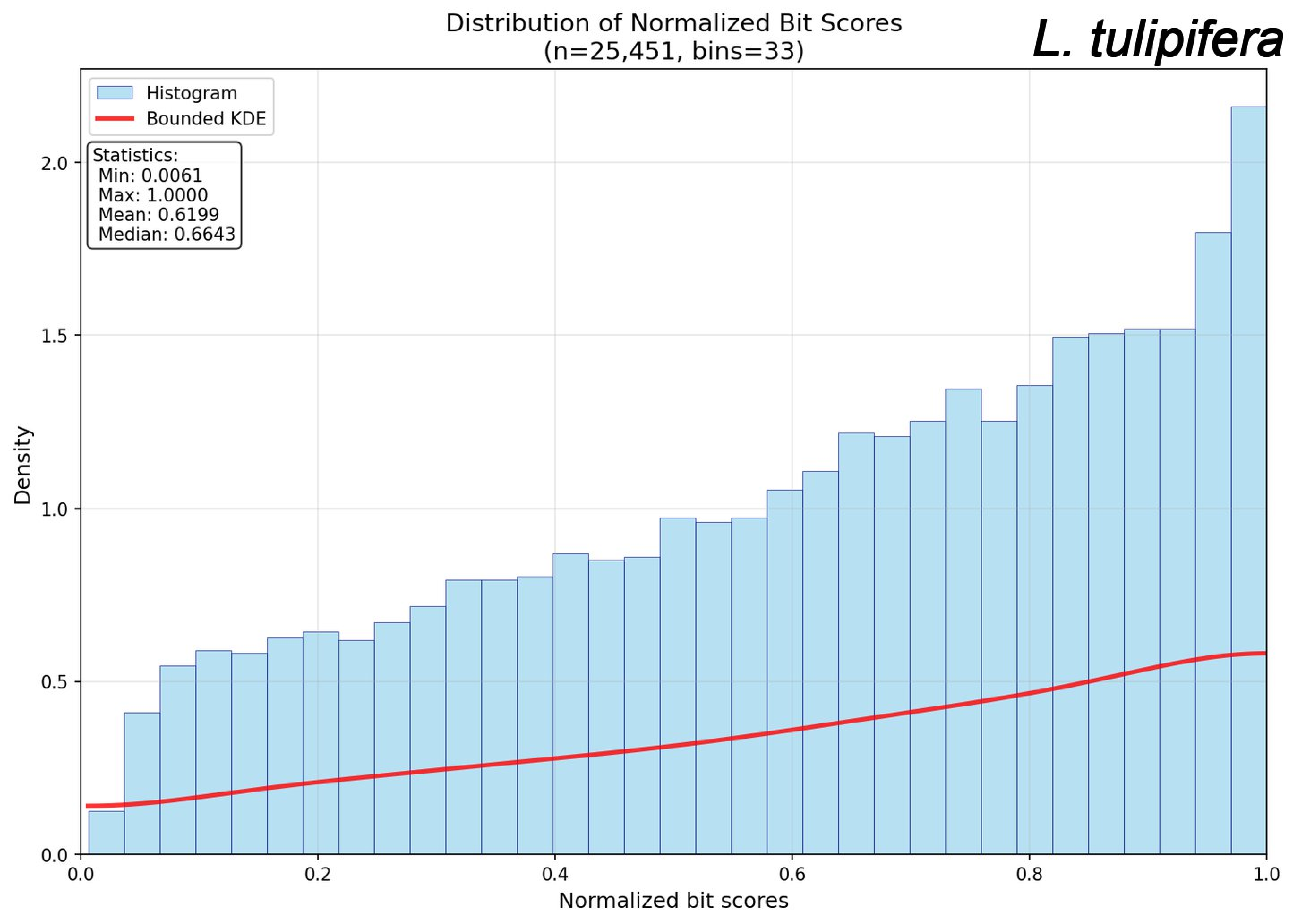

Distribution of Normalized Bit Scores
(n=25,451, bins=33)
L. tulipifera
Histogram
Bounded KDE
Statistics:
Min: 0.0061
Max: 1.0000
Mean: 0.6199
Median: 0.6643
Density
Normalized bit scores
0.0
0.5
1.0
1.5
2.0
0.2
0.4
0.6
0.8
1.0

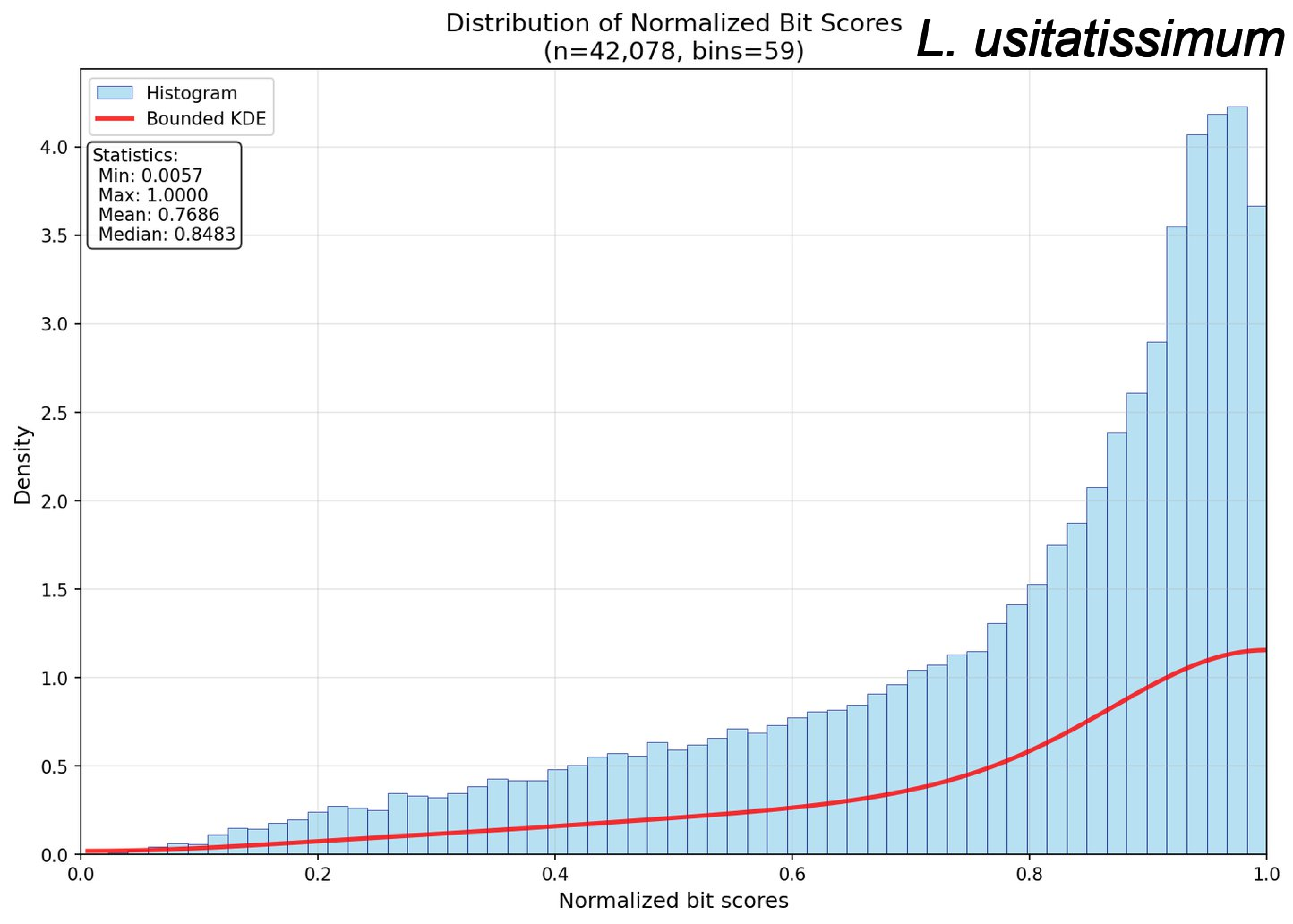

Distribution of Normalized Bit Scores
(n=42,078, bins=59)
L. usitatissimum
Histogram
Bounded KDE
Statistics:
Min: 0.0057
Max: 1.0000
Mean: 0.7686
Median: 0.8483
Density
Normalized bit scores
0.0
0.5
1.0
1.5
2.0
2.5
3.0
3.5
4.0
0.2
0.4
0.6
0.8

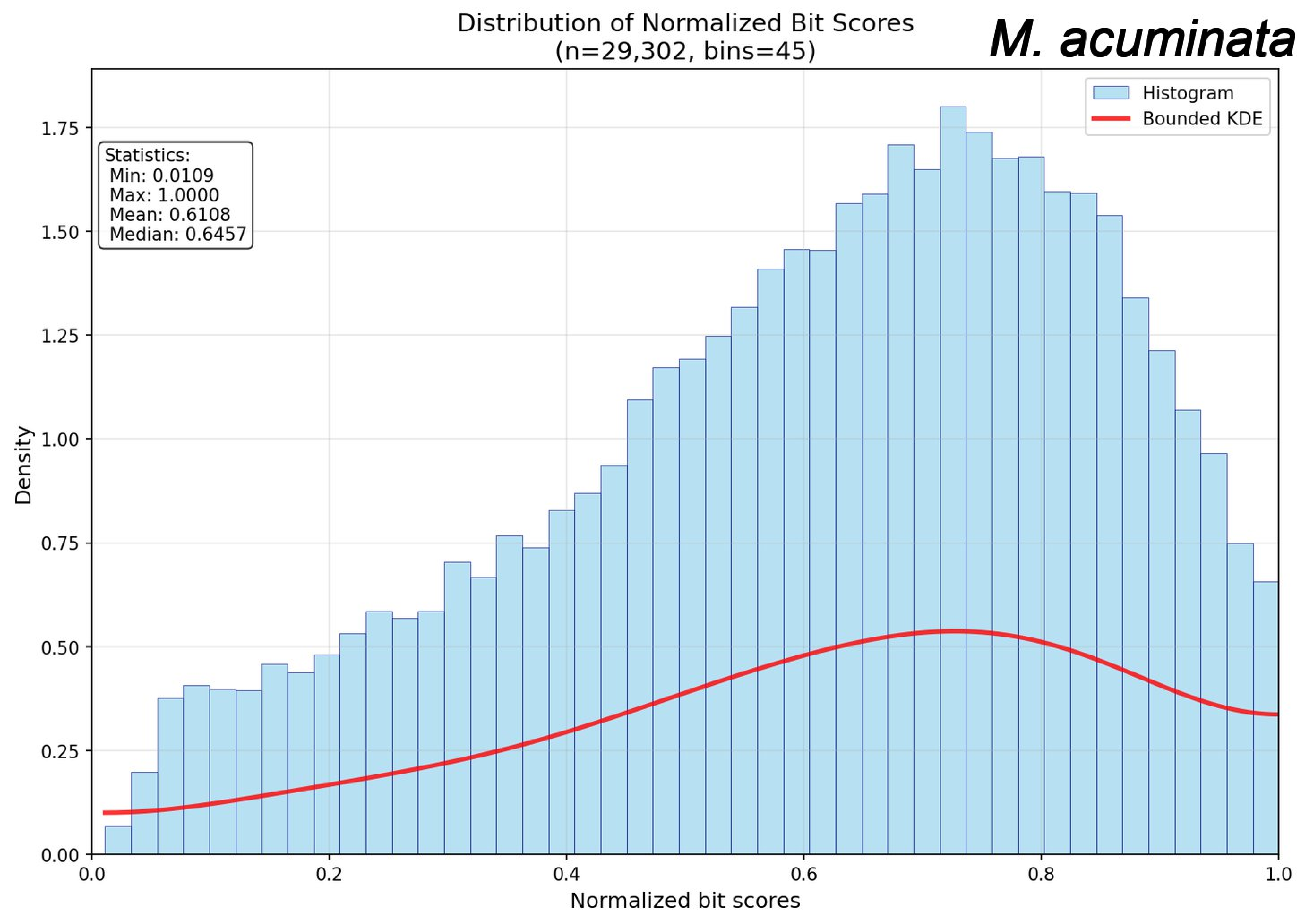
Distribution of Normalized Bit Scores
(n=29,302, bins=45)
M. acuminata
Histogram
Bounded KDE
Statistics:
Min: 0.0109
Max: 1.0000
Mean: 0.6108
Median: 0.6457
Density
Normalized bit scores
0.00
0.25
0.50
0.75
1.00
1.25
1.50
1.75
0.0
0.2
0.4
0.6
0.8
1.0

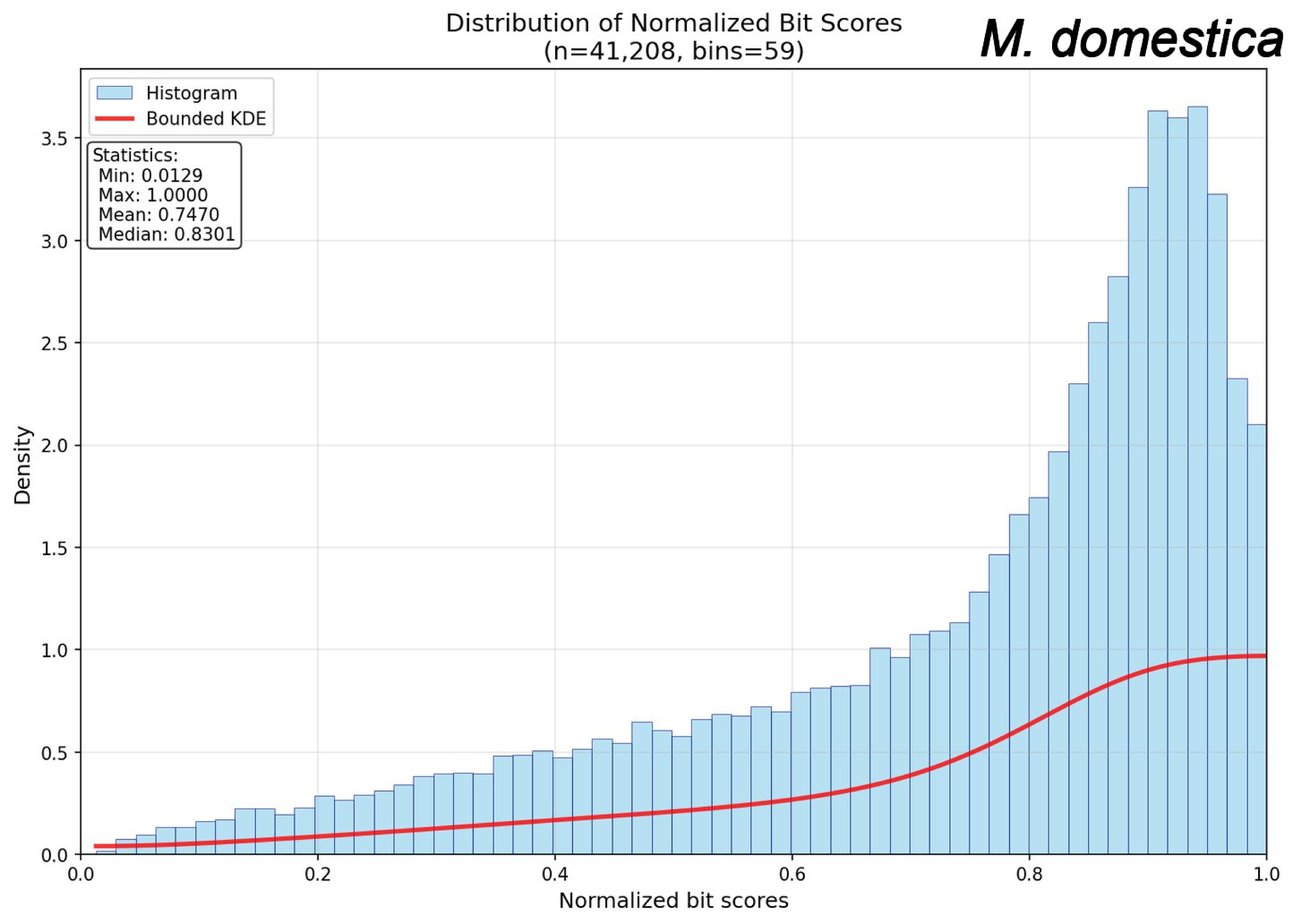
Distribution of Normalized Bit Scores
(n=41,208, bins=59)
M. domestica
Histogram
Bounded KDE
Statistics:
Min: 0.0129
Max: 1.0000
Mean: 0.7470
Median: 0.8301
Density
Normalized bit scores
0.0
0.5
1.0
1.5
2.0
2.5
3.0
3.5
0.2
0.4
0.6
0.8
1.0

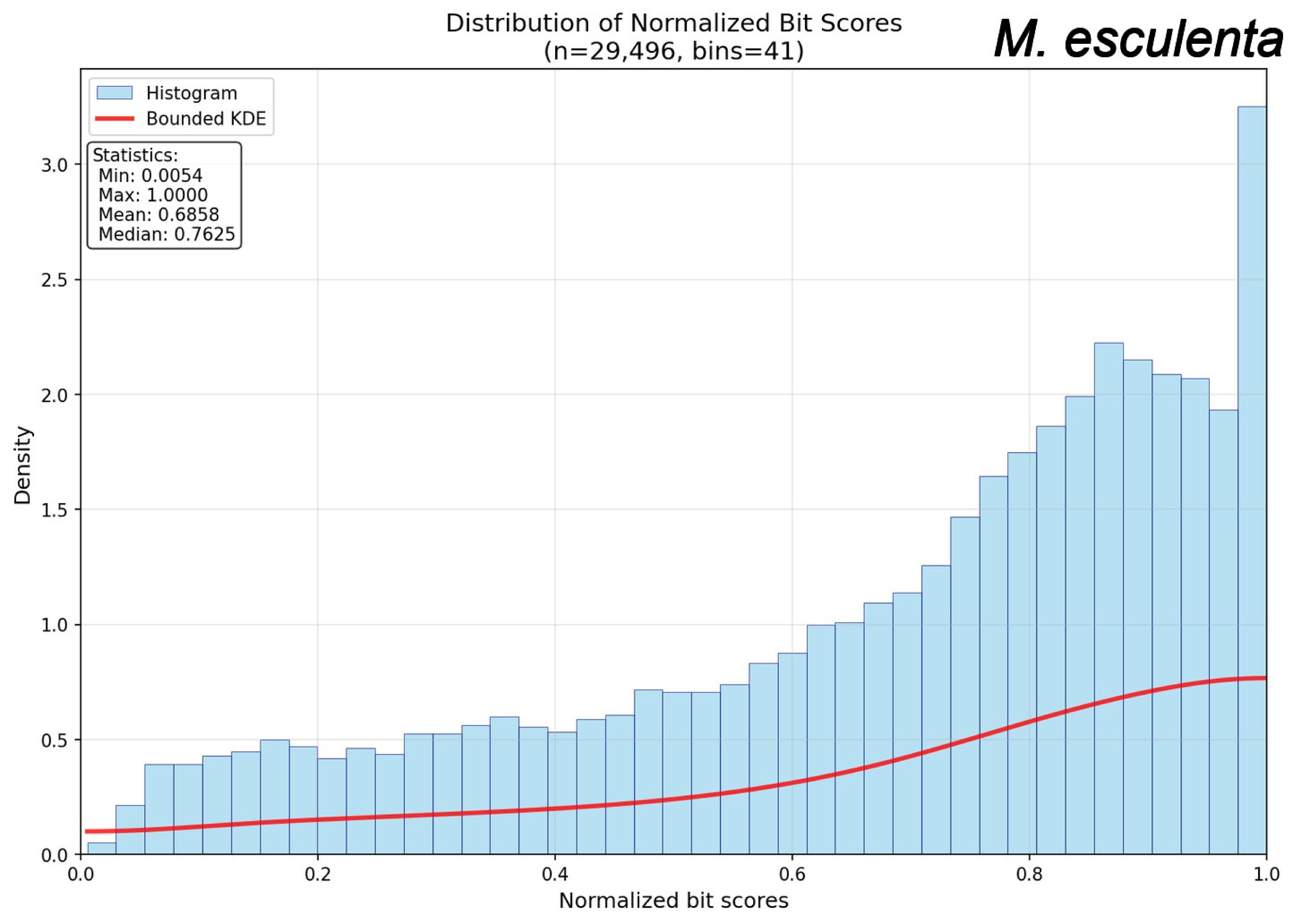
Distribution of Normalized Bit Scores
(n=29,496, bins=41)
M. esculenta
Histogram
Bounded KDE
Statistics:
Min: 0.0054
Max: 1.0000
Mean: 0.6858
Median: 0.7625
Density
Normalized bit scores
0.0
0.5
1.0
1.5
2.0
2.5
3.0
0.0
0.2
0.4
0.6
0.8
1.0

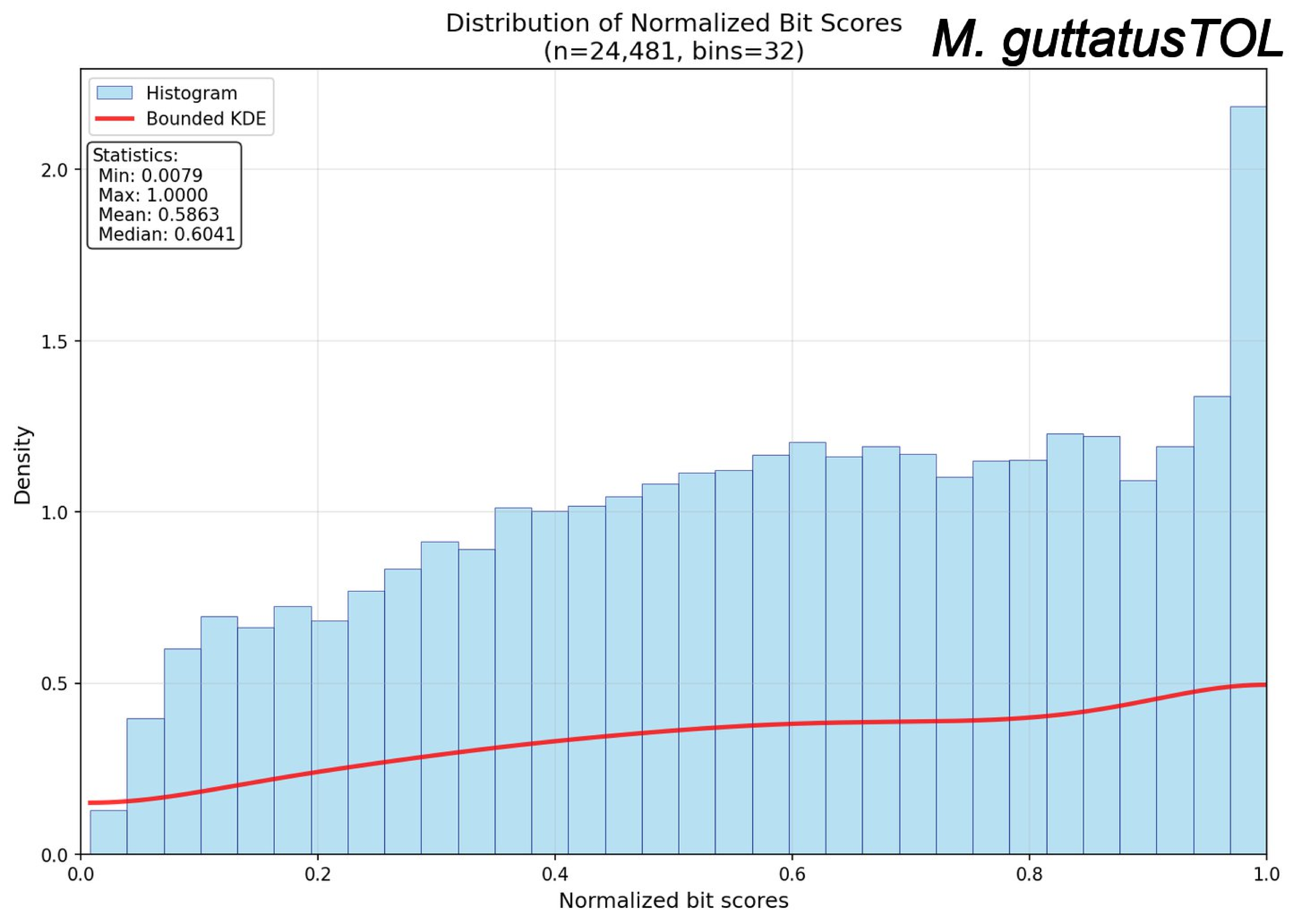
Distribution of Normalized Bit Scores
(n=24,481, bins=32)
M. guttatusTOL
Histogram
Bounded KDE
Statistics:
Min: 0.0079
Max: 1.0000
Mean: 0.5863
Median: 0.6041
Density
Normalized bit scores
0.0
0.5
1.0
1.5
2.0
0.0
0.2
0.4
0.6
0.8
1.0

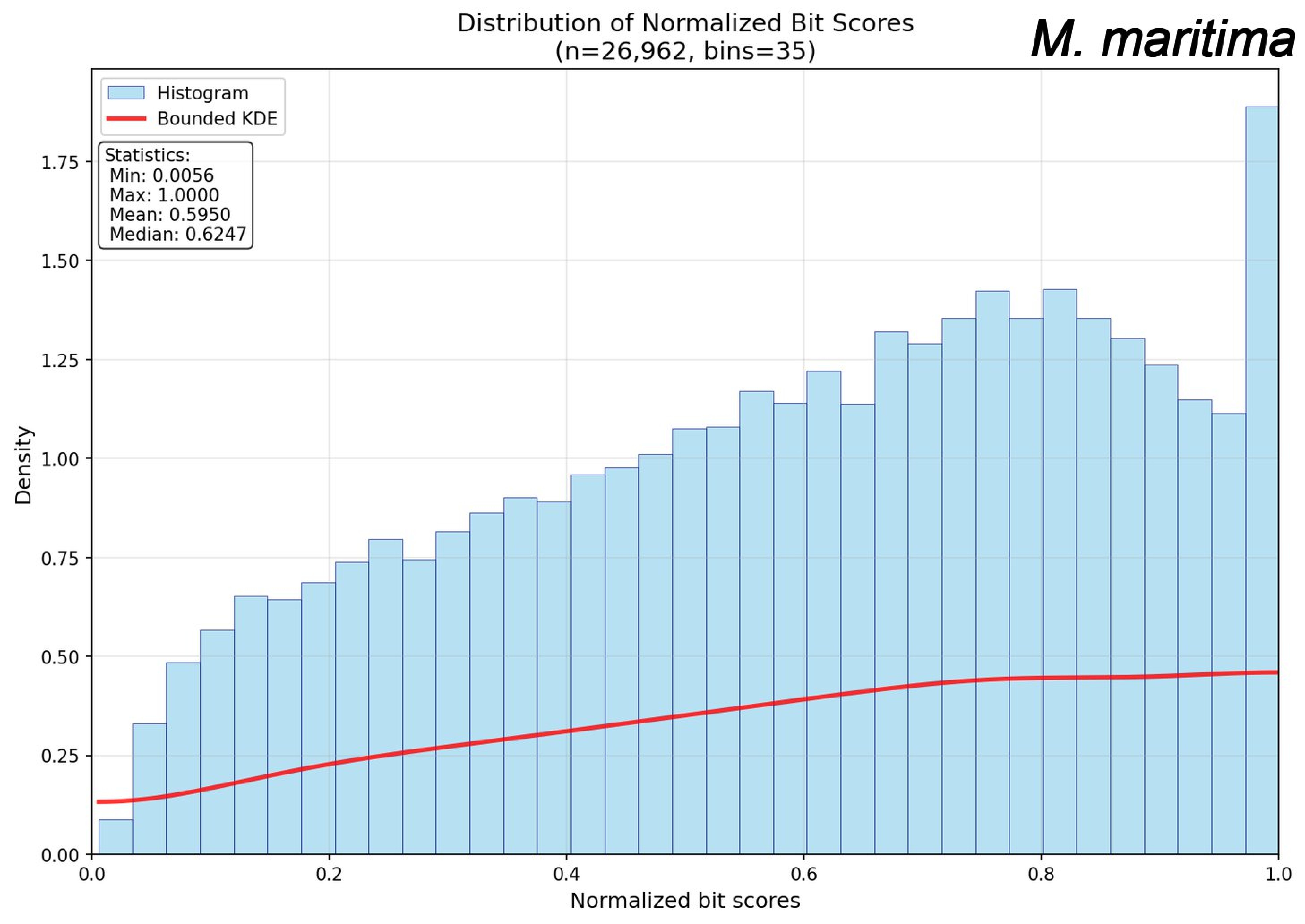
Distribution of Normalized Bit Scores
(n=26,962, bins=35)
M. maritima
Histogram
Bounded KDE
Statistics:
Min: 0.0056
Max: 1.0000
Mean: 0.5950
Median: 0.6247
Density
Normalized bit scores
0.00
0.25
0.50
0.75
1.00
1.25
1.50
1.75
0.0
0.2
0.4
0.6
0.8
1.0

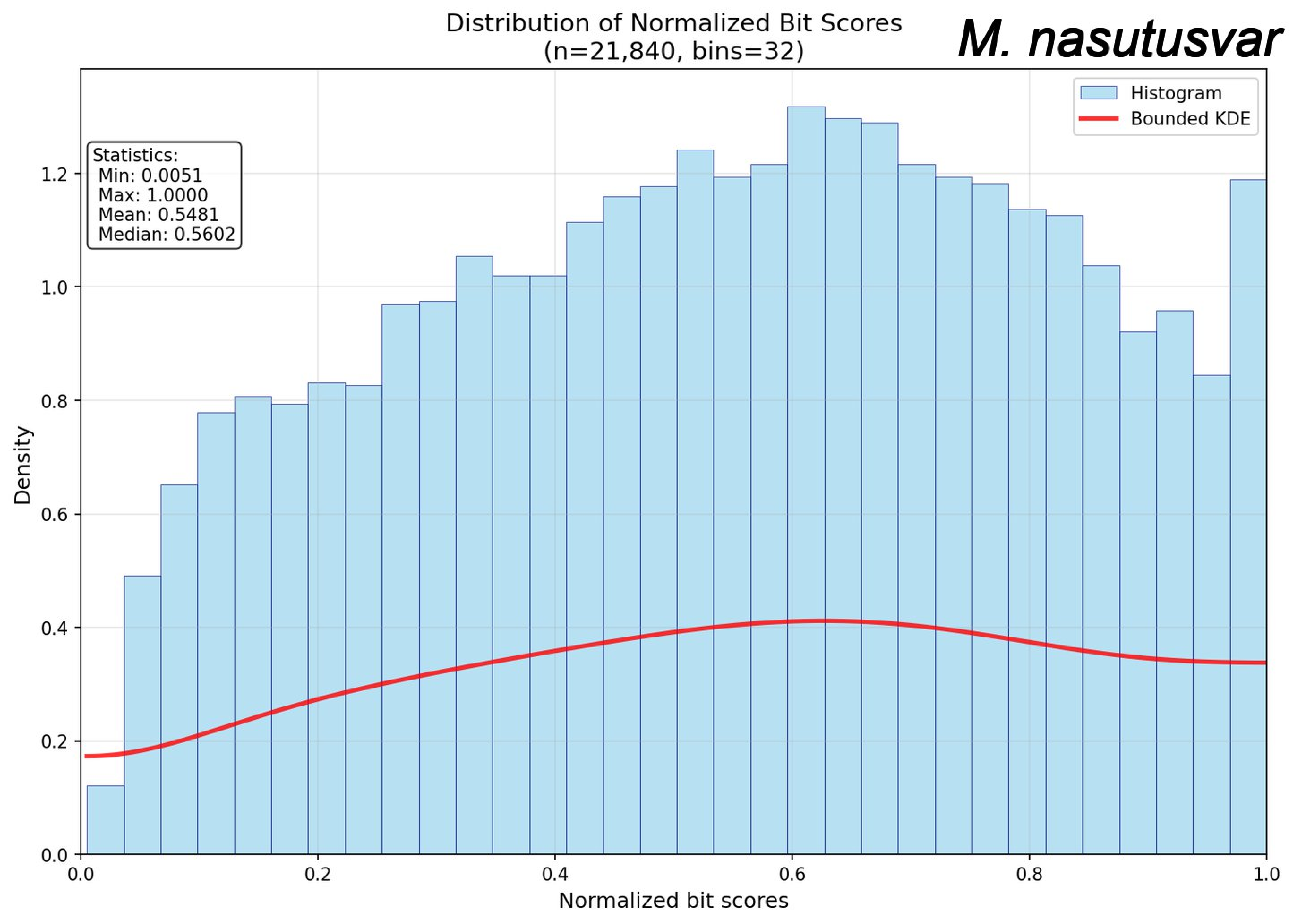
Distribution of Normalized Bit Scores
(n=21,840, bins=32)
M. nasutusvar
Histogram
Bounded KDE
Statistics:
Min: 0.0051
Max: 1.0000
Mean: 0.5481
Median: 0.5602
Density
Normalized bit scores
0.0
0.2
0.4
0.6
0.8
1.0
1.2

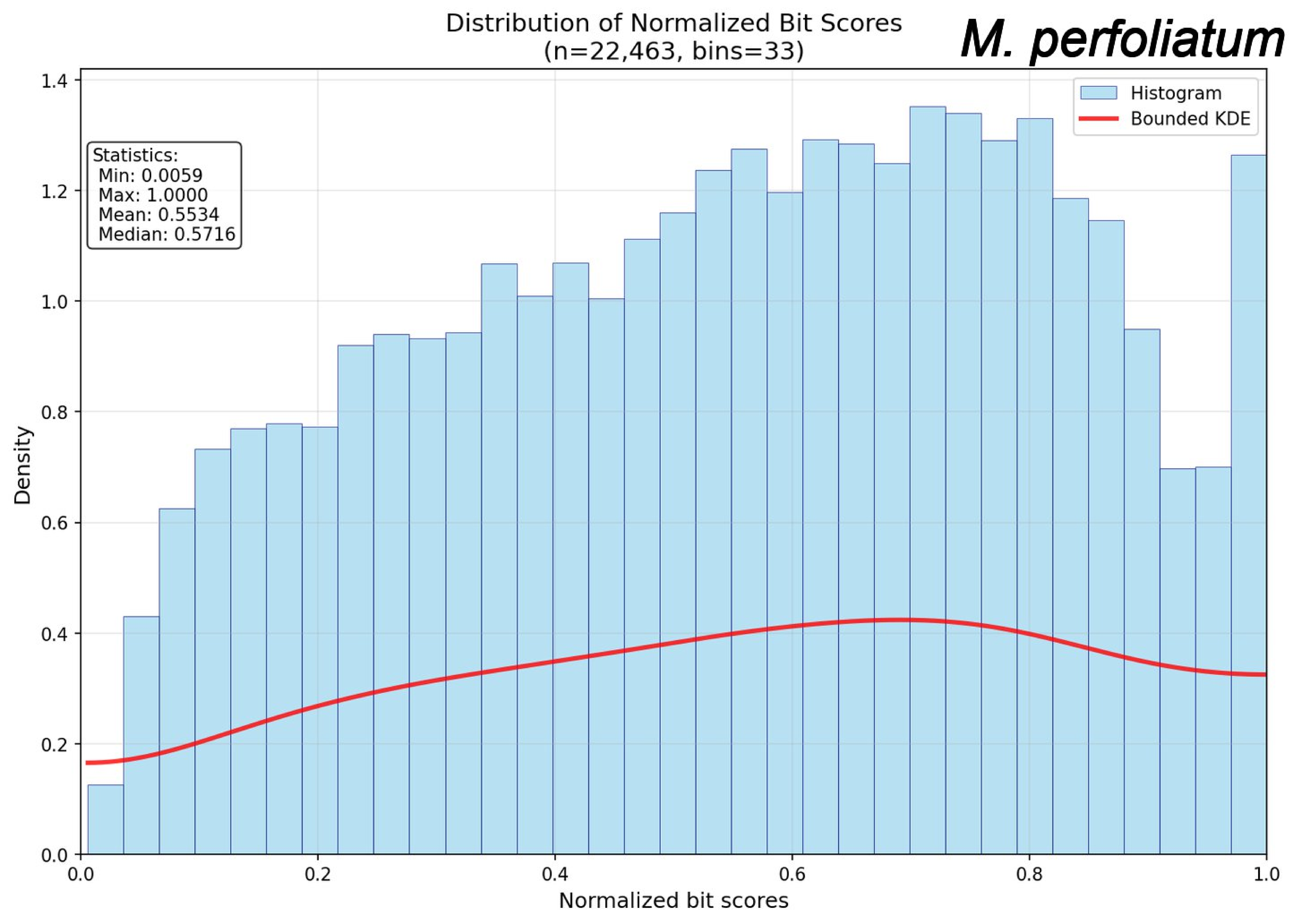
Distribution of Normalized Bit Scores
(n=22,463, bins=33)
M. perfoliatum
Histogram
Bounded KDE
Statistics:
Min: 0.0059
Max: 1.0000
Mean: 0.5534
Median: 0.5716
Density
Normalized bit scores
0.0
0.2
0.4
0.6
0.8
1.0
1.2
1.4

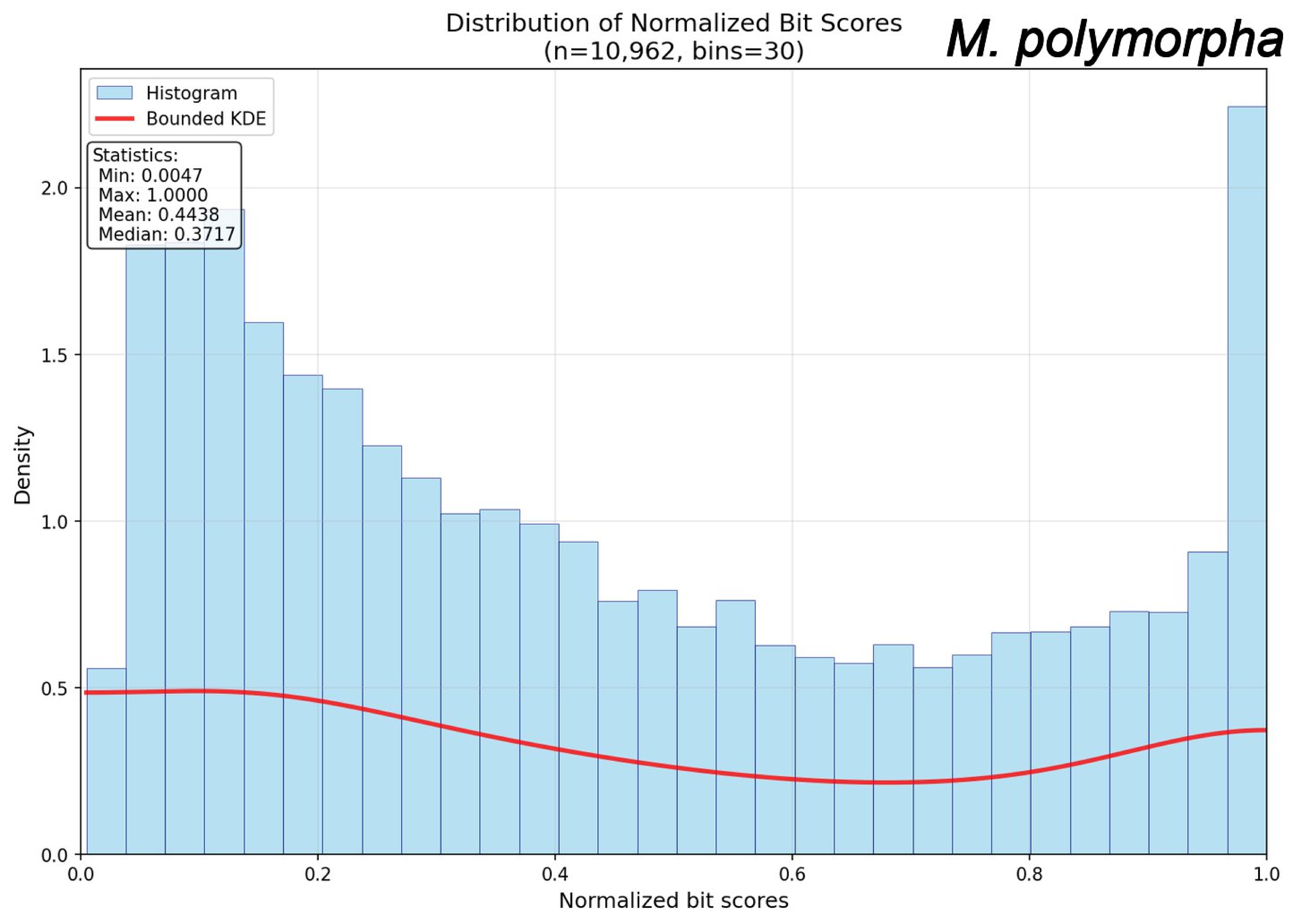
Distribution of Normalized Bit Scores
(n=10,962, bins=30)
M. polymorpha
Histogram
Bounded KDE
Statistics:
Min: 0.0047
Max: 1.0000
Mean: 0.4438
Median: 0.3717
Density
Normalized bit scores
0.0
0.5
1.0
1.5
2.0
0.0
0.2
0.4
0.6
0.8
1.0

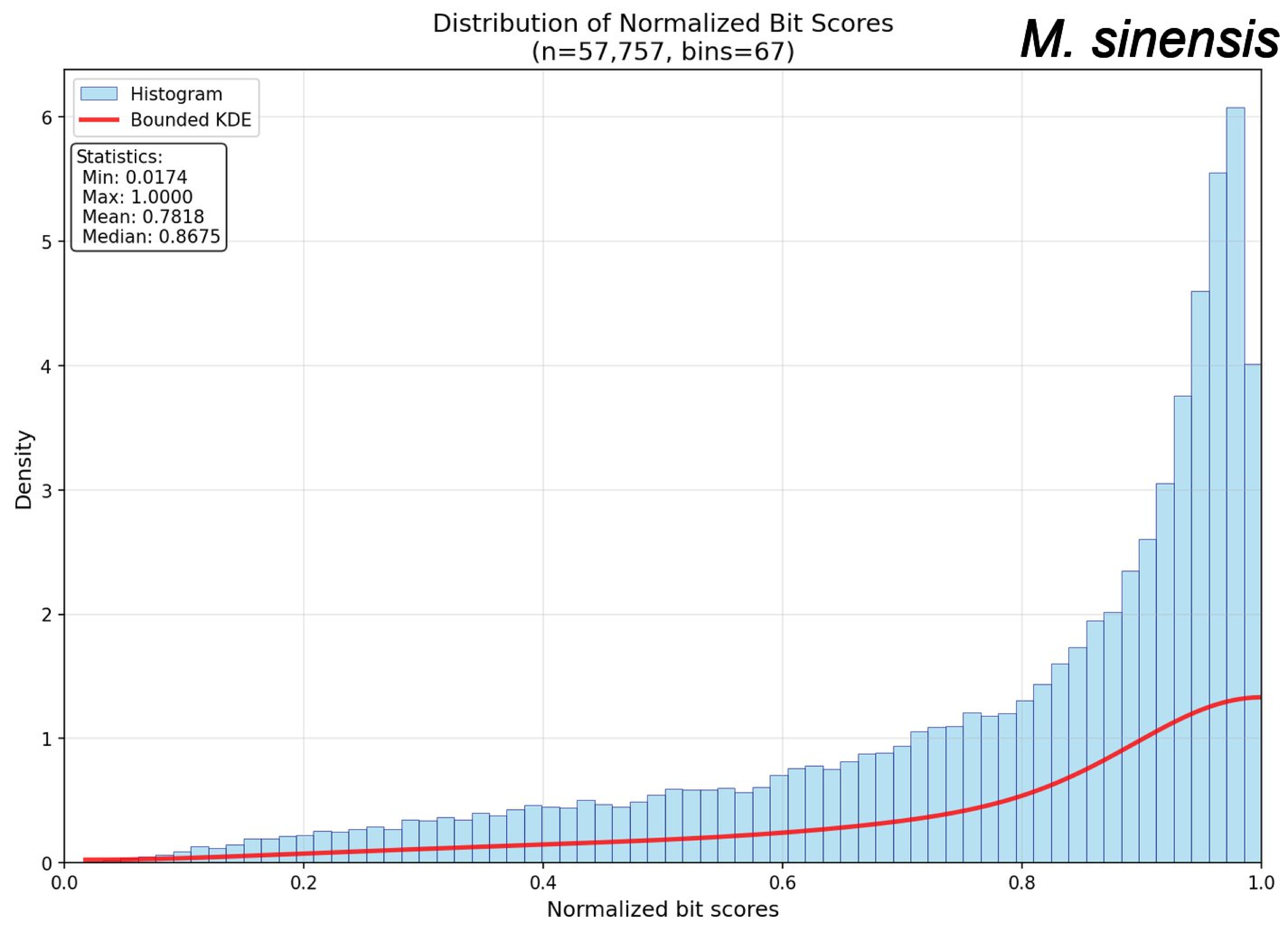
Distribution of Normalized Bit Scores
(n=57,757, bins=67)
M. sinensis
Histogram
Bounded KDE
Statistics:
Min: 0.0174
Max: 1.0000
Mean: 0.7818
Median: 0.8675
Density
Normalized bit scores
0
1
2
3
4
5
6
0.0
0.2
0.4
0.6
0.8
1.0

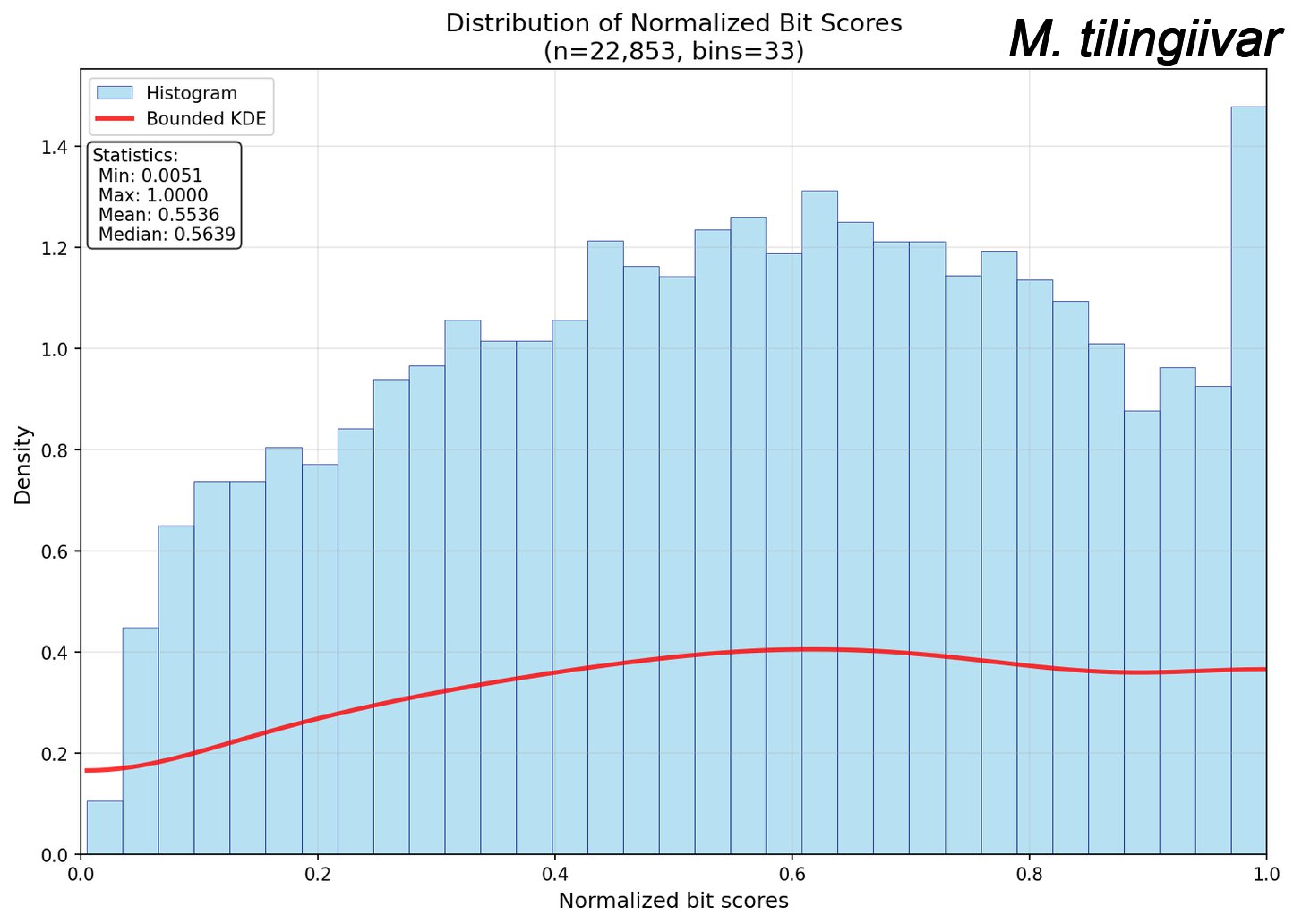
Distribution of Normalized Bit Scores
(n=22,853, bins=33)
M. tilingiivar
Histogram
Bounded KDE
Statistics:
Min: 0.0051
Max: 1.0000
Mean: 0.5536
Median: 0.5639
Density
Normalized bit scores
0.0
0.2
0.4
0.6
0.8
1.0
1.2
1.4

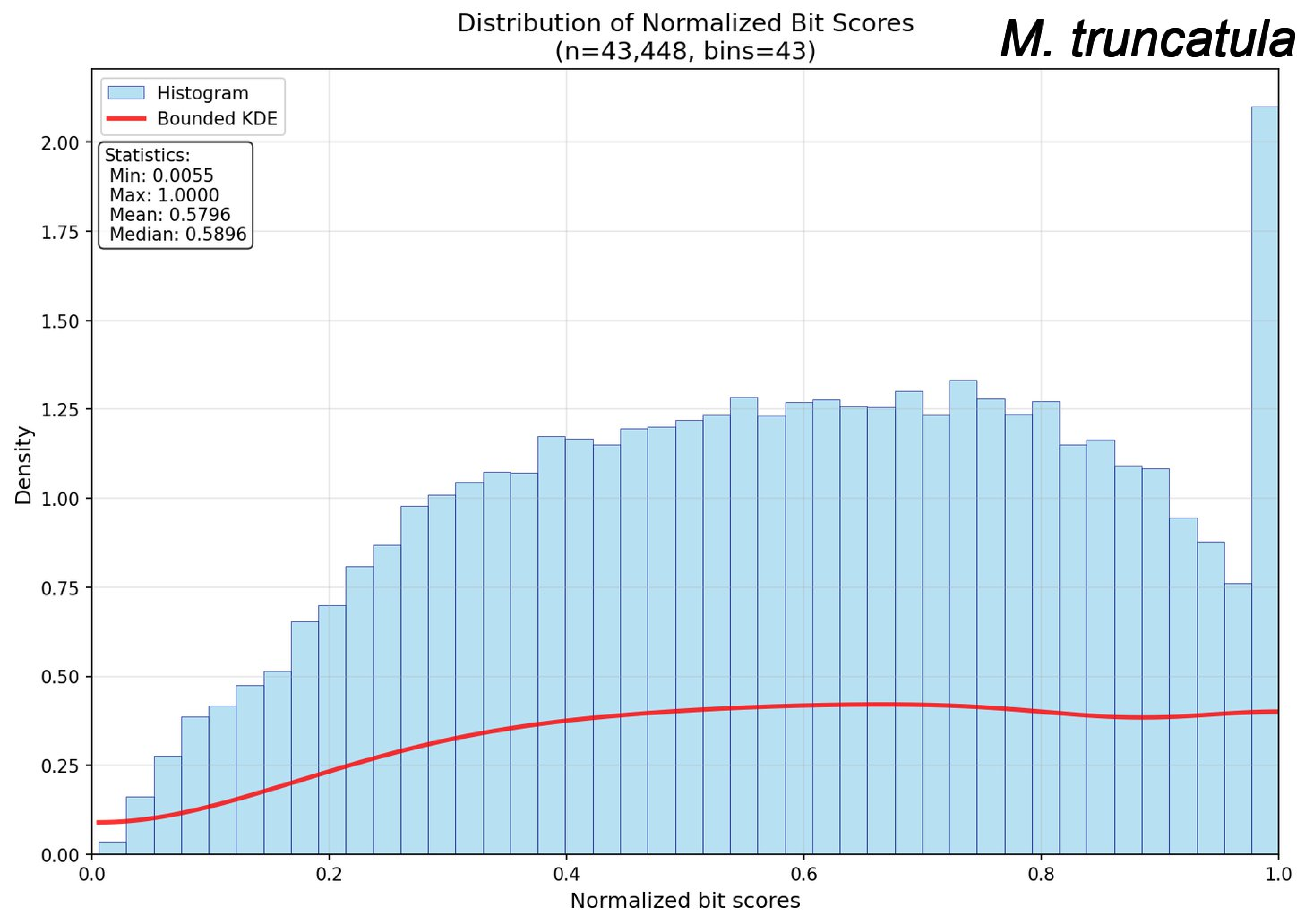

Distribution of Normalized Bit Scores
(n=43,448, bins=43)
M. truncatula
Histogram
Bounded KDE
Statistics:
Min: 0.0055
Max: 1.0000
Mean: 0.5796
Median: 0.5896
Density
Normalized bit scores
0.00
0.25
0.50
0.75
1.00
1.25
1.50
1.75
2.00
0.0
0.2
0.4
0.6
0.8
1.0

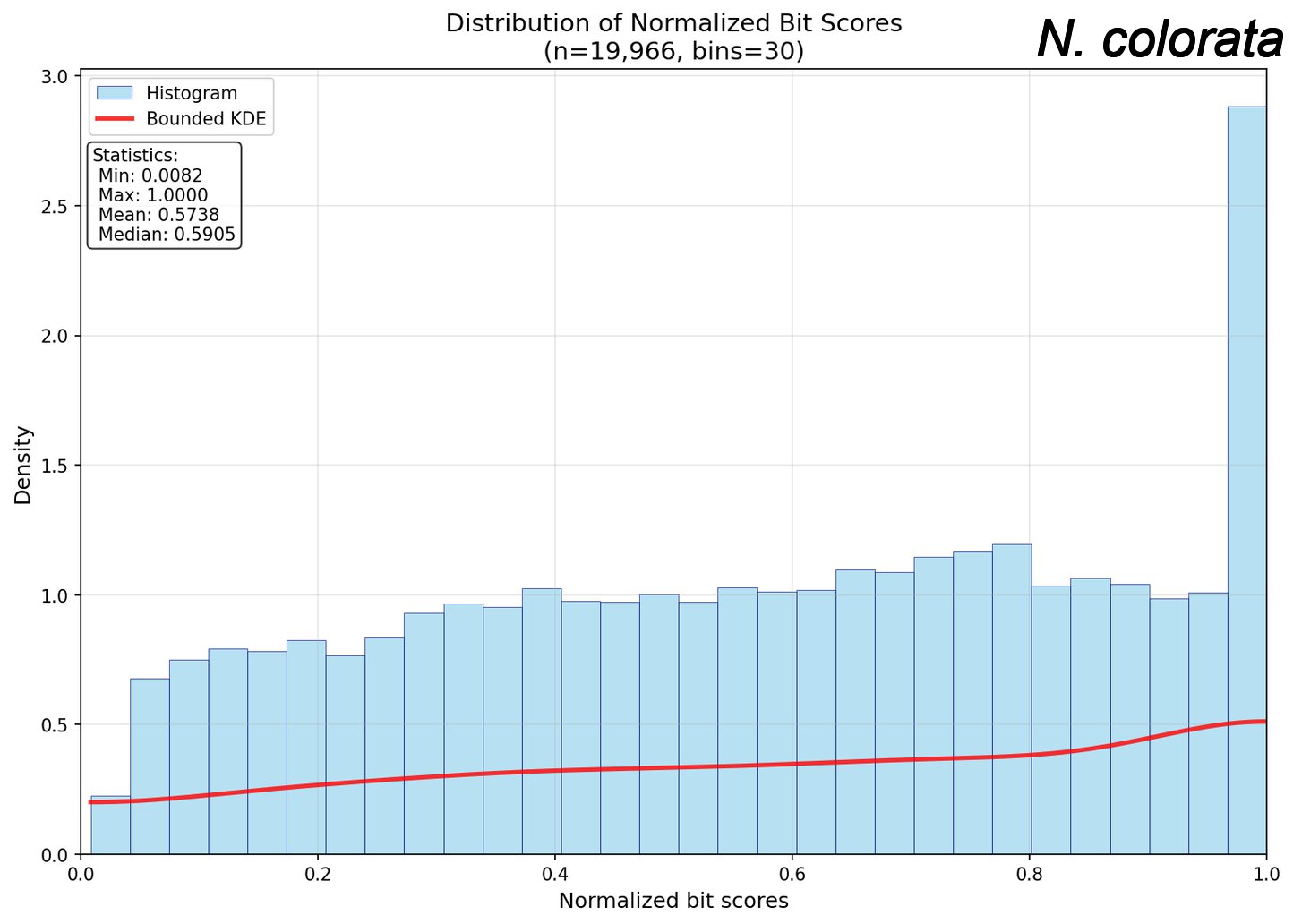
Distribution of Normalized Bit Scores
(n=19,966, bins=30)
N. colorata
Histogram
Bounded KDE
Statistics:
Min: 0.0082
Max: 1.0000
Mean: 0.5738
Median: 0.5905
Density
Normalized bit scores
0.0
0.5
1.0
1.5
2.0
2.5
3.0
0.2
0.4
0.6
0.8
1.0

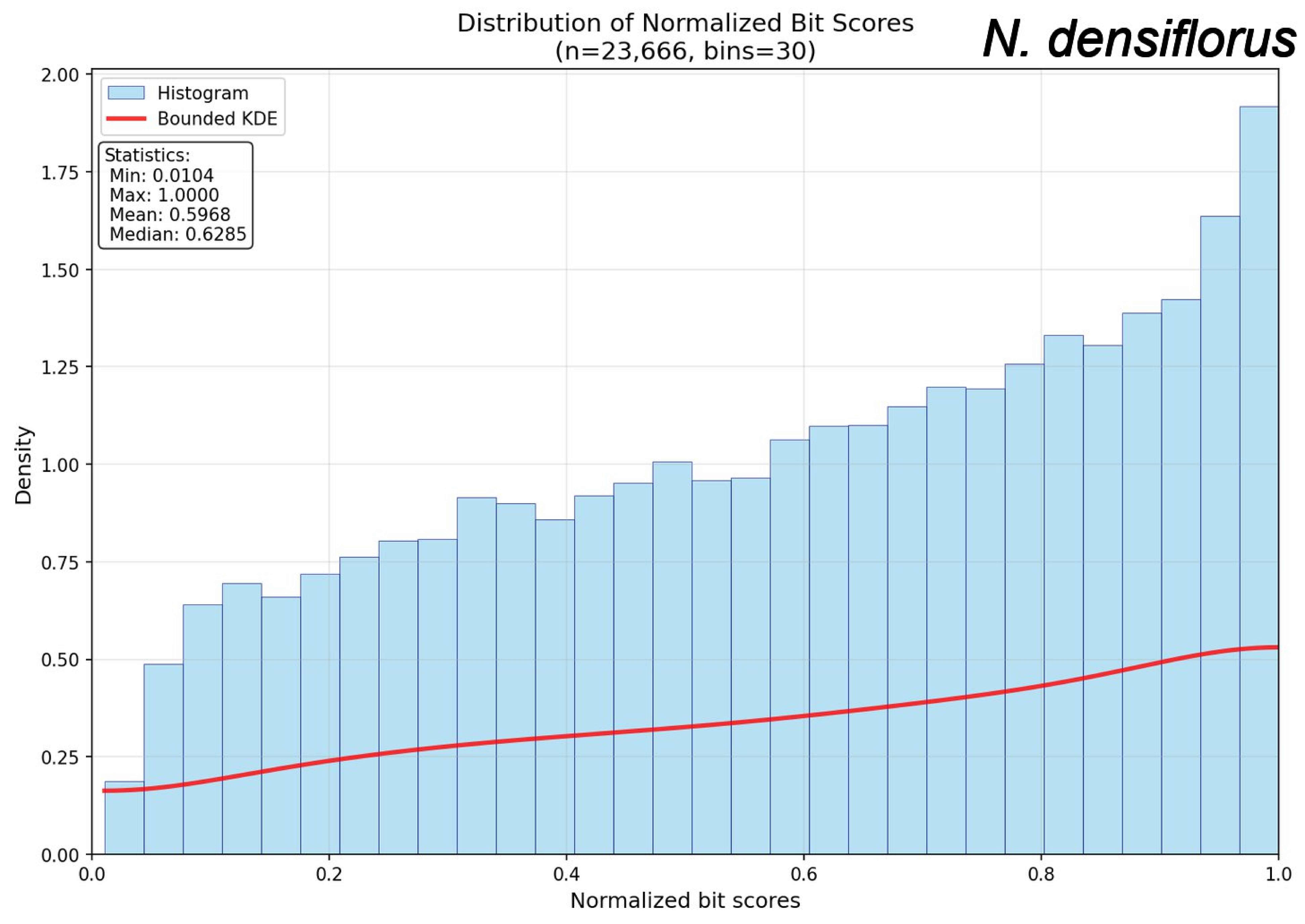
Distribution of Normalized Bit Scores
(n=23,666, bins=30)
N. densiflorus
Histogram
Bounded KDE
Statistics:
Min: 0.0104
Max: 1.0000
Mean: 0.5968
Median: 0.6285
Density
Normalized bit scores
0.00
0.25
0.50
0.75
1.00
1.25
1.50
1.75
2.00
0.0
0.2
0.4
0.6
0.8
1.0

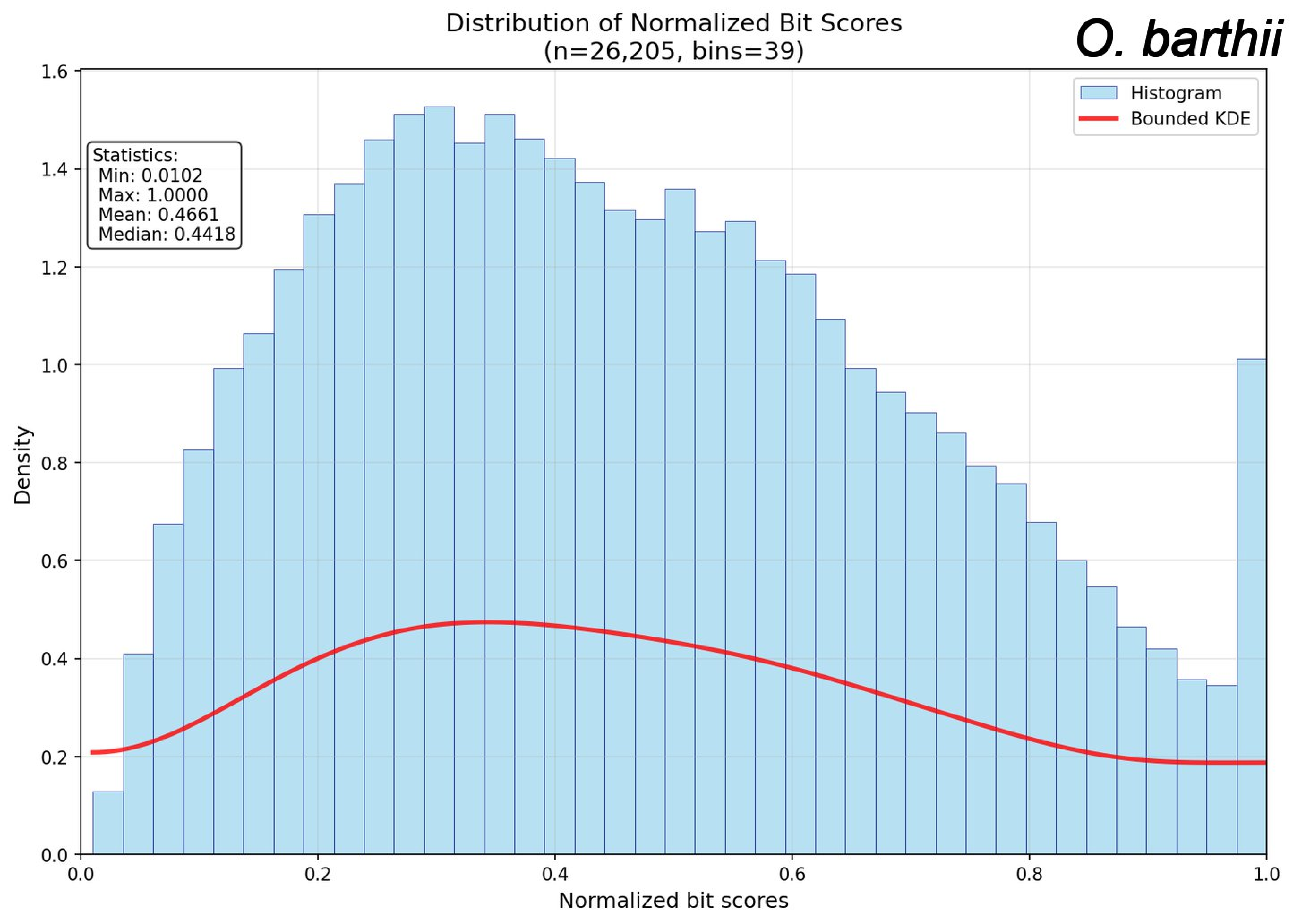
Distribution of Normalized Bit Scores
(n=26,205, bins=39)
O. barthii
Histogram
Bounded KDE
Statistics:
Min: 0.0102
Max: 1.0000
Mean: 0.4661
Median: 0.4418
Density
Normalized bit scores
0.0
0.2
0.4
0.6
0.8
1.0
1.2
1.4
1.6

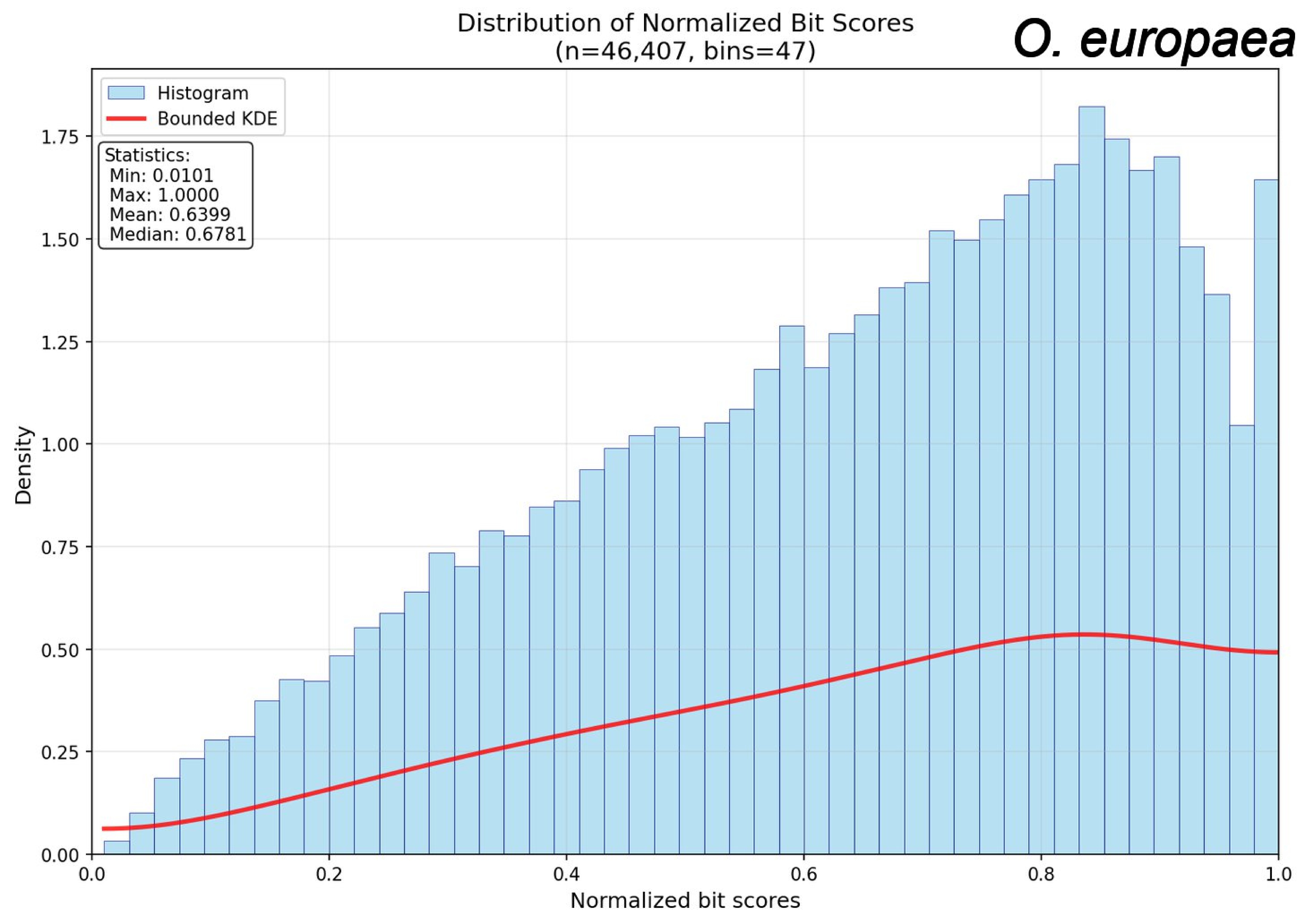
Distribution of Normalized Bit Scores
(n=46,407, bins=47)
O. europaea
Histogram
Bounded KDE
Statistics:
Min: 0.0101
Max: 1.0000
Mean: 0.6399
Median: 0.6781
Density
Normalized bit scores
0.00
0.25
0.50
0.75
1.00
1.25
1.50
1.75
0.0
0.2
0.4
0.6
0.8
1.0

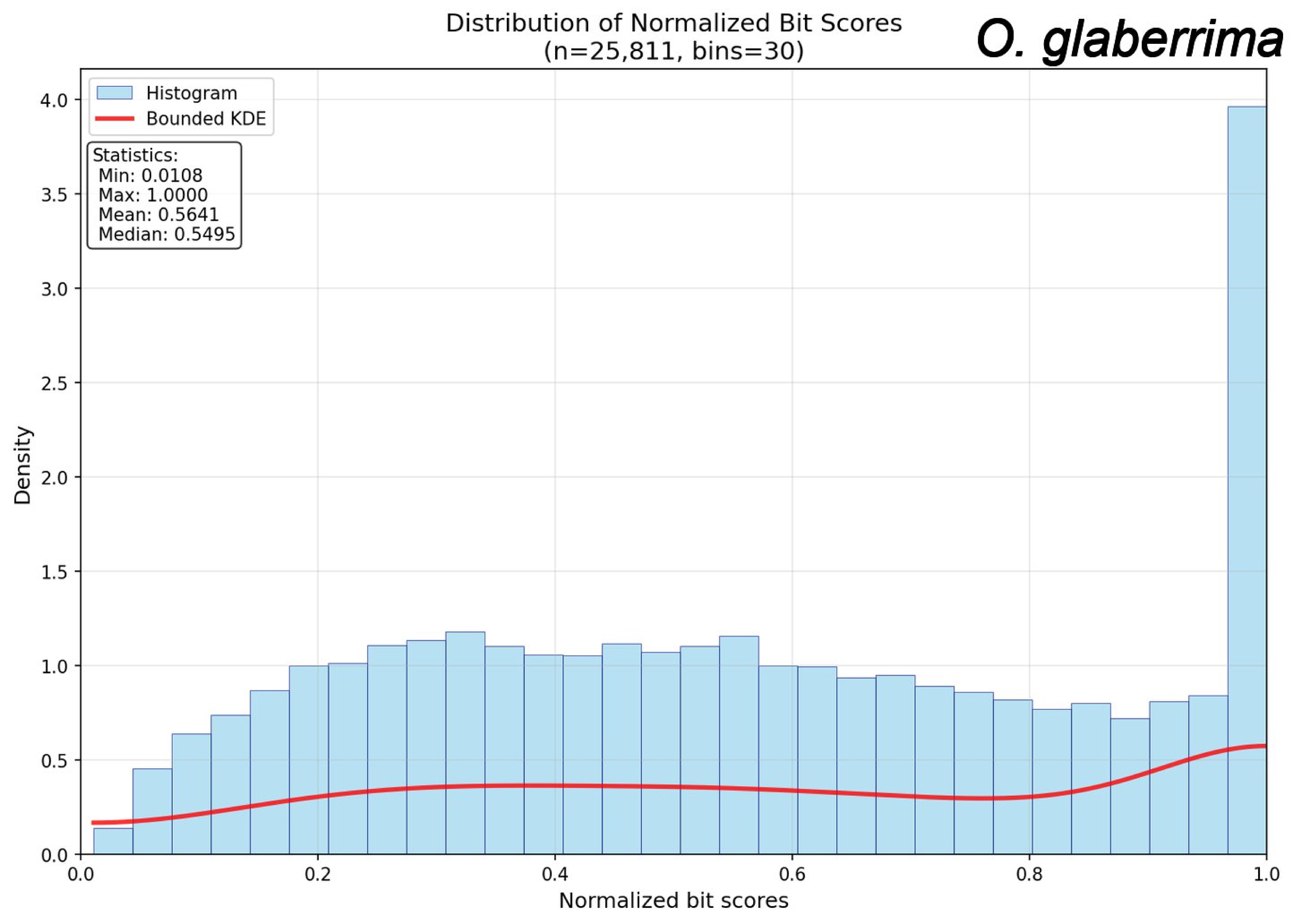
Distribution of Normalized Bit Scores
(n=25,811, bins=30)
O. glaberrima
Histogram
Bounded KDE
Statistics:
Min: 0.0108
Max: 1.0000
Mean: 0.5641
Median: 0.5495
Density
Normalized bit scores
0.0
0.5
1.0
1.5
2.0
2.5
3.0
3.5
4.0
0.0
0.2
0.4
0.6
0.8
1.0

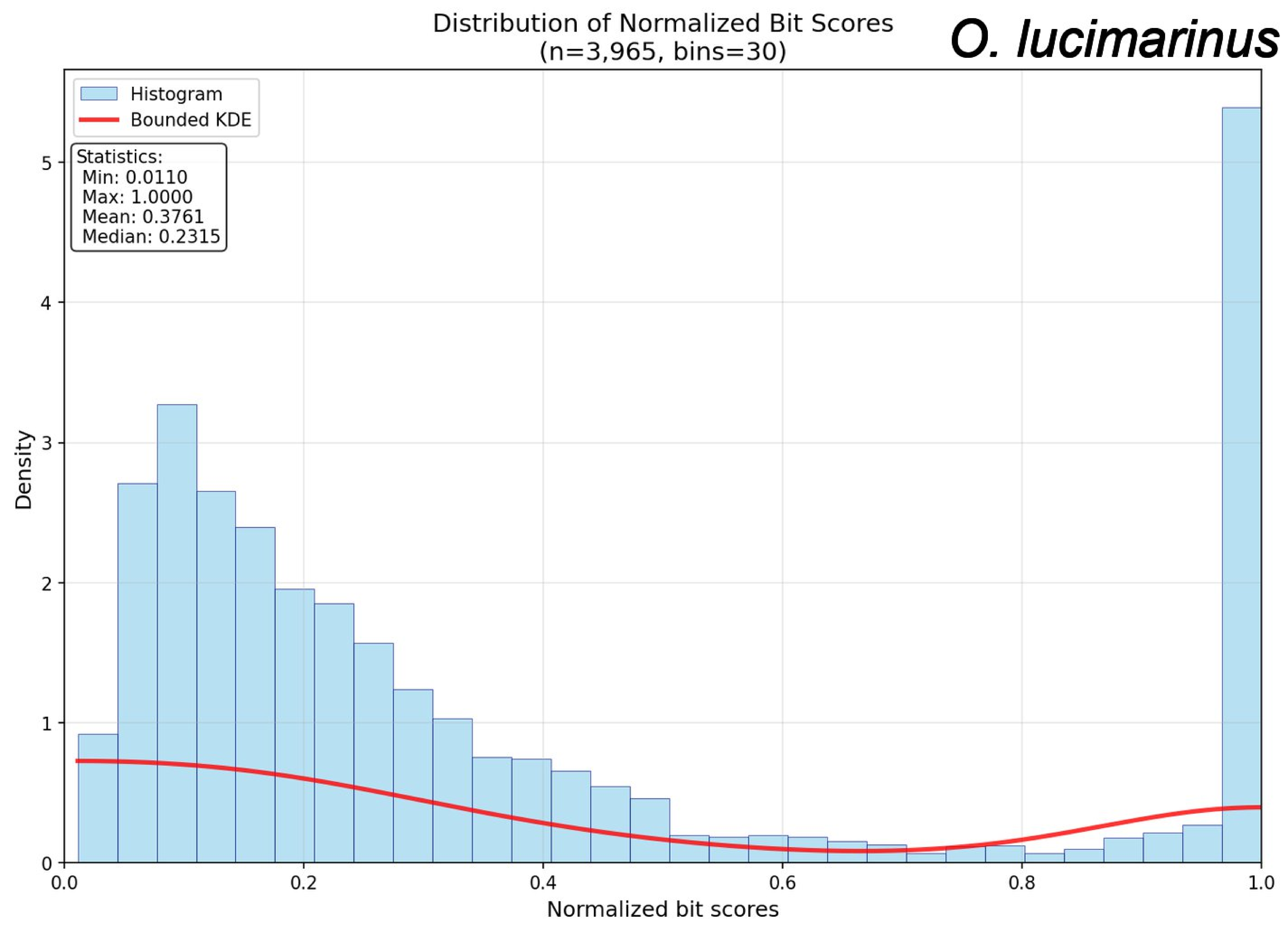

Distribution of Normalized Bit Scores
(n=3,965, bins=30)
O. lucimarinus
Histogram
Bounded KDE
Statistics:
Min: 0.0110
Max: 1.0000
Mean: 0.3761
Median: 0.2315
Density
Normalized bit scores
0
1
2
3
4
5
0.0
0.2
0.4
0.6
0.8
1.0

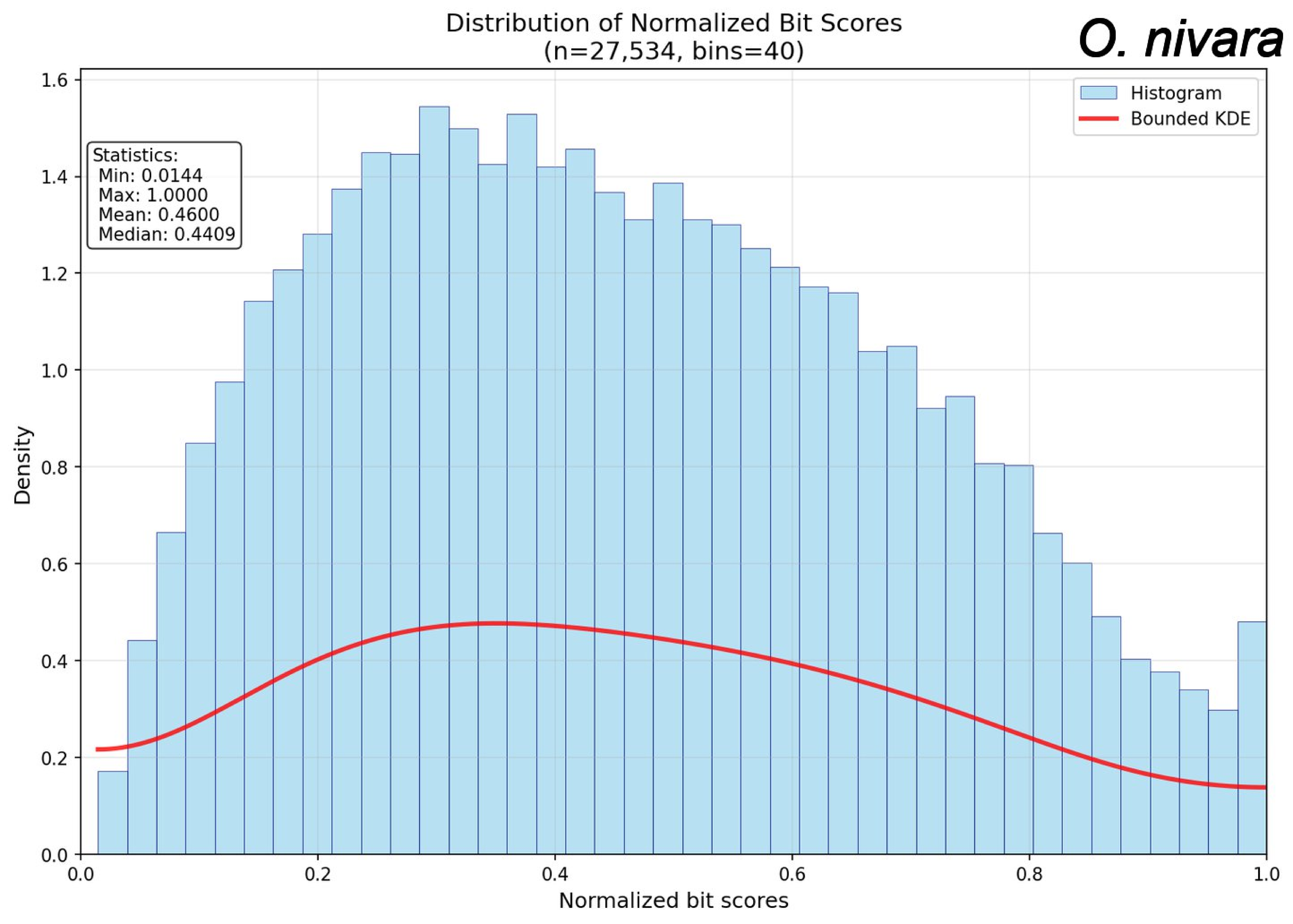
Distribution of Normalized Bit Scores
(n=27,534, bins=40)
O. nivara
Statistics:
Min: 0.0144
Max: 1.0000
Mean: 0.4600
Median: 0.4409
Histogram
Bounded KDE
Density
Normalized bit scores
0.0
0.2
0.4
0.6
0.8
1.0
1.2
1.4
1.6

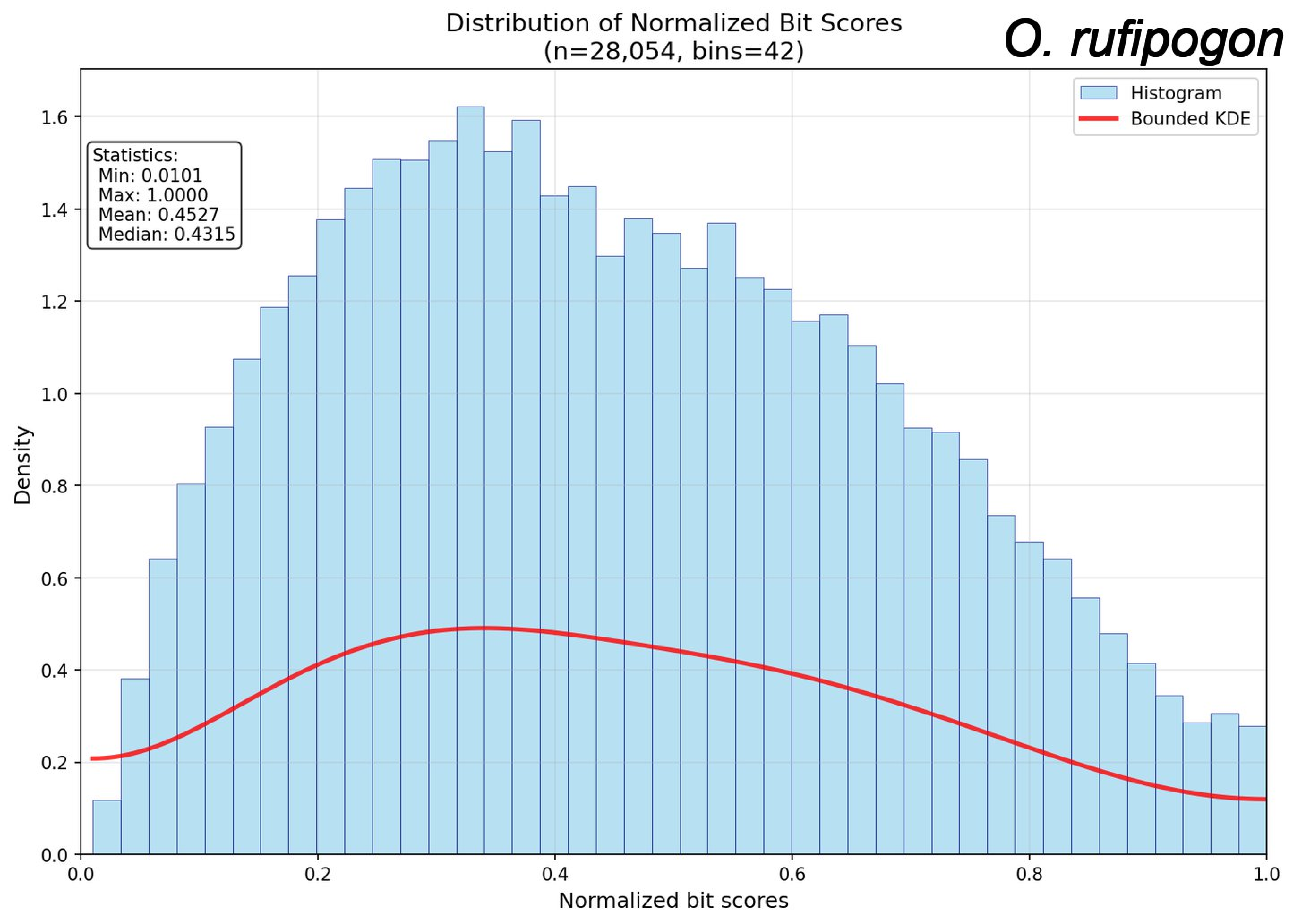

Distribution of Normalized Bit Scores
(n=28,054, bins=42)
O. rufipogon
Statistics:
Min: 0.0101
Max: 1.0000
Mean: 0.4527
Median: 0.4315
Histogram
Bounded KDE
Density
Normalized bit scores
0.0
0.2
0.4
0.6
0.8
1.0
1.2
1.4
1.6

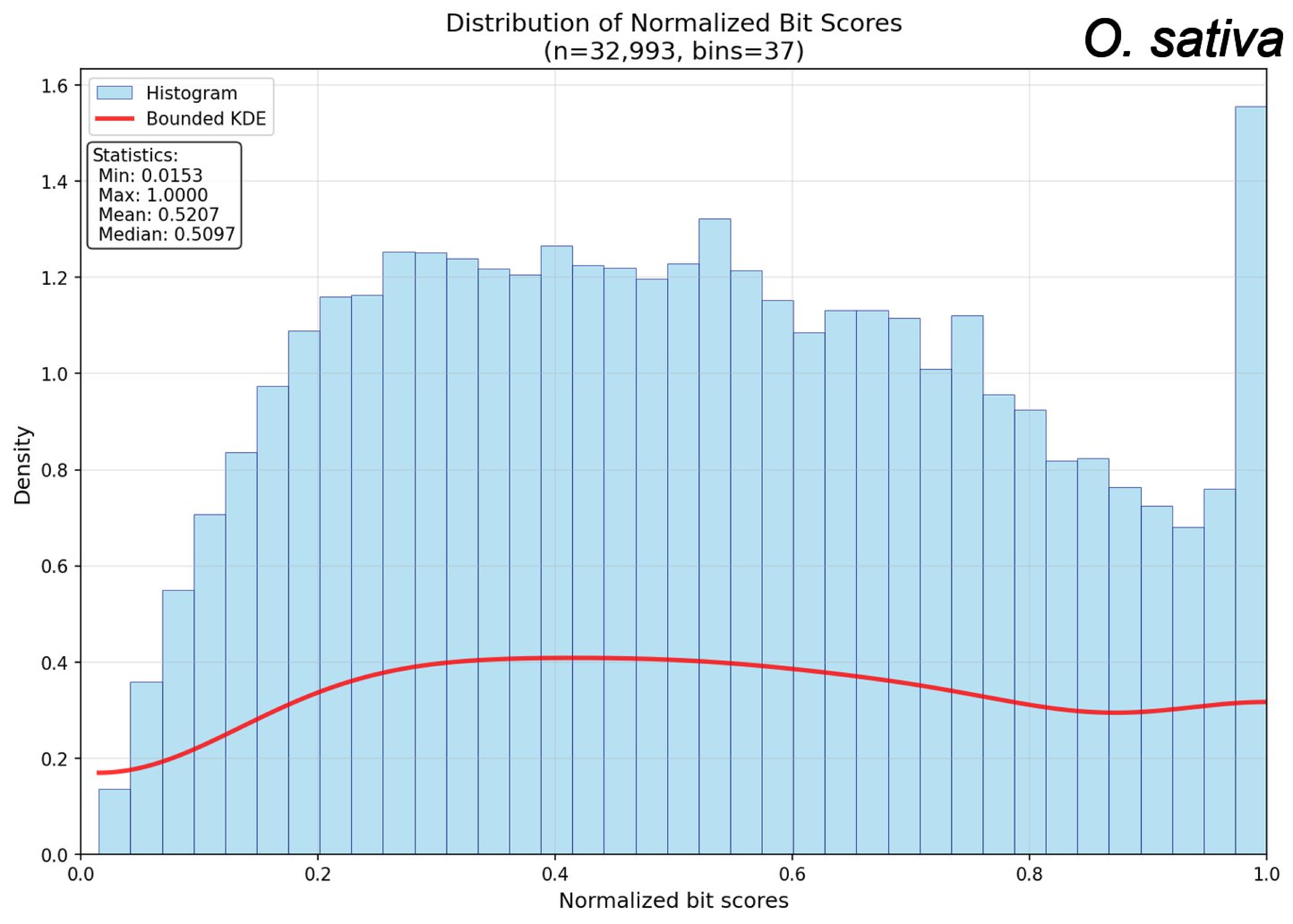
Distribution of Normalized Bit Scores
(n=32,993, bins=37)
O. sativa
Histogram
Bounded KDE
Statistics:
Min: 0.0153
Max: 1.0000
Mean: 0.5207
Median: 0.5097
Density
Normalized bit scores
0.0
0.2
0.4
0.6
0.8
1.0
1.2
1.4
1.6

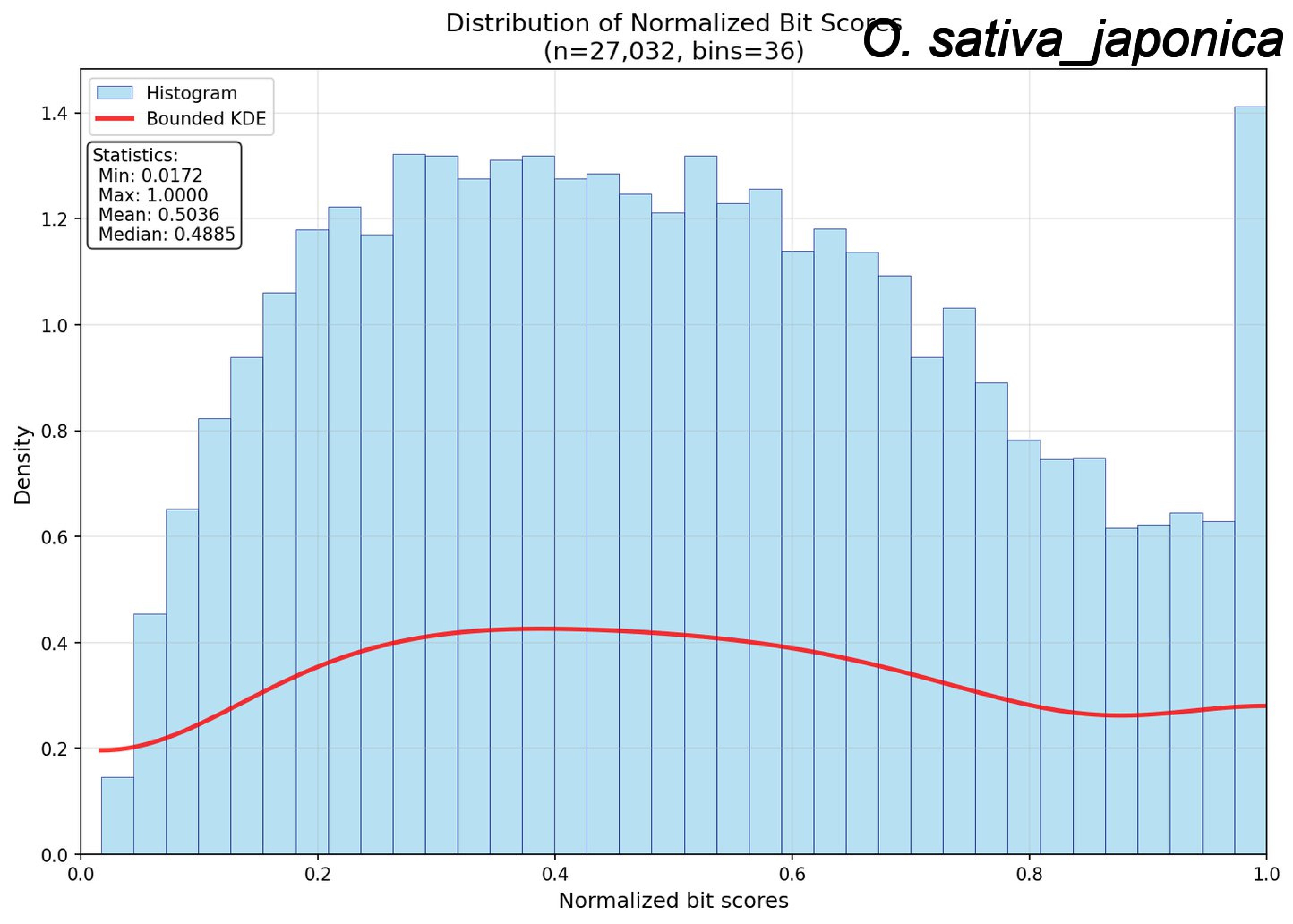
Distribution of Normalized Bit Scores
(n=27,032, bins=36)
O. sativa_japonica
Histogram
Bounded KDE
Statistics:
Min: 0.0172
Max: 1.0000
Mean: 0.5036
Median: 0.4885
Density
Normalized bit scores
0.0
0.2
0.4
0.6
0.8
1.0
1.2
1.4

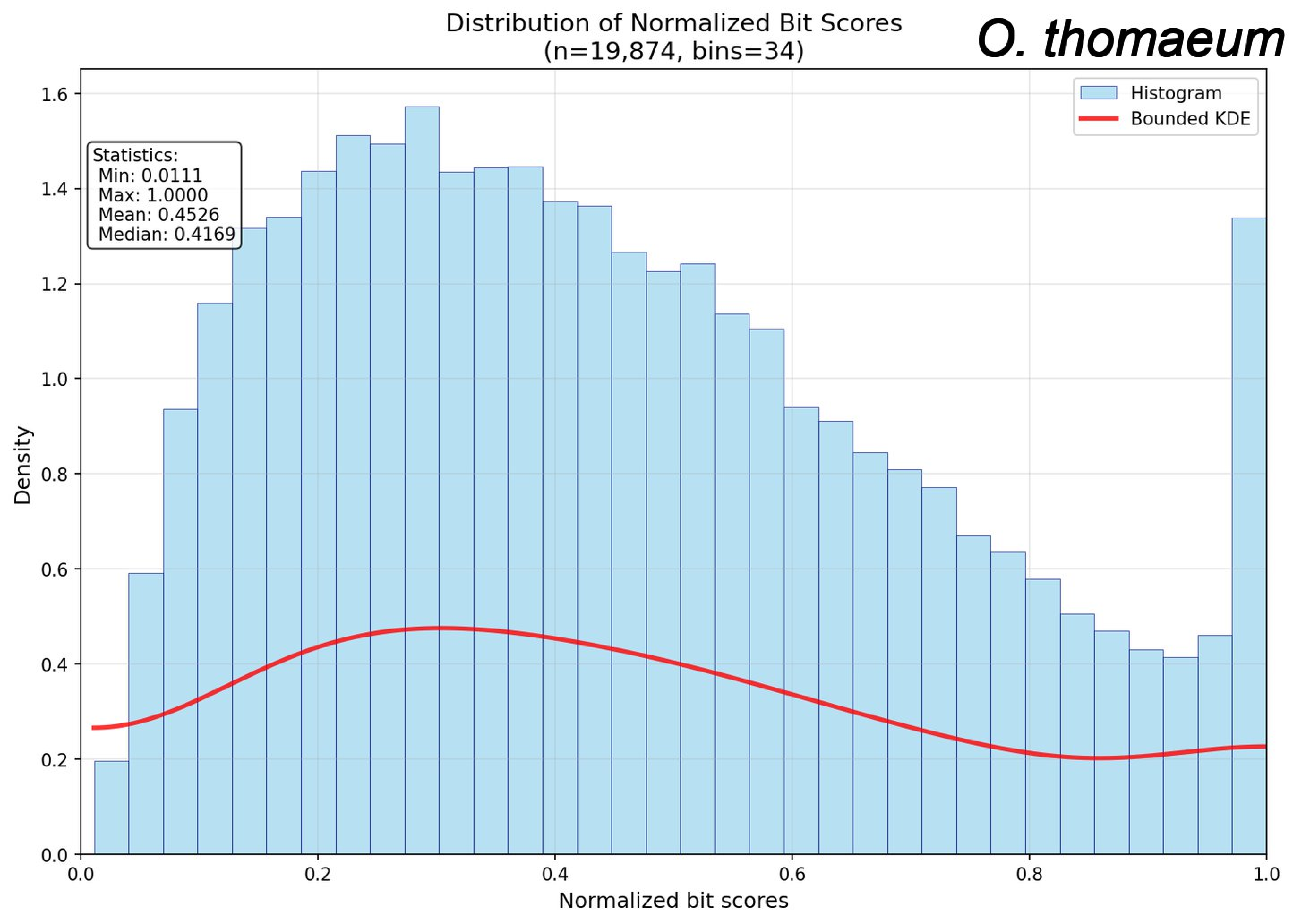
Distribution of Normalized Bit Scores
(n=19,874, bins=34)
O. thomaeum
Histogram
Bounded KDE
Statistics:
Min: 0.0111
Max: 1.0000
Mean: 0.4526
Median: 0.4169
Density
Normalized bit scores
0.0
0.2
0.4
0.6
0.8
1.0
1.2
1.4
1.6

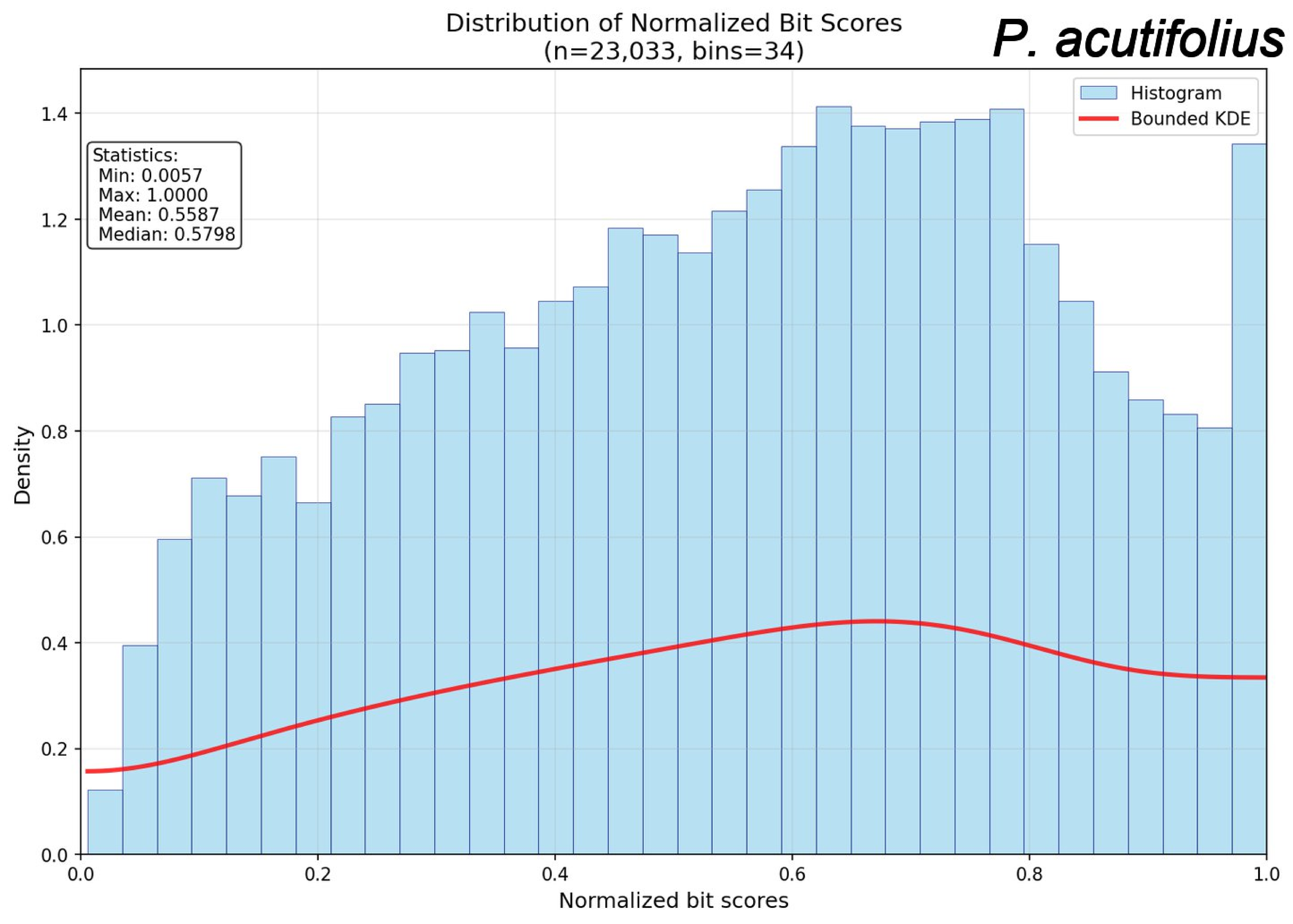

Distribution of Normalized Bit Scores
(n=23,033, bins=34)
P. acutifolius
Histogram
Bounded KDE
Statistics:
Min: 0.0057
Max: 1.0000
Mean: 0.5587
Median: 0.5798
Density
Normalized bit scores
0.0
0.2
0.4
0.6
0.8
1.0
1.2
1.4

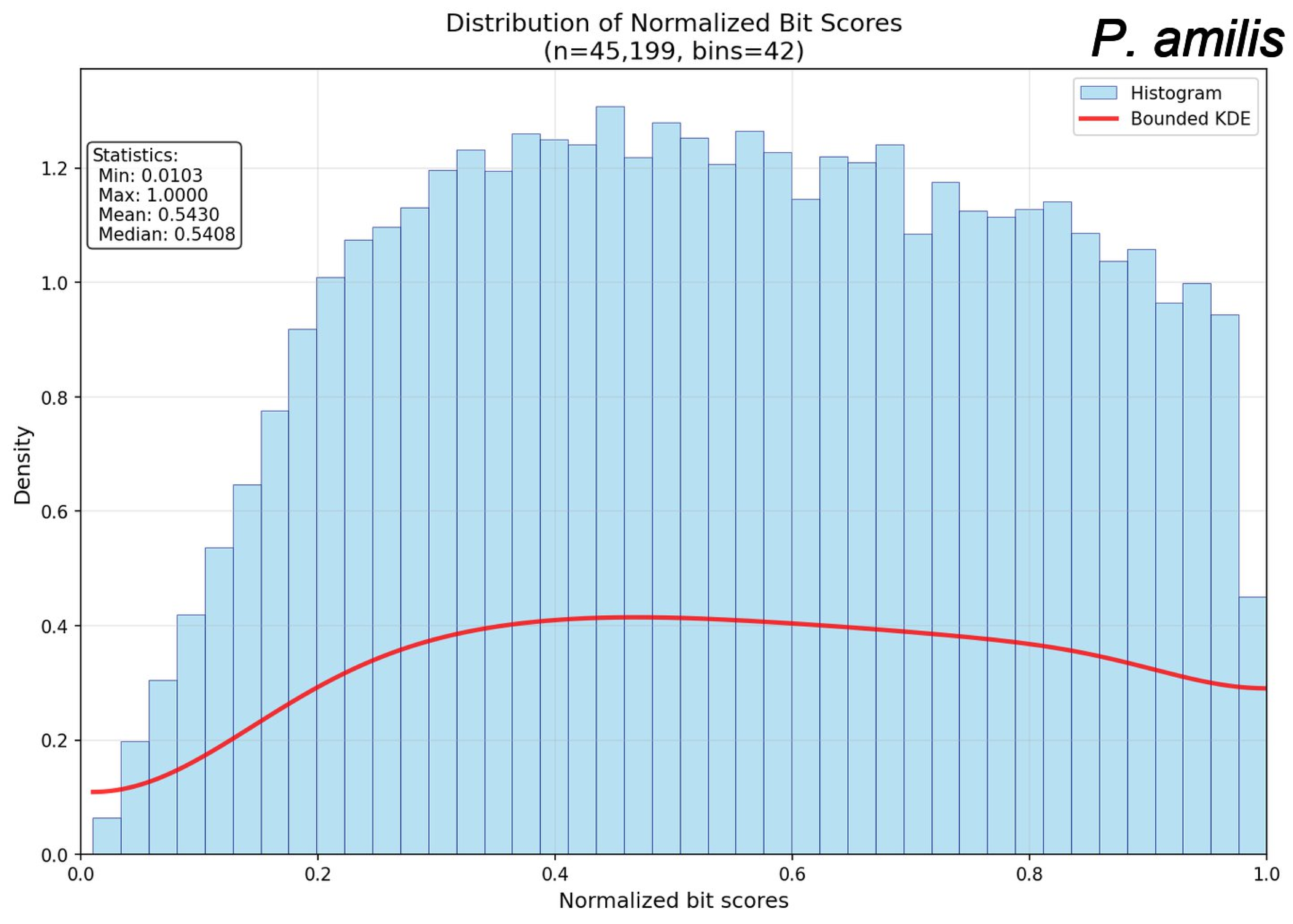

Distribution of Normalized Bit Scores
(n=45,199, bins=42)
P. amilis
Statistics:
Min: 0.0103
Max: 1.0000
Mean: 0.5430
Median: 0.5408
Histogram
Bounded KDE
Density
Normalized bit scores
0.0
0.2
0.4
0.6
0.8
1.0
1.2

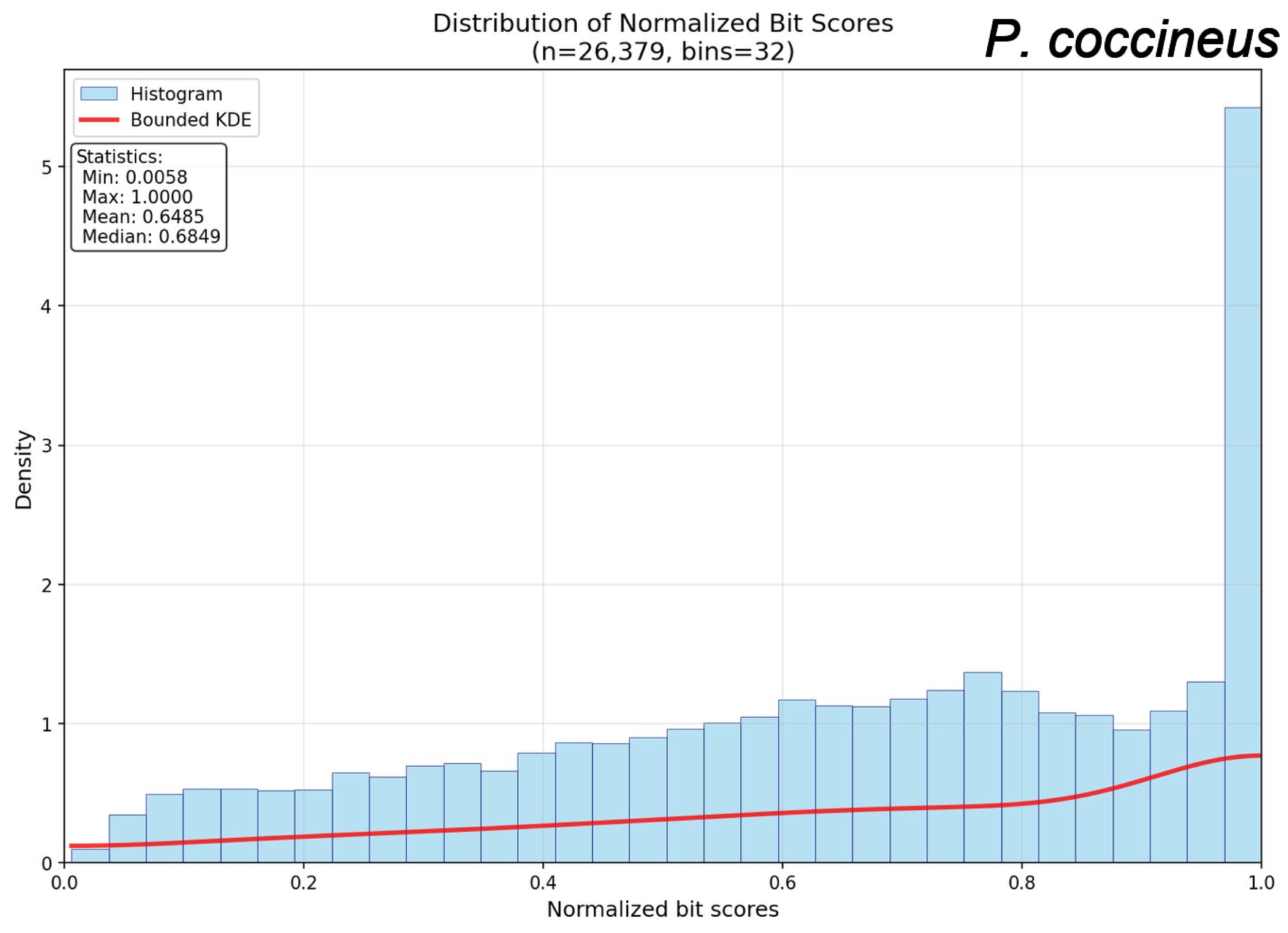

Distribution of Normalized Bit Scores
(n=26,379, bins=32)
P. coccineus
Histogram
Bounded KDE
Statistics:
Min: 0.0058
Max: 1.0000
Mean: 0.6485
Median: 0.6849
Density
Normalized bit scores
0
1
2
3
4
5
0.0
0.2
0.4
0.6
0.8
1.0

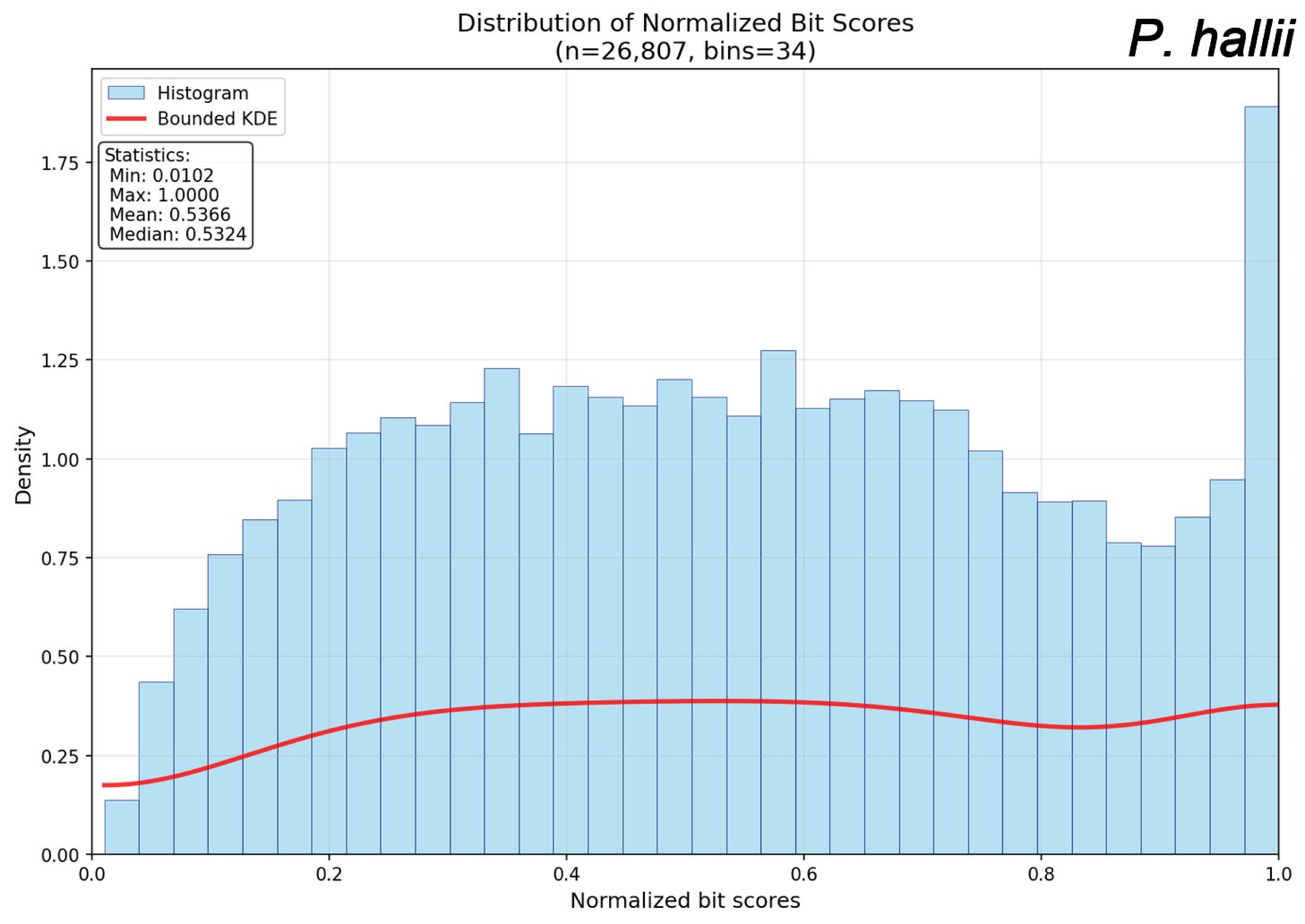
Distribution of Normalized Bit Scores
(n=26,807, bins=34)
P. hallii
Histogram
Bounded KDE
Statistics:
Min: 0.0102
Max: 1.0000
Mean: 0.5366
Median: 0.5324
Density
0.00
0.25
0.50
0.75
1.00
1.25
1.50
1.75
0.0
0.2
0.4
0.6
0.8
1.0
Normalized bit scores

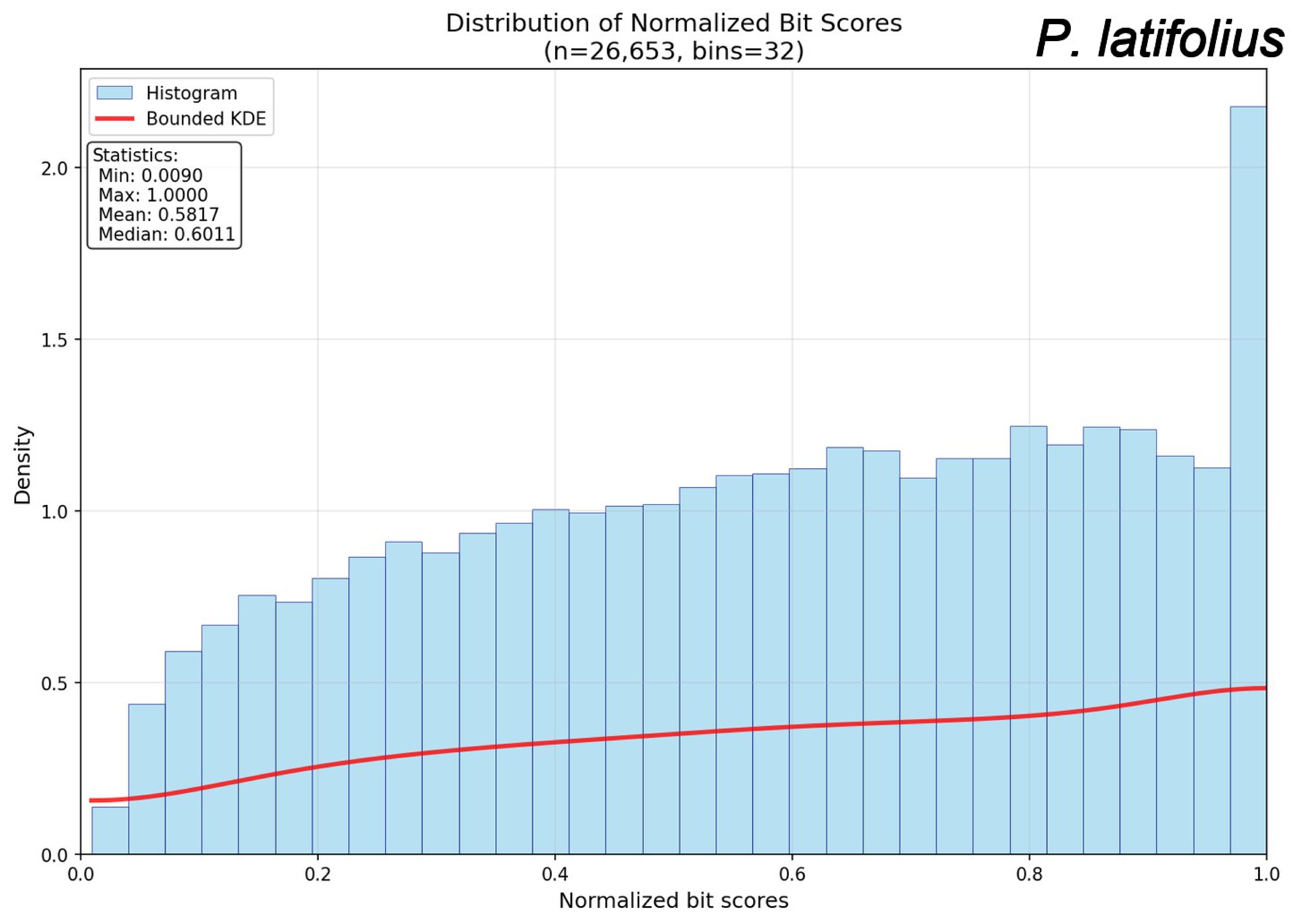

Distribution of Normalized Bit Scores
(n=26,653, bins=32)
P. latifolius
Histogram
Bounded KDE
Statistics:
Min: 0.0090
Max: 1.0000
Mean: 0.5817
Median: 0.6011
Density
0.0
0.5
1.0
1.5
2.0
0.0
0.2
0.4
0.6
0.8
1.0
Normalized bit scores

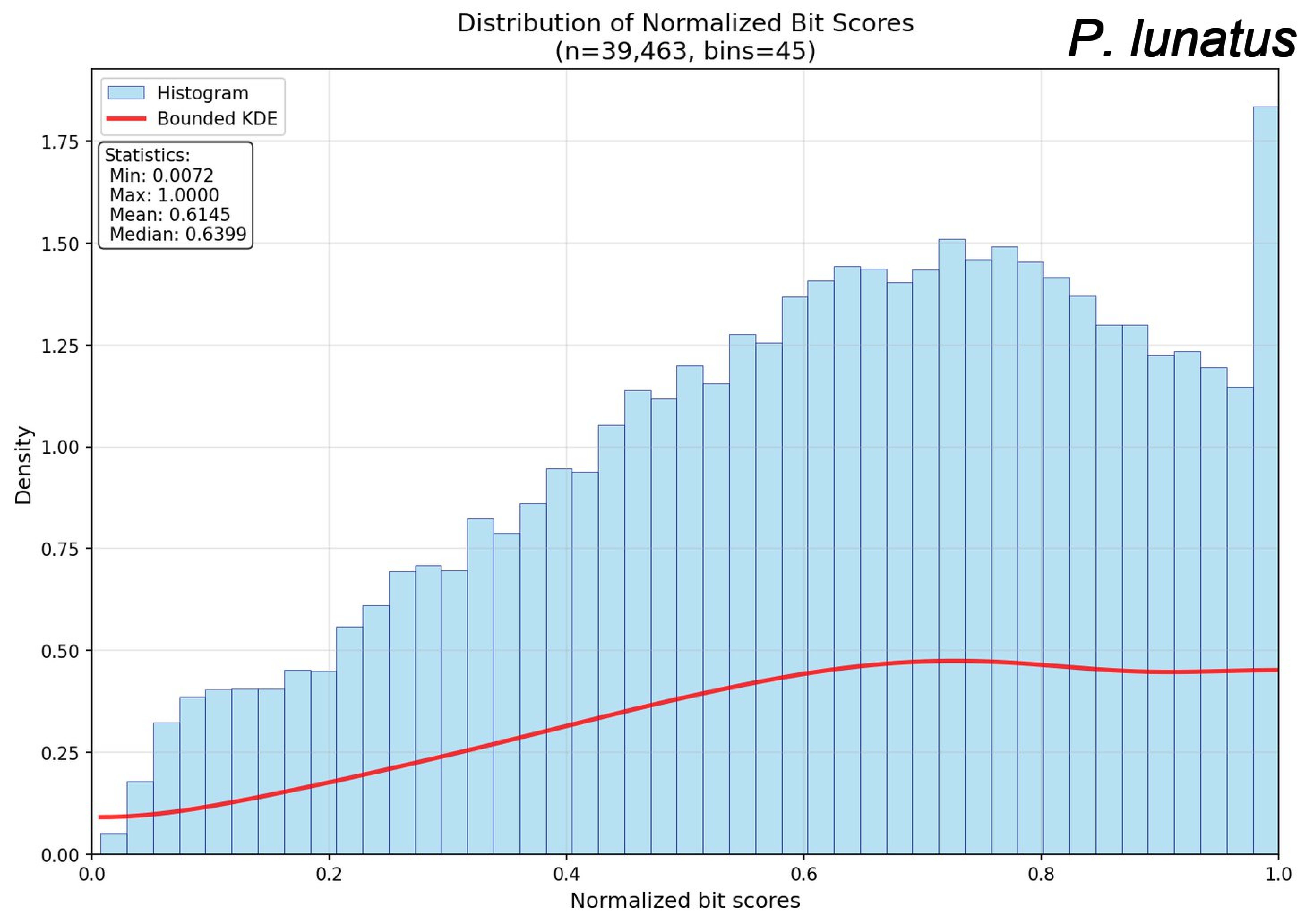
Distribution of Normalized Bit Scores
(n=39,463, bins=45)
P. lunatus
Histogram
Bounded KDE
Statistics:
Min: 0.0072
Max: 1.0000
Mean: 0.6145
Median: 0.6399
Density
Normalized bit scores
0.00
0.25
0.50
0.75
1.00
1.25
1.50
1.75
0.0
0.2
0.4
0.6
0.8
1.0

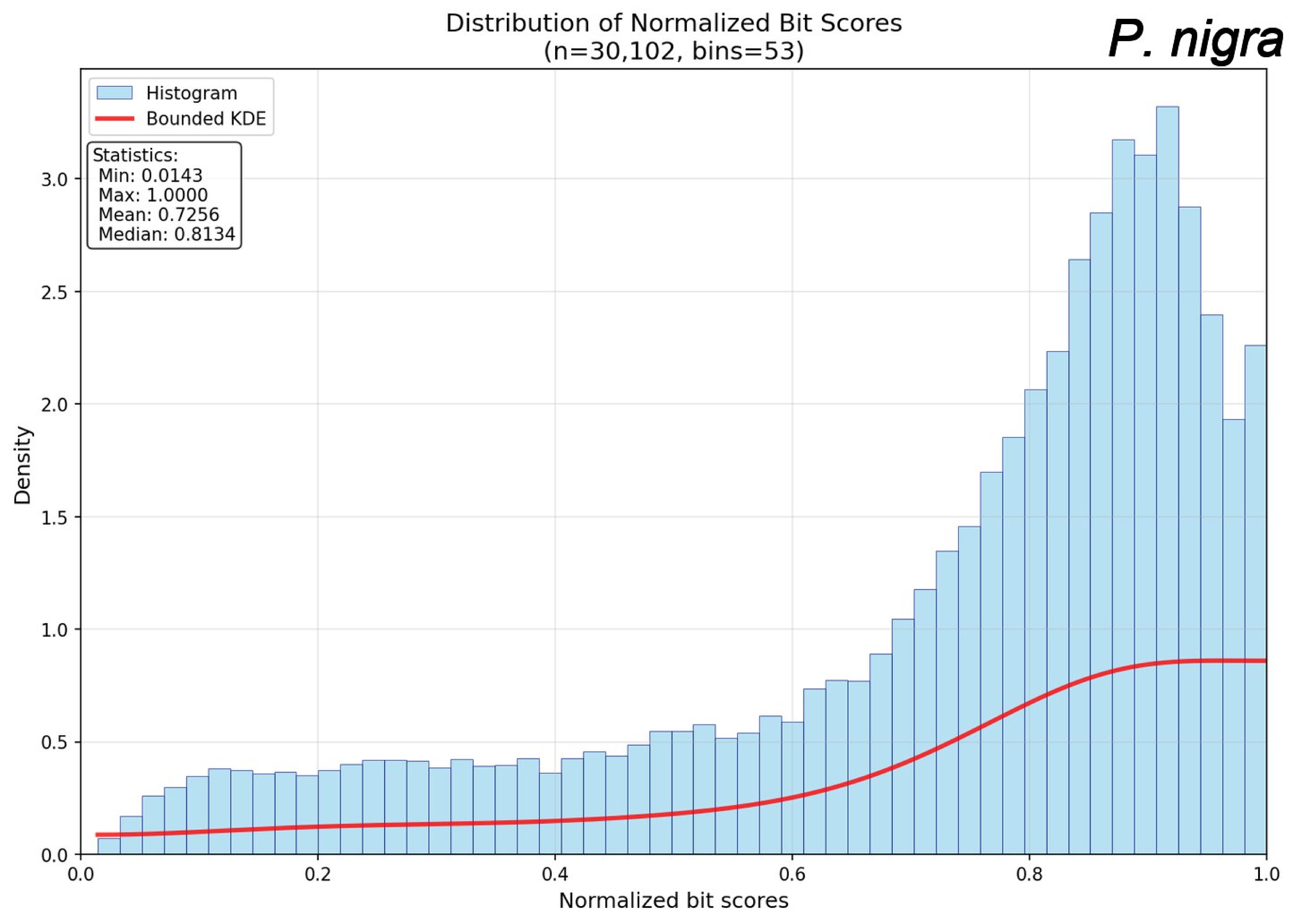
Distribution of Normalized Bit Scores
(n=30,102, bins=53)
P. nigra
Histogram
Bounded KDE
Statistics:
Min: 0.0143
Max: 1.0000
Mean: 0.7256
Median: 0.8134
Density
Normalized bit scores
0.0
0.5
1.0
1.5
2.0
2.5
3.0
0.0
0.2
0.4
0.6
0.8
1.0

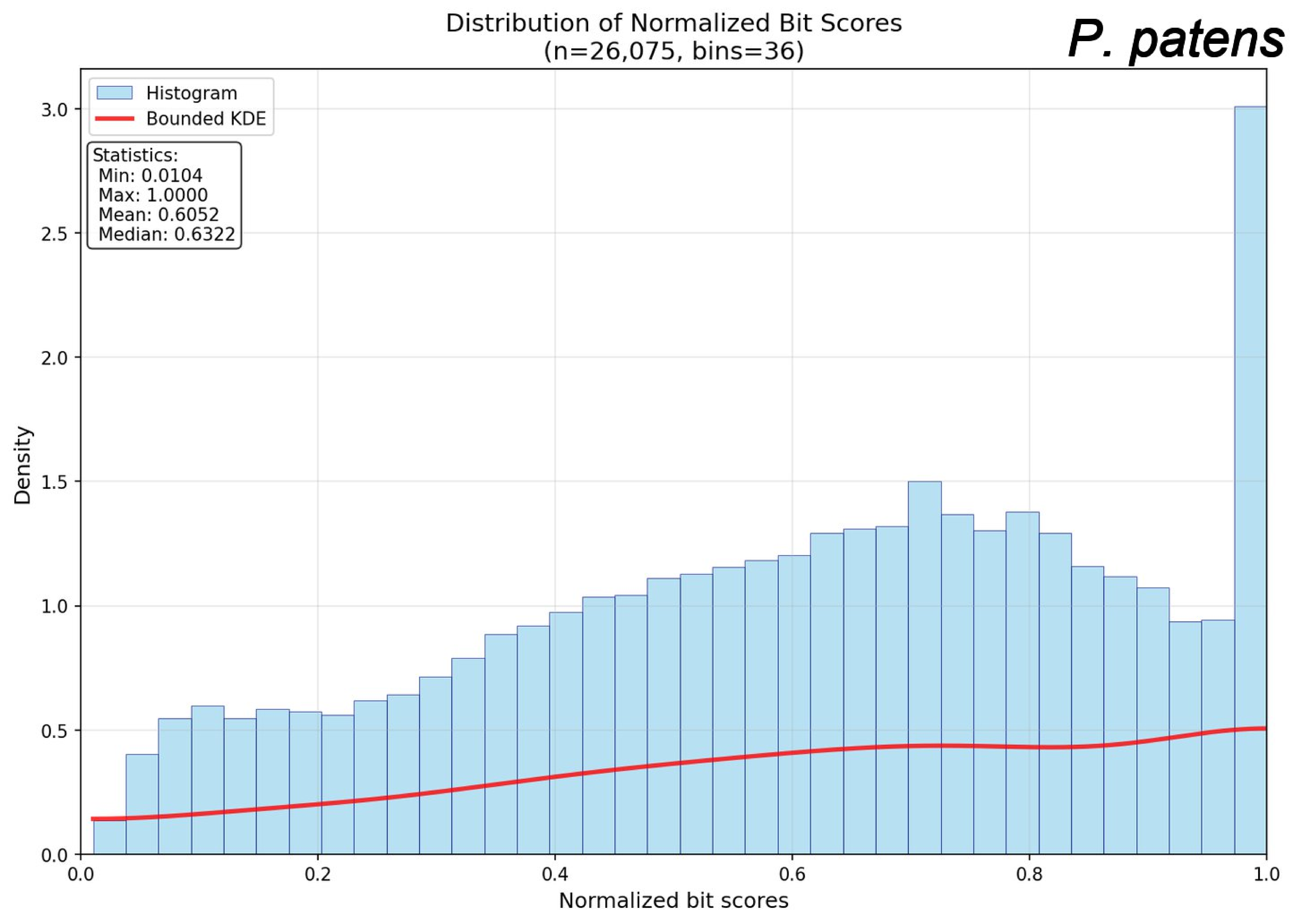
Distribution of Normalized Bit Scores
(n=26,075, bins=36)
P. patens
Histogram
Bounded KDE
Statistics:
Min: 0.0104
Max: 1.0000
Mean: 0.6052
Median: 0.6322
Density
Normalized bit scores
0.0
0.5
1.0
1.5
2.0
2.5
3.0
0.2
0.4
0.6
0.8

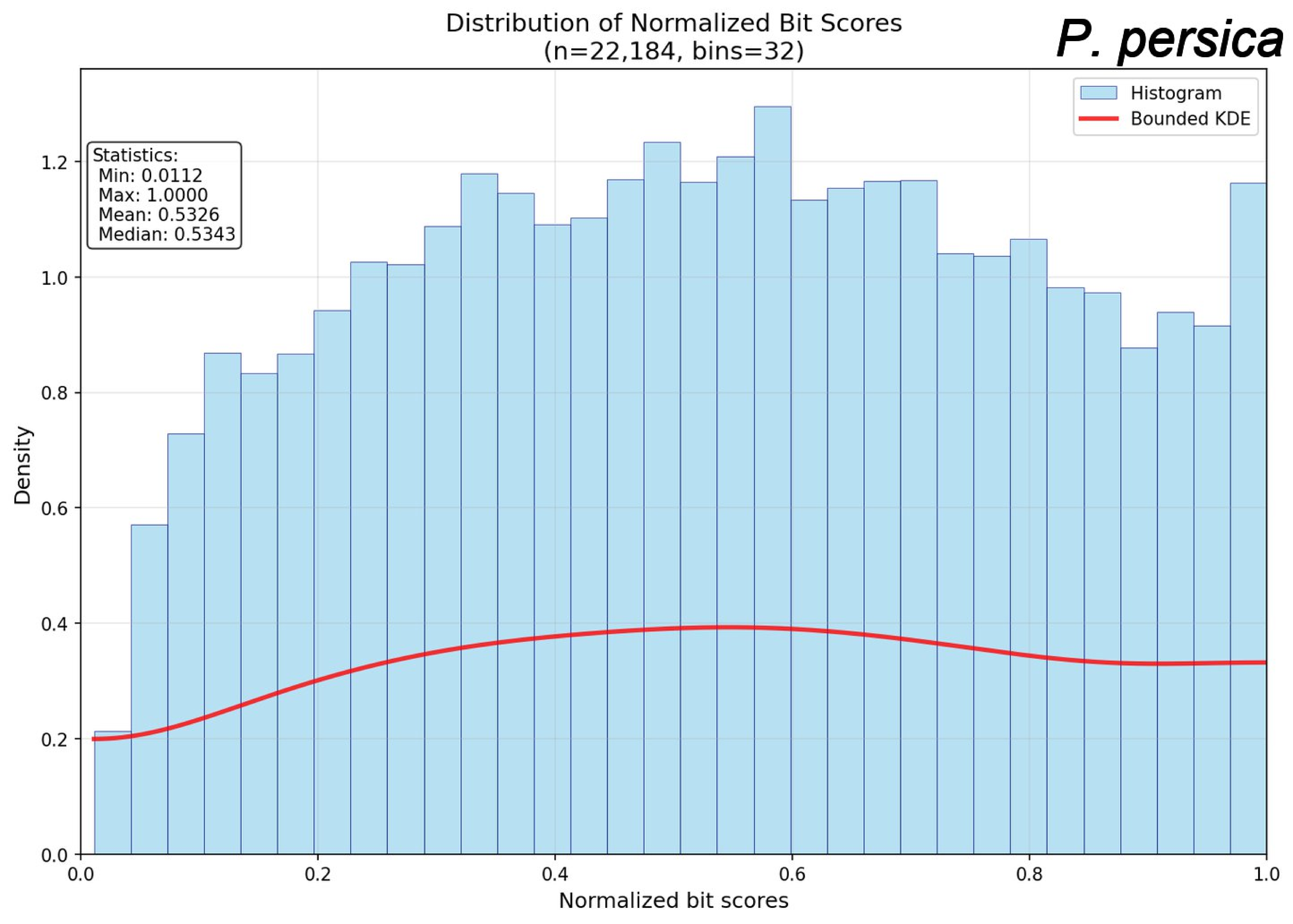

Distribution of Normalized Bit Scores
(n=22,184, bins=32)
P. persica
Histogram
Bounded KDE
Statistics:
Min: 0.0112
Max: 1.0000
Mean: 0.5326
Median: 0.5343
Density
Normalized bit scores
0.0
0.2
0.4
0.6
0.8
1.0
1.2

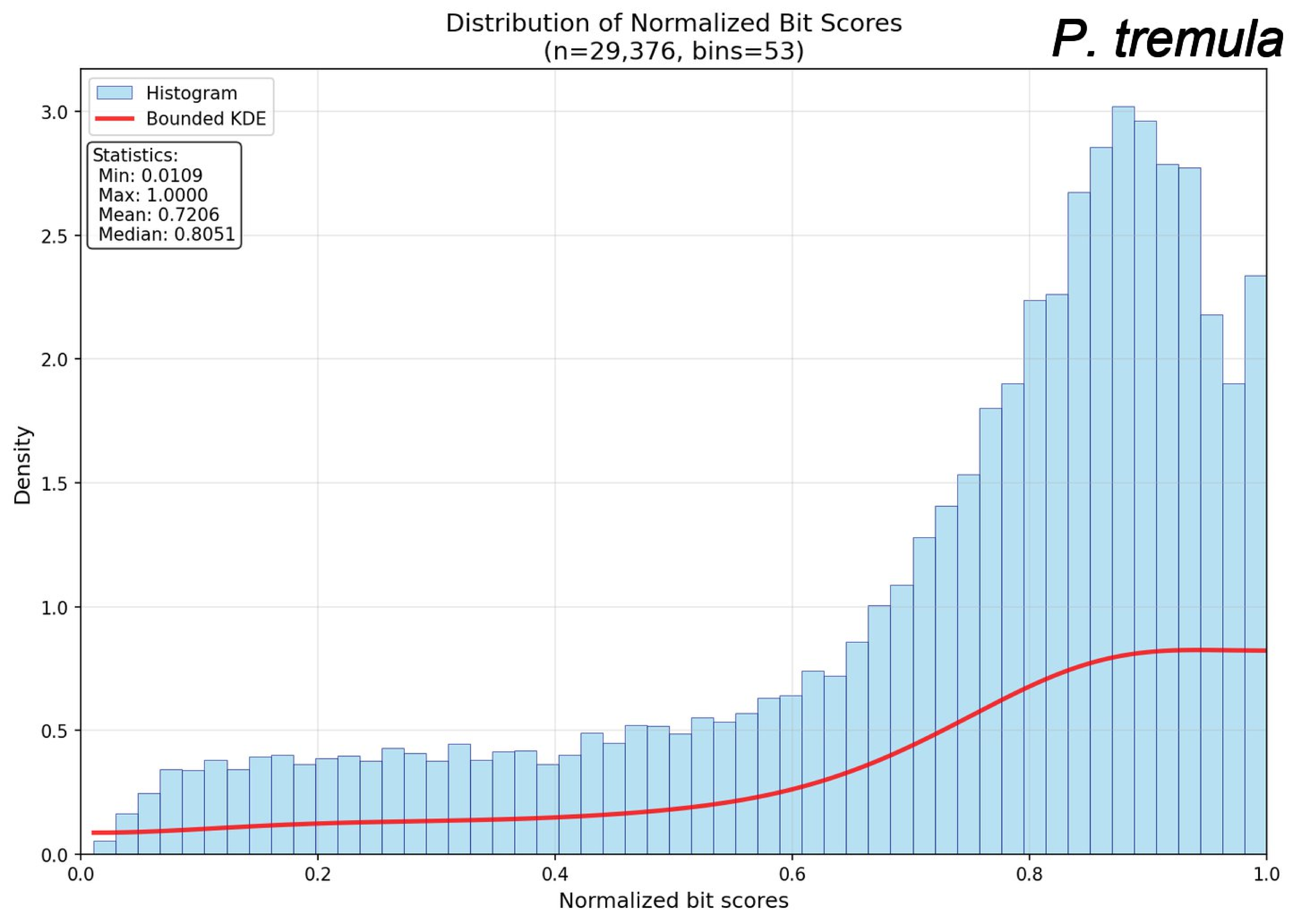
Distribution of Normalized Bit Scores
(n=29,376, bins=53)
P. tremula
Histogram
Bounded KDE
Statistics:
Min: 0.0109
Max: 1.0000
Mean: 0.7206
Median: 0.8051
Density
Normalized bit scores
0.0
0.5
1.0
1.5
2.0
2.5
3.0
0.0
0.2
0.4
0.6
0.8
1.0

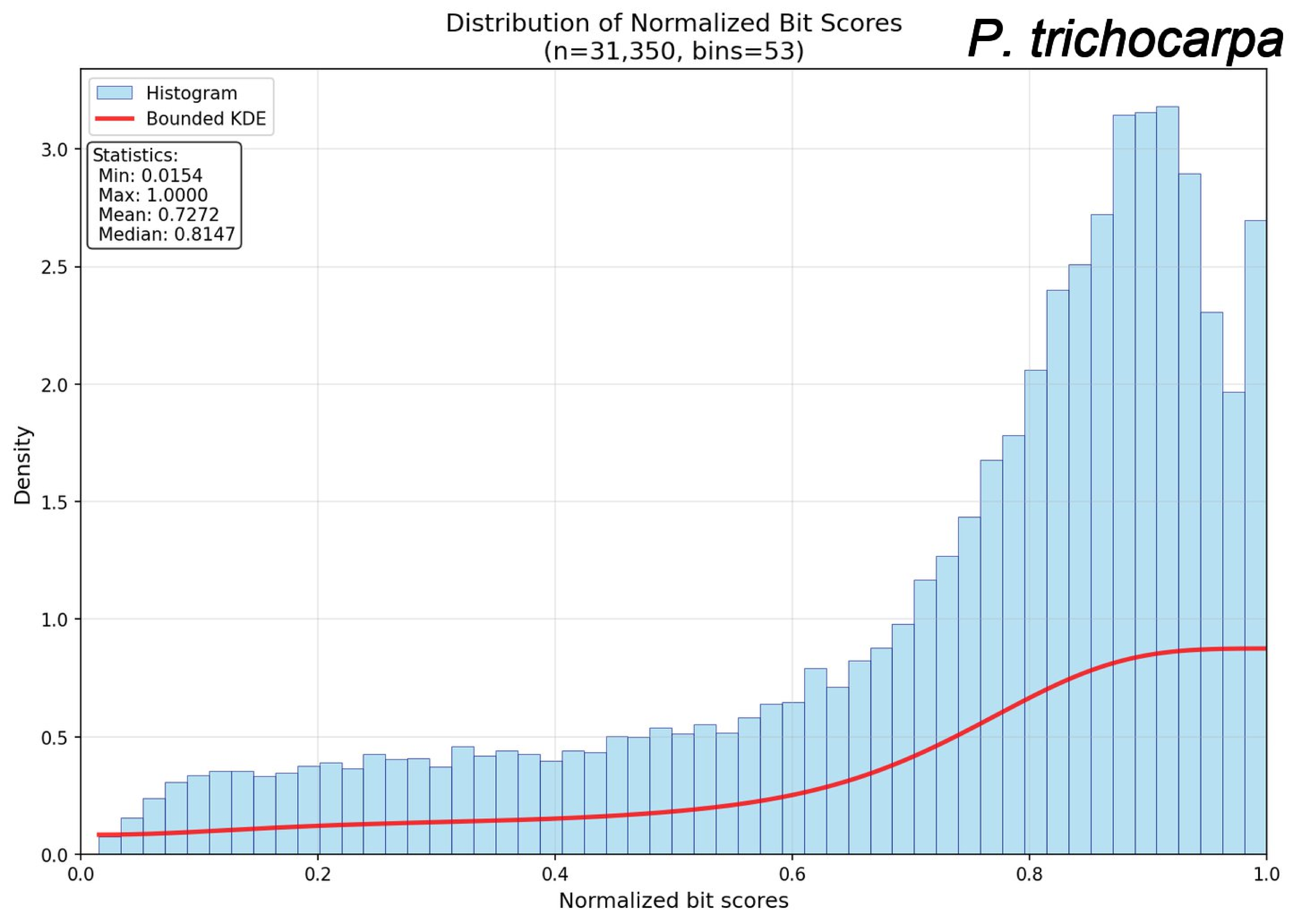
Distribution of Normalized Bit Scores
(n=31,350, bins=53)
P. trichocarpa
Histogram
Bounded KDE
Statistics:
Min: 0.0154
Max: 1.0000
Mean: 0.7272
Median: 0.8147
Density
Normalized bit scores
0.0
0.5
1.0
1.5
2.0
2.5
3.0
0.2
0.4
0.6
0.8

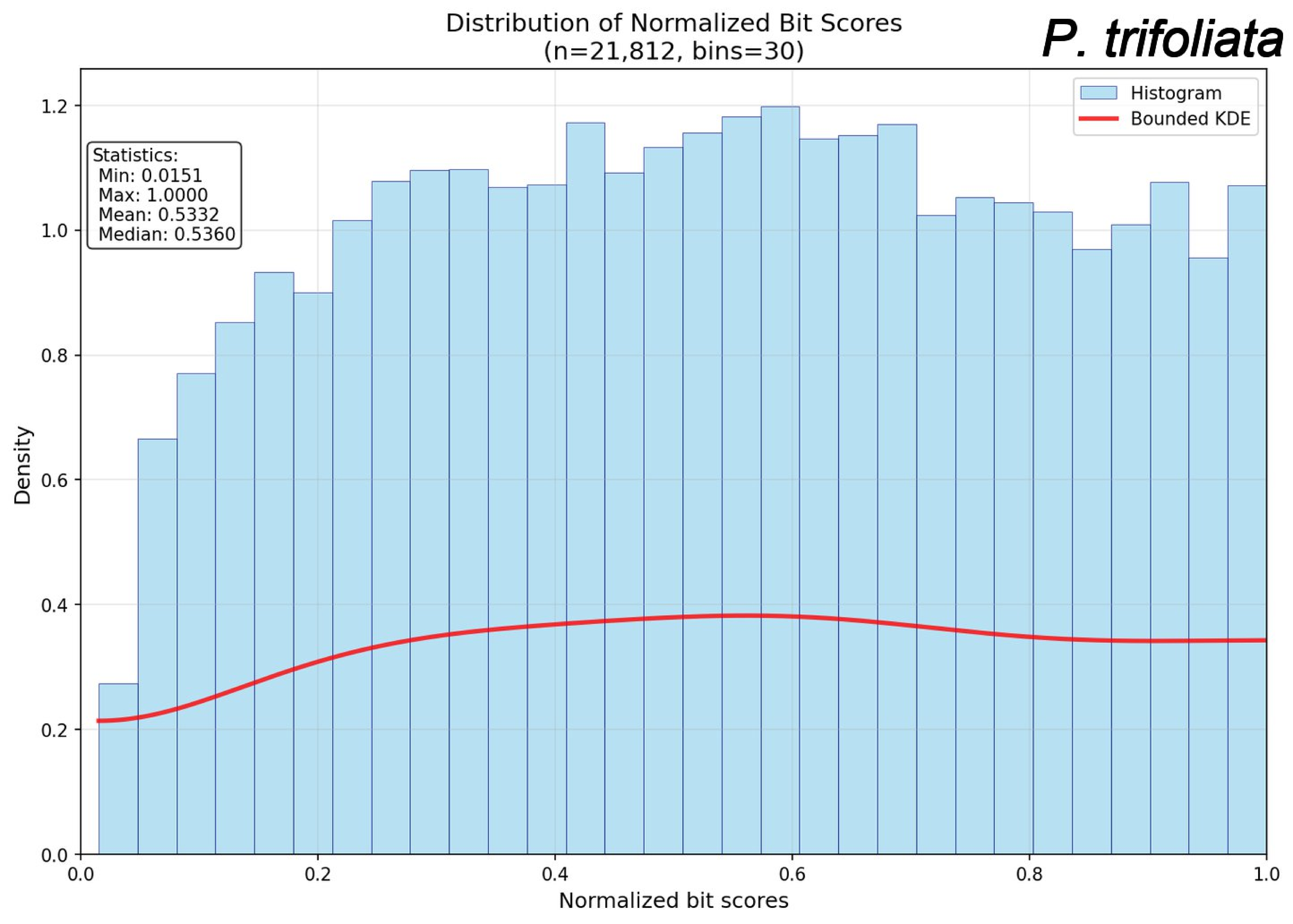
Distribution of Normalized Bit Scores
(n=21,812, bins=30)
P. trifoliata
Histogram
Bounded KDE
Statistics:
Min: 0.0151
Max: 1.0000
Mean: 0.5332
Median: 0.5360
Density
Normalized bit scores
0.0
0.2
0.4
0.6
0.8
1.0
1.2

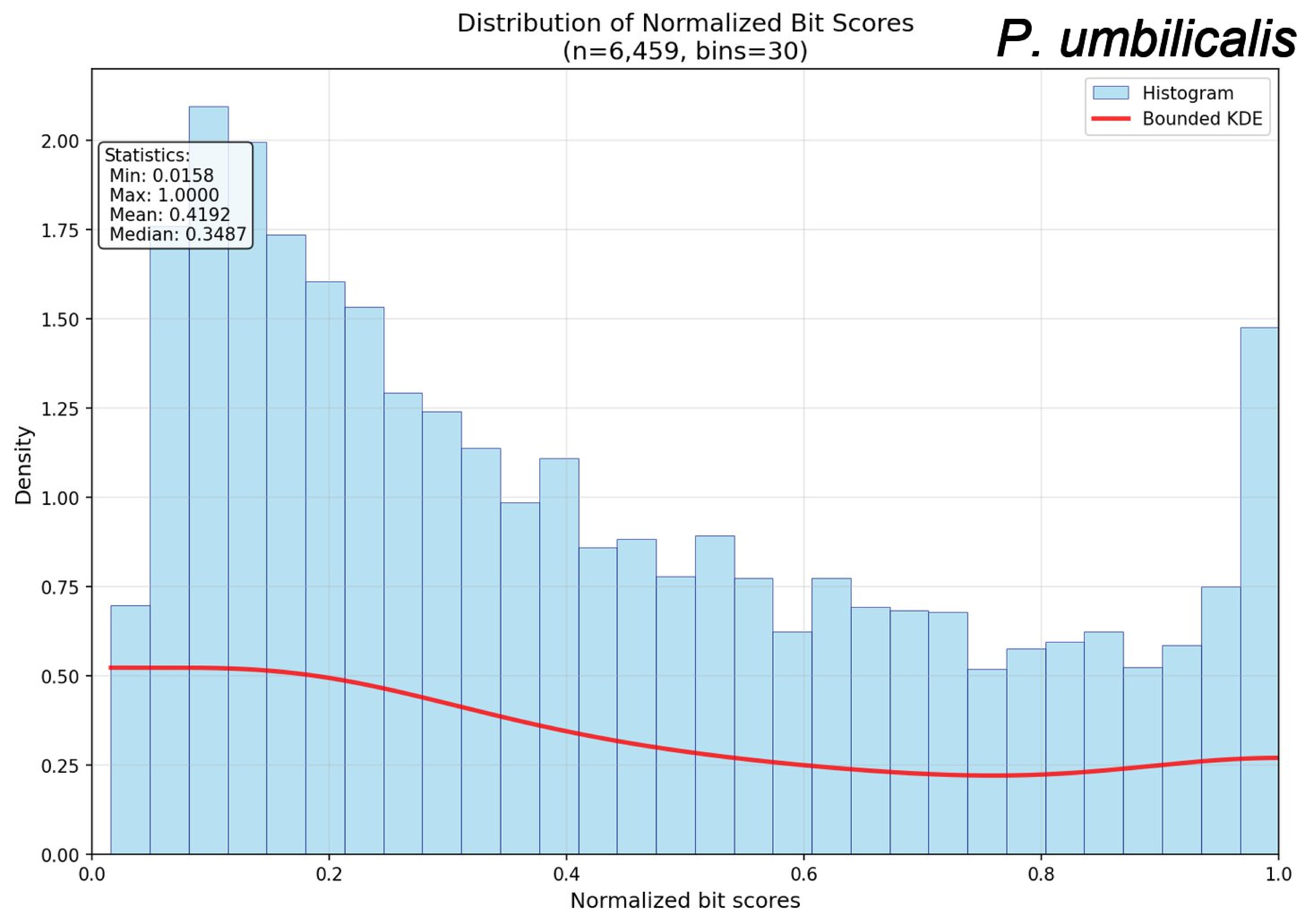
Distribution of Normalized Bit Scores
(n=6,459, bins=30)
P. umbilicalis
Histogram
Bounded KDE
Statistics:
Min: 0.0158
Max: 1.0000
Mean: 0.4192
Median: 0.3487
Density
Normalized bit scores
0.00
0.25
0.50
0.75
1.00
1.25
1.50
1.75
2.00
0.0
0.2
0.4
0.6
0.8
1.0

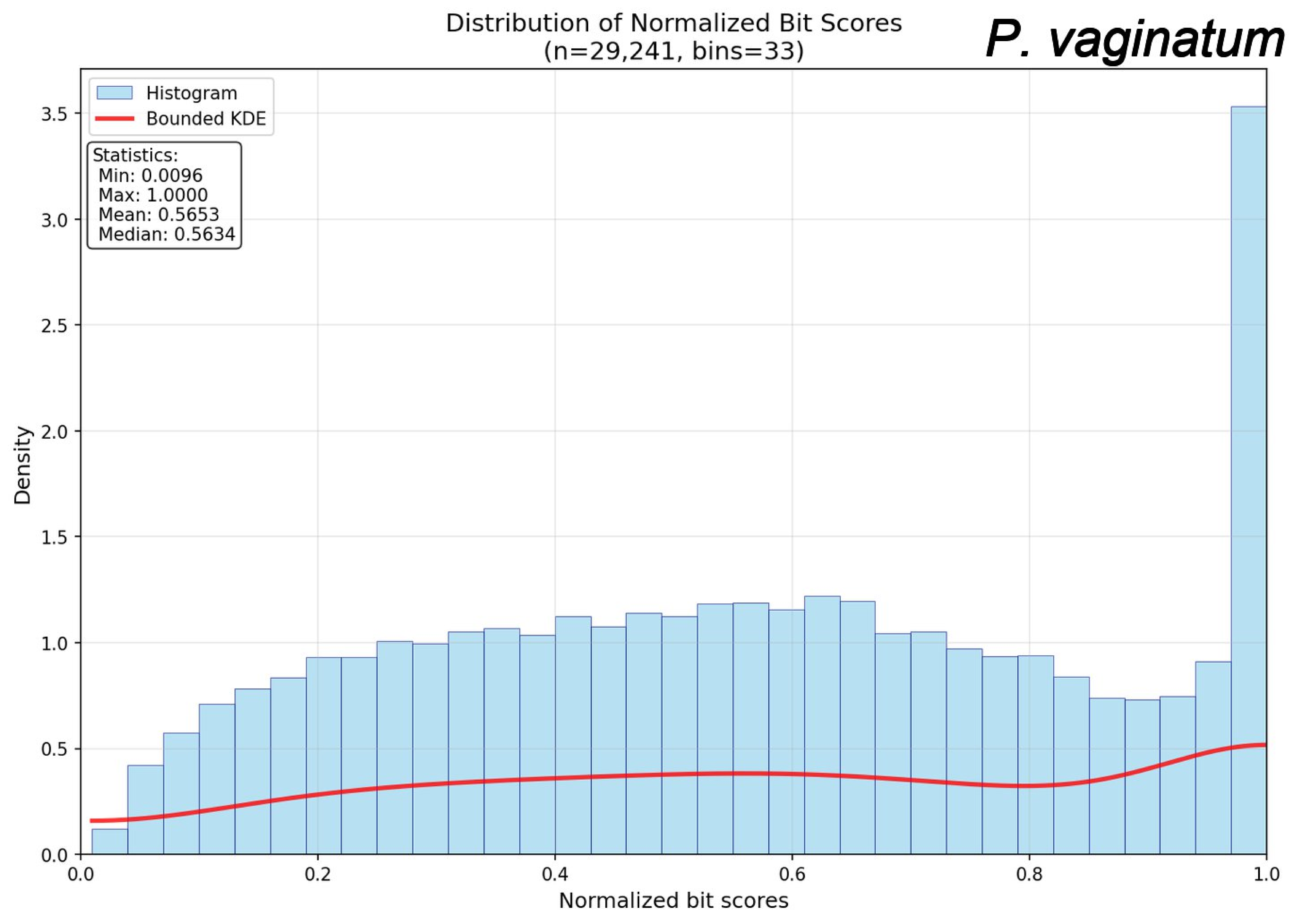
Distribution of Normalized Bit Scores
(n=29,241, bins=33)
P. vaginatum
Histogram
Bounded KDE
Statistics:
Min: 0.0096
Max: 1.0000
Mean: 0.5653
Median: 0.5634
Density
Normalized bit scores
0.0
0.5
1.0
1.5
2.0
2.5
3.0
3.5
0.0
0.2
0.4
0.6
0.8
1.0

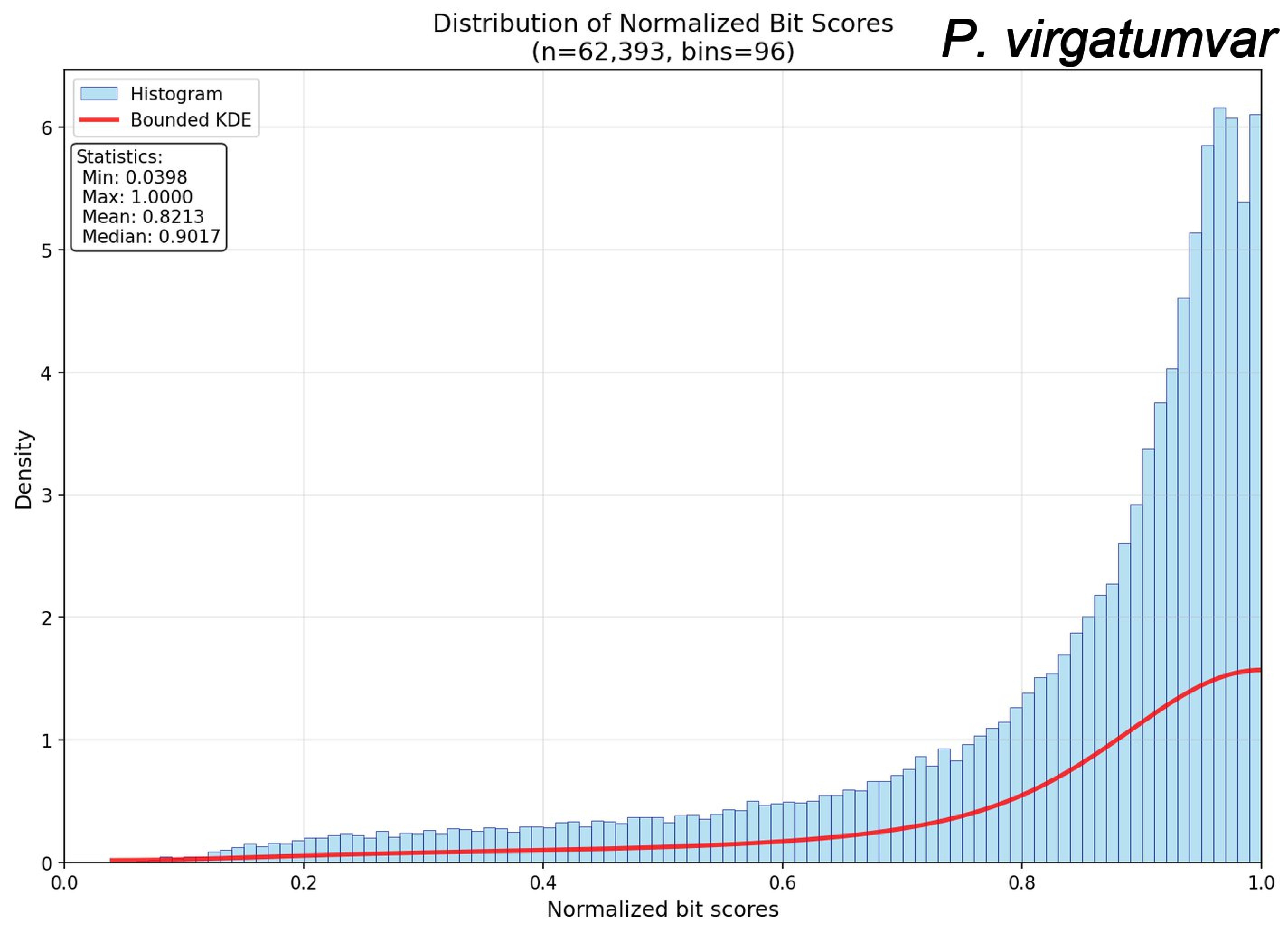
Distribution of Normalized Bit Scores
(n=62,393, bins=96)
P. virgatumvar
Histogram
Bounded KDE
Statistics:
Min: 0.0398
Max: 1.0000
Mean: 0.8213
Median: 0.9017
Density
Normalized bit scores
0
1
2
3
4
5
6
0.0
0.2
0.4
0.6
0.8
1.0

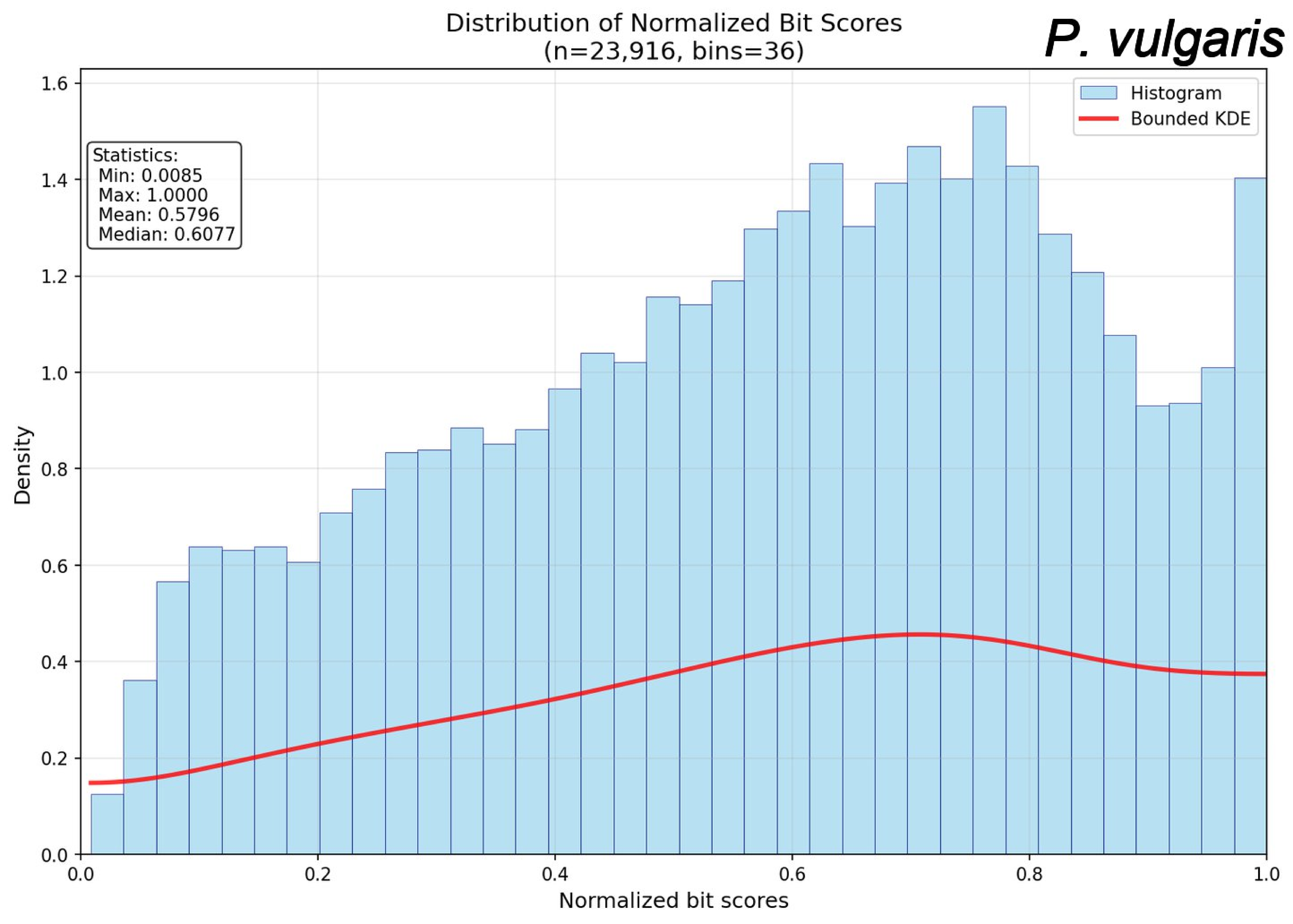
Distribution of Normalized Bit Scores
(n=23,916, bins=36)
P. vulgaris
Histogram
Bounded KDE
Statistics:
Min: 0.0085
Max: 1.0000
Mean: 0.5796
Median: 0.6077
Density
Normalized bit scores
0.0
0.2
0.4
0.6
0.8
1.0
1.2
1.4
1.6

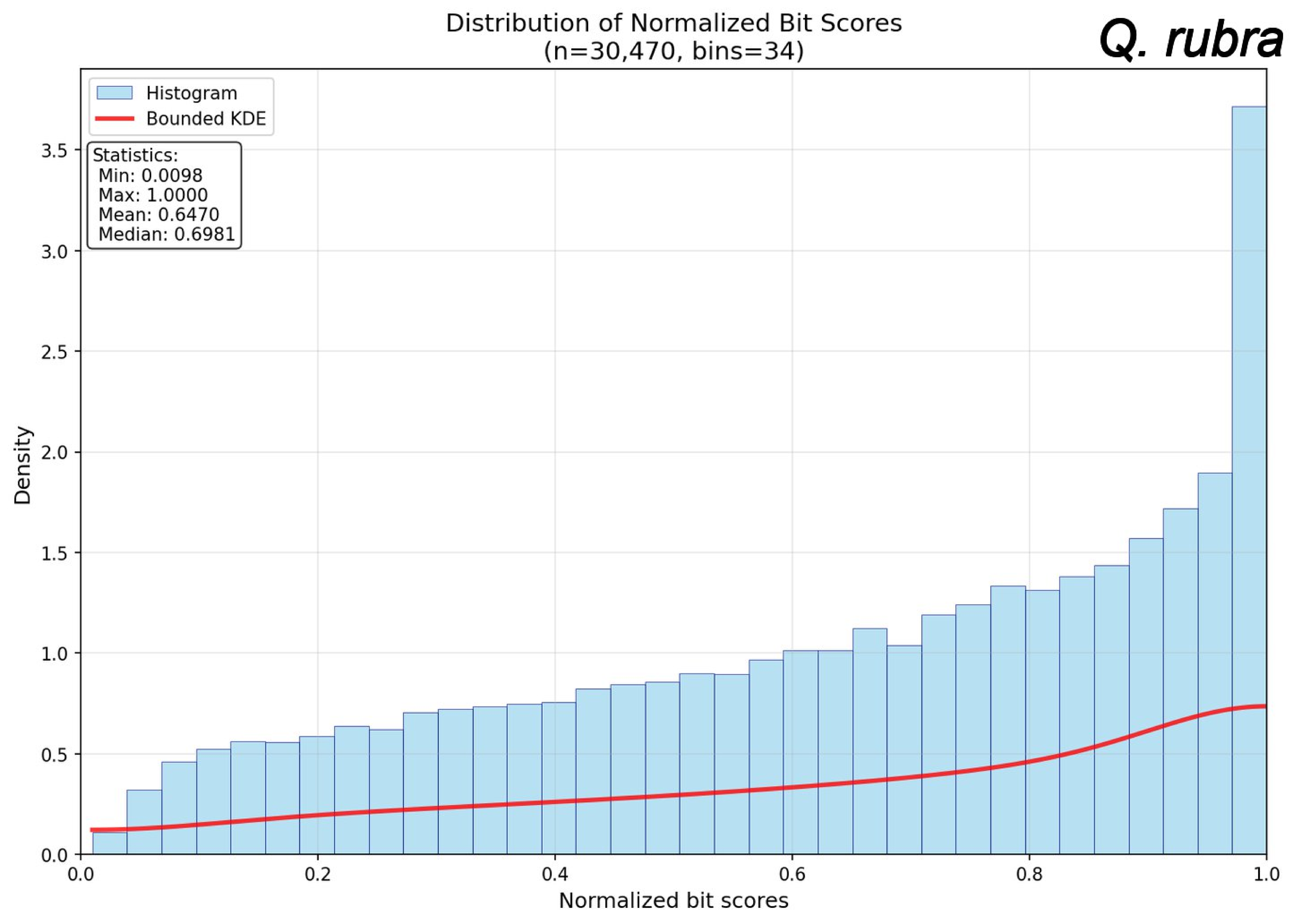
Distribution of Normalized Bit Scores
(n=30,470, bins=34)
Q. rubra
Histogram
Bounded KDE
Statistics:
Min: 0.0098
Max: 1.0000
Mean: 0.6470
Median: 0.6981
Density
Normalized bit scores
0.0
0.5
1.0
1.5
2.0
2.5
3.0
3.5
0.0
0.2
0.4
0.6
0.8
1.0

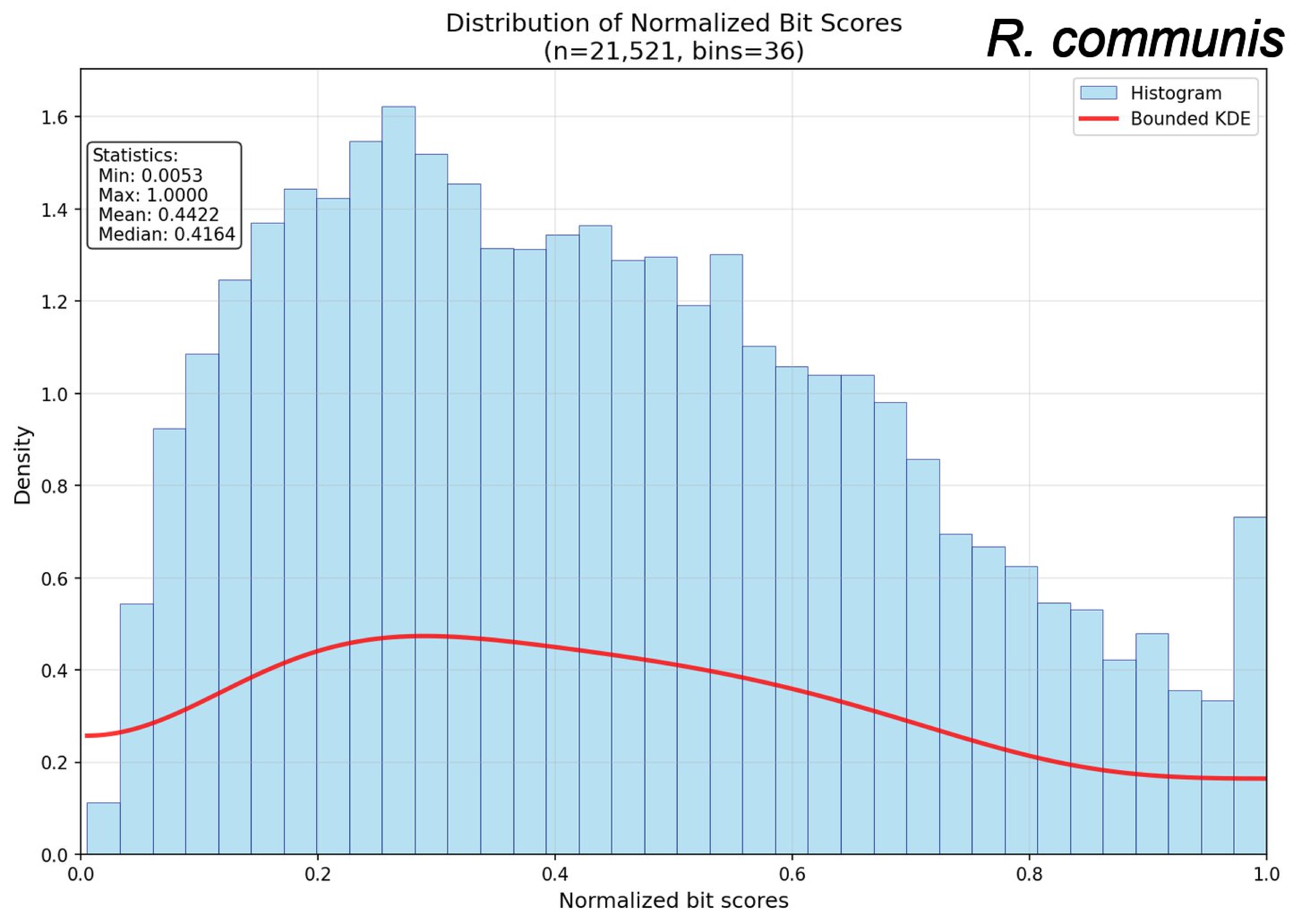

Distribution of Normalized Bit Scores
(n=21,521, bins=36)
R. communis
Histogram
Bounded KDE
Statistics:
Min: 0.0053
Max: 1.0000
Mean: 0.4422
Median: 0.4164
Density
Normalized bit scores
0.0
0.2
0.4
0.6
0.8
1.0
1.2
1.4
1.6

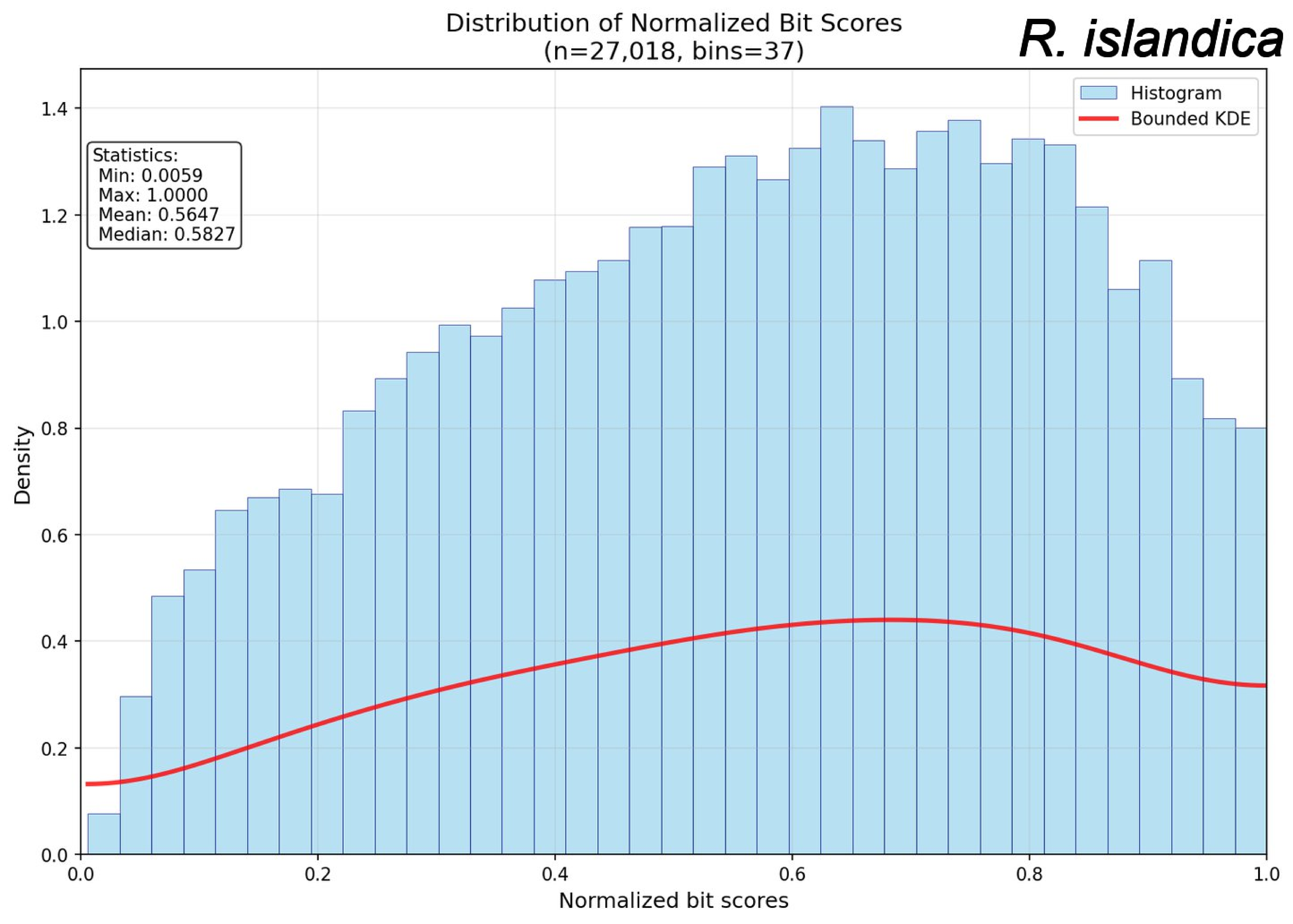
Distribution of Normalized Bit Scores
(n=27,018, bins=37)
R. islandica
Histogram
Bounded KDE
Statistics:
Min: 0.0059
Max: 1.0000
Mean: 0.5647
Median: 0.5827
Density
Normalized bit scores
0.0
0.2
0.4
0.6
0.8
1.0
1.2
1.4

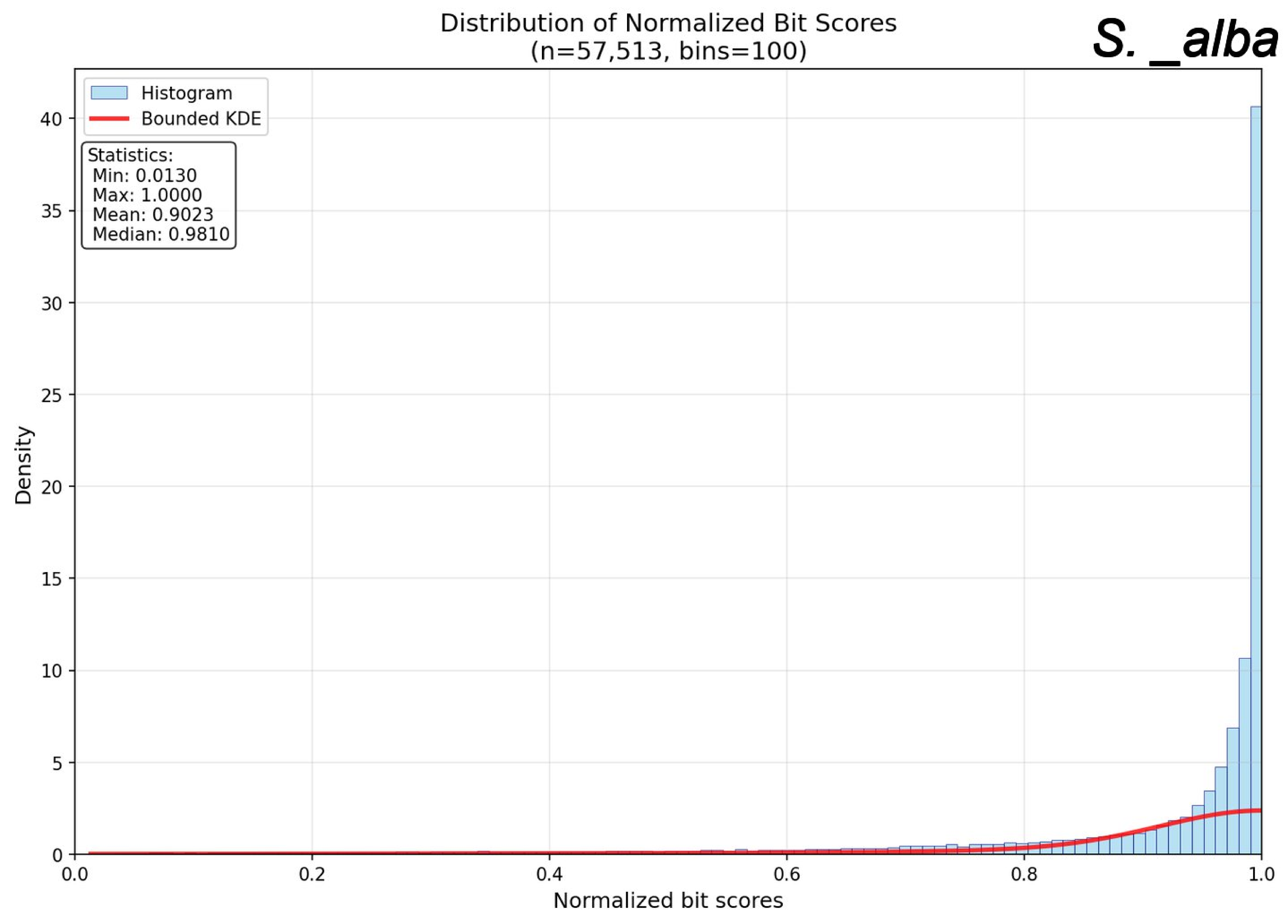
Distribution of Normalized Bit Scores
(n=57,513, bins=100)
S. _alba
Histogram
Bounded KDE
Statistics:
Min: 0.0130
Max: 1.0000
Mean: 0.9023
Median: 0.9810
Density
0
5
10
15
20
25
30
35
40
0.0
0.2
0.4
0.6
0.8
1.0
Normalized bit scores

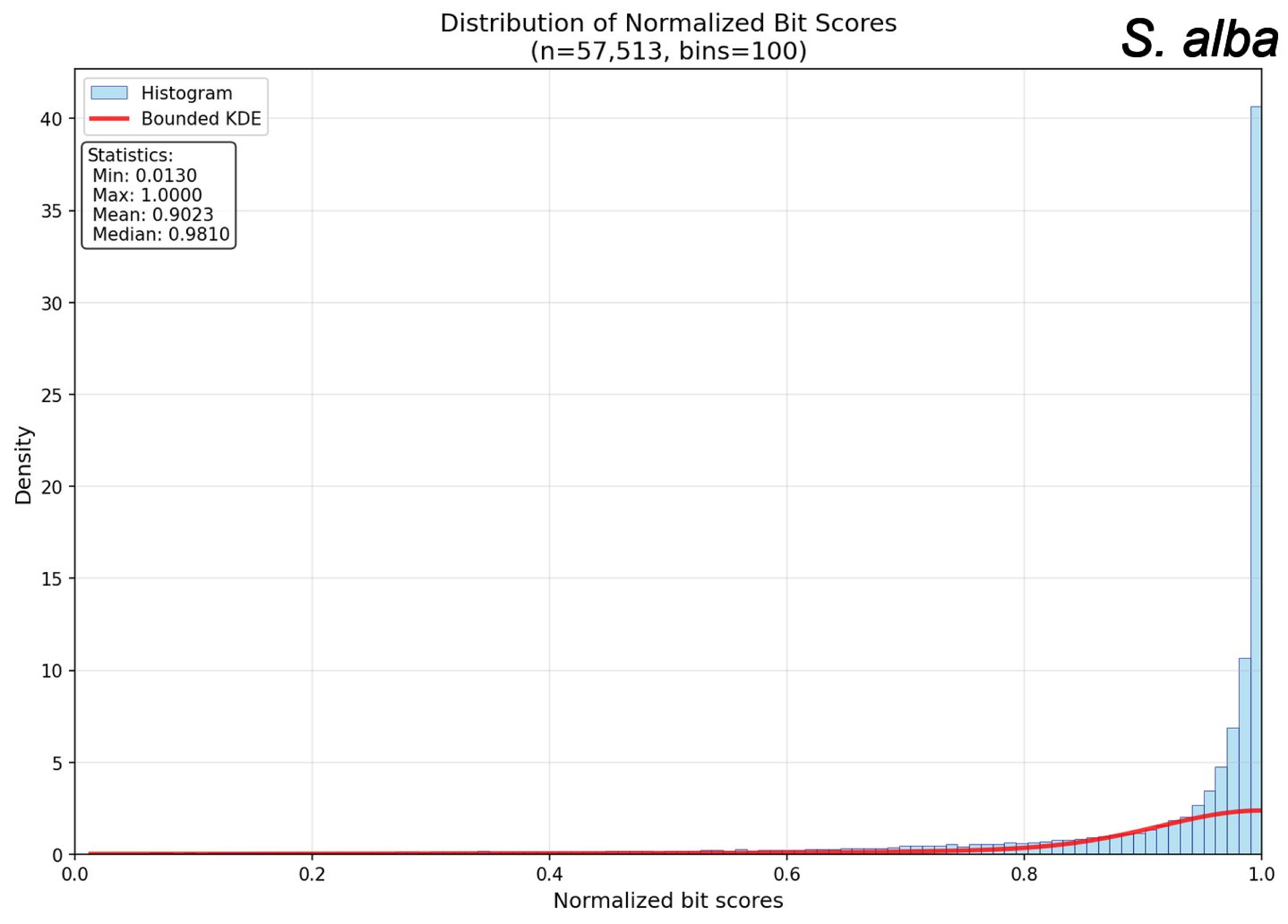
Distribution of Normalized Bit Scores
(n=57,513, bins=100)
S. alba
Histogram
Bounded KDE
Statistics:
Min: 0.0130
Max: 1.0000
Mean: 0.9023
Median: 0.9810
Density
Normalized bit scores
0
5
10
15
20
25
30
35
40
0.0
0.2
0.4
0.6
0.8
1.0

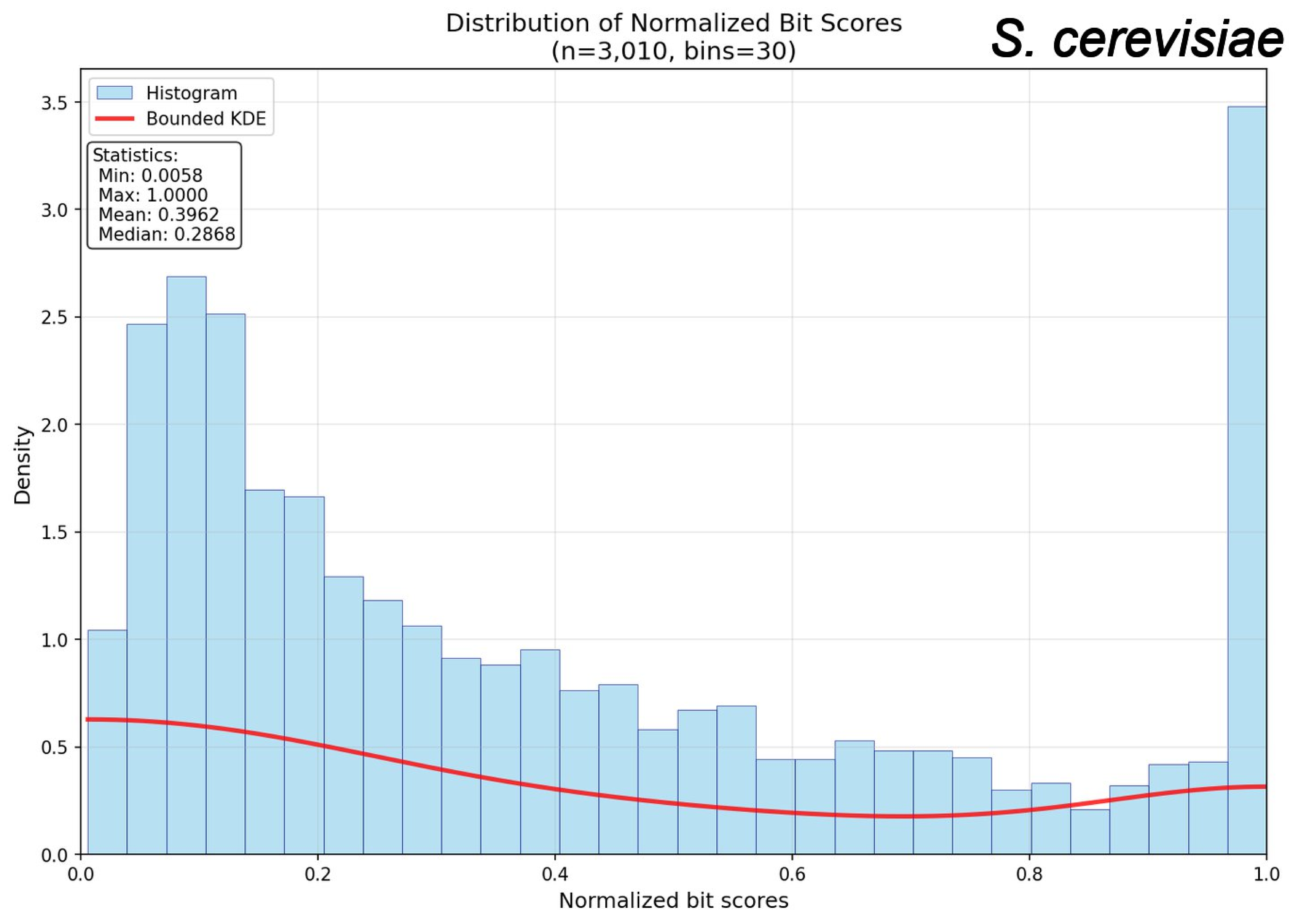
S. cerevisiae
Distribution of Normalized Bit Scores
(n=3,010, bins=30)
Histogram
Bounded KDE
Statistics:
Min: 0.0058
Max: 1.0000
Mean: 0.3962
Median: 0.2868
Density
Normalized bit scores
0.0
0.5
1.0
1.5
2.0
2.5
3.0
3.5
0.0
0.2
0.4
0.6
0.8
1.0

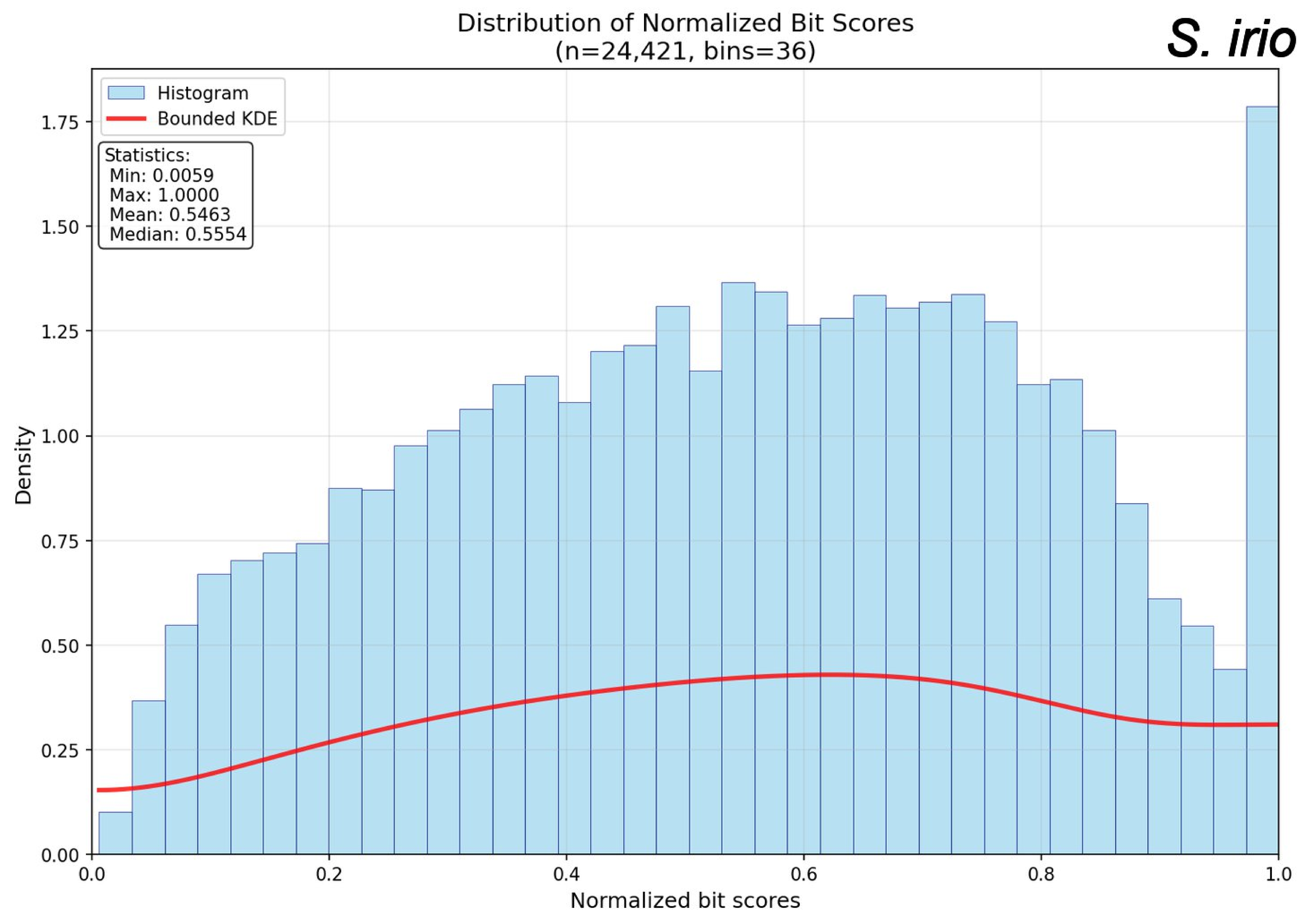
Distribution of Normalized Bit Scores
(n=24,421, bins=36)
S. irio
Histogram
Bounded KDE
Statistics:
Min: 0.0059
Max: 1.0000
Mean: 0.5463
Median: 0.5554
Density
Normalized bit scores
0.00
0.25
0.50
0.75
1.00
1.25
1.50
1.75
0.0
0.2
0.4
0.6
0.8
1.0

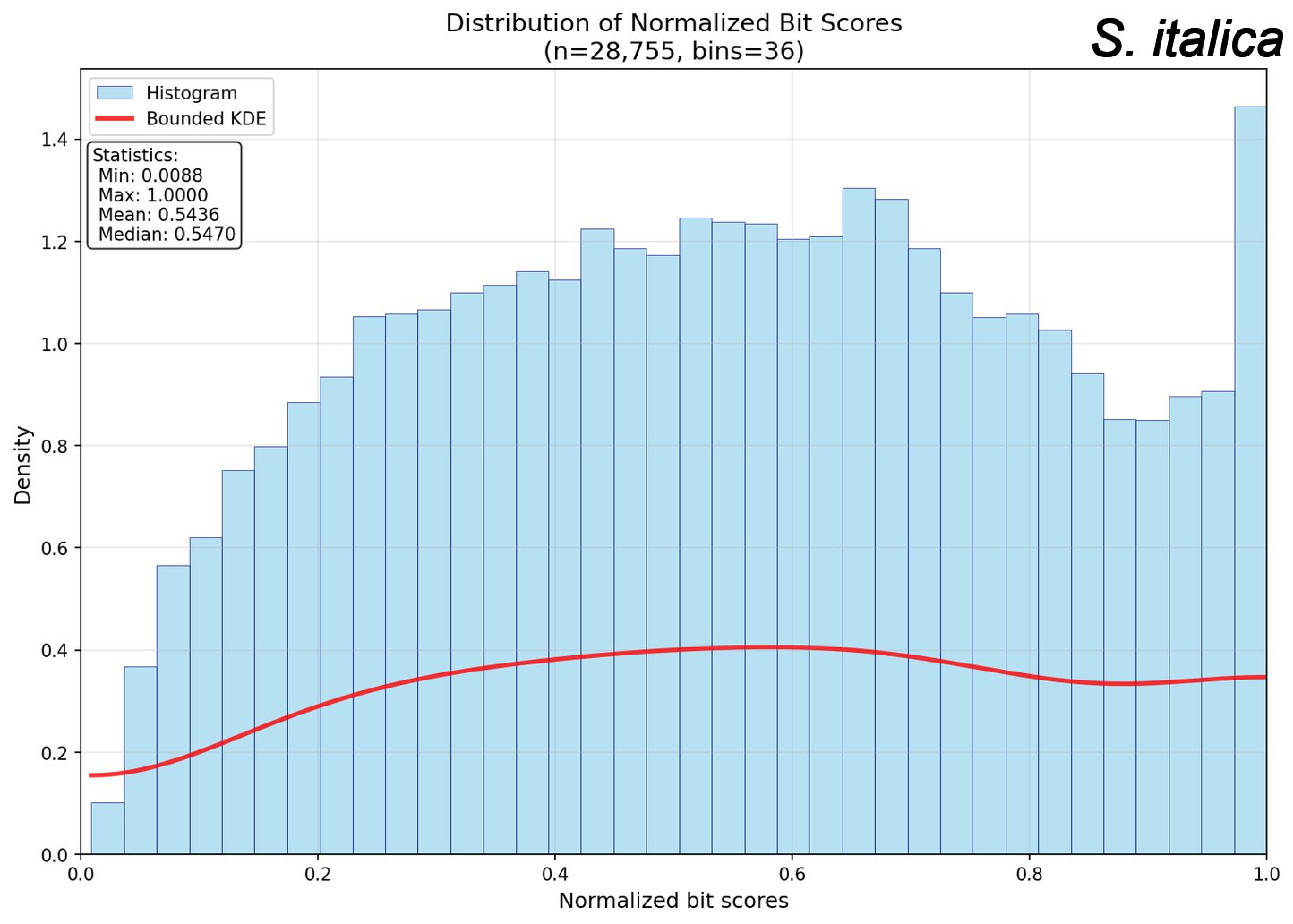
Distribution of Normalized Bit Scores
(n=28,755, bins=36)
S. italica
Histogram
Bounded KDE
Statistics:
Min: 0.0088
Max: 1.0000
Mean: 0.5436
Median: 0.5470
Density
Normalized bit scores
0.0
0.2
0.4
0.6
0.8
1.0
1.2
1.4

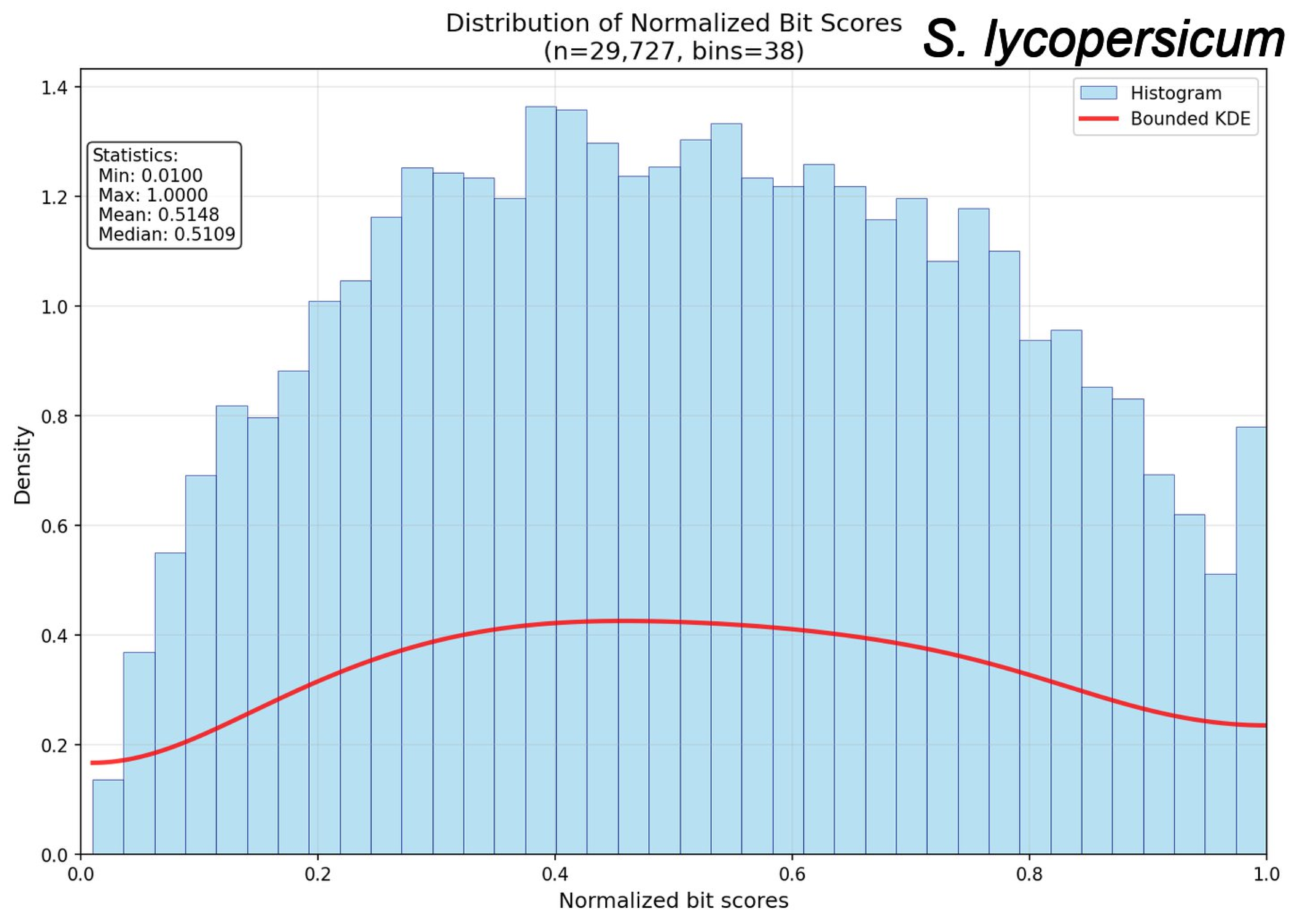
Distribution of Normalized Bit Scores
(n=29,727, bins=38)
S. lycopersicum
Histogram
Bounded KDE
Statistics:
Min: 0.0100
Max: 1.0000
Mean: 0.5148
Median: 0.5109
Density
Normalized bit scores
0.0
0.2
0.4
0.6
0.8
1.0
1.2
1.4

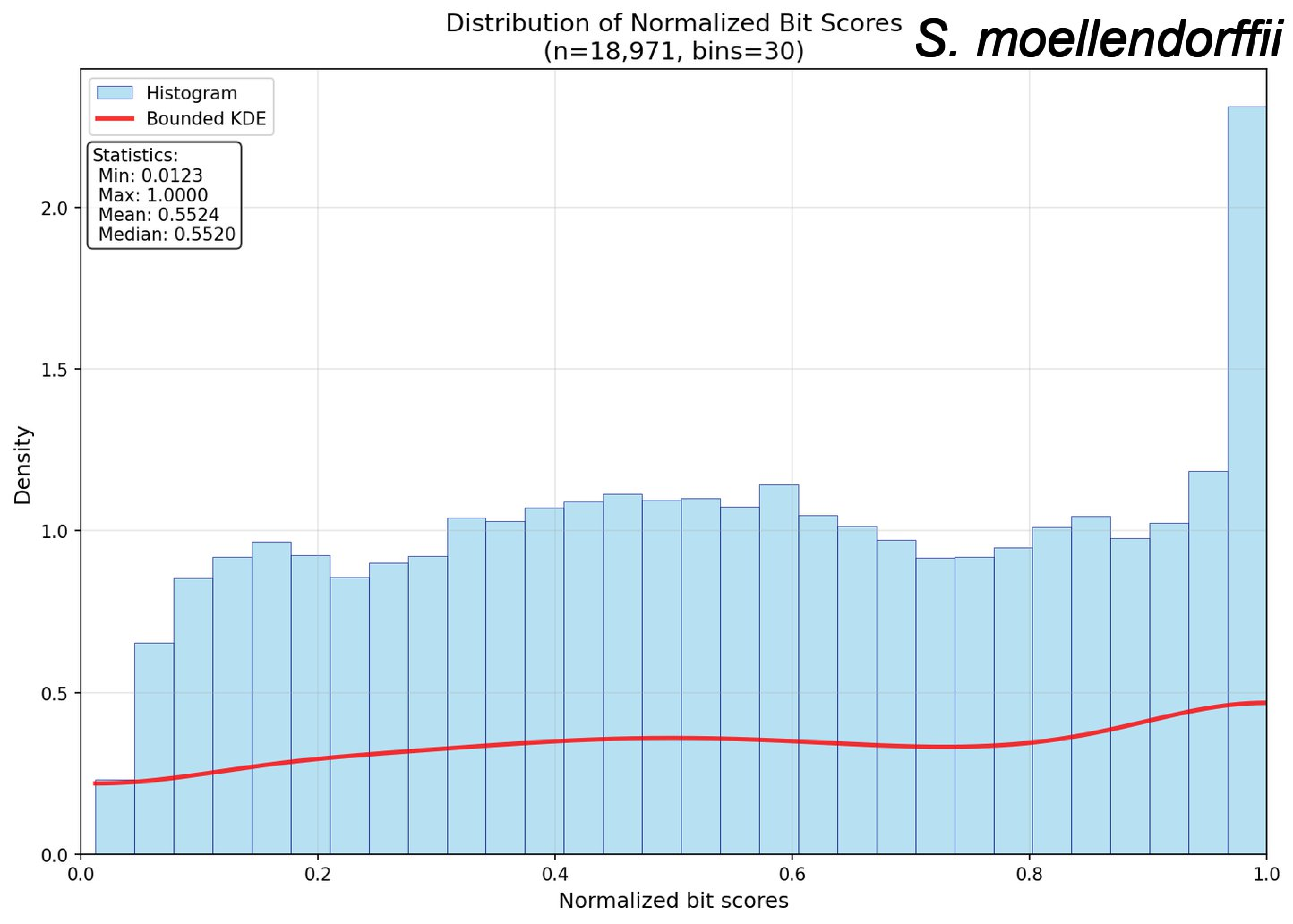
Distribution of Normalized Bit Scores
(n=18,971, bins=30)
S. moellendorffii
Histogram
Bounded KDE
Statistics:
Min: 0.0123
Max: 1.0000
Mean: 0.5524
Median: 0.5520
Density
Normalized bit scores
0.0
0.5
1.0
1.5
2.0
0.2
0.4
0.6
0.8

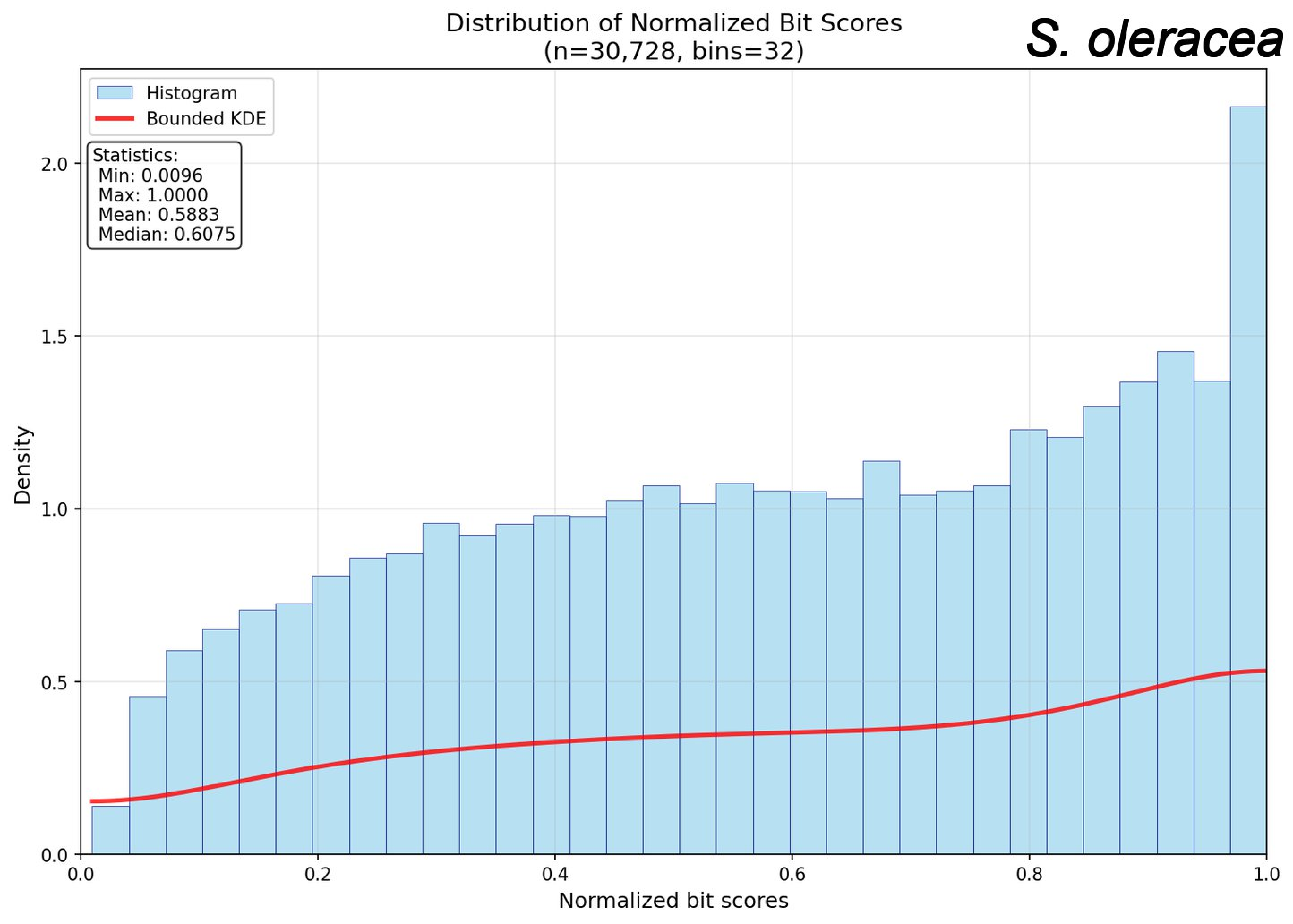

Distribution of Normalized Bit Scores
(n=30,728, bins=32)
S. oleracea
Histogram
Bounded KDE
Statistics:
Min: 0.0096
Max: 1.0000
Mean: 0.5883
Median: 0.6075
Density
Normalized bit scores
0.0
0.5
1.0
1.5
2.0
0.0
0.2
0.4
0.6
0.8
1.0

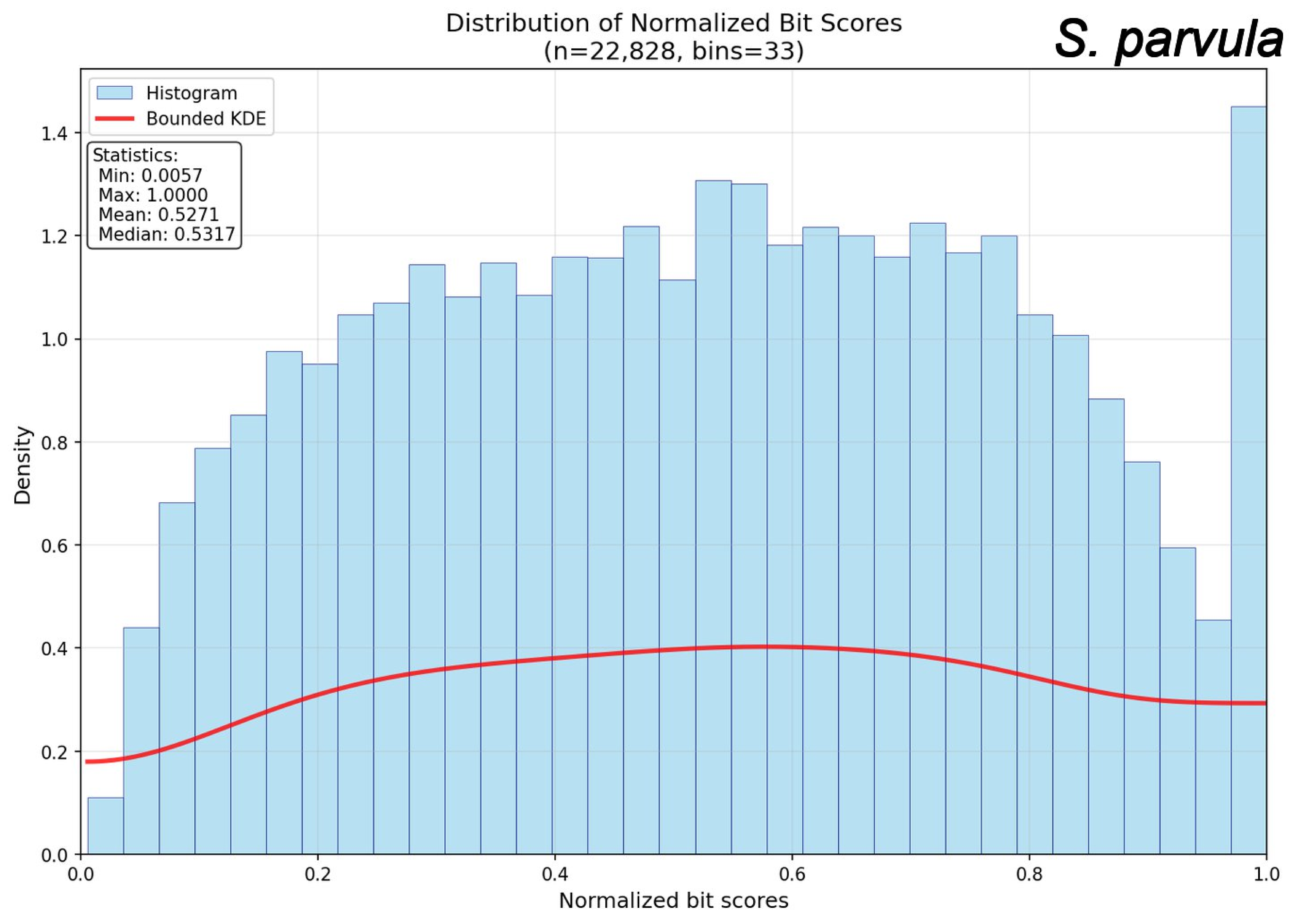
Distribution of Normalized Bit Scores
(n=22,828, bins=33)
S. parvula
Histogram
Bounded KDE
Statistics:
Min: 0.0057
Max: 1.0000
Mean: 0.5271
Median: 0.5317
Density
Normalized bit scores
0.0
0.2
0.4
0.6
0.8
1.0
1.2
1.4

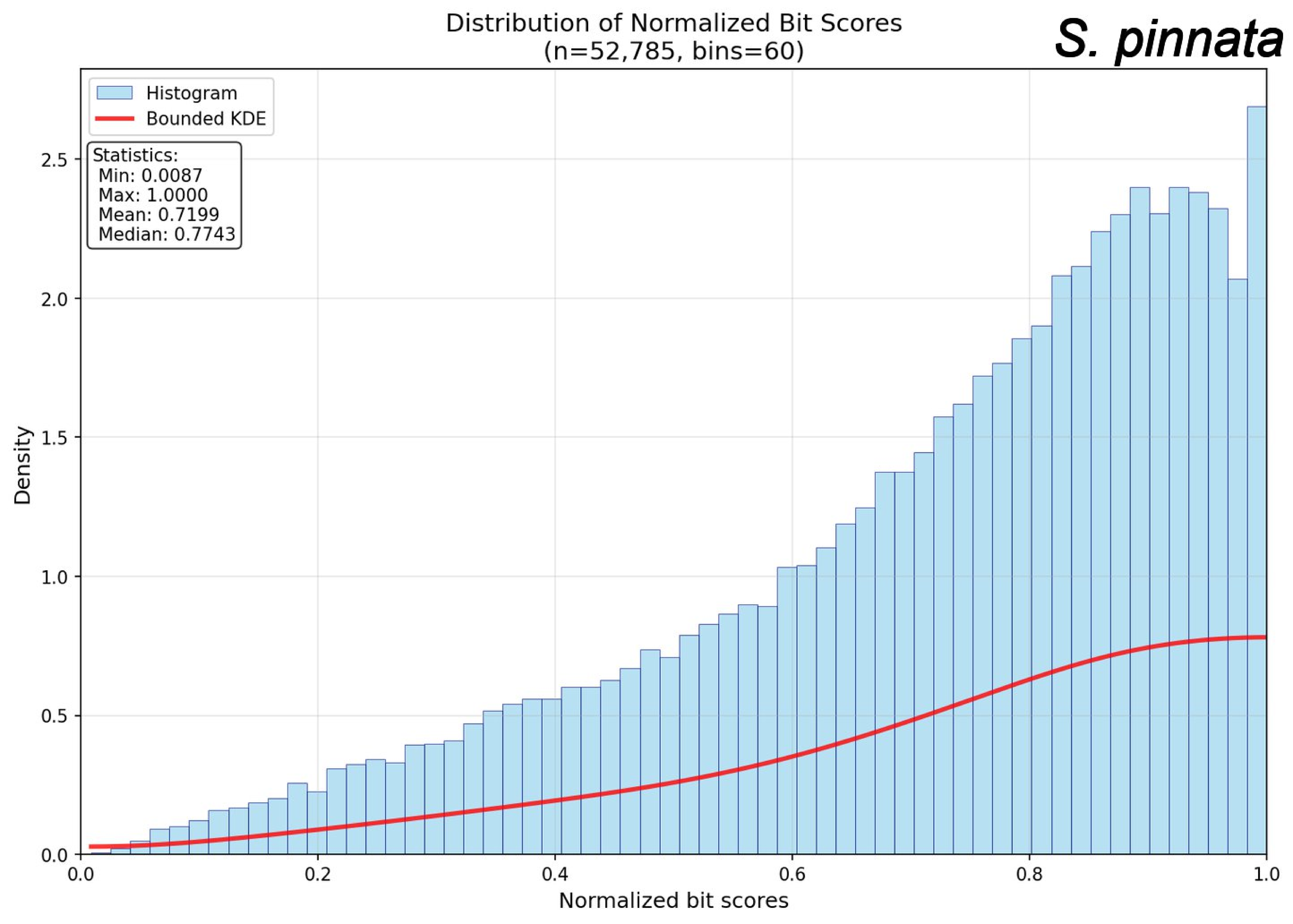
Distribution of Normalized Bit Scores
(n=52,785, bins=60)
S. pinnata
Histogram
Bounded KDE
Statistics:
Min: 0.0087
Max: 1.0000
Mean: 0.7199
Median: 0.7743
Density
Normalized bit scores
0.0
0.5
1.0
1.5
2.0
2.5
0.0
0.2
0.4
0.6
0.8
1.0

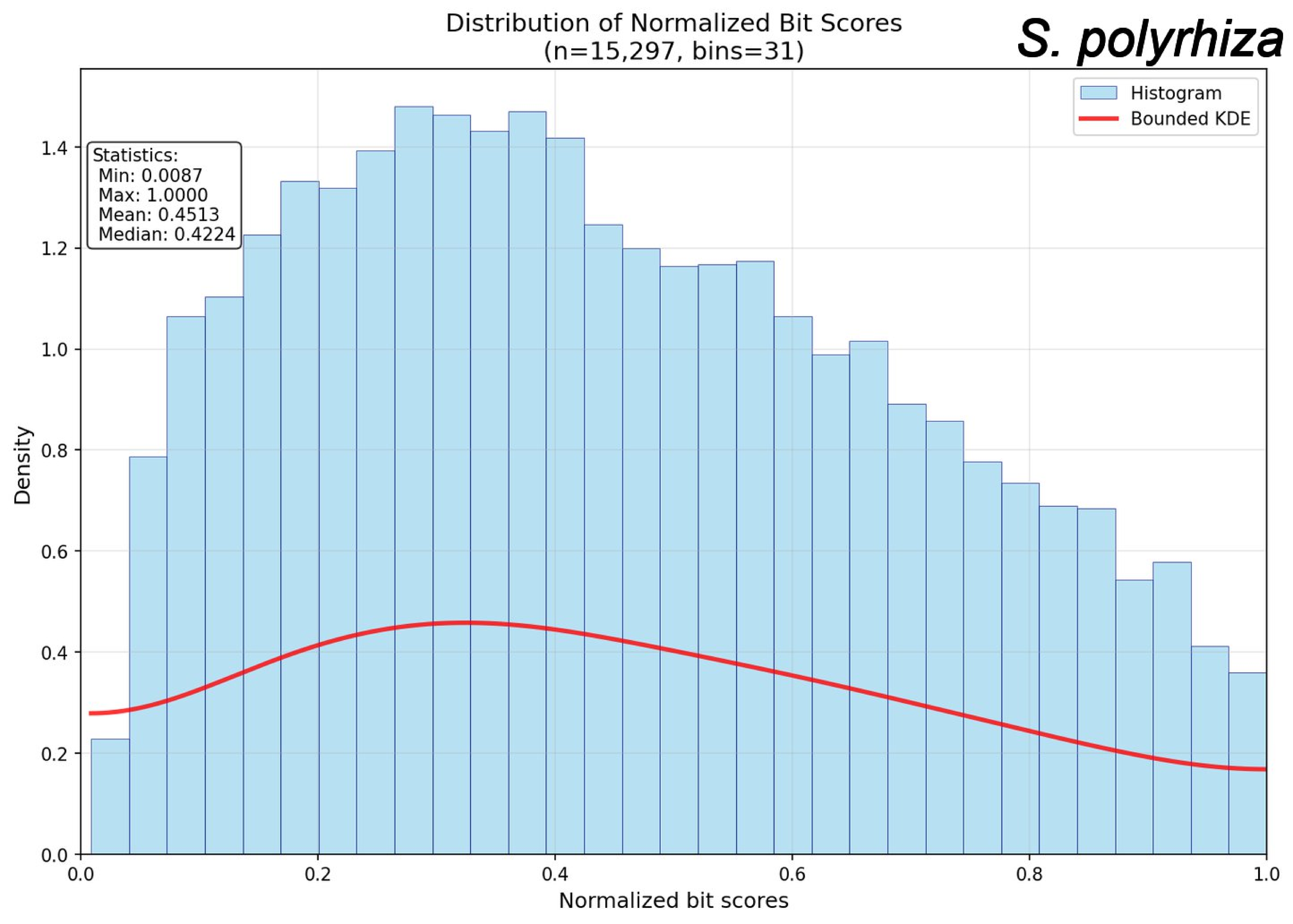

Distribution of Normalized Bit Scores
(n=15,297, bins=31)
S. polyrhiza
Histogram
Bounded KDE
Statistics:
Min: 0.0087
Max: 1.0000
Mean: 0.4513
Median: 0.4224
Density
Normalized bit scores
0.0
0.2
0.4
0.6
0.8
1.0
1.2
1.4

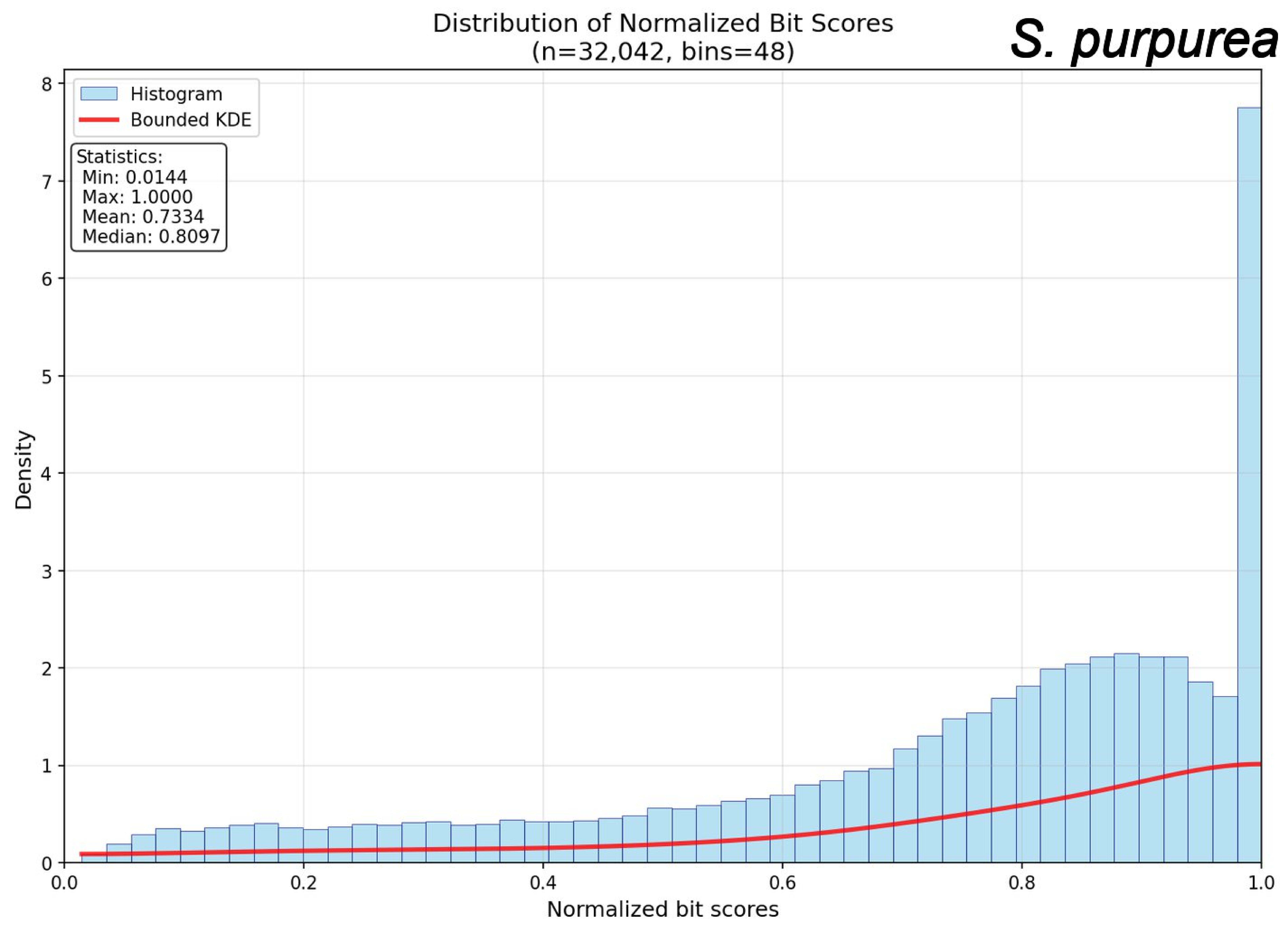
Distribution of Normalized Bit Scores
(n=32,042, bins=48)
S. purpurea
Histogram
Bounded KDE
Statistics:
Min: 0.0144
Max: 1.0000
Mean: 0.7334
Median: 0.8097
Density
Normalized bit scores
0
1
2
3
4
5
6
7
8
0.0
0.2
0.4
0.6
0.8
1.0

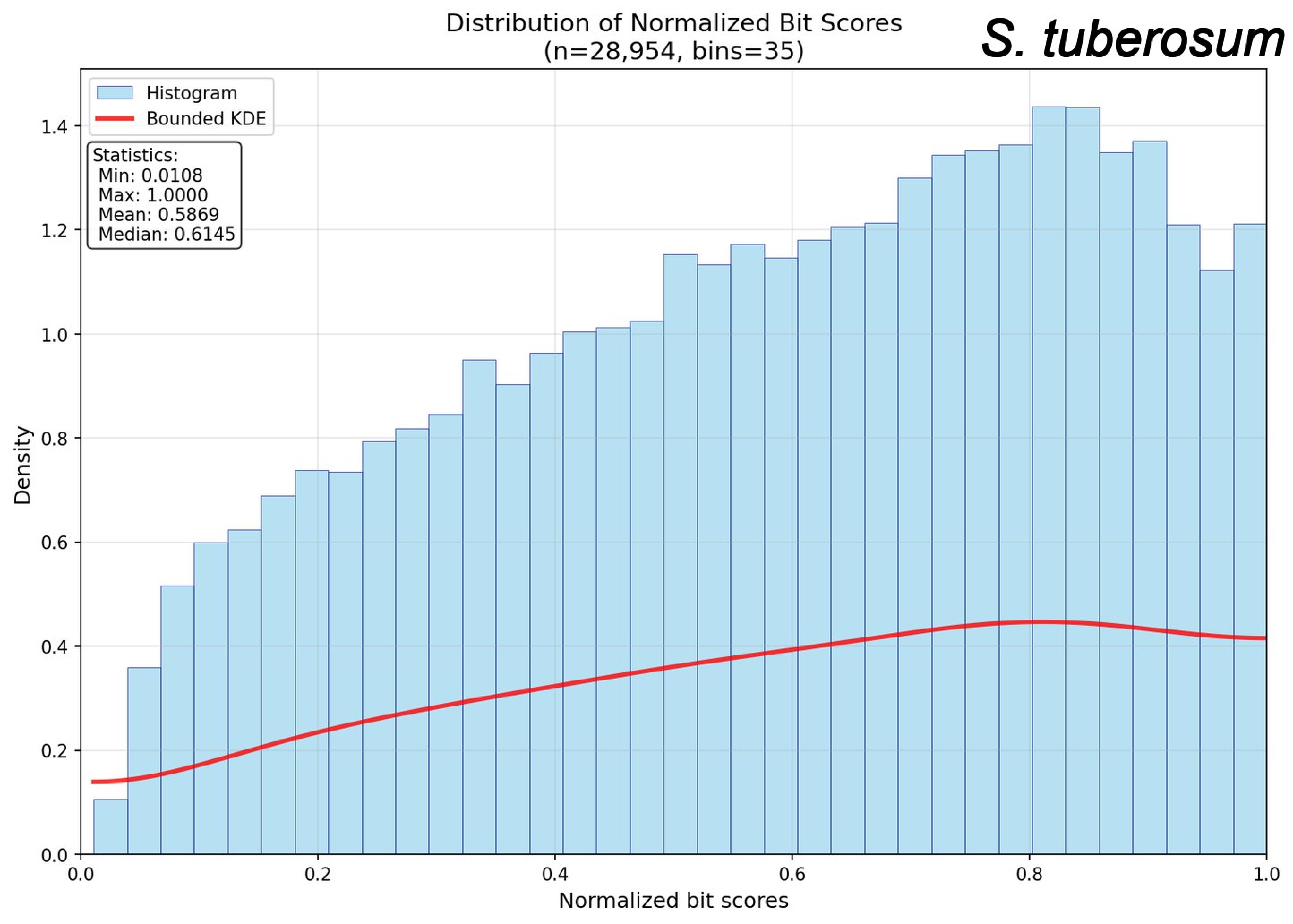

Distribution of Normalized Bit Scores
(n=28,954, bins=35)
S. tuberosum
Histogram
Bounded KDE
Statistics:
Min: 0.0108
Max: 1.0000
Mean: 0.5869
Median: 0.6145
Density
Normalized bit scores
0.0
0.2
0.4
0.6
0.8
1.0
1.2
1.4

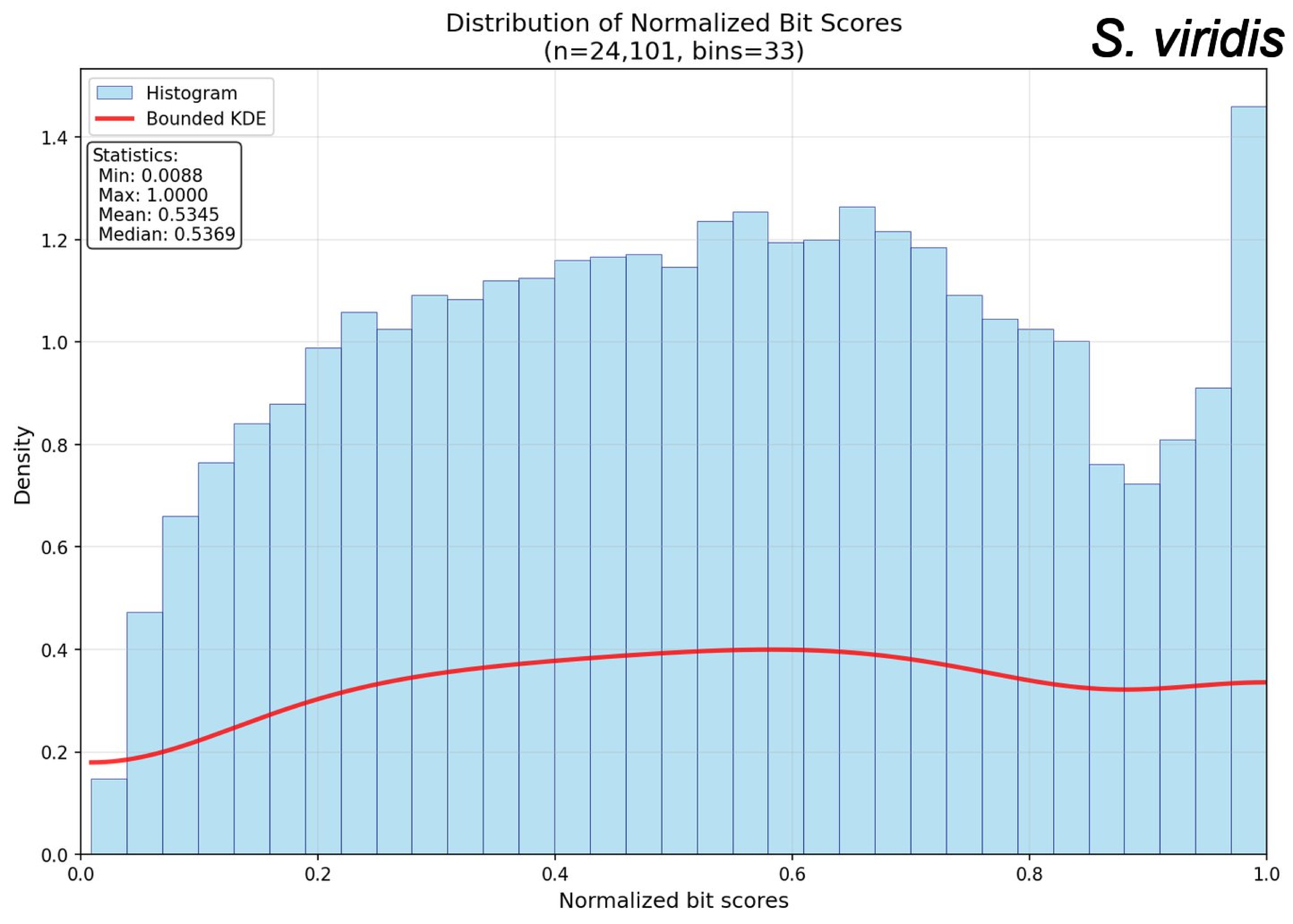
Distribution of Normalized Bit Scores
(n=24,101, bins=33)
S. viridis
Histogram
Bounded KDE
Statistics:
Min: 0.0088
Max: 1.0000
Mean: 0.5345
Median: 0.5369
Density
Normalized bit scores
0.0
0.2
0.4
0.6
0.8
1.0
1.2
1.4

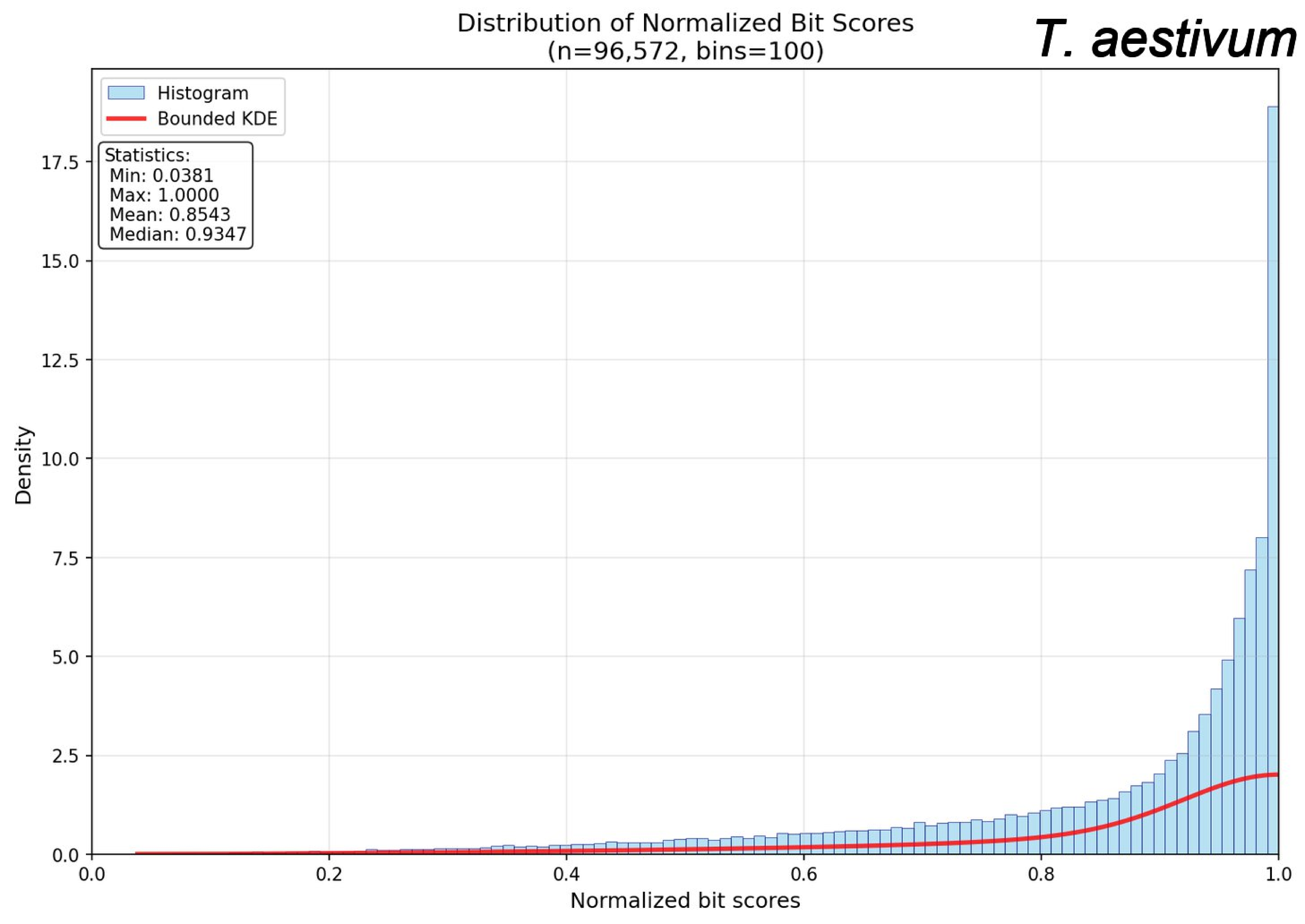
Distribution of Normalized Bit Scores
(n=96,572, bins=100)
T. aestivum
Histogram
Bounded KDE
Statistics:
Min: 0.0381
Max: 1.0000
Mean: 0.8543
Median: 0.9347
Density
Normalized bit scores
0.0
2.5
5.0
7.5
10.0
12.5
15.0
17.5
0.2
0.4
0.6
0.8
1.0

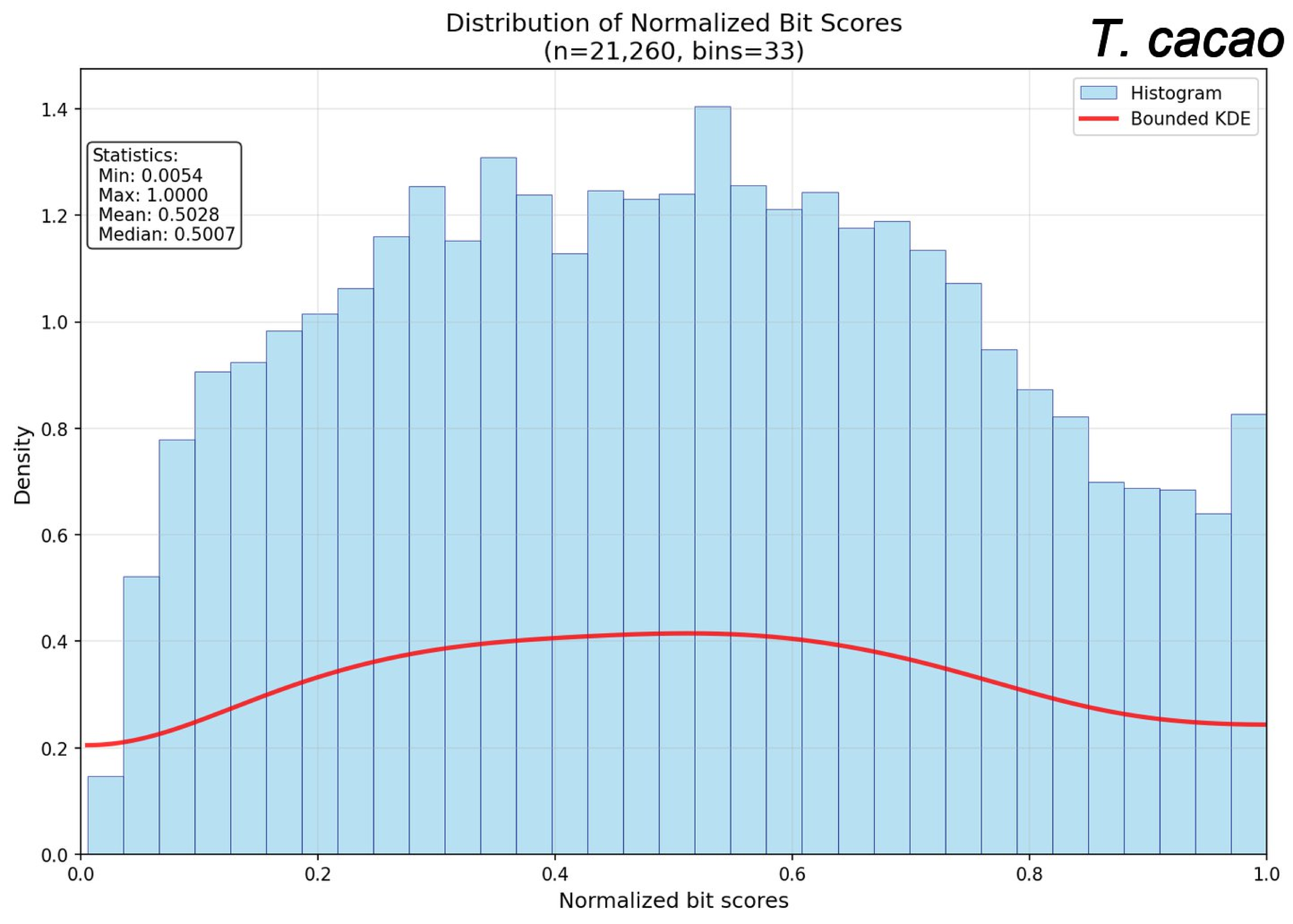

Distribution of Normalized Bit Scores
(n=21,260, bins=33)
T. cacao
Histogram
Bounded KDE
Statistics:
Min: 0.0054
Max: 1.0000
Mean: 0.5028
Median: 0.5007
Density
Normalized bit scores
0.0
0.2
0.4
0.6
0.8
1.0
1.2
1.4

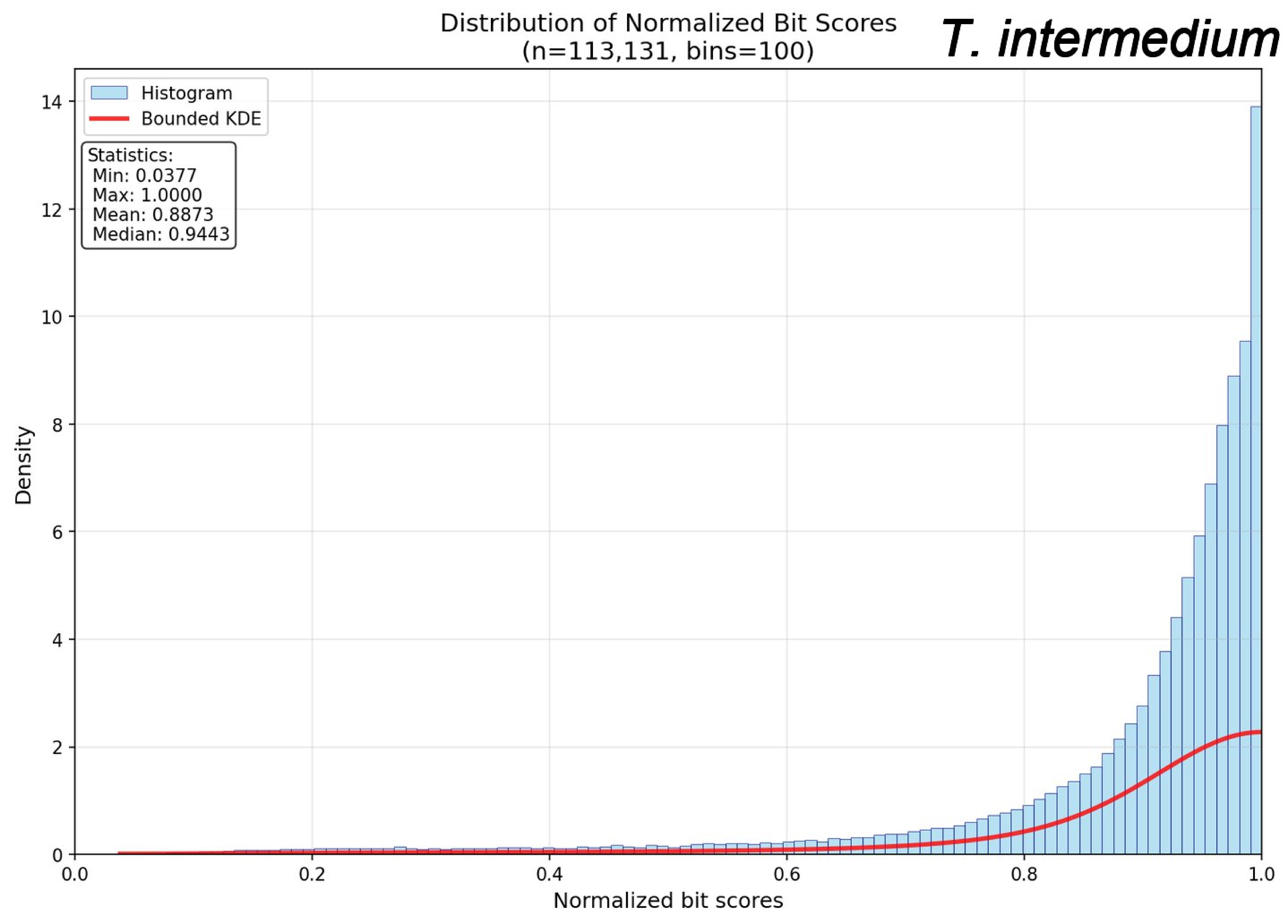

Distribution of Normalized Bit Scores
(n=113,131, bins=100)
T. intermedium
Histogram
Bounded KDE
Statistics:
Min: 0.0377
Max: 1.0000
Mean: 0.8873
Median: 0.9443
Density
Normalized bit scores
0
2
4
6
8
10
12
14
0.0
0.2
0.4
0.6
0.8
1.0

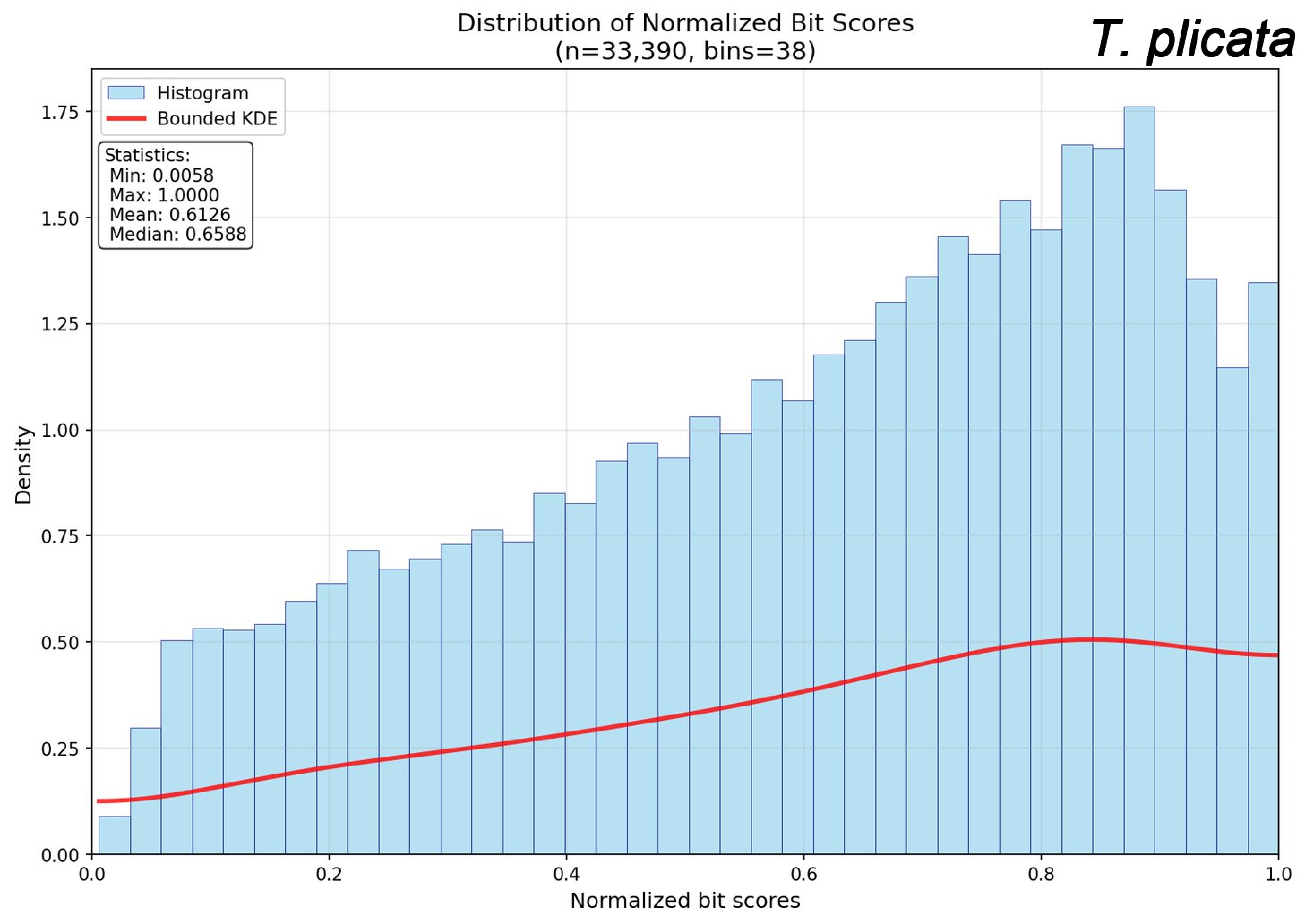
Distribution of Normalized Bit Scores
(n=33,390, bins=38)
T. plicata
Histogram
Bounded KDE
Statistics:
Min: 0.0058
Max: 1.0000
Mean: 0.6126
Median: 0.6588
Density
Normalized bit scores
0.00
0.25
0.50
0.75
1.00
1.25
1.50
1.75
0.0
0.2
0.4
0.6
0.8
1.0

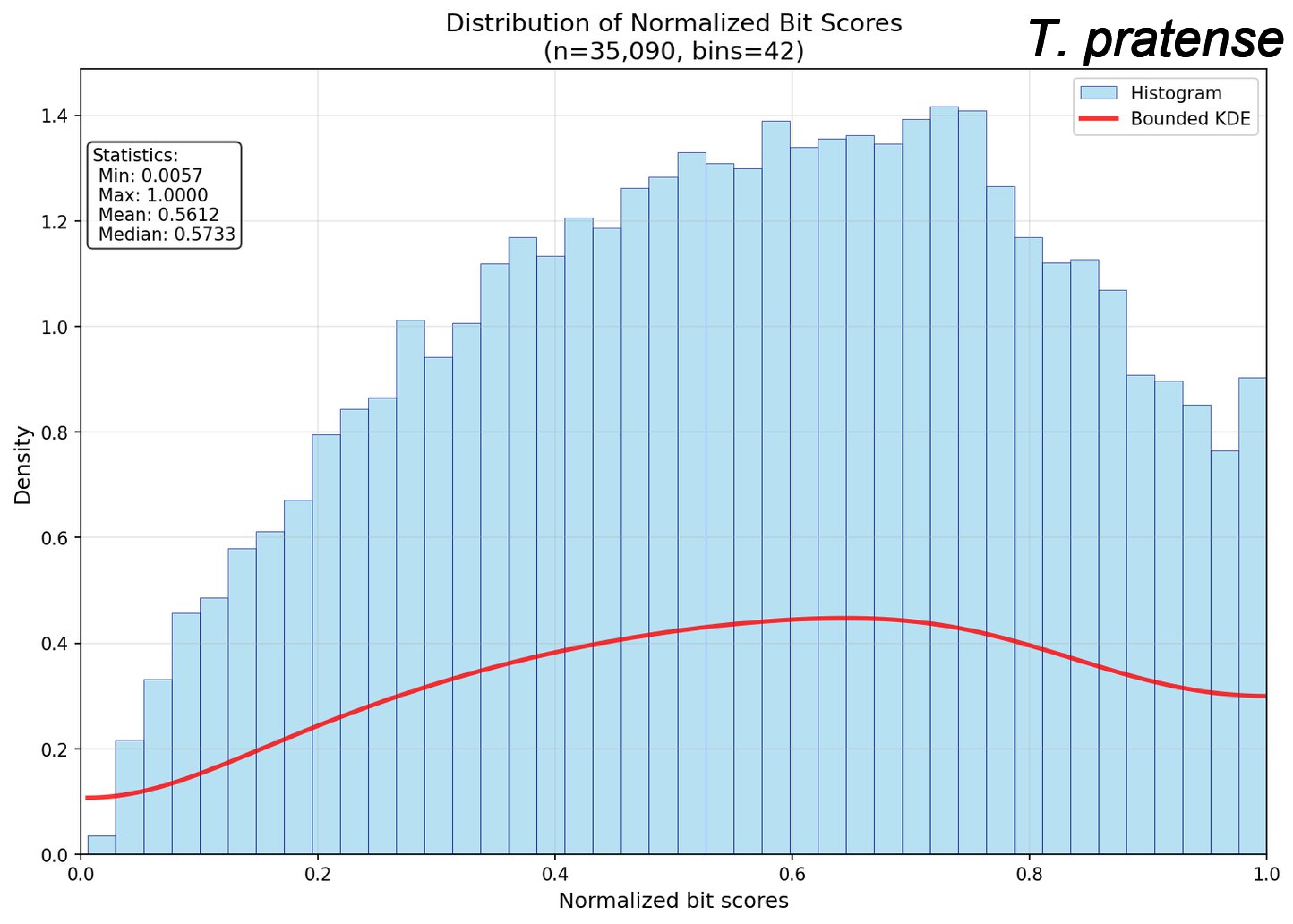

Distribution of Normalized Bit Scores
(n=35,090, bins=42)
T. pratense
Histogram
Bounded KDE
Statistics:
Min: 0.0057
Max: 1.0000
Mean: 0.5612
Median: 0.5733
Density
0.0
0.2
0.4
0.6
0.8
1.0
1.2
1.4
0.0
0.2
0.4
0.6
0.8
1.0
Normalized bit scores

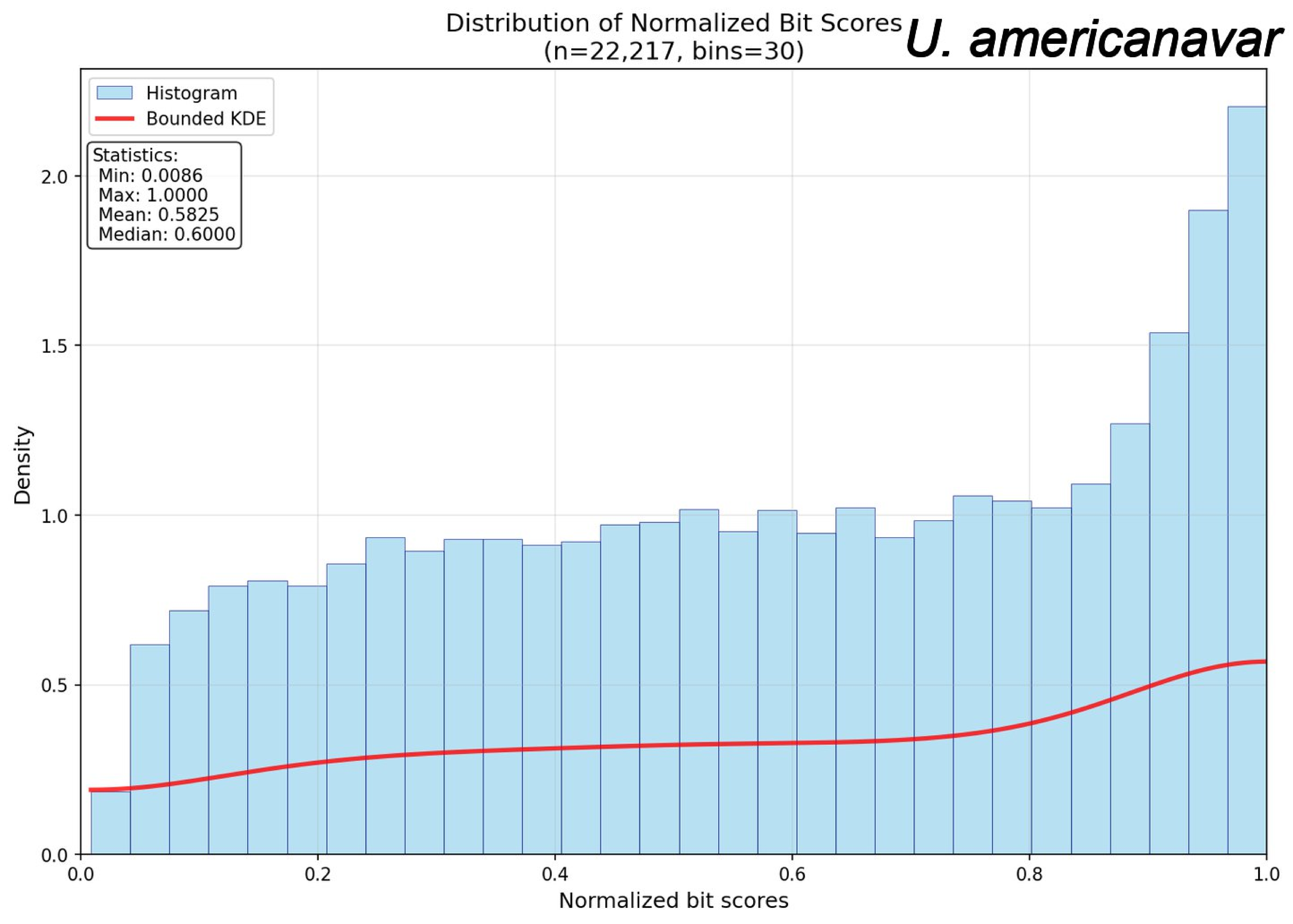

Distribution of Normalized Bit Scores
(n=22,217, bins=30)
U. americanavar
Histogram
Bounded KDE
Statistics:
Min: 0.0086
Max: 1.0000
Mean: 0.5825
Median: 0.6000
Density
0.0
0.5
1.0
1.5
2.0
0.0
0.2
0.4
0.6
0.8
1.0
Normalized bit scores

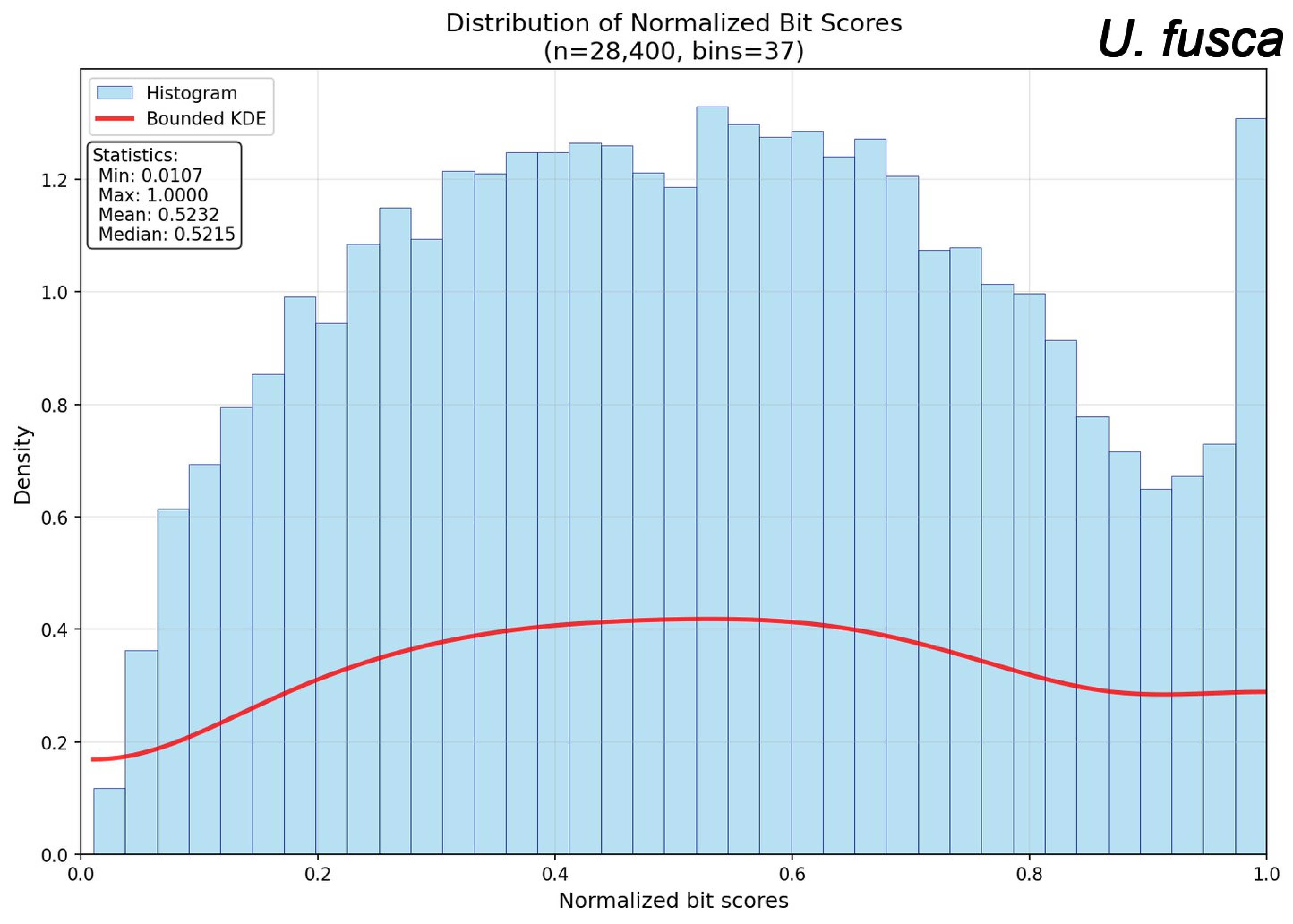
Distribution of Normalized Bit Scores
(n=28,400, bins=37)
U. fusca
Histogram
Bounded KDE
Statistics:
Min: 0.0107
Max: 1.0000
Mean: 0.5232
Median: 0.5215
Density
Normalized bit scores
0.0
0.2
0.4
0.6
0.8
1.0
1.2

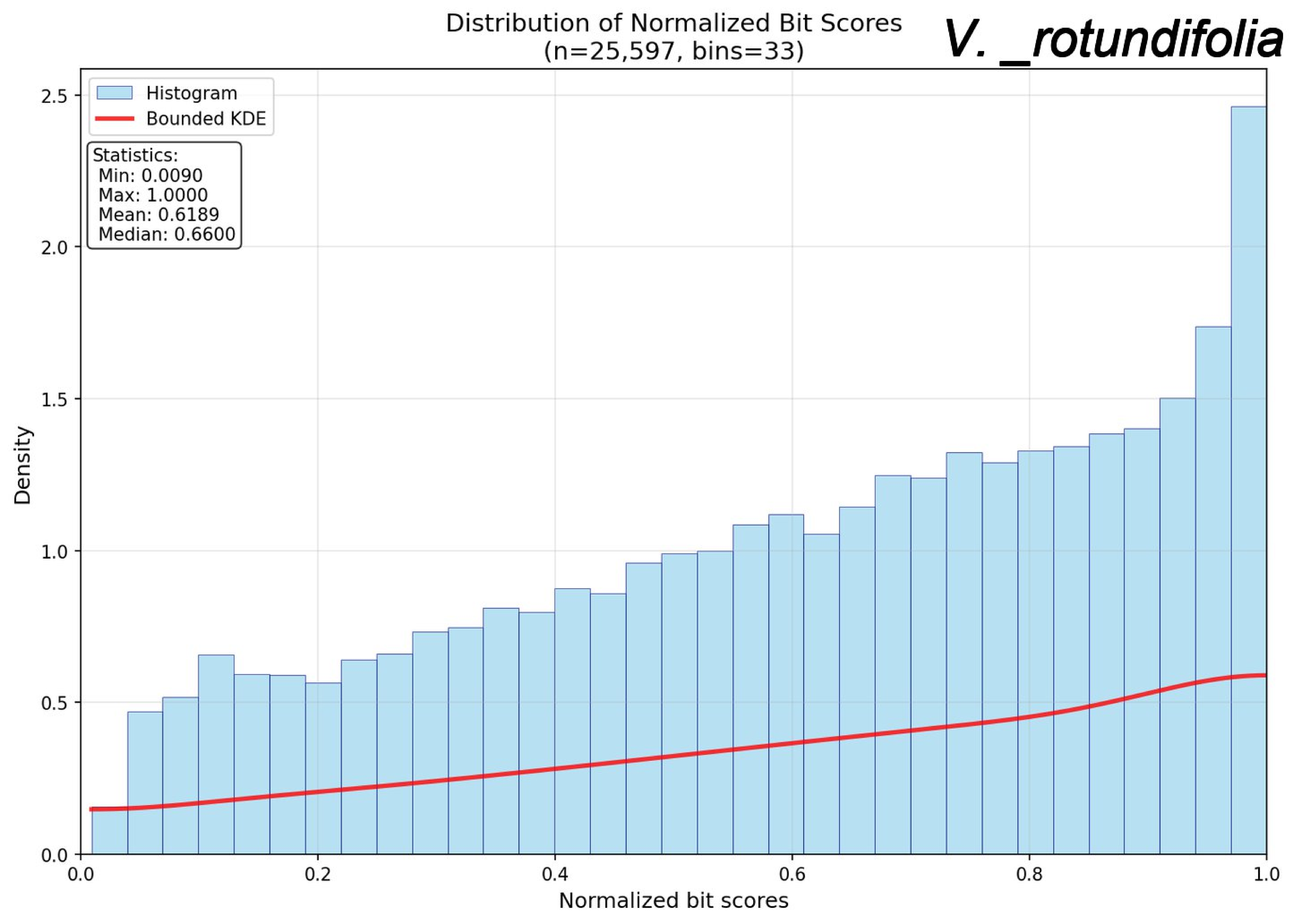

Distribution of Normalized Bit Scores
(n=25,597, bins=33)
V. _rotundifolia
Histogram
Bounded KDE
Statistics:
Min: 0.0090
Max: 1.0000
Mean: 0.6189
Median: 0.6600
Density
Normalized bit scores
0.0
0.5
1.0
1.5
2.0
2.5
0.0
0.2
0.4
0.6
0.8
1.0

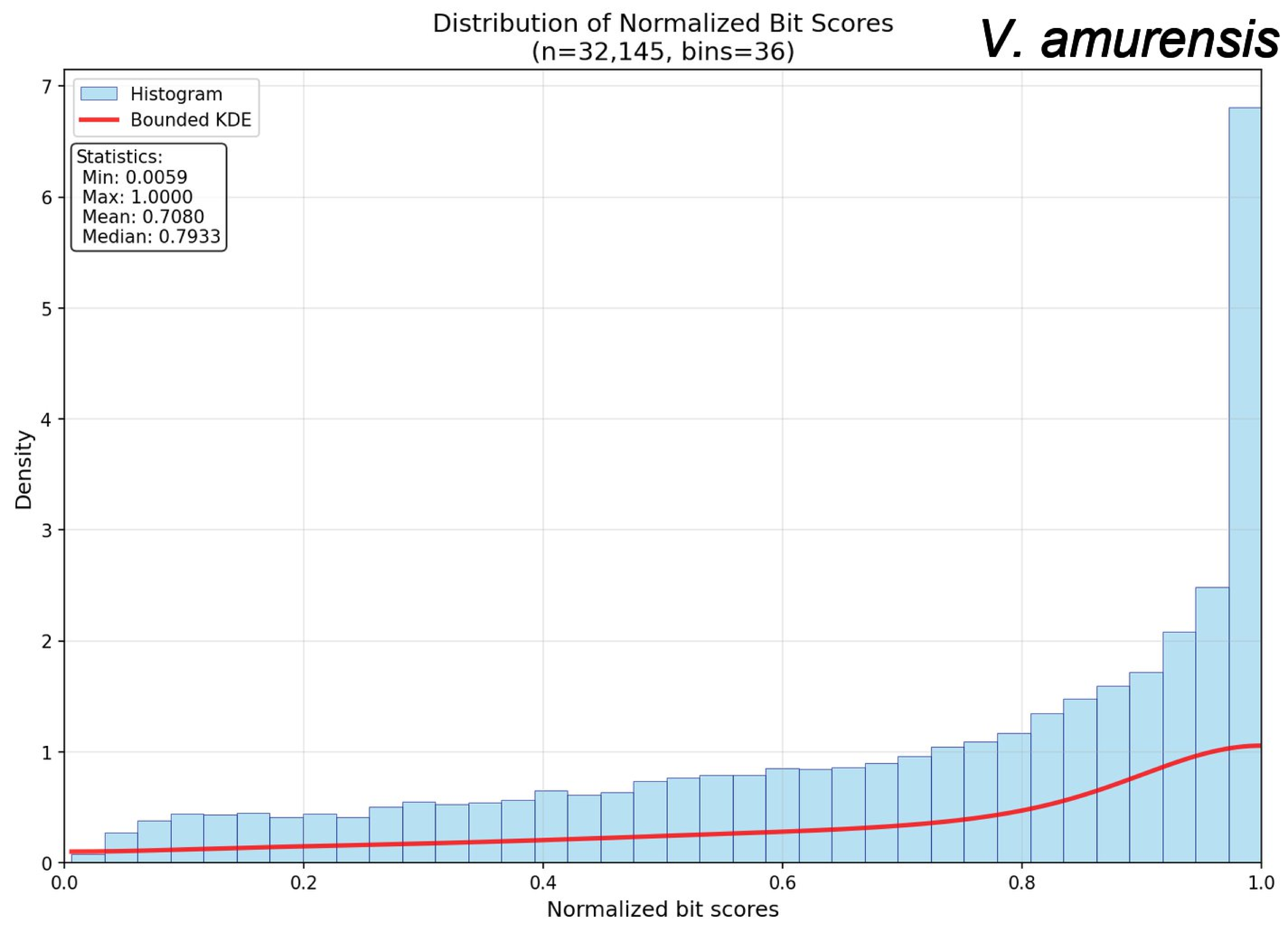

Distribution of Normalized Bit Scores
(n=32,145, bins=36)
V. amurensis
Histogram
Bounded KDE
Statistics:
Min: 0.0059
Max: 1.0000
Mean: 0.7080
Median: 0.7933
Density
Normalized bit scores
0
1
2
3
4
5
6
7
0.0
0.2
0.4
0.6
0.8
1.0

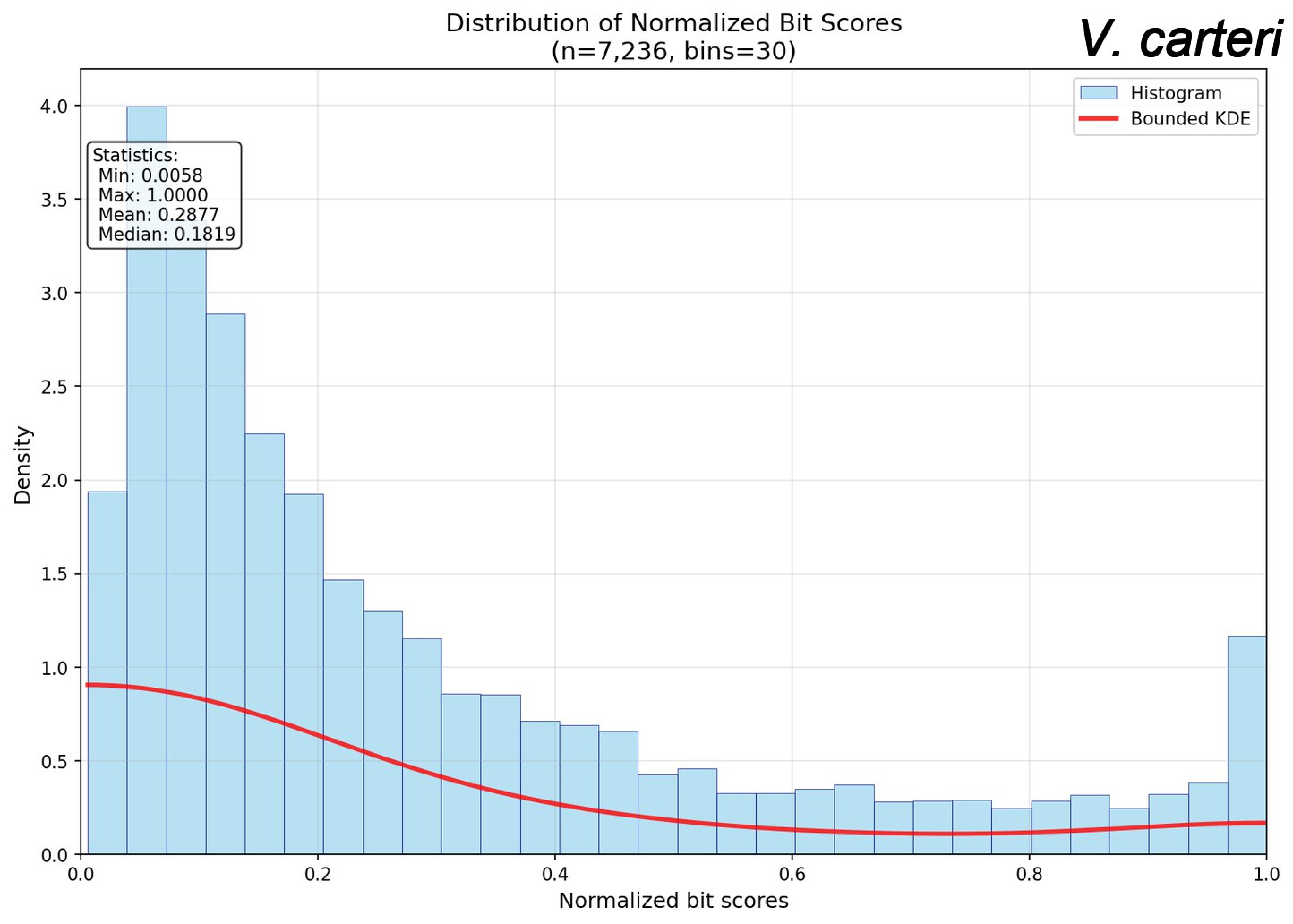
Distribution of Normalized Bit Scores
(n=7,236, bins=30)
V. carteri
Histogram
Bounded KDE
Statistics:
Min: 0.0058
Max: 1.0000
Mean: 0.2877
Median: 0.1819
Density
Normalized bit scores
0.0
0.5
1.0
1.5
2.0
2.5
3.0
3.5
4.0
0.0
0.2
0.4
0.6
0.8
1.0

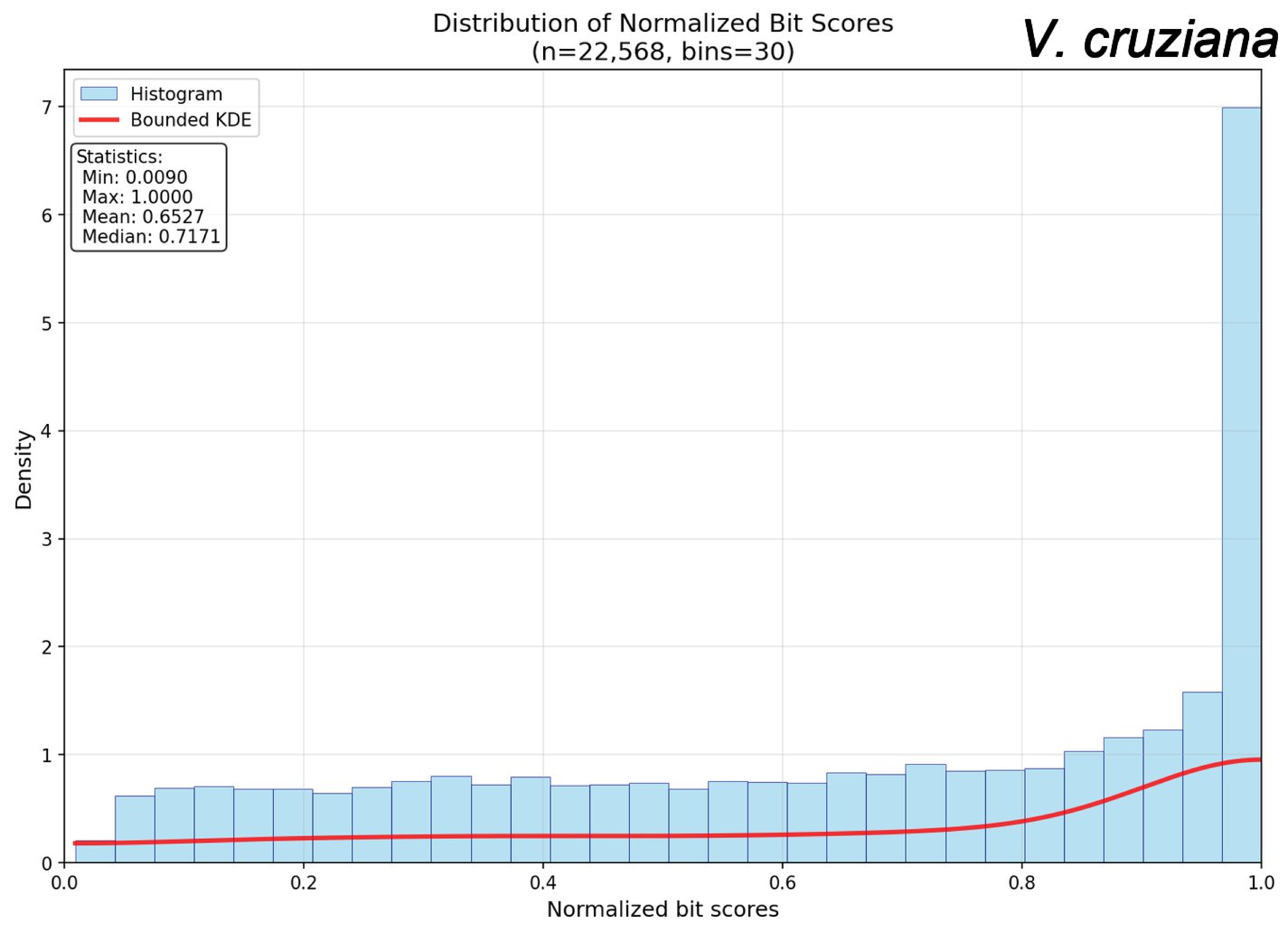

Distribution of Normalized Bit Scores
(n=22,568, bins=30)
V. cruziana
Histogram
Bounded KDE
Statistics:
Min: 0.0090
Max: 1.0000
Mean: 0.6527
Median: 0.7171
Density
Normalized bit scores
0
1
2
3
4
5
6
7
0.0
0.2
0.4
0.6
0.8
1.0

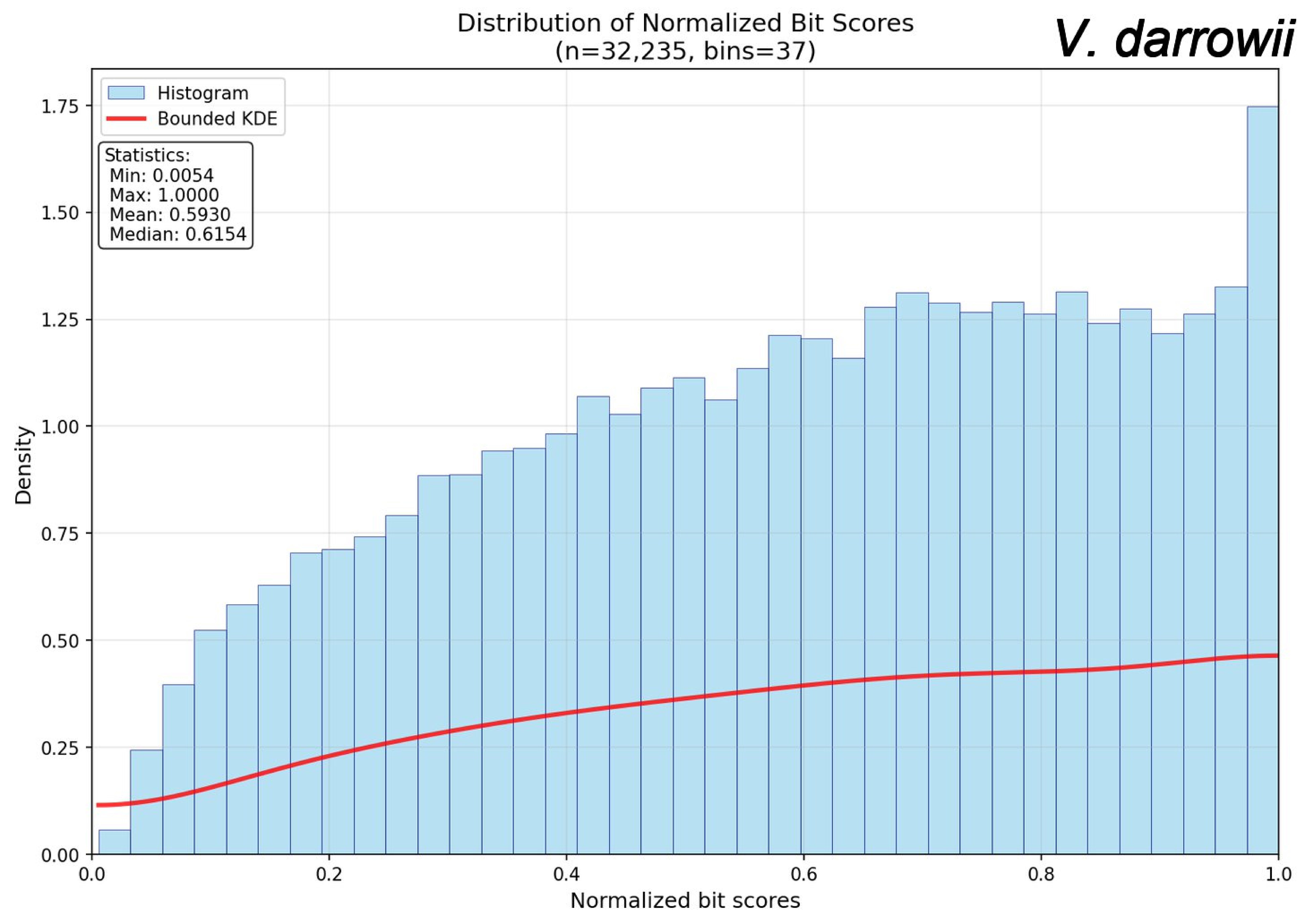

Distribution of Normalized Bit Scores
(n=32,235, bins=37)
V. darrowii
Histogram
Bounded KDE
Statistics:
Min: 0.0054
Max: 1.0000
Mean: 0.5930
Median: 0.6154
Density
Normalized bit scores
0.00
0.25
0.50
0.75
1.00
1.25
1.50
1.75
0.0
0.2
0.4
0.6
0.8
1.0

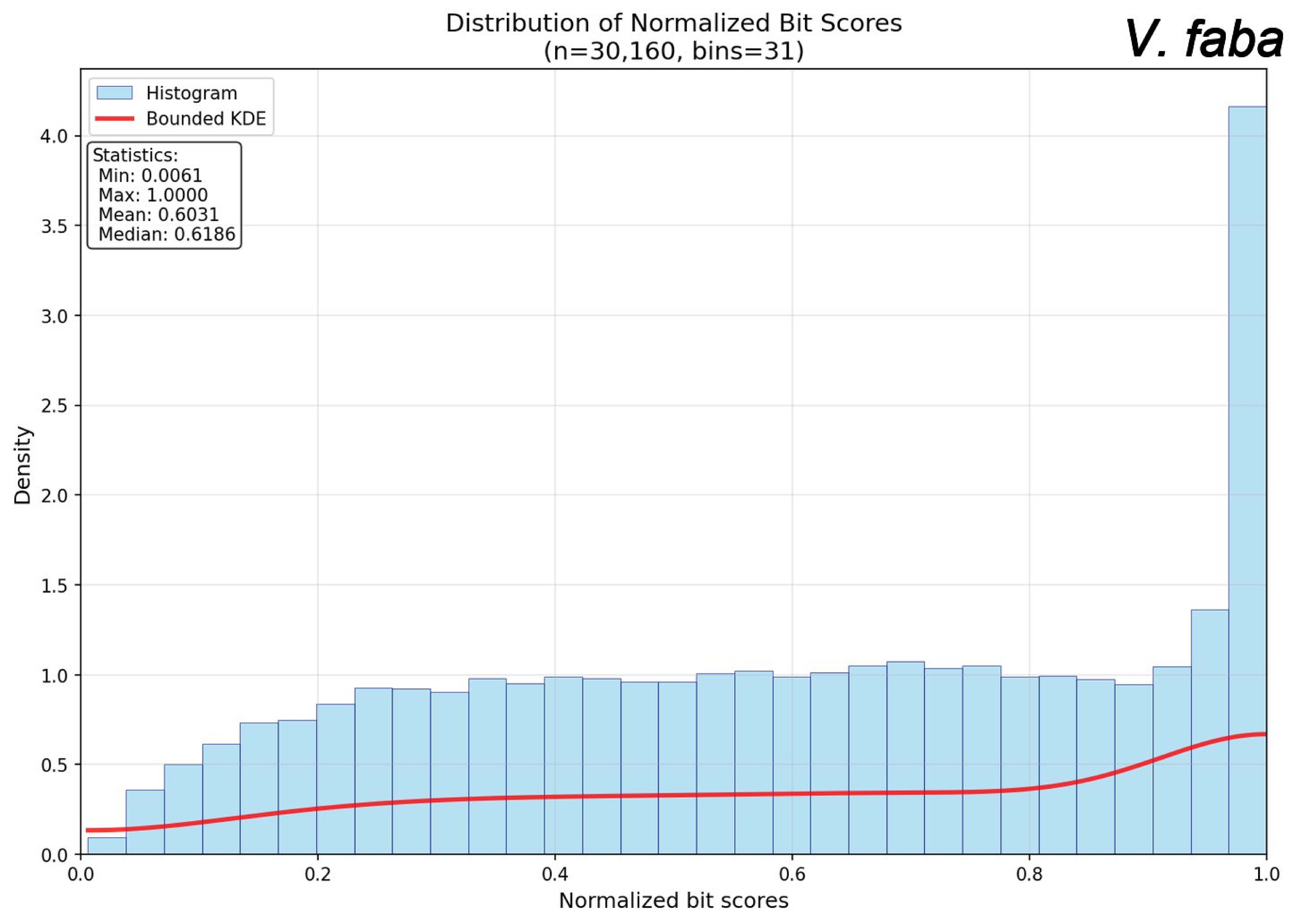
Distribution of Normalized Bit Scores
(n=30,160, bins=31)
V. faba
Histogram
Bounded KDE
Statistics:
Min: 0.0061
Max: 1.0000
Mean: 0.6031
Median: 0.6186
Density
Normalized bit scores
0.0
0.5
1.0
1.5
2.0
2.5
3.0
3.5
4.0
0.2
0.4
0.6
0.8

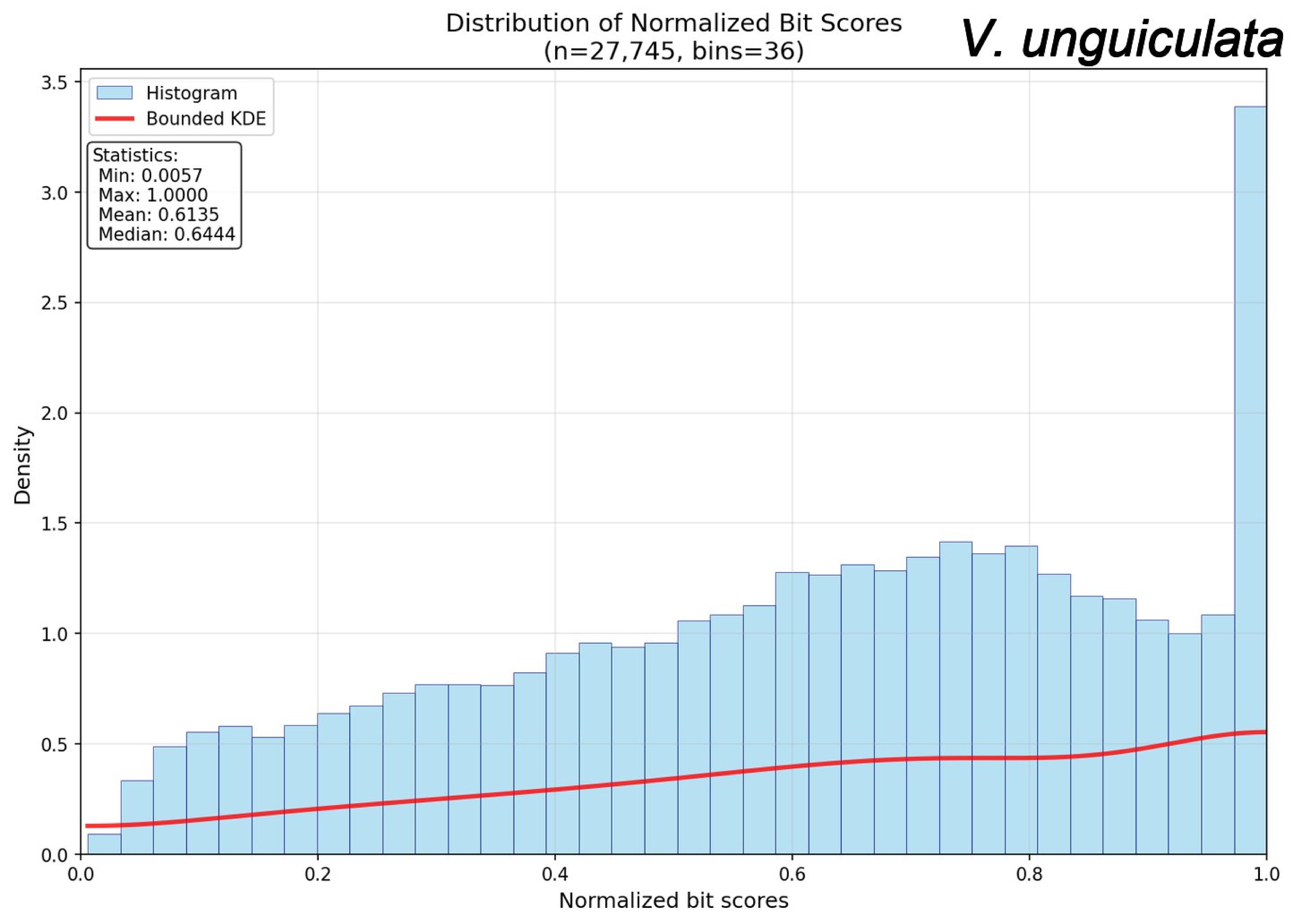

Distribution of Normalized Bit Scores
(n=27,745, bins=36)
V. unguiculata
Histogram
Bounded KDE
Statistics:
Min: 0.0057
Max: 1.0000
Mean: 0.6135
Median: 0.6444
Density
Normalized bit scores
0.0
0.5
1.0
1.5
2.0
2.5
3.0
3.5
0.0
0.2
0.4
0.6
0.8
1.0

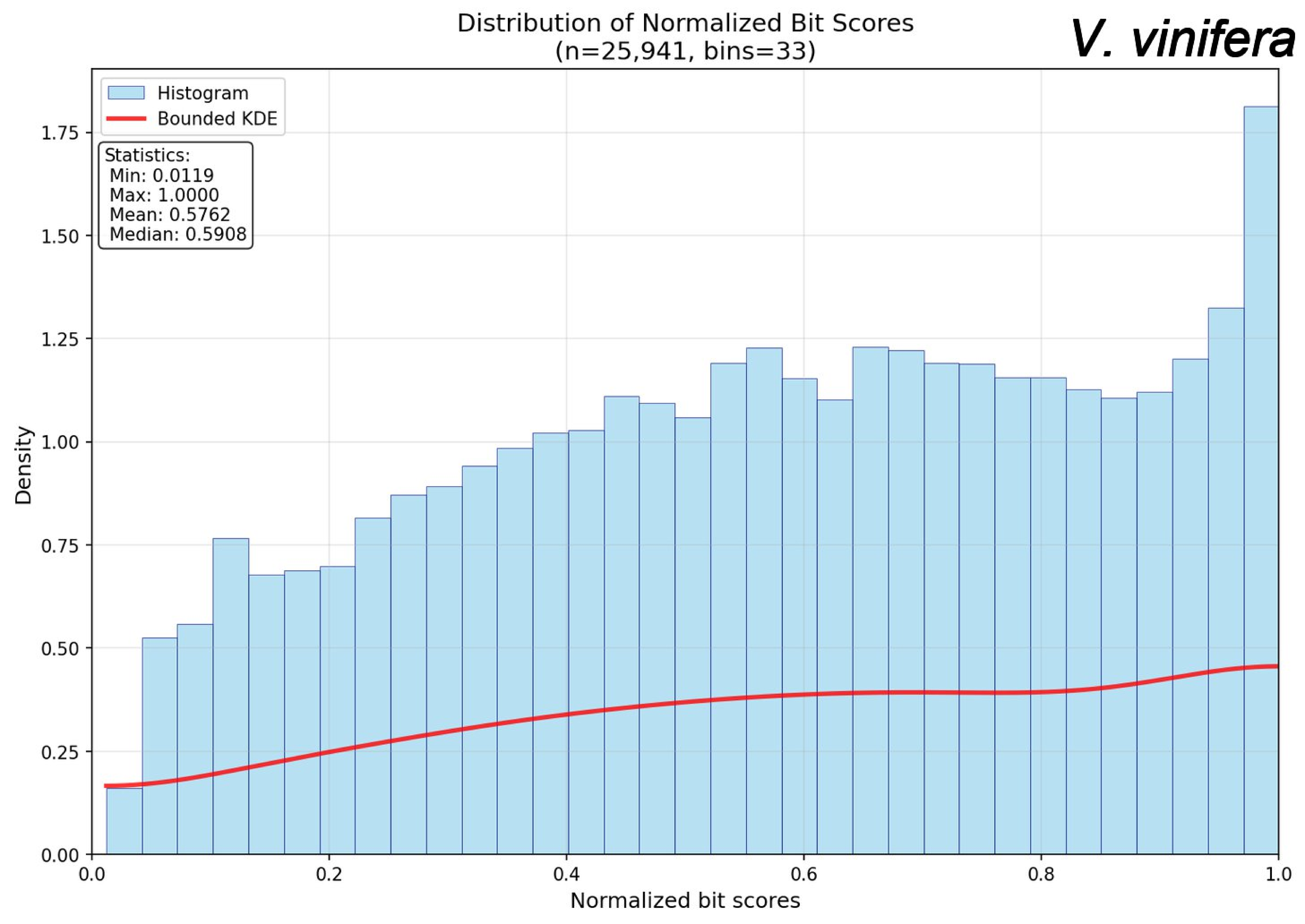

Distribution of Normalized Bit Scores
(n=25,941, bins=33)
V. vinifera
Histogram
Bounded KDE
Statistics:
Min: 0.0119
Max: 1.0000
Mean: 0.5762
Median: 0.5908
Density
Normalized bit scores
0.00
0.25
0.50
0.75
1.00
1.25
1.50
1.75
0.0
0.2
0.4
0.6
0.8
1.0

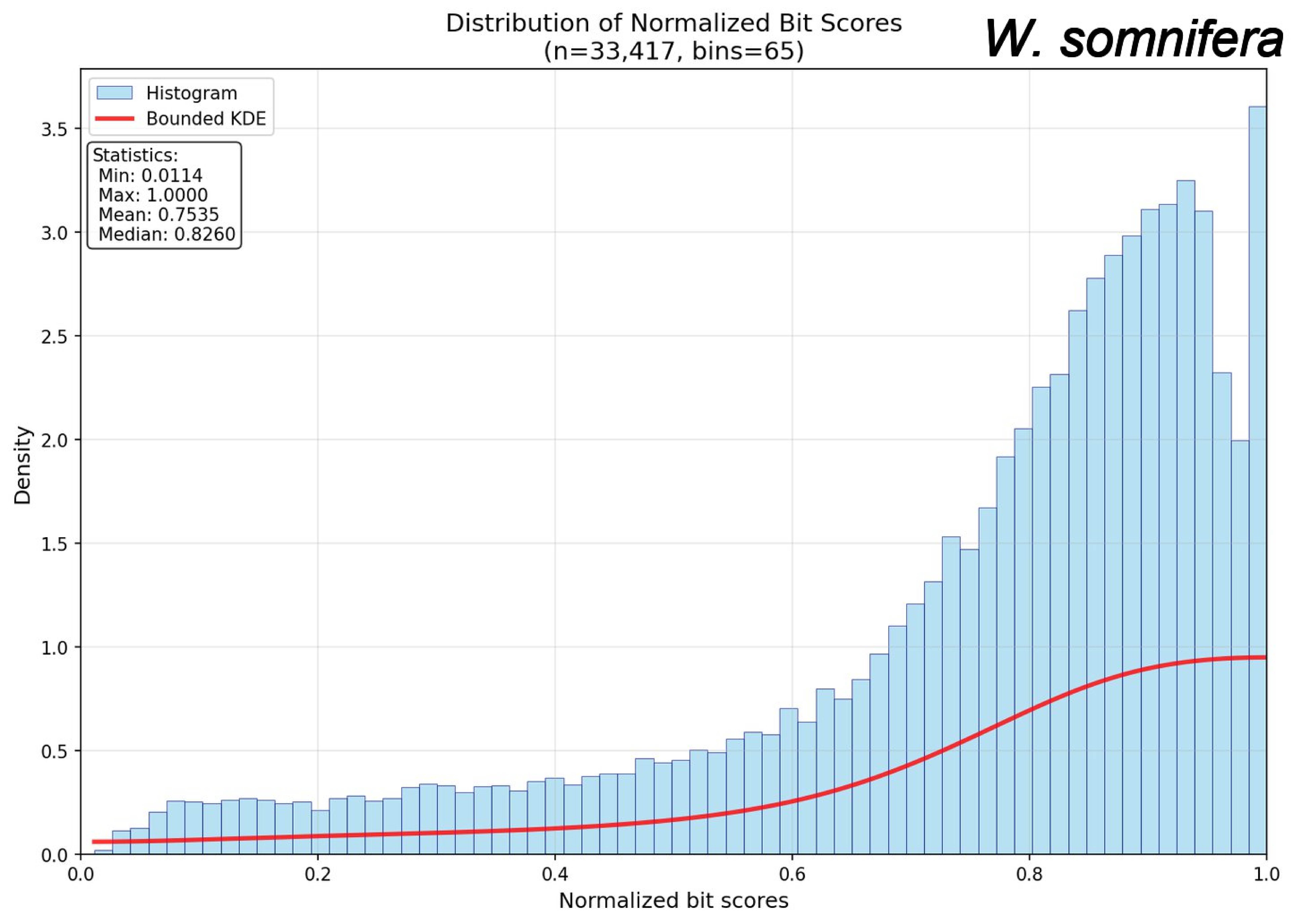
Distribution of Normalized Bit Scores
(n=33,417, bins=65)
W. somnifera
Histogram
Bounded KDE
Statistics:
Min: 0.0114
Max: 1.0000
Mean: 0.7535
Median: 0.8260
Density
Normalized bit scores
0.0
0.5
1.0
1.5
2.0
2.5
3.0
3.5
0.0
0.2
0.4
0.6
0.8
1.0

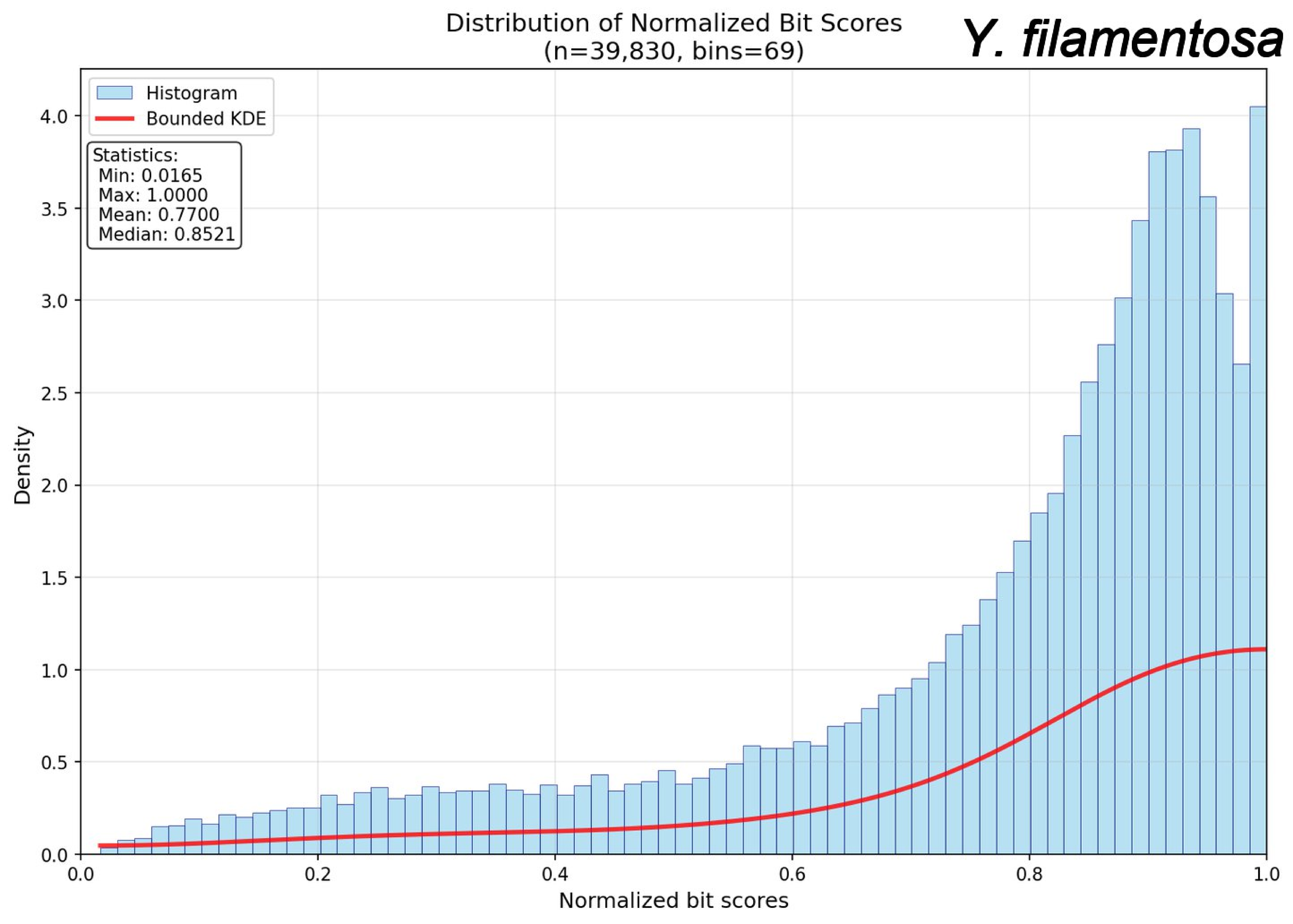
Distribution of Normalized Bit Scores
(n=39,830, bins=69)
Y. filamentosa
Histogram
Bounded KDE
Statistics:
Min: 0.0165
Max: 1.0000
Mean: 0.7700
Median: 0.8521
Density
Normalized bit scores
0.0
0.5
1.0
1.5
2.0
2.5
3.0
3.5
4.0
0.2
0.4
0.6
0.8

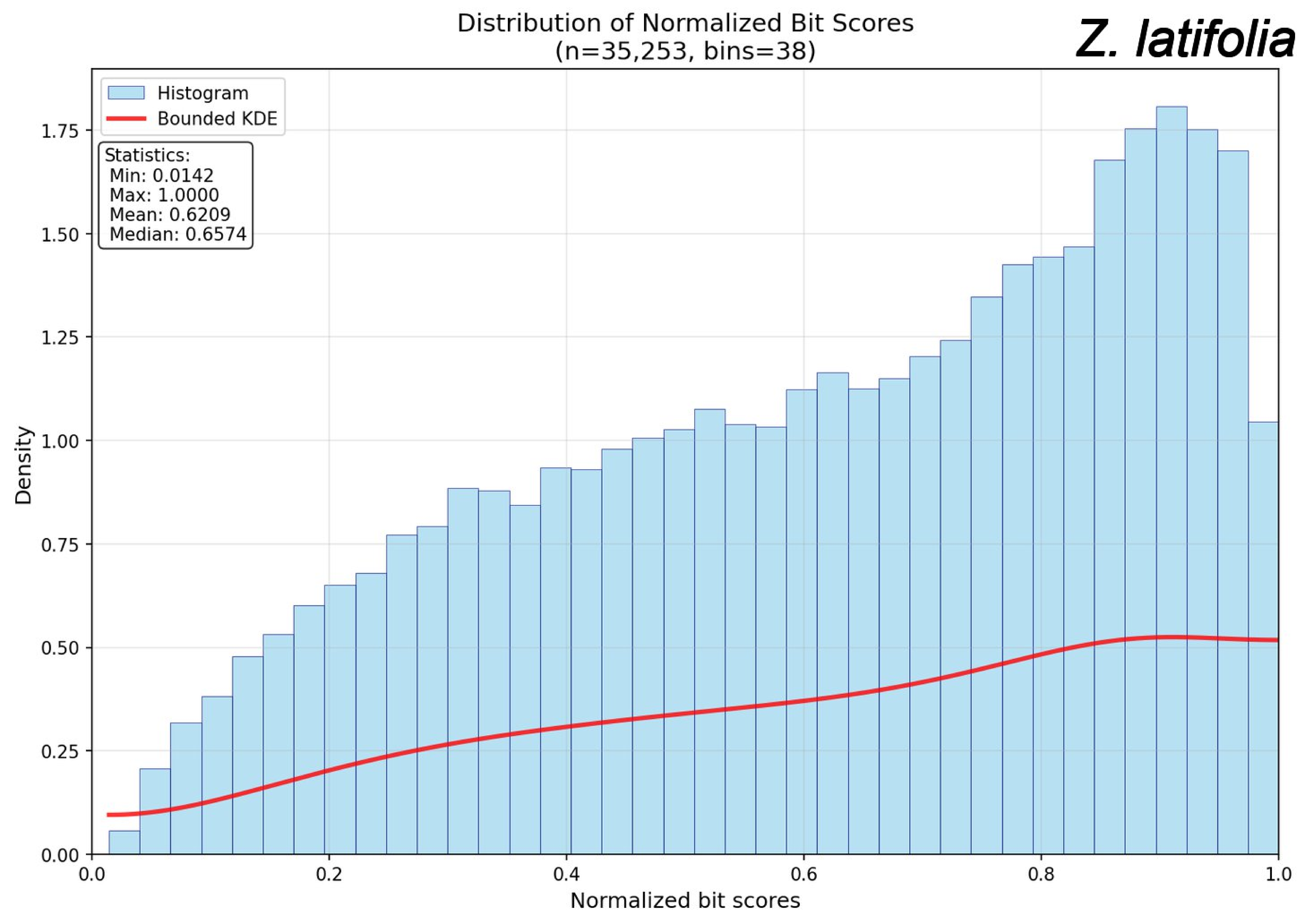
Distribution of Normalized Bit Scores
(n=35,253, bins=38)
Z. latifolia
Histogram
Bounded KDE
Statistics:
Min: 0.0142
Max: 1.0000
Mean: 0.6209
Median: 0.6574
Density
Normalized bit scores
0.00
0.25
0.50
0.75
1.00
1.25
1.50
1.75
0.0
0.2
0.4
0.6
0.8
1.0

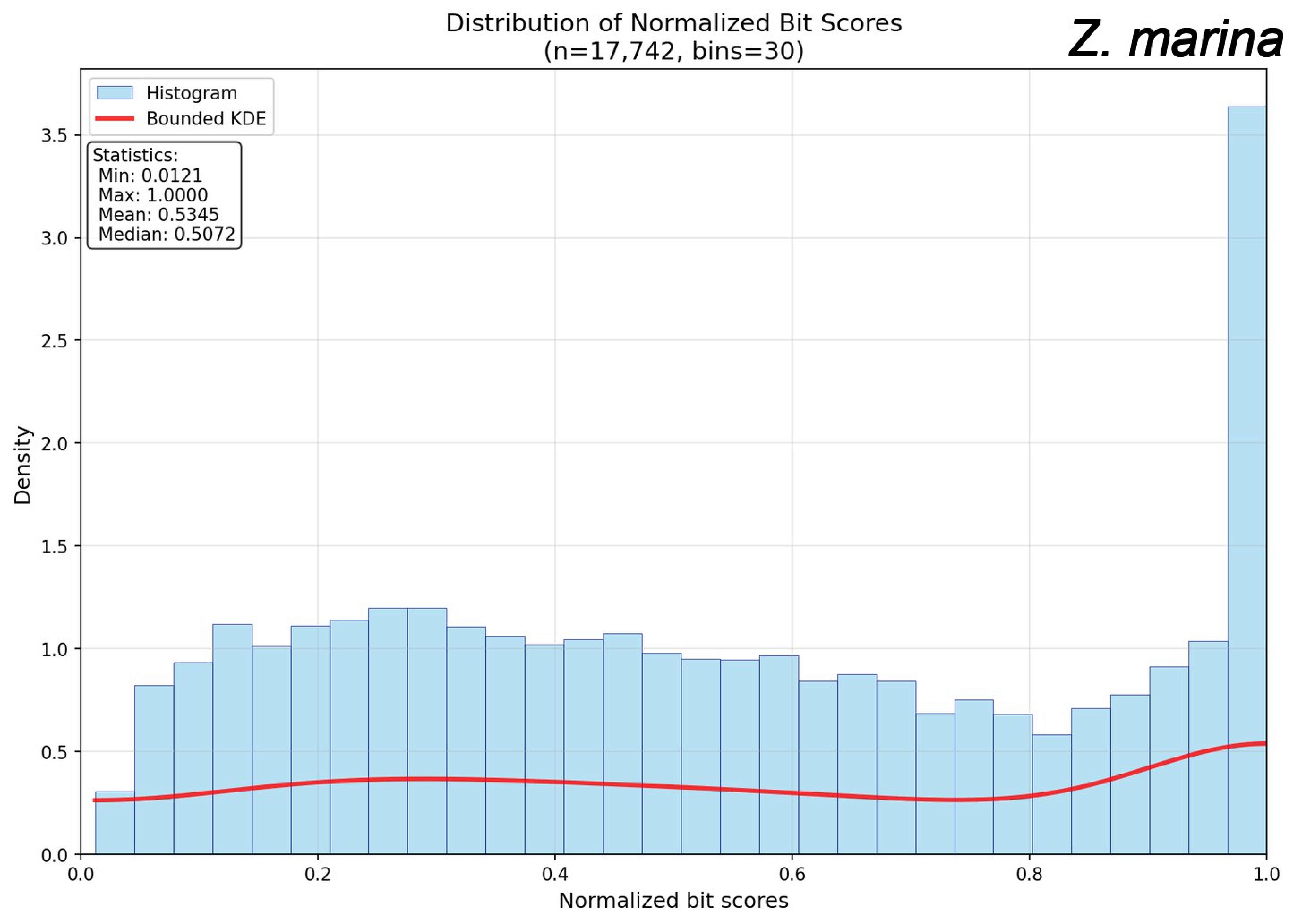
Distribution of Normalized Bit Scores
(n=17,742, bins=30)
Z. marina
Histogram
Bounded KDE
Statistics:
Min: 0.0121
Max: 1.0000
Mean: 0.5345
Median: 0.5072
Density
Normalized bit scores
0.0
0.5
1.0
1.5
2.0
2.5
3.0
3.5
0.0
0.2
0.4
0.6
0.8
1.0

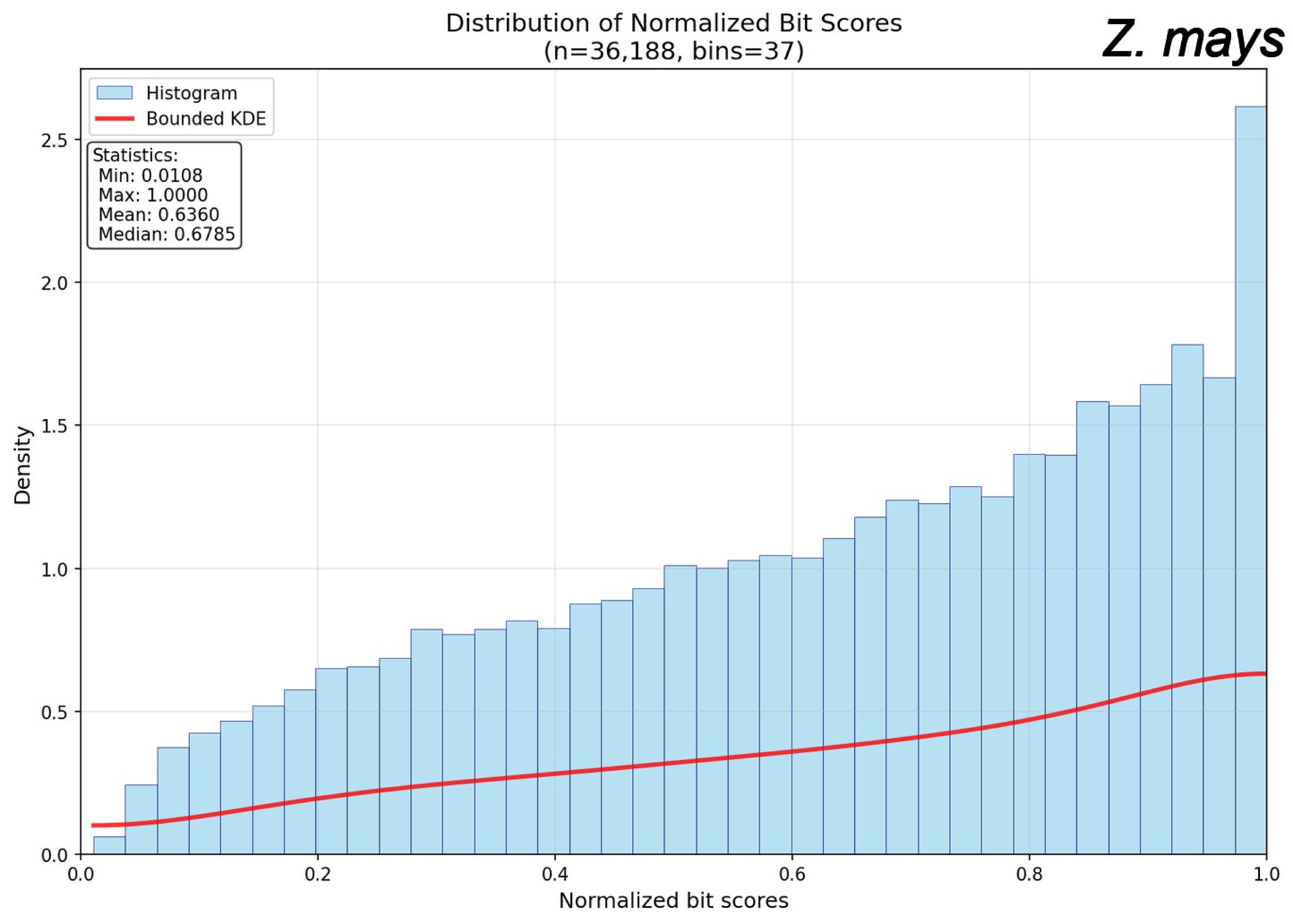
Distribution of Normalized Bit Scores
(n=36,188, bins=37)
Z. mays
Histogram
Bounded KDE
Statistics:
Min: 0.0108
Max: 1.0000
Mean: 0.6360
Median: 0.6785
Density
Normalized bit scores
0.0
0.5
1.0
1.5
2.0
2.5
0.0
0.2
0.4
0.6
0.8
1.0
