## Supplementary Results 2 for "DupyliCate: mining, classifying, and characterizing gene duplications"

### Ka/Ks computation correlation analysis results

A correlation analysis was performed using Ka/Ks values computed for the gene duplicates in DuypliCate using the reimplemented Python code in the script and the Ka/Ks values computed for the gene duplicates by KaKs_Calculator. The *A. thaliana* Nd-1 dataset was used for the analysis without the BUSCO-based thresholding mode to obtain more gene duplicates for Ka/Ks computation for a confident correlation analysis. Nd-1 accession was chosen over Col-0 since it has slightly more protein coding genes than Col-0 and hence could be useful for a better correlation analysis. The statistical results of the correlation analyses were as follows:

**Nei Gojobori implementation correlation analysis**

Final dataset size: 20273 genes
Number of valid points: 18895
Unique values in KaKs_Calculator implementation of NG : [0.027645 0.194868 0.343314 0.411139]
Unique values in DupyliCate implementation of NG: [0.027645 0.194868 0.343314 0.411139]
Spearman correlation: 1.000 (p = 0)
Percentage of problematic genes: 0.0%

Difference Distribution Analysis:
Total genes: 18895
Differences < 0.001: 18895 (100.0%)
Differences < 0.01: 18895 (100.0%)
Differences < 0.05: 18895 (100.0%)
Differences > 0.05: 0 (0.0%)

Correlation comparison:
Spearman (rank-based): 1.000000
Pearson (value-based): 1.000000
Suggests proportional differences

**Modified Yang Nielsen implementation correlation analysis**

Final dataset size: 20273 genes
Number of valid points: 20092
Unique values in KaKs_Calculator implementation of MYN : [0.018121 0.125167 0.379578 0.443544]
Unique values in DupyliCate implementation of MYN: [0.017748 0.120787 0.379578 0.437956]
Spearman correlation: 0.989 (p = 0)

Difference Distribution Analysis:
Total genes: 20092

Differences < 0.001: 4046 (20.1%)
Differences < 0.01: 14510 (72.2%)
Differences < 0.05: 19136 (95.2%)
Differences > 0.05: 956 (4.8%)

Correlation comparison:
Spearman (rank-based): 0.988844
Pearson (value-based): 0.993521
Suggests proportional differences

Biological Significance Assessment:
Genes with changed biological interpretation: 152/19138
Percentage with changed interpretation: 0.8%

The Ka/Ks values are plotted and the correlation line for NG and MYN methods are shown in the supplementary file . As seen from the plots, DupyliCate NG implementation exactly mimics the NG implementation in KaKs_Calculator while the DupyliCate MYN implementation is an approximate implementation of the MYN method implemented in KaKs_Calculator.
